# A revision of the Neotropical *Costaceae*: results from sixty years of taxonomic struggle

**DOI:** 10.1101/2025.01.15.633188

**Authors:** P.J.M. Maas, H. Maas-van de Kamer, T. André, D. Skinner, E. Valderrama, C.D. Specht

## Abstract

Here we present a taxonomic revision of all neotropical *Costaceae* represented by 90 species, distributed in four monophyletic genera, *Chamaecostus* (9 spp.), *Costus* (76 spp.), *Dimerocostus* (5 spp.), and *Monocostus* (1 sp.). The work resulted in three new combinations, several nomenclatural changes, and four insufficiently known species are mentioned to demonstrate the need for continued and ongoing work on this charismatic yet often cryptic lineage of plants. We highlight the seventeen species of *Costus* and one species of *Chamaecostus* that were recently described (Maas et al. 2023) based on knowledge that emerged during this complete revisionary work, 60 years in the making. Included in this revision are details concerning the morphology, distribution, ecology, and phylogenetic relationships among the species studied. All currently recognized species are fully described, including notes on taxonomic synonymy, lectotypification, information on distribution and ecology, conservation status (either as evaluated and published in the IUCN Red List of Threatened Species or as estimated for this publication), vernacular names (when given) and short taxonomic notes. For each species, a distribution map is prepared and at least one colour photograph is included to assist in species identification. Dichotomous keys that include all species are provided, as is a list of all (c. 11.000) exsiccatae studied and an index to scientific names.

## INTRODUCTION

The *Costaceae* or “spiral gingers” are a charismatic lineage of tropical monocots, one of eight families in the order *Zingiberales* which contains economically and horticulturally important families such as the *Musaceae* (bananas), *Zingiberaceae* (gingers), *Cannaceae* (cannas), *Marantaceae* (prayer plants) and *Heliconiaceae* (lobster claws) among others. Recognizable by their characteristic monistichous spiral phyllotaxy, tubular sheathing leaf bases with a pronounced ligule, and showy terminal inflorescences with imbricate bracts arranged in several series of parastiches. These rhizomatous herbs are often abundant in moist low-lands, varzea or flooded forest, clearings and stream beds in the lowland tropics. The *Costaceae* comprise seven monophyletic genera and as a whole are pantropical, with four genera being native to the Neotropics. Of those, three (*Monocostus* K.Schum, *Dimerocostus* Kuntze, *Chamaecostus* C.D.Specht & D.W.Stev.) are endemic to the Neotropics while *Costus* L. is most speciose in the Neotropics but is also found in Africa. Several recent publications have highlighted the rapid diversification of the Neotropical taxa (Andre et al. 2016), particularly of the genus *Costus* with its diversity of flowers that reflect either bee or bird pollination syndromes (Salzman et al. 2015; Valderrama et al. 2020) and among the Brazilian species of *Chamaecostus* which demonstrate the potential for critpic speciation (André et al. 2015). This monograph is an opportunity to highlight species diversity across this charismatic lineage of tropical plants and to provide a context for future phylogenetic, population genetic, and comparative genomic studies that characterize the role of ecological and evolutionary processes in generating patterns of speciation and diversification.

### History of this revision

In 1962 Paul Maas started his taxonomic study of the family *Zingiberaceae* (which at the time included the *Costoideae*) as a graduate student at Utrecht University (The Netherlands). In the framework of Pulle’s “Flora of Suriname”, he prepared descriptions for species of the genera *Costus*, *Dimerocostus*, and *Renealmia.* This resulted in the 1972 publication of Monograph No. 8 *Costoideae* (*Zingiberaceae*) treatment for Flora Neotropica as well as the *Zingiberaceae* treatment for the Flora of Surinam (Maas 1979). Since that time, additional data concerning the *Costaceae* lineage has come to light especially in the form of novel observations based on floral and vegetative characters from living material either in living collections or from field expeditions. These data resulted in the publication of 18 additional species between 1976–1997 (Maas 1976, 1977; Maas & Maas-van de Kamer 1990, 1997). Most recently, 18 new species were described by Maas and colleagues (Maas et al. 2023). A few additional species have been published by other taxonomists including *Costus dirzoi* García-Mendoza & G.Ibarra (García-Mendoza & Ibarra-Manríquez 1991), *Dimerocostus cryptocalyx* N.R.Salinas & Betancur (Salinas & Betancur 2004), *Costus atlanticus* E.M.Pessoa & M.Alves (Pessoa et al. 2020), and *Costus flammulus* K.M.Kay & P.Juárez (Juárez et al. 2023). In the present publication we present a full treatment of the total of c. 90 species currently described and recognized for the neotropical members of the family *Costaceae*. In addition, several nomenclatural changes are made as necessary to keep with the code, as for example in a group of species now recognized as the *Costus guanaiensis* complex and the lineage that what was up until now referred to as *Costus laevis* Ruiz & Pav.

Most important to the current revision are data collected on living material and during field work, as the 1972 Monograph (Maas 1972) had been based almost exclusively on the study of herbarium material. The difference between studying a dry herbarium specimen and investigating a living plant is enormous. As such, many collecting trips between 1972 and 2022 have added insights, observations, and comparative data to our understanding of species delimitations across the *Costaceae*.

Over the past more-than-60 years, many collections have been made throughout the Neotropics. As a result, our knowledge of the distribution of species and species complexes has been considerably advanced. The number of specimens has also multiplied: as the 1972 Monograph was based on c. 3000 specimens, the most recent list of observed specimens now comprises over 11.000 entries.

In addition to new accessions represented in living collections and as herbarium vouchers, many photographs sent to us by colleagues from all over tropical America added valuable characters for consideration in this revision. This includes characters such as colour of the floral organs, orientation of the floral opening with respect to the axis of the inflorescence, colour of the bracts, coloration of the stem, and nature and colour of hairs on stem and leaf surfaces. Thus, we recognize that this work would not have been possible without our network of collaborators: collectors, botanists, researchers, and naturalists.

During recent collecting trips, samples of leaf material were collected directly in silica and stored in dried conditions for optimal extraction of unfragmented DNA. Each DNA extraction is linked to an herbarium voucher, includes a photograph, and in some cases these specimens are also being retained in a living collection – all following best practices to build an “extended specimen” framework for *Costaceae*. The results of these molecular phylogenetic studies are ongoing as we continually add new accessions to test the monophyly of the proposed species, to help delimit new taxonomic groupings, and to understand the evolutionary relationships of the species and their populations. A molecular phylogeny that includes samples from all species here presented and with multiple accessions to test monophyly of species in addition to relationships among species will be used to futher refine taxonomic decisions left unanswered in this revision.

## GENERAL MORPHOLOGY

Our observations of general morphology build on information included in Maas (1972) and Maas-van de Kamer et al. (2016). Following are some additional remarks.

### Ligule

The shape of the ligule, which is a quite important feature to distinguish between species both in the field and in the herbarium, is well illustrated in Maas-van de Kamer et al. (2016: fig. 1A). The length of the ligule is measured from the junction of the petiole to the sheath.

**Fig. 1.**
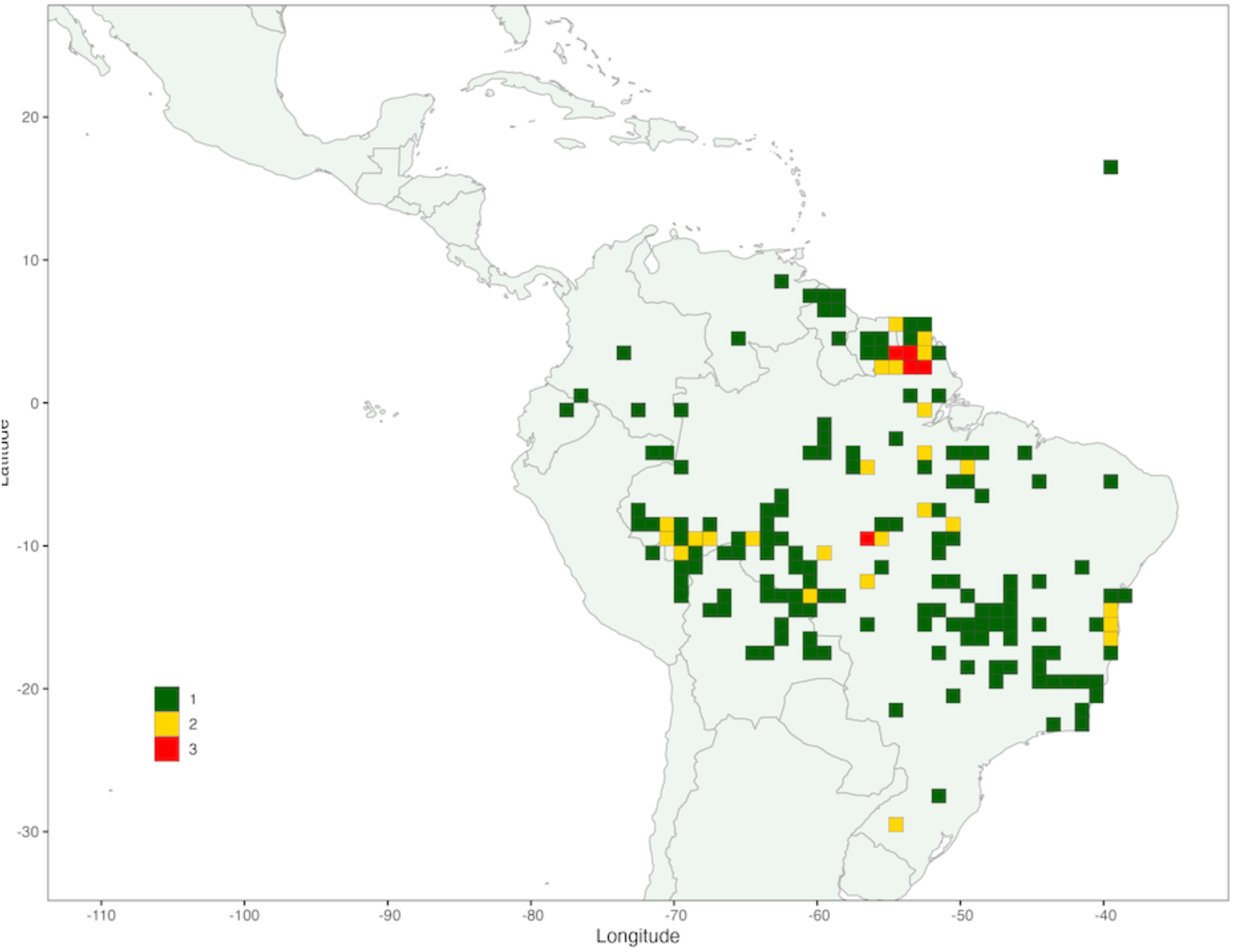
Species Richness of *Chamaecostus*. The maximum number of species occurring in a grid cell is 3 (red) with cells containing 2 (yellow) or 1 (green) species located throughout the distribution of the genus.

### Pedicel

In almost all species of neotropical *Costaceae* the flowers are sessile. Exceptions to this rule are *Chamaecostus cuspidatus* (Nees & Mart.) C.D.Specht & D.W.Stev. and *Monocostus uniflorus* (Poepp. Ex. Petersen) Maas, where a distinct pedicel is present.

### Leaves

Leaves of *Costaceae* comprise a sheathing leaf base, a petiole, and a leaf blade or lamina. The petiole of the *Zingiberales* is sometimes referred to as a “pseudopetiole”, however, given its position between the sheathing leaf base and the leaf lamina it is more appropriate to refer to it as a petiole. This is consistent with previous species descriptions specific to *Costaceae*. The petiole is continuous with the mid-rib of the leaf lamina.

The shape of the leaf lamina is generally narrowly elliptic to narrowly obovate, but there are also a few species that can be quickly identified by their narrow leaves (i.e. *Costus stenophyllus* Standl. & L.O.Williams, *Costus zingiberoides* J.F.Macbr.). While most species have leaves that are green on both upper and lower surfaces, some species of *Chamaecostus* and *Costus* have a leaf that is reddish to dark purple on the lower surface; this is never found in *Dimerocostus* or *Monocostus*. Another important leaf feature encountered in both *Chamaecostus* and *Costus* is the plication or undulation of the lamina, as is clearly shown in *Costus plicatus* Maas among others.

### Indument

Describing the indument of the plants continues to be quite difficult. Comparing the description of hairs observed in dried material to those observed in living material of the same species or even individual tends to cause problems: e.g. erect hairs can look appressed after drying. Moreover, and perhaps most critically, for some indument characters we observe variation within species that could range from glabrous to densely hairy and everything in between, especially in some of the more widespread species. While some species are quite consistent in indument, others tend to be more variable. In some cases, there are particular features of indument (e.g. cilate margin on a ligule) that can be consistent and used for species identification. These are noted throughout the monograph. Future work will focus on correlating indument variation within a given species to ecological or environmental features.

To describe the diversity and varation of indument characters, we use the following terminology:

**Puberulous (**including **minutely puberulous)**: erect to semi-appressed, soft, very short hairs < 0.5 mm long.

**Sericeous**: appressed, soft hairs, often somewhat silky to the touch, of variable length but longer than 0.5 mm.

**Velutinous**: erect, soft, very short hairs, < 0.5 mm long, forming a dense indument velvety to the touch.

**Villose**: erect, straight to more or less flexible, soft to stiff hairs 0.5–5 mm long; this includes the formerly used terms **hirsute** and **strigose**, as the difference between soft and stiff hairs is not always easy to distinguish.

**Scabrid**: appressed, very short hairs with thickened base, rough to the touch; this type is found only in a few species, e.g. *Costus glaucus* Maas and *Costus lima* K.Schum.

Apart from these terms, the colour and the density (sparsely, rather densely or densely) of the hairs are used in the description of the indument. When possible, we indicate when traits are consistent or are variable within a given species.

### Bracts

The bract types are clearly illustrated in fig. 3 in Maas (1972). The figure further demonstrates that in species with appendaged bracts, the callus (a vertical line of nectar secreting cells) is reduced to absent.

### Bracteole

The bracteole is almost always boat-shaped in the genus *Costus*, except for *C. osae* Maas & H.Maas and *C. ricus* Maas & H.Maas, where it is 2-keeled. In *Chamaecostus*, *Dimerocostus*, and *Monocostus*, the bracteole is tubular and typically 2-keeled at the adaxial side.

### Flowers

In most genera of *Costaceae* the flowers are oriented abaxially; i.e. the curvature and opening of the flower are directed away from the axis of the inflorescence. In *Costus*, however, there are a few species with flowers oriented adaxially to the effect that the open flower faces toward the axis of the inflorescence (e.g. *Costus spiralis* (Jacq.) Roscoe). The flower is not resupinate; instead, the floral organs grow such that the opening faces inward rather than outward.

### Calyx

Calyx and bracteole types are illustrated in fig. 4 of Maas (1972).

### Corolla

The corolla forms a tube held together by the calyx, with 3 lobes of variable size, the dorsal lobe being slightly larger than the 2 lateral lobes. The petals are not fused except perhaps at the base, having emerged from a ring primordium.

### Androecium

The androecium consists of 2 whorls (inner and outer) each with 3 androecial members. A labellum is formed from the fusion of 5 staminodes that arise from both the outer (3) and inner (2) androecial whorls (see Kirchoff 1988). When the labellum is lobed as in many species of neotropical *Costus*, the lobes emerge from the 3 outer androecial members such that there is one ventral and two lateral lobes. A single fertile stamen is laminar (not filamentous) and petaloid and is a member of the inner androecial whorl. The labellum and stamen together form a tube that is fused at the base. All measurements recorded in this monograph include only the free part of the labellum or the laminar stamen (above the fused section).

#### Labellum

The labellum of *Chamaecostus* is entire, broadly angular obovate to almost circular, horizontally spreading to reflexed, with dentate or fimbriate margin.

The labellum of *Costus* shows either melittophilous features or ornithophilous features. The melittophilous features include a horizontally spreading lip with recurved apical part and reddish striped lateral lobes together forming a landing platform and nectar guide, with the entry to the throat blocked by the thecae of the single fertile stamen. These flowers are generally pollinated by bees or other flying insects, with pollen deposition occurring on the dorsal surface of the bee. The labellum can also show ornithophilous features such as a tubular labellum without lateral markings and lacking a recurved apical surface (i.e. not forming a landing platform), and the entry to the throat is not blocked by the stamen. These flowers are generally pollinated by hummingbirds. Pollen may be deposited on the forehead or chin of the bird, depending on the orientation of the flower.

Some flowers are not distinctly melittophilous nor ornithophilous. The labellum in these species is funnel-shaped (e.g. *Costus atlanticus* E.Pessoa & M.Alves). The upper free part of the labellum (lip) is erect (not horizontally spreading) and the lateral lobes are often reddish striped and small.

The flower of *Costus guanaiensis* is different from all the other species of *Costus*. The labellum is erect, without distinct lateral lobes, with a relatively narrow recurved margin all around, and 4–6 distinct apical teeth. The labellum is not tubular, nor spreading nor funnel-shaped. The fertile stamen is relatively wide and closing the throat.

Some species of *Dimerocostus* have so-called “hooded” flowers, meaning that 1 erect large corolla-lobe forms a hood that covers the entrance to the flower (e.g. *D. rurrenabaqueanus* (Rusby) Maas & H.Maas and *D. argenteus* (Ruiz & Pav.) Maas). The lateral lobes of the labellum in this floral type are reflexed. The hooded corolla lobe and the labellum together create the appearance of a 2-lipped flower. In the other species of *Dimerocostus,* the 3 corolla lobes are about equal in length, the labellum is much wider, and the lateral lobes of the labellum are not reflexed.

The flowers of *Monocostus* have a horizontally spreading almost circular labellum similar to some of the species of *Chamaecostus*.

#### Laminar Stamen

The single fertile stamen has 2 thecae and is laminar (and petaloid) in all species. The apex of the stamen is irregularly dentate, acute or obtuse, extending above the position of the thecae. The stamen length can be such that the stamen distinctly exceeds the floral tube, as seen in *Costus pulverulentus* C.Presl (Figure 12c) and *C. fissicalyx* N.R.Salinas, Clavijo & Betancur B (Figure 8g).

### Stigma & Style

There are several stigma types illustrated in the additions to the Neotropical Monograph of *Costoideae* (Maas 1977: 162, fig. 62). The shape of the stigma of the genus *Costus* is 2-lamellate with a 2-lobed dorsal appendage that hooks between the 2 thecae of the stamen. In the other 3 genera, the stigma is funnel-shaped and the appendage is less developed. In all species the style is extremely thin and filamentous and is supported between the thecae of the fertile stamen.

### Fruit & Seeds

The fruits of all *Costaceae* are fibrous dry capsules, bilocular in *Monocostus* and *Dimerocostus*, and trilocular in *Chamaecostus* and *Costus*. They either dehisce loculicidally by 3 longitudinal slits or are indehiscent and decompose to reveal the seeds. Seeds are black or dark brown with a white aril.

## GEOGRAPHICAL DISTRIBUTION AND ECOLOGY

Neotropical *Costaceae* are distributed from Mexico and the Greater and the Lesser Antilles in the North to Bolivia, Argentina, Paraguay, and Southern Brazil in the South. The genus *Chamaecostus* occurs in tropical South America, with its main center of distribution located in Southern Amazonia: it is completely absent from Central America (Fig. 1). By far the largest and most widely distributed genus is *Costus*, occurring throughout the Neotropics. *Costus* is most common in Central America (Costa Rica and Panama) and the Pacific region of Colombia and Ecuador with a secondary center in the Guianas and adjacent Amazonian Brazil. *Costus* is found from Mexico (Sinaloa, c. 24⁰ N) in the North to Paraguay (the department of Paraguarí, c. 26⁰ S lat.) and Argentina (Misiones, c. 27⁰ 30’S) in the South (Fig. 2). Of the 4 genera occurring in the Neotropics, the monotypic genus *Monocostus* is very rare and restricted to just a few localities in Amazonian Peru (Fig. 3). *Dimerocostus* occurs in Central America (to Honduras in the North) and the western part of tropical South America (to Bolivia in the South) and reaches eastward to Venezuela and Suriname (Fig. 3). Species richness maps demonstrate centres of diversity for each genus (Figs 1–3).

**Fig. 2.**
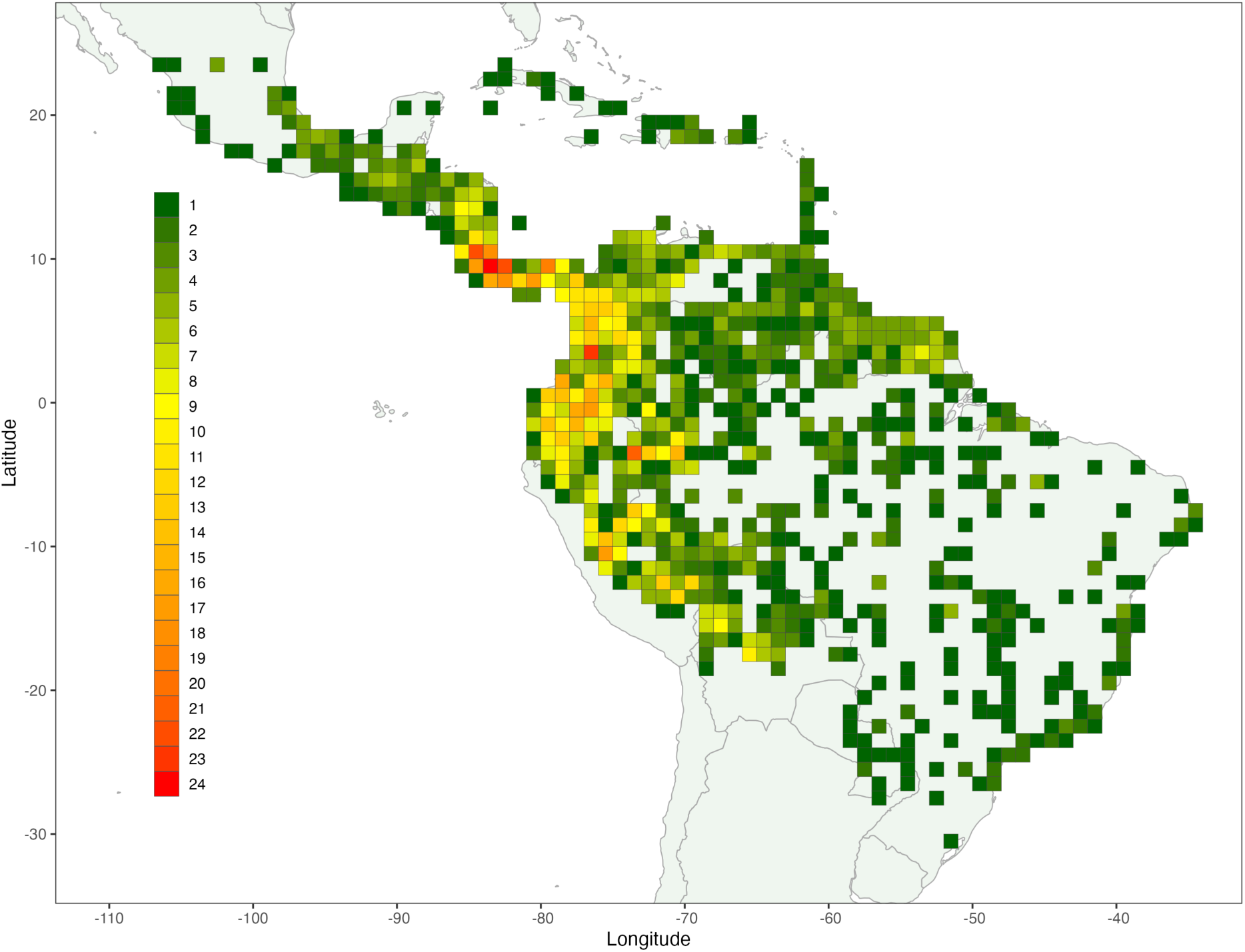
Species Richness of *Costus*. The maximum number of species occurring in a grid cell is 24 (red) with a gradient of richness down to a single species (green) located at the extremes of the distribution of the genus. The northern Andes and Central America harbour the greatest species richness for *Costus*.

**Fig. 3.**
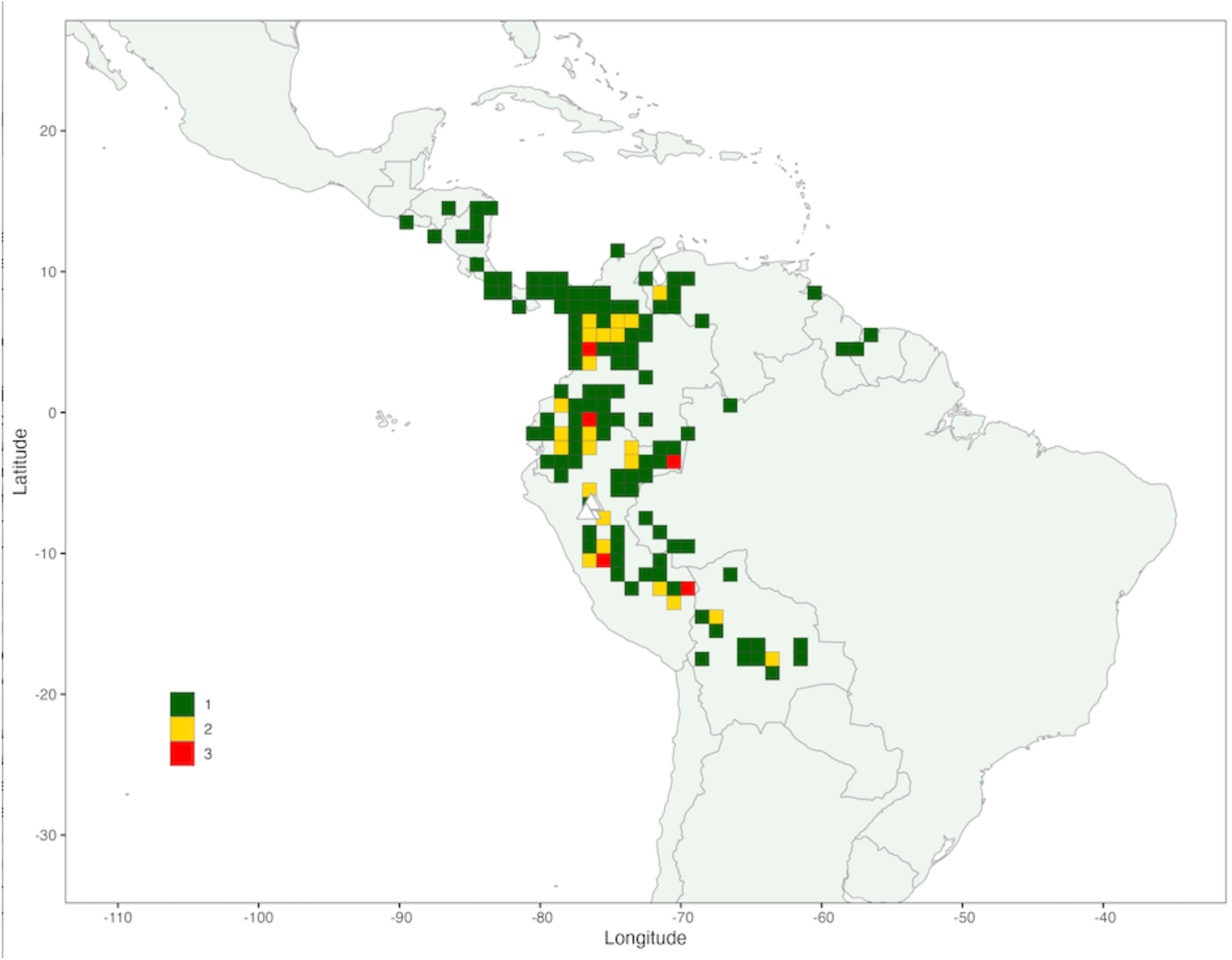
Species richness of *Dimerocostus* and the distribution of *Monocostus*. The maximum number of species occurring in a grid cell for *Dimerocostus* is 3 (red) with cells containing 2 (yellow) or 1 (green) species located throughout the distribution of the genus. The distribution of the monotypic genus *Monocostus* (*M. uniflorus*) is indicated with white triangles.

Most *Costaceae* inhabit various forest types, ranging from non-inundated terra firme rain forest to seasonally dry forests to submontane or even sometimes montane forest. A few species occur in periodically inundated forest, with *Costus varzearum* Maas providing a nice example. There are a few exceptions to this rule: for example, *Costus woodsonii* Maas inhabits sandy beaches and coastal dunes, and *Chamaecostus subsessilis* (Nees & Mart.) C.D.Specht & D.W.Stev., with its tuberous roots, can inhabit savannah vegetation or the understory of seasonally dry forest. Many species of *Costaceae* are pioneer species and are found in habitats that include secondary vegetation such as roadsides, creek margins, river margins, etc. By far the majority of the neotropical species of *Costaceae* occur at low elevations below 1500 m. The exceptions to this rule are the Central American species *Costus montanus* Maas and some populations of *C. wilsonii* Maas (occurring between 1000 and 1800 m), the Mexican *Costus alticolus* Maas & H.Maas (1200–1900 m), and the Peruvian species *Costus rubineus* D.Skinner & Maas (1400–2200 m). The “recordholder” is a specimen of *Costus guanaiensis* Rusby from the department of Cuzco, Peru, which according to the label of *Valenzuela et al. 10471* (MO) was collected at the amazing (incredible?) elevation of 2700 m!

## MATERIAL AND METHODS

Material from the following 127 herbaria was studied (for herbarium abbreviations see Thiers continuously updated): AAU, AMAZ, AS, B, BBS, BH, BHR, BISH, BM, BR, BRG, BRH, BRIT, C, CAS, CAY, CR, CEN, CEPEC, COAH, COL, CR, CUVC, CUZ, DAV, DUKE, E, EAP, ECON, ECUAMZ, ENAG, ENCB, F, FAUC, FMB, FTG, G, GB, GH, GOET, HB, HBR, HEM, HERZU, HLA, HOXA, HRCB, HSB, HUA, HULE, HUT, IAC, IAN, IBE, ICESI, IJ, INB, INPA, JAUM, JBSD, K, KIEL, L, LAM, LBB, LE, LIL, LOJA, LY, M, MA, MBM, MBML, MEDEL, MER, MEXU, MG, MICH, MISS, MO, MY, NA, NHA, NY, P, PH, PMA, PORT, PSO, PTBG, QAME, QAP, QCA, QCNE, R, RB, S, SEL, SMU, SP, TAMPA, TEFH, TRIN, U, UB, UC, UCWI, UDBC, UEC, UECU, UFP, UMO, UNA, UNAH, USCG, USJ, USM, USMS, USZ, USZU, VALLE, VEN, W, WAG, WIS, Z.

Living collections, which were essential for this study, were cultivated and maintained by the garden staff of Burgers’ Bush (the greenhouse of Burgers’ Zoo, Arnhem, The Netherlands), the greenhouses of the Utrecht University, Netherlands, and the Botanic Gardens at Meise, Belgium.

Additional field work was completed by Dave Skinner in Mexico, Guatemala, Honduras, Costa Rica, Panama, Colombia, Ecuador, Peru, Bolivia, Brazil, and Guyana. Photos and detailed measurements in the field were taken and used in part for the individual species descriptions. Visits were made to the type localities of several species. These observations have been recorded at https://www.inaturalist.org/observations/selvadero.

Conservation assessments include published IUCN assessments for 19 neotropical species, each formally assessed by D.Skinner in 2014 and 2015 (https://www.iucnredlist.org/search?query=Costus&searchType=species). For those species where no formal assessment is available, preliminary assessments were completed by D.Skinner combining data from our list of specimens examined with recent field work and verified observations recorded on iNaturalist (https://www.inaturalist.org/).

Species distribution maps were generated in RStudio version 2023.12.1+402 (https://dailies.rstudio.com/version/2023.12.1+402/). The required R packages, including “raster”, “ggplot2”, “maptools”, “ggspatial”, “sf”, and “ggnewscale” were installed. A base world map was constructed using functions within these packages. Subsequently, elevation data was projected onto the map using SRTM elevation data obtained from WorldClim.org. Occurrence data representing specimen collection sites were then plotted onto the map to visualize the distribution of each species. The “ggplot2” package was utilized for data visualization, allowing for the customization of the aesthetics of the map such as political boundaries and axis markings.

Species richness maps were generated using the same version of RStudio. In addition to the packages that were necessary to generate the species distribution maps, “dplyr”, “sp”, and “rgdal” were installed to facilitate the creation of species richness maps. After the packages were installed, a base world map was constructed using functions within the packages. Species richness was calculated per 1° grid cell, beginning with reading in the *Costaceae* occurrence data. To calculate species richness from the occurrence data, it was filtered by genus, longitudinal and latitudinal boundaries were established, and the coordinates of the occurrences data were rounded and centered to group the points. The grouped points were used to calculate the number of species within 1° grid cells. After computing richness, “ggplot2” was used to overlay the calculated species richness onto the world map and to make aesthetic customizations.

## TAXONOMIC TREATMENT

### *Costaceae* Nakai

#### *Costaceae* Nakai (1941) 203. — Type: *Costus* L

Perennial, rhizomatous, terrestrial, non-aromatic, tall, gigantic, low, or acaulescent herbs; shoot erect, terete, straight or spirally contorted, unbranched or rarely with leafy shoots arising from the base of the inflorescence or from the axils of the leaves. *Leaves* many to few, spirally arranged along the shoot, shootless species with few rosulate leaves (*Chamaecostus subsessilis*); leaf sheaths closed around the shoot, tubular; ligule present, truncate to 2-lobed at the apex; petiole (region between sheathing leaf base and lamina) short; lamina generally more or less elliptic in outline, sometimes shiny or plicate, with mostly acuminate apex and acute, rounded, or rarely cordate base, glabrous or hairy. *Inflorescence* a many- to few-flowered spike, terminating a leafy shoot or terminating a separate, short leafless shoot, or flowers solitary in the axils of the upper leaves (*Monocostus*); bracts imbricate, spirally arranged, each carrying 1 flower, coriaceous, sometimes herbaceous or chartaceous, mostly with a linear, nectariferous callus below the apex; foliaceous, apical appendages of bracts absent or present, mostly ascending to horizontally spreading; each flower enclosed by one boat-shaped or tubular and 1–2-keeled bracteole. *Flowers* epigynous, perfect, zygomorphic, abaxially or rarely adaxially orientated, sessile, very rarely pedicellate; calyx tubular, shortly 3-lobed; corolla tubular at the base, tube equalling the calyx, 3-lobed, lobes imbricate, the dorsal lobe somewhat larger than the 2 lateral ones; *labellum* petaloid, equalling or often much exceeding the corolla, more or less 3-lobed, either large and horizontally flattened, broadly obovate when spread out, middle lobe often recurved, with a yellow honey mark, lateral lobes erect and very large; or *labellum* small, tubular, narrowly oblong-obovate when spread out, lateral lobes small, rolled inwards and forming a tube; *stamen* 1, laminar and petaloid, anther usually attached at the middle with 2 longitudinally dehiscing thecae; base of stamen and labellum joined into a tube; *style* 1, filiform, supported between the thecae of the anther; s*tigma* 1, 2-lamellate, provided with a 2-lobed dorsal appendage, or unappendaged and funnel-shaped, margin ciliate; septal nectary glands 2, at the apex of the ovary, secreting nectar in the base of the flower tube; *ovary* inferior, 3-locular (*Chamaecostus*, *Costus*) or 2-locular (*Dimerocostus*, *Monocostus*), ovules many, placentation axillary, organized in 1 or 2 rows per locule, anatropous. *Fruit* a 3-locular (*Chamaecostus*, *Costus*) or 2-locular (*Dimerocostus*, *Monocostus*) capsule, crowned by the persistent calyx, dehiscing loculicidally by 3 longitudinal slits, or indehiscent and irregularly breaking when old. *Seeds* many, irregularly angular, reflecting tight packing in fruit, black to brown, shiny; aril white, lacerate or cushion-like; embryo straight, in copious endosperm.

## KEY TO THE NEOTROPICAL GENERA OF *COSTACEAE*

1. Flowers solitary in the axils of the upper leaves 4. *Monocostus*

1. Flowers subtended by bracts arranged in a many-flowered spike 2

2. Ovary 2-locular 3. *Dimerocostus*

2. Ovary 3-locular 3

3. Plant mostly < 1 m tall, sometimes acaulescent; bracts herbaceous to chartaceous, green to yellow; bracteole tubular, 2-keeled at the adaxial side; stigma cup or funnel-shaped 1. *Chamaecostus*

3. Plant > 1 m tall; bracts coriaceous, red, orange, yellow to green; bracteole boat-shaped, very rarely 2-keeled; stigma 2-lamellate 2. *Costus*

**1.** ***Chamaecostus*** C.D.Specht & D.W.Stev.

*Chamaecostus* C.D.Specht & D.W.Stev. (2006) 157. — Type: *Chamaecostus subsessilis* (Nees & Mart.) C.D.Specht & D.W.Stev.

Low to acaulescent herbs, often with tuberous roots and thick stem nodes. *Leaves:* ligule short, truncate to 2-lobed; lamina generally narrowly ovate to narrowly obovate, sometimes slightly plicate to undulate, lower side sometimes purplish red, base acute to obtuse, rarely cordate, apex shortly acuminate to acute. *Inflorescence* spike-like to capitate, terminating a leafy shoot, or rarely terminating a separate, short, leafless shoot. *Flowers* abaxially orientated, sessile or becoming pedicellate in fruit; bracts herbaceous to chartaceous, green or yellow, generally ovate to elliptic or narrowly so, sometimes provided with foliaceous, apical appendages; bracteole tubular and/or abaxially split with age, 2-keeled at the adaxial side; calyx small, tubular; corolla white, yellow, orange, or purple-red, not or rarely exceeding the bracts; labellum white, yellow, orange, pink, or purple-red, large, entire, spreading to reflexed, margin irregularly dentate, sometimes fimbriate; stamen petaloid with the anther attached at the center of the upper half; stigma funnel-shaped; ovary 3-locular, ovules organized in 2 rows per locule. *Capsule* 3-locular, mostly ellipsoid, membranous, tardily dehiscent. *Seeds* with a lacerate aril.

Distribution — A genus with 9 species restricted to tropical South America.

Habitat & Ecology — Forest understory, terrestrial or rupicolous, usually at shallow soils with rocky outcrops, particularly on calcareous soils and in riparian vegetation along streams; most species have tuberous roots and show intermittent growth following rain seasonality, mostly without living shoots throughout driest months of the year.

## KEY TO THE SPECIES

1. Bracts appendaged, or their apices reflexed, recurved or ascending 2

1. Bracts not appendaged 5

2. Calyx 1–1.1 cm long; bracts yellow to orange. — French Guiana 3. *C. curcumoides*

2. Calyx 0. 6–4.8 cm long; bracts green 3

3. Flowers orange, rarely red-orange; anther irregularly dentate at the apex. — Atlantic Forests of Eastern Brazil 4. *C. cuspidatus*

3. Flowers yellow or whitish; anther obtuse or irregularly lobulate at the apex 4

4. Plants acaulescent to very low (< 0.25 m tall); leaf blades oblong-obovate, upper side densely puberulous. — Cerrados from Western Brazil and seasonally dry Amazonian forests of Brazil, Bolivia and Peru 1. *C. acaulis*

4. Plants caulescent (0.25–1 m tall), erect; leaf blades elliptic, upper side sparsely puberulous to glabrous. — Cerrados from Eastern Brazil and Atlantic Coastal Forest 9. *C. subsessilis*

5. Inflorescence terminating a separate leafless shoot; bracts obovate to ellipic. — Amazonian Brazil, Colombia, and Ecuador 5. *C. fragilis*

5. Inflorescence terminating a leafy stem; bracts ovate to narrowly triangular 6

6. Bracts imbricate, yellow, 1–2 cm wide. Flowers yellow. — Amazonian Brazil and Peru, and Suriname 6. *C. fusiformis*

6. Bracts non-overlapping, green or orange, 0.5–1.2 cm wide. Flowers white, red, pink, or orange 7

7. Flowers pink, red or orange; corolla glabrous to puberulous. — Throughout the Amazon Region, French Guiana and Suriname 7. *C. lanceolatus*

7. Flowers white; corolla puberulous to villose8

8. Labellum with fimbriate margin; inflorescence broadly obovoid, bracts 0.5–1.2 cm wide. — Amazonian Brazil and Venezuela, and the Guianas 2. *C. congestiflorus*

8. Labellum with crenulate margin; inflorescence narrowly cylindric; bracts 0.3–0.5 cm wide. — Central Amazonia, Brazil 8. *C. manausensis*

**1.** ***Chamaecostus acaulis*** (S.Moore) T.André & C.D.Specht — Fig. 4a; Map 1

**Fig. 4.**
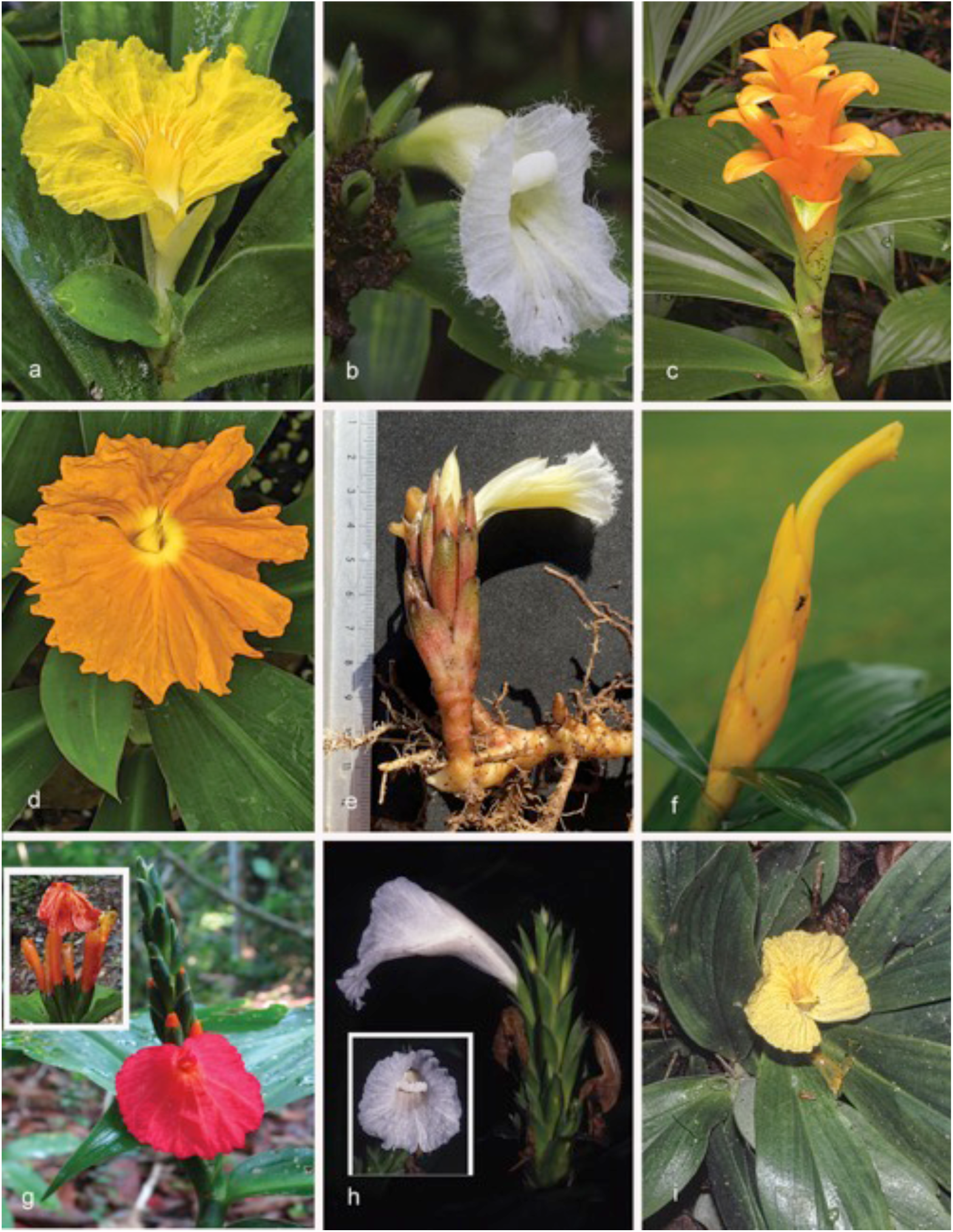
Species of *Chamaecostus*: a. *Ch. acaulis* (S.Moore) T.André & C.D.Specht (*D. Skinner R3391*, Santa Cruz, Bolivia); b. *Ch. congestiflorus* (Rich. ex Gagnep.) C.D.Specht & D.W.Stev. (photo only, French Guiana); c. *Ch. curcumoides* (Maas) C.D.Specht & D.W.Stev. (*Gonzalez 2133* from French Guiana, Mont Itoupé-Sommet Tabulaire); d. *Ch. cuspidatus* (Nees & Mart.) C.D.Specht & D.W.Stev. (*D. Skinner R3301*, Espirito Santo, Brazil); e: *Ch. fragilis* (Maas) C.D.Specht & D.W.Stev. (*Anonymous s.n*., near Leticia, Colombia); f: *Ch. fusiformis* (Maas) C.D.Specht & D.W.Stev. (*Zappi 896*, Brazil, Mato Grosso, Parque Estadual Cristalino); g. *Ch. lanceolatus* (Petersen) C.D.Specht & D.W.Stev. subsp. *lanceolatus* (*D. Skinner R3249*, Alta Floresta, Brazil (inset of subsp. *pulchriflorus* (Ducke) C.D.Specht & D.W.Stev.) (*C.Bhikhi 1239*, Surinam, Sipaliwini area, Kasikasima) h. *Ch. manausensis* Maas & H.Maas (*J. Miralha 296*, Brazil, Amazonas, Reserva Ducke, Manaus-Itacoatiara); i. *Ch. subsessilis* (Nees & Mart.) C.D.Specht & D.W.Stev. (*D.Skinner R3247*, Brazil, DF, Brasilia). Photos a, d, g & i by D. Skinner; b & h by P.J.M. Maas; c by S. Gonzalez; e by A. Colmenares; f by W. Milliken; g inset by C. Bhikhi.

*Chamaecostus acaulis* (S.Moore) T.André & C.D.Specht in André et al. (2015) 272 — *Costus acaulis* S.Moore (1895) 480, t. 33, f. 1–5. — Type: *S. Moore 679* (holo BM! [BM000923831]; iso F! [F0BN009889]), Brazil, Mato Grosso, Santa Cruz, Nov. 1891.

*Costus kaempferoides* Loes. (1929) 714. — Lectotype (designated by Maas 1972: 30): *Ule 9197* (holo B, destroyed; lecto K [K000586760]; isolecto F [F0BN009899], G [G00168446]), Peru, Madre de Dios, Seringal São Francisco, Sept. 1911.

*Costus steinbachii* Loes. (1929) 714. — Lectotype (designated by Maas 1972: 30): *Steinbach 7386* (holo B, destroyed; lecto F [F0047183F]; isolecto BM [BM000923832], G [G00168413], GH [GH00030662], GOET [GOET011649], K [K000586761], LIL [LIL000233], MO [MO-1515243], S [S-R-1263]), Bolivia, Santa Cruz, Prov. Sara, Buenavista, 31 Dec. 1925.

Acaulescent to very short-stemmed plants (< 25 cm tall), roots fleshy, ending in fusiform to ellipsoid tubers. *Leaves:* 4–6, rosulate: sheaths membranous, obtuse at the apex, 1.5–8.5 by 0.7– 3.0 cm, densely hirsute, sparsely to rather densely puberulous; ligule < 1 mm long; petiole absent; lamina oblong-obovate, 3–36 by 4–15 cm, densely puberulous on both sides, base acute, apex mucronate or shortly acuminate (acumen 0–1.8 cm long). *Inflorescence* terminal on a leafy shoot, formed in the centre of the rosette, 1–4-flowered; bracts, appendages of bracts, bracteole, pedicels, and calyx glabrous to densely puberulous, ovary densely sericeous to glabrous; bracts green, herbaceous, 3–7 by 0.6–1.3 cm, appendages green, foliaceous, narrowly triangular to deltate, apex mucronate; bracteole tubular, 2–3 cm long, 2-lobed at the apex, lobes narrowly triangular to deltate, 2–15 mm long. *Flowers:* pedicels c. 5 mm long; calyx 0.1–1.2 cm long, lobes narrowly triangular to deltate, 1–20 mm long, mucronate; corolla yellow, 55–70 mm long, glabrous to densely puberulous, lobes narrowly elliptic, 30–40 by 6–12 mm; labellum yellow to pale yellow, striped with orange in the throat, broadly obovate when spread out, 60–70 by 70–95 mm, margin irregularly crenulate; stamen yellow, 20–50 by 6–20 mm, apex reflexed, obtuse or irregularly lobulate, anther 7–8 mm long. *Capsule* unknown.

Distribution — Peru (Madre de Dios, Ucayali), Bolivia (Beni, Pando, Santa Cruz), Brazil (Acre, Goiás, Mato Grosso, Minas Gerais, Pará, Rondônia, São Paulo, Tocantins).

Habitat & Ecology — In non-inundated (terra firme) forests of Amazonia or in the understory of gallery forests of Cerrado savannas, at elevations of 0–1000 m. Flowering in the rainy season, mainly from October to January across most of its range.

IUCN Conservation Status — Not evaluated. There are 80 collection records examined of which all specimens were collected within the past 50 years. This species is also often reported in iNaturalist recent observation data (https://www.inaturalist.org/observations?place_id=any&taxon_id=537965). The species is widely distributed in Brazil, Peru and Bolivia. Our observations indicate that this is a rather common species within its range and some of these areas are protected areas with no known specific threats. Population and threat data is not available but based on the number of collection records and our general field observations, this species would probably score a status of Least Concern.

Vernacular names — Bolivia: Pathu chico de flor grande. Brazil: Ara’iã, Campo alegre, Cana de macaco, Cana fistula, Flor de ouro, Sororoca baixinha.

Notes — *Chamaecostus acaulis* is very similar to *C. subessilis*, but it is distinguished by its oblong-obovate leaf blades instead of elliptic, with upper side densely puberulous instead of sparsely puberulous to glabrous, and usually very low to acaulescent habit. See André et al. (2015) for a discussion on these populations.

2. ***Chamaecostus congestiflorus*** (Rich. ex L.F.Gagnep.) C.D.Specht & D.W.Stev. — Fig. 4b; Map 2

*Chamaecostus congestiflorus* (Rich. ex Gagnep.) C.D.Specht & D.W.Stev. (2006) 158. —

*Costus congestiflorus* Rich. ex Gagnep. (1902) 97. — Lectotype (designated here): *G.W. Martin s.n.* (lecto P [P00686613]; isolecto BM [BM000923833], P [P00686614]), French Guiana, unknown locality.

*Costus congestiflorus* Rich. ex Gagnep. var. *glabrior* K.Schum. (1904) 401. — Syntype: *Poiteau s.n.* (K [K000586751]), French Guiana, unknown locality, Jul. 1824.

Low herbs, 0.5–1 m tall. *Leaves:* sheaths 5–10(–15) mm diam; ligule 1–2 mm long; petiole 1–5 mm long; sheaths, ligule and petiole glabrous to densely puberulous; lamina narrowly elliptic to narrowly obovate, 5–8-undulate, 10–20 by 2.5–7 cm, upper side glabrous, lower side often purplish red, glabrous to densely puberulous, base acute, obtuse, rarely cordate, apex acuminate (acumen 5–15 mm long). *Inflorescence* broadly obovoid, 3–10 by 1.5–4 cm, terminating a leafy shoot; bracts, bracteoles, calyx, ovary, and capsule glabrous to densely puberulous; bracts green, chartaceous, narrowly triangular to triangular, often mucronate, 1.5–3.5 by 0.5–1.2 cm, callus 2– 7 mm long; bracteole tubular, often deeply split on the abaxial side, 13–20 mm long, 2-lobed, lobes deltate, 2–3 mm long. *Flowers:* calyx green, (20–)25–35 mm long, lobes subulate, 1–10 mm long, tube often deeply split by the flower bud during anthesis; corolla cream, 50–65 mm long, densely puberulous to villose, lobes narrowly obovate, 30–40 mm long; labellum cream, throat yellow, upper part horizontally spreading, broadly angular-obovate, c. 35 by 40 mm, margin distinctly fimbriate; stamen cream, c. 25 by 7–12 mm, apex reflexed, irregularly dentate to entire, anther 6–8 mm long. *Capsule* ellipsoid, 10–15 mm long.

Distribution — Venezuela (Delta Amacuro), Guyana, Suriname, French Guiana, Brazil (Amapá, Amazonas, Pará).

Habitat & Ecology — In lowland rain forests, on sandy, lateritic or clayey soils, sometimes growing on granitic rocks, at elevations of 0–500 m. Flowering during the rainy season with peak flowering in third and fourth months of the rainy season.

IUCN Conservation Status — Not evaluated. There are 100 collection records examined of which 71 specimens were collected within the past 50 years. They are widely distributed in Brazil, French Guiana, Guyana, Suriname, and one specimen in Venezuela. Population and threat data is not available but it is not a common species and based on the number of collection records would probably score a status of Near Threatened or Vulnerable.

Vernacular names — French Guiana: Ago singa afu (Boni), Canne congo (Créole), Canne l’eau (Créole), Kapiya (Wayãpi), Kapiya-pila (Wayãpi), Kapiya-si (Wayãpi), Pikin singa afu (Boni), Yapusĩ (Wayãpi). Guyana: White congo cane. Suriname: Sangrafoe.

Notes — *Chamaecostus congestiflorus* is recognizable by having white flowers, a distinctly fimbriate labellum and hairy petals. From *C. fragilis* it differs by chartaceous instead of membranous bracts and by an inflorescence terminating a leafy shoot instead of a separate leafless shoot. For differences with another white-flowered species of *Chamaecostus*, namely *C. manausensis* see under that species.

3. ***Chamaecostus curcumoides*** (Maas) C.D.Specht & D.W.Stev. — Fig. 4c; Map 3

*Chamaecostus curcumoides* (Maas) C.D.Specht & D.W.Stev. (2006) 158. — *Costus curcumoides* Maas (1972) 34, f. 17. — Type: *Oldeman B.553* (holo P [P00686615]; iso P [P00686616]), French Guiana, Crique Anis, near Mapaou, Approuague River, 11 June 1966.

Low herbs, 0.4–0.7 m tall. *Leaves:* sheaths 3–9 mm diam; ligule 1–2 mm long; petiole 0–5 mm long; sheaths, ligule and petiole glabrous; lamina narrowly elliptic to narrowly obovate, 5–6- plicate, 5–22 by 3–8 cm, upper side glabrous, lower side sparsely puberulous to glabrous, base acute, apex acuminate (acumen 5–10 mm long). *Inflorescence* cylindric, 4–7 by 1–3 cm, terminating a leafy shoot; bracts, bracteole, calyx, ovary, and capsule glabrous to sparsely puberulous; bracts yellow to orange, membranous, narrowly ovate-triangular to ovate-triangular, 2.5–7 by 1–4.5 cm, apical part recurved or ascending, triangular, 3–20 by 5–12 mm, callus obscure, 0.5–5 mm long; bracteole 4–8 mm long, 2-lobed, lobes deltate, 1–2 mm long*. Flowers:* calyx colour not recorded, 10–11 mm long, lobes deltate, 1–2 mm long; corolla yellow to orange, glabrous, c. 60 mm long, glabrous, lobes narrowly elliptic, c. 45 mm long; labellum yellow to orange, lateral lobes rolled inwards and forming a curved tube of c. 8 mm diam, oblong-obovate when spread out, 6–8 by 6–8 mm (in bud); stamen orange to yellow, 8–10 mm long, apex orange, obtuse, anther 4–5 mm long (in bud). *Capsule* ellipsoid, 10–14 mm long.

Distribution — French Guiana.

Habitat and Ecology — In non-inundated primary or secondary forests, on lateritic clay, one collection “growing on granite”, at elevations of 0–700 m. Flowering only in the rainy season with peak flowering in February to April.

IUCN Conservation Status — Not evaluated. There are 18 collection records examined of which 16 specimens were collected within the past 50 years. They are narrowly distributed in French Guiana. Population and threat data is not available but it is considered a very rare species, and based on the small known extent of occupancy (EOO) from a single collection site, this species is likely to be Vulnerable or possibly Endangered.

Vernacular names — French Guiana: Canne congo (Créole), Yapusĩ (Wayãpi).

Notes — *Chamaeostus curcumoides*, endemic to French Guiana, is unique in the genus by its yellow to orange bracts, with recurved or ascending tips, resembling those of the genus *Curcuma* (*Zingiberaceae*).

4. ***Chamaecostus cuspidatus*** (Nees & Mart.) C.D.Specht & D.W.Stev. — Fig. 4d; Map 3

*Chamaecostus cuspidatus* (Nees & Mart.) C.D.Specht & D.W.Stev. (2006) 158. — *Globba cuspidata* Nees & Mart. (1823) 28. — *Costus cuspidatus* (Nees & Mart.) Maas (1976) 469. — Syntype: *Wied zum Neuwied s.n.* (BR [BR0000005188741]), Brazil, Bahia, “Ad ripas fluv. Ilhéos circa viam Felisbertiam”, anno 1817.

*Costus igneus* N.E.Br. (1884) 25, pl. 511. — Type: *Hort. Kew., January 1844* (holo K [K000586749]), cultivated from seeds sent by the Compagnie Continentale d’Horticulture Gand, Belgium, Jan. 1884, from Bahia, Brazil.

Low herbs, 0.2–0.5 m tall, roots fleshy, somewhat thickened at the end. *Leaves:* sheaths 5–10 mm diam; ligule 1–2 mm long; petiole 1–3 mm long; sheaths, ligule and petiole glabrous to sparsely puberulous; lamina narrowly elliptic to narrowly obovate or ovate to elliptic, 8–18 by 3– 6(–7.5) cm, upper and lower side glabrous to sparsely puberulous, base acute, apex acuminate (acumen 5–10 mm long), ending in a 1–3 mm long filiform point. *Inflorescence* 3–8-flowered, ellipsoid, 2–4 by 1.5–2 cm, terminating a leafy shoot; bracts, appendages of bracts, bracteole, and pedicels glabrous to sparsely puberulous, calyx, ovary, and capsule densely puberulous; bracts pale green, herbaceous, ovate-elliptic, 1–2 by 0.5–1 cm; appendages green, foliaceous, narrowly triangular to deltate, 2–11 by 1.5–4.5 cm, apex acute; bracteole tubular, often deeply split on the abaxial side, 25–30 mm long, 2-lobed at the apex, lobes narrowly triangular to deltate, 2–8 mm long. *Flowers*: pedicels fleshy, green, 1–5 by 3–5 mm, elongating up to c. 30 mm in fruit; calyx pale green, 30–40 mm long, lobes triangular, 6–10 mm long, usually with a 2– 3 mm long callus; corolla orange, c. 60 mm long, glabrous to sparsely puberulous, lobes narrowly ovate-elliptic, c. 35 mm long; labellum orange, throat yellow, upper part horizontally spreading, suborbicular, 50–70 by 50–70 mm, margin irregularly dentate; stamen yellow, c. 15 by 5–8 mm, apex orange, reflexed, irregularly dentate, anther 5–7 mm long. *Capsule* ellipsoid, c. 15 mm long.

Distribution — Southeastern and Northeastern Brazil (Bahia, Espírito Santo, Minas Gerais, Rio de Janeiro).

Habitat & Ecology — In Atlantic rain forests, at elevations of 0–200 m. Flowering primarily in the rainy season.

IUCN Conservation Status — Not evaluated. There are 29 collection records examined of which 15 specimens were collected within the past 50 years. They are narrowly distributed in the eastern states of Brazil (Bahia, Espirito Santo, and Minas Gerais). The Brazilian authorities have listed this as an endangered species so the scoring of its Global Red List status would probably be one of the endangered categories.

Vernacular names — Brazil: Costus de fogo.

Notes — *Chamaecostus cuspidatus*, restricted to Southeastern and Northeastern Brazil, is much cultivated all over the world for its showy orange flowers. In two collections from Bahia, Brazil, however, the flower colour is not given as orange, but as red (“vermelho”) (*dos Santos 1654*) and golden yellow (“amarelo-ouro”) (*dos Santos 893*), respectively.

A typical feature of this species is the often presence of ellipsoid to ovoid bulbils developing in the axil of the lowermost bracts.

In all species of *Chamaecostus* and *Costus* the flowers are sessile, but *C. cuspidatus* is unique as the ripe fruit is clearly elevated by a distinctly long pedicel (up to 30 mm).

5. ***Chamaecostus fragilis*** (Maas) C.D.Specht & D.W.Stev. — Fig. 4e; Map 3

*Chamaecostus fragilis* (Maas) C.D.Specht & D.W.Stev. (2006) 158. – *Costus fragilis* Maas (1972) 37, f. 19. — Type: *Ducke RB 14127* (holo RB, 3 sheets [RB00567441, RB00544556, RB00568979]), Brazil, Amazonas-Pará, Cachoeira da Montanha, Rio Tapajós, 18 Dec. 1919.

Low herbs, 0.15–0.45 m tall; rhizomes 3–5 mm thick, roots fusiform, 20–80 by c. 2 mm. *Leaves:* sheaths brownish red, membranous, 3–8 mm diam; ligule 1–2 mm long; petiole 0–5 mm long; sheaths, petiole, and ligule glabrous to rather densely puberulous; lamina narrowly elliptic to narrowly obovate, 10–22 by 3–6 cm, upper side glabrous, lower side glabrous to rather densely puberulous (particularly along the margin and primary vein), base acute, apex acute to acuminate (acumen 10–20 mm long). *Inflorescence*: cylindric, 4–18 by 2–4 cm, terminating a separate, leafless shoot, 3–30 cm long, sheaths obliquely truncate, membranous, 5–40 mm long, sparsely puberulous; bracts, bracteole, calyx, ovary, and capsule glabrous to rather densely puberulous; bracts green, greenish pink or red, membranous, obovate to elliptic, 1.5–3.5 by 0.6–1.7 cm, apex obtuse to acute and mucronate, callus 1–5 mm long, apex and margin falling apart into fibers; bracteole tubular, 6–10 mm long, shortly 2-lobed, lobes 1–3 mm long. *Flowers*: calyx (reddish) green, 20–33 mm long, often unilaterally split and unequally 3-lobed, lobes triangular to deltate, 3–8 mm long, mucronate; corolla white to yellowish white, 50–60 mm long, glabrous, lobes narrowly obovate, 20–30 mm long; labellum white, throat yellow, upper part horizontally spreading, broadly obovate to transversely broadly elliptic, 30–40 by 30–50 mm, margin irregularly dentate and distinctly fimbriate; stamen white, not measurable, anther 7–8 mm long. *Capsule* ovoid to narrowly ovoid, 17–23 mm long.

Distribution — Colombia (Amazonas, Caquetá, Vaupés), Ecuador (Sucumbios), Brazil (Amazonas, Pará).

Habitat and Ecology — In non-inundated (terra firme) rain forests, on clayey soil, at elevations up to 265 m. Flowering in months with high rainfall.

IUCN Conservation Status — Not evaluated. There are 15 collection records examined of which 11 specimens were collected within the past 50 years. They are distributed in Brazil, Colombia and Ecuador. Population and threat data is not available but based on the number of collection records this species is likely either Near Threatened or Vulnerable.

Notes — *Chamaecostus fragilis* is easily distinguished from all other species of *Chamaecostus* by an inflorescence terminating a separate leafless shoot, membranous bracts, and a relatively long calyx up to 33 mm long. It shares the white and fimbriate labellum with *C. congestiflorus*.

6. ***Chamaecostus fusiformis*** (Maas) C.D.Specht & D.W.Stev. — Fig. 4f; Map 3

*Chamaecostus fusiformis* (Maas) C.D.Specht & D.W.Stev. (2006) 158. — *Costus fusiformis* Maas (1972) 37, f. 18. – Type: *J.G. Kuhlmann 1916* (holo U [U0007241]; iso RB [RB00544557, RB00568981, RB00567442]), Brazil, Pará, Varadouro de Periquito, near Pimental, Rio Tapajós, 5 Apr. 1924.

Low herbs, 0.75–1 m tall, rhizomes with runners up to 50 cm by 1–3 mm, beset with membranous sheaths 5–10 cm long. *Leaves*: sheaths membranous, 3–10 mm diam; ligule 1–3 mm long; petiole 2–4 mm long; sheaths, ligule and petiole glabrous to sparsely puberulous; lamina narrowly obovate to narrowly elliptic, 10–23 by 4–8 cm, 5–8-undulate, upper side glabrous, lower side rather densely puberulous (particularly along the primary vein) to glabrous, base obtuse to acute, apex acuminate (acumen 5–20 mm long). *Inflorescence* fusiform, 6–11 by 1–2 cm, terminating a leafy shoot; bracts, bracteole, calyx, ovary, and capsule glabrous to sparsely puberulous; bracts yellow, chartaceous, ovate-triangular, 2–3.5 by 1–2 cm, apex obtuse to acute, sometimes mucronate, callus 3–7 mm long; bracteole tubular, 5–8 mm long, 2-lobed, lobes 1–2 mm long. *Flowers*: calyx yellow, 11–15 mm long, lobes deltate, c. 1 mm long; corolla yellow, c. 60 mm long, glabrous, lobes narrowly elliptic, c. 20 mm long; labellum yellow, lateral lobes rolled inwards and forming a curved tube; stamen not seen. *Capsule* narrowly ellipsoid, 11–15 mm long.

Distribution — Suriname, Brazil (Amazonas, Mato Grosso, Pará), Peru (Loreto). Habitat & Ecology— In non-inundated (terra firme) forests, at elevations up to 150 m. Flowering late in the rainy season.

IUCN Conservation Status — Not evaluated. There are 13 collection records examined of which 9 specimens were collected within the past 50 years. All recent specimens are narrowly distributed in the Mato Grosso and Pará states of Brazil. Population and threat data is not available but based on the number of collection records this species is likely either Near Threatened or Vulnerable.

Vernacular names — Brazil: Caninho-do-brejo. Peru: Puca Urquillo (Witoto).

Notes — *Chamaecostus fusiformis* is easily distinguished from all other species of the genus by a fusiform and yellow inflorescence and the presence of superficial runners.

7. ***Chamaecostus lanceolatus*** (Petersen) C.D.Specht & D.W.Stev. — Fig. 4g; Map 4

*Chamaecostus lanceolatus* (Petersen) C.D.Specht & D.W.Stev. (2006) 158. — *Costus lanceolatus* Petersen (1890) 56. — Lectotype (designated here): *L.C. Richard s.n.* (lecto [P00686617]), Brazil, locality unknown.

*Costus phlociflorus* Rusby (1902) 694. — Lectotype (designated here): *Rusby 2229* (lecto NY [NY00320342]; isolecto F [F0047177F], GH [GH00030656], NY [NY00320340, NY00320343], PH [PH00006741], US [US00340637, US00092970]), Brazil, Amazonas, Rio Madeira near falls, Oct. 1886.

Low herbs, 0.3–0.5 m tall. *Leaves*: sheaths 3–8 mm diam; ligule 1–4 mm long; petiole 2-4 mm long; sheaths, ligule and petiole glabrous to rather densely puberulous; lamina narrowly elliptic to narrowly obovate, 6–20(–27) by 2–7 cm, upper side glabrous, lower side sometimes purplish red, rather densely puberulous, base acute to obtuse, apex acuminate (acumen 5–20 mm long). *Inflorescence* subcapitate, broadly obovoid to fusiform, 3–15 by 2–4 cm, terminating a leafy shoot; bracts, bracteole, calyx, ovary, and capsule glabrous to rather densely puberulous; bracts green or orange, herbaceous to chartaceous, narrowly triangular to triangular, 0.5–3 by 0.5–0.8 cm, callus 1–5 mm long; bracteole tubular, often deeply split on the abaxial side, 10–17 mm long, 2-lobed, lobes deltate, 1–3 mm long. *Flowers:* calyx green, 10–30 mm long, lobes deltate, 2–5 mm long, callus c. 2 mm long, tube often deeply split during anthesis; corolla white or orange, tube yellowish white, 50–60 mm long, glabrous to rather densely puberulous, lobes narrowly ovate, 25–35 mm long; labellum carmine red to orange, throat yellow, upper part horizontally spreading, reflexed, suborbicular, 25–30 by 30–35 mm, margin irregularly dentate; stamen red to orange, 12–15 by 3–6 mm, erect, apex red, irregularly dentate, anther c. 5 mm long. *Capsule* ellipsoid, 10–11 mm long.

Distribution — see subspecies.

Habitat & Ecology — see subspecies.

IUCN Conservation Status — Not evaluated. There are 107 collection records examined of which 87 specimens were collected within the past 50 years. This species is also commonly reported in iNaturalist (https://www.inaturalist.org/observations?place_id=any&taxon_id=504228) with 38 recent observations. The species is widely distributed in Brazil and the Guianas. Our observations indicate that this is a rather common species within its range and some of these areas are protected areas with no known specific threats. Population and threat data is not available but based on the number of collection records and our general field observations, this species would probably score a status of Least Concern.

## KEY TO THE SUBSPECIES

1. Bracts 1.5–3 cm long; calyx shorter or as long as the bracts; inflorescence strongly elongate to fusiform a. subsp. *lanceolatus*

1. Bracts 0.5–1.5 cm long; calyx longer than the bracts; inflorescence subcapitate b. subsp. *pulchriflorus*

**a.** subsp. ***lanceolatus*** — Fig. 4g

*Inflorescence* strongly elongate to fusiform; bracts 1.5–3 cm long. *Calyx* shorter or as long as the bracts.

Distribution — Colombia (Meta), Venezuela (Amazonas), Peru (Madre de Dios), Brazil (Acre, Amazonas, Mato Grosso, Pará, Rondônia).

Habitat & Ecology — In non-inundated (terra firme) forests, one collection from llanos vegetation, on clayey or sandy soil, at elevations of 0–250 m. Flowering and fruiting: mainly from October to March.

Vernacular name — Brazil: Caninho-do-brejo.

Notes — In two collections from Acre, Brazil (*Lowrie 485* and *Maas et al. 9153*), the presence of ellipsoid tubers (10–20 by c. 5 mm) in the roots is mentioned.

**b.** subsp*. **pulchriflorus*** (Ducke) C.D.Specht & D.W.Stev. – Fig. 4g, inset

*Chamaecostus lanceolatus* (Petersen) C.D.Specht & D.W.Stev. subsp*. pulchriflorus* (Ducke) C.D.Specht & D.W.Stev. (2006) 158. — *Costus pulchriflorus* Ducke (1922) 22, t. 2 a–c. — *Costus lanceolatus* subsp. *pulchriflorus* (Ducke) Maas (1972) 41, f. 20. — Lectotype (designated by Maas 1972: 41): *Ducke MG 15649* (lecto G [G00168444]; isolecto BM [BM001010494], F [F0047180F], G [G00168444], P [P00686618], RB [RB00544558], US [US00092974]), Brazil, Pará, Alcobaça, Rio Tocantins, 7 Jan. 1915.

*Inflorescence* subcapitate; bracts 0.5–1.5 cm long. *Calyx* longer than the bracts.

Distribution — Suriname, French Guiana, Brazil (Amazonas, Goiás, Maranhão, Mato Grosso, Pará).

Habitat & Ecology — In non-inundated (terra firme) forests, sometimes on rock savannas, on sandy, clayey or lateritic soils, at elevations of 0–750 m. Flowering and fruiting: all year through.

Vernacular names — Brazil: Beiradão, Cana de macaco. French Guiana: Ampuku singaafu (Boni), Pikin singa afu (Boni).

Notes — Both subspecies look quite similar to *C. congestiflorus* and *C. fragilis*, but they are distinguished from both species by the red to orange instead of white flowers.

8. ***Chamaecostus manausensis*** Maas & H.Maas — Fig. 4h; Map 5

**Fig. 5.**
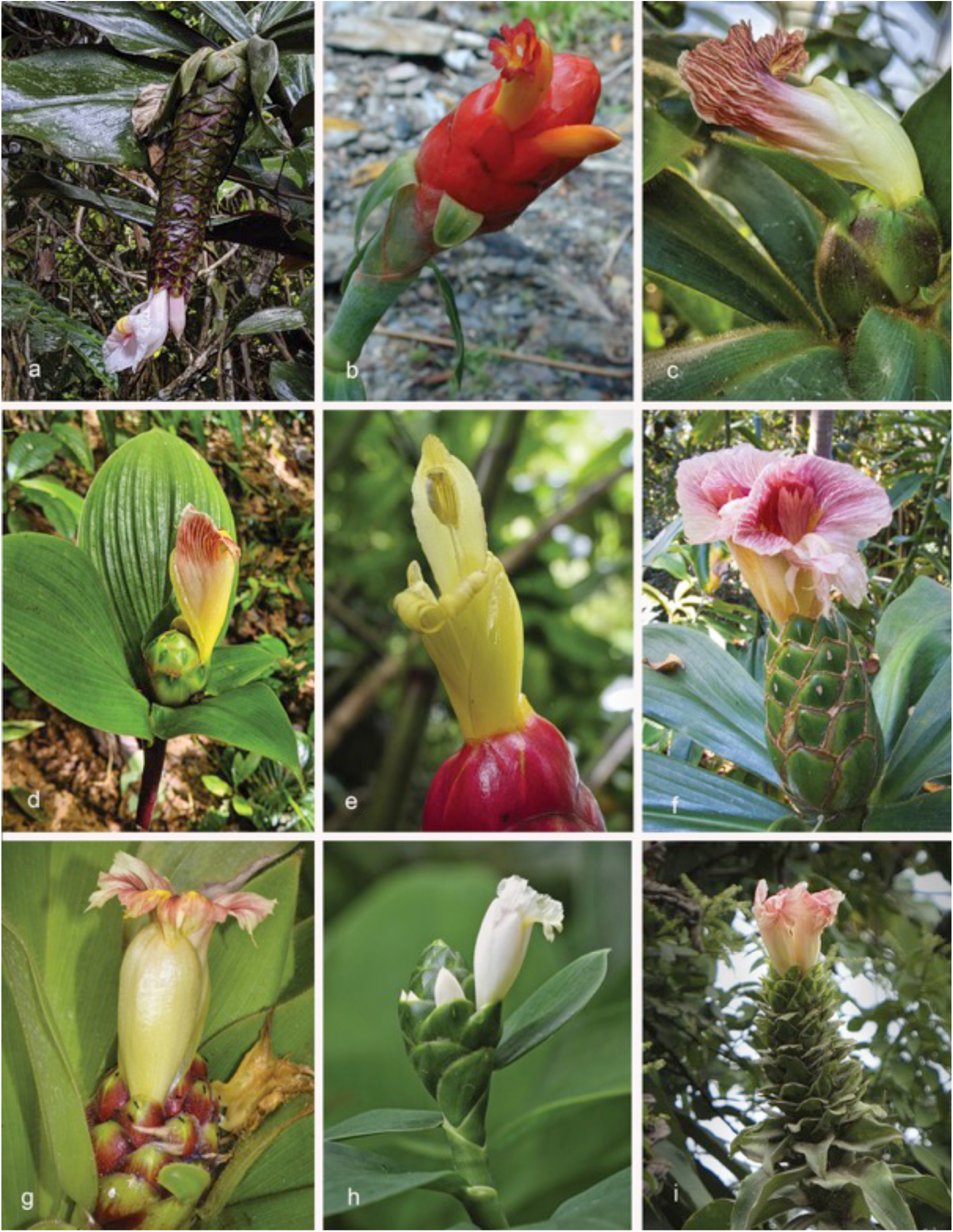
*Costus* species: *C. acreanus – C.asplundii*: a. *C. acreanus* (Loes.) Maas (*D.Skinner R3486*, from Rio Tarauacá, Acre, Brazil; b. *C. alfredoi* Maas & H.Maas (*Fuentes 6216* from Bolivia; c. *C. allenii* Maas (*D.Skinner R3350*, from Gamboa, Panama); d. *C. alleniopsis* Maas & D.Skinner (*D. Skinner R3498*, from Veraguas, Panama); e. *C. alticolus* Maas & H.Maas (*D. Skinner R3400* from Oaxaca, Mexico); f. *C. amazonicus* (Loes.) J.F.Macbr. (*D. Skinner R3198*, Rio Pastaza, Ecuador); g. *C. antioquiensis* Maas & H.Maas (cultivated plant at Burgers Bush, the Netherlands); h. *C. arabicus* L. (*D. Skinner R3124*, from Rio Mazon, Loreto, Peru); i. *C. asplundii* (Maas) Maas (not collected, photographed in Putumayo, Colombia). Photos: a, c–f, h & i by D. Skinner; b by A. Fuentes; g by P.J.M. Maas.

*Chamaecostus manausensis* Maas & H.Maas in Maas et al. (2023) 77, f. 1. — Type: *Prance et al. 11529* (holo U, 2 sheets [U1219716, U1219717]; iso BR, COL [COL128233], CR, DAV, E, F [F175333, F1842269], GH, INPA, K, MO, NY, P, S, US, VEN, W), Brazil, Amazonas, Mauá Road, 22 March 1971.

Low herbs, 0.4–1 m tall. *Leaves:* sheaths 2–15 mm diam; ligule 1–2 mm long; petiole 0–5 mm long; sheaths, ligule and petiole densely puberulous; lamina 13–25 by 3–6 cm, narrowly elliptic, adaxially sparsely puberulous to glabrous, abaxially sometimes reddish-purple, sparsely to densely puberulous, particularly the primary vein, base obtuse, apex acuminate, acumen 5–15 mm long. *Inflorescence* narrowly cylindric, 4–10 by 1.5–3 cm, up to c. 13 by 4 cm in fruit, terminating a leafy shoot; bracts, bracteoles, calyx, ovary, and capsule rather densely to densely puberulous; bracts 2.5–3.5 by 0.3–0.5 cm, narrowly triangular, green, chartaceous, apex often mucronate, callus 3–6 mm long; bracteole 15–20 mm long, tubular, often deeply split on the abaxial side, 2-lobed, lobes 3–7 mm long, deltate. *Flowers:* calyx 20–33 mm long, green, tube often deeply split during anthesis, lobes 5–10 mm long, subulate; corolla 50–65 mm long, cream, densely puberulous, lobes 30–40 mm long narrowly, obovate; labellum c. 35 by 40 mm, cream, distal edge horizontally spreading, broadly angular-obovate, margin crenulate; stamen 25–30 by 10 mm, cream, apex reflexed, irregularly dentate, anther 5–6 mm long. *Capsule* ellipsoid, 11–13 mm long.

Distribution — Brazil (Amazonas, Pará).

Habitat & Ecology — In non-inundated rain forests or campinarana vegetation, on sandy or clayey soils, around sea level. Flowering and fruiting: October to March, July.

IUCN Conservation Status — Not evaluated. There are 8 collection records examined of which 6 specimens were collected within the past 50 years. This species is narrowly distributed in the states of Amazonas and Pará in Brazil. Population and threat data is not available but based on the number of collection records this species is probably either Near Threatened or Vulnerable.

Notes — *Costus manausensis* looks quite similar to *C. congestiflorus* and has been regularly confused with that species, but can be distinguished by a narrowly cylindric (instead of broadly obovoid) inflorescence and a non-fimbriate labellum. For additional field data see Costa et al. (2011), where this species has been treated under the name *Costus congestiflorus*.

9. ***Chamaecostus subsessilis*** (Nees & Mart.) C.D.Specht & D.W.Stev. — Fig. 4i; Map 1

*Chamaecostus subsessilis* (Nees & Mart.) C.D.Specht & D.W.Stev. (2006) 158. — *? Globba subsessilis* Nees & Mart. (1823) 29. — *Costus subsessilis* (Nees & Mart.) Maas (1976) 469. —Type: *Wied zum Neuwied s.n.* (holo BR, 2 sheets [BR0000005188031, BR0000005188369]), Brazil, Bahia, “ad viam Felisbertiam”, Dec. 1816. See note 2.

*Costus warmingii* Petersen (1890) 57, t. 14. — Lectotype (designated here): *Warming 502* (C, 2 sheets [C10017745, C10017746] (coloured drawings!)), Brazil, Minas Gerais, Lagoa Santa, 9 Dec. 1865

*Costus pumilus* Petersen (1890) 58. — Lectotype (designated by Maas 1972: 30): *Burchell 6365* (B, destroyed; isolecto K [K000586745]), Brazil, Goiás, between Retiro and Sapezal, 29 Oct. 1827.

*Costus paucifolius* Gagnep. (1902)100. — Type: *Glaziou 22180* (holo P [P00686623]; iso BR, C [C10017747], K, LE [LE00001316], P [P00686624, P00686625]), Brazil, Goiás, Capelinha de Sao Antonio near Lage, 25 Oct. 1894.

*Costus rosulifer* Gagnep. (1902) 101. — Syntypes: *Weddell 2832* (P [P000686630, P00686631]), Brazil, Goiás, Sertão do Amaro Leite, Sept. – Oct. 1844.

*Costus latifolius* Gagnep. (1902) 101. — Syntypes: *A. de Saint*-*Hilaire 13* (F [F0047175F], P [P00686627, P00686628, P00686629], U [U0007220]), Brazil, Minas Gerais, Olho d’Agua, anno 1816.

*Costus pumilus* Petersen var. *pilosissimus* Gagnep. (1902) 102. — *Costus pilosissimus* (L.F.Gagnep.) K.Schum. (1904) 420. — Lectotype (designated by Maas 1972: 30): *Glaziou 22181* (lecto P [P00686626]; isolecto BR [BR0000006942694], C, G [G00168414], K [K000586747]), Brazil Goiás, between Guariroba and São Antonio, 20 Oct. 1894.

*Costus gagnepainii* K.Schum. (1904) 420. — Lectotype (designated by Maas 1972: 30): *Glaziou 15666* (lecto C; isolecto C, K [K000586748], LE [LE00001315], P [P00686622]), Brazil, locality unknown.

Acaulescent or very low, rosulate herbs, 0.25–1 m tall; roots fleshy, ending in fusiform to ellipsoid tubers, 0.5–3 cm long. *Leaves* 4–6, rosulate; sheaths membranous, obtuse at the apex, 2.5–6 by 0.5–2 cm, densely hirsute, sparsely to rather densely puberulous, or glabrous; ligule < 1 mm long; petiole absent; lamina elliptic or narrowly so, 5.5–34 by 2.4–13.3 cm, upper side sparsely puberulous or glabrous, lower side densely hirsute, rather densely to sparsely puberulous, or rarely glabrous, base acute, apex mucronate or shortly acuminate (acumen 0–22 mm long). *Inflorescence* terminal, formed in the centre of the rosette, 1–4-flowered; bracts, appendages of bracts, bracteole, pedicels, and calyx glabrous to densely puberulous, ovary densely sericeous to glabrous; bracts green, herbaceous, 1–10 by 0.2–2.6 cm; appendages green, foliaceous, narrowly triangular to deltate, apex mucronate; bracteole tubular, 25–32 mm long, 2-lobed at the apex, lobes narrowly triangular to deltate, 2–15 mm long. *Flowers*: pedicels c. 5 mm long; calyx 6–48 mm long, lobes narrowly triangular to deltate, 4–15 mm long, mucronate; corolla yellow or whitish, 55–70 mm long, glabrous to densely puberulous, lobes narrowly elliptic, 30–40 mm long; labellum yellow to pale yellow, striped with orange in the throat, broadly obovate when spread out, 60–70 by 70–95 mm, margin irregularly crenulate; stamen yellow, 35–47 by 6–20 mm, apex reflexed, obtuse or irregularly lobulate, anther 7–8 mm long. *Capsule* unknown.

Distribution — Brazil (Bahia, Ceará, Distrito Federal, Espírito Santo, Goiás, Maranhão, Minas Gerais, Rio de Janeiro, São Paulo, Tocantins).

Habitat & Ecology — In the understory of gallery forests of Cerrado savannas and the Atlantic rain forest of Southeastern Brazil, at elevations of 0–1000 m. Flowering in the rainy season.

IUCN Conservation Status — Not evaluated. There are 71 collection records examined of which 33 specimens were collected within the past 50 years. They are distributed in the eastern states of Brazil (Bahia, Goiás, Mato Grosso, Minas Gerais, and the Distrito Federal). Population and threat data is not available but based on the number of collection records this species would probably score a status of Least Concern.

Vernacular names — Brazil: Campo alegre, Cana de macaco, Cana fistula, Erva mole amarela.

Notes — 1. *Chamaecostus subsessilis* is very similar to *C. acaulis*, but it is distinguished by its elliptic leaves instead of oblong-obovate, with upper side sparsely puberulous to glabrous instead of densely puberulous, and usually low (from 25 cm up to 1 m tall) and erect growth habit. For additional discussion on this complex see André *et al*. (2015).

2. The name *Globba subsessilis* is published with a question mark. We interpret it as an accepted taxon of which the placement in *Globba* is questionable. Therefore, Art. 36.1 (Turland et al. 2018) does not apply and the name is validly published.

***2. Costus*** L.

*Costus* L. (1753) 2. — Type: *Costus arabicus* L.

*Gissanthe* Salisb. (1812) 279, nom. inval., no description. — *Glissanthe* Steud. (1840) 688 (likely spelling error) — Based on: *Alpinia spiralis* Salisb., nom. inval.

*Jacuanga* T.Lestib. (1841) 329, 341. — Type: *Costus pisonis* Lindl.

Tall, gigantic, or low herbs, occasionally branched. *Leaves*: ligule short or long, truncate to 2-lobed; lamina generally narrowly elliptic to narrowly obovate, sometimes plicate, lower side sometimes purplish red, base acute, rounded, or occasionally cordate, apex acuminate, rarely acute. *Inflorescence* a spike, terminating a leafy shoot, or terminating a separate, short, leafless shoot. *Flowers* abaxially or rarely adaxially orientated; bracts coriaceous, rarely chartaceous, green, yellow, orange, or red, narrowly to broadly ovate-triangular, often with a nectariferous callus, often provided with foliaceous, apical appendages; bracteole boat-shaped, very rarely 2-keeled at the adaxial side; calyx small, rarely exceeding the bracts; corolla white, yellow, orange, or red; labellum white, yellow, orange, or red, 3-lobed, large and horizontally flattened, lateral lobes often erect, with reddish venation, middle lobe often recurved and with a honey mark, or labellum small and tubular; basal part of labellum in nearly all species papillate; stamen with the anther attached at the middle or rarely at the base; stigma 2-lamellate, with a dorsal 2-lobed appendage; ovary 3-locular, ovules in 2 rows per locule. *Capsule* 3-locular, ellipsoid to globose, chartaceous, loculicidally dehiscent or irregularly breaking up with age. *Seeds* with a large and lacerate aril, all seeds of one locule often cohering by their arils at dehiscence.

Distribution — Throughout the Neotropics.

Habitat & Ecology — In various forest types, ranging from rain forest, submontane forest to montane forest, generally occurring at low elevations, but sometimes up to 2500 m.

## KEY TO THE SPECIES

Two dichotomous keys are provided: one to the species of Central America and a separate key for the species of tropical South America and the Antilles.

## KEY TO THE SPECIES OF CENTRAL AMERICA

1. All bracts provided with a leafy appendage or apical part of bracts slightly curved outwards 2

1. All bracts without a leafy appendage (or only the lowermost provided with a leafy appendage), apical part of bracts not curved outwards 15

2. Appendages of bracts green; labellum horizontally spreading 3

2. Appendages of bracts reddish, rarely yellow or green; labellum tubular 7

3. Ligule 1–4 mm long; most parts of the plant, including the corolla, densely velutinous. – Mexico 45. *C. mollissimus*

3. Ligule 2–22 mm long: plants villose, puberulous, or glabrous; corolla glabrous 4

4. Labellum completely yellow; leaves mostly cordate at the base. — Nicaragua, Costa Rica, Panama 69. *C. villosissimus*

4. Labellum white, yellow, or pink, lateral lobes mostly striped with red; leaves mostly acute to rounded at the base (if cordate, then labellum not yellow) 5

5. Leaves generally 10–20-plicate; inflorescence mostly nodding. — Nicaragua, Costa Rica, Panama 14. *C. bracteatus*

5. Leaves not plicate; inflorescence erect 6

6. Most parts of the plants whitish villose, puberulous, or glabrous; upper side of leaves scabrid to the touch. — Mexico (Chiapas) to Panama 43. *C. macrostrobilus*

6. Most parts of the plants brownish villose; upper side of leaves smooth to the touch. — Costa Rica, Panama 62. *C. sinningiiflorus*

7. Average ligule length 15–40 mm 8

7. Average ligule length 1–15 mm 11

8. Inflorescence terminating a separate leafless shoot. — Costa Rica, Panama 15. *C. callosus*

8. Inflorescence terminating a leafy shoot — Costa Rica 9

9. Lobes of ligule triangular acute; most parts of the plant, including the corolla, densely velutinous. 46. *C. montanus*

9. Lobes of ligule rounded to obtuse; most parts of the plants not velutinous, but varying from villose, puberulous, sericeous to glabrous 10

10. Margins of bracts often covered with hair-like fibers; bracteole boat-shaped 13. *C. barbatus*

10. Margins of bracts not covered with hair-like fibers; bracteole 2-keeled 58. *C. ricus*

11. Plants generally < 1 m tall; leaves obovate to elliptic, densely to rather densely villose on both sides; bracteole 2-keeled. — Costa Rica 50. *C. osae*

11. Plants generally > 1 m tall; leaves narrowly obovate to narrowly elliptic, not densely villose on both sides; bracteole boat-shaped (but 2-keeled in some Panamanian material of *C. comosus*) 12

12. Bracts orange, yellow, or red, their apex slightly curved outwards; upper part of ligule with a minute salient rim 13

12. Bracts red, rarely green, apical part leafy, often reflexed; upper part of ligule without salient rim 14

13. Inflorescence terminating a separate, leafless shoot; callus of bracts 8—25 mm long. — Costa Rica, Panama 15. *C. callosus*

13. Inflorescence terminating a leafy shoot; callus of bracts absent or inconspicuous. — Costa Rica, Panama 23. *C. curvibracteatus*

14. Ligule 1–3(–6) mm long, upper side of leaves sometimes slightly scabrid to the touch; appendages of bracts ovate-triangular to narrowly triangular. — Mexico (Chiapas) to Panama 19. *C. comosus*

14. Ligule 5–15 mm long, upper side of leaves scabrid to the touch; appendages of bracts ovate to broadly ovate. — Honduras to Panama 40. *C. lima*

15. Inflorescence terminating a separate leafless shoot 16

15. Inflorescence terminating a leafy shoot (but see also *C. alleniopsis* which rarely has an inflorescence terminating a separate leafless shoot) 20

16. Leaves linear, to 2 cm wide, sheaths whitish with a reddish-brown upper margin. — Costa Rica. 66. *C. stenophyllus*

16. Leaves > 2 cm wide, sheaths green 17

17. Labellum tubular, yellow to orange; lateral lobes not striped 18

17. Labellum horizontally spreading; lateral lobes of labellum striped with red 19

18. Ligule 4—12 mm long; bracts red to bright crimson, apex appressed. — Mexico (Oaxaca) 5. *C. alticolus*

18. Ligule 10—30 mm long; bracts orange, red to yellow, apex often curved outwards. — Costa Rica, Panama 15. *C. callosus*

19. Ligule 3—12 mm long; lower side of leaves densely velutinous. — Mexico (Veracruz) 24*. C. dirzoi*

19. Ligule 15—22 mm long; lower side of leaves densely villose. — Mexico, Guatemala, Honduras 61. *C sepacuitensis*

20. Average ligule length 0.5–20 mm 21

20. Average ligule length 15–100 mm 32

21. Ligule < 1 mm long; leaves densely puberulous to villose on both sides; bracts green; labellum horizontally spreading, yellow, lateral lobes striped with red. — Nicaragua, Costa Rica 44. *C. malortieanus*

21. Ligule 2–20 mm long; leaves with a different indument; bracts red, orange, yellow, or green; labellum horizontally spreading or tubular, variously coloured 22

22. Inflorescence fusiform (also rarely found in *C. lasius*), with a distinctly pointed apex; margin of bracts covered with hair-like fibers; labellum tubular, red to yellow. — Mexico to Panama 57. *C. pulverulentus*

22. Inflorescence more or less ovoid to globose, apex obtuse to acute, never pointed; margin of bracts not covered with hairy fibers; labellum tubular or horizontally spreading, variously coloured 23

23. Apex of bracts slightly curved outwards; upper part of ligule with a minute salient rim. — Costa Rica, Panama 23. *C. curvibracteatus*

23. Apex of bracts appressed; upper margin of ligule without a salient rim 24

24. Ligule 2–4 mm long; bracts green, callus inconspicuous; labellum horizontally spreading. — Mexico to Costa Rica 51. *C. pictus*

24. Ligule 2–20 mm long; bracts green, orange, yellow, to red; callus conspicuous; labellum tubular or horizontally spreading 25

25. Lower side of leaves densely velutinous. — Mexico (Veracruz) 24. *C. dirzoi*

25. Lower side of leaves never velutinous 26

26. Leaves and sheaths mostly densely brownish villose 27

26. Leaves and sheaths glabrous to puberulous (sheaths sometimes villose in *C. wilsonii*) 29

27. Bracts green; flowers adaxially orientated to erect, with a horizontally spreading labellum 28

27. Bracts yellow to pale orange; flowers abaxially orientated, labellum tubular. — Nicaragua (?) to Panama 38. *C. lasius*

28. Leaves not or only very slightly plicate; ligule 2–10 mm long. — Costa Rica, Panama 3. *C. allenii*

28. Leaves distinctly plicate; ligule 12–20 mm long. — Panama 4. *C. alleniopsis*

29. Flowers adaxially orientated, with a horizontally spreading labellum 50–70 mm long. — Guatemala to Panama 35. *C. kuntzei*

29. Flowers abaxially orientated, with a tubular labellum 20–35 mm long 30

30. Plants completely glabrous; leaf base cordate; growing on or near sandy beaches. — Nicaragua to Panama 74. *C. woodsonii*

30. Plants not completely glabrous; leaf base acute to rounded; never growing on or near sandy beaches 31

31. Bracts orange-red to red, never yellow; labellum tubular, lateral lobes never striped with red; apex of leaves acuminate, but margin of acumen never rolled inwards. — Mexico to Panama 60*. C. scaber*

31. Bracts yellow, sometimes red, orange, or green; labellum tubular, lateral lobes striped with red; apex of leaves long-acuminate with margin rolling inwards. — Costa Rica, Panama 73. *C. wilsonii*

32. Ligule 70–100 mm long; sheaths, ligule, and lower side of leaves wine-red; basal sheaths more or less inflated with inflexed upper margin. — Panama 70. *C. vinosus*

32. Ligule up to 65 mm long; sheaths, ligule, and lower side of leaves green; basal sheaths not inflated 33

33. Plants with glaucous stems, leaves, and bracts; bracts scabrid to the touch. — Nicaragua to Panama 32. *C. glaucus*

33. Plants not glaucous; bracts not scabrid to the touch 34

34. Leaf sheaths glabrous, rarely sparsely sericeous 35

34. Leaf sheaths villose to velutinous, or puberulous 37

35. Ligule 5–20 mm long; flowers adaxially orientated; labellum large and horizontally spreading, mostly dark purple with white to yellow striped lateral lobes. — Guatemala to Panama 35. *C. kuntzei*

35. Ligule 20–60 mm long; flowers abaxially orientated; labellum small and tubular, yellow 36

36. Leaves not plicate, mostly shiny; corolla 60–75 mm long. — Costa Rica, Panama 47. *C. nitidus*

36. Leaves strongly 6–8-plicate, not shiny; corolla 45–55 mm long. — Costa Rica, Panama 53. *C. plicatus*

37. Labellum large and horizontally spreading, lateral lobes striped with red, yellow to orange; flowers adaxially orientated; corolla glabrous; ligule 12–20 mm long. — Panama 4. *C. alleniopsis*

37. Labellum small and tubular, completely yellow to orange; flowers abaxially orientated; corolla puberulous to velutinous; ligule 20–60 mm long 38

38. Lobes of ligule rounded to obtuse; margin of bracts often covered with hair-like fibers; most parts of the plant puberulous to villose. — Costa Rica 13. *C. barbatus*

38. Lobes of ligule acute to triangular; margin of bracts not covered with hair-like fibers; most parts of the plant velutinous. — Costa Rica 46. *C. montanus*

## KEY TO THE SPECIES OF *COSTUS* FROM TROPICAL SOUTH AMERICA AND THE ANTILLES

### Key to the four groups of species of *Costus*

1. All bracts provided with a leafy appendage or apical part of bracts slightly curved outwards 2

1. All bracts without a leafy appendage, apical part of bracts not curved outwards 3

2. Labellum small, tubular, lateral lobes indistinct Group 1

2. Labellum large, composed of a short tube and a distinct horizontally spreading limb, lateral lobes distinct and often striped with red Group 2

3. Labellum small, tubular, lateral lobes indistinct Group 3

3. Labellum large, composed of a short tube and a distinct horizontally spreading limb, lateral lobes distinct and often striped with red Group 4

### Key to the species of *Costus* Group 1 (appendaged bracts, labellum small, tubular)

1. Inflorescence terminating a separate leafless shoot. — Colombia. 15. *C. callosus*

1. Inflorescence terminating a leafy shoot (but see also *C. comosus*) 2

2. Ligule 1–3 mm long. — Throughout the western part of tropical South America 19. *C. comosus*

2. Ligule 5–60 mm long 3

3. Leaves obovate to elliptic; bracteole 2-keeled; low plants generally < 1 m tall. — Colombia, Ecuador 50. *C. osae*

3. Leaves narrowly ovate to narrowly obovate (rarely obovate in *C. juruanus*); bracteole boat-shaped; plants generally > 1 m tall 4

4. Upper side of leaves more or less scabrid to the touch, lower side densely villose. — throughout tropical South America 40. *C. lima*

4. Upper side of leaves smooth to the touch, lower side glabrous to densely villose 5

5. Lower side of leaves green, glabrous, but primary vein densely villose; leaves not plicate. — Bolivia (Cochabamba) 18. *C. cochabambae*

5. Lower side of leaves often reddish purple, sparsely to densely villose; leaves 4–8-plicate 6

6. Ligule 10–15 mm long; corolla orange to yellow, densely villose; lower side of leaves densely villose; labellum orange-red. — Colombia (Putumayo) 22. *C. cupreifolius*

6. Ligule 5–60 mm long; corolla mostly pale yellow, glabrous; lower side of leaves glabrous, rarely sparsely to rather densely villose; labellum mostly yellow. — throughout tropical South America 34. *C. juruanus*

### Key to the species of *Costus* Group 2 (appendaged bracts, labellum large, composed of a short tube and a distinct horizontally spreading limb)

1. Ligule 20–70 mm long 2

1. Ligule 2–20 mm long 8

2. Bracts red, without a distinct appendage, but apex curved outwards and almost pungent. — Amazonian Colombia, Ecuador, Peru, Brazil and Guyana 41. *C. longibracteolatus*

2. Bracts green, with a distinct leafy appendage 3

3. Labellum including lateral lobes almost completey white 4

3. Labellum salmon-coloured, cream to white, lateral lobes always striped with red to pink 5

4. Lobes of ligule acute; margin of bracts glabrous. — Pacific side of Colombia and Ecuador 39. *C. leucanthus*

4. Lobes of ligule obtuse; margin of bracts densely villose. — Amazonian Colombia 52. *C. pitalito*

5. Bracts without a distinct appendage, but apex curved outwards and almost pungent. — Amazonian Colombia, Ecuador, Peru, Brazil and Guyana 41. *C. longibracteolatus*

5. Bracts with a distinct leafy appendage 6

6. Appendages of bracts horizontally spreading, 5–15 mm long, apex rounded; calyx 7–10 mm long. — Amazonian Peru and adjacent Brazil, often in periodically inundated várzea forest 68. *C. varzearum*

6. Appendages of bracts ascending, 25–60 mm long, apex acute; calyx 11–20 mm long 7

7. Upper side of leaves green; leaves mostly distinctly 6–7-plicate; ligule unequally 2-lobed. — Colombia, Ecuador, Peru, Brazil (Acre) 27. *C. erythrophyllus*

7. Upper side of leaves dark olive green; leaves not or only very slightly plicate; ligule obliquely truncate. — Colombia, Ecuador 72. *C. whiskeycola*

8. Flowers completely yellow. — throughout tropical South America 69. *C. villosissimus*

8. Flowers differently coloured, but never completely yellow 9

9. Calyx 25–27 mm long; lower side of leaves dark purple. — Peru 48. *C. obscurus*

9. Calyx 9–22 mm long; lower side of leaves greenish10

10. Labellum erect, with a narrow, recurved margin, with 4—6 apical teeth, without distinct lateral lobes. — Peru, Bolivia, Brazil (Acre) 33. *C. guanaiensis*

10. Labellum horizontally spreading, without apical teeth, with distinct lateral lobes 11

11. Leaf sheaths densely to sparsely villose 12

11. Leaf sheaths glabrous to puberulous or sericeous 14

12. Inflorescence often nodding. — Colombia, Ecuador, Peru 9. *C. asplundii*

12. Inflorescence always erect 13

13. Upper side of leaves scabrid to the touch, most vegetative parts whitish villose. — Throughout tropical South America 43. *C. macrostrobilus*

13. Upper side of the leaves smooth to the touch; most vegetative parts brownish villose. — Throughout tropical South America 62. *C. sinningiiflorus*

14. Margin of lower leaf sheaths distinctly swollen; ligule often with a horizontal ridge near the apex. — Ecuador 31. *C. gibbosus*

14. Margin of lower leaf sheaths not swollen; ligule without a horizontal ridge near the apex 15

15. Inflorescence terminating a separate leafless shoot 16

15. Inflorescence terminating a leafy shoot 17

16. Ligule 5–15 mm long, truncate or slightly 2-lobed; appendages of bracts horizontally spreading, apex rounded. — The 3 Guianas 17. *C. claviger*

16. Ligule 10–40 mm long, very deeply and unequally 2-lobed; appendages of bracts ascending, apex acute. — Colombia, Ecuador, Peru, Brazil (Acre) 27. *C. erythrophyllus*

17. Leaves often 6–7-plicate, often dark purple-red at the lower side —Colombia, Ecuador, Peru, Brazil (Acre) *C. erythrophyllus*

17. Leaves not plicate, greenish at the lower side. — Colombia, Venezuela, Ecuador, Peru 37. *C. laevis*

### Key to the species of *Costus* Group 3 (non-appendaged bracts, tubular flowers)

1. Flowers adaxially orientated, pinkish red to salmon-red 2

1. Flowers abaxially orientated to erect, variously coloured 3

2. Bracts generally red to pinkish red; the upper leaves not forming a bowl-shaped involucrum at the base of the inflorescence. — throughout tropical South America 64. *C. spiralis*

2. Bracts green; the upper leaves forming a bowl-shaped involucrum at the base of the inflorescence. — Peru, Bolivia 71. *C. weberbaueri*

3. Leaves linear 4

3. Leaves not linear 6

4. Ligule < 1 mm long; calyx 15–16 mm long. — Bolivia (La Paz) 2. *C. alfredoi*

4. Ligule 1–3 mm long; calyx 7–13 mm long 5

5. Flowers red, pink, or pinkish orange; calyx 9–13 mm long; leaves 2–7 cm wide, lower side glabrous to sparsely puberulous. — Amazonian Peru and Brazil 25. *C. douglasdalyi*

5. Flowers yellow; calyx 7–9 mm long; leaves 1.5–2.5 cm wide, lower side densely puberulous. — Amazonian Colombia, Ecuador, Peru 76. *C. zingiberoides*

6. Inflorescence fusiform and slightly pointed; margin of bracts mostly covered with hair-like fibers; corolla lobes often recurved. — Throughout the western part of South America 57. *C. pulverulentus*

6. Inflorescence ovoid, cylindric, globose, or ellipsoid (but see also *C. fissicalyx*); margin of bracts rarely covered with hair-like fibers 7

7. Leaves bullate; stamen far exceeding the labellum. — Amazonian Colombia 29. *C. fissicalyx*

7. Leaves never bullate (but sometimes slightly bullate in *C. cordatus*); stamen not or only slightly exceeding the labellum 8

8. Inflorescence terminating a separate leafless shoot 9

8. Inflorescence terminating a leafy shoot 18

9. Leaves 3–12-plicate 10

9. Leaves generally not plicate 13

10. Calyx 15–20 mm long; corolla c. 100 mm long; labellum 50–70 mm long. — Colombia, Ecuador 30. *C. geothyrsus*

10. Calyx 8–14 mm long; corolla 45–75 mm long; labellum 30–45 mm long 11

11. Leaves 10–12-plicate, base cordate to obtuse; corolla 65–75 mm long. — Pacific Colombia and Ecuador 21. *C. cordatus*

11. Leaves 3–10-plicate, base acute to obtuse; corola 45–55 mm long 12

12. Lower side of the leaves sparsely puberulous to glabrous; corolla red, reddish pink, or pinkish orange. — The 3 Guianas, Amazonian Brazil and Peru 28. *C. erythrothyrsus*

12. Lower side of the leaves densely to rather densely villose; corolla yellow. — throughout Colombia 54. *C. plowmanii*

13. Calyx 17–22 mm long; ligule 1–2 mm long; plants completely glabrous; young sheaths somewhat glaucous. — Peru (Puno) 10. *C. asteranthus*

13. Calyx 6–18 mm long; ligule 2–60 mm long; plants never completely glabrous; young sheaths not glaucous (but see also *C. vargasii*) 14

14. Ligule 25–60 mm long. — Peru, Bolivia 13. *C. beckii*

14. Ligule 2–35 mm long 15

15. Leaf sheaths often glaucous; petiole often reddish; plants completely glabrous. — Peru, Bolivia67. *C. vargasii*

15. Leaf sheaths not glaucous; petiole not reddish; plants always partly covered with hairs 16

16. Ligule 10–30 mm long, upper part of ligule with a minute salient rim; lower side of leaves rather densely villose. — Colombia 15. *C. callosus*

16. Ligule 2–10 mm long, upper part of ligule without a salient rim; lower side of leaves puberulous, sericeous, or glabrous 17

17. Ligule 2–3 mm long; leaves 2–7 cm wide, lower side glabrous to sparsely puberulous; margin of bracts not covered with hair-like fibers; corolla 45–55 mm long, pink to orange. — Amazonian Brazil and Peru 25. *C. douglasdalyi*

17. Ligule 6–10 mm long; leaves 7–13 cm wide, lower side densely to rather densely whitish sericeous; margin of bracts covered with hair-like fibers; corolla 60–65 mm long, yellow. — Amazonian Colombia, Ecuador, and Peru 26. *C. erythrocoryne*

18. Bracts generally yellow; leaf sheaths brownish villose. — throughout tropical South America 38. *C. lasius*

18. Bracts mostly red to orange, never yellow; leaf sheaths glabrous to hairy, sometimes brownish villose (*C. chartaceus*) 19

19. Plants completely glabrous 20

19. Plants always covered with some hairs 21

20. Ligule 5–35 mm long; leaf base acute to obtuse; leaf sheaths and upper leaf side often glaucous. — Peru, Bolivia 67. *C. vargasii*

20. Ligule 2–6 mm long; leaf base cordate; leaf sheaths and upper leaf side green and never glaucous. — Pacific Colombia, often growing along beaches 74. *C. woodsonii*

21. Lateral lobes of labellum distinct and often reddish striped. — Greater and Lesser Antilles 63. *C. spicatus*

21. Lateral lobes of labellum indistinct and rarely reddish striped 22

22. Labellum funnel-shaped. — Brazil (Pernambuco) 11. *C. atlanticus*

22. Labellum tubular 23

23. Calyx 3–7 mm long; primary vein of the upper side of the leaves covered with a row of erect hairs. — throughout tropical South America 60. *C. scaber*

23. Calyx 6–15 mm long; primary vein of the upper side of the leaves mostly not covered with a row of erect hairs 24

24. Leaves 3–12-plicate, lower side sometimes purplish red 25

24. Leaves not plicate, lower side green 27

25. Calyx 15–20 mm long; corolla c. 100 mm long. —Colombia, Ecuador 30. *C. geothyrsus*

25. Calyx 8–13 mm long; corolla 45–75 mm long 26

26. Leaves 10–12-plicate, base cordate to obtuse; bracts red to orange; corolla 65–75 mm long. — Pacific Colombia and Ecuador 21. *C. cordatus*

26. Leaves 3–8-plicate, base acute to obtuse; bracts carmine red; corola 45–55 mm long. — The 3 Guianas, Amazonian Brazil and Peru 28. *C. erythrothyrsus*

27. Leaf sheaths densely sericeous to glabrous. — Peru (Huánuco and Pasco), high elevations 59. *C. rubineus*

27. Leaf sheaths densely villose to glabrous 28

28. Bracts reddish pink to dark pink, without an appndage; ligule 5–15 mm long. — Amazonian Colombia, Ecuador, Peru 16. *C. chartaceus*

28. Bracts red to dark red, sometimes provided with a small, deltate appendage; ligule 2–8 mm long 29

29. Flowers yellow to orange. — Amazonian Brazil and Peru 55. *C. prancei*

29. Flowers pinkish 30

30. Leaves 6–22 by 3–7 cm; inflorescence 2–3 cm wide; bracteole 8–13 mm long; labellum c. 17 by 7 mm when spread out. — Brazil (Pará) 65. *C. sprucei*

30. Leaves 20–43 by 4–12 cm; inflorescence 3–6 cm wide; bracteole 15–20 mm long; labellum 30–35 by 20–25 mm when spread out. — Bolivia (Beni) and Brazil (Rondônia) 56. *C. pseudospiralis*

### Key to the species of *Costus* Group 4 (non-appendaged bracts, labellum large, composed of a short tube and a distinct horizontally spreading limb

1. Leaf sheaths and young leaves with a glaucous cover; ligule 15–55 mm long 2

1. Leaf sheaths and young leaves without a glaucous cover; ligule mostly much shorter 3

2. Bracts reddish, distinctly convex, smooth to the touch; upper leaves tightly wrapping and completely hiding the young inflorescence; bracteole 40–45 mm long; calyx 15–25 mm long. — Amazonian Colombia and Ecuador 20. *C. convexus*

2. Bracts pale green, not convex, scabrid to the touch; upper leaves not tightly wrapping and hiding the young inflorescence; bracteole 22–27 mm long; calyx 7–16 mm long. — Colombia 32. *C. glaucus*

3. Flowers adaxially orientated to erect 4

3. Flowers abaxially orientated 8

4. Leaf sheaths and leaves mostly completely glabrous; labellum mostly dark purple-red to reddish brown 5

4. Leaf sheaths and leaves mostly hairy (but see also glabrous forms of *C. antioquiensis*); labellum never dark purple-red to reddish brown 6

5. Inflorescence terminating a leafy shoot; leaves with a long-acuminate apex, lower side often purplish; calyx 6–15 mm long. — Colombia and Ecuador, W of the Andes 36. *C. kuntzei*

5. Inflorescence terminating a leafless shoot, rarely terminating a leafy shoot; leaves mostly with a long-acute apex, lower side green; calyx 12–17 mm long. — Amazonian Ecuador and adjacent Colombia, often at high elevations 75. *C. zamoranus*

6. Leaf sheaths densely to sparsely puberulous; corolla lobes recurved. — Amazonian Ecuador, high elevations 49. *C. oreophilus*

6. Leaf sheaths densely to sparsely villose, sometimes glabrous; corolla lobes erect 7

7. Base of the ligule provided with a ring of brown, erect and rather stiff hairs, sometimes absent; ligule green; upper side of leaves glabrous to densely villose; upper leaves tightly wrapping and completely hiding the inflorescence. — Colombia, N Venezuela, Ecuador, Peru 7. *C. antioquiensis*

7 Base of the ligule without a ring of hairs; ligule often reddish; upper side of leaves densely villose; upper leaves not tightly wrapping and completely hiding the inflorescence. — Pacific side of Colombia 3. *C. allenii*

7. Margin of bracts covered with hair-like fibers; corolla densely to rather densely puberulous to villose. — Amazonian Colombia, Ecuador, Peru 6. *C. amazonicus*

8b. Margin of bracts entire; corolla mostly glabrous (but see also *C. acreanus*) 9

8. Flowers often completely white; leaf base mostly cordate; lateral branches often arising from the base of the inflorescence. — throughout tropical South America 8. *C. arabicus*

9. Flowers never completely white; leaf base acute to obtuse (sometimes also cordate in *C. krukovii*); lateral branches absent 10

10. Inflorescence nodding to erect, mostly terminatng a leafy shoot; leaves not plicate; calyx 17–25 mm long. — throughout the Amazon region 1. *C. acreanus*

10. Inflorescence erect, mostly terminating a separate leafless shoot, sometimes terminating a leafy shoot; leaves mostly 8–10-plicate; calyx 10–17 mm long. — throughout the Amazon region and Guyana 35. *C. krukovii*

**1. *Costus acreanus*** (Loes.) Maas — Fig. 5a; Map 6

**Fig. 6.**
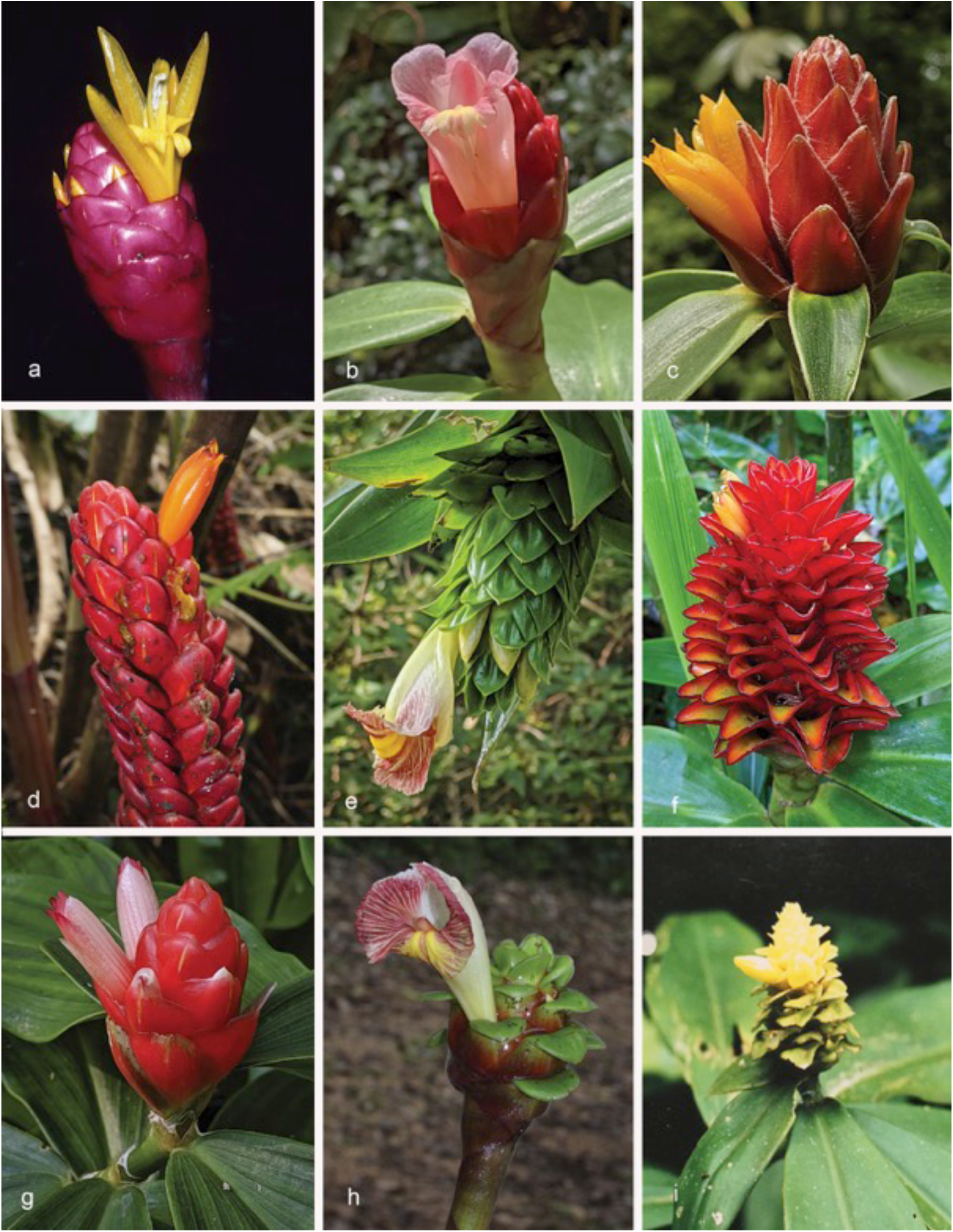
*Costus* species: *C. asteranthus – C. cochabambae*: a. *C. asteranthus* Maas & H.Maas (*Maas 6086* from Peru, Puno, San Gabán); b. *C. atlanticus* E.Pessoa & M.Alves (*D.Skinner R2907*, cultivated plant ‘Tropicais’ from Brazil); c. *C. barbatus* Suess. (*D.Skinner R3213* from Costa Rica, San José); d. *C. beckii* Maas & H.Maas (*D.Skinner R3392* from Bolivia, Cochabamba, Villa Tunari); e. *C. bracteatus* Rowlee (*D.Skinner R3361* from Panama, Guna Yala, Nusigandi); f. *C. callosus* Maas & H.Maas (cultivated at Lyon Arboretum, Hawaii, L- 88.0426 from *Kress 86-1978*); g. *C. chartaceus* Maas (*D.Skinner R2708*, cultivated plant from USBRG Kress collection); h. *C. claviger* Benoist (*Maas 10622* from French Guiana); i. *C. cochabambae* Maas & H.Maas (photo found in voucher folder of *Beck 7320* at Leiden). Photos: a & h by P.J.M. Maas; b–g, & i by D. Skinner.

*Costus acreanus* (Loes.) Maas (1972) 66, f. 13. — *Costus cylindricus* Jacq. var. *acreanus* Loes. (1929) 712. — Lectotype (designated by Maas 1972): *Ule 9195* (lecto K [K000586764]; isolecto MG [MG014054]), Peru, Madre de Dios, Seringal Auristella, Oct. 1911.

Herbs, 1–4.5 m tall. *Leaves*: sheaths 5–28 mm diam; ligule truncate to obliquely truncate, 1–15 mm long; petiole 1–20 mm long; sheaths, ligule and petiole densely to rather densely puberulous, sometimes glabrous; lamina narrowly elliptic to narrowly obovate, 12–55 by (4–)7–18 cm, upper side glabrous, but primary vein sparsely covered with a row of erect hairs, lower side often purplish to dark red, densely to rather densely puberulous, rarely glabrous, base acute to rounded, apex acuminate (acumen 5–10 mm long) to long-acute. *Inflorescence* nodding to erect, ovoid to cylindric, 4.5–12 by 3–8 cm, enlarging to c. 20 by 8–10 cm in fruit, terminating a leafy shoot or rarely terminating a separate leafless shoot c. 35 cm long, sheaths obliquely truncate, 3–5 cm long, sparsely to densely puberulous; bracts, bracteoles, calyx, ovary, and capsule densely to rather densely puberulous. *Flowers* abaxially orientated; bracts green, dark purple, or red, coriaceous, ovate-triangular, 3.5–6 by 2–3.5 cm, apex acute to obtuse, callus 3–11 mm long; bracteole boat-shaped, 20–40 mm long; calyx red, 17–25 mm long, sometimes exceeding the bracts, lobes deltate to triangular, 3–6 mm long; corolla white to pink, 70–100 mm long, glabrous or densely puberulous, lobes narrowly elliptic, 60–75 mm long; labellum white to pink, upper part horizontally spreading, broadly obovate, 80–105 by 70–90 mm, lateral lobes more or less striped with red, middle lobe recurved with yellow honey mark, margin irregularly dentate to lobulate; stamen white to pink, 45–70 by 15–17 mm, not exceeding the labellum, apex pinkish red to white, acute, anther 10–13 mm long. *Capsule* ellipsoid, 12–17 mm long.

Distribution — Colombia (Amazonas, Putumayo), Ecuador (Morona-Santiago, Zamora-Chinchipe), Peru (Cuzco, Loreto, Madre de Dios, Puno, Ucayali), Bolivia (Beni, Pando), Brazil (Acre, Amazonas, Rondônia).

Habitat & Ecology — In periodically inundated (restinga, várzea), non-inundated, or premontane moist forests, or marshes, at elevations of 0–700(–2800) m. Flowering all year round.

IUCN Conservation Status — Not evaluated. There are 77 collection records examined of which 58 specimens were collected within the past 50 years. This species is also reported in ∼30 recent iNaturalist observations (https://www.inaturalist.org/observations?place_id=any&taxon_id=607641). The species is widely distributed in Bolivia, Brazil, Colombia, Ecuador and Peru. Our observations indicate that this is a common species within its range and some of these areas are protected areas with no known specific threats. Population and threat data is not available but based on the number of collection records and our general field observations, this species would probably score a status of Least Concern.

Vernacular names — Peru: Caña-caña, Caña-caña morada.

Notes — *Costus acreanus* is one of the few species of *Costus* with an often nodding inflorescence. It can also be distinguished by the often densely puberulous bracts, bracteole, and calyx. The lower side of the leaves of this species is often purplish to dark red-coloured, and the corolla and labellum white to pink. Another feature of this species is the long calyx, which is often almost exceeding the bracts.

*Huamantupa C. 2485* (CUZ, MO) from Huaytampo, Cuzco, Peru is quite remarkable as it has been collected at an elevation of no less than 2800 m! Phylogenetic results using whole genome data indicate that the plants in Ecuador and Colombia may belong to a separate species from the plants of Brazil and Peru, although morphological analyses continue to place them in *Costus acreanus*.

2. ***Costus alfredoi*** Maas & H.Maas — Fig. 5b; Map 6

*Costus alfredoi* Maas & H.Maas in Maas et al. (2023) 79, f. 2. — Type: *Fuentes Claros et al. 6216* (holo MO [MO-2098346]; iso U [U1212213, U1212214]), Bolivia, La Paz, Prov. Bautista Saavedra, Area Natural de Manejo Integrado Apolobamba, between Paujeyuyo and Calzada, 1140 m, 16 Nov. 2003.

Herbs, c. 2 m tall. *Leaves*: sheaths 4–5 mm diam; ligule obliquely truncate, <1 mm long; petiole 5–7 mm long; sheaths, ligule and petiole glabrous; lamina linear, 15–23 by 1.5–2.5 cm, upper and lower side glabrous except for some hairs along the margin, base acute to obtuse, apex long-acute. *Inflorescence* ovoid, 11–12 by 5–6 cm, terminating a separate leafless shoot 50–85 cm long, sheaths obliquely truncate, 6–8 cm long, glabrous; bracts, bracteole, calyx, ovary, and capsule glabrous. *Flowers* abaxially orientated; bracts red, coriaceous, ovate, 3–4 by 1.5–2.5 cm, apex obtuse, callus 5–7 mm long; bracteole boat-shaped, 25–27 mm long; calyx 15–16 mm long, lobes shallowly ovate-triangular, 2–3 mm long; corolla yellow to orange, c. 50 mm long, glabrous, lobes narrowly elliptic, c. 30 mm long; labellum yellow, lateral lobes rolled inwards and forming a curved tube c. 10 mm diam, oblong-elliptic when spread out, c. 35 by 15 mm, middle lobe with yellow honey mark, 5–7–lobulate, lobules red, 5–6 mm long; stamen yellow, c. 35 by 7 mm, slightly exceeding the labellum, apex red, slightly dentate. *Capsule* ellipsoid, c. 15 mm long.

Distribution — Bolivia (La Paz).

Habitat & Ecology — In wet, subandine forests with *Oenocarpus bataua* Mart. (*Arecaceae*), at an elevation of c. 1140 m. Flowering in November.

IUCN Conservation Status — Not evaluated.

Notes — *Costus alfredoi* can be identified by its very narrow, grass-like leaves, a very small ligule not exceeding 1 mm in length, combined with a basal inflorescence, red, unappendaged bracts and yellow, tubular flowers. Another remarkable feature, rarely met with in the genus *Costus*, is the complete absence of any indument, except for a few scattered hairs bordering the leaf margin. It shares the last feature with *C. vargasii*, but from that species it differs by a shorter ligule (< 1 mm v. 5–35 mm long) and a much longer calyx (15–16 v. 6–10 mm) and bracteole (25–27 v. 12–22 mm).

3. ***Costus allenii*** Maas — Fig. 5c; Map 6

*Costus allenii* Maas (1972) 61, f. 28. — Type: *P.H. Allen 3587* (holo G [G00168450]; iso EAP [EAP103295]), Panama, Colón, Camp Piña, 25 m, 11 July 1946.

*Costus sp. B*: Maas (1972) 124.

Herbs, 0.5–4 m tall. *Leaves*: sheaths 8–20 mm diam; ligule obliquely truncate, 2–10 mm long, often reddish; petiole 5–25 mm long; sheaths, ligule and petiole densely brownish villose; lamina narrowly elliptic to (narrowly) obovate, 15–40(–60) by 4–14(–17) cm, lower side sometimes reddish, not or only very slightly plicate, upper and lower side densely brownish villose, base cordate to acute, apex acuminate (acumen 5–10 mm long). *Inflorescence* ovoid to subglobose, 4– 10 by 3–5 cm, enlarging to c. 30 by 12 cm in fruit, terminating a leafy shoot; bracts rather densely to densely brownish villose, bracteole, calyx, ovary, and capsule rather densely to densely puberulous to glabrous*. Flowers* adaxially orientated to erect; bracts green to yellowish green, coriaceous, broadly ovate, 3–5 by 3–5 cm, apex obtuse, callus 2–4 mm long; bracteole boat-shaped, 17–25 mm long; calyx pinkish red, 5–10 mm long, lobes shallowly triangular to deltate, 2–4 mm long; corolla yellowish white or salmon-coloured, 70–80 mm long, glabrous, lobes narrowly elliptic, 50–60 mm long; labellum yellow, upper part horizontally spreading, broadly obovate, 50–70 by 40–50 mm, lateral lobes striped with red, middle lobe recurved with a yellow honey mark, margin irregularly 5-lobed; stamen white, slightly tinged with yellow, 30–35 by 10–13 mm, not exceeding the labellum, apex red, obtuse to irregularly dentate, anther 8–10 mm long. *Capsule* ellipsoid, 15–20 mm long.

Distribution — Costa Rica, Panama, Colombia (Antioquia, Chocó, Valle del Cauca).

Habitat & Ecology — In moist rain forests, at elevations of 0–1200 m. Flowering in the rainy season.

IUCN Conservation Status — Not evaluated. There are 74 collection records examined of which 46 specimens were collected within the past 50 years. This species is also reported in 8 recent iNaturalist observations (https://www.inaturalist.org/observations?place_id=any&taxon_id=503943). The species is distributed in Colombia and Panama. Our observations indicate that this is a common species within its range and some of these areas are protected areas with no known specific threats. Population and threat data is not available but based on the number of collection records and our general field observations, this species would probably score a status of Least Concern.

Notes — *Costus allenii* is characterized by a dense, brownish villose indument of sheaths and leaves, unappendaged, green bracts and a yellow, red-striped labellum. Another very typical feature of *C. allenii* is the adaxial orientation of the flowers, a feature relatively rare in the genus.

We have united *C. sp. B* (Maas 1972: 124) (*Allen 3595* (F, G, U), Panama, Colón, Camp Piña, 25 m, 11 July 1946, the type locality of *C. allenii*) into *C. allenii* as most features of indument, leaves, and inflorescence fitted this species quite well.

4. ***Costus alleniopsis*** Maas & D.Skinner — Fig. 5d; Map 7

**Fig. 7.**
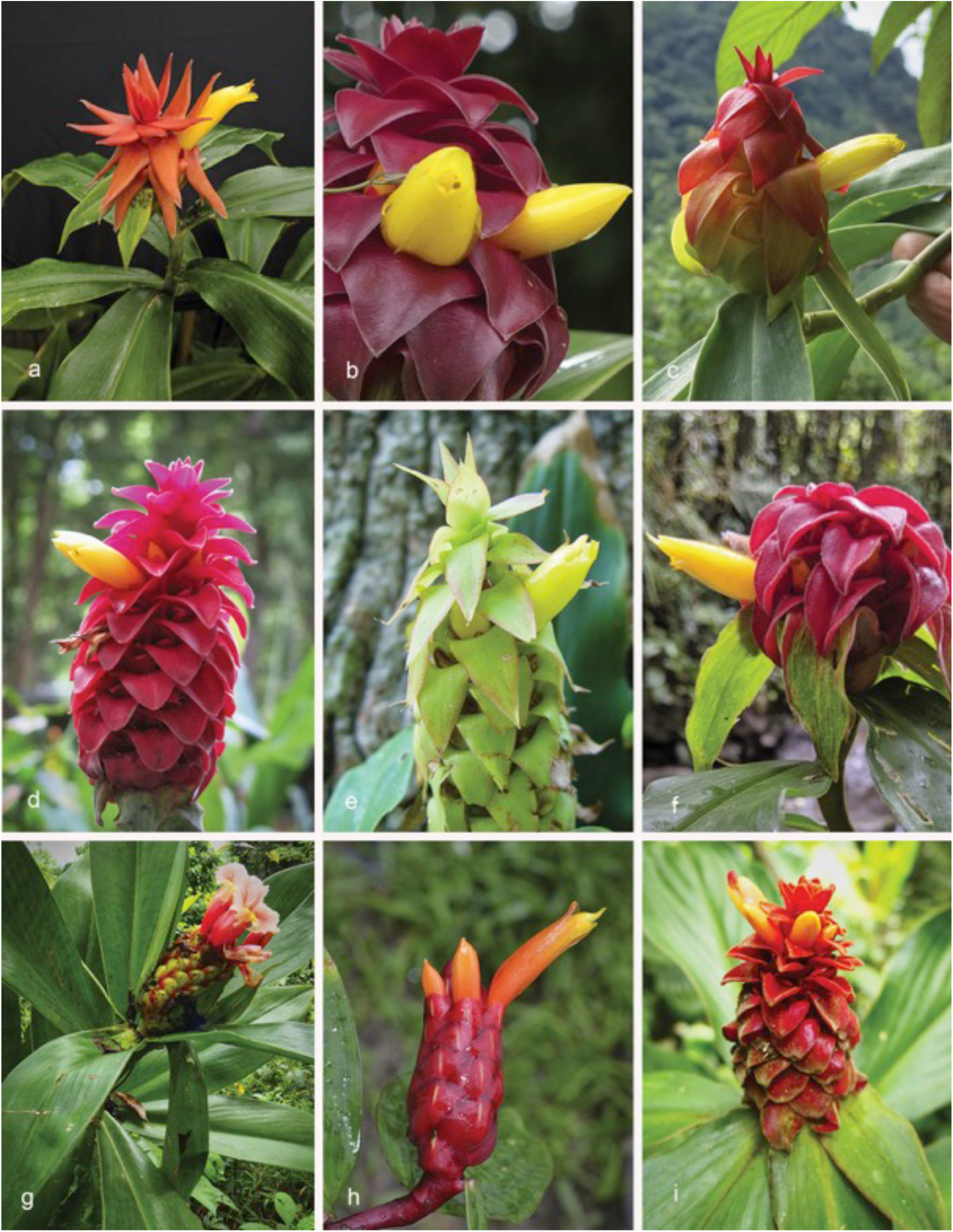
*Costus* species: *C. comosus – C. cupreifolius*: a*–*f. *C. comosus* (Jacq.) Roscoe: a. bakeri form (*D.Skinner R3312*, from Chiapas, Mexico); b. (*D.Skinner R3396*, from wild collection, Cochabamba, Bolivia); c. (*D.Skinner R3308*, Santa Maria de Boyaca, Colombia); d. horticultural form often listed as “*C. barbatus*” (*D.Skinner R2947*, cultivated plant, origin unknown); e. maritimus form (*D.Skinner R3208*, from Osa Peninsula, Costa Rica); f. (*D.Skinner R3500*, from Fortuna area, Chiriquí, Panama); g. *C. convexus* Maas & D.Skinner (photo in habitat in Putumayo, Colombia, not collected); h. *C. cordatus* Maas (*Maas 10658*, from Colombia); i. *C. cupreifolius* Maas (*D.Skinner R3462*, from type locality Putumayo, Colombia). Photos a–g & i by D. Skinner; h by P.J.M. Maas.

*Costus alleniopsis* Maas & D.Skinner in Maas et al. (2023) 81, f. 3. — Type: *Maas & Dressler 1652* (holo U [U1231561]; iso F, MO [MO-2193025]), Panama, Veraguas, Río Segundo Brazo, 700–750 m, 8 Sept. 1974.

Herbs, 0.5–1.2 m tall. *Leaves*: sheaths 10–20 mm diam; ligule obliquely truncate, 12–20 mm long, often reddish; petiole 10–25 mm long; sheaths, ligule and petiole densely brownish villose; lamina ovate to obovate, 25–32 by 13–16 cm, lower side sometimes reddish, 10–12-plicate, upper and lower side densely brownish villose, base cordate, acute, or obtuse, apex acuminate (acumen 10–15 mm long). *Inflorescence* ovoid, 6–7 by 3–5 cm, terminating a leafy shoot or rarely terminating a separate leafless shoot; bracts rather densely to densely brownish villose, bracteole, calyx, ovary, and capsule rather densely to densely puberulous to glabrous*. Flowers* adaxially orientated to erect; bracts green, rarely red, coriaceous, broadly ovate to ovate, 3–5 by 2–5 cm, apex obtuse, callus 5–7 mm long; bracteole boat-shaped, 20–30 mm long; calyx pink to red, 8–10 mm long, lobes shallowly triangular, 2–3 mm long; corolla yellow, 60–70 mm long, glabrous, lobes narrowly elliptic, 50–60 mm long; labellum yellow, upper part horizontally spreading, broadly obovate, 50–70 by 40–50 mm, lateral lobes striped with red, middle lobe recurved with a yellow honey mark, margin crenulate; stamen yellow, 40–50 by 12–15 mm, not exceeding the labellum, apex red, obtuse to irregularly dentate, anther 7–10 mm long. *Capsule* ellipsoid, 15–20 mm long.

Distribution — Panama (Coclé, Colón, Veraguas).

Habitat & Ecology — In forests, at elevations of 200–1300 m. Flowering in the rainy season.

IUCN Conservation Status — Not evaluated. There are only 17 collection records examined of which all 17 specimens were collected within the past 50 years. This species is also reported in 4 recent iNaturalist observations (https://www.inaturalist.org/observations?place_id=any&taxon_id=1460159). The species is distributed only in the provinces of Veraguas and Coclé in Panama. Our observations indicate that this is a fairly common species within its narrow range and is found in protected areas with no known specific threats. Population and threat data is not available but based on the number of collection records and our general field observations, this species is probably Near Threatened or Vulnerable.

Notes — *Costus alleniopsis* is restricted to Panama. It is morphologically closest to *C. allenii*, differing by a longer ligule (12–20 mm v. 2–10 mm) and distinctly plicate leaves. The orientation of the flowers is adaxial to erect.

5. ***Costus alticolus*** Maas & H.Maas — Fig. 5e; Map 7

*Costus alticolus* Maas & H.Maas in Maas et al. (2023) 83, f. 4. — Type: *Solheim & Reisfeld 1367* (holo WIS), Mexico, Oaxaca, Sierra Mazateca, Mun. Santa María Chilchotla, Cuahtemoc, c. 4 km NE of Santa María Chilchotla, near Clemencia and Santa Rosa, 1200 m, 16 Jan. 1984.

Herbs, 2–3.5 m tall. *Leaves*: sheaths 10–30 mm diam, green; ligule truncate, 4–12 mm long; petiole 5–20 mm long; sheaths, ligule and petiole densely to sparsely puberulous; lamina narrowly elliptic, 25–47 by 6–14 cm, upper side glabrous, lower side densely (particularly along the primary vein) sericeous to almost glabrous, base acute, apex acuminate (acumen 5–10 mm long). *Inflorescence* cylindric, 11–18 by 5–9 cm, terminating a separate leafless shoot 20–95 cm long, sheaths obliquely truncate, 4–7 cm long, rather densely puberulous to glabrous; bracts, bracteole, calyx, ovary, and capsule glabrous to densely puberulous*. Flowers* abaxially orientated; bracts red to bright crimson, coriaceous, ovate-oblong, 4–6 by 3–4 cm, apex obtuse and appressed, callus c. 10 mm long; bracteole boat-shaped, 30–35 mm long; calyx red, 13–21 mm long, lobes deltate to shallowly triangular, 3–6 mm long; corolla yellow to orange, 50–60 mm long, glabrous, lobes narrowly elliptic, c. 40 mm long; labellum yellow, lateral lobes rolled inwards and forming a slightly curved tube 18–20 mm diam, obovate when spread out, c. 60 by 35 mm, irregularly lobulate, lobules recurved, c. 4 mm long; stamen yellow, c. 65 by 11 mm, far exceeding the labellum, apex acute, anther c. 8 mm long. *Capsule* ellipsoid, 17–18 mm long.

Distribution — Mexico (Oaxaca).

Habitat & Ecology — In primary, wet montane to premontane cloud forests, with some mosses on trunks, drier on ridge crests, at elevations of 1040–1900 m. Flowering all year round.

IUCN Conservation Status — Not evaluated. There are only 6 collection records examined, all of which were collected within the past 50 years. This species is also reported in 11 recent iNaturalist observations (https://www.inaturalist.org/observations?place_id=any&taxon_id=1460168). The species is distributed only in the mountainous area of Oaxaca, Mexico. Our observations indicate that this is a fairly common species within its narrow range and is found in a protected indigenous area. It has a unique habitat for *Costus*, growing only in montane cloud forest. Population and threat data is not available but based on the number of collection records and our general field observations, this species would probably score as Near Threatened or Vulnerable.

Notes — *Costus alticolus* can be recognized by leafless flowering shoots up to almost 1 m long combined with large, red, unappendaged bracts, and tubular, yellow flowers in which the stamen far exceeds the labellum.

**6. *Costus amazonicus*** (Loes.) J.F.Macbr. — Fig. 5f; Map 7

*Costus amazonicus* (Loes.) J.F.Macbr. (1931) 13. — *Costus malortieanus* Wendl. var. *amazonicus* Loes. (1929) 710. —*Costus amazonicus* (Loes.) J.F.Macbr. subsp. *amazonicus* Maas (1972) 69. — Lectotype (designated by Maas 1972): *Tessmann 4903* (holo B, destroyed; lecto G [G00168449]; isolecto NY [NY00320338]), Peru, Loreto, lower Río Morona near confluence with Río Marañon, 160 m, 9 Jan. 1924.

Herbs, 1–3 m tall. *Leaves*: sheaths 10–30 mm diam; ligule obliquely truncate, 2–9 mm long; petiole 5–20 mm long; sheaths, ligule and petiole densely to sparsely villose, rarely glabrous; lamina narrowly obovate to narrowly elliptic, sometimes ovate to elliptic, 15–55 by 6–16 cm, sometimes 5–10-plicate, upper and lower side rather densely to densely villose, rarely glabrous, base acute to obtuse, apex acuminate (acumen 10–20 mm long). *Inflorescence* ellipsoid, cylindric, or subglobose, 6.5–15 by 3–7 cm, terminating a leafy shoot or terminating a separate leafless shoot 15–25 cm long, sheaths truncate, 4–5 cm long, rather densely villose to glabrous; bracts, bracteole, and calyx densely to sparsely puberulous to villose, ovary and capsule densely villose*. Flowers* abaxially orientated to erect; bracts green, rarely dark red, coriaceous, broadly ovate to broadly obovate, 2.5–4 by 2.5–4 cm, apex obtuse, margin falling apart into fibers, callus 5–6 mm long; bracteole boat-shaped, 20–32 mm long; calyx red, 10–19 mm long, lobes shallowly triangular to deltate, 1–5 mm long; corolla yellowish white to pink, 75–80 mm long, densely to rather densely puberulous to villose, lobes elliptic, 55–65 mm long; labellum white, pale yellow, or pink, upper part horizontally spreading, broadly obovate, 60–70 by 65–75 mm, lateral lobes striped with dark red to purple, middle lobe recurved with a yellow honey mark, margin crenulate to irregularly dentate; stamen pink to reddish, 50–55 by 12–15 mm, not exceeding the labellum, apex red, irregularly dentate, anther 5–9 mm long. *Capsule* ellipsoid, 10–12 mm long.

Distribution — Colombia (Amazonas, Caquetá, Cauca, Cundinamarca, Putumayo), Ecuador (Morona-Santiago, Napo, Orellana, Pastaza, Sucumbios, Zamora-Chinchipe), Peru (Amazonas, Loreto, Pasco, San Martín).

Habitat & Ecology — In non-inundated, lowland or submontane rain forests, at elevations of 200–1700 m. Flowering all year round.

IUCN Conservation Status — Not evaluated. There are 75 collection records examined of which 63 specimens were collected within the past 50 years. This species is also reported in 44 recent iNaturalist observations (https://www.inaturalist.org/observations?place_id=any&taxon_id=427388). The species is distributed in Colombia, Ecuador and Peru. Our observations indicate that this is a common species within its range and some of these areas are protected areas with no known specific threats. Population and threat data is not available but based on the number of collection records and our general field observations, this species would probably score a status of Least Concern.

Vernacular names — Colombia: Caña, Cañagria, Cañaguate, No ba cha. Ecuador: Gatunkagui (Huaorani), Gunequimonkagui (Huaorani). Peru: Shiwanuk, Shiwanuk-sinkusuk.

Notes — *Costus amazonicus* had been split into 2 subspecies by Maas (1972), we now consider them as two separate species: *C. amazonicus* with a hairy corolla and the margin of the bracts falling apart into fibers and *C. krukovii* with a glabrous corolla and bracts with an entire margin.

7. ***Costus antioquiensis*** Maas & H.Maas — Fig. 5g; Map 8

**Fig. 8.**
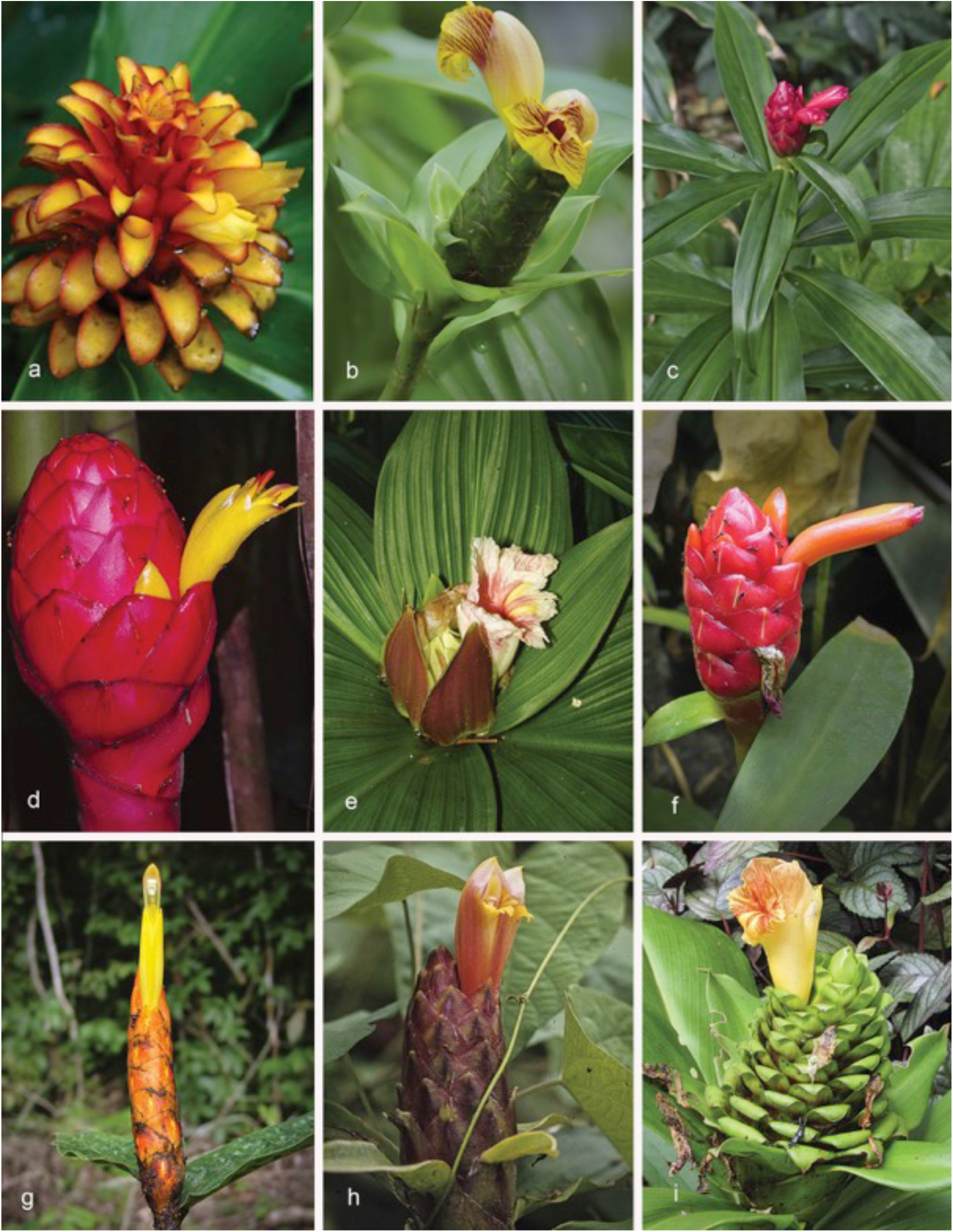
*Costus* species: *C. curvibracteatus – C. gibbosus*: a. *C. curvibracteatus* Maas (in cultivation by Carla Black, Fortuna, Panama); b. *C. dirzoi* García-Mend. & G.Ibarra (*D.Skinner R3448*, from type locality Los Tuxtlas, Veracruz, Mexico); c. *C. douglasdalyi* Maas & H.Maas (*D.Skinner R3482*, from Cruzeiro do Sul, Acre, Brazil); d. *C. erythrocoryne* K.Schum. (not collected, photographed in Ecuador); e. *C. erythrophyllus* Loes. (Photo from cultivated plant at Singapore Botanic Gardens); f. *C. erythrothyrsus* (*Maas 10623* from French Guiana); g. *C. fissicalyx* N.R.Salinas, Clavijo & Betancur B. (photo from original description in Caldasia 29(2) Bogotá July/Dec. 2007); h. *C. geothyrsus* K.Schum. (*Maas 2916*, Ecuador); i. *C. gibbosus* D.Skinner & Maas (*D.Skinner R3012*, cultivated at Waimea Gardens, Hawaii, 74S2037 from Plowman collection in Ecuador). Photos: a by C. Black; b, c & i by D. Skinner; d by G. Gerlach; e by J. Leong-Škornicková; f & h by P.J.M. Maas; g by L. Clavijo.

*Costus antioquiensis* Maas & H.Maas in Maas et al. (2023) 84, f. 5 & 6. — Type: *Maas et al. 10488* (holo L, 2 sheets [L.4496463, L.4496464]; iso JAUM, K, MO [MO-3159234, MO-3159235]), Colombia, Antioquia, Mun. San Rafael, along road from Guatapé to San Rafael, Río El Bizcocho, Finca El Castillo, 1059 m, 12 Oct. 2013.

Herbs, 0.5–5 m tall. *Leaves*: sheaths 10–20 mm diam; ligule truncate, 5–10 mm long, basally densely covered with a ring of brown, erect and rather stiff hairs (villose), sometimes absent; petiole 5–20 mm long; sheaths and petiole sparsely or sometimes densely villose; lamina narrowly elliptic, 30–50 by 9–13 cm, upper side glabrous, sometimes densely villose, lower side densely to sometimes sparsely villose or glabrous, base obtuse, apex acuminate (acumen 5–10 mm long). *Inflorescence* ovoid, 8–16 by 5–6 cm, enlarging to c. 25 by 7–8 cm in fruit, terminating a leafy shoot and wrapped tightly by the upper leaves; bracts, bracteoles, calyx, ovary, and capsule densely puberulous to villose, rarely glabrous*. Flowers* adaxially orientated; bracts green, coriaceous, broadly ovate, 3–4 by 3–4 cm, apex obtuse, callus 8–10 mm long; appendages rarely present; bracteole boat-shaped, 20–27 mm long; calyx (pale) red, 7–11 mm long, lobes deltate to shallowly triangular, c. 3 mm long; corolla yellow, pale orange, or pink, 60–70 mm long, glabrous, lobes elliptic, 40–60 mm long; labellum yellowish white, upper part horizontally spreading, broadly obovate, 50–70 by 40–60 mm, lateral lobes striped with red, middle lobe recurved with a dark yellow honey mark, margin irregularly dentate; stamen yellowish white to pink, c. 40 by 10–12 mm, not exceeding the labellum, apex dark red, entire to dentate, anther c. 8 mm long. *Capsule* obovoid, 25–28 mm long.

Distribution — Colombia (Amazonas, Antioquia, Arauca, Boyacá, Caldas, Cauca, Chocó, Córdoba, Cundinamarca, Guajira, Guaviare, Huila, Meta, Norte de Santander, Quindío, Risaralda, Santander, Tolima, Valle del Cauca, Vichada), Venezuela (Mérida), Ecuador (Azuay, Carchi, Pastaza, Zamora-Chinchipe), Peru (Ucayali).

Habitat & Ecology — In forests (a.o. “selva baja/alta perennifolia” or wet montane forests) or forested roadsides, at elevations of 50–1700 m. Flowering all year round.

IUCN Conservation Status — Not evaluated. There are 131 collection records examined of which 106 specimens were collected within the past 50 years. This species is also reported in 7 recent iNaturalist observations (https://www.inaturalist.org/observations?place_id=any&taxon_id=1460447). The species is widely distributed in Colombia and is also found in Ecuador and Venezuela. Our observations indicate that this is a common species within its range and some of these areas are protected areas with no known specific threats. Population and threat data is not available but based on the number of collection records and our general field observations, this species would probably score a status of Least Concern.

Vernacular names — Colombia: Caña agria, Cañagria, Cañaguate. Peru: Sacha huiro.

Notes — *Costus antioquiensis* is recognizable by adaxially orientated flowers and a ring of brown, erect and rather stiff hairs at the base of the ligule, which is often present; the lower leaf side and the bracts are often densely villose. *Costus antioquiensis* could be confused with *C. allenii*, but in that species the entire surfaces of the sheaths, ligule and petiole are villose. The ring of erect hairs at the base of the ligule is almost always present in this species, but lacking in *C. allenii*. Moreover, the upper side of the lamina in *C. antioquiensis* is mostly glabrous, whereas it is permanently villose in *C. allenii*. This species also looks similar to *C. kuntzei* (the former “*Costus laevis*”) and was misidentified as such in the past. It shares with that species the adaxially orientated flowers, but they are much darker red in *C. kuntzei*. Moreover, there are big differences in the indument, which is almost completely lacking in *C. kuntzei* whereas in *C. antioquiensis* the vegetative parts are mostly densely villose.

1. *C. antioquiensis* is highly variable in its indument. Most material has a quite dense villose indument all over the plant, but some specimens in the same population can be almost completely glabrous.

**8. *Costus arabicus*** L. — Fig. 5h; Map 9

**Fig. 9.**
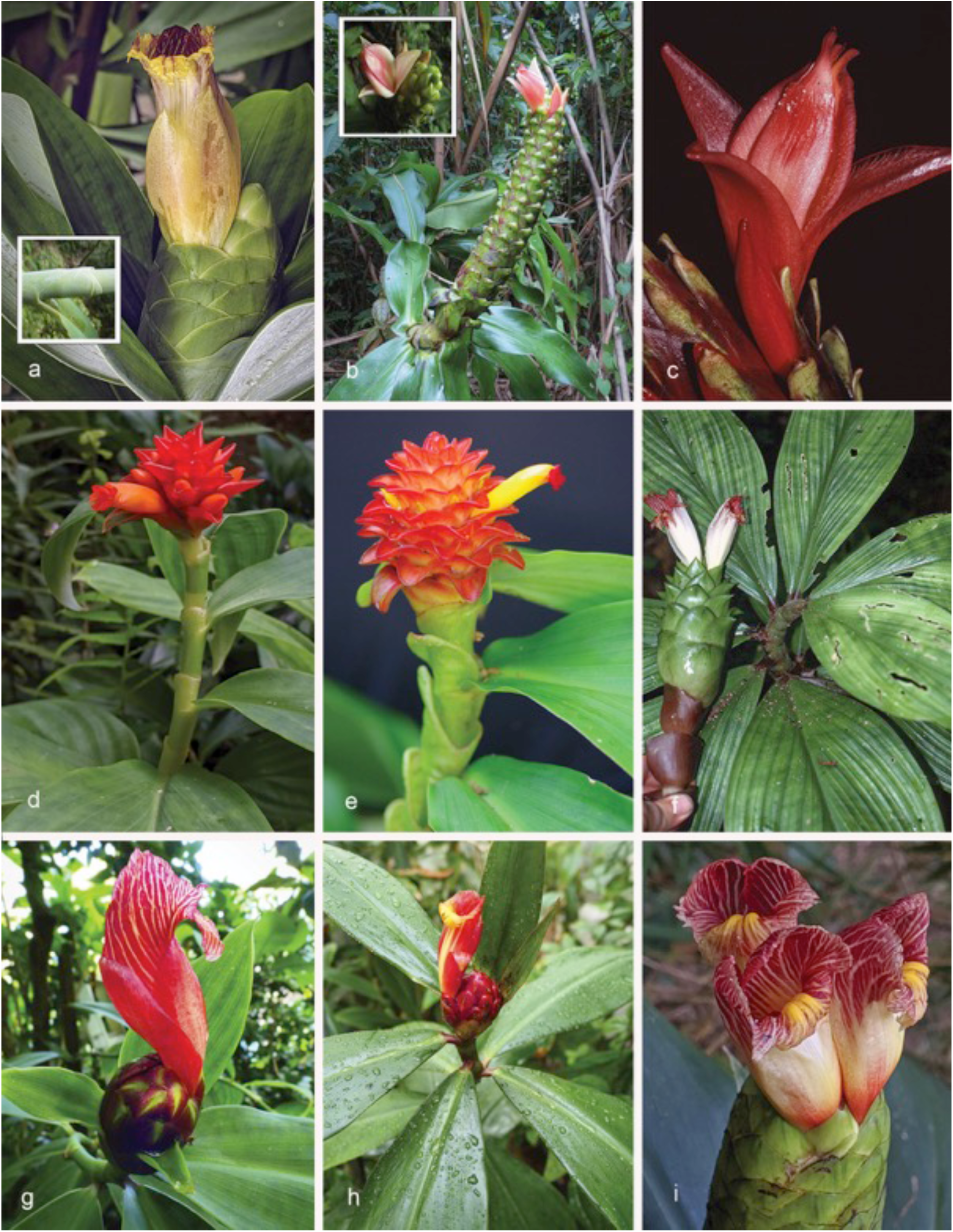
*Costus* species: *C. glaucus – C. kuntzei:* a. *C. glaucus* Maas (*D.Skinner R3363*, collected in Costa Rica at type locality CATIE (inset = close up of stem)); b. *C. guanaiensis* Rusby (*D.Skinner R3263* from Peru, Madre de Dios, Manu Learning Centre (inset = close up of flower)); c. *C. guanaiensis* Rusby, red floral form (*Maas 9933* from Bolivia); d. *C. juruanus* K.Schum. (*D.Skinner R3488* from near type locality in Brazil, Acre, Rio Tarauacá); e. *C. juruanus* K.Schum., cupped ligule form (*D.Skinner R3415* from Peru, Pasco, Bella Esperanza); f. *C. krukovii* (Maas) Maas & H.Maas (not collected, photo in Ecuador); g–i. *C. kuntzei* K.Schum.: g. (*D.Skinner R3086* from Costa Rica, Puntarenas, Reserva Durika); h. (cultivar ‘Lita Red’ *D.Skinner R3201* from Ecuador, Imbabura, Lita); i. (*D.Skinner R3508* from Honduras, Atlántica, Rio Cangrejal-Omega). Photos: a, b, d, e, g–i by D. Skinner; c by P.J.M. Maas; f by R. Foster.

*Costus arabicus* L. (1753) 2. — Type: *Drawing by Ehret of a plant cultivated in the Hortus Cliffortianus* (holo BM).

*Costus glabratus* Sw. (1788) 11. — Type. *Masson s.n*. (not seen), St. Lucia, probably not preserved.

*Costus niveo-purpureus* Jacq. (1809) 55, t. 67, f. 2 & t. 79. — *Costus sextus* Roem. & Schult. (1817) 26, nom. nud., nom. superfl. – *Costus glabratus* Sw. var. *niveo-purpureus* (Jacq.) Petersen (1890) 53. — Lectotype (designated here): *Plate 32* in Plumier, Msc. 5, based on material collected in Martinique.

*Costus niveus* G.Mey. (1818) 1 (‘*nivea*’). — Lectotype (designated here): *Rodschied s.n.* (lecto GOET [GOET011645]), Suriname, plantation Hamburg.

*Costus discolor* Roscoe (1825) t. 81. — Lectotype (designated here): *Table 81* of Roscoe 1825 (nothing of the original material introduced by Newman from Maranhão, Brazil, and cultivated at Liverpool has been preserved).

*Costus arabicus* Vell. (1829) 2, nom. inval., non L. (1753). — *Costus arrabidae* Steud. (1840) 426. — Lectotype (designated here): *Tab. 5* in Vellozo (1827); no specimen preserved.

*Costus verschaffeltii* Lem. (1854) t. 381. (‘*verschaffeltianus*’). — Lectotype (designated here): *Table 381* in Lemaire (1854), since nothing of the original material, cultivated in Gand, Belgium by Mr. F. de Vos from Ilha Sta. Catarina, Brazil, has been preserved.

*Costus spiralis* (Jacq.) Roscoe var. *hirsutus* Petersen (1890) 55. — Lectotype (designated here): *Regnell III 1213* (lecto S [S-R-1262]; iso US [US00092978]), Brazil, Minas Gerais, Uberaba, Jan. 1849.

*Costus pubescens* S.Moore (1895) 481. — Lectotype (designated here): *Spencer Moore 813* (holo B [BM000923836]), Brazil, Mato Grosso, banks of Rio Paraguay between Santa Cruz and Villa Maria, “1891, 1892”.

*Costus brasiliensis* K.Schum. (1904) 403. — Lectotype (designated here): *Lindman A 2767* (lecto S [S-R-1254]; isolecto S [S-R-6269]), Brazil, Mato Grosso, Matto do Curupira, Febr. 1894.

*Costus pilgeri* K.Schum. (1904) 404. — Lectotype (designated by Maas 1972): *Hassler 5974* (lecto G [G00009249]; isolecto BM [BM000087262], G [G0004769], K [K000586753], NY [NY00320344], US), Paraguay, Río Tapiraguay, Yerbales, Sierra de Maracayu, 1900.

*Costus validus* Loes. (1929) 711. — Lectotype (designated by Maas 1972): *Ule 9196* (holo B, destroyed; lecto K [K000586773]), Peru, Madre de Dios, Seringal Auristella, Apr. 1911.

*Costus gracilis* Loes. (1929) 711. – Type: *Ule 95 br* (holo B, destroyed), Peru, Madre de Dios, Seringal Auristella, Apr. 1911.

*Costus pubescens* S.Moore forma *fibrillosus* Loes. (1929) 711 (‘*fibrillosa*’). — Type: *F.C. Hoehne 1634* (not seen), Brazil, Mato Grosso, Tapiraguã, May 1909.

*Costus ramosus* Woodson (1939) 171. — Type: *A.C. Smith 2869* (holo MO, 2 sheets [MO-1515252 & MO-1515253]; iso F [F0047182F], GH [GH00030660], K [K000586752], NY [NY00320350, NY00320351], U [U0007227], US [US00092975]), Guyana, Basin of Shodikar Creek, Essequibo tributary, 8–22 Jan. 1938.

Herbs, 1–3 m tall, often branched at the base of the inflorescence. *Leaves*: sheaths 5–15 mm diam; ligule truncate, 2–10 mm long; petiole 2–7 mm long; sheaths, ligule and petiole glabrous to densely puberulous to villose; lamina narrowly ovate to narrowly obovate, 10–25 by 3–10 cm, upper side glabrous, lower side glabrous to densely puberulous to villose, base cordate or rarely rounded, apex acuminate (acumen 5–15 mm long). *Inflorescence* ovoid to cylindric, 3–15 by 2.5–5 cm, enlarging to c. 20 by 7 cm in fruit, terminating a leafy shoot or rarely terminating a separate leafless shoot 35–50 cm long, sheaths obliquely truncate, 3–6 cm long, densely puberulous, lateral branches often arising from the base of the inflorescence; bracts, appendages of bracts, bracteole, calyx, ovary, and capsule glabrous to sparsely or rarely densely puberulous*. Flowers* abaxially orientated to erect; bracts green, coriaceous, broadly ovate to ovate, 2.5–4.5 by 1.5–3 cm, apex acute or obtuse, callus 3–7 mm long; appendages mostly absent, when present green, foliaceous, ascending to reflexed, broadly to shallowly triangular, 1–2 by 1–1.5 cm, apex acute; bracteole boat-shaped, very rarely 2-keeled, 20–35 mm long; calyx pinkish red or slightly tinged with green, (10–)12–22(–27) mm long, often exceeding the bracts, lobes deltate, 2–6 mm long; corolla white, 60–70 mm long, glabrous, lobes elliptic to obovate, 40–50 mm long; labellum white, upper part horizontally spreading, broadly obovate, 50–70 by 50–70 mm, lateral lobes sometimes tinged with purple or red, middle lobe recurved with a yellow honey mark, margin irregularly lobed to dentate; stamen white, 40–50 by 10–15 mm, not or slightly exceeding the labellum, apex sometimes tinged with purple, irregularly dentate, anther 6–12 mm long. *Capsule* ellipsoid, 10–18 mm long.

Distribution — The Greater Antilles (Dominican Republic, Haiti), the Lesser Antilles (Guadeloupe, Martinique, Santa Lucia), Colombia (Amazonas, Antioquia, Bolívar, Caquetá, Casanare, Cesar, Guainía, Guaviare, Meta, Santander, Vaupés, Vichada), Venezuela (Amazonas, Anzoátegui, Apure, Barinas, Bolívar, Carabobo, Cojedes, Delta Amacuro, Mérida, Miranda, Monagas, Nueva Esparta, Sucre, Trujillo, Zulia), Trinidad and Tobago, Guyana, Suriname, French Guiana, Ecuador (Los Ríos, Napo, Orellana, Pastaza, Sucumbios), Peru (Huánuco, Loreto, Madre de Dios, Tumbes, Ucayali), Bolivia (Beni, Cochabamba, La Paz, Santa Cruz), Brazil (Acre, Amapá, Amazonas, Bahia, Distrito Federal, Espírito Santo, Goiás, Maranhão, Mato Grosso, Mato Grosso do Sul, Minas Gerais, Pará, Paraná, Rio de Janeiro, Rondônia, Roraima, Santa Catarina, São Paulo, Tocantins), Paraguay (Alto Paraguay, Amambay, Canindeyú, Central, Concepción, Cordillera, Paraguari, San Pedro), Argentina (Misiones).

Habitat & Ecology — In periodically inundated forests (restinga, tahuampa), margins of forests, forest islands, or savannas, in swamps, or in wet secondary vegetation, mostly at low elevations, rarely up to 1000 m. Flowering all year round.

IUCN Conservation Status – Not evaluated. There are 789 collection records examined of which 472 specimens were collected within the past 50 years. This species is also reported in 158 recent iNaturalist observations (https://www.inaturalist.org/observations?place_id=any&taxon_id=504230). The species is widely distributed throughout the Amazonian region of South America, the Guianas and as far south as Argentina and Paraguay. Our observations indicate that this is a common species within its range and some of these areas are protected areas with no known specific threats. Population and threat data is not available but based on the number of collection records and our general field observations, this species would certainly score a status of Least Concern.

Vernacular names — Bolivia: Buga, Cana fistula, Caña agria, Shico. Brazil: Anana brava, Cana de macaco, Cana de macaco branco, Cana do brejo, Caña fistula, Canafistula, Canarana, Canha braba, Canha brava, Grana de macaco, Pacó caatinga. Colombia: Abebe blanco, Cañagria, Kaw-shak (Puinare), Nane, Naneiboto. Ecuador: Sacha chibilia (Quichua). French Guiana: Ago singaafu (Boni), Canne congo (Créole), Kapiwa asikalu (Wayãpi), Kopiya (Wayãpi), Maikaman (Wayana), Singafu, Tuiu-cinõ (Palikur), Tuiu waik mwiunõ (Palikur). Guyana: Congo cane, Eseyundu, White congo cane. Paraguay: Caña brava. Peru: Agrio-wiru, Caña agria de la blanca, Caña caña colorada, Caña gamboa, Sacha huiro, Sachahuiro, Sachahuiro blanco. Suriname: Reddi sangrafoe, Sanganafoe, Sangrafoe, Siewiejoeloe (Carib), Weti sangaafu (Saramaccan). Venezuela: Caña de india, Caña de la india, Caña guinea.

Notes — *Costus arabicus* is rather easily recognizable by the combination of green, generally unappendaged bracts, often completely white flowers, and leaves which very often have a cordate base. Another feature which is often observed is the presence of lateral, axillary branches arising all along the shoot.

In part of the distribution area (Bolivia (Beni), Brazil (Acre, Roraima, and Rondônia)) the bracts are sometimes provided with leafy appendages. In an earlier stage (Maas 1977: 189) we assumed that this material might represent an undescribed species and provisionally placed it under *Costus* aff. *claviger*. After having seen and studied this form in the field (in Beni, Bolivia) we came to the conclusion that it perfectly matches *C. arabicus* (in features of leaves, bracts, flowers, etc.) and that it should be included in that species.

As previously discussed (Maas 1972: 60, 61), the indument of the lower side of the leaves is rather variable: in plants of the northern part of the area the lower leaf side usually is glabrous, but in the southern part of the area (Bolivia, Paraguay, and southeastern Brazil) it is often puberulous to villose and soft to the touch. As the transition from glabrous to villose is gradual, with all kinds of intergradations,we have not used this character for taxonomic distinction.

The leaves of this species can be variegated in cultivated plants.

9. ***Costus asplundii*** (Maas) Maas — Fig. 5i; Map 10

**Fig. 10.**
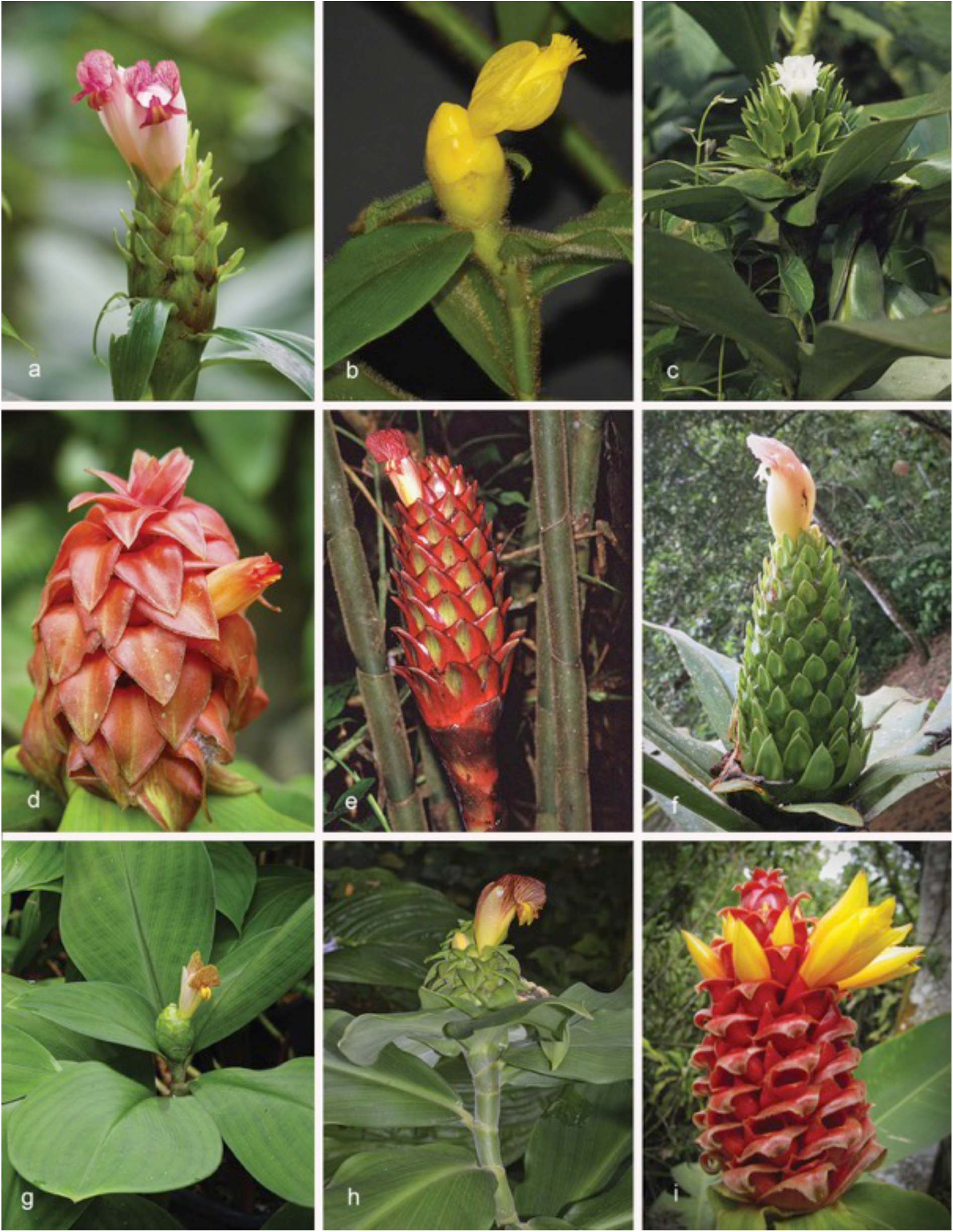
*Costus* species: *C. laevis – C. montanus*: a. *C. laevis* Ruiz & Pav. (*D.Skinner R3414* from Peru, Junín, Puente Perené); b. *C. lasius* Loes. (*D.Skinner R3127* from Peru, Loreto, Urco Mirano); c. *C. leucanthus* Maas (*Maas & Plowman 1920* from Colombia, Valle de Cauca); d. *C. lima* K.Schum. (not collected, photo from Costa Rica, Puntarenus, Ro Barrigones); e. *C. longibracteolatus* Maas (not collected, photo from Ecuador); f. *C. macrostrobilus* K.Schum. (not collected, photo in habitat near Bahia Solano, Chocó, Colombia); g. *C. malortieanus* H.Wendl. (*D.Skinner R1562* origin unknown, plant in cultivation); h. *C. mollissimus* Maas & H.Maas (plant in cultivation at Burgers Bush); i. *C. montanus* Maas (*D.Skinner R2972*, from Lancester Gardens, Costa Rica). Photos: a, b, d, f, g & i by D. Skinner; c & h by P.J.M. Maas; e by R. Foster.

*Costus asplundii* (Maas) Maas (1976) 470. — *Costus guanaiensis* Rusby var. *asplundii* Maas (1972) 52. — Type: *Asplund 19115* (holo S [S-R-6749]; iso B [B100108558], NY [NY00320332], R [R000125356], U [U0007228]), Ecuador, Pastaza, Mera, Río Pastaza, 1000 m, 30 Jan. 1956.

Herbs, 1.5–5 m tall. *Leaves*: sheaths 10–35 mm diam; ligule truncate, 2–20 mm long; petiole 5– 10 mm long; sheaths, ligule and petiole densely or sometimes sparsely brownish villose; lamina narrowly elliptic, 20–50 by 4–10 cm, lower side sometimes reddish, upper side glabrous, sometimes sparsely to rather densely brownish villose, lower side densely to rather densely brownish villose, base acute to obtuse, apex long-acute to acuminate (acumen 10–15 mm long). *Inflorescence* often nodding, ovoid, 6–15 by 6–7 cm, terminating a leafy shoot; bracts, appendages of bracts, bracteole, calyx, ovary, and capsule glabrous to densely puberulous*. Flowers* abaxially orientated; bracts red or green, coriaceous, broadly ovate, 2.5–4 by 2.5–4 cm; appendages green or sometimes reddish brown and basally red, foliaceous, reflexed or horizontally spreading to slightly recurved in living plants, broadly triangular, 1–4 by 1–4 cm, apex acute; bracteole boat-shaped, 25–30 mm long; calyx bright red, 13–20 mm long, lobes deltate to shallowly triangular, 3–7 mm long; corolla white, pale pink, or pale yellow, 70–90 mm long, glabrous, lobes elliptic, 60–75 mm long; labellum white, yellow, or pink, upper part horizontally spreading, broadly obovate, c. 75 by 70–80 mm, lateral lobes white, yellow to pink, sometimes striped with red, middle lobe recurved with a yellow honey mark, margin irregularly dentate; stamen pale pink to pale red, 45–60 by 15–20 mm, not exceeding the labellum, apex yellow to red, irregularly dentate, anther 8–15 mm long. *Capsule* ellipsoid, 15–20 mm long.

Distribution — Colombia (Caquetá, Cauca, Putumayo), Ecuador (El Oro, Los Rios, Morona-Santiago, Napo, Orellana, Pastaza, Sucumbios, Tungurahua, Zamora-Chinchipe), Peru (Amazonas, Loreto, San Martín).

Habitat & Ecology — In lowland or premontane forests, or along roadsides and streams, at elevations of 0–2000 m. Flowering all year round.

IUCN Conservation Status — Not evaluated. There are 97 collection records examined of which 84 specimens were collected within the past 50 years. This species is also reported in 28 recent iNaturalist observations (https://www.inaturalist.org/observations?place_id=any&taxon_id=505461). The species is widely distributed in Colombia, Ecuador and Peru. Our observations indicate that this is a common species within its range and some of these areas are protected areas with no known specific threats.Population and threat data is not available but based on the number of collection records and our general field observations, this species would probably score a status of Least Concern.

Vernacular names — Colombia: Caña agria, Caña agria negra lanudo, Caña lanuda. Ecuador: Ayahuiru, Caña agria, Kauruntuntup (Shuar), Kauruntuntu (Shuar), Wanoka unki (Huaorani). Peru: Mun siwanuk, Unt untunt (Huambisa), Untuntu (Huambisa).

Notes — *Costus asplundii* can be recognized by bracts provided with often reflexed, green appendages, large, relatively narrow leaves, and densely brownish villose leaf sheaths (particularly the ligule). The form found in Ecuador in Napo and Orellana differs from the type form by having red bract appendages and pink flowers as well as a reddish sheath and petioles. This beautiful form of *C. asplundii* is cultivated around the world.

**10. *Costus asteranthus*** Maas & H.Maas — Fig. 6a; Map 10

*Costus asteranthus* Maas & H.Maas (1990) 307, f. 1 & 2. — Type: *Maas et al. 6086* (holo U, 3 sheets [U0007229, U0007230, U0007231]; iso F [F0040740F], K [K000586768], MO [MO-258427], NY [NY00039319], USM [USM001035]), Peru, Puno, Prov. Carabaya, vicinity of San Gabán (= Lanlacuni Bajo), 600–800 m, 18 Oct. 1984.

Herbs, c. 2 m tall. *Leaves*: sheaths 8–15 mm diam, young ones somewhat glaucous; ligule truncate, 1–2 mm long; petiole 10–20 mm long; sheaths, ligule and petiole glabrous; lamina (narrowly) elliptic to (narrowly) obovate, 18–32 by 6–13 cm, upper side glabrous, lower side subglabrous, base acute, apex acute to acuminate (acumen to c. 20 mm long). *Inflorescence* ellipsoid to ovoid, 6–10 by 3–4 cm, terminating a separate leafless shoot to c. 85 cm long, sheaths obliquely truncate, to c. 6 cm long, glabrous; bracts, bracteole, calyx, ovary, and capsule glabrous*. Flowers* abaxially orientated to erect; bracts red, coriaceous, obovate-triangular, 3.5– 4.5 by 2–2.5 cm, apex obtuse, callus 5–10 mm long; bracteole boat-shaped, 22–35 mm long; calyx pink, 17–22 mm long, lobes shallowly triangular, c. 2 mm long; corolla yellow, 75–80 mm long, longer than all other flower parts, glabrous, lobes narrowly elliptic, spreading, 65–70 mm long; labellum yellow, lateral lobes rolled inwards and forming a tube 10–15 mm diam, obtriangular when spread out, 40–45 by 20–28 mm, 5-lobulate, lobules recurved, 3–10 mm long; stamen yellow, 50–55 by 10 mm, exceeding the labellum, apex dentate, anther c. 10 mm long. *Capsul*e ellipsoid, c. 12 mm long.

Distribution — Peru (Puno).

Habitat & Ecology — In forests, on steep slopes, at elevations of 600–800 m. Flowering in October.

IUCN Conservation Status — Not evaluated. Only known from the type specimen collected in 1984. This species is only reported in 1 recent iNaturalist observation (https://www.inaturalist.org/observations?place_id=any&taxon_id=700309) made by D.Skinner at the same site as the type locality. The species is distributed only in a small area near the town of San Gabán in the province of Puno, Peru. This site was visited by Skinner in 2018 and found to be under severe threat due to frequent avalanches and a large bridge construction project in its locality. This species would probably be assessed as Critically Endangered if formally assessed for IUCN status.

Notes — *Costus asteranthus*, a narrow endemic of the Peruvian department of Puno, is well recognizable by an inflorescence terminating a separate leafless shoot, red bracts, a long calyx (17–22 mm), but particularly by a ligule of only 1–2 mm long, and spreading corolla lobes, creating a star-like appearance (the feature after which this species was named). Another peculiarity of this species is that the corolla is distinctly exceeding the labellum and the stamen in length, a feature not often seen in the genus *Costus.* The young sheaths and leaves are covered with a glaucous coating.

**11. *Costus atlanticus*** E.Pessoa & M.Alves — Fig. 6b; Map 10

*Costus atlanticus* E.Pessoa & M.Alves in Pessoa et al. (2020) 50, f. 5. — Type: *Pessoa et al. 1346* (holo UFP; iso RB, SP), Brazil, Pernambuco, Caruaru, Brejo dos Cavalos, Parque Ecológico Municipal, 1100 m, 3 Jul 2017.

Herbs, 1–2 m tall. *Leaves*: sheaths 8–15 mm diam; ligule truncate, 2–10 mm long; petiole 2–8 mm long; sheaths sparsely puberulous to villose; lamina narrowly elliptic to narrowly obovate, 6–21 by 3–8 cm, upper side glabrous or hairy along the primary vein, lower side glabrous or sparsely puberulous to villose, base acute, rounded, or slightly cordate, apex acute to acuminate (acumen c. 10 mm long). *Inflorescence* globose, ovoid, or ellipsoid, 5–11 by 2.5–7 cm, terminating a leafy shoot; bracts, bracteole, calyx, ovary, and capsule sparsely puberulous*. Flowers* abaxially orientated to erect; bracts red, coriaceous, broadly ovate, 3–4 by 2.5–4.5 cm, apex obtuse, callus 8–10 mm long; bracteole boat-shaped, 22–26 mm long; calyx red, 9–16 mm long, lobes shallowly triangular to deltate, 2–3 mm long; corolla pinkish to white, 60–65 mm long, glabrous, lobes narrowly obovate, 50–55 mm long; labellum pink to white, obovate when spread out, funnel-shaped, 45–50 by 35–50 mm, upper margin reflexed, middle part with yellow honey mark; stamen pale pink, 30–35 by 10–12 mm, not exceeding the labellum, apex acute to 3-lobed, anther c. 10 mm long. *Capsule* globose, 9–12 mm diam.

Distribution — Brazil (Pernambuco).

Habitat & Ecology — In semideciduous seasonal forests, at elevations of 200–1100 m. Flowering all year round.

IUCN Conservation Status — Not evaluated. This species is widely cultivated in Brazil but there are only 3 collection records from the wild, all of which are in the state of Pernambuco in Brazil. There is still some doubt as to whether this is a valid species or a hybrid of *Costus arabicus* with *Costus spiralis*. There are 17 observations of this species posted to iNaturalist (https://www.inaturalist.org/observations?place_id=any&taxon_id=1232590), but none of them can be clearly determined as growing in the wild vs. being cultivated. Based on these issues, it is difficult to assess the probable IUCN status. If this is truly a natural species and not a hybrid, it would probably score as Critically Endangered due to the lack of known specimens growing wild.

Notes — *Costus atlanticus* looks quite similar to *C. spiralis* by its inflorescence. The funnel-shaped flowers are, in our opinion, intermediate between the tubular flowers of *C. spiralis* and the horizontally spreading labellum of *C. arabicus*.

*Costus atlanticus* is widely cultivated throughout Brazil, and its form has been registered with the cultivar name ‘Tropicais’. Hundreds of inaturalist observations are from urbanized areas in Brazil where the plants are growing in cultivation; the only known naturally occuring plants of this species are from the state of Pernambuco.

**12 *Costus barbatus*** Suess. — Fig. 6c; Map 11

**Fig. 11.**
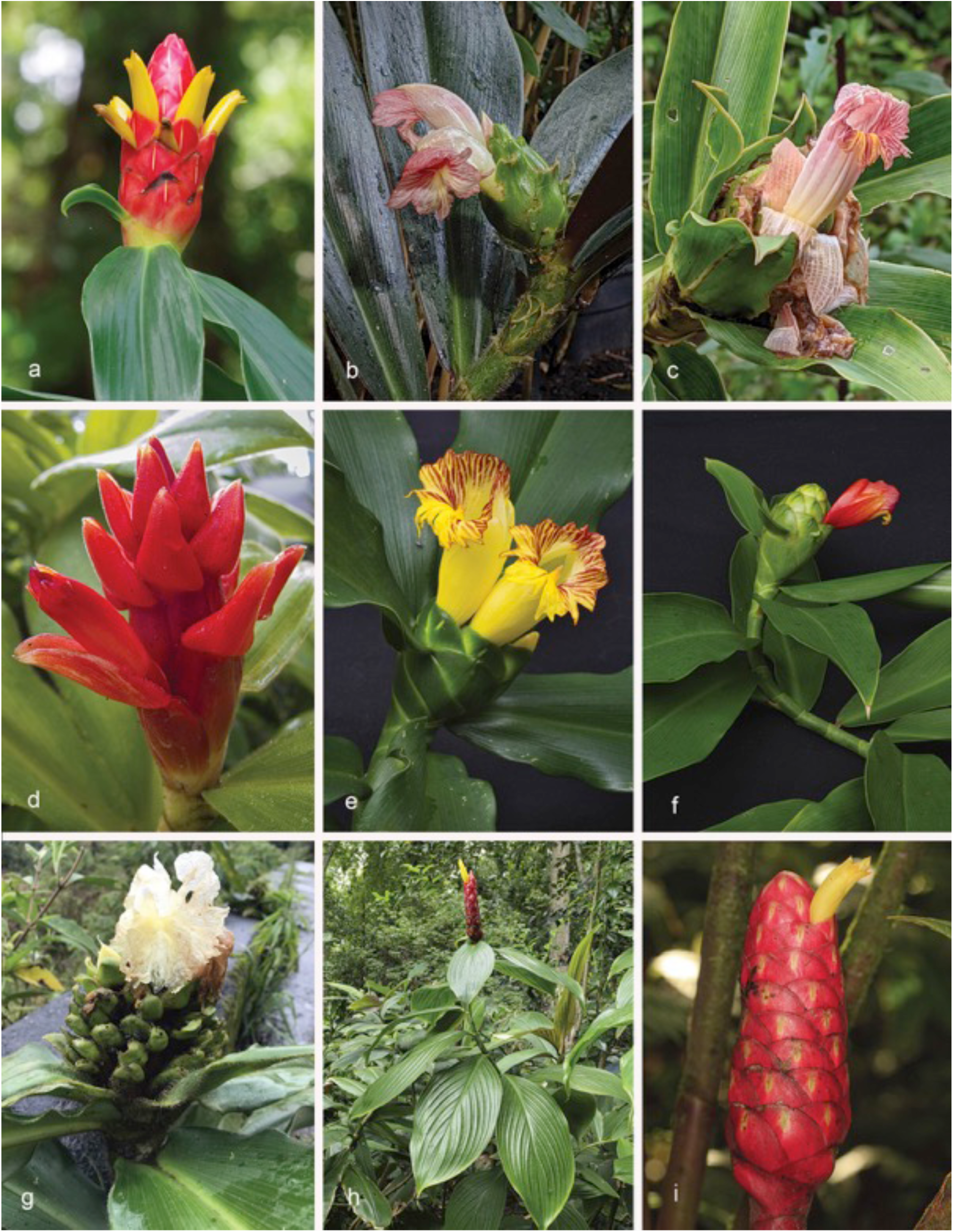
*Costus* species: *C. nitidus – Costus plowmanii*: a. *C. nitidus* Maas (*D.Skinner R3088* from Costa Rica, Limón); b. *C. obscurus* D.Skinner & Maas (*D.Skinner R3322* from Peru, Madre de Dios, Pantiacolla); c. *C. oreophilus* Maas & D.Skinner (*D.Skinner R3523* from Ecuador, Tungurahua); d. *C. osae* Maas & H.Maas (*D.Skinner R2566*, origin unknown, from plant in cultivation); e, f. *C. pictus* D.Don (e) pattern stem form (*D.Skinner R3453* from Mexico, Veracruz); f. red flower Pacific coastal form (*D.Skinner R3453* from Mexico, Chiapas); g. *C. pitalito* C.D.Specht & H.Maas (photographed in Colombia, Huila, near Pitalito); h. *C. plicatus* Maas (*D.Skinner R3211* from Costa Rica, Puntarenas); i. *C. plowmanii* Maas (*J.Betancur 14570* from Colombia). Photos: a–f & h by D. Skinner; g by C.D. Specht; i by J. Betancur.

*Costus barbatus* Suess. (1942) 295. — Type: *Kupper 947* (holo M0-242449), Costa Rica, San José, Curridabat, c. 1200 m, 2 Apr. 1932.

Herbs, 1.5–3 m tall. *Leaves*: sheaths 10–20 mm diam; ligule (10–)15–30 mm long, truncate to unequally 2-lobed, lobes rounded to obtuse, upper margin red; petiole 6–12 mm long; sheaths, ligule and petiole densely to sparsely puberulous to villose; lamina narrowly elliptic, 13–26 by 5–8 cm, upper side glabrous, sometimes sparsely puberulous, lower side rather densely to densely puberulous to villose, base acute to obtuse, apex acuminate (acumen 5–15 mm long). *Inflorescence* cylindric, ovoid to fusiform, 4–10 by 2.5–4.5 cm, terminating a leafy shoot; bracts, appendages of bracts, bracteole, calyx, ovary, and capsule densely to sparsely puberulous to villose*. Flowers* abaxially orientated; bracts red to reddish orange, coriaceous, broadly ovate, 3–4 by 2–4 cm, apex acute or slightly rounded and sometimes reflexed, margin often falling apart into fibers, callus absent or inconspicuous; appendages red to green, patent to recurved, or absent, 0.2–1.5 by 0.5–1.5 cm (lowermost), apex acute; bracteole boat-shaped, 22–25 mm long, margin falling apart into fibers; calyx red, 12–15 mm long, lobes shallowly triangular, 1–3 mm long, margin falling apart into fibers; corolla yellow tinged with orange, 30–40 mm long, rather densely to sparsely puberulous, lobes narrowly elliptic, 25–30 mm long; labellum yellow, lateral lobes rolled inwards and forming a slightly curved tube 8–9 mm diam, broadly ovate when spread out, 25–30 by 15–25 mm, irregularly 5-lobulate, lobules 1–2 mm long; stamen pale yellow, 25–30 by 8–10 mm, not exceeding the labellum, apex obtuse to irregularly dentate, anther 5–9 mm long. *Capsule* obovoid, 10–12 mm long.

Distribution — Costa Rica.

Habitat & Ecology — Along rivers or in wet forests, at elevations of 1200–1700 m. Flowering all year round.

IUCN Conservation Status — Critically endangered.

Notes — *Costus barbatus*, endemic to Costa Rica, is morphologically probably closest to *C. pulverulentus* and *C. montanus*, differing from the widely spread *C. pulverulentus* in having a hairy corolla and a much longer ligule, and from *C. montanus*, another Costa Rican endemic,in the shape and length of its ligule.

For additional information of this species see Skinner (2011).

**13. *Costus beckii*** Maas & H.Maas — Fig. 6d; Map 12

**Fig. 12.**
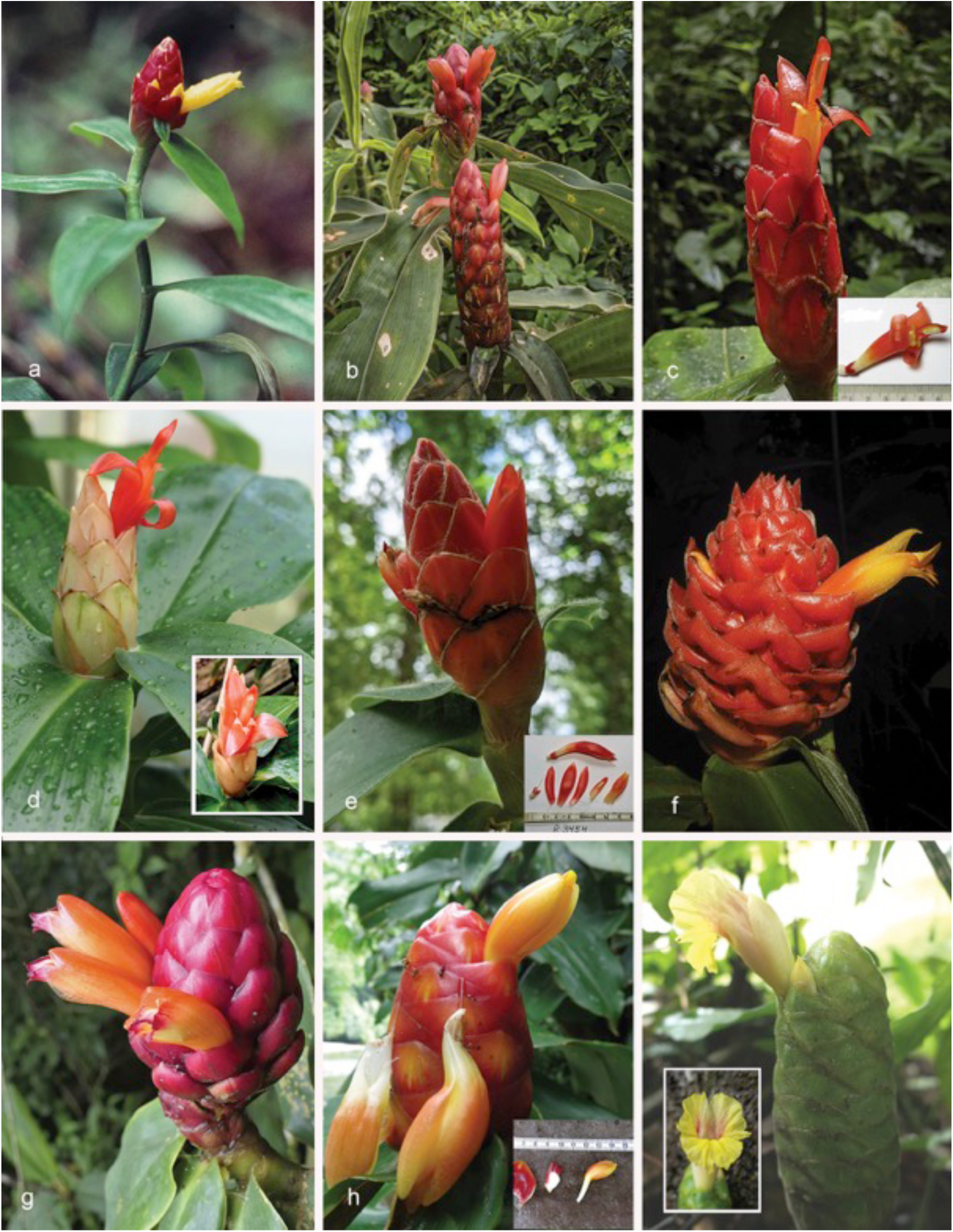
*Costus* species: *C. prancei – C. sepacuitensis*: a. *C. prancei* Maas & H.Maas (*Prance 12265* from Brazil, Acre, Serra do Moa); b. *C. pseudospiralis* Maas & H. Maas (not collected, photo in habitat Brazil, Acre, Xapurí); c–e: *C. pulverulentus* C.Presl: c. Central American form (not collected, photo from Panama, Veraguas (inset = dissected floral organs)); d. short, pink-bract form (D.Skinner R3231 cultivated in Colombia, Quindio Botanical Garden (inset = dissected floral organs)); e. Mesoamerican form (*D.Skinner R3454* from Mexico, Oaxaca near La Esmeralda (inset = flower)); f. *C. ricus* Maas & H.Maas (photo from Costa Rica, Puntarenas, Osa Peninsula); g. *C. rubineus* D.Skinner & Maas (*D.Skinner R3421* from Peru, Pasco, Oaxapampa); h. *C. scaber* Ruiz & Pav. (*D.Skinner R3484* from type locality in Peru, Huanuco, Rio Derrepente (inset = dissected floral organs)); i. *C. sepacuitensis* Rowlee (*D.Skinner R3402* from Mexico, Oaxaca (inset = closeup of flower)). Photos: a by P.J.M. Maas; b–e & g–i by D. Skinner; f by R. Aguilar.

*Costus beckii* Maas & H.Maas (1990) 309, f. 3. — Type: *Beck 7324* (holo U, 2 sheets [U0007233, U0007234]; iso K [K000586770], LPB [LPB0000393], NY [NY00381820, NY00381824]), Bolivia, Cochabamba, Prov. Chapare, Villa Tunari, 14 km towards Palmar, foot of mountains, 550 m, 24 Nov. 1981.

Herbs, 1.5–3.5 m tall. *Leaves*: sheaths 10–30 mm diam; ligule unequally 2-lobed, 25–60 mm long; petiole 10–20 mm long; sheaths, ligule and petiole densely to rather densely villose; lamina narrowly elliptic to narrowly obovate, 30–43 by 7–15 cm, upper side glabrous to densely villose, lower side densely villose, very rarely glabrous, base acute, apex acuminate (acumen 15–20 mm long). *Inflorescence* narrowly oblong to ellipsoid, 9.5–20 by 4–5.5 cm, terminating a separate leafless shoot 50–110 cm long or rarely terminating a leafy shoot, sheaths obliquely truncate, 3–6.5 cm long, glabrous to densely puberulous; bracts, bracteole, calyx, ovary, and capsule densely to rather densely puberulous*. Flowers* abaxially orientated; bracts red to dark red, coriaceous, concave and bulging, broadly obovate to ovate, 3–3.5 by 2–3.5 cm, apex obtuse, sometimes acute, callus 5–9 mm long; bracteole boat-shaped, 23–30 mm long; calyx reddish, 11–17 mm long, lobes very shallowly triangular, 1–3 mm long; corolla yellow to orange, c. 50 mm long, densely to rather densely puberulous, lobes elliptic, 30–35 mm long; labellum yellow to orange, lateral lobes rolled inwards and forming a curved tube c. 10 mm diam, elliptic when spread out, c. 45 by 25 mm, 5-lobulate, lobules red, c. 5 mm long; stamen yellow to orange, 35–45 by c. 9 mm, not or slightly exceeding the labellum, apex red, acute, anther 8–10 mm long. *Capsul*e obovoid, 12–18 mm long.

Distribution — Peru (Madre de Dios, Puno), Bolivia (Beni, Cochabamba, La Paz).

Habitat & Ecology — On forested ridges and along roadsides, at elevations of 250-1800 m. Flowering all year round.

IUCN Conservation Status — Not evaluated. There are 14 collection records examined of which all 14 specimens were collected within the past 50 years. This species is also reported in 5 recent iNaturalist observations (https://www.inaturalist.org/observations?place_id=any&taxon_id=537963). The species is widely distributed in Bolivia and Peru. Our observations indicate that this is a common species within its range, including protected areas, with no known specific threats. Population and threat data is not available but based on the number of collection records and our general field observations, this species would probably score a status of Least Concern.

Vernacular names — Bolivia: Caña agria, Ucuno (Trinitario), Yushashta (Yuracare).

Notes — *Costus beckii* is characterized by an inflorescence usually terminating a separate leafless shoot, densely to rather densely villose sheaths, red bracts, a yellow and curved, tubular labellum with five red lobules, and a long ligule of 25–60 mm long.

*Seidel & Hirschle 2690* (U) from Bolivia, La Paz, Alto Beni, Concesión de Sapecho, 550 m, is aberrant from *C. beckii* by the glabrous lower leaf side.

**14. *Costus bracteatus*** Rowlee — Fig. 6e; Map 12

*Costus bracteatus* Rowlee (1922) 285, pl. 12. — Type: *Rowlee & Stork 675* (holo BH, 2 sheets), Costa Rica, Limón, 1 mile S of Siquirres, July 1920.

Herbs, 1–5 m tall. *Leaves*: sheaths 5–15 mm diam; ligule truncate to unequally 2-lobed, 3–15 mm long; petiole 5–15 mm long; sheaths, ligule and petiole densely brownish villose; lamina narrowly elliptic to narrowly obovate, 15–40 by 4–13 cm, generally 10–20-plicate, upper and lower side densely brownish villose, base acute, rounded, or cordate, apex long-acute to acuminate (acumen 10–15 mm long). *Inflorescence* generally nodding, cylindric, 5–19 by 3–9 cm, terminating a leafy shoot or terminating a separate leafless shoot 40–70 cm long, sheaths obliquely truncate, 5–8 cm long, densely puberulous to villose; bracts, appendages of bracts, bracteole, calyx, ovary, and capsule densely to sparsely puberulous to villose*. Flowers* abaxially orientated; bracts pale green, coriaceous, elliptic to ovate to broadly ovate, 2–3 by 1.5–3 cm, callus 3–7 mm long; appendages green, foliaceous, ascending, narrowly ovate-triangular to ovate-triangular, 0.5–3.5 by 0.5–1.5 cm, apex acute; bracteole boat-shaped, 15–28 mm long; calyx white to greenish white, 15–20 mm long, lobes triangular, 2–5 mm long; corolla pale yellow, 45–65 mm long, glabrous, lobes narrowly ovate to elliptic, 35–50 mm long; labellum yellow to cream, upper part horizontally spreading, obovate, 40–75 by 30–40 mm, lateral lobes striped with red, middle lobe recurved with a yellow honey mark, margin irregularly crenate; stamen pale yellow, 35–40 by 10–12 mm, not exceeding the labellum, apex reddish, irregularly 2–3-dentate, anther 8–10 mm long. *Capsule* narrowly obovoid, 10–20 mm long.

Distribution — Nicaragua, Costa Rica, Panama.

Habitat & Ecology — In primary forests, at elevations of 0–800 m. Flowering all year round.

IUCN Conservation Status — Least Concern.

Vernacular names — Panama: Binwe (Kuna), Sapar pinwe (Kuna).

Notes — *Costus bracteatus* has been incorrectly included by Maas (1972: 51) in the synonymy of *C. guanaiensis* var. *guanaiensis*, but in a later publication (Maas & Maas-van de Kamer 2003) it was accepted as a good species of its own. It can be distinguished by a combination of a villose indument of the vegetative parts, generally plicate leaves, a mostly nodding inflorescence, and bracts with green, narrowly ovate-triangular to ovate-triangular appendages.

**15. *Costus callosus*** Maas & H.Maas — Fig. 6f; Map13

*Costus callosus* Maas & H.Maas in Maas et al. (2023) 89, f. 7. — Type: *E. Forero et al. 6668* (holo COL, 3 sheets; iso MO [MO-1456278], U [U1607067, U1607068, U1607069, U1610613]), Colombia, Chocó, Mun. San José del Palmar, mouth of Río Torito (affluent of Río Hábita), Finca Los Guaduales, 730–830 m, 4 March 1980.

Herbs, 0.5–3 m tall. *Leaves*: sheaths 15–30 mm diam, green; ligule obliquely truncate, 10–30 mm long, upper part black to dark purple, with a minute, salient rim; petiole 5–20 mm long; sheaths, ligule and petiole glabrous, sparsely puberulous, or densely villose; lamina narrowly elliptic, 25–50 by 7–16 cm, upper side sparsely to rather densely villose to puberulous, lower side rather densely villose, base acute, apex acuminate (acumen 5–15 mm long). *Inflorescence* ovoid to subglobose, 6–23 by 6–12 cm, terminating a separate leafless shoot 20–90 cm long, rarely terminating a leafy shoot (*Knapp & Vodicka 5524*), sheaths withering and completely or partly falling off with age, obliquely truncate, 4–10 cm long, glabrous, sparsely puberulous to densely villose; bracts, bracteole, calyx, ovary, and capsule glabrous to rather densely puberulous*. Flowers* abaxially orientated; bracts orange-red, red, or yellow, coriaceous, ovate-triangular, 4–7 by 2.5–4 cm, apex acute to obtuse, more or less curved outwards to patent in living plants, callus 8–25 mm long, margins woolly in living material; bracteole boat-shaped, 20–30 mm long; calyx red, 10–18 mm long, lobes deltate, 2–8 mm long; corolla orange to yellow, c. 50 mm long, glabrous, lobes elliptic, c. 30 mm long; labellum yellow, lateral lobes rolled inwards and forming a tube c. 10 mm diam, elliptic when spread out, c. 35 by 25 mm, 3-lobulate, lobules red, 5–6 mm long; stamen yellow, 35–40 by 12–15 mm, slightly exceeding the labellum, apex orange-red, acute, anther 9–10 mm long. *Capsule* ellipsoid to obovoid, 8–15 mm long.

Distribution — Costa Rica, Panama, Colombia (Antioquia, Chocó).

Habitat & Ecology — In forests, at elevations of 0–1300 m. Flowering all year round.

IUCN Conservation Status – Not evaluated. There are 24 collection records examined of which 21 specimens were collected within the past 50 years. This species is also reported in 2 recent iNaturalist observations (https://www.inaturalist.org/observations?place_id=any&taxon_id=1460215). Based upon our observations of this recently described species it is rather rare, found in Colombia, Panama and Costa Rica. Population and threat data is not available but based on the number of collection records and our general field observations, this species might score as Near Threatened or Vulnerable.

Vernacular names — Panama: Caña agria (Kuna).

Notes — *Costus callosus* is most similar to *C. curvibracteatus,* differing in that the inflorescence of *C. callosus* is mostly produced on a separate leafless shoot with soon withering sheaths, and the inflorescence bracts have a distinct very long callus. In 1977, Maas had the suspicion that this *C. callosus* was different from *C. curvibracteatus* Maas, although it shares characters of ligule and bracts with it.

Material recently collected (*Maas et al. 10688*) in the Reserva Natural Nirvana, Valle del Cauca, Colombia, made it possible to add all flower details to the description (Maas et al. 2023).

**16. *Costus chartaceus*** Maas — Fig. 6g; Map 13

**Fig. 13.**
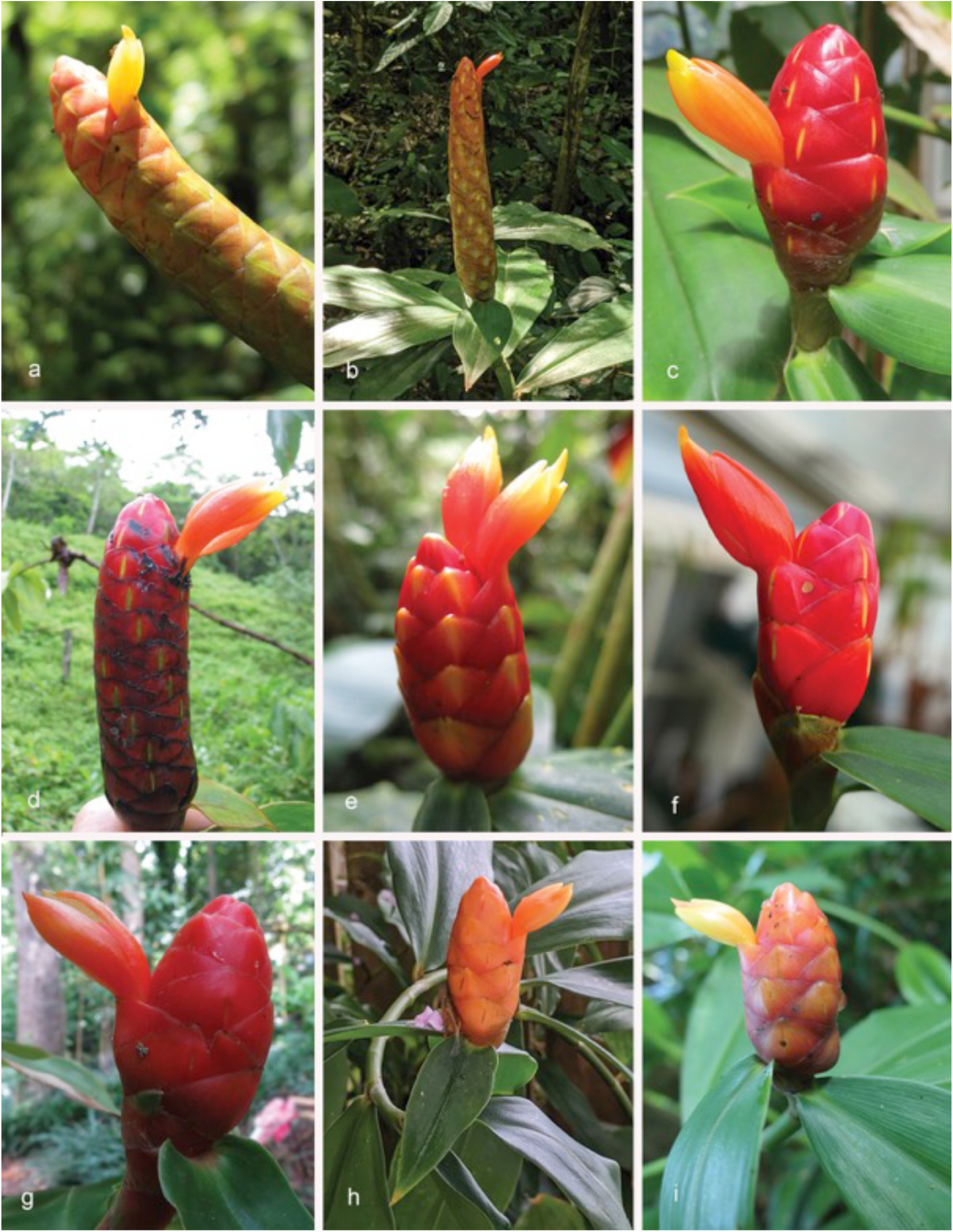
Some of the observed variation in the inflorescence of *Costus scaber* Ruiz & Pav.: a. not collected, photo in Oaxaca, Mexico; b. not collected, photo in Atlantca, Honduras; c. *D.Skinner R3076* from Ecuador, Zamora-Chinchipe; d. *D.Skinner R2971* from Costa Rica, Puntarenas, Cerro Nara; e. not collected, photo in Monteverde, Costa Rica; f. not collected, photo in Costa Rica, Osa Peninsula, Rio Barrigones; g. *D.Skinner R3220* from Guyana, Iwokrama; h. *D.Skinner R3386* from Bolivia, Santa Cruz, La Chonta; i. *D.Skinner R3254* from Brazil, Mato Grosso, Alta Floresta. Photos by D. Skinner.

*Costus chartaceus* Maas (1972) 98, f. 45. — Type: *Schultes 3540* (holo GH [GH00030653]), Colombia, Putumayo, Río San Miguel ó Sucumbios, Conejo y sus alrededores, en frente de la Quebrado Conejo, 2 Apr. 1942.

Herbs, 0.3–1(–3) m tall. *Leaves*: sheaths 5–10 mm diam; ligule unequally 2-lobed to truncate, 5– 15 mm long; petiole 3–10 mm long; sheaths, ligule and petiole subglabrous to densely brownish villose; lamina narrowly elliptic to narrowly obovate, 15–25 by 4–10 cm, upper and lower side glabrous to rather densely brownish villose, particularly the primary vein, base acute, rounded, or cordate, apex acuminate (acumen 5–15 mm long). *Inflorescence* broadly ovoid to subcylindric, 3.5–12 by 2.5–4 cm, terminating a leafy shoot; bracts, bracteole, calyx, ovary, and capsule glabrous*. Flowers* abaxially orientated; bracts reddish pink to dark pink, chartaceous, broadly ovate, 2–3 by 1.5–2.5 cm, apex obtuse, callus 2–7 mm long; bracteole boat-shaped, 13–18 mm long; calyx reddish pink, 7–11 mm long, lobes depressed ovate-triangular, 1–2 mm long; corolla pinkish white, 30–35 mm long, glabrous, lobes narrowly elliptic, 20–25 mm long; labellum pink, lateral lobes rolled inwards and forming a tube c. 4 mm diam, oblong-elliptic when spread out, 25–30 by 11–12 mm, 5-lobulate, lobules red, recurved, c. 1 mm long; stamen pink to pinkish white, 20–25 by 6–10 mm, slightly exceeding the labellum, apex obtuse, anther 6–10 mm long. *Capsul*e subglobose, c. 8 mm long.

Distribution — Colombia (Amazonas, Caquetá, Guaviare, Nariño, Putumayo, Valle del Cauca,Vaupés), Ecuador (Carchi, Napo, Orellana, Pastaza), Peru (Loreto).

Habitat & Ecology — In forests, at low elevations of 0–1100(–1570) m. Flowering all year round.

IUCN Conservation Status — Not evaluated. There are 86 collection records examined of which 72 specimens were collected within the past 50 years. This species is also reported in 4 recent iNaturalist observations (https://www.inaturalist.org/observations?place_id=any&taxon_id=828875). Population and threat data is not available but based on the number of collection records, this species would probably score a status of Least Concern.

Vernacular names — Colombia: Caña agria. Ecuador: Allpala-shangu (Quichua), Tentemokagi (Huaorani), Úntuntup (Achuar, Jivaro), Virucaspi.

Notes — *Costus chartaceus* closely resembles *C. prancei* and *C. sprucei*, species being characterized by chartaceous bracts; it can be distinguished from *C. sprucei* by pinkish white instead of yellow to orange flowers, and an unequally 2-lobed ligule of 5–15 mm long, versus a truncate ligule of 2–7 mm long in *C. prancei*.

Several Colombian collections from Amazonas (Villa Azul and Araracuara) and Caquetá (Sierra Chiribiquete) are aberrant by a somewhat longer ligule of 30–40 mm long. As all other features match *C. chartaceus* fairly well we have included them here.

**17. *Costus claviger*** Benoist — Fig. 6h; Map 13

*Costus claviger* Benoist (1927) 270. — Type: *Benoist 534* (holo P [P00686612]), French Guiana, Camp Charvein, 29 June 1914.

*Costus bracteatus* Gleason (1929) 20, nom. illeg., non Rowlee (1922). — *Costus guianicus* Loes. (1930) 632. — Type: *Jenman 6504* (holo BRG; iso K [K000586755]), Guyana, Mazaruni River, March 1892.

Herbs, 0.5– 2.5 m tall. *Leaves*: sheaths often reddish purple, 5–20 mm diam; ligule often reddish purple, obliquely truncate or slightly 2-lobed, (2–)5–15(–30) mm long; petiole 3–15 mm long; sheaths, ligule and petiole sparsely to densely puberulous to sericeous, rarely glabrous; lamina narrowly elliptic to narrowly obovate, 10–50 by 3–13 cm, lower side often reddish purple, rarely slightly plicate, upper side glabrous, rarely sparsely puberulous, lower side glabrous to densely puberulous, base acute to rounded, rarely cordate, apex acuminate (acumen 5–15 mm long) to acute. *Inflorescence* ovoid, (4–)6–16 by 3.5–7 cm, enlarging to c. 30 by 10 cm in fruit, terminating a separate leafless shoot 15–50 cm tall, sheaths often apically inflated, obliquely truncate, 3–5 cm long, sparsely to densely puberulous to sericeous, rarely villose or glabrous; bracts, bracteole, calyx, ovary, and capsule densely puberulous to sericeous, rarely glabrous*. Flowers* abaxially orientated; bracts dark red, coriaceous, broadly ovate, 3–6 by 2–5 cm, callus green, obsolete, up to 10 mm long; appendages green, foliaceous, horizontally spreading in living plants, usually reflexed when dry, broadly triangular-ovate, 0.5–2(–3.5) by 0.5–2(–3.5) cm, apex rounded; bracteole boat-shaped, (20–)25–30 mm long; calyx red, (10–)12–18 mm long, lobes deltate, 3–7 mm long; corolla yellowish white, 60–85 mm long, glabrous, lobes narrowly obovate, 50–70 mm long; labellum yellow to white, upper part horizontally spreading, broadly obovate, 50–60 by 50–60 mm, lateral lobes usually striped with dark red, middle lobe recurved with a yellow honey mark, margin irregularly crenate; stamen white, 40–50 by 7–13 mm, not exceeding the labellum, apex tinged with red and yellow, irregularly dentate, anther 10–12 mm long. *Capsule* ellipsoid, 12–20 mm long.

Distribution — Guyana, Suriname, French Guiana.

Habitat & Ecology — In rain forests, on sandy to clayey soils and often on granitic outcrops, at elevations of 0–700(–1250 m). Flowering in the rainy season.

IUCN Conservation Status — Not evaluated. There are 103 collection records examined of which 76 specimens were collected within the past 50 years. This species is also reported in 21 recent iNaturalist observations (https://www.inaturalist.org/observations?place_id=any&taxon_id=504271). The species is distributed throughout the Guianas. Population and threat data is not available but based on the number of collection records and our general field observations, this species would probably score a status of Least Concern.

Vernacular names – French Guiana: Ago chiga afou (Boni), Kapiyã (Wayãpi), Kapiya-pila (Wayãpi), Wilapa poa (Wayãpi), Yavelemoh (Oyampi). Guyana: Congo cane. Suriname: Ango sangrafoe, Apoekoe sangrafoe, Baku sãga afu (Saramaccan), Maikamin (Oyana), Sangrafoe.

Notes — *Costus claviger* is a quite common species in the 3 Guianas. It can be recognized by the inflorescence always produced on a separate leafless shoot, bracts provided with green appendages (with a rounded apex), and large flowers of which the labellum is yellow with reddish striped lateral lobes.

Material identified by Maas (1977) as *Costus* aff. *claviger* is nothing else but a form of *C. arabicus* from Bolivia (Beni) and Brazil (Acre, Rondônia) with appendaged bracts (see under that species).

**18. *Costus cochabambae*** Maas & H.Maas — Fig. 6i; Map 13

*Costus cochabambae* Maas & H.Maas in Maas et al. (2023) 92, f. 8 & 9. — Type: *Beck 7320* (holo LPB, specimen without barcode; iso L [L1480367, L1480368] and spirit collection: L [L0303088]), Bolivia, Cochabamba, Prov. Chapare, Villa Tunari, 40 kms hacia el Beni, por Chipiriri, 400 m, 24 Nov. 1981.

Herbs, c. 1.7 m tall. *Leaves*: sheaths 5–8 mm diam; ligule 15–25 mm long, unequally lobed, lobes obtuse; petiole c. 5 mm long; sheaths, ligule and petiole densely to rather densely puberulous to villose; lamina narrowly elliptic to narrowly obovate, 17–30 by 6–8 cm, upper side glabrous, lower side glabrous, but primary vein densely villose, strongly raised and prolonged as a prominent rim into the sheaths, base obtuse, apex acuminate (acumen 5–10 mm long). *Inflorescence* ovoid, 5–8 by 3–5 cm, terminating a leafy shoot; bracts, bracteole, and calyx densely to rather densely puberulous, ovary and capsule densely puberulo-sericeous*. Flowers* abaxially orientated; bracts green, coriaceous, broadly ovate, 1.5–2 by 1.5–2 cm; appendages green, foliaceous, reflexed when dry, broadly to shallowly ovate-triangular, 0.5–1.2 by 1.2–2 cm, apex obtuse to acute; bracteole boat-shaped, 15–17 mm long; calyx 9–12 mm long, lobes shallowly ovate-triangular, 2–3 mm long; corolla yellow, 30–40 mm long, subglabrous, lobes elliptic, 20–25 mm long; labellum yellow, lateral lobes rolled inwards and forming a tube c. 10 mm diam, oblong-elliptic when spread out, c. 20 by 15 mm, dentate; stamen yellow, c. 20 by 5 mm, not exceeding the labellum, apex dentate, anther 8–9 mm long. *Capsule* not seen.

Distribution — Bolivia (Cochabamba).

Habitat & Ecology — In high forests (“pie de la ladera con bosque alto”); at an elevation of c. 400 m. Flowering in November.

IUCN Conservation Status — Not evaluated. There is only 1 specimen, collected in 2012 that is documented for this species. There are no observations of this species on iNaturalist and the species has not been seen in the field by the authors of this manuscript. There is insufficient information to make a tentative assessment of IUCN status other than “Data Deficient”.

Note — *Costus cochabambae* superficially resembles *C. comosus*, but differs by greenish appendaged bracts, a subglabrous corolla, a glabrous rather than hirsute lower leaf side (except for the hairy primary vein), and a much longer ligule up to 25 mm long.

**19. *Costus comosus*** (Jacq.) Roscoe — Fig. 7a–f; Map 14

**Fig. 14.**
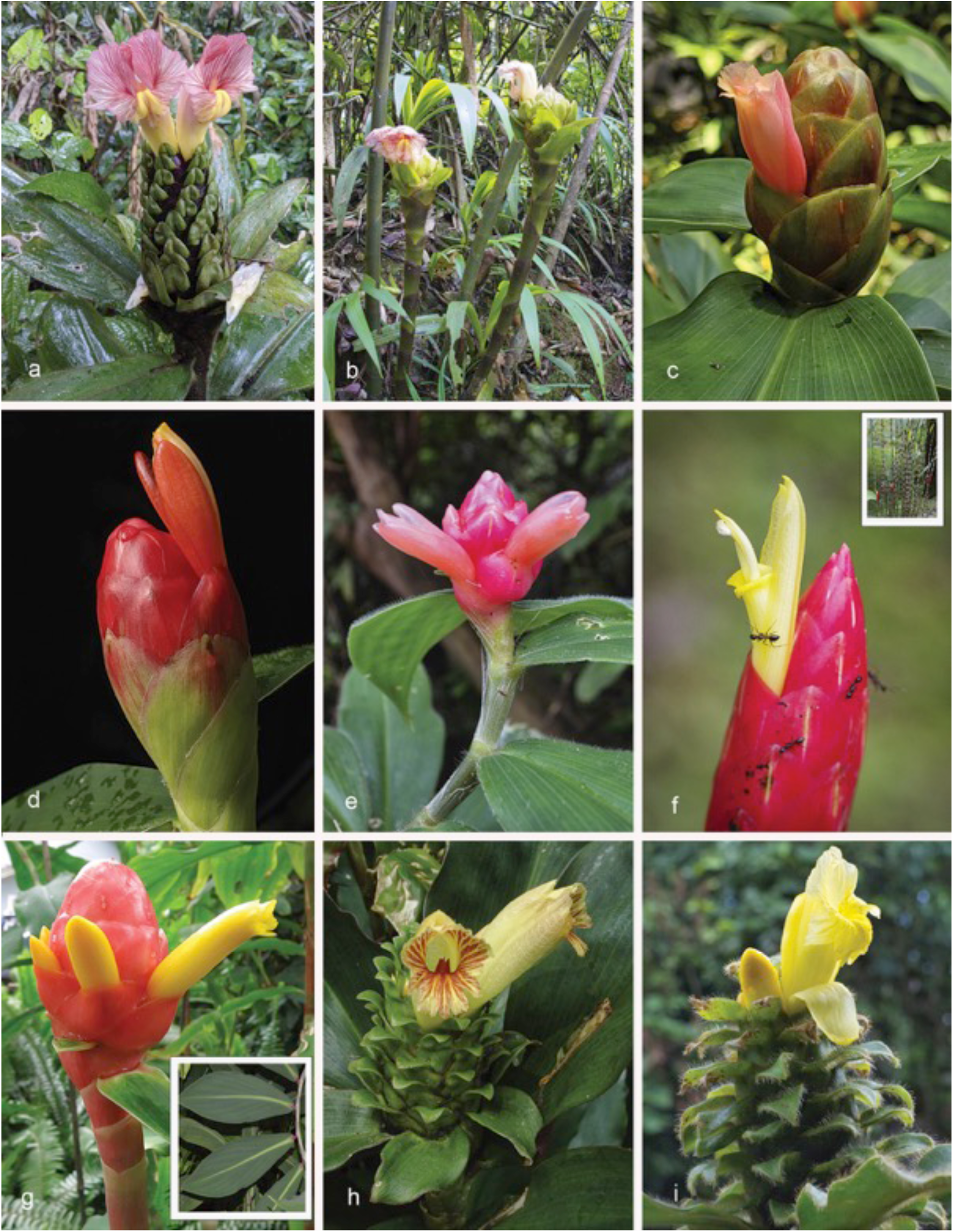
*Costus* species: *C. sinningiiflorus – C. villosissimus*: a, b. *C. sinningiiflorus* Rusby: a. common form (not collected, photo from Brazil, Acre near Cruzeiro do Sul); b. form with globular, basal inflorescence (*D.Skinner R3380* from Peru, Huanuco, Pumahuassi); c. *C. spicatus* (Jacq.) Sw. (*D. Skinner R3385*, Dominica); d. *C. spiralis* (Jacq.) Roscoe (*D. Skinner R3218*, Guyana, Iwokrama); e. *C. sprucei* Maas (*HSTM 13993 Coleta: D.B. Rodrigues* from Rurópolis, Pará, Brasil (Floresta Nacional do Tapajós)); f. *C. stenophyllus* Standl. & L.O.Williams (not collected, photo from La Gamba area, Puntarenas, Costa Rica (inset = habit showing stem coloration and characteristic narrow basal inflorescences)); g. *C. vargasii* Maas & H.Maas (*D.Skinner R3264* from Peru, Madre de Dios, Pantiacolla (inset = characteristic leaf and stem coloration –photo of plant in cultivation at Burgers Bush)); h. *C. varzearum* Maas (cultivated plant at Singapore Botanic Gardens); i. *C. villosissimus* Jacq. (not collected, photo from Costa Rica, Puriscal, La Cangrejá). Photos: a–d, f, g, i by D. Skinner; e by T. André; g inset by P.J.M. Maas; h by J. Leong-Škornicková.

*Costus comosus* (Jacq.) Roscoe (1807) 350. — *Alpinia comosa* Jacq. (1791) 112; (1793) 12, t. 202. — Lectotype (designated here): *Alpinia comosa* Jacq., ic. 2. tab. 202. collect. 4. p. 112. Hort. Vindob. (lecto BM [BM001000834]). Maas (1972) designated *Plate 202* from Jacquin (1793) as lectotype, originating from Venezuela, Distrito Federal, Caracas, and cultivated in the “Hortus Schoenbrunnensis” at Vienna because the original material at W was destroyed in 1943. Since then original material BM0010000834 was found and is here properly designated as the lectotype.

*Costus bakeri* K.Schum. (1904) 387. — *Costus comosus* (Jacq.) Roscoe var. *bakeri* (K.Schum.) Maas (1972) 85. —Lectotype (designated by Maaas 1972): *Donnell Smith 2802* (holo B, destroyed; lecto M [M0242450]; isolecto GH [GH00030645], K [K000586731], NY [NY00320322], U [U0007236], US [US00092956]), Guatemala, Amatitlán, Barranca de Eminencia, 1400 ft, Febr. 1892.

*Costus quasi-appendiculatus* Woodson ex Maas (1972) 85, f. 40. — Type: *Krukoff 10487* (holo U [U0007217]; iso BM [BM000923835], F [F0047181F], G [G00168416], GH [GH00030659], K [K000586766], LPB [LPB0000394], MO [MO-21132268], NY [NY00320349], S [S-R-1260]), Bolivia, La Paz, Basin of Río Bopi, near Calisaya, 750–900 m, 1 Jul. 1939.

*Costus maritimus* Standl. & L.O.Williams (1951) 233. — Lectotype (designated here): *P.H. Allen 5625* (lecto US [US00092967]; isolecto EAP [EAP103301], F [F0047167F, F0047168F], MO [MO-120587], US [US00338233]), Costa Rica, Puntarenas, delta of Río Esquinas, Golfo Dulce, 0 m, 28–29 Aug. 1951.

Herbs, 0.5–3 m tall. *Leaves*: sheaths 5–20 mm diam; ligule truncate, 1–3(–6) mm long; petiole 2–10 mm long; sheaths, ligule and petiole densely to rather densely puberulous to villose, rarely glabrous; lamina narrowly elliptic to narrowly obovate, 10–37 by 3–10 cm, upper side sparsely to rather densely puberulous to villose, sometimes slightly scabrid to the touch or glabrous, lower side densely puberulous to villose and soft to the touch, base acute, obtuse, or cordate, apex long-acuminate (acumen 10–30 mm long). *Inflorescence* ovoid to cylindric, 6–15 by 4.5–6 cm, terminating a leafy shoot, rarely terminating a separate leafless shoot 30–100 cm tall, sheaths obliquely truncate, 3–5 cm long, densely puberulous; bracts, appendages of bracts, bracteole and calyx rather densely puberulous to villose, sometimes glabrous, appendages of bracts sometimes slightly scabrid to the touch, ovary and capsule densely puberulous to villose*. Flowers* abaxially orientated; bracts dark red, pink, rarely pale green, coriaceous, broadly ovate-triangular, 2–3 by 2–3 cm; appendages red, pink, rarely pale green, foliaceous, horizontally spreading or ascending in the living plants, usually reflexed when dry, ovate-triangular to narrowly triangular, 1.5–4 by 0.5–2 cm (the lower ones to c. 8 cm long), apex acute, sometimes obtuse; bracteole boat-shaped, rarely 2-keeled, 14–20 mm long; calyx red to green, 9–18 mm long, lobes deltate to shallowly triangular, 2–6 mm long; corolla yellow, very rarely reddish, 30–45 mm long, densely puberulous to villose, sometimes glabrous, lobes narrowly elliptic, 20–30 mm long; labellum yellow, lateral lobes rolled inwards and forming a tube 8–10 mm diam, oblong-obovate when spread out, 20–30 by 8–20 mm, irregularly lobulate, lobules reflexed, deltate, 1–5 mm long; stamen yellow, 18–30 by 5–15 mm, slightly or not exceeding the labellum, apex rarely red, obtuse to acute, anther 5–10 mm long. *Capsule* ellipsoid, 8–12 mm long.

Distribution — Mexico (Chiapas), El Salvador, Guatemala, Panama, Costa Rica, Colombia (Antioquia, Bolívar, Boyacá, Caquetá, Casanare, Cesar, Cundinamarca, Magdalena, Meta, Santander), Venezuela (Aragua, Distrito Federal, Falcón, Lara, Miranda, Portuguesa, Zulia), Peru (Cuzco, Huánuco, Junín, Pasco), Bolivia (Cochabamba, La Paz, Santa Cruz).

Habitat & Ecology — In moist forests, montane forests, riverbank forests, especially in clearings, or on wooded slopes of volcanos, at elevations of 0–2200 m. Flowering period depending on variety: the former *Costus bakeri* flowers only in the dry season, other varieties flower in the rainy season.

IUCN Conservation Status — Not evaluated. There are 187 collection records examined of which 130 specimens were collected within the past 50 years. This species is also reported in 169 recent iNaturalist observations although its possible many are horticultural and the bakeri form is reported as a separate taxon. The species in its various forms is widely distributed from southern Mexico to southern Bolivia. Our observations indicate that this is a very common species with no known specific threats. Population and threat data is not available but based on the number of collection records and our general field observations, this species would almost certainly score a status of Least Concern as it is currently defined.

Vernacular names — Bolivia: Cañagria. Colombia: Caña agria, Cañagria, Caña agria pequeña. Guatemala: Caña Cristo, Caña Cristo del agua, Caña de Cristo, Caña de la Virgen. Mexico: Caña de la Virgen, Palo cristo. Peru: Sachahuiro. Horticultural: Red Tower Ginger.

Notes — *Costus comosus* is characterized by bracts mostly bearing reddish appendages, a tubular labellum, and an almost completely yellow, hairy corolla, and a very short ligule mostly not exceeding 3 mm in length. It could be confused with *C. lima*, a species which also has reddish appendaged bracts and hairy flowers, but that species has a much longer ligule (5–15 mm long) and the upper leaf surface and appendages of the bracts are mostly scabrid to the touch.

Although in the present treatment, the various forms of *Costus comosus* are here combined, there are indications of at least three separate linages based on geography (Bolivia/Peru, Panama, and Mexico), morphology, and molecular phylogenetic results (see Fig 7 a–f). Further research is needed before these differences can be fully resolved.

**20. *Costus convexus*** Maas & D.Skinner — Fig. 7g; Map 15

**Fig. 15.**
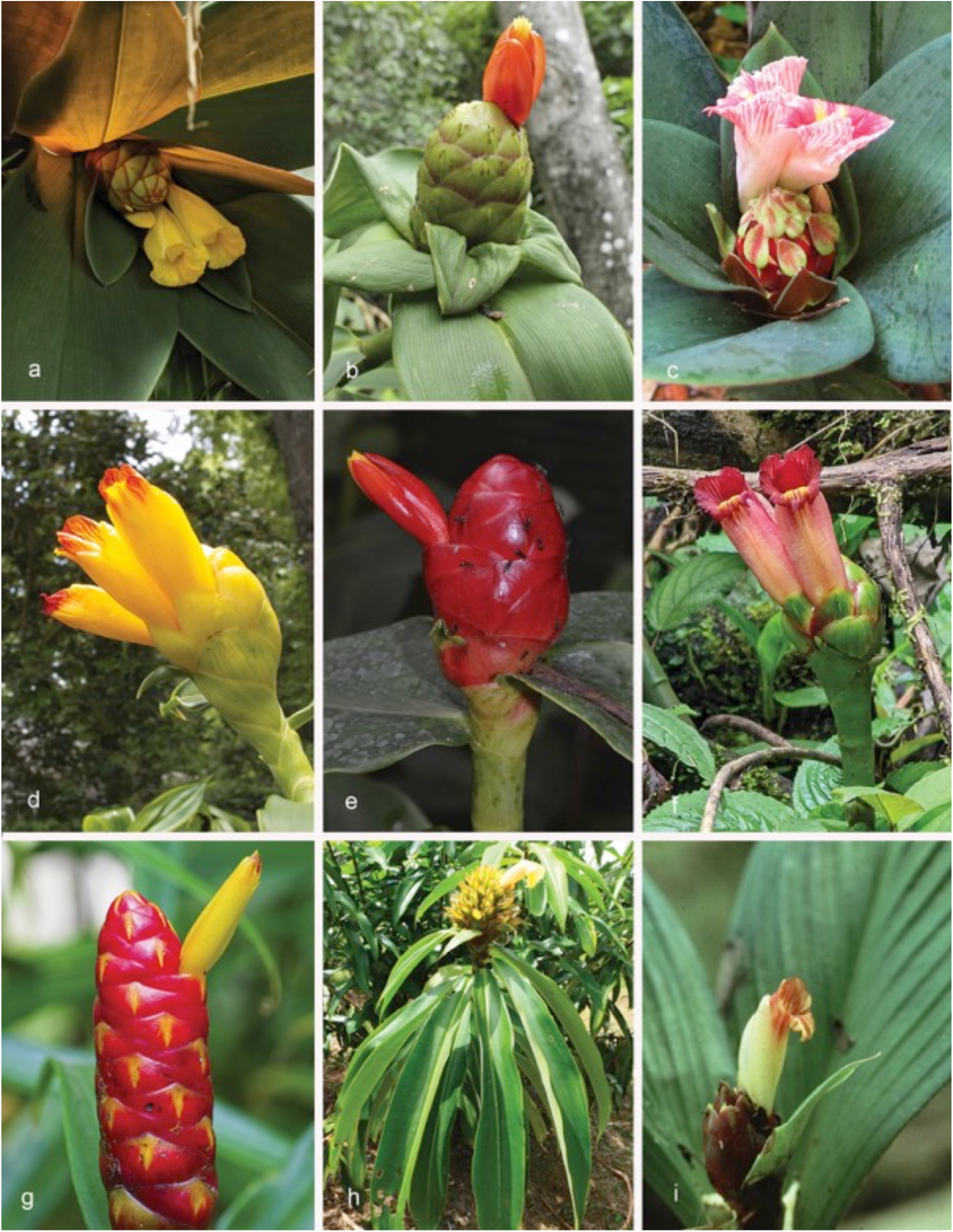
*Costus* species: *C. vinosus – C. sp. nov. “Gentry”*: a. *C. vinosus* Maas (*D.Skinner R3352* from Panama, Colón, Santa Rita ridge); b. *C. weberbaueri* Loes. (photo in habitat near La Merced, Junín, Peru); c. *C. whiskeycola* Maas & H.Maas (Collected by *Ray Baker* in Ecuador, Tena, cultivated at Lyon Arboretum L-82.1415); d. *C. wilsonii* Maas (*D.Skinner R3092* from Costa Rica, Puntarenas, La Amistad); e. *C. woodsonii* Maas (cultivated at Burgers’ Bush, Arnhem, The Netherlands); f. *C. zamoranus* Steyerm. (not collected, photo in Ecuador, Tena, Narupa region); g. *C. zingiberoides* J.F.Macbr. (*D.Skinner R3406* from Burgers’ Bush, Arnhem, The Netherlands); h. *C. sp. nov. “Chambi”* (*R3431* from Peru, Cusco, Quincemil); i. *C. sp. nov. “Gentry”* (*A.H.Gentry 24096* from Colombia, Chocó. Photos: a–d, f, g, h by D. Skinner; e by P.J.M. Maas; i by A.H. Gentry

*Costus convexus* Maas & D.Skinner in Maas et al. (2023) 92, f. 10. — Type: *Maas et al 10707* (holotype L, 2 sheets [L4468395, L4468396]), Colombia, Putumayo, 15 km S of Villagarzón on road to Puerto Umbria, 360 m, 17 July 2018.

Herbs, 3–5 m tall, young shoots and young leaves glaucous. *Leaves*: sheaths c. 20 mm diam; ligule truncate to obliquely truncate, 20–45 mm long; petiole 20–22 mm long, blushed in pink; sheaths, ligule and petiole glabrous; lamina narrowly elliptic, 31– 57 by 12–16 cm, upper side pale green, lower side green, or purple in very young plants, upper side glabrous, lower side glabrous to sparsely puberulous, base acute to slightly cordate, apex acuminate (acumen c. 15 mm long). *Inflorescence* globose to ovoid, 8–20 by 5–8 cm, wrapped tightly by the upper leaves, terminating a leafy shoot; bracts, bracteole, calyx, ovary, and capsule glabrous*. Flowers* adaxially orientated to erect; bracts distinctly convex, reddish on the outer margins, greenish at the center, coriaceous, ovate, 5.5–6.5 by 4–4.5 cm, apex obtuse, callus green, indistinct; bracteole boat-shaped, 40–45 mm long; calyx red, 15–25 mm long, lobes shallowly triangular to deltate, 4–10 mm long; corolla pink, 80–85 mm long, glabrous, lobes 50–60 mm long; labellum yellow, upper part horizontally spreading, broadly obovate, 60–70 by 60–70 mm, lateral lobes strongly striped with pink or red, middle lobe recurved with a yellow honey mark, margin irregularly dentate to lobulate; stamen dark pink to pinkish white, 30–40 by 12–15 mm, not exceeding the labellum, apex obtuse, anther 12–15 mm long. *Capsule* ellipsoid, 12–30 mm long.

Distribution — Colombia (Putumayo), Ecuador (Morona-Santiago, Zamora-Chinchipe).

Habitat & Ecology — In forests, in open shade or sunny areas, often found along roadsides and other disturbed areas, at elevations of 360–1500 m. Flowering all year round.

IUCN Conservation Status — Not evaluated. There are 8 collection records examined, all of which were collected within the past 50 years. This species is also reported in 11 recent iNaturalist observations (https://www.inaturalist.org/observations?place_id=any&taxon_id=1460220). The species is widely distributed in Colombia and Ecuador. Our observations indicate that this recently described species is very common within its range with no known specific threats. Population and threat data is not available but based on the number of collection records and our general field observations, this species would probably score a status of Least Concern.

Notes — *Costus convexus* can be recognized by its glaucous young parts, a globose inflorescence with the upper leaves wrapped tightly around it, and by distinctly convex bracts. Plants of *Costus convexus* can become huge, up to 5 m tall, with large inflorescences and flowers. The younger shoots and leaves usually have a glaucous covering making them appear similar to *C. glaucus* but having an inflorescence with distinctly convex bracts that are reddish or pink, instead of the pale green and glaucous bracts in *C. glaucus*.

This species has been observed in the field to be very common and nearly continuous along the eastern flanks of the mountains from Putumayo, Colombia, south through Ecuador to Zamora-Chinchipe. There are many more recent observations of this species recorded on inaturalist.com in Napo and Pastaza, Ecuador.

**21. *Costus cordatus*** Maas — Fig. 7h; Map 15

*Costus cordatus* Maas (1976) 472. — Type: *Maas & Plowman 1965* (holo U, 2 sheets U [U0007237, U0007238]; iso COL), Colombia, Valle del Cauca, El Boquerón, km 85 of new road Cali-Buenaventura, 150 m, 6 Oct. 1974.

Herbs, 0.75–2 m tall. *Leaves*: sheaths 5–10 mm diam; ligule 3–10(–20) mm long, unequally 2-lobed to truncate, lobes rounded; petiole 3–10(–15) mm long; sheaths, ligule and petiole glabrous to densely puberulous; lamina narrowly obovate to narrowly elliptic, (15–)25–40 by (5–)9–10(–13) cm, 10–12-plicate, sometimes slightly bullate, upper side glabrous, but primary vein covered with a row of erect hairs, lower side sparsely sericeous to glabrous, base cordate to obtuse, rarely acute, apex acuminate (acumen 5–20 mm long). *Inflorescence* cylindric, 4–11 by 2–5 cm, terminating a leafy shoot or sometimes terminating a separate leafless shoot 50–70 cm tall, sheaths obliquely truncate, 4–6 cm long, glabrous; bracts, bracteole, calyx, ovary, and capsule sparsely puberulous to glabrous*. Flowers* abaxially orientated; bracts red to orange, coriaceous, broadly ovate-triangular, 2–4 by 1.5–3 cm, apex obtuse, callus 4–9 mm long; bracteole boat-shaped, 14–23 mm long; calyx red, 9–13 mm long, lobes shallowly triangular, 3–4 mm long; corolla yellow, orange, or pink, 65–75 mm long, glabrous, lobes narrowly oblong-elliptic, 50–55 mm long; labellum yellow, orange, or pink, lateral lobes rarely striped with red, rolled inwards and forming a curved tube c. 12 mm diam, oblong-obovate when spread out, 40–45 by 25–27 mm, irregularly 7–9-1obulate; stamen yellow, 35–40 by 11–12 mm, not or slightly exceeding the labellum, apex obtuse, anther 10–11 mm long. *Capsule* ellipsoid, 10–15 mm long.

Distribution — Colombia (Cauca, Chocó, Nariño, Valle del Cauca) and Ecuador (Carchi, Esmeraldas, Los Ríos).

Habitat & Ecology — In non-inundated forests or secondary roadside vegetation, at elevations of 0–1200(–1800) m. Flowering all year round.

IUCN Conservation Status — Not evaluated. There are 35 collection records examined of which 30 specimens were collected within the past 50 years. This species is also reported in 4 recent iNaturalist observations (https://www.inaturalist.org/observations?place_id=any&taxon_id=846856). This region has been extensively deforested for coca growing and has had limited field observations in recent decades; so it is possible that there has been a substantial decrease in populations. Population and threat data is not available but based on the number of collection records and known destruction of habitat in Pacific coastal Colombia, this species is likely to be Vulnerable and it is quite possible that a full IUCN assessment would result in a category of Endangered for this species.

Vernacular names — Ecuador: Cana agria, Cañagrie, Guish (Coaiquer).

Notes — *Costus cordatus* can be distinguished by an inflorescence with red to orange, unappendaged bracts, plicate leaves with often a cordate base, and curved, yellow, orange, or pink, tubular flowers.

**22. *Costus cupreifolius*** Maas — Fig. 7i; Map 16

**Fig. 16.**
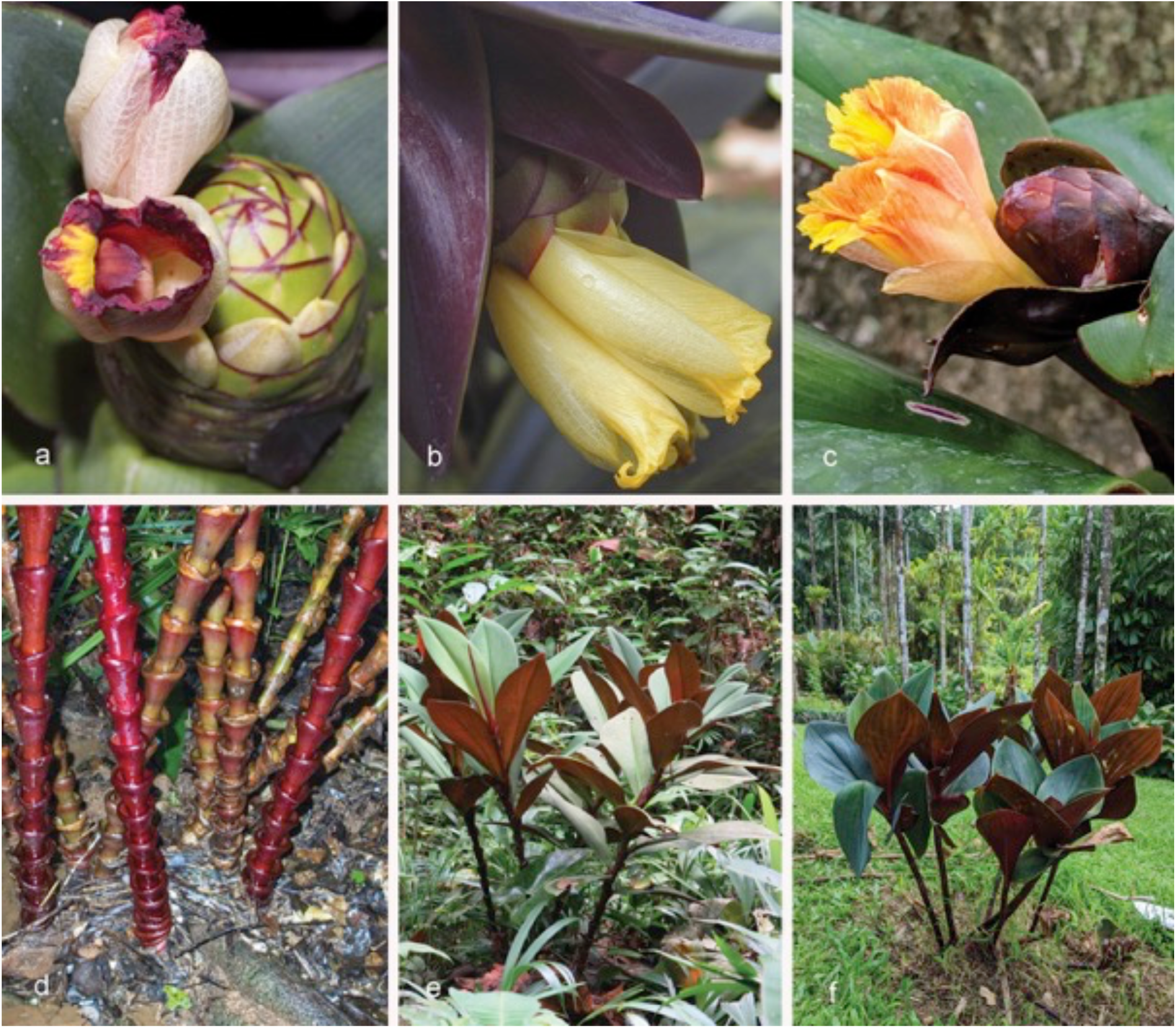
Variation in floral form and details on habit of *Costus vinosus* Maas in comparison with *C. whiskeycola* Maas & H.Maas: a. *C. vinosus* form 1 (*Dressler 4406* cultivated at Waimea Arboretum, from Panama, Colón, Rio Guanche); b. *C. vinosus* form 2 (*Dressler s.n.* cultivated at Marie Selby Gardens, from Panama, Colón, Rio Guanche); c. *C. vinosus* form 3 (*D. Skinner R3353* from Panama, Colón, Santa Rita Ridge), plant may represent a hybrid; d. cupped ligules (Santa Rita Ridge, Panama); e. in habitat at Santa Rita Ridge, Panama; f. *C. whiskeycola* (Ray Baker collection growing at Lyon Arboretum). Photos: a by J. Mood; b–f by D. Skinner.

*Costus cupreifolius* Maas (1976) 471. — Type: *Ewan 16748* (holo US [US00092958]; iso NY [NY00320333]), Colombia, Putumayo, near San Diego de Colorado, tributary of Río Putumayo, above Puerto Asis, 300 (“650”) m, 11 Jan. 1945.

Herbs, 1–2 m tall. *Leaves*: sheaths c. 10 mm diam; ligule obliquely truncate, 10–15 mm long; petiole c. 10 mm long; sheaths, ligule and petiole densely whitish villose; lamina narrowly elliptic to narrowly obovate, 25–30 by 8–11 cm, lower side red to reddish green, 4–7-plicate, upper side glabrous, but primary vein covered with a row of erect hairs, lower side densely villose, base acute to obtuse, apex acuminate (acumen 5–15 mm long). *Inflorescence* cylindric, 6–12 by 4–6 cm, terminating a leafy shoot; bracts, appendages of bracts, bracteole, calyx, ovary, and capsule densely whitish villose*. Flowers* abaxially orientated; bracts red, coriaceous, broadly ovate, 1.5–2 by 1.5–2 cm, callus indistinct, c. 1 mm long; appendages red, foliaceous, patent to reflexed, deltate, 0.7–2 by 0.7–2 cm, apex acute; bracteole boat-shaped, 13–20 mm long; calyx reddish, 10–20 mm long, lobes shallowly ovate-triangular, 2–5 mm long; corolla orange to yellow, 25–35 mm long, densely villose, lobes narrowly elliptic c. 25 mm long; labellum orange-red, lateral lobes rolled inwards and forming a tube c. 8 mm in diam, oblong-obovate when spread out, c. 25 by 20 mm, irregularly lobulate; stamen orange-red, c. 25 by 6 mm, slightly exceeding the labellum, apex obtuse, anther c. 5 mm long. *Capsule* ellipsoid, 5–20 mm long.

Distribution — Colombia (Putumayo).

Habitat & Ecology — In forests, at an elevation of 300–385 m. Flowering period is uncertain, but found in flower in December.

IUCN Conservation Status — Not evaluated. There are only 3 collection records examined of which all were in the province of Putumayo, Colombia, and there were only 2 collections within the last 50 years. This species is only reported in 2 recent iNaturalist observations both made by D.Skinner (https://www.inaturalist.org/observations?place_id=any&taxon_id=829195). The holotype site was visited by D.Skinner in 2018 and the only plants of this species found were in a small forest patch in the midst of areas that had been cleared for coca growing. This species would probably be assessed as Critically Endangered if formally assessed for IUCN status.

Notes – *Costus cupreifolius* is characterized by the often reddish (“coppery-red”) colour of the lower side of its plicate leaves, red-appendaged bracts, yellow to orange-red, tubular flowers, and a villose indument on many parts of the plant.

*Costus cupreifolius* is morphologically closest to *C. comosus* and *C. lima,* sharing the red-appendaged bracts and a hairy corolla, but it differs from both species by a villose indument on many parts of the plant and 4–7-plicate leaves, of which the lower side is often reddish (“coppery-red”).

This very rare species has been studied and photographed at its type locality at Vereda San Diego (the current name for San Diego de los Colorados), in the Municipality of Calceido, Putumayo, Colombia.

**23. *Costus curvibracteatus*** Maas — Fig. 8a; Map 16

*Costus curvibracteatus* Maas (1976) 473. — Type: *Maas & Cramer 1371* (holo U, 2 sheets U [U0007239, U0007240]; iso A [A00030647], COL [COL000000312], CR, E [E00319721], F [F0047164F], K [K000586735], MO [MO-120588]), Costa Rica, Cartago, Tapantí, 1200 m, 19 Aug. 1974.

*Costus spec. A*: Maas (1972) 124.

Herbs, 1–3 m tall. *Leaves*: sheath 10–20(–25) mm diam; ligule 5–10(–15) mm long, obliquely truncate to slightly 2-lobed, lobes rounded, upper part black to dark purple with a minute, salient rim; petiole 5–15 mm long; sheath, ligule and petiole densely villose to glabrous; lamina narrowly elliptic to narrowly obovate, (13–)20–35 by (4–)6–10(–12) cm, sometimes slightly plicate, upper side glabrous, but primary vein and margin often villose, lower side sparsely to densely villose, particularly the primary vein, base acute to rounded, sometimes cordate, apex acute to acuminate (acumen to c. 20 mm long). *Inflorescence* ovoid to broadly ovoid, 3–15 by 3– 5 cm, enlarging to c. 20 by 9 cm in fruit, terminating a leafy shoot; bracts, bracteole, calyx, ovary, and capsule glabrous to densely villose*. Flowers* abaxially orientated; bracts orange, red, or yellowish, apex sometimes green, coriaceous, ovate to ovate-triangular, (2.5–)4–5.5 by (1–)2– 3.5 cm, apex acute to obtuse, more or less curved outwards to patent in living plants, margin incurved, callus absent or inconspicuous; bracteole boat-shaped, 18–28 mm long; calyx dark red to reddish orange, 9–15 mm long, lobes shallowly triangular to deltate, 2.5–6 mm long; corolla yellow, rarely orange, 37–40 mm long, glabrous, lobes concave, narrowly elliptic, 27–30 mm long; labellum yellow tinged with orange, lateral lobes rolled inwards and forming a tube c. 10 mm diam, oblong-ovate when spread out, 25–40 by 20–22 mm, 5-lobulate, lobules linear to deltate, 3–6 mm long; stamen yellow, tinged with orange, 28–35 by 10–12 mm, slightly exceeding the labellum, apex red, obtuse, anther 8–10 mm long. *Capsule* ellipsoid, 10–20 mm long. round.

Distribution — Costa Rica, Panama.

Habitat & Ecology — In forests, at elevations of 600–1500(–2100) m. Flowering all year

IUCN Conservation Status — Least Concern.

Vernacular names — Panama: Caña agria.

Notes — In the field *C. curvibracteatus* can hardly be confused with any other species of *Costus* because of its slightly patent, orange, red, to yellow bracts. In that aspect it looks superficially similar to the *Renealmia cernua* (Sw. ex Roem. & Schult.) J.F.Macbr. of the *Zingiberaceae*. It has some characters in common with *C. barbatus*, but diverges from that species in the shape and colour of its bracts, smaller ligule, and a glabrous corolla. A characteristic feature of *C. curvibracteatus* is the upper part of the ligule which is black to dark purple and provided with a minute, salient rim. Morphologically it is closest to *C. callosus* (see under that species).

**24. *Costus dirzoi*** García-Mend. & G.Ibarra — Fig. 8b; Map 17

**Fig. 17.**
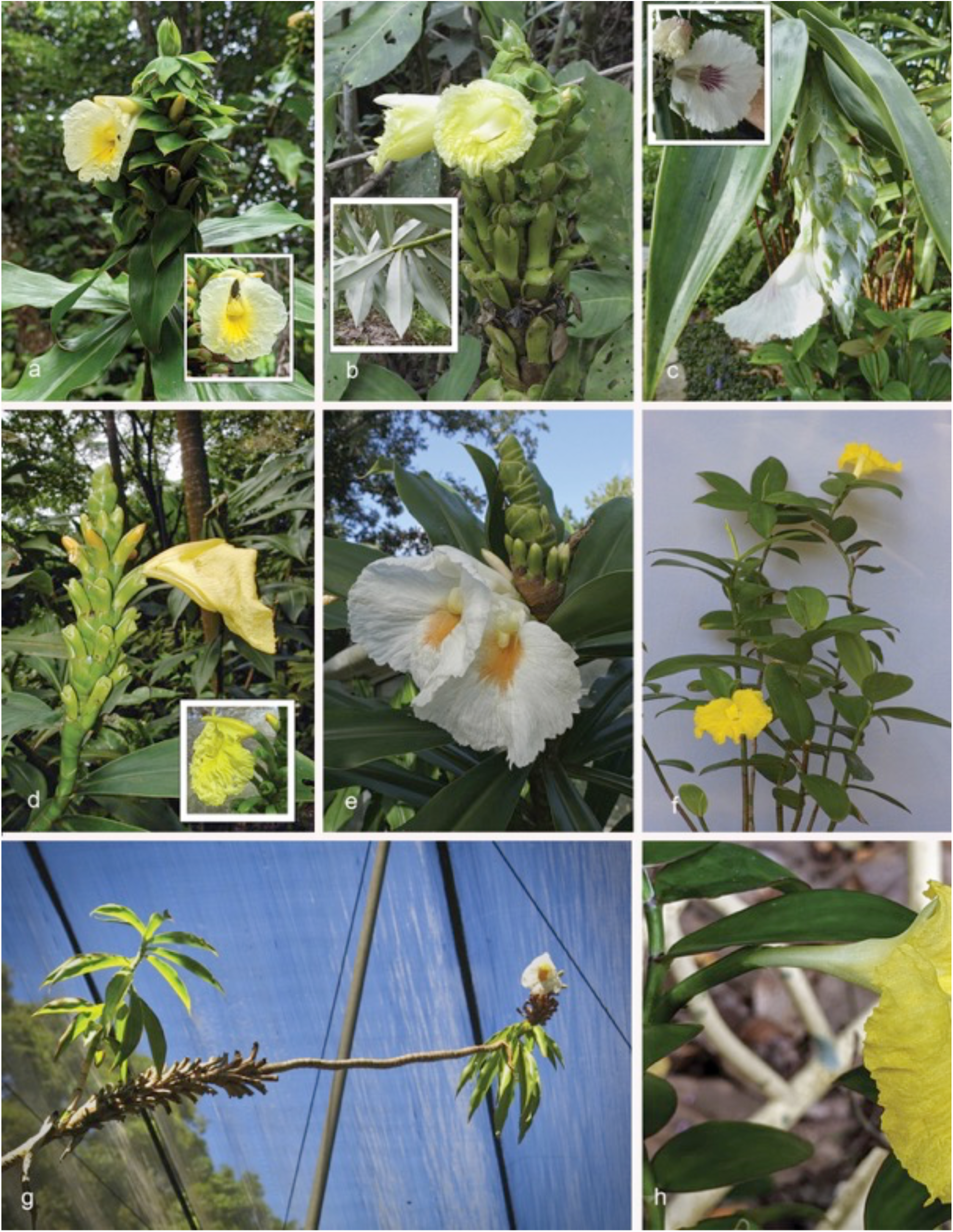
*Dimerocostus* and *Monocotus*: a. *D. appendiculatus* (Maas) Maas & H.Maas (not collected, photo from Peru, Loreto, Madre Selva (inset = closeup of flower)); b. *D. argenteus* (Ruiz & Pav.) Maas (*D.Skinner R3376* from Peru, Huanuco, La Gloria Road near type locality (inset = abaxial surface of leaves showing silvery indument)); c. *D. cryptocalyx* N.R.Salinas & Bentacur B. (not collected from Colombia (inset = closeup of flower showing labellum markings)); d. *D. rurrenabaqueanus* (Rusby) Maas & H.Maas (Singapore Botanic Gardens (inset = flower showing hooded structure from Peru, Madre de Dios, Manu Learning Centre)); e. *D. strobilaceus* Kuntze (*D.Skinner R3239* from Anchicaya, Colombia); f. *M. uniflorus* (Poepp. Ex Petersen) (*D.Skinner R3171* plant in cultivation, origin unknown); g. *D. strobilaceus* Kuntze (shoot continues demonstrating that inflorescence is not terminal, cultivated at Sitio Burle Marx, Brazil); h. *M. uniflorus* (Poepp. ex Petersen) showing axillary flower (*D.Skinner R3171* cultivated). Photos: a, b, d(inset), f–h by D. Skinner; c by C. Black; d by J. Leong-Škornicková

*Costus dirzoi* García-Mend. & G.Ibarra (1991) 1081, f. 1. — Type: *Ibarra Manríquez 3400* (holo MEXU [MEXU00798472]; iso BM [BM000923826, BM000923827], EAP [EAP103304, EAP103305], ENCB [ENCB003686], K [K000586736, K000586737], LE, MEXU [MEXU00798473], MO [MO-102944], US [US00386092], XAL [XAL423948]), Mexico, Veracruz, Mun. San Andrés Tuxtla, Estación de Biología Tropical Las Tuxtlas, 200 m, 7 June 1989. Herbs, 0.5–1.5(–2) m tall. *Leaves*: sheaths 4–8 mm diam, green; ligule obliquely truncate, 3–12 mm long; petiole 3–15 mm long; sheaths, ligule and petiole densely puberulous; lamina elliptic to narrowly elliptic, 20–27 by 8–15 cm, slightly c. 5-plicate, upper side glabrous, lower side densely velutinous, base acute, apex acuminate (acumen 20–30 mm long). *Inflorescence* ovoid to subglobose, 3–10 by 2.5–4 cm, enlarging to c. 16 by 5.5 cm in fruit, terminating a separate leafless shoot 20–40(–70) cm long, sometimes terminating a leafy shoot, sheaths obliquely truncate, 3–5 cm long, rather densely puberulous; bracts, bracteole and calyx densely puberulous, ovary and capsule densely villose*. Flowers* abaxially orientated; bracts green, coriaceous, broadly ovate, 2.5–4 by 2–3 cm, callus 5–8 mm long; bracteole boat-shaped, 15–20 mm long; calyx red, 5–9 mm long, lobes deltate, 2–4 mm long; corolla yellow, 50–60 mm long, glabrous, lobes narrowly obovate, 40–50 mm long; labellum yellow, upper part horizontally spreading, obovate, 50–65 by 30–35 mm, lateral lobes generally striped with dark red, middle lobe recurved with a yellow honey mark, irregularly 3–5-lobulate and dentate; stamen yellowish white, 33–45 by 10–12 mm, not exceeding the labellum, apex dark red, irregularly lobed, anther 8–10 mm long. *Capsule* broadly obovoid, 6–11 mm long.

Distribution — Mexico (Veracruz).

Habitat & Ecology — In primary, tropical rain forests, at elevations of 160–300(–700) m. Flowering only in the late dry season, May and June. Fruits and seeds are eaten and depredated (perhaps dispersed) by mice” (García-Mendoza & Ibarra Manríquez 1991)

IUCN Conservation Status — Not evaluated. There are 16 collection records examined of which 13 specimens were collected within the past 50 years. This species is also reported in 9 recent iNaturalist observations (https://www.inaturalist.org/observations?place_id=any&taxon_id=276564). The species is distributed only in Veracruz, Mexico, where it is a fairly common species. This region has been extensively deforested resulting in decreased habitat. Population and threat data is not available but based on the number of collection records and our general field observations, this species is likely to be Vulnerable based mainly upon its limited area of occupancy.

Vernacular names — Mexico: Bordón, Caña agria, Caña de venado.

Notes — *Costus dirzoi* can be distinguished by a velutinous indument of the lower side of the lamina, green, unappendaged bracts, and an inflorescence mostly on a separate leafless shoot. For the differences with *C. mollissimus* see under that species.

**25. *Costus douglasdalyi*** Maas & H. Maas — Fig. 8c; Map 16

*Costus douglasdalyi* Maas & H.Maas in Maas et al. (2023) 97, f. 11. — Type: *Daly et al. 8606* (holo U [U1225319], iso NY), Brazil, Acre, Mun. Tarauacá, Basin of Rio Juruá, Rio Tarauacá, Seringal Sumaré, 19 Nov. 1995.

Herbs, 1–4 m tall. *Leaves*: sheaths 5–20 mm diam; ligule truncate, 2–3 mm long; petiole 3–8 mm long; sheaths, ligule and petiole glabrous to sparsely puberulous; lamina narrowly elliptic to linear, 20–35 by 2–7 cm, lower side sometimes purple, upper side glabrous, lower side glabrous to sparsely puberulous, primary vein sparsely covered with a row of erect hairs, base acute to rounded, apex long-acute. *Inflorescence* ovoid to cylindric, 8–9 by c. 4 cm, enlarging to 13–17 by 5–7 cm in fruit, terminating a separate leafless shoot 45–70 cm long or rarely terminating a leafy shoot, sheaths obliquely truncate, 3–5 cm long, glabrous; bracts, bracteole, calyx, ovary, and capsule glabrous, apex of ovary and fruit sometimes sparsely puberulous*. Flowers* abaxially orientated; bracts red to dark red, coriaceous, broadly ovate, 2.5–4 by 2–3 cm, apex obtuse, callus 5–7 mm long; bracteole boat-shaped, 20–25 mm long; calyx red, 9–13 mm long, lobes shallowly triangular, 1–2 mm long; corolla red, pink, or pinkish orange, 45–55 mm long, glabrous, lobes (narrowly) ovate-elliptic, 35–45 mm long; labellum pink to red, lateral lobes rolled inwards and forming a curved tube c. 8 mm diam, broadly obovate to elliptic when spread out, 18–40 by 12–30 mm, irregularly 3–7-lobulate, lateral lobes striped with red, middle lobe yellow, 3–5 mm long, irregularly crenulate; stamen pale pink, 15–40 by 7–10 mm, not exceeding the labellum, apex acute, anther 8–10 mm long. *Capsule* ellipsoid, 10–15 mm long.

Distribution — Peru (Ucayali), Brazil (Acre, Rondônia).

Habitat & Ecology — In non-inundated (terra firme) forest, campinarana, river margins, or roadsides, at elevations of 0–350 m. Flowering all year round.

IUCN Conservation Status — Not evaluated. There are 17 collection records examined of which 14 specimens were collected within the past 50 years. This species is only known from western Brazil. Observations indicate that this is a fairly common species within its range and some of these areas are protected areas with no known specific threats. Population and threat data is not available, but based on the number of collection records and our general field observations, this species would probably score a status of Least Concern or possibly Near Threatened.

Notes — *Costus douglasdalyi* looks similar to *C. erythrothyrsus* in its floral characters, but differs by having very narrow, often linear leaves.

**26. *Costus erythrocoryne*** K.Schum. — Fig. 8d; Map 17

*Costus erythrocoryne* K.Schum. (1904) 410. — Lectotype (designated by Maas 1972): *Ule 6188* (holo B, destroyed; lecto MG [MG006059]), Peru, Loreto, Iquitos, July 1902.

Herbs, 1–6 m tall. *Leaves*: sheaths 10–20 mm diam; ligule obliquely truncate to slightly 2-lobed, 6–10 mm long, dark red for most of its length; petiole 5–12 mm long; sheaths, ligule and petiole densely to sparsely puberulous, rarely densely villose; lamina narrowly elliptic, 18–55 by 7–13 cm, upper side glabrous to sparsely puberulous, primary vein often sparsely covered with a row of erect hairs, lower side densely to rather densely whitish sericeous, rarely glabrous, base acute to obtuse, apex acuminate (acumen 10–20 mm long). *Inflorescence* ovoid to cylindric, 8–15 by 4–5 cm, enlarging to c. 30 by 8 cm in fruit, terminating a separate leafless shoot 30–90 cm long, sheaths obliquely truncate, 4–8 cm long, densely to rather densely puberulous, rarely densely villose; bracts, bracteole, calyx, ovary, and capsule sparsely to densely puberulous, margin of bracts, bracteole and calyx falling apart into fibers particularly when young*. Flowers* abaxially orientated; bracts red, coriaceous to chartaceous, broadly ovate, 3–5 by 3–5 cm, apex obtuse or acute, callus 7–15 mm long; bracteole boat-shaped, 25–30 mm long; calyx white to pink, 10–15 mm long, lobes shallowly triangular to deltate, 1– 2 mm long; corolla yellow, 60–65 mm long, densely puberulous in bud, becoming glabrous with age, lobes narrowly elliptic, 30–50 mm long; labellum yellow, lateral lobes rolled inwards and forming a slightly curved tube c. 10 mm diam, oblong-obovate when spread out, 35–40 by 20 mm, irregularly 5-lobulate, lobules red, 2–10 mm long; stamen yellow, 35–40 by 6–10 mm, slightly exceeding the labellum, apex red, acute, anther 8–10 mm long. *Capsule* ellipsoid, 18–20 mm long.

Distribution — Colombia (Amazonas), Ecuador (Napo, Orellana, Pastaza), Peru (Loreto).

Habitat & Ecology — In primary or secondary rain forests, sometimes in swampy places, at elevations of 0–500 m. Flowering all year round.

IUCN Conservation Status — Not evaluated. There are 57 collection records examined of which 47 specimens were collected within the past 50 years. This species is also reported in 9 recent iNaturalist observations (https://www.inaturalist.org/observations?place_id=any&taxon_id=505386). The species is distributed in Amazonian regions of Colombia, Ecuador and Peru. Our observations indicate that this is a fairly common species within its range and some of these areas are protected areas with no known specific threats. Population and threat data is not available but based on the number of collection records and our general field observations, this species would probably score a status of Least Concern.

Vernacular names — Colombia: Nebujae (Ticunas). Ecuador: Gonkemo.

Notes — *Costus erythrocoryne* is characterized by a separate leafless flowering shoot, red, unappendaged bracts, yellow flowers bearing a tubular labellum with red apical lobules, and a ligule 6–10 mm long. Another feature is the margin of the bracts, bracteole, and calyx falling apart into fibers, particularly when young.

**27. *Costus erythrophyllus*** Loes. — Fig. 8e; Map 18

*Costus erythrophyllus* Loes. (1929) 707; Maas (1977) 185, f. 71. —Type: *Tessmann 4813* (B, destroyed), Peru, Loreto, mouth of Río Apaga, Río Marañon Region, 145 m, Dec. 1924. Neotype (designated by Maas 1972): *Hortus Botanicus Utrecht 72-359* (neo U, 5 sheets [U0007829, U0007830, U0007831, U0007832, U0007833]), cultivated from seeds of *Prance et al. 12064a* (U [U1219636)], Brazil, Acre, Rio Moa, between Igarapé Ipiranga and Aquidabã, 4 July 1974.

Herbs, 0.5–2 m tall. *Leaves*: sheaths 8–20 mm diam; ligule (5–)10–40 mm long, unequally and very deeply 2-lobed, lobes narrowly ovate-elliptic to ovate-elliptic; petiole 5–15 mm long; sheaths, ligule and petiole glabrous, rarely rather densely to densely villose; lamina narrowly elliptic, some of the uppermost ones ovate-elliptic, (14–)20–32 by (5–)8–11 cm, often dark purple-red below, often 6–7-plicate, upper and lower side glabrous, rarely sparsely to densely villose, base acute, apex acute to acuminate (acumen to c. 10 mm long). *Inflorescence* ovoid, 4–8 by 3.5–6 cm, terminating a leafy shoot or terminating a short, separate shoot with few (2–7) well-developed leaves 7–27 by 4–9 cm just below the inflorescence, sheaths green to reddish, obliquely truncate, 3.5–11 cm long, rather densely to sparsely puberulous to villose; bracts and appendages of bracts rather densely to sparsely puberulous to villose, bracteole, calyx, ovary, and capsule sparsely to rather densely puberulous*. Flowers* abaxially orientated; bracts dark red, coriaceous, broadly ovate, 2–3.5 by 2–3.5 cm, callus 2–4 mm long; appendages green, foliaceous, ascending, triangular-ovate, 2.5–6 by 2–4 cm, apex acute; bracteole boat-shaped, 20– 27 mm long; calyx red, 11–18 mm long, lobes very shallowly triangular, 2–3 mm long; corolla white, 70–85 mm long, glabrous, lobes narrowly elliptic to narrowly obovate, 55–70 mm long; labellum white to salmon-coloured, upper part horizontally spreading, broadly obovate, 60–65 by 60–65 mm, lateral lobes striped with dark reddish pink, middle lobe recuved with a yellow honey mark, 3-lobulate, lobules 15–17 mm long, margin irregularly dentate; stamen pinkish white, 40– 50 by 12–16 mm, not exceeding the labellum, apex 3-dentate, anther 8–10 mm long. *Capsule* ellipsoid, 10–11 mm long.

Distribution — Colombia (Boyacá, Cundinamarca, Meta, Putumayo), Ecuador (Napo, Orellana, Pastaza, Sucumbios), Peru (Amazonas, Cuzco, Loreto, Madre de Dios, Pasco, Ucayali), Brazil (Acre).

Habitat & Ecology — In primary, non-inundated or periodically inundated (várzea) forests, at elevations of 0–900 m. Flowering throughout the rainy season.

IUCN Conservation Status — Not evaluated. There are 49 collection records examined of which 38 specimens were collected within the past 50 years. This species is also reported in 9 recent iNaturalist observations (https://www.inaturalist.org/observations?place_id=any&taxon_id=427386). The species is distributed in Amazonian regions of Colombia, Ecuador and Peru as well as one collection (the type collection) in Acre, Brazil. Our observations indicate that this is a common species within its range and some of these areas are protected areas with no known specific threats. Population and threat data is not available but based on the number of collection records and our general field observations, this species would probably score a status of Least Concern.

Vernacular names — Peru: Untuntu (Huambisa).

Notes — *Costus erythrophyllus* is characterized by often distinctly plicate leaves with a purplish lower side, an unequally 2-lobed ligule up to 40 mm long, and by an irregularly dentate labellum. Plants from Colombia and Ecuador that are included in this species have green lower sides of the leaves instead of purplish and the plication can appear less distinct.

**28. *Costus erythrothyrsus*** Loes. — Fig. 8f; Map 19

*Costus erythrothyrsus* Loes. (1929) 713. — Lectotype (designated by Maas 1972): *Tessmann 4262* (holo B, destroyed; lecto G [G00168447]), Peru, Loreto, western end of the Pongo de Manseriche near mouth of Río Santiago, Upper Río Marañon, 160 m, Oct. 1924.

Herbs, 1–3.5 m tall. *Leaves*: sheaths 8–13 mm diam; ligule slightly 2-lobed to obliquely truncate, 5–15(–20) mm long; petiole 5–15 mm long; sheaths, ligule and petiole sparsely puberulous to glabrous; lamina (narrowly) elliptic to (broadly) obovate, 8–35 by 5–15 cm, lower side sometimes purple, 3–8-plicate, upper side glabrous, but primary vein sparsely covered with a row of erect hairs, lower side sparsely puberulous to glabrous, base acute to obtuse, apex acuminate (acumen 5–15 mm long). *Inflorescence* (narrowly) ovoid to cylindric, 3–10 by 2–7 cm, enlarging to c. 20 by 8 cm, terminating a separate leafless shoot 15–100 cm long or rarely terminating a leafy shoot, sheaths obliquely truncate, 1.5–10 cm long, glabrous; bracts, bracteole, calyx, ovary, and capsule glabrous to densely puberulous*. Flowers* abaxially orientated; bracts carmine red, coriaceous, (very) broadly ovate, 2–4.5 by 2–4.5 cm, apex obtuse, callus 0–10 mm long; bracteole boat-shaped, 18–35 mm long; calyx pink to red, 8–13(–20) mm long, lobes shallowly triangular to deltate, 1–3 mm long; corolla red, reddish pink, or pinkish orange, 45–55 mm long, glabrous, lobes (narrowly) ovate-elliptic, 35–45 mm long; labellum pale pink, lateral lobes rolled inwards and forming a curved tube 7–8 mm diam, broadly obovate to elliptic when spread out, 30–40 by 15–30 mm, irregularly 3–7-lobulate, lateral lobes slightly striped with red, middle lobe yellow, 3–5 mm long, irregularly crenulate; stamen pale pink, 30–40 by 7–10 mm, not exceeding the labellum, apex red, acute, anther 7–10 mm long. *Capsule* ellipsoid to obovoid, 10–12 mm long.

Distribution —– Guyana, Suriname, French Guiana, Peru (Amazonas, Loreto, Ucayali), Brazil (Acre, Amazonas, Pará, Rondônia).

Habitat & Ecology — In forest, at elevations of 0–1000 m. Flowering all year round.

IUCN Conservation Status — Not evaluated. There are 140 collection records examined of which 120 specimens were collected within the past 50 years. This species is also reported in 26 recent iNaturalist observations (https://www.inaturalist.org/observations?place_id=any&taxon_id=870074). The species is distributed in Amazonian regions of Brazil and Peru as well as in the Guianas. Our observations indicate that this is a common species within its range and some of these areas are protected areas with no known specific threats. Population and threat data is not available but based on the number of collection records and our general field observations, this species would probably score a status of Least Concern.

Vernacular names — French Guiana: Canne congo (Créole), Kapiya-pila (Wayapi), Tuiu marikasmatgene (Palikur), Tuiwu axawukune (Palikur), Yanwilembo pilan. Guyana: Congo cane, Eseyundu (Carib), Red congo cane. Peru: Ashamat untugtú (Huambisa), Askamat ugtuntu (Huambisa), Kapantu utalu, Muuka chian (Huambisa), Uchiuntuntú (Huambisa), Untuntu (Huambisa). Suriname: Rode sangrafoe, Sangrafoe.

Notes — *Costus erythrothyrsus* is recognizable by the pink to red, tubular flowers mostly produced from a leafless shoot, combined with non-appendaged carmine red bracts. In that aspect it resembles *C. spiralis*, with which it was confused by Maas (1972), but in that species the flowers always are adaxially orientated and not abaxially as in *C. eythrothyrsus*.

**29. *Costus fissicalyx*** N.R.Salinas, Clavijo & Betancur — Fig. 8g; Map 20

*Costus fissicalyx* N.R.Salinas, Clavijo & Betancur (2007) 196, f. 1 & 2. — Type: *Clavijo-R. & Cordero-P. 415* (holo COL [COL000255697]; iso COAH [COAH78425]), Colombia, Vaupés, Taraira, Estación Biológica Mosiro Itajura (Caparú), 200 m, 20 Febr. 2004.

Herbs, 0.5–1.5 m tall. *Leaves*: sheaths 5–11 mm diam; ligule 2-lobed, 3–12 mm long; petiole 3– 10 mm long; sheaths, ligule and petiole glabrous to sparsely puberulous; lamina narrowly elliptic to elliptic, 15–30 by 7–11 cm, bullate, upper side glabrous, lower side glabrous to densely velutinous, base acute to rounded, apex acuminate (acumen 5–20 mm long). *Inflorescence* narrowly cylindric to fusiform, 4–14 by 2–3 cm, terminating a leafy shoot; bracts, bracteole, calyx, ovary, and capsule sparsely puberulous to glabrous*. Flowers* abaxially orientated to erect; bracts red, orange or yellow, broadly ovate, chartaceous, 2–3 by 1.5–2.5 cm, callus 5–8 mm long; bracteole boat-shaped, 12–20 mm long; calyx whitish, 10–14 mm long, unilaterally split, lobes indistinct, shallowly ovate-triangular, <1 mm long; corolla yellow, c. 60 mm long, glabrous, lobes narrowly elliptic, 32–38 mm long; labellum yellow, margin red, lateral lobes rolled inwards and forming an erect tube c. 5 mm diam, oblong-obovate when spread out, c. 35 by 13–15 mm, 5-lobulate, lobules acute to obtuse, 3–4 mm long; stamen yellow, c. 55 by 7 mm, far exceeding the labellum, apex obtuse, anther c. 6 mm long. *Capsule* not seen.

Distribution — Colombia (Amazonas, Vaupés).

Habitat & Ecology — In forests at elevations of 100–250 m, flowering year-round.

IUCN Conservation Status — Not evaluated. There are 7 collection records examined of which 5 specimens were collected within the past 50 years. There are no iNaturalist observations of this species. The species is distributed only in the western part of Colombia in Amazonas and Vaupés. The authors have not observed this species in the wild and it is not known how common it is, but it is possible that a full IUCN assessment would result in a status of Vulnerable or Endangered.

Vernacular names —Colombia: Jiyari-jiji (Yukuna).

Notes – *Costus fissicalyx*, endemic to Amazonian Colombia, is well marked by a very narrow inflorescence and yellow, tubular flowers of which the stamen far exceeds the labellum, and by slightly bullate leaves (a feature otherwise only seen in some specimens of *C. cordatus*).

**30. *Costus geothyrsus*** K.Schum. — Fig. 8h; Map 20

*Costus geothyrsus* K.Schum. (1904) 410. — Lectotype (designated by Maas 1972): *Von Eggers 14857* (holo B, destroyed; lecto K [K000586767]; isolecto PR), Ecuador, Manabi, El Recreo, 4 Aug. 1893.

Herbs, 1.5–4 m tall. *Leaves*: sheaths 15–30 mm diam; ligule truncate, 3–12 mm long; petiole 18– 25 mm long; sheaths, ligule and petiole glabrous to rather densely whitish puberulous; lamina narrowly elliptic to narrowly obovate, 36–57 by 13–20 cm, 7–8-plicate, upper side glabrous, lower side rather densely whitish puberulous, base acute to rounded, apex acuminate (acumen 10–20 mm long). *Inflorescence* cylindric to ovoid, 7–17 by 5–6 cm, enlarging to c. 32 by 9 cm in fruit, apex often slightly pointed, terminating a separate leafless shoot 75–100 cm long or sometimes terminating a leafy shoot, sheaths obliquely truncate, 2.5–5 cm long, rather densely to sparsely puberulous; covered part of bracts rather densely puberulous, exposed part subglabrous, bracteole, calyx, ovary, and capsule sparsely to densely puberulous*. Flowers* abaxially orientated to erect; bracts greenish red to wine-red, coriaceous, broadly ovate-triangular, 4–6 by 3–5 cm, apex acute or obtuse, margin entire or falling apart into fibers, callus 5–10 mm long; bracteole boat-shaped, 28–32 mm long; calyx reddish, 15–20 mm long, lobes deltate to shallowly triangular, 3–5 mm long; corolla reddish orange, c. 100 mm long, rather densely puberulous to glabrous, lobes narrowly obovate, c. 80 mm long; labellum yellow, lateral lobes rolled inwards and forming a tube 13–15 mm diam, obovate when spread out, 50–70 by c. 25 mm, 5–7-lobulate, lobules yellow, 8–10 mm long; stamen yellow, 55–65 by c. 10 mm, exceeding the labellum, apex obtuse, anther 10–11 mm long. *Capsule* obovoid, 15–20 mm long.

Distribution — Colombia (Nariño), Ecuador (Bolívar, Esmeraldas, Guayas, Imbabura, Los Ríos, Manabí).

Habitat & Ecology — In primary or secondary forests, and one collection from savanna, at elevations of 0–1000 m. Flowering in the rainy season.

IUCN Conservation Status — Critically endangered. Note that subsequent to the 2015 IUCN assessment, Skinner has found records of additional subpopulations of this species.

Notes — *Costus geothyrsus* looks somewhat similar to *C. pulverulentus* by its slightly pointed inflorescence; it is strongly aberrant, however, in having the inflorescence often produced on a separate leafless shoot and by much larger leaves and inflorescence. Moreover, the stamen in *C. pulverulentus* is far exceeding the corolla lobes whereas it is shorter than the corolla lobes in *C. geothyrsus*. Another remarkable feature is the length of the corolla lobes which are c. 100 mm long!

**31. *Costus gibbosus*** D.Skinner & Maas — Fig. 8i; Map 21

*Costus gibbosus* D.Skinner & Maas in Maas et al. (2023) 99, f. 12. — Type: *Plowman & Davis 4455* (holo U [U1212018]; iso GH, USM), Ecuador, Pichincha, along the road from Aloag to Santo Domingo, 1150 m (“3500 ft”), 17 Nov. 1974.

Herbs, 1.5–5 m tall. *Leaves*: sheaths 15–35 mm diam, the margin of the lower ones swollen; ligule truncate to slightly 2-lobed, often with a horizontal ridge near the apex, 7–14 mm long; petiole 10–15 mm long; sheaths, ligule and petiole glabrous to densely puberulous; lamina narrowly elliptic, 30–50 by 10–20 cm, slightly shiny above, pale green to slightly glaucous below, upper side glabrous, rarely puberulous, lower side glabrous, sometimes densely puberulous to villose, base acute to cordate, apex acuminate (acumen 15–20 mm long) to acute. *Inflorescence* ovoid to cylindric, 11–18 by 5–9 cm, terminating a leafy shoot; bracts, bracteole, and calyx glabrous, rarely rather densely puberulous, ovary and capsule glabrous to densely puberulo-sericeous*. Flowers* abaxially orientated; bracts red, coriaceous, broadly ovate, 4–5 by 3–4 cm; appendages green, foliaceous, erect, broadly triangular to triangular, 1.5–6 by 1–3 cm, often incurved at the apex; bracteole boat-shaped, 22–32 mm long; calyx red to pink, rarely white, (7–)12–20 mm long, lobes deltate to shallowly triangular, 2–5 mm long; corolla pale yellow, white, or pink, 60–70 mm long, glabrous, lobes ellliptic, 45–60 mm long; labellum pale yellow, orange, pinkish, or white, upper part horizontally spreading, broadly obovate, 60–85 by 50–70 mm, lateral lobes erect, more or less striped with pink or red, middle lobe recurved with a yellow honey mark, margin irregularly dentate to lobulate; stamen pink, red, yellow, or white, 40–50 by 12–15 mm, not exceeding the labellum, apex irregularly dentate, anther 9–13 mm long. *Capsule* ellipsoid to narrowly ellipsoid, 12–20 mm long.

Distribution — Ecuador (El Oro, Esmeraldas, Guayas, Loja, Manabi, Pichincha).

Habitat & Ecology — Forested roadsides, dry forest, or gallery forest, at elevations of 0– 1500 m. Flowering all year round.

IUCN Conservation Status – Not evaluated. There are 36 collection records examined of which 28 specimens were collected within the past 50 years. This species is also reported in 22 recent iNaturalist observations (https://www.inaturalist.org/observations?place_id=any&taxon_id=1460280). The species is distributed only in the western provinces of Ecuador. Population and threat data is not available but based on the number of collection records and our general field observations, this species would probably score a status of Least Concern.

Notes — *Costus gibbosus* can readily be recognized by the swollen margins of the lower leaf sheaths; the inflorescence is similar to that of *C. macrostrobilus* from which it differs by the horizontal orientation of the bract appendages with an often distinctly incurved apex and the mostly glabrous leaves, bracts, and calyx. For differences with *C. laevis* see under that species.

**32. *Costus glaucus*** Maas — Fig. 9a; Map 21

*Costus glaucus* Maas (1976) 470. — Type: *Maas 1535* (holo U, 2 sheets [U0060321, U0060322]; iso CR, F [F0047165F], US), Costa Rica, Cartago, forested hills of Río Reventazón Valley, Turrialba, near grounds of IICA, 600 m, 30 Aug. 1974.

Herbs, 1.5–5 m tall, young vegetative parts glaucous. *Leaves*: sheaths 6–20 mm diam; ligule 15– 55 mm long, unequally 2-lobed, lobes rounded; petiole 5–20 mm long; sheaths, ligule and petiole glabrous; lamina narrowly ovate to narrowly obovate, 20–43 by 5–14 cm, both sides glaucous when young, in older plants only lower side glaucous, upper and lower side glabrous, base acute to rounded, apex acute to acuminate (acumen 10–15 mm long). *Inflorescence* ovoid to cylindric, 5–16 by 3–8 cm, enlarging to c. 22 by 6 cm in fruit, terminating a leafy shoot; bracts scabrid to the touch, covered with 1-celled, appressed hairs 0.1–0.3 mm long, rarely glabrous, bracteole, calyx, ovary, and capsule sparsely to rather densely puberulous, rarely glabrous*. Flowers* adaxially orientated to erect; bracts glaucous to pale green with red margin, coriaceous, ovate to broadly ovate, 4.5–6 by 3–4.5 cm, apex obtuse, callus inconspicuous, just a vague line; bracteole boat-shaped, 22–27 mm long; calyx pink, 7–16 mm long, lobes very shallowly triangular to deltate, 1–6 mm long; corolla yellowish red, 65–70 mm long, glabrous, lobes narrowly elliptic, 50–55 mm long; labellum yellowish white, upper part horizontally spreading, broadly obovate, 60–70 by 50–55 mm, lateral lobes striped with pale to dark red, middle lobe recurved with a yellow honey mark, 3-lobulate, lobules c. 15 mm long; stamen pinkish to yellowish white, 40–50 by 13–15 mm, not exceeding the labellum, apex red, irregularly dentate, anther c. 13 mm long. *Capsule* ellipsoid, 15–25 mm long.

Distribution — Nicaragua, Costa Rica, Panama, Colombia (Antioquia, Chocó, Meta, Valle del Cauca).

Habitat & Ecology — In secondary and primary forests, at elevations of 0–800(–1700?) m. Flowering mostly in the rainy season.

IUCN Conservation Status — Near Threatened.

Notes — *Costus glaucus* is very easily recognizable by a glaucous cover on all young vegetative parts and bracts; other characteristics are the long and unequally 2-lobed ligule (15–55 mm long), the scabrid bracts, and the adaxially orientated flowers.

**33. *Costus guanaiensis*** Rusby — Fig. 9b, c; Map 21

*Costus guanaiensis* Rusby (1902) 694 (‘*guanaiense*’). — Lectotype (designated here): *Rusby 2225* (lecto NY, 2 sheets [NY00320334, NY00320335]; iso NDG [NDG11936], PH [PH00006740], US [US934292]), Bolivia, La Paz, Guanay, 2000 ft, May 1886.

*Costus longifolius* Rusby (1934) 50. — Lectotype (designated here): *Tate 442* (lecto NY [NY00320337]), Bolivia, La Paz, Mapiri, 10 Apr. 1926.

Herbs, 1.5–3 cm tall. *Leaves*: sheaths 10–25 mm diam; ligule 2–5 mm long, 2-lobed, lobes rounded; petiole 1–10 mm long; sheaths, ligule, and petiole densely to rather densely villose, rarely puberulous; lamina narrowly elliptic, 25–40 by 4–10 cm, upper side glabrous, but primary vein often rather densely covered with a row of erect hairs, lower side densely villose, rarely puberulous, base cordate, sometimes rounded, apex long-acute to acuminate (acumen 5–20 mm long). *Inflorescence* ovoid to cylindric, 6–25 by 3–5 cm, terminating a leafy shoot; bracts, appendages of bracts and bracteole glabrous, rarely sparsely puberulous, calyx often sparsely puberulous at the base, ovary and capsule densely sericeous*. Flowers* abaxially orientated; bracts red or green, coriaceous, broadly ovate, 2–4 by 2–3 cm, callus 5–13 mm long; appendages green, foliaceous, erect or recurved, triangular-ovate to broadly triangular-ovate, 0.5–3 by 0.5–1.5 cm, apex acute to obtuse; bracteole boat-shaped, 20–32 mm long; calyx red, 11–17 mm long, lobes deltate, 3–5 mm long; corolla red, pink, sometimes white, 60–80 mm long, glabrous, lobes narrowly elliptic, often patent, 45–60 mm long; labellum red to pink, upper part erect, obovate to quadrangular, 35–40 by 20–25 mm, without distinct lateral lobes, with a relatively narrow recurved margin all around and with 4–6 apical teeth, rarely the 2 central teeth yellow (honey mark); stamen red, 35–40 by 10–12 mm, slightly exceeding the labellum, apex entire or sometimes 3-dentate, anther 10–12 mm long. *Capsule* ellipsoid, 15–20 mm long.

Distribution — Peru (Cajamarca, Cuzco, Huánuco, Madre de Dios, Pasco, Puno, Ucayali), Bolivia (Beni, Cochabamba, La Paz, Santa Cruz), Brazil (Acre).

Habitat & Ecology — In moist, premontane or montane Yungas forests at elevations of 250–2700 m. Flowering in the rainy season.

IUCN Conservation Status — Not evaluated. There are 37 collection records examined of which 31 specimens were collected within the past 50 years. A proper observation count cannot be done on iNaturalist because it will include the observations made under the prior nomenclature before this species was reinterpreted. Under the interpretation proposed in this manuscript, this species is not very common, and its distribution is limited to Bolivia and adjacent parts of Peru and Brazil. Population and threat data is not available but based on the number of collection records and our general field observations, this species would probably score a status of Least Concern, but it is possible that a full assessment would score this species as Near Threatened.

Vernacular names— Peru: Caña caña, Caña-caña blanca.

Notes — *Costus guanaiensis* has long been a source of error as it had been, unfortunately, misinterpreted by Maas (1972). During a recent visit to the New York Herbarium (NY), Paul and Hiltje Maas and Chelsea Specht carefully reexamined the poor type collection of this species and came to a rather surprising conclusion. The specimen appeared to fit rather well in a species which they were about to describe. Therefore this “undescribed” Bolivian species is the real *C. guanaiensis*. The plants in the former *C. guanaiensis* have now been replaced by four different species names: *C. gibbosus, C. laevis* (also reinterpreted), *C. macrostrobilis, and C. sinningiiflorus*.

*Costus guanaiensis* is immediately recognized when in flower: they are red to pink, with an erect, not spreading labellum of which the margin is recurved. It shows other distinguishing characters like bracts which are provided with very small and green (sometimes red) appendages and also the mostly densely villose leaf sheaths and lower side of the (quite narrow) leaves.

We have studied the type and only collection of *C. longifolius*, collected in Bolivia not that far from the type of locality of *C. guanaiensis*. In our opinion it fits quite well in *C. guanaiensis*, but is aberrant by extremely narrowly leaves of c. 25 by 4 cm.

**34. *Costus juruanus*** K.Schum. — Fig. 9d, e; Map 22

*Costus juruanus* K.Schum. (1904) 389. — Lectotype (designated here): *Ule 5740* (holo B, destroyed; lecto HBG), Brazil, Acre, Rio Juruá Mirim, Sept. 1901.

*Costus productus* Gleason ex Maas (& var. *productus*) (1977) 193. — Type: *Killip & Smith 25317* (holo US [US00092973]; iso NY [NY00320346]), Peru, Junín, Perene Bridge, Río Paucartambo Valley, 700 m, 19 June 1929.

*?Costus juruanus* K.Schum. var. *strigosus* Maas (1972) 90, *syn. nov*. — *Costus productus* Gleason ex Maas var. *strigosus* (Maas) Maas (1976) 472. — Type: *Cuatrecasas 10684* (holo US [US00092961]; iso COL [COL000000314], F [F0047179F]), Colombia, Putumayo, Puerto Ospina, towards la Loma, 230–250 m, 19 Nov. 1940.

Herbs, 0.3–3 m tall. *Leaves*: sheaths 10–30 mm diam, sometimes inflated in young sprouts; ligule 5–60 mm long, obliquely truncate to unequally 2-lobed, lobes rounded, membranous and fragile when dry; petiole 2–20 mm long; sheaths, ligule and petiole sparsely to densely puberulous to villose or glabrous; lamina narrowly elliptic, narrowly obovate to obovate, 15– 30(–40) by 6–13(–20) cm, lower side often reddish purple, slightly 5–8-plicate, upper side glabrous, rarely sparsely to rather densely villose, primary vein sometimes covered with a row of erect hairs, lower side glabrous, rarely sparsely villose, base acute, obtuse, or cordate, apex acuminate (acumen 5–20 mm long). *Inflorescence* ovoid to cylindric, 3–14 by 3–7 cm, terminating a leafy shoot; bracts, appendages of bracts, bracteole and calyx glabrous, sometimes sparsely to rather densely puberulous to villose, ovary and capsule densely sericeous to villose*. Flowers* abaxially orientated to erect; bracts bright red, orange, or pale yellow, chartaceous to coriaceous, broadly ovate, 2.5–4 by 2–4 cm, callus 3–12 mm long, often inconspicuous; appendages of bracts often present, bright red to green, foliaceous, usually reflexed when dry, very broadly ovate, deltate, or shallowly triangular, 0.5–3.5 by 0.5–3 cm, apex acute to obtuse; bracteole boat-shaped, 15–25 mm long; calyx white to red, 9–19 mm long, lobes shallowly ovate-triangular to deltate, 2–7 mm long; corolla pale yellow, rarely orange, 30–50 mm long, glabrous, rarely sparsely sericeous, lobes narrowly obovate to narrowly elliptic, 25–35 mm long; labellum yellow, rarely orange, lateral lobes rolled inwards and forming a slightly curved tube 7– 9 mm diam, oblong-obovate to narrowly so when spread out, 25–35 by 8–25 mm, irregularly 5-lobulate, lobules red, 3–8 mm long; stamen yellow, rarely orange, 25–35 by 4–10 mm, not exceeding the labellum, apex red, irregularly dentate, anther 5–8 mm long. *Capsule* ellipsoid, 10–14 mm long.

Distribution — Colombia (Norte de Santander, Putumayo), Venezuela (Táchira), Guyana, Peru (Cuzco, Huánuco, Junín, Loreto, Madre de Dios, Pasco, Puno, San Martín, Ucayali), Brazil (Acre).

Habitat & Ecology — In forests, at elevations of 0–1700 m. Flowering all year round.

IUCN Conservation Status — Not evaluated. Collection records that previously were listed as *C. productus* are now listed as *C. juruanus* as that is now known to be the earliest name for the species. There are 72 collection records examined of which 56 specimens were collected within the past 50 years. This species is also reported in 12 recent iNaturalist observations under the previous name *C. productus*. This species is mostly found in Peru where it has been considered to be endemic, but there are a few disjunct records interpreted as this species in Venezuela and Guyana as well as in the type locality of Acre, Brazil. Population and threat data is not available but based on the number of collection records and our general field observations, this species would probably score a status of Least Concern.

Vernacular names — Brazil: Cana de macaco. Peru: Enano sachahuiro, Sachahuiro blanco, Sachahuiro colorado, Sachahuiro rojo, Sacha huiro colorado, Sacha huiro rojo.

Notes — *Costus juruanus* is easily recognizable by a very well developed, obtusely lobed, often membranous ligule to 60 mm long, a red to orange (or rarely yellow) inflorescence in which the small appendages of the bracts (if present) are often reflexed. The labellum is tubular and mostly yellow with red apical lobules. Variation of the ligule is noted especially in living specimens.

After having traced a duplicate of the type collection of *C. juruanus* (*Ule 5740* from HBG) we were quite sure about the identity of that species and included *C. productus* in its synonymy.

We have included here material from Colombia (Norte de Santander), adjacent Venezuela (Táchira), and Guyana. Initially we considered this material as a distinct species, but after having studied it in detail we found that the presence of a long, membranous ligule, the red-appendaged bracts and tubular flowers made it clear that this material should belong in *C. juruanus*. By doing this, the range of distribution of *C. juruanus* is logically extended.

**35. *Costus krukovii*** (Maas) Maas & H.Maas, *comb. et stat. nov.* — Fig. 9f; Map 23

*Costus amazonicus* (Loes.) J.F.Macbr. subsp. *krukovii* Maas, Fl. Neotrop. Monogr. 8 (1972) 71, f. 32 g–i. — Type: *Krukoff 7165* (holo S [S-R-1253]; iso BM [BM000923837, BM000923838], F [F0047174F], G [G00168448], GH [GH00030652], K [K000586754], LE [LE00001314], MICH [MICH1192017], MO [MO-202811], NY [NY00320331], U [U0007226], US [US00092955]), Brazil, Amazonas, Mun. Humaitá, on plateau between Rio Livramento and Rio Ipixuna, 11 Nov. 1934.

Herbs, 0.5–3.5 m tall. *Leaves*: sheaths 10–20 mm diam; ligule obliquely truncate, 5–15 mm long; petiole 3–10 mm long; sheaths, ligule and petiole densely to sparsely villose, rarely subglabrous; lamina narrowly obovate to narrowly ovate, sometimes ovate to elliptic, 25–40 by 7–15(–19) cm, lower side sometimes reddish purple, mostly 8–10-plicate, upper and lower side densely villose, rarely glabrous, base obtuse to cordate, apex acuminate (acumen 5–20 mm long), rarely acute.

*Inflorescence* ellipsoid to cylindric, 7–15 by 3–6 cm, enlarging to c. 17 by 6 cm in fruit, terminating a leafy shoot or mostly terminating a separate leafless shoot 10–60 cm long, sheaths obliquely truncate, often inflated, 1.5–8 cm long, glabrous, rarely densely villose; bracts, bracteole, and calyx glabrous, sometimes densely puberulous, ovary and capsule glabrous, except for the densely villose apex*. Flowers* abaxially orientated to erect; bracts green or greenish red, coriaceous, broadly ovate to broadly obovate, 3–4 by 2.5–4 cm, apex obtuse, sometimes acute, sometimes pungent and slightly recurved, callus 4–10 mm long; bracteole boat-shaped, 20–30 mm long; calyx red, 10–17 mm long, lobes shallowly triangular to deltate, 2–4 mm long; corolla white to yellow, 60–85 mm long, glabrous, lobes narrowly elliptic, 40–65 mm long; labellum white, upper part horizontally spreading, broadly obovate, 50–70 by 50 mm, lateral lobes striped with red-purple, middle lobe recurved with a yellow, red-margined honey mark, margin irregularly dentate; stamen white to yellow, 35–45 by 7–14 mm, not exceeding the labellum, apex pinkish white, 3-dentate, anther 7–13 mm long. *Capsule* ellipsoid, 10–15 mm long.

Distribution — Colombia (Amazonas, Caquetá, Guaviare, Putumayo, Vaupés), Ecuador (Napo, Orellana, Sucumbios), Peru (Amazonas, Cuzco, Loreto, Pasco, San Martín), Guyana, Brazil (Acre, Amazonas, Mato Grosso, Pará, Rondônia).

Habitat & Ecology — In non-inundated (terra firme) or periodically inundated (várzea) forests, at elevations of 0–500(–1150) m. Flowering all year round.

IUCN Conservation Status — Not evaluated. This species was formerly considered a subspecies of *Costus amazonicus*. There are 84 collection records examined of which 59 specimens were collected within the past 50 years. This species is also reported in 7 recent iNaturalist observations under its former status as a subspecies of *C. amazonicus*. The species is distributed in Colombia, Ecuador and Peru. Our observations indicate that this is a common species within its range and some of these areas are protected areas with no known specific threats. Population and threat data is not available but based on the number of collection records and our general field observations, this species would probably score a status of Least Concern.

Vernacular names — Brazil: Camafita. Colombia: Caña agria, Nebujae (Ticunas), Nebujae (Ticunas), Nebujé (Ticunas), Senje zé’nze, Tapaculo. Ecuador: Undundu (Shuar). Peru: Conaigre, Sachahuiro morado.

Notes — *Costus krukovii*, which has been previously included as a variety of *C. amazonicus*, can be recognized by greenish, non-appendaged bracts of which the apex is often pungent and slightly recurved, a villose indument on its vegetative parts, and the inflorescence mostly produced on a leafless shoot with often inflated sheaths. In the field this species is most easily identified by squeezing the sheath and finding a spongy feeling.

**36. *Costus kuntzei*** K.Schum. — Fig. 9g–i; Map 24

*Costus kuntzei* K.Schum. (1899) 422. — *Costus giganteus* Kuntze (1891) 687, nom. illeg., non Welw. ex Ridl. (1887) 131. — *Costus maximus* K.Schum. (1904) 405, nom. illeg., superfl. — Lectotype (designated here): *Kuntze 2140* (lecto NY [NY00320323]), Costa Rica, Limón, Puerto Limón, Baguar, 2500 ft, Jan. 1874.

*Costus splendens* Donn.Sm. & Türckh. (1902) 260. — Lectotype (designated here): *Von Türckheim 8015* (holo US, 3 sheets [US00092979, US00092980, US00092981]), Guatemala, Alta Verapaz, Cubilguïtz, 350 m, July 1901.

*Costus skutchii* C.V.Morton (1937) 306. — Type: *Skutch 2690* (holo US [US0092977]; iso GH [GH00030650], K [K000586730], MICH [MICH1192173], MO [MO-129051], NY [NY00320329], S [S-R-1261]), Costa Rica, San José, El General, 850 m, July 1936.

Herbs, 0.5–6 m tall, generally drying reddish brown. *Leaves*: sheaths 10–20(–40) mm diam; ligule obliquely truncate, 5–20(–35) mm long; petiole 5–20(–30) mm long; sheaths, ligule and petiole glabrous, rarely sparsely sericeous; lamina narrowly elliptic to narrowly obovate, 15–50 by 4–16 cm, young leaves and lower side of older leaves often purplish, upper side glabrous, rarely sparsely puberulous, primary vein often covered with a row of erect hairs, lower side glabrous, rarely densely puberulous, base acute, sometimes rounded or cordate, apex long-acuminate (acumen to c. 30 mm long). *Inflorescence* cylindric to ovoid, 5–10 by 3–7 cm, enlarging to c. 25 by 9 cm in fruit, often enclosed by some upper leaves, terminating a leafy shoot; bracts, bracteole, calyx, ovary, and capsule glabrous to rather densely puberulous*. Flowers* adaxially orientated; bracts green to yellowish green, occasionally dark red, coriaceous, broadly ovate, 2.5–5(–6.5) by 3–4.5 cm, apex obtuse, callus to c. 15 mm long; bracteole boat-shaped, 20– 30(–35) mm long; calyx red, 6–15(–20) mm long, lobes deltate to shallowly triangular, 2–5 mm long; corolla pale yellow to red, 50–75 mm long, glabrous, lobes narrowly obovate, 35–50 mm long; labellum basally white, upper part horizontally spreading, broadly obovate, 50–70 by 40– 55 mm, lateral lobes dark purple-red with white to yellow venation, middle lobe recurved with a yellow honey mark, 3–5-lobulate, margin crenulate, sometimes fimbriate; stamen red, 35–50 by 10–13 mm, not exceeding the labellum, apex red to purple, obtuse, anther 6–12 mm long. *Capsule* ellipsoid, 8–20 mm long.

Distribution — Guatemala, Belize, Honduras, Nicaragua, Costa Rica, Panama, Colombia (Antioquia, Caquetá, Cauca, Chocó, Cundinamarca, Magdalena, Nariño, Putumayo, Santander, Valle de Cauca), Ecuador (Bolívar, Carchi, Esmeraldas, Imbabura, Los Rios, Manabí, Napo, Pichincha.

Habitat & Ecology — In rain forests, often in clearings, in riverbank forests, or in swamps, at elevations of 0–2000 m. Flowering all year round.

IUCN Conservation Status — Not evaluated. This species was formerly considered a form of *Costus laevis* under its previous interpretation. There are 378 collection records examined of which 273 specimens were collected within the past 50 years. This species is also reported in many recent iNaturalist observations under its former classification as a form of *C. laevis*. The species is distributed in Central America and western provinces of Colombia and Ecuador. Our observations indicate that this is an extremely common species within its range and some of these areas are protected areas with no known specific threats. Population and threat data is not available but based on the number of collection records and our general field observations, this species would almost certainly score a status of Least Concern.

Vernacular names — Colombia: Caña agria, Caña agria blanca lisa, Cañagria, Cañagria de playa, Caña guate, Cañeja, Guira. Ecuador: Ajijimbre, Caña agria, Papatatih pil (Awápit), Sjivingola-tapé (Cayapa). Guatemala: Sangre de Criste. Peru: Sacha huiro.

Notes — *Costus kuntzei* can be distinguished by having adaxially orientated flowers with a large labellum with mostly red or dark-purple lateral lobes, and by having almost completely glabrous leaves. The colour of the exposed bracts is very dark red or mixed red/green where visible; the hidden parts of the bracts are always red.

Material of this species has thus far been wrongly identified as *C. laevis*, a species from Peru. Recent studies of the type collections of *C. laevis* made it clear that the real *C. laevis* is a very different species with appendaged instead of non-appendaged bracts. For further explanation see under *C. laevis*.

*Maas & McAlpin 1461* (U) from Costa Rica, Puntarenas, Esquinas Forest is aberrant in having a ligule which is densely villose at its apex.

**37. *Costus laevis*** Ruiz & Pav. — Fig. 10a; Map 25

*Costus laevis* Ruiz & Pav. (1798) 3. — Lectotype (designated here): *Ruiz López & Pavón s.n.* (lecto MA [MA810654]; isolecto BM [BM000923834]), Peru, Huánuco, near Pillao and Chacahuassi, 1787. For the discussion about the typification see notes.

*Costus tarmicus* Loes. (1929) 709. — *Costus guanaiensis* var. *tarmicus* (Loes.) Maas (1972) 56. — Lectotype (designated by Maas 1972): *Weberbauer 1856* (holo B, destroyed; lecto USM; isolecto MOL [MOL00008247]), Peru, Junín, Prov. Tarma, Chanchamayo valley, near La Merced, 700–1000 m, Dec. 1902.

*?Costus tarapotensis* J.F.Macbr. (1931) 50. — Type: *L. Williams 6529* (holo F [F0041534F]), Peru, San Martín, Tarapoto, Alto Río Huallaga, 12 Dec. 1929.

Herbs, 1.5–6 m tall. *Leaves*: sheaths 5–30 mm diam; ligule truncate to obliquely truncate, 3–12 mm long; petiole 5–15 mm long; sheaths, ligule and petiole glabrous, rarely sparsely to rather densely puberulous to villose; lamina narrowly elliptic to narrowly obovate, 15–50 by 4–16 cm, upper side glabrous, lower side glabrous, rarely sparsely to densely puberulous to villose, base cordate to rounded, rarely acute, apex acuminate (acumen 5–20 mm long). *Inflorescence* ovoid to cylindric, 6–20 by 4–12 cm, terminating a leafy shoot; bracts, appendages of bracts, bracteole, and calyx glabrous, rarely sparsely puberulous, ovary and capsule glabrous to densely puberulous*. Flowers* abaxially orientated; bracts red to green, coriaceous, broadly ovate to ovate, 3–5 by 1.5–4 cm, callus 3–10 mm long; appendages green, foliaceous, ascending, triangular to deltate, 0.8–2.5 by 0.5–1.5 cm; bracteole boat-shaped, 20–35 mm long; calyx red, 9–22 mm long, lobes triangular to shallowly triangular, 2–6 mm long; corolla white, pale yellow, or pinkish, 60–75 mm long, glabrous, lobes narrowly elliptic to narrowly obovate, 45–50 mm long; labellum white, pale yellow, or pinkish, upper part horizontally spreading, broadly obovate, 60– 70 by c. 50 mm, lateral lobes striped with pale to dark brownish red or violet, middle lobe recurved with a yellow honey mark, margin crenate; stamen white to yellow, 40–50 by 10–15 mm, not exceeding the labellum, apex red, dentate, anther 8–12 mm long. *Capsule* obovoid, 15– 20 mm long.

Distribution — Colombia (Antioquia, Caldas, Quindío, Valle del Cauca), Venezuela (Barinas), Ecuador (Bolívar, Cotopaxi, El Oro, Esmeraldas, Guayas, Loja, Los Rios, Manabí, Pichincha, Sucumbios), Peru (Amazonas, Cajamarca, Cuzco, Huánuco, Junín, Loreto, Madre de Dios, Pasco, San Martín, Ucayali).

Habitat & Ecology — In rain forests, often in clearings, riverbank forests, usually at middle elevations of 300–2000 m. Flowering mostly in the rainy season.

IUCN Conservation Status — Not evaluated. There are 76 collection records examined that are now included in this reinterpreted species of which 54 specimens were collected within the past 50 years. This species is also reported in 19 recent iNaturalist observations under its former name of *C. guanaiensis* var. *tarmicus*. Coauthor Skinner found this species to be very common in the departments of Junín and Pasco in Peru. Population and threat data is not available but based on the number of collection records and our general field observations, this species would probably score a status of Least Concern.

Vernacular names — Ecuador: Airo cte (Secoya). Peru: Caña agria.

Notes — The name *C. laevis* has, unfortunately, been wrongly applied by Maas (1972), because of a wrong interpretation of the type material of Ruiz and Pavón. This same error was first made by Robert Woodson (Woodson & Schery 1942). When Woodson looked at the specimen in Madrid with unappendaged bracts, he applied the name *Costus laevis* to the Central American plants we are now considering to be *Costus kuntzei*. Likewise in 1972, the *Costus laevis* that Maas had in mind was a specimen with **unappendaged** bracts.

Recently we studied the 4 syntypes of the original *Costus laevis* material of Ruiz & Pavón (“Habitat in Peruviae nemoribus ad vicum Pillao juxta Chacahuassi locis umbrosis humidisque”). Some essential characters from the original description by Ruiz & Pavón (1798; see Skinner 2016) are: “Culmi glabri…folia glabra…petioli ore purpurei…bracteae inferne rubrae, superne virides, **incurvae**…petala luteo-carnea…nectarium maximum, in centro rubrum, lineis carneo-lutescentibus striatum…” Upon revision of the syntypes, we came to the following results:

a. A collection in the Barcelona Herbarium (BA 872966): This specimen, collected in 1787 near Chacahuassi, does **not** have the appendaged bracts (“bracteae **incurvae**”) mentioned by Ruiz & Pavón. This specimen is quite similar to plants found near Chacahuassi by Dave Skinner in 2015 with green and **unappendaged** bracts and pinkish-red, tubular flowers.

b. A Ruiz & Pavón specimen in the BM Herbarium (marked “Costus laevis, Fl Per”) matches the original description of Ruiz & Pavón and clearly shows **appendaged**, recurved bracts. Unfortunately, this specimen has no exact location data.

c. A specimen in the Madrid Herbarium (MA810653) has **unappendaged** bracts. At the label it reads: “Costus laevis..Chacahuassi… 1787”.

d. A second specimen in the Madrid Herbarium (MA810654) has a well-developed inflorescence, and the bracts are **appendaged**. The label indicates: “Costus laevis Flor peruv… ex Herbario Flor. Peruv.”

Conclusion: The BA syntype and the MA syntype MA 810653 do not correspond with the original description of Ruiz & Pavón by having **unappendaged** bracts. They instead seem to represent a species later described as *Costus weberbaueri* Loes. The BM specimen and the MA specimen MA810654 both correspond to the original description of Ruiz & Pavón and are herewith selected for lectotypification of *C. laevis*. As the MA Herbarium houses many of the original collections of Ruiz & Pavón, we have selected MA810654 as the lectotype.

The conclusion of our studies is that the material previously identified as *Costus laevis* with **unappendaged** bracts should be given the second oldest name, *Costus kuntzei* K.Schum*. C. kuntze*i is a common species in Central America and the western part of tropical South America. The “real” *Costus laevis,* as typified by Ruiz & Pavón, occurs in western tropical South America and has - in contrast to *C. kuntzei* - **appendaged** bracts. It is consistent with C. *tarmicus*, which was described in 1929 by Loesener; thus *C. tarmicus* is presented as a synonym of *C. laevis*.

*Costus laevis* (the former *C. tarmicus*) occurs all over western South America, from Colombia to Peru. It is marked by bracts, which have weakly developed, green appendages. Another feature of this species is often the absence of any indument. As in all the three species of the former “*Costus guanaiensis*-group” (*C. laevis*, *C. macrostrobilus*, and *C. sinningiiflorus*), additional work is needed to understand the differentiation within this species complex.

Most material of *C. laevis* has completely glabrous leaves, but several specimens from Peru (Loreto, San Martín, and Ucayali) are aberrant by having a villose, white indument on the lower side of the lamina. In that aspect they resemble plants from the now-included *C. tarapotensis.* It is possible that *C. tarapotensis* is a species separate from the newly circumscribed *C. laevis*, in which case the material from Loreto, San Martín, and Ucayali, Peru should be considered for inclusion in *C. tarapotensis*. Future work will focus on understanding the relationship of these two taxa.

*Clark 3967* (MO, QCNE) and *Bass et al. 180* (MO, QCNE, U) collected at the Bilsa Biological Station, Quinindé, Esmeraldas, Ecuador, also have densely hairy leaves and the flowers look quite different (also smaller) from typical *C. laevis*. This material needs additional attention.

**38. *Costus lasius*** Loes. — Fig. 10b; Map 26

*Costus lasius* Loes. (1929) 710. — Neotype (designated by Maas 1972): *Allard 22060* (neo US [US00092962]), Peru, San Martín, Arroyo Bravo on road to Divisoria, 40 km from Tingo Maria on highway to Pucallpa, 1250 m, 1 Nov. 1950. A neotype had been selected by Maas (1972) as the holotype *Ule 6180* from Peru, Loreto, Leticia, June 1902, was destroyed in Berlin (B) in 1943.

*Costus nutans* auct. non K.Schum.: Rowlee (1922) 291; Woodson in Woodson & Schery (1945) 71.

Herbs, 0.5–2 m tall. *Leaves*: sheaths 3–8 mm diam; ligule truncate, 3–9 mm long; petiole 3–8 mm long; sheaths, ligule and petiole densely to sparsely brownish villose; lamina elliptic to obovate or mostly narrowly so, 6–28 by 2–6(–8.5) cm, upper side glabrous to rather densely brownish villose, lower side densely brownish villose, rarely glabrous, base acute to slightly rounded, apex acuminate (acumen 5–15 mm long). *Inflorescence* erect or nodding, ovoid to fusiform, 3–7(–10) by 2–3(–4) cm, terminating a leafy shoot; bracts subglabrous, the lower ones densely brownish villose, bracteole, calyx, and ovary glabrous to densely puberulous, capsule glabrous*. Flowers* abaxially orientated; bracts yellow to pale orange, rarely red, coriaceous (to chartaceous), broadly ovate, 2–3 by 2–3 cm, apex acute or obtuse, callus 3–5 mm long; bracteole boat-shaped, 8–12 mm long; calyx pale yellow, sometimes upper part red, 2–5 mm long, lobes very shallowly triangular, c. 1 mm long; corolla yellow, sometimes pale orange, 30–50 mm long, glabrous, lobes concave, obovate, 15–30 mm long; labellum yellow, lateral lobes rolled inwards and forming a tube 5–6 mm diam, oblong-elliptic when spread out, 17–25 by 10–15 mm, irregularly 3–7-lobulate, margin dentate; stamen yellow, 15–25 by 5–8 mm, slightly exceeding the labellum, apex irregularly dentate, anther 4–7 mm long. *Capsule* subglobose, 5–7 mm diam.

Distribution — Nicaragua (?), Costa Rica, Panama, Colombia (Amazonas, Antioquia, Caldas, Caquetá, Cauca, Chocó, Guajira, Meta, Putumayo, Quindío, Santander, Tolima, Vaupés), French Guiana, Ecuador (Carchi, Morona-Santiago, Napo, Pastaza, Sucumbíos), Peru (Cuzco, Huánuco, Loreto, Madre de Dios, Pasco, San Martín, Ucayali), Brazil (Acre, Amazonas, Pará, Paraíba, Rondônia).

Habitat & Ecology — In moist rain forests, foothill or mountain forests, at elevations of 0–1900 m. Flowering all year round.

IUCN Conservation Status – Least Concern.

Vernacular names — Colombia: Caña agria, Cañagüate, Kuendrumebema (Emberá), Rollinoginai (Witoto), Varela. Panama: Pinu puruiwat (Bayano Cuna). Peru: Sachahuire amarillo, Sacha huirito amarillo, Sachahuirito amarillo, Sacha huiro, Sachahuiro enano, Sacha huiro amarillo.

Notes — *Costus lasius* is one of the most distinctive species by its very slender shoots, a very small, generally yellow inflorescence, yellow flowers with concave petals, and a brownish villose indument of all vegetative parts.

**39. *Costus leucanthus*** Maas — Fig. 10c; Map 27

*Costus leucanthus* Maas (1976) 469. — Type: *Maas & Plowman 1920* (holo U, 3 sheets [U0007242, U0007243, U0007244]; iso COL [COL60631], F [F0047176F], GH, K [K000586762, K000586763], MO [MO-202809], US [US00092963, US00092964]), Colombia, Valle del Cauca, Limón, near km 71 of new road from Cali to Buenaventura, 150 m, 6 Oct. 1974.

Herbs, 2.5–4 m tall. *Leaves*: sheaths 20–30 mm diam; ligule 55–70 mm long, unequally and very deeply 2-lobed, lobes deltate to triangular, acute; petiole 10–15 mm long; sheaths, ligule and petiole densely to rather densely villose; lamina narrowly elliptic, 40–55 by 11–16 cm, upper and lower side densely to rather densely villose, base acute to rounded, apex acute to acuminate (acumen to c. 30 mm long). *Inflorescence* ovoid to cylindric, 10–17 by 9–10 cm, enlarging to c. 35 by 12 cm in fruit, terminating a leafy shoot; appendages of bracts densely to rather densely villose; bracts, bracteole, calyx, ovary, and capsule sparsely to rather densely puberulous*. Flowers* abaxially orientated to erect; bracts red, coriaceous, broadly ovate, 3–5 by 2.5–4 cm, callus 3–8 mm long; appendages green, foliaceous, the lower ones horizontally spreading, the upper ones vertically ascending, narrowly triangular to triangular, 4–8 by 1.5–3 cm, apex acute; bracteole boat-shaped, 28–31 mm long; calyx red, apex green, 16–20 mm long, lobes deltate to triangular, 6–7 mm long; corolla white, 80–90 mm long, glabrous, lobes narrowly elliptic, 60–70 mm long; labellum white, upper part horizontally spreading, broadly obovate, 60–70 by 45–65 mm, lateral lobes white, middle lobe recurved with a pale yellow honey mark, margin irregularly lobed and fimbriate; stamen white, 50–55 by 15 mm, not exceeding the labellum, apex irregularly dentate, anther 13–15 mm long. *Capsule* ellipsoid, 15–17 mm long.

Distribution — Colombia (Antioquia, Chocó, Nariño, Valle del Cauca), Ecuador (Esmeraldas, Imbabura).

Habitat & Ecology — Along forested roadsides, at elevations of 0–800 m. Flowering all year round.

IUCN Conservation Status — Not evaluated. There are 20 collection records examined of which 18 specimens were collected within the past 50 years. This species is also reported in 6 recent iNaturalist observations (https://www.inaturalist.org/observations?place_id=any&taxon_id=504887). The species is found only in southwest Colombia and northwest Ecuador. Our observations indicate that this is a somewhat uncommon species and much of this area has become deforested. Population and threat data is not available but based on the number of collection records and our general field observations, this species would probably score a status of Vulnerable or possibly Endangered.

Notes — *Costus leucanthus* is a species very well marked by large, and almost completely white flowers, a very long, acutely 2-lobed ligule (55–70 mm long), and a villose indument on most of its vegetative parts.

**40. *Costus lima*** K.Schum. — Fig. 10d; Map 28

*Costus lima* K.Schum. (1904) 388. — Type: *Von Scherzer s.n*. (holo B, destroyed, very small leaf fragment in F [F0047166F]), Costa Rica, Puntarenas, Jan. 1854; epitype (designated here, in concordance with Maas 1972): *Grijpma s.n.* (epi U [U0007209]), Costa Rica, Limón, Monteverde, c. 30 km W of Puerto Limón, in coastal plain, on clayey soil, 15 m, 28 Aug. 1968.

*Costus lima* K.Schum. var. *wedelianus* Woodson (1939) 277. — Type: *Woodson, Allen & Seibert 1926* (holo MO [MO-120590]; iso F, GH [GH00030655], NY [NY00320326]), Panama, Bocas del Toro, Río Cricamola, Finca St. Louis, between St. Louis and Konkintoë, 10–50 m, 12–16 Aug. 1938.

*Costus lima* K.Schum. var. *scabrimarginatus* Maas (1972) 81, f. 37a–g (‘*scabremarginatus*’). — Type: *Wessels Boer 2462* (holo U [U0007211]; iso A [A00030654], NY [NY00320336], US [US00092965]), Venezuela, Zulia, Perijá, near Misión de los Angeles del Tucuco, 900 m, 8 Apr. 1968.

Herbs, 2–6 m tall. *Leaves*: sheaths 10–20 mm diam; ligule obliquely truncate, 5–15 mm long, often with a purple margin; petiole 5–15 mm long; sheaths, ligule and petiole densely to sparsely puberulous to sericeous; lamina narrowly elliptic, 18–60 by 5–13 cm, upper side rather densely to sparsely villose, more or less scabrid by the thickened bases of the hairs, rarely glabrous, lower side densely villose and very soft to the touch, base acute to rounded, rarely cordate, apex long-acuminate (acumen 10–40 mm long). *Inflorescence* broadly ovoid to cylindric, 6–18 by 4–6 cm, enlarging to c. 30 by 8 cm in fruit, terminating a leafy shoot; bracts, bracteole, calyx, ovary, and capsule densely to sparsely puberulous to villose, appendages of bracts rather densely villose and scabrid to the touch, sometimes glabrous*. Flowers* abaxially orientated; bracts red to green, coriaceous, broadly ovate, 2–2.5 by 2–2.5 cm; appendages dark red, sometimes dark green, often with a yellow line in the middle, foliaceous, horizontally spreading to recurved in living plants, usually reflexed when dry, ovate to broadly ovate, 1.5–3 by 1–3.5 cm, apex acute to obtuse; bracteole boat-shaped, 15–23 mm long; calyx red, 10–15 mm long, lobes broadly ovate-triangular, 2–4 mm long; corolla deep pink, red, or yellow, basally white, 35–50 mm long, densely puberulous to sericeous, lobes narrowly obovate, 25–40 mm long; labellum basally whitish pink, upper part red, lateral lobes striped with red, rolled inwards and forming a slightly curved tube c. 10 mm in diam, narrowly oblong-obovate when spread out, 30–40 by 10–15 mm, 5-lobulate, lobules dark red, central lobules with yellow honey mark; stamen red to yellow, 30– 40 by 6–10 mm, not exceeding the labellum, apex red, irregularly dentate, anther 8–10 mm long. *Capsule* ellipsoid, 10–15 mm long.

Distribution — Honduras, Nicaragua, Costa Rica, Panama, Colombia (Antioquia, Cauca, Chocó, Magdalena, Meta, Nariño, Norte de Santander, Valle del Cauca),Venezuela (Zulia), Ecuador (Bolívar, Cañar, El Oro, Esmeraldas, Guayas, Imbabura, Los Rios, Morona-Santiago, Napo, Pichincha), Peru (Ayacucho, Cuzco, Junín).

Habitat & Ecology — In moist forests, wet thickets, swamps, or along streams, at elevations of 0–2000 m. Flowering all year round.

IUCN Conservation Status — Least Concern.

Vernacular names — Colombia: Cagaloduro, Cañagria, Canaquí, Hilo proprio, Pángala, Pangana. Ecuador: F/iba-ljuin-chi-tapé (Cayapa), Piní cola (Colorado). Nicaragua: Caña agria. Panama: Ginger Plant.

Notes — *Costus lima* is recognizable at first glance by the long-acuminate leaves, which are more or less scabrid to the touch at the upper side and densely villose and very soft to the touch at the lower side. It shares the red-appendaged bracts with *C. comosus* but differs from that species by the scabrid leaf indument and a much longer ligule (5–15 mm versus 1–3 mm long).

The many intermediates between var. *lima* and var. *scabrimarginatus* convinced us to unite both varieties.

**40. *Costus longibracteolatus*** Maas — Fig. 10e; Map 29

*Costus longibracteolatus* Maas (1972) 72, f. 33 (‘*longebracteolatus*’). — Type: *Asplund 19765* (holo S [S-R-1258]; iso U [U0007213]), Ecuador, Morona-Santiago, Macas, 900 m, 14 March 1956.

Herbs, 2–6 m tall. *Leaves*: sheaths 10–15 mm diam; ligule truncate, 10–30 mm long; petiole 5– 25 mm long; sheaths, ligule and petiole densely brownish villose; lamina narrowly elliptic to narrowly obovate, 20–48 by 6–18 cm, not or only very sightly plicate, upper side glabrous to densely brownish villose, lower side densely, sometimes sparsely brownish villose, base acute to obtuse, apex acuminate (acumen 5–20 mm long). *Inflorescence* ovoid, 7–25 by 5–10 cm, terminating a separate leafless shoot 50–80 cm long, sheaths obliquely truncate, 3–6 cm long, densely brownish villose; bracts, bracteole, calyx, and ovary glabrous to densely brownish villose, capsule glabrous*. Flowers* abaxially orientated to erect; bracts red, the central zone often green, coriaceous, broadly ovate-triangular to ovate-triangular, 3.5–7.5 by 3–5 cm, callus 5–20 mm long, sometimes inconspicuous, apex curved outwards in living plants and almost pungent, lower bracts often provided with a more or less distinct broadly ovate-triangular to deltate appendage 0.5–2 cm long, apex acute; bracteole boat-shaped, 32–42 mm long; calyx red to pink, 15–22 mm long, lobes triangular to deltate, 5–10 mm long; corolla yellowish white, reddish tinged, 70–100 mm long, glabrous, lobes narrowly elliptic, 50–70 mm long; labellum cream at the base, upper part horizontally spreading, broadly obovate, 60–90 by 50–70 mm, lateral lobes with wine-red venation or completely red, middle lobe recurved with a yellow honey mark, margin crenulate; stamen white to pale yellow, 35–50 by 12–14 mm, not exceeding the labellum, apex red to pink, irregularly dentate, anther 10–12 mm long. *Capsule* ellipsoid, 10–25 mm long.

Distribution — Colombia (Amazonas, Caquetá, Putumayo), Guyana, Ecuador (Morona-Santiago, Napo, Orellana, Pastaza, Sucumbios), Peru (Loreto, Pasco), Brazil (Amazonas).

Habitat & Ecology — In primary or secondary, non-inundated or periodically inundated (tahuampa or várzea) forests, at elevations of 0–500(–1000) m. Flowering all year round. In Peru (Loreto), the seeds are eaten by a bird (“palomita”; *Balick et al. 991*, U).

IUCN Conservation Status — Not evaluated. There are 77 collection records examined of which 65 specimens were collected within the past 50 years. This species is also reported in 26 recent iNaturalist observations (https://www.inaturalist.org/observations?place_id=any&taxon_id=505462). The species is distributed in amazonian regions of Brazil, Colombia, Ecuador and Peru. Our observations indicate that this is a common species within its range and some of these areas are protected areas with no known specific threats. Population and threat data is not available but based on the number of collection records and our general field observations, this species would probably score a status of Least Concern.

Vernacular names — Colombia: Caña. Ecuador: Sacha chiguilla (Kichwa). Peru: Caña agria, Kiwácyo (Bora).

Notes — *Costus longibracteolatus* is very well characterized by brownish villose shoots, a very long bracteole and calyx, bracts almost pungent at the apex, and an extremely long callus, up to c. 20 mm long, combined with having an inflorescence produced on a separate leafless shoot. It looks quite similar to *C. krukovii*, the main difference being the pungent bracts with a more pronounced callus and the indument of the leaf sheaths, which is generally more dense than found in *C. krukovii*.

**41. *Costus lucanusianus*** J.Braun & K.Schum. — Map 29

*Costus lucanusianus* J.Braun & K.Schum. (1889) 151. — Type: *Braun s.n.* (B, destroyed in 1943), Cameroon, South Province, Batanga. Neotype (designated by Maas-van de Kamer & Maas in Maas-van de Kamer et al. 2016): *Van Andel et al. 3406* (neo WAG [WAG0145874]; isoneo KRIBI), Cameroon, South Province, Campo-Ma’an National Park, Ntem River, Ebianemeyong, at the foot of Asuagale falls, 350 m, 6 May 2001.

*Costus dussii* K.Schum. (1904) 402, f. 45B. — Lectotype (designated by Maas 1972): *Duss 2109 b* (holo B, destroyed; lecto NY [NY00073335]), Martinique, ‘Hauteurs du Carbet. Fonds Saint Denis’, anno 1880.

Herbs, 1–2 m tall. *Leaves*: sheaths 5–15 mm diam; ligule truncate to slightly 2-lobed, 1–2 mm long, with a basal horizontal rim 1–2 mm high provided with a prominent row of needle-like hairs 2–6 mm long; petiole 4–10 mm long; sheaths, ligule and petiole glabrous; lamina narrowly elliptic, 12–30 by 5–7 cm, upper side glabrous, lower side densely silvery sericeo-puberulous, base obtuse to cordate, apex acuminate (acumen 10–20 mm long). *Inflorescence* broadly ovoid to globose, 4–12 by 4–9 cm, sometimes elongating to c. 20 cm in fruit, terminating a leafy shoot; bracts, bracteole, calyx, ovary, and capsule glabrous*. Flowers* abaxially orientated; bracts green, coriaceous, broadly to very broadly ovate-triangular, 1.5–3 by 1.5–3 cm, apex obtuse, callus 1–3 mm long; bracteole boat-shaped, 15–20 mm long; calyx green, 15–20 mm long, lobes broadly ovate-triangular to triangular, 5–7 mm long, horizontally spreading to reflexed, in fruit distinctly exceeding the bracts; corolla white, 30–45 mm long, glabrous, lobes elliptic, 25–30 mm long; labellum white, upper part horizontally spreading, broadly obovate, 40–50 by 40–45 mm, lateral lobes striped with dark red, middle lobe recurved with a yellow to orange honey mark, margin slightly crenate; stamen white, 30–35 by 10–15 mm, not exceeding the labellum, apex dark pink, irregulary 3-dentate, anther 6–10 mm long*. Capsule* globose, 14–15 mm diam.

Distribution — Cultivated and possibly naturalized in the island of Martinique. An African species, very common throughout Northeast, West, Central, East and Southern Africa.

Habitat & Ecology — This species is native to forests and forest edges of tropical Africa.

Notes — In contrast to all neotropical species of *Costus*, which have a single flower per bract, inflorescences of *C. lucanusianus* bear two flowers in the axil of each bract as do several other African species. As this species is native to Africa and is only found cultivated in the neotropics, it is not included in the key. More information about this species can be found in a monograph of African Costaceae (Maas-van de Kamer et al. 2016).

**43. *Costus macrostrobilus*** K.Schum. — Fig. 10f; Map 30

*Costus macrostrobilus* K.Schum. in Urban (1903) 159. — *Costus guanaiensis* Rusby var. *macrostrobilus* (K.Schum.) Maas (1972) 52, f. 25 & 26. —Lectotype (designated by Maas 1972): *Von Eggers 1289* (lecto US [US00092966]; isolecto C), Puerto Rico, El Sobrante, Sierra de Luquillo, 1800 ft, May 1883.

*Costus argenteus* auct. non Ruiz & Pav.: Gagnep. (1902) 157; K.Schum. (1904) 389.

*Costus villosissimus* auct. non Jacq.: Rowlee (1922) 287, in part.

*Costus scaber* auct. non Ruiz & Pav.: Loes. (1931) 89.

*Costus friedrichsenii* auct. non Petersen: Woodson in Woodson & Schery (1942) 329.

Herbs, 1–5 m tall. *Leaves*: sheaths 5–20(–40) mm diam; ligule truncate to obliquely truncate, 5– 10 mm long; petiole 5–20 mm long; sheaths, ligule and petiole densely to sparsely villose, rarely sparsely puberulous or glabrous; lamina narrowly elliptic to narrowly obovate, 25–70 by 5–15 cm, upper side rather densely to sparsely villose, hairs whitish, scabrid to the touch, rarely glabrous, lower side densely villose, rarely sparsely so to glabrous, base acute, rarely rounded to cordate, apex acuminate (acumen 10–15 mm long). *Inflorescence* ovoid to cylindric, 8–20 by 5– 12 cm, enlarging to c. 35 by 15 cm in fruit, erect, terminating a leafy shoot; bracts, appendages of bracts, bracteole, and calyx sparsely puberulous to glabrous, ovary and capsule densely puberulous, rarely glabrous*. Flowers* abaxially orientated; bracts red, coriaceous, broadly ovate to ovate, 2–6 by 2–4(–6) cm; appendages green, foliaceous, ascending, triangular, 0.5–4 by 0.3–2 cm; bracteole boat-shaped, 25–40 mm long; calyx red, 9–18(–22) mm long, lobes triangular to shallowly triangular, 1–8 mm long; corolla white to yellowish or pinkish white, 70–100 mm long, glabrous, sometimes rather densely puberulous, lobes narrowly elliptic, 50–80 mm long; labellum white, upper part horizontally spreading, broadly obovate to obovate, 60–110 by 50–80 mm, lateral lobes striped with red, middle lobe recurved with a yellow honey mark, margin irregularly crenate; stamen white, 45–60 by 10–20 mm, not exceeding the labellum, apex red, irregularly dentate, anther 8–12 mm long. *Capsule* obovoid, 15–25 mm long.

Distribution — Mexico (Chiapas), Greater Antilles (Puerto Rico), Guatemala, Belize, Nicaragua, Costa Rica, Panama, Colombia (Antioquia, Atlantico, Bolívar, Caquetá, Cauca, Chocó, Guajira, Guaviare, Magdalena, Meta, Nariño, Norte de Santander, Valle del Cauca, Vaupés), Trinidad, Venezuela (Amazonas, Apure, Aragua, Barinas, Bolívar, Carabobo, Lara, Mérida, Miranda, Portuguesa, Sucre, Táchira, Trujillo, Yaracuy, Zulia), Guyana, Ecuador (Carchi, Esmeraldas, Guayas, Los Rios, Manabí), Peru (Cuzco, Huánuco, Junín, Loreto, Ucayali), Brazil (Acre, Amazonas, Roraima).

Habitat & Ecology — In moist forests, lowland thickets, or in swamps, at elevations of 0–1500 m. Flowering all year round.

IUCN Conservation Status — Not evaluated. There are 327 collection records examined of which 181 specimens were collected within the past 50 years. This species is also reported in 57 recent iNaturalist observations as a variety of *C. guanaiensis* from which it was split off as a separate species (https://www.inaturalist.org/observations?place_id=any&taxon_id=314881). The species is distributed over a wide range from Mexico to Peru and Brazil as well as the islands of Puerto Rico and Trinidad. Population and threat data is not available but based on the number of collection records and our general field observations, this species would probably score a status of Least Concern.

Vernacular names — Belize: W’eh-te (Maya, Kekchí). Brazil: Canafiche, Canafistula, Canafixe, Naxurumeake (Yanomami). Colombia: Caña agria, Cañagria, Cañaguate, Ñá-ka (Taiwano). Costa Rica: Caña agria. Ecuador: Caña agria. Peru: Sacha huiro. Puerto Rico: Caña agria, Cana di India. Venezuela: Auti (Piapora), Caña de la India.

Notes — *Costus macrostrobilus* was formerly treated by Maas (1972) as a variety of *C. guanaiensis*. It is characterized by bracts with distinct, green, thick and leafy appendages, which are concave and somewhat pungent at the apex. Moreover, the leaves are very well recognizable as both sides are more or less densely hairy. An additional feature that is particularly useful in recognizing herbarium material is that the adaxial surface of the leaves is somewhat scabrid to the touch, a feature found in only a few other species (e.g., *C. lima*).

The typical form is encountered in Central America and western South America, including Trinidad. The type collection is from Puerto Rico, but it is probable that it does not occur naturally there.

Within this species we see a lot of variation, and additional field work is needed to determine the extent of variation within this species versus potential for designating separate species. For example, a form with plicate leaves which are purplish on the abaxial surface is found in NW Ecuador (near Bilsa and Lita). Additional material is needed to confirm if these collections remain included in *C. macrostrobilus* or instead represent an undescribed species.

**44. *Costus malortieanus*** H.Wendl. — Fig. 10g; Map 31

*Costus malortieanus* H.Wendl. (1863) 30. — Neotype (designated here): *McDowell 126* (neo DUKE; isoneo F, MO [MO-356580], NY), Costa Rica, Heredia, Finca La Selva, the OTS Field Station on the Río Puerto Viejo, just E of its junction with the Río Sarapiquí, 100 m, 17 Sept. 1982. A neotype is designated as there is no original herbarium material preserved of Wendland’s species (from Costa Rica, Heredia, near La Muelle and near Río Sarapiquí, Aug. 1857).

[*Costus elegans* Hortul. ex Petersen (1890) 52, nom. nud., non Veitch ex J.Dix (1860)].

Herbs, 0.5–2 m tall. *Leaves* 7–10, concentrated toward the top of the shoot; sheaths fragile when dry, 10–20 mm diam; ligule truncate, hardly 1 mm long; petiole to c. 2 mm long; sheaths, ligule and petiole densely villose, puberulous to glabrous; lamina obovate to elliptic, 25–35 by 11–18 cm, pale green with dark green bands converging from base to apex above, glaucous below, upper side densely villose, lower side densely puberulous, base acute, apex acuminate (acumen 2–10 mm long) and apiculate. *Inflorescence* globose, sometimes cylindric, 4–9 by 4 cm, terminating a leafy shoot; bracts, bracteole, calyx, ovary, and capsule glabrous*. Flowers* abaxially orientated to erect; bracts green, coriaceous, broadly ovate, 2–4 by 2–4 cm, apex obtuse, callus 3–6 mm long; bracteole boat-shaped, 14–20 mm long; calyx red to green, 5–9 mm long, lobes shallowly triangular, 2–3 mm long; corolla yellow to pinkish white, 50–70 mm long, glabrous, lobes narrowly obovate, 40–50 mm long; labellum yellow, upper part horizontally spreading, broadly obovate, 50–70 by 40–50 mm, lateral lobes usually striped with dark red, middle lobe recurved with a dark yellow honey mark, 3-lobulate, lobules 5–10 mm long, margin slightly crenulate; stamen white, slightly tinged with purple, 35–45 by 10–15 mm, not exceeding the labellum, apex yellow, pink, or dark red, obtuse to obscurely 3-lobulate, anther 5–7 mm long. *Capsule* ellipsoid, c. 12 mm long.

Distribution — Nicaragua, Costa Rica.

Habitat & Ecology — In forests, at elevations of 0–950 m. Flowering all year round. *Frankie 49a* (MO) from Costa Rica, Finca La Selva, noticed that the flowers are visited by Euglossine bees.

IUCN Conservation Status — Least Concern.

Vernacular names — Costa Rica: Chitania. Nicaragua: Cañagria.

Notes — *Costus malortieanus* is generally a small herb very well characterized by obovate to elliptic leaves to 18 cm wide, an extremely short ligule (hardly 1 mm long), an often globose inflorescence with green bracts, and a yellow labellum striped with red. Its somewhat resembles *C. pictus*, from which it differs by the densely hairy and much wider leaves.

**45. *Costus mollissimus*** Maas & H.Maas — Fig. 10h; Map 31

*Costus mollissimus* Maas & H.Maas in Maas et al. (2023) 102, f. 13. — Type: *Ishiki, Maas, Maas-van de Kamer et al. 2400* (holo L, 2 sheets [L4368349 & L4368350]; iso ECOSUR), Mexico, Chiapas, Mun. Chilon, 22 km from Temo in the direction of Palenque, 2 km from Patathel, near El Chich, 850 m, 4 Nov. 1998.

Herbs, 0.5–2.5 m tall. *Leaves*: sheaths 12–15 mm diam; ligule truncate, 1–4 mm long; petiole 10–15 mm long; sheaths, ligule and petiole densely velutinous; lamina narrowly elliptic, sometimes elliptic, 18–36 by 6–13 cm, slightly 5–8-plicate, upper side shiny, glabrous, lower side densely velutinous, base acute, obtuse, to cordate, apex acuminate (acumen 5–15 mm long). *Inflorescence* cylindric to subglobose, 5–14 by 4–5 cm, terminating a leafy shoot; bracts, appendages of bracts, bracteole, calyx, ovary, and capsule densely velutinous*. Flowers* abaxially orientated; bracts green, red, or pink, coriaceous, broadly elliptic, 2–2.5 by 2–3 cm; appendages green, foliaceous, patent to slightly reflexed, triangular to broadly triangular, 0.5–3.5 by 1.2–2.5 cm, apex acute to obtuse; bracteole boat-shaped, 16–20 mm long; calyx red to pink, 9–10 mm long, lobes broadly to shallowly triangular, 2–5 mm long, with a hairy tuft at the apex; corolla cream, yellow, or pale orange, 50–60 mm long, densely velutinous, lobes narrowly elliptic, 30– 40 mm long; labellum yellow, upper part horizontally spreading, broadly obovate, 30–40 by 40– 50 mm, lateral lobes dark red with yellow venation, middle lobe recurved with a yellow honey mark, margin dentate; stamen yellow, 30–35 by 10–12 mm, not exceeding the labellum, apex red, slightly 3-dentate, anther c. 8 mm long. *Capsule* broadly obovoid to obovoid, 12–17 mm long.

Distribution — Mexico (Chiapas, Oaxaca, Tabasco, Veracruz).

Habitat & Ecology — In forests (a.o. “selva baja/alta perennifolia”) at elevations of 200– 1000(–1500) m. Flowering all year round.

IUCN Conservation Status — Not evaluated. There are only 5 collection records examined for this recently described species, found only in Chiapas and Oaxaca in Mexico. This species is also reported in 4 recent iNaturalist observations (https://www.inaturalist.org/observations?place_id=any&taxon_id=1460499), 3 of which are in Chiapas and one in Tabasco. The species is currently too data deficient to base a formal assessment, but with few records over a limited range we believe a full assessment would result in a status of Vulnerable or possibly Endangered.

Vernacular name — Mexico: Apagafuego.

Notes — *Costus mollissimus* is easily recognizable by a velutinous indument on most parts of the plant, appendaged bracts and a very short ligule. It superficially resembles *C. dirzoi*, both sharing the velutinous indument all over the plant, but it is aberrant in having bracts which are provided with foliaceous, green appendages and by a shorter ligule (1–4 mm versus 3–12 mm long). Moreover, *C. dirzoi* has bracts with a distinct callus which is absent in *C. mollissimus*.

**46. *Costus montanus*** Maas — Fig. 10i; Map 32

*Costus montanus* Maas (1972) 119, f. 57. — Type: *Brenes 22607* (holo NY [NY00320327]), Costa Rica, Alajuela, San Pedro de San Ramón, 4 May 1937.

Herbs, 1–2 m tall. *Leaves*: sheaths 5–12(–20) mm diam; ligule (5–)15–40 mm long, acute, inequally 2-lobed, largest lobe triangular, c. 10 mm long, smallest lobe depressed triangular, 3–5 mm long; petiole 8–20 mm long; sheaths, ligule and petiole densely brownish velutinous; lamina narrowly elliptic, 13–28 by 4–8.5 cm, upper side glabrous, but primary vein covered with a row of erect hairs, lower side densely brownish velutinous, base acute, apex acuminate (acumen 10– 20 mm long). *Inflorescence* ovoid, 4–10 by 2.5–4.5 cm, terminating a leafy shoot; bracts, appendages of bracts, bracteole, calyx, ovary, and capsule densely brownish velutinous*. Flowers* abaxially orientated; bracts red, coriaceous, broadly ovate, to c. 2.5 by 2.5 cm, apex acute; appendages when present red, foliaceous, recurved or reflexed, triangular to narrowly triangular, to c. 2 by 1 cm (lowermost to c. 6 by 1.5 cm), apex acute; bracteole boat-shaped, 17–25 mm long; calyx red, 11–16 mm long, lobes depressed triangular, 2–4 mm long; corolla orange to yellowish orange, 40–55 mm long, densely brownish velutinous, lobes narrowly obovate, 35–45 mm long; labellum orange to yellowish orange, lateral lobes striped with dark orange to red, rolled inwards and forming a tube 10–12 mm diam, obovate when spread out, 25–35 by 15–20 mm, middle lobe recurved with a yellow honey mark, irregularly lobulate, lobules 1–2 mm long; stamen yellow to orange, 25–35 by 6–8 mm, not exceeding the labellum, apex acute to obtuse, anther 7–8 mm long. *Capsule* globose to obovoid, 10–15 mm long.

Distribution — Costa Rica.

Habitat & Ecology — In moist cloud forests, along streams, at elevations of (250–)1300– 1800(–2100) m. Flowering all year round.

IUCN Conservation Status — Near Threatened.

Vernacular names — Costa Rica: Caña agria.

Notes — *Costus montanus* is a species endemic to Costa Rica and restricted to high elevations. It is readily recognized by a very long, unequally acutely 2-lobed ligule and a brownish velutinous indument on most parts of the plant, including the corolla. Another striking feature of this species is the fact that the calyx is often exceeding the bracts during fructification.

**47. *Costus nitidus*** Maas — Fig. 11a; Map 33

*Costus nitidus* Maas (1976) 473. — Type: *Maas & Cramer 1345* (holo U, 3 sheets [U0007214, U0007215, U0007216]), Costa Rica, Cartago, forested hills of Río Reventazón, near grounds of IICA, Turrialba, 600 m, 16 Aug. 1974.

Herbs, 2–4 m tall. *Leaves*: sheaths shiny, 10–20 mm diam; ligule obliquely truncate to slightly 2-lobed, 20–40 mm long; petiole 8–15 mm long; sheaths, ligule and petiole glabrous; lamina narrowly elliptic, 30–45 by 7–10 cm, slightly 5–6-plicate, mostly shiny on both sides, upper and lower side glabrous, margin densely covered with erect, brownish hairs, base acute, apex acute to acuminate (acumen 15–25 mm long). *Inflorescence* ovoid, 7–16 by 3–5.5 cm, enlarging to c. 22 by 8 cm in fruit, terminating a leafy shoot; bracts, bracteole, calyx, ovary, and capsule glabrous*. Flower* abaxially orientated; bracts pink, orange, or red, coriaceous, broadly ovate, 3–5 by 3–5 cm, apex obtuse, callus 8–18 mm long; bracteole boat-shaped, 18–20 mm long; calyx whitish, 7– 10 mm long, lobes shallowly to very shallowly triangular, 1–3 mm long; corolla yellow, 60–75 mm long, glabrous, lobes narrowly elliptic, 40–45 mm long; labellum yellow, lateral lobes rolled inwards and forming a slightly curved tube 10–12 mm diam, oblong-obovate when spread out, 30–35 by 15–20 mm, 5-lobulate, lobules narrowly triangular, 2–5 mm long; stamen yellow, 30– 35 by 10–13 mm, slightly exceeding the labellum, apex acute, anther 9–10 mm long. *Capsule* ellipsoid, 10–20 mm long.

Distribution — Costa Rica, Panama.

Habitat & Ecology — In forests, at elevations of 0–900 m. Flowering in the rainy season.

IUCN Conservation Status — Endangered.

Notes — *Costus nitidus* is well recognizable by strongly shiny leaves and bracts, yellow, tubular flowers, and a ligule 20–40 mm long. It closely resembles *C. plicatus*, sharing the long ligule and most flower characters, but in *C. plicatus* the leaves are strongly plicate whereas they mostly lack any plicae in *C. nitidus*.

**48. *Costus obscurus*** D.Skinner & Maas — Fig. 11b; Map 34

*Costus obscurus* D.Skinner & Maas in Maas et al. (2023) 104, f. 14. — Type: *Plowman & Davis 5054* (holo USM; iso GH, L [L1480369, L1480371, L1480372]), Peru, Madre de Dios, Prov. Manu, near Shintuya, on road to Salvación, 600 m, 9 Feb 1974.

Herbs, 1–1.5 m tall. *Leaves*: sheaths 10–20 mm diam; ligule truncate, 10–15 mm long; petiole 8– 12 mm long; sheaths, ligule and petiole densely to rather densely whitish villose; lamina narrowly elliptic, 32–45 by 9–17 cm, upper side dark green, lower side dark purple, upper side rather densely to densely whitish villose, lower side densely to sparsely whitish puberulous, base acute to cordate, apex acuminate (acumen 5–10 mm long). *Inflorescence* ovoid, 8–13 by 5–7 cm, terminating a leafy shoot; bracts, appendages of bracts, bracteole, calyx, ovary, and capsule rather densely to sparsely puberulous*. Flowers* abaxially orientated; bracts green in the exposed part, red or green in the covered part, coriaceous, broadly ovate-triangular, 3.5–4.5 by 3–3.5 cm; appendages green, erect, broadly ovate-triangular, 1.5–2 by 1.8–2 cm, or almost absent, apex rounded; bracteole boat-shaped, 35–37 mm long; calyx red, 25–27 mm long, deeply split on one side, lobes deltate to broadly triangular, c. 2 mm long; corolla cream to yellow, c. 75 mm long, glabrous, lobes narrowly elliptic, 55–60 mm long; labellum pale yellow, upper part horizontally spreading, broadly obovate, c. 60–70 by 60–70 mm, lateral lobes strongly striped with pink or red, middle lobe recurved with a yellow honey mark, margin irregularly dentate to lobulate; stamen white to pink, 35–45 by 12–15 mm, not exceeding the labellum, apex pinkish red, deeply and irregularly lobed, anther 7–8 mm long. *Capsule* ellipsoid, 12–15 mm long.

Distribution — Peru (Cuzco, Huánuco, Madre de Dios, San Martín).

Habitat & Ecology — In forests, in wet, shady areas, often in disturbed places, at elevations of 450–1300 m. Flowering in the rainy season.

IUCN Conservation Status — Not evaluated.

There are only 9 collection records examined for this recently described species, found only in Peru. This species is also reported in 3 recent iNaturalist observations (https://www.inaturalist.org/observations?place_id=any&taxon_id=1460541). This recently described species has been found growing only in deep shade in very wet and muddy soil and might require this specific habitat. Our field observations indicate that this is a very uncommon species and would probably score as Vulnerable or Endangered in a full assessment.

Notes — *Costus obscurus* can be recognized by its large, dark green leaves with dark purple undersides and densely whitish villose sheaths. The dark green upper sides of the leaves and contrasting purple undersides are striking in appearance. Plants of *C. obscurus* have a compact appearance with leaves closely spaced together along the shoot and generally pointed upward rather than horizontal or drooping. *Costus obscurus* can be confused with *C. erythrophyllus*, but can be distinguished from that species by its shorter, truncate ligule instead of a long and deeply 2-lobed ligule as well as by having smooth rather than (mostly) plicate leaves.

**49. *Costus oreophilus*** Maas & D.Skinner — Fig. 11c; Map 34

*Costus oreophilus* Maas & D.Skinner in Maas et al. (2023) 107, f. 15. — Type: *Asplund 20068* (holo S, 2 sheets [S06-1367, S06-13643]; iso GB), Ecuador, Tunguruhua, Río Verde Grande, 1500 m, 30 March 1956.

Herbs, 3–4 m tall. *Leaves*: sheaths 15–20 mm diam; ligule truncate, 3–10 mm long; petiole 5–10 mm long; sheaths, ligule and petiole densely to sparsely puberulous; lamina narrowly elliptic, 20–38 by 7–11 cm, upper side shiny, lower side green, upper side glabrous, sometimes rather densely puberulous, lower side densely to sparely puberulous to velutinous, base cordate to acute, apex acuminate (acumen 5–10 mm long). *Inflorescence* cylindric to ovoid, 7–12 by 3–6 cm, wrapped tightly by the upper leaves, terminating a leafy shoot; bracts, bracteole, calyx, ovary, and capsule sparsely puberulous to glabrous*. Flowers* adaxially orientated to erect; bracts green in the exposed part, red in the covered part, sometimes completely red, coriaceous, broadly ovate, 3.5–4.5 by 3–4 cm, apex obtuse, callus inconspicuous; bracteole boat-shaped, 18–31 mm long; calyx red, 6–12 mm long, lobes shallowly triangular, 3–4 mm long; corolla white to yellow, 50–60 mm long, glabrous, lobes recurved, narrowly obovate, 40–55 mm long; labellum white, upper part horizontally spreading, broadly obovate, 70–75 by 70–75 mm, lateral lobes striped with red, middle lobe recurved with a yellow honey mark, margin irregularly crenulate; stamen red to pink, 35–40 by 12–13 mm, not exceeding the labellum, apex red, irregularly dentate, anther 10–12 mm long. *Capsule* ellipsoid, c. 15 mm long.

Distribution — Ecuador (Pastaza, Tungurahua, Zamora-Chinchipe).

Habitat & Ecology — In forests, at elevations of 1150–1600 m. Flowering in the rainy season.

IUCN Conservation Status — Not evaluated. There are only 3 collection records examined for this recently described species, which is found only in mountainous parts of Ecuador. This species is also reported in 4 recent iNaturalist observations (https://www.inaturalist.org/observations?place_id=any&taxon_id=1460541). Our field observations indicate that this is a very uncommon species and would probably score as Vulnerable or Endangered in a full assessment.

Notes — Most material of *Costus oreophilus* was identified by Maas (1972) with a note under the previous circumscription of *Costus laevis* (now *C. kuntzei*): “Three Ecuadorian collections deviated by being puberulous to velutinous on the lower side of the leaves. The flower material being very incomplete I could not determine their taxonomic status, i.e., whether it is a variety of *C. laevis* or a distinct species”. Fifty years later, we decided that these collections merit specific rank (Maas et al. 2023).

*Costus oreophilus* differs from *C. convexus* by having hairy instead of glabrous sheaths and leaves, a smaller calyx (6–12 versus 15–25 mm long), a smaller ligule (3–10 versus 20–45 mm long). It can be confused with *C. kuntzei*, with which it shares the mostly green bracts and adaxially orientated flowers, but it is distinguished by the puberulous sheaths and leaves and a smaller ligule (3–10 mm versus 5–20 mm long); moreover the corolla lobes are often recurved, a feature rarely seen in *Costus*.

**50. *Costus osae*** Maas & H.Maas — Fig. 11d; Map 35

*Costus osae* Maas & H.Maas (1997) 277, f. 2. — Type: *Herrera C. 3979* (holo U [U0006854]; iso CR, F, K, MO [MO-902895]), Costa Rica, Puntarenas, Osa Peninsula, Parque Nacional Vaquedano, Cerro Brujo, Quebrada Vaquedano, 400 m, 20 July 1990.

Herbs, 0.5–1 m tall. *Leaves* 3–7, concentrated toward the top of the shoot; sheaths 5–15 mm diam; ligule truncate, (2–)5–9 mm long; petiole 4–18 mm long; sheaths, ligule and petiole densely to rather densely villose; lamina obovate to elliptic, (10–)13–31 by (6–)8–16 cm, upper and lower side densely to rather densely villose, base acute, apex acuminate (acumen 5–10 mm long) to acute. *Inflorescence* cylindric, 5–10 by 4–6 cm, elongating to c. 10 cm in fruit, terminating a leafy shoot; bracts, appendages of bracts, bracteole, calyx, ovary and capsule densely to rather densely puberulous*. Flowers* abaxially orientated; bracts red, orange, or pink, chartaceous to coriaceous, obovate to broadly ovate, 2.5–4.5 by 1–2.5 cm, callus inconspicuous; appendages red, red-orange, or yellow, foliaceous, more or less distinctly separated from the basal part of the bract proper which is erect and ascending, upper part more or less patent and slightly concave, triangular-ovate to deltate, 1–4 by 1.5–2 cm, apex acute; bracteole 2-keeled, 25–35 mm long, 2-keeled; calyx reddish to red-orange, 12–20 mm long, lobes ovate to broadly ovate, 2–5 mm long; corolla pink, red, or yellow with red venation, 45–50 mm long, densely to sparsely puberulous, lobes ovate to narrowly ovate, 35–40 mm long; labellum basally white, lateral lobes rolled inwards and forming a tube 5–10 mm diam, obovate when spread out, 27–35 by 15–20 mm, irregularly lobulate, lobules dark red, shallowly triangular to triangular, 2–3 mm long, middle lobe with yellowish honey mark; stamen white, 25–35 by 6–8 mm, only slightly exceeding the labellum, apex dark red, irregularly dentate, anther 6–8 mm long. *Capsul*e obovoid, c. 13 mm long.

Distribution — Costa Rica, Colombia (Antioquia), Ecuador (Cotopaxi, Los Rios).

Habitat & Ecology — In high forests (often on steep rocky slopes), at elevations of 0–400 m. Flowering in the rainy season.

IUCN Conservation Status — Vulnerable.

Vernacular names — Colombia: Cañagria.

Notes — *Costus osae* is a small herb with few leaves concentrated at the top of the shoot, like in *C. malortieanus*. It is well characterized by obovate to elliptic, broad leaf blades with both sides densely to rather densely villose, and an inflorescence composed of reddish bracts with concave and more or less patent appendages. Another feature of this species is the 2-keeled bracteole.

**51. *Costus pictus*** D.Don — Fig. 11e, f; Map 36

*Costus pictus* D.Don (1833) t. 1594. — Type: *Plate 1594* in Don (1833). Nothing of the original material introduced by Deppe from Mexico, and cultivated and flowering in 1832 in Boyton, England, has been preserved.

*Costus mexicanus* Liebm. ex Petersen (1893) 261, t. 16. — Type: *A plant cultivated in Hort. Bot. Haun., flowering in September 1891* (holo C, spirit collection). Probably collected by Liebmann.

*Costus congestus* Rowlee (1922) 291, pl. 14, p.p., excluding descriptions of ovary and calyx. — Lectotype (designated here): *Donnell Smith 2036* (lecto US [US00092957]), Guatemala, Prov. Escuintla, Escuintla, 1100 ft, March 1890.

Herbs, 0.5–4 m tall. *Leaves*: sheaths 5–15 mm diam; ligule truncate, 2–4(–8) mm long; petiole 10–18 mm long, sometimes dark purple; sheaths, ligule and petiole glabrous, rarely ligule covered with erect, long and stiff hairs; lamina narrowly elliptic to narrowly obovate, often with wavy margin, 10–50 by 3–14 cm, upper side glabrous, rarely sparsely villose, lower side glabrous, sometimes rather densely puberulous, rarely sparsely villose, base acute, rounded, or slightly cordate, apex acuminate (acumen 10–20 mm long) to long-acute. *Inflorescence* globose to ovoid, 3–8 by 3–4 cm, elongating to c. 13 cm in fruit, terminating a leafy shoot; bracts, bracteole, calyx, ovary, and capsule glabrous to sparsely puberulous*. Flowers* abaxially orientated; bracts green, rarely orange-red, coriaceous, broadly ovate, 2–5 by 2–6 cm, apex obtuse, callus inconspicuous, 0–7 mm long; bracteole boat-shaped, 16–22 mm long; calyx red, (4–)6–9(–10) mm long, lobes deltate to shallowly triangular, 2–3.5 mm long; corolla yellow, orange, or red, 60–65 mm long, glabrous, lobes narrowly elliptic to narrowly ovate, 45–50 mm long; labellum yellow, white, or red, upper part horizontally spreading, broadly obovate, 45–65 by 40–50 mm, lateral lobes striped with dark red, middle lobe recurved with a yellow honey mark, margin irregularly crenate; stamen yellow to white, 35–45 by 10–20 mm, not exceeding the labellum, apex dark red, acute, irregularly dentate, anther 7–8 mm long. *Capsule* subglobose, 11– 15 mm diam.

Distribution — Mexico (Chiapas, Guerrero, Hidalgo, Jalisco, Michoacán, Nayarit, Oaxaca, Puebla, Quintana Roo, San Luis Potosí, Sinaloa, Tabasco, Tamaulipas, Veracruz, Yucatán), Guatemala, El Salvador, Belize, Honduras, Nicaragua, Costa Rica.

Habitat & Ecology — In rain forests, clearings, and hill forests, along watercourses and roads, at elevations down to sea level, but mainly from 300–1800 m. Flowering all year round but plants on the Pacific side, where there is a distinct and strong dry season, are generally flowering during the rainy season. On *Bailey 196* (U) from Belize it is mentioned that the flowers are visited by hummingbirds.

IUCN Conservation Status — Not evaluated. There are 187 collection records examined of which 107 specimens were collected within the past 50 years. This species is also reported in 185 recent “Research Grade” iNaturalist observations (https://www.inaturalist.org/observations?place_id=any&taxon_id=276565). The species is distributed over a wide range from Mexico to Costa Rica and the island of Puerto Rico. Population and threat data is not available but based on the number of collection records and our general field observations, this species would almost certainly score a status of Least Concern.

Vernacular names — El Salvador: Caña de Cristo. Guatemala: Caña Criste, Caña de Cristo. Honduras: Caña de Cristo. Mexico: Baj tsa, Cañagria, Caña agrio, Caña Cristo, Caña de agua, Caña de javali, Caña de Indio, Caña del Indio, Caña de puerco, Caña de venado, Flor de lanten, Masa–owat, Riñonera. Nicaragua: Cañagria, Caña agria.

Notes — *Costus pictus* can be recognized by a mostly green inflorescence in which the callus of the bracts is inconspicuous or even absent, and even more importantly by a very short ligule; the flowers have a spreading labellum, and are yellow to white with red stripes. The shoots of this species are sometimes red or green with red stripes or dots in plants on the Atlantic side.

In most collections studied the sheaths, petiole, and ligule are completely glabrous, but in various collections the ligule is covered with erect, long, and stiff hairs (*McDade 77-706* (F) from Mexico, *Standley 90115* (F) from Guatemala, and *Allen 6780* (U) from Honduras). This feature is nicely shown in *Maas & Maas-van de Kamer 10540* (U) (origin unknown), which was photographed in the botanical greenhouses of München. This collection was aberrant, however, by having the inflorescence produced from a separate leafless shoot. Plants seen in Guatemala and Honduras on the Atlantic side often have inflorescences only on separate, leafless shoots.

As already noted by Maas (1972), several collections from Guatemala and Mexico are possibly hybrids between *C. pictus* and *C. pulverulentus* or have introgression with *C. pulverulentus*. There is also a form included in *Costus pictus* that produces red bracts instead of green bracts and occurs along the Pacific coast of Central America, from southern Mexico to Costa Rica. The inflorescences of these plants appear exclusively terminally on a leafy shoot. Phylogenetic results (Valderrama et al. 2020) indicate that this is a separate species from the plants found on the Atlantic side, but further research is needed to determine the status of this form.

**52. *Costus pitalito*** C.D.Specht & H.Maas — Fig. 11g; Map 37

*Costus pitalito* C.D.Specht & H.Maas in Maas et al. (2023) 109, f. 16. — Type: *Maas, Maas-van de Kamer, Valderrama, Specht, Erkens & Rodriguez 10733* (holo L, 2 sheets [L4468397, L4468398]), Colombia, Cauca, forested roadside from San Juan de Villalobos to Pitalito, 1590 m, 19 July 2018.

Herbs, 1.7– 2.5 m tall. *Leaves*: sheaths c. 35 mm diam; ligule deeply 2-lobed, 45–60 mm long, lobes rounded, obtuse; petiole 10–20 mm long; sheaths, ligule and petiole densely, brownish villose; lamina narrowly elliptic, 30–45 by 12.5–14 cm, upper side glabrous, but sparsely brownish villose when young, lower side densely brownish villose, base acute, apex acuminate (acumen c. 10 mm long). *Inflorescence* suglobose, 11–12 by 9–11 cm, terminating a leafy shoot; bracts rather densely villose, appendages of bracts sparsely villose to glabrous, its margins densely brownish villose, bracteole and calyx sparsely puberulous, ovary and capsule rather densely sericeous*. Flowers* abaxially orientated to erect; bracts red, coriaceous, broadly ovate, 3– 4 by 2–3 cm; appendages green, foliaceous, cup-shaped, erect to patent, broadly ovate-triangular, 1–1.5 by 1–1.5 cm, apex obtuse to acute; bracteole 1- to 2-keeled, 30–32 mm long; calyx red, apex green, 20–22 mm long, lobes ovate-triangular, 10–12 mm long; corolla pale yellow to white, base pinkish, 55–60 mm long, glabrous, lobes narrowly elliptic, 35–40 mm long, with filiform apex; labellum pale yellow to white, yellow in throat, upper part horizontally spreading, broadly obovate, c. 70 by 70–80 mm, margin irregularly dentate to crenate; stamen cream, c. 50 by 15 mm, apex reflexed, irregularly dentate, not exceeding the labellum, anther c. 13 mm long. *Capsule* ellipsoid, c. 25 mm long.

Distribution — Colombia (Cauca, Putumayo).

Habitat & Ecology — Along forested but disturbed roadsides, at an elevation of c. 1600 m. Flowering and fruiting: May and July.

IUCN Conservation Status — Not evaluated. There are only 2 collection records examined for this recently described species, which has been found only in Colombia at the type location near Pitalito and another herbarium specimen from Putumayo. No iNaturalist observations of this species have been recorded. Our field observations indicate that this is a very uncommon species and would probably score as Endangered and possibly Critically Endangered in a full assessment.

Notes — *Costus pitalito* looks superficially like *C. leucanthus* based on the shared light-coloured flowers and a very long ligule, but it differs from *C. leucanthus* by markedly obtuse ligule lobes and bracts with densely brownish villose margins. The flower of *C. pitalito* is also more cream-colored and hyaline than the bright white flowers of *C. leucanthus*. This species appears to be relatively rare given the few collections despite being a large and conspicuous plant.

**53. *Costus plicatus*** Maas — Fig. 11h; Map 37

*Costus plicatus* Maas (1976) 472. — Type: *Maas & Dressler 1619* (holo U, 2 sheets [U123415, U123416]; iso MO [MO-476242]), Panama, Veraguas, quebrada near Río Segundo Brazo, 700 m, 7 Sept. 1974.

Herbs, 0.5–4 m tall. *Leaves*: sheaths 5–20 mm diam; ligule very obliquely truncate to 2-lobed, 25–60 mm long; petiole 15–20 mm long; sheaths, ligule and petiole glabrous; lamina narrowly elliptic to narrowly obovate, 25–30(–48) by (7–)8.5–13(–19) cm, rather asymmetrical, distinctly 6–8-plicate, upper side glabrous, lower side glabrous to sparsely puberulous, margin densely covered with long, silky hairs, often becoming glabrous with age, base acute, apex long-acuminate (acumen 15–25 mm long). *Inflorescence* cylindric to ovoid, 5–15 by 3–5 cm, terminating a leafy shoot; bracts, bracteole, calyx, ovary, and capsule glabrous*. Flowers* abaxially orientated to erect; bracts pink to red, coriaceous, broadly ovate-deltate, 2.5–4 by 2–3 cm, apex acute, callus 5–11 mm long; bracteole boat-shaped, 17–18 mm long; calyx pink, 7–10 mm long, lobes very shallowly triangular, 0.5–1.5 mm long; corolla yellow, 45–55 mm long, glabrous, lobes narrowly elliptic, 30–40 mm long; labellum yellow, lateral lobes rolled inwards and forming a tube 9–10 mm diam, oblong-ovate when spread out, 30–35 by 15–20 mm, 5-lobulate, lobules shallowly triangular to triangular, 2–5 mm long; stamen yellow, 35–40 by 10– 12 mm, slightly exceeding the labellum, apex acute, anther c. 11 mm long. *Capsul*e subglobose, 12–13 mm diam.

Distribution — Costa Rica, Panama.

Habitat & Ecology — In forests, at elevations of 0–850 m. Flowering mainly in the rainy season.

IUCN Conservation Status – Least Concern.

Notes — *Costus plicatus* is a Central American species, which is easily distinguished by a combination of strongly plicate leaves, a long petiole (15–20 mm), and a long ligule (25–60 mm). It closely resembles another Central American species, *C. nitidus*, but differs by having plicate leaves.

**54. *Costus plowmanii*** Maas — Fig. 11i; Map 37

*Costus plowmanii* Maas (1976) 470. — Type: *Maas & Plowman 1900* (holo U, 3 sheets [U0007890, U0007891, U0007892]; iso AAU, COL [COL606918, COL606934], E [E00319722], F [F0047178F], GH [GH00030657], K [K000586765], MO [MO-1456319, MO-202810], NY [NY00320345], P, US [US00092971], VEN [VEN121940], Z), Colombia, Valle del Cauca, old road from Cali to Buenaventura, steep forested slopes near Queremal, Cordillera Occidental, 1350 m, 3 Oct. 1974.

Herbs, 1.5–4 m tall. *Leaves*: sheaths 8–20 mm diam; ligule obliquely truncate, 2–15(–20) mm long; petiole 5–20 mm long; sheaths, ligule and petiole densely to sparsely villose, rarely sparsely puberulous; lamina narrowly elliptic to narrowly obovate, sometimes elliptic, 20–47 by 7–18 cm, 5–10-plicate, upper side glabrous, but primary vein sometimes covered with a row of erect hairs, lower side densely to rather densely villose, base acute to obtuse, apex acuminate (acumen 5–20 mm long). *Inflorescence* ovoid to cylindric, 7–25 by 4–8.5 cm, terminating a separate leafless shoot 40–75 cm long, sheaths obliquely truncate, 4–8 cm long, sparsely villose to glabrous; bracts, bracteole, calyx, ovary, and capsule glabrous, sometimes sparsely puberulous*. Flowers* abaxially orientated; bracts red, coriaceous, broadly obovate, 3.5–5 by 2.5– 4 cm, apex obtuse, callus 2–8 mm long; bracteole boat-shaped, 20–29 mm long; calyx red, 8–14 mm long, lobes deltate to shallowly triangular, 2–4 mm long; corolla yellow, 45–50 mm long, glabrous, lobes narrowly elliptic, c. 35 mm long; labellum yellow, lateral lobes rolled inwards and forming a curved tube 8–10 mm diam, oblong-elliptic when spread out, 30–35 by 17–25 mm, 5-lobulate, the middle lobule filiform, c. 4 mm long, the others irregularly triangular, 2–4 mm long; stamen yellow, 33–40 by 10–11 mm, slightly exceeding the labellum, apex acute, anther 7–11 mm long. *Capsule* ellipsoid, 8–15 mm long.

Distribution — Colombia (Antioquia, Cauca, Chocó, Nariño, Risaralda, Valle del Cauca).

Habitat & Ecology — In steep slopes in premontane forests, at elevations of 950–1950 m. Flowering mainly in the rainy season.

IUCN Conservation Status — Not evaluated. There are 29 collection records examined for this species, which is found only in the western cordillera of Colombia. This species is also reported in 7 recent iNaturalist observations (https://www.inaturalist.org/observations?place_id=any&taxon_id=504249). The population data for this species is not available but based on its limited range and specific habitat it would probably score as Vulnerable in a full assessment.

Vernacular names — Colombia: Cañaguate.

Notes — *Costus plowmanii* is characterized by an inflorescence produced on a separate leafless shoot, red bracts, yellow and tubular flowers, and a villose lower side of the lamina. It resembles *C. beckii*, but that species has a much longer ligule (55–60 mm long) and a hairy corolla.

*Franco Rosselli et al. 4879* (COL), from Colombia, Nariño, Mun. Barbacoas, Corregimiento Altaquer, Vereda El Barro, Reserva Natural Ñambi, 1325 m, may belong to *C. plowmanii*, but its leaves are almost completely glabrous, and the corolla is reddish (“corolla roja nacarada”) instead of yellow.

**55. *Costus prancei*** Maas & H.Maas — Fig. 12a; Map 37

*Costus prancei* Maas & H.Maas in Maas et al. (2023) 111, f. 17. — Type: *Prance, Maas et al. 12265* (holo U, 2 sheets [U1607120, U1603130 (spirit collection)]; iso INPA, NY [NY00867011]), Brazil, Acre, vicinity of Serra da Moa, 22 Apr. 1971.

Herbs, 1–1.5 m tall. *Leaves*: sheaths 8–10 mm diam; ligule truncate, 2–7 mm long; petiole 2–8 mm long; sheaths, ligule, and petiole densely to sparsely villose; lamina narrowly elliptic, 11–25 by 4–7 cm, upper side densely villose to glabrous, with a dense row of hairs along the primary vein, lower side densely puberulous to villose, base acute, rounded, or cordate, apex acuminate (acumen c. 15 mm long). *Inflorescence* cylindric, 3–15 by 1.5–4 cm, terminating a leafy shoot; bracts, bracteole, calyx, ovary, and fruit densely puberulo-villose to glabrous*. Flowers* abaxially orientated; bracts red, coriaceous to chartaceous, broadly to depressed ovate, 1.5–2.5 by 1.5–3 cm, callus 3–5 mm long; bracteole boat-shaped, 9–15 mm long; calyx red to pale orange-red, 6– 12 mm long, lobes shallowly triangular, c. 1 mm long; corolla yellow to orange (2 labels indicated cream and pink to orange!), 20–30 mm long, glabrous, lobes narrowly elliptic, 20–25 mm long; labellum yellow to orange, lateral lobes involute and forming a straight tube 7–8 mm diam, oblong-obovate when spread out, 20–25 by 12–16 mm, irregularly lobulated, lobules 0.5-5 mm long; stamen yellow, 23–25 by 7–8 mm, slightly exceeding the labellum, apex rounded, anther c. 6 mm long. *Capsule* ellipsoid to subglobose, 6–7 by 5–7mm.

Distribution — Brazil (Acre, Amazonas, Rondônia), Peru (Loreto, San Martín).

Habitat & Ecology — In non-inundated (terra firme) or periodically inundated (várzea) forests, at elevations of 0–260 m. Flowering all year round.

IUCN Conservation Status — Not evaluated. There are 28 collection records examined of which 17 specimens were collected within the past 50 years. This species is also reported in 4 recent iNaturalist observations (https://www.inaturalist.org/observations?place_id=any&taxon_id=1460765) with a distribution limited to the Brazil-Peru border region of the amazon ecoregion. Population and threat data is not available but based on the number of collection records, this species would probably score a status of Near Threatened or Vulnerable.

Vernacular names —Brazil: Canafiche.

Notes — *Costus prancei* looks quite similar to *C. sprucei* both sharing most of the vegetative and floral characters, but *C. prancei* has yellow to orange flowers whereas the flowers of *C. sprucei* are pinkish red. *Costus sprucei* and *C. prancei* closely resemble a third species, *C. chartaceus,* which occurs in Amazonian Colombia, Ecuador, and Peru. They share many of the vegetative and floral features, but *C. chartaceus* mostly has an unequally lobed ligule of 5–15 mm long, while that of *C. sprucei* and *C. prancei* is truncate and 2–7 mm long. The essential difference can be found in the flower colour, which is pinkish white in *C. chartaceus* (versus pinkish red in *C. sprucei* and yellow to orange in *C. prancei*).

Two Peruvian collections (*Gentry et al. 52203* and *Uliana et al. 1342*) look quite similar to this species and agree in most of the floral measurements but the leaves are quite large for this species (25–35 by 9–13 cm).

Additional field collections and phylogenetic work is needed to verify if the diffences between the above discussed three species, in particular between *C. chartaceus* and *C. sprucei*, are indeed sufficient for recognition as separate species.

**56. *Costus pseudospiralis*** Maas & H. Maas —Fig. 12b; Map 38

*Costus pseudospiralis* Maas & H. Maas in Maas et al. (2023) 114, f. 18. — Type: *Maas et al. 8751* (holo U, 2 sheets [U1610003, U1610004]; iso USZ), Bolivia, Beni, km 4 of road from Riberalta to Santa María, 150 m, 22 Jan. 1999.

Herbs, 1.5–2 m tall. *Leaves*: sheaths 10–15 mm diam; ligule obliquely truncate, 5–8 mm long; petiole 3–5 mm long; sheaths, ligule and petiole densely villose; lamina narrowly elliptic, 20–43 by 4–12 cm, upper side densely to rather densely villose, lower side densely villose, base cordate, apex acuminate (acumen 10–15 mm long). *Inflorescence* ovoid, 4–12 by 3–6 cm, terminating a leafy shoot; bracts, bracteole, calyx, ovary, and capsule glabrous*. Flowers* abaxially orientated; bracts red to dark red, coriaceous, broadly ovate, 2.5–3.5 by 2.5–3.5 cm, apex obtuse, callus 5–7 mm long, sometimes with a small, deltate appendage c. 5 by 5 mm; bracteole boat-shaped, 15–20 mm long; calyx red, 8–11 mm long, lobes shallowly ovate-triangular to very shallowly ovate-triangular deltate, 1–2 mm long; corolla dark salmon pink, 45– 50 mm long, glabrous, lobes narrowly obovate-elliptic, 30–35 mm long; labellum pale salmon pink, apex and center white, upper part of middle lobe yellowish, lateral lobes rolled inwards and forming a slightly curved tube 6–12 mm diam, oblong-obovate when spread out, 30–35 by 20–25 mm, apex irregularly 5-lobulate; stamen salmon pink, c. 30 by 7 mm, not exceeding the labellum, apex white, rounded, anther 8–10 mm long. *Capsule* ellipsoid, 10–20 mm long.

Distribution — Bolivia (Beni), Brazil (Rondônia).

Habitat & Ecology — In secondary roadside vegetation, at elevations of 150–230 m. Flowering all year round.

IUCN Conservation Status — Not evaluated. There are only 8 collection records examined for this recently described species, found only in western Brazil and adjacent Bolivia. This species is also reported in 7 recent iNaturalist observations (https://www.inaturalist.org/observations?place_id=any&taxon_id=1460767). Our field observations indicate that this species is quite common within its range and that includes protected areas of western Brazil. Population and threat data is not available but based on the number of collection records and our general field observations, this species would probably score a status of Least Concern.

Notes — *Costus psuedospiralis* looks superficially quite similar to *C. spiralis*, but differs by the orientation of the flowers (floral opening facing abaxially versus adaxially) and a cordate leaf base. It differs from *C. douglasdalyi* by the inflorescence terminating a leafy shoot instead of terminating a shorter leafless shoot (referred to as basal in the literature), the presence of a villose indument on most of its vegetative parts (versus glabrous or sparsely puberulous), and a cordate (versus acute to rounded) leaf base.

**57. *Costus pulverulentus*** C.Presl — Fig. 12c–e; Map 39

*Costus pulverulentus* C.Presl (1827) 111. — Lectotype (designated here): *Haenke s.n.* (lecto PR [PR495638]), Nicaragua (“Mexico”), locality unknown, anno 1790. See the note about the typification.

*Costus ruber* C.Wright ex Griseb. (1866) 256. — Lectotype (designated here): *Wright 1514* (lecto GOET [GOET011647]; isolecto BM [BM000923830], F [F0047173], G [G00168417, G00168418], GH [GH00030658], K [K000586741, K000586742, K000586743], LE, MO [MO-202812], NY [NY73336], P [P00686611]), Cuba, Monte Verde, 20 Jul. 1859.

*Costus laxus* Petersen (1890) 56. — Lectotype (designated here): *Oersted s.n.* (lecto C), Costa Rica, without location, Nov. 1847.

*Costus sanguineus* Donn.Sm. (1901) 122. — Lectotype (designated here): *Von Tuerckheim 7686* (lecto US [US00092976]; isolecto GH [GH00030648], K [K000586733], LE [LE00001318], M0 [MO-242451], NY [NY00320328], US [US00338338]), Guatemala, Alta Verapaz, Cubilguïtz, 350 m, May 1900.

*Costus formosus* C.V.Morton (1937) 305. — Type: *Skutch 2775* (holo US [US00092959]; iso GH [GH00030649], K [K000586734], MICH [MICH1192172], MO [MO-129050], NY [NY00320321], S [S-R-1256]), Costa Rica, San José, El General, 850 m, July 1936.

Herbs, 0.5–2.5(–3.5) m tall. *Leaves*: sheaths 5–15 mm diam; ligule truncate or slightly 2-lobed, 3–15(–25) mm long; petiole 2–10 mm long; sheaths, ligule and petiole glabrous, rarely densely puberulous or villose; lamina narrowly elliptic to narrowly obovate, (7–)10–30 by (3.5–)6–12 cm, upper side glabrous or rarely densely villose, primary vein densely covered with a row of erect hairs, lower side glabrous, rarely densely puberulous to villose, base acute to rounded, rarely cordate, apex acuminate (acumen 5–15 mm long). *Inflorescence* fusiform, 3–7 by 1.5–4.5 cm, enlarging to c. 20 by 6 cm in fruit, apex usually distinctly pointed, terminating a leafy shoot; bracts and bracteole sparsely puberulous, sericeous or glabrous, calyx, ovary, and capsule glabrous*. Flowers* abaxially orientated to erect; bracts red, orange-red, occasionally dark green or yellowish green, coriaceous, broadly ovate-triangular, 2.5–4.5 by 2.5–4.5 cm, apex acute, margin falling apart into fibers, but often becoming glabrous with age, callus 5–10 mm long; bracteole boat-shaped, 18–23 mm long, margin usually falling apart into fibers; calyx reddish, 6–10 mm long, lobes deltate, 1–3 mm long, margin often falling apart into fibers; corolla red to yellow, 50–70 mm long, glabrous, lobes often recurved, narrowly obovate, 40–50 mm long; labellum yellow, red, rarely white or pink, lateral lobes rolled inwards and forming a slightly curved tube c. 8 mm diam, broadly oblong-obovate when spread out, 30–40 by 30–40 mm, 5-lobulate, lobules recurved, narrowly triangular to deltate, 2–6 mm long; stamen red, 35–50 by 4–10 mm, far exceeding the labellum, apex acute to obtuse, anther 4–8 mm long. *Capsule* ellipsoid, 10–15 mm long.

Distribution — Mexico (Chiapas, Oaxaca, Puebla, San Luis Potosí, Tabasco, Veracruz), Cuba, Guatemala, El Salvador, Belize, Honduras, Nicaragua, Costa Rica, Panama, Colombia, Venezuela, Ecuador.

Habitat & Ecology — In undergrowth of moist, dense forests, in clearings and along creeks, in plantations and wooded ravines, at elevations of 0–1600 m. Flowering all year round.

IUCN Conservation Status — Not evaluated. There are 1,121 collection records examined of which 705 specimens were collected within the past 50 years. This species is also reported in 709 recent “Research Grade” iNaturalist observations (https://www.inaturalist.org/observations?place_id=any&taxon_id=276566). The species is widely distributed from Mexico to Ecuador. Population and threat data is not available but based on the number of collection records and our general field observations, this species would certainly score a status of Least Concern.

Vernacular names — Belize: Caño de Cristo, Ch’-uun (Maya, Kekchí), Spiral ginger. Colombia: Cañagria, Cañaguate, Rollinoginai (Witoto). Costa Rica: Cañagria, Caña agria. Cuba: Cañuela santa. Ecuador: Cago-lachi-tapé (Cayapá), Caña agria, Caña brava, San Juanillo. Guatemala: Pacuite, Pak ku stet, Platanillo macho, Platanillo xib’al (Itza, Maya). Honduras: Caña de Cristo. Mexico: Apagafuego, Cañagria, Caña ágria, Chilalaga, Choschogo, Cimmaróna, Flor de laguna, Flor de lantén. Nicaragua: Cañagria, Caña agria. Panama: Yar binwe (Kuna).

Notes — *Costus pulverulentus*, one of the most common and widespread species in the genus, is easily recognizable by the fusiform, pointed inflorescence with bracts of which the margin is often falling apart into fibers and by tubular flowers in which the stamen is always far exceeding the labellum. Moreover, the corolla lobes are often recurved.

*Costus pulverulentus* is a highly variable species, particularly with respect to vegetative characters like indument and leaf shape. As the variation of both characters is rather continuous and showing many transitional forms, they are not used for delimitating taxa at any level. Also, the plants in Mesoamerica do not have the fully exposed thecae or the recurved corolla lobes as found in the plants in Central America and western South America. Recent molecular phylogenetic analyses indicate that the Mesoamerican populations may form a distinct lineage; further work is necessary to determine if separation of the observed morphological variation into different taxonomic species is required.

The type of this species was collected in 1790 by the Czech botanist Thaddeus Haenke during his voyage from Panama to Acapulco, Mexico. During this voyage Haenke stopped at Realejo in present-day Nicaragua and made an expedition collecting plants in the region of Volcán Viejo. All plants collected in this part of his journey were tagged as being from Mexico, but it is likely that his collection was actually made in Nicaragua.

**58. *Costus ricus*** Maas & H.Maas — Fig. 12f; Map 40

*Costus ricus* Maas & H.Maas (1997) 274. — Type: *Maas et al. 7867* (holo U, 2 sheets [U0006984, U0006985]; iso CR, K [K000586738]), Costa Rica, Puntarenas, Cantón de Sierpe, W of Rancho Quemado on road to Drake and new logging road, along dry creek in high forest, 300 m, 7 Feb. 1991.

Herbs, 0.5–2 m tall. *Leaves*: sheaths 7–15 mm diam; ligule unequally and bluntly 2-lobed, 15–30 mm long; petiole 0–5 mm long; sheaths, ligule and petiole densely to sparsely puberulous, often villose towards the ligule and petiole; lamina narrowly obovate to narrowly elliptic, 24–44 by 6– 12 cm, upper side shiny, glabrous, rarely sparsely puberulous to villose, margin densely villose, lower side densely to sparsely puberulous, rarely rather densely villose, base acute to rounded, apex long-acuminate (acumen (10–)25–55 mm long). *Inflorescence* globose to broadly ellipsoid, 4.5–5 by 4–5 cm, enlarging to c. 17 by 6 cm in fruit, terminating a leafy shoot; bracts, appendages of bracts, bracteole, calyx, ovary, and capsule densely puberulous*. Flowers* abaxially orientated to erect; bracts dark red to dark greenish red, chartaceous, broadly elliptic, 1.4–2.5 by 0.7–2.5 cm, callus 2–4 mm long; appendages dark red to dark greenish red, patent to reflexed, concave, shallowly to very shallowly ovate-triangular, 0.7–1.5 by 0.8–2.7 cm, apex acute; bracteole 2-keeled, (12–)18–26 mm long; calyx red, (12–)17–20 mm long, lobes very shallowly triangular to delate, 2–8 mm long; corolla basally white, middle part red to orange, apex yellow, 40–50 mm long, densely puberulous, lobes narrowly elliptic to elliptic, 32–35 mm long; labellum orange to red, lateral lobes rolled inwards and forming a slightly curved tube 7–10 mm diam, elliptic when spread out, (20–)30–35 by 15–20 mm, 5-lobulate, lobules dark red, shallowly triangular to triangular, 4–5.5 mm long; stamen white, 30–35 by 8 mm, not exceeding the labellum, apex dark red, acute, anther c. 8 mm long. *Capsul*e obovoid, 10–13 mm long.

Distribution — Pacific coastal Costa Rica.

Habitat & Ecology — In undisturbed forests often on steep slopes, at elevations of 0– 1200 m. Flowering only during the dry season from late December to May.

IUCN Conservation Status — Least Concern.

Vernacular names — Costa Rica: Caña agria.

Notes — *Costus ricus* is characterized by a rather long unequally and bluntly 2-lobed ligule 15–30 mm long, an inflorescence composed of reddish bracts, with more or less patent to concave appendages, a hairy corolla, and an orange to red, tubular labellum. A peculiar feature, very rarely seen in other species of *Costus*, is the presence of a 2-keeled bracteole. This is one of the few species of *Costus* that flowers exclusively in the dry season which occurs in late December until May in Pacific coastal Costa Rica.

**59. *Costus rubineus*** D.Skinner & Maas — Fig. 12g; Map 40

*Costus rubineus* D.Skinner & Maas in Maas et al. (2023) 115, f. 19. — Type: *Rojas G. et al. 3574* (holo L [L4196133]; iso HUT, MO [MO-2152325], USM), Peru, Pasco, Prov. Oxapampa, Gramazú, 1961 m, 17 March 2005.

Herbs, 0.5–4 m tall. *Leaves*: sheaths 15–30 mm diam; ligule truncate to slightly lobed, 2–7 mm long; petiole 3–8 mm long; sheaths, ligule and petiole densely sericeous to glabrous; lamina narrowly elliptic, 20–37 by 5–13 cm, upper side glabrous, lower side rather densely villose to almost glabrous, base obtuse to slightly cordate, apex acuminate (acumen 10–15 mm long) to acute. *Inflorescence* ovoid, 7–11 by 4–6 cm, terminating a leafy shoot; bracts, bracteole, and calyx rather densely puberulous to glabrous, ovary and capsule densely sericeous*. Flowers* abaxially orientated; bracts dark ruby red to red, coriaceous, broadly ovate, 3–4 by 2.5–4 cm, apex obtuse, callus 6–10 mm long; bracteole boat-shaped, 20–28 mm long; calyx red to pink, 9– 15 mm long, lobes very shallowly triangular, 2–3 mm long; corolla pink to orange, 50–60 mm long, glabrous, lobes narrowly elliptic, 40–50 mm long; labellum yellow, upper part red, lateral lobes rolled inwards and forming a curved tube c. 10 mm diam, oblong-obovate when spread out, 30–50 by 18–20 mm, 5-lobulate, lobules narrowly ovate-triangular, c. 4 mm long; stamen yellow, 30–45 by 7–10 mm, not or slightly exceeding the labellum, apex red, irregularly dentate, anther 8–9 mm long *Capsule* obovoid, c. 13 mm long.

Distribution — Peru (Huánuco, Pasco).

Habitat & Ecology — In primary or secondary, premontane or montane forests, at elevations of (300–)1400–2200 m. Flowering all year round.

IUCN Conservation Status — Not evaluated. There are 13 collection records examined for this recently described species, which is found only in mountainous parts of San Martín, Huanuco and Pasco, Peru. This species is also reported in 10 recent iNaturalist observations (https://www.inaturalist.org/observations?place_id=any&taxon_id=1460778). Our field observations indicate that this is a fairly common species within its range and mountainous habitat, with at least one large population in a protected area. This new species would probably score as Near Threatened or Vulnerable in a full assessment.

Notes — *Costus rubineus* is a species mostly restricted to relatively high elevations (except for *Foster 8950 A* found at 300 m). It looks at first glance somewhat similar to *C. spiralis*, but it differs from that species by its abaxially orientated flowers and a very short ligule. Living plants also look somewhat similar to *C. scaber*, but differ from that species by a larger calyx (9–15 versus 3–7 mm), larger bracteole (20–28 versus 9–12 mm), larger corolla (50–60 versus 35–40 mm), and a labellum which is distinctly lobulate at the apex versus entire in *C. scaber*; furthermore it is restricted to high elevations, whereas *C. scaber* is mostly a lowland species; *C. rubineus* also lacks the row of hairs on the upper side of the lamina, which is so typical for *C. scaber*. There are often several flowers simultanously at anthesis, whereas in *C. scaber* there is usually only one flower fully open at a time.

**60. *Costus scaber*** Ruiz & Pav. — Fig. 12h, 13; Map 41

*Costus scaber* Ruiz & Pav. (1798) 2, t. 3. — Lectotype (designated here): *Ruiz López & Pavón s.n.* (lecto BM, fragment of the inflorescence, herewith designated; the MA specimen MA00810655 is *C. laevis*), Peru, Huánuco, anno 1787.

*Costus anachiri* Jacq. (1809) 55, t. 78. — [*Costus quintus* Roem. & Schult. (1817) 26, nom. nud.] — *Costus spicatus* (Jacq.) Sw. β *anthocono purpurascente*… Horan. (1862) 37, nom inval. — *Costus cylindricus* Jacq. var. *anachiri* (Jacq.) Petersen (1890) 54. — Lectotype (designated here): Plumier msc. 5: plate 31 (lecto P), based on material collected in St. Vincent.

*Costus ciliatus* Miq. (1844) 73. — *Costus cylindricus* Jacq. var. *ciliatus* (Miq.) Petersen (1890) 54. — Lectotype (designated here): *Focke 123* (lecto U [U0007218]), Suriname.

*Costus scaberulus* Rich. ex Gagnep. (1902) 99. — Lectotype (designated here): *L.C. Richard s.n.* (lecto P [P00686619]), French Guiana, Gabriel.

*Costus nutans* K.Schum. (1904) 407. — Type: *Hoffmann 727* (holo B, destroyed; photograph in F [F0B009906]), Costa Rica, Aguacate, Sept. 1857.

*Costus puchucupango* J.F.Macbr. (1931) 49. — Type: *L. Williams 4570* (holo F [F004153F]), Peru, Loreto, Lower Río Huallaga, San Ramón, Yurimaguas, 155–200 m, 4 Nov. 1929.

*Costus tatei* Rusby (1934) 51. — Type: *Tate 523* (holo NY [NY00320354]), Bolivia, La Paz, Río Chimate, 1900 ft, 1–14 Apr. 1936.

*Costus flammulus* K.M.Kay & P.Juárez in Juárez et al. (2023) 152, f. 4 & 5. — Type: P. Juárez & C. Girvin 1620 (holo US; iso CR, MO), Costa Rica, Puntarenas, Monteverde, E. B. Monteverde, Quebarada Máquina, 1524 m, 18 Aug. 2022.

*Costus cylindricus* auct. non Jacq.: Roscoe (1825 ‘1828’) t. 78.

*Costus spicatus* auct. non Jacq.: Standl. (1928) 118, pl. 15.

*Costus igneus* auct. non N.E.Br.: K.Goebel (1931) 161, f. 142.

Herbs, 0.5–3 m tall. *Leaves*: sheaths 5–15(–20) mm diam; ligule obliquely truncate, 2–12(–15) mm long; petiole 2–10 mm long; sheaths, ligule and petiole glabrous to densely puberulous, rarely villose; lamina narrowly elliptic, 10–32 by 3–11 cm, upper side glabrous to sparsely puberulous, primary vein densely covered with a row of erect hairs, rarely glabrous, lower side glabrous to rather densely puberulous, base acute to rounded, rarely cordate, apex acuminate (acumen 5–30 mm long). *Inflorescence* ovoid to narrowly cylindric, 4–10 by 1.5–3.5 cm, enlarging to c. 27 by 5 cm in fruit, terminating a leafy shoot; bracts, bracteole, calyx, ovary, and capsule glabrous to densely puberulous, bracts rarely densely villose*. Flowers* abaxially orientated; bracts orange-red to red, rarely green, coriaceous, broadly ovate, 2–3.5 by 2–3.5 cm, apex obtuse, callus 2–10 mm long; bracteole boat-shaped, 9–12(–17) mm long; calyx reddish, 3– 7 mm long, lobes very shallowly triangular, c. 1 mm long; corolla orange-red to yellow, 35–40 mm long, glabrous, lobes narrowly obovate, c. 30 mm long; labellum yellow, lateral lobes rolled inwards and forming a slightly curved tube 5–10 mm diam, oblong-obovate when spread out, 20–30 by 15–20 mm, irregularly dentate; stamen red, orange-red, or yellow, 20–25 by 6–8 mm, slightly exceeding the labellum, apex yellow to red, obtuse, anther 5–7 mm long. *Capsule* ellipsoid to subglobose, 7–12 mm diam.

Distribution — Mexico (Chiapas, Veracruz), The Greater Antilles, The Lesser Antilles (Grenada, St. Vincent and the Grenadines), Central America (Guatemala, Honduras, Belize, Nicaragua, Costa Rica, Panama), Colombia, Trinidad and Tobago, Venezuela, Guyana, Suriname, French Guiana, Ecuador, Peru, Bolivia, Brazil.

Habitat & Ecology — Rather common in clearings and along forest edges, or banks of streams, at elevations of 0–800(–1800) m. Flowering all year round.

IUCN Conservation Status — Not evaluated. There are 1,740 collection records examined of which 1,187 specimens were collected within the past 50 years. This species is also reported in 503 recent “Research Grade” iNaturalist observations (https://www.inaturalist.org/observations?place_id=any&taxon_id=283424). The species is distributed over a wide range from Mexico to Bolivia and east into Brazil and the Guianas. Population and threat data is not available but based on the number of collection records and our general field observations, this species would certainly score a status of Least Concern.

Vernacular names — Bolivia: Blua (Chimane), Cañagria, Caña agria, Caña agria de flor naranja, Chupete, Quiatime (Siriono), Yushashta (Yuracare). Brazil: Banana de macaco, Cañagria, Cana de los Indios, Cana ficha, Cana de macaco, Canarana, Consolha de viuva, Odolokakana (Paumaris), Orelha de onca, Orelha de anta, Orelha de macaco. Colombia: Caña, Caña agria, Caña agria morada, Caña fuerte, Caña selvatica, Cañagria, Cañagria de monte, Cañaguate, Caño unguyá, Flor candelabro, Jebuyae (Ticuna), Jee buyae (Ticuna), Kaguiba (Miranda), Ñapanga, Rollino ginai (Witoto), Sĕ-dyá (Makuna), Tuonoguiyo (Huitoto). Costa Rica: Caña de Cristo. Ecuador: Caña, Gumenkimonkagui (Huaorani), Odengimonkagi (Geke), Toro quilon (Quichua), Untuntup (Shuar). French Guiana: Canne congo (Créole), Canne de l’eau (Créole), Gaan singaafu (Boni), Kan cõgo (Créole), Kapiya e (Wayãpi), Kapiya e’e (Wayãpi), Kapiya-pila (Wayãpi), Maikaman (Wayana), Maya-kongo, Sãga afu (Boni), Scalarima (Wayana), Sinegafou (Surinam), Tsilulu (Emerillon), Wanayecou (Wayana). Guyana: Cana laina, Congo cane, Eseyundu (Carib), Red congo cane. Honduras: Caña de Cristo. Peru: Campanciode sigkusuk, Caña agre rojo, Caña agria, Canagre colorado, Caño agrio, Cavu, Karo, Papelillo sacha huiro, Puchucu panga, Puntuntup, Sachahuiro, Sacha huiro, Sacha huiro colorado, Sachahuiro macho, Sacha huiro negro, Sinkusuk, Siwanúk, Uchi shiwamuk, Uci siwanuk, Untúntu, Untúntup. Suriname: Fico fico, Sangaafu (Saramaccan), Sangrafoe, Sangrafoe swiejoeroe (Carib), She-we-ru-e-mah (Tirio). Singafuu (Aucan). Trinidad: Canne rivière. Venezuela: Cabu-simoro, Cana de la India, Caña de la India, Caña la India, Kabishimoro (Winikina Warao), Kabisimoru (Winikina Warao), Mata de la India, Prucunuma (Yanomami).

Notes — *Costus scaber* is the most common species of the genus *Costus* and occurs all over the Neotropics. It is well marked by the primary vein of the upper side of the lamina, which is almost always covered with a row of erect hairs, a cylindric, often orange-red inflorescence, and small orange-red to yellow, tubular flowers. Another feature is the very small calyx of 3–7 mm long.

Although *C. scaber* can very well be recognized it shows quite a geographical variation in the field as was noticed by Dave Skinner. Additional studies on this aspect are needed in the near future. *Costus scaber* probably hybridizes with *C. pulverulentus* and may form hybrid species swarms in areas where this happens frequently. A recently proposed new species, *C. flammulus*, may have resulted from such hybridization and is currently occupying a range that overlaps with that of *C. wilsonii* in Monteverde, Blasa and Tilarán, Costa Rica. Upon further examination of specimens fitting the description of *C. flammulus*, we believe they should be placed instead in *Costus scaber* based on being indistinguishable from this species despite its reported phylogenetic proximity to *C. pulverulentus*: this includes *Burger 9720*, *9656*; *Caravjal Ugalde 191*; *Grayum & Garcia 12580*; *Grayum & Rojas 12835*; *Feinsinger 408*, *Haber 10798*; *Haber & Bello 2027*, *2725*, *3161*; *Hammel & Kennedy 21078*; *Maas 1218*; *Weston 5064*.

A few collections from Amazonian Ecuador and Peru are aberrant in having very narrow, almost linear leaves (*Croat 19923* (MO, U), *Pitman 247* (MO, QCNE, U), *Plowman 2581* (GH, K, U, USM), and *Wallnöfer & Aichinger 14-41287* (U, USM)).

**61. *Costus sepacuitensis*** Rowlee — Fig. 12i; Map 42

*Costus sepacuitensis* Rowlee (1922) 286, pl. 13. — Type: *O.F. Cook & Griggs 596* (holo US [US408305]), Guatemala, Alta Verapaz, Finca Sepacuité, c. 1000 m, 13 Apr. 1902.

Herbs, (0.5–)2.5–5 m tall. *Leaves*: sheaths 8–18 mm diam, green; ligule truncate to obliquely truncate, 15–22 mm long; petiole 10–30 mm long; sheaths, ligule and petiole densely villose; lamina narrowly elliptic, 27–43 by 7–13.5 cm, upper side sparsely to densely villose, lower side densely villose, base acute, apex acuminate (acumen 5–15 mm long). *Inflorescence* cylindric to ovoid, 7–13 by 3–5 cm, terminating a separate leafless shoot 20–30 cm long, sheaths obliquely truncate, 2.5–5 cm long, densely villose to glabrous (in the type collection); bracts, bracteoles, calyx, and ovary densely to rather densely villose*. Flowers* abaxially orientated to erect; bracts green, margin reddish, coriaceous, broadly to depressed obovate, 2.5-5 by 2.5–4 cm, apex obtuse, callus c. 4 mm long; bracteole boat-shaped, 25–30 mm long; calyx dark red or pale green to whitish, 11–15 mm long, lobes deltate to shallowly triangular, 3–8 mm long; corolla pale yellow to white, c. 50 mm long, glabrous, lobes narrowly elliptic, 30–40 mm long; labellum pale yellow, upper part horizontally spreading, broadly obovate, 45–50 by 60–70 mm, lateral lobes and throat striped with red, middle lobe recurved with a yellow honey mark, margin irregularly lobed; stamen white, 25–30 by c. 10 mm, not exceeding the labellum, apex yellow and red, irregularly lobed, anther 6–7 mm long. *Capsule* ellipsoid, 15–20 mm long.

Distribution — Mexico (Oaxaca), Guatemala (Alta Verapaz), Honduras (Yoro, Malacatón).

Habitat & Ecology — In steep, forested slopes or disturbed areas near coffee fields, at elevations of (600–)1000–1400 m. Flowering in the rainy season.

IUCN Conservation Status — Not evaluated. There are only 7 collection records examined for this species, which is known only in single localities in Oaxaca, Mexico, Alta Verapaz, Guatemala and Yoró, Honduras. The only field observations posted to iNaturalist to date are made by the authors of this manuscript and visits at these sites found the species to be very rare with only a single subpopulation or plant in each locality. These populations were not in protected areas and probably this species would score as Critically Endangered if a full assessment was completed.

Notes — *Costus sepacuitensis* is different from any Central American species of *Costus* by a unique combination of the inflorescence with unappendaged bracts emerging on a separate leafless shoot, and the villose indument of the vegetative parts. It might be confused with *C. dirzoi*, a species only known from Veracruz in Mexico, but that species has a much shorter ligule (3–12 versus 15–22 mm long) and a velutinous to puberulous instead of villose indument of its vegetative parts. Moreover, *C. sepacuitensis* is much taller than *C. dirzoi* and has an inflorescence on a completely leafless shoot, whereas *C. dirzoi* usually has a few leaves around the base of the inflorescence.

**62. *Costus sinningiiflorus*** Rusby — Fig. 14a, b; Map 42

*Costus sinningiiflorus* Rusby (1927) 219 (‘*sinningiaeflorus*’). — Lectotype (designated here): *Cárdenas Hermosa 1851* (holo NY [NY00320353]), Bolivia, Beni, Rurrenabaque, 1000 ft, 26 Nov. 1921.

Herbs, 1–5 m tall. *Leaves*: sheaths 5–25 mm diam; ligule truncate to obliquely truncate, 5–10 mm long; petiole 5–20 mm long; sheaths, ligule and petiole densely brownish villose, rarely sparsely so to almost glabrous; lamina narrowly elliptic to narrowly obovate, 15–55 by 4–18 cm, upper side densely brownish villose to almost glabrous, smooth to the touch, lower side densely brownish villose, rarely sparsely so, base acute to rounded, rarely cordate, apex acuminate (acumen 10–20 mm long) to acute. *Inflorescence* ovoid to cylindric, 5–20 by 4–8 cm, erect, terminating a leafy shoot or terminating a separate leafless shoot 35–65 cm long, sheaths obliquely truncate, 3–9 cm long, glabrous, sometimes sparsely to densely brownish villose; bracts sparsely puberulous, villose, or glabrous, appendages of bracts, bracteole, and calyx sparsely, rarely densely puberulous, or glabrous, ovary and capsule rather densely puberulous to sericeous*. Flowers* abaxially orientated; bracts red, coriaceous, broadly ovate to ovate, 3–5 by 2– 3 cm, callus mostly inconspicuous to c. 5 mm long; appendages green, foliaceous, ascending, rarely reflexed, broadly ovate-triangular to ovate-triangular, often provided with a longitudinal impression over their whole length, 1–3 by 1–3 cm, base often slightly cordate and convex; bracteole boat-shaped, 22–30 mm long; calyx red, 10–18 mm long, lobes triangular to shallowly triangular, 2–4 mm long; corolla pink to white, 80–90 mm long, glabrous, lobes narrowly elliptic, 60–70 mm long; labellum white, yellow, or pink, upper part horizontally spreading, broadly obovate, 80–100 by 45–70 mm, lateral lobes striped with pink or red, middle lobe recurved with a yellow honey mark, margin crenate; stamen dark red, 50–60 by 10–15 mm, not exceeding the labellum, apex dentate, anther 10–12 mm long. *Capsule* ellipsoid, 15–25 mm long.

Distribution — Costa Rica, Panama, Colombia (Amazonas, Caquetá), Ecuador (Napo), Peru (Ayacucho, Cuzco, Huánuco, Junín, Loreto, Madre de Dios, Pasco, Puno, San Martín, Ucayali), Bolivia (Beni, Cochabamba, La Paz, Santa Cruz), Brazil (Acre).

Habitat & Ecology — In non-inundated forests, riverbank forests, or swamps, at elevations of 0–1500 m. Flowering all year round.

IUCN Conservation Status — Not evaluated. There are 126 collection records examined of which 100 specimens were collected within the past 50 years. This is the species formerly classified as *Costus guanaiensis*, which has been reported in 289 recent iNaturalist observations, and from which *C. sinningiiflorus* was split off as a separate species. The species is mostly concentrated in Bolivia, Peru and western Brazil with a few records from other countries currently attributed to this species. Population and threat data is not available but based on the number of collection records and our general field observations, this species would probably score a status of Least Concern.

Vernacular names — Bolivia: Bu’u (Chimane). Peru: Caña agria del altura, Caña agre blanco, Cañagre, Cañagria, Chacia, Sachahuiro, Sacha huiro, Sacha huiro negro, Sachahuiro pardo, Sacha huiro pardo, Sacha wiro.

Notes — *Costus sinningiiflorus* was wrongly put by Maas (1972) in the synonymy of *C. guanaiensis* (see under that species). *Costus sinningiiflorus* is very distinct by the presence of brown, erect hairs (villose) all over the vegetative parts of the plant, combined with greenish appendaged bracts, which are, moreover, often provided with a longitudinal impression over their whole length.

Various forms can be distinguished: the typical form occurs in western South America (Peru, Bolivia, and Brazil). A second form occurs in the Iquitos region of Peru sharing most of the features of the typical form, but differing by the bract appendages, which have a crinkly appearance. A third form occurring in the Peruvian department of Huánuco (the Tingo Maria region) almost always produces its inflorescence on a separate leafless shoot.

The species also occurs in Central America (mainly in Costa Rica, the Tapantí region). Material from that region differs by closely appressed appendages, which are creased or ruffled.

*Costus sinningiiflorus* needs much additional study in order to investigate if more species are involved.

**63. *Costus spicatus*** (Jacq.) Sw. — Fig. 14c; Map 42

—

*Costus spicatus* (Jacq.) Sw. (1788) 11. — *Alpinia spicata* Jacq. (1760) t. 1. — *Amomum petiolatum* Lam. (1783) 136, nom. illeg., superfl. — *Costus conicus* Stokes (1812) 75, nom. illeg., superfl. — Lectotype (designated by Maas 1972): Jacquin (1763) t. 1 (the original material collected by Jacquin in Martinique (“parochia St. Maria, Cabesterre”) was destroyed in the Vienna Herbarium (W) in 1943).

*Costus cylindricus* Jacq. (1809) 54, t. 77. — *Costus quartus* Roem. & Schult. (1817) 26, nom. nud. — Lectotype (designated here): Plumier (msc.) Pl. 30 (lecto P), based on material collected in Martinique.

*Costus micranthus* Gagnep. (1903) 586. — Type: *Hort. Bot. Paris, Oct. 1903* (holo P [P00686621]; iso BM [BM000923829], K [K000586740]), probably originating from Martinique.

*Costus arabicus* auct. non L.: Aubl. (1775) 2.

Herbs, 1–3 m tall. *Leaves*: sheaths 10–20 mm; ligule truncate, 2–10 mm long; petiole 2–12 mm long; sheaths, ligule and petiole rather densely puberulous to glabrous; lamina narrowly elliptic to narrowly obovate, 7–33 by 3–9(–12.5) cm, upper and lower side subglabrous, but primary vein on upper side often covered with a row of erect hairs, base rounded, rarely cordate, apex acuminate (acumen 5–15 mm long). *Inflorescence* ovoid to cylindric, 5–25 by 3–5 cm, elongating to c. 35 by 6 cm in fruit, terminating a leafy shoot; bracts, bracteole, calyx, ovary, and capsule sparsely puberulous to glabrous*. Flowers* abaxially orientated; bracts greenish red to red, coriaceous, broadly ovate, 2–3 by 2–3.5 cm, apex obtuse, callus 5–10 mm long; bracteole boat-shaped, 16–30 mm long; calyx pink to red, 9–16 mm long, lobes shallowly triangular, 1–3 mm long; corolla white, yellow, pink, or red, 40–50 mm long, glabrous, lobes narrowly elliptic to narrowly obovate, 30–40 mm long; labellum yellow, white, or pink, lateral lobes often reddish striped, rolled inwards and forming a slightly curved tube c. 10 mm diam, middle lobe with recurved with a yellow honey mark, broadly oblong-obovate when spread out, 30–50 by 30–50 cm, margin irregularly lobulate to crenulate; stamen yellow, white, to pink, 30–40 by 8–10 mm, slightly exceeding the labellum, apex obtuse to acute, anther 6–8 mm long. *Capsule* ellipsoid, 10–15 mm long.

Distribution — The Greater Antilles (Dominican Republic, Guadeloupe, Martinique, Puerto Rico) and Lesser Antilles (Dominica).

Habitat & Ecology — In wet forests, swamps, or along watercourses, at elevations of 0– 500(–1200) m. Flowering in the rainy season.

IUCN Conservation Status — Not evaluated. There are 74 collection records examined of which only 30 specimens were collected within the past 50 years. There are only 8 recent “Research Grade” iNaturalist observations from within the confirmed range of this species. The species is distributed in the windward islands in Martinique, Guadeloupe, Dominica and the Dominica Republic as well as two collections in Puerto Rico. This data indicates that the species has suffered a significant decline and that would likely cause the scoring close to the level of Endangered if a full assessment was completed.

Vernacular names — Dominican Republic: Cumani, Gengibre cimarron; Gengibrillo. Martinique: Canne d’eau, La canne de rivière.

Notes — *Costus spicatus* is the only endemic *Costaceae* in the Antilles; it is apparently a rather common plant in the Dominican Republic, Guadeloupe, Dominica, and Martinique.

*Costus spicatus* could be confused with *C. arabicus*, differing however, by the shape of the labellum which is tubular in *C. spicatus* versus horizontally spreading in *C. arabicus*. Furthermore the flowers of *C. arabicus* are often pure white (except for the yellow honey mark), a colour never seen in *C. spicatus*. Moreover, the leaf base of *C. arabicus* is mostly cordate whereas that of *C. spicatus* is mostly rounded and only rarely cordate.

*Costus spicatus* has been confused in the past with *C. scaber*, but from that species it differs by a much longer calyx (9–16 versus 3–7 mm), a much larger labellum when spread out (30–50 by 30–50 mm versus 20–30 by 15–20 mm), and the presence of often reddish striped lateral lobes of the labellum, which are absent in the labellum of *C. scaber*.

**64. *Costus spiralis*** (Jacq.) Roscoe — Fig. 14d; Map 43

*Costus spiralis* (Jacq.) Roscoe (1807) 350. — *Alpinia spiralis* Jacq. (1797) 1, t. 1. — *Costus spiralis* (Jacq.) Roscoe var. *jacquinii* Griseb. (1864) 602, nom. inval., not autonym. — Lectotype (designated by Maas 1972): Jacquin (1797) t. 1 (the original material in the Vienna Herbarium (W) was destroyed in 1943, which was collected near Caracas, Venezuela). See note 2. *Costus pisonis* Lindl. (1825) t. 899. — *Costus spiralis* (Jacq.) Roscoe var. *pisonis* (Lindl.) Griseb. (1864) 602. — Lectotype (designated here): *Hesketh s.n.* (holo CGE), Brazil, Maranhão, anno 1824.

*Costus spiralis* (Jacq.) Roscoe var. *roscoei* Griseb. (1864) 602. — Type: Not indicated.

*Costus spiralis* var. *villosus* Maas (1972) 109, f. 50 d–g. — Type: *Florschütz & Maas 2690* (holo U [U0007219]; iso F, K, NY), Suriname, Heidoti, Coppename River, 26 Jan. 1965.

Herbs, 0.5–-3.5(–5) m tall. *Leaves*: sheaths 5–20 mm diam; ligule truncate, 2–10 mm long; petiole 2–17 mm long; sheaths glabrous, rarely sparsely villose, ligule and petiole glabrous to densely villose; lamina narrowly elliptic to narrowly obovate, 8–43 by 5–14 cm, upper side glabrous to sparsely villose, lower side glabrous to densely villose, base acute to rounded, rarely cordate, apex acuminate (acumen 5–15 mm long). *Inflorescence* ovoid, 4–11 by 2–5 cm, enlarging to 20–40 by 6–8 cm in fruit, terminating a leafy shoot, rarely terminating a separate leafless shoot 40–60 cm long, sheaths truncate, c. 5 cm long, glabrous; bracts, bracteole, calyx, ovary, and capsule sparsely puberulous to glabrous*. Flowers* adaxially orientated to erect; bracts red, pinkish red, rarely green, coriaceous, broadly ovate, 2–4 by 2.5–4.5 cm, apex obtuse, rarely acute, callus 5–10 mm long; bracteole boat-shaped, 15–29 mm long; calyx purplish red, 7–12(– 15) mm long, lobes shallowly triangular to deltate, 2–4 mm long; corolla pinkish red to salmon-red, 45–60 mm long, glabrous, lobes narrowly obovate-elliptic, 35–50 mm long; labellum pinkish red to salmon-red, upper part of middle lobe yellowish, lateral lobes rolled inwards and forming a tube 7–10 mm diam, broadly oblong-obovate when spread out, 25–30 by 20–25 mm, apex slightly irregularly crenulate to dentate; stamen red, 25–30 by 5–10 mm, not exceeding the labellum, apex emarginate, anther 7–9 mm long. *Capsule* ellipsoid, 10–13 mm long.

Distribution —Colombia, Venezuela, Guyana, Suriname, French Guiana, Peru, Bolivia, Brazil.

Habitat & Ecology — In moist rain forests, savanna forests, or on granitic outcrops, at elevations of 0–800(–1900) m. Flowering all year round.

IUCN Conservation Status — Not evaluated. There are 636 collection records examined of which 378 specimens were collected within the past 50 years. This species is also reported in 183 recent iNaturalist observations (https://www.inaturalist.org/observations?place_id=any&taxon_id=465976). The species is widely distributed throughout the Amazonian region of South America, the Guianas and as far south as southern Brazil. Our observations indicate that this is a common species within its range and some of these areas are protected areas with no known specific threats. Population and threat data is not available but based on the number of collection records and our general field observations, this species would almost certainly score a status of Least Concern.

Vernacular names — Bolivia: Bu’a’ (Chimane), Budubd’ui (Tacana), Caña agria. Brazil: Cana brava, Cana de macaco, Cana de macao, Cana do mato, Canafiche, Canafistula, Canarana branca, Capão de mata, Jacuanga, Kani-oho. Colombia: Caña de la India, Cañagria, Cañaguate, Cañejo, Naney (Guahibo), Rollinoginai (Witoto), Timonogina. French Guiana: Canne congo (Créole), Canne olivier (Créole), Kapiya (Wayãpi), Kapiya iwa (Wayãpi), Kapiya u (Wayãpi), Kapiya yowa, (Wayãpi), Kapiyua (Wayãpi), Kapiyuwa asikalu (Wayãpi), Maikanon (Wayana), Segafu (Boni), Tapatilulu (Émerillon), Tuiu (Palikur), Tuiu awaigl (Palikur). Guyana: Ka-mar-u-mang. Suriname: Canne congo, Rode sangrafoe, Sangrafoe, Syiviulu (Carib), Tuiu (Palikur). Venezuela: Caña de India, Caña de la India, Caña del Indo, Caña salada, Prucunuma (Yanomami).

Notes — 1. *Costus spiralis* can easily be recognized in the field by a red inflorescence and pinkish red to salmon-red, adaxially orientated flowers. The last feature is rarely encountered elsewhere in the genus.

2. Grisebach (1864) distinguishes three varieties, of var. *jacquinii* we assume that it contains the type of the species as Jacquin (1797) is the author of the *C. spiralis*. Under the present code (Turland et al. 2018) Grisebach’s name is invalid as it is not the autonym.

**65. *Costus sprucei*** Maas — Fig. 14e; Map 44

*Costus sprucei* Maas (1972) 96, f. 44. — Type: *Spruce 439* (holo K [K000586759]; iso K [K000586758], P [P01740589]), Brazil, Pará, Santarém, Nov. 1849.

Herbs, to 1 m tall. *Leaves*: sheaths 5–10 mm diam; ligule truncate, 2–5(–12) mm long; petiole 2– 8 mm long; sheaths, ligule and petiole densely to sparsely villose; lamina narrowly elliptic to narrowly obovate, 6–22 by 3–7 cm, upper side glabrous to rather densely villose, primary vein densely villose, lower side densely to rather densely villose, rarely glabrous, base acute, rounded, or cordate, apex acuminate (acumen 5–15 mm long). *Inflorescence* ovoid to cylindric, 4–11 by 2–3 cm, terminating a leafy shoot; bracts, bracteole, calyx, ovary, and capsule sparsely to rather densely puberulous to villose, sometimes glabrous*. Flowers* abaxially orientated; bracts red, rarely red-orange, chartaceous to coriaceous, broadly ovate, 1.5–2.5 by 1.5–2 cm, callus 2–6 mm long; bracteole boat-shaped, 8–13 mm long; calyx pinkish white, 6–9 mm long, lobes shallowly ovate-triangular, 1–2 mm long; corolla pale pink, 25– 30 mm long, glabrous, lobes narrowly elliptic, 15–25 mm long; labellum pinkish red, lateral lobes involute and forming a straight tube 5–6 mm diam, oblong-obovate when spread out, c. 17 by 7 mm when spread out, irregularly dentate, diam 5–6 mm; stamen pinkish red, c. 15 by 3 mm, exceeding the labellum, anther c. 4 mm long. *Capsul*e globose to ellipsoid, 5–9 by 5–9 mm diam.

Distribution — Brazil (Pará).

Habitat & Ecology — In non-inundated (terra firme) forests, often on sandy soils, at elevations of 0–225 m. Flowering in the rainy season, November to January.

IUCN Conservation Status — Not evaluated. There are only 6 collection records examined; of which only 2 were collected within the past 50 years and all were collected in a small region of Pará, Brazil near Santarém. This species is only reported in 3 recent iNaturalist observations (https://www.inaturalist.org/observations?place_id=any&taxon_id=911031). Population and threat data is not available but based on the number of collection records this species is likely Endangered and might score as Critically Endangered.

Notes — *Spruce 439*, the type collection of *C. sprucei*, was collected near Santarém in the Brazilian state of Pará. Co-author on this revision Thiago André lives in the city of Santarém, and reports that *C. sprucei* is one of the few species of *Costus* that is quite common in that region. Photos of a plant growing in his garden revealed many additional characters, such as pinkish red flowers; not yellow as described by Maas (1972: 96, f. 44). After studying *C. sprucei* sensu lato and incorporating the data provided by Thiago André, we have composed an adapted description of *Costus sprucei*, a species restricted to the Brazilian state of Pará. The material with yellow to orange flowers from the states of Acre, Amazonas, and Rondônia now is included in the new species *C. prancei*.

**66. *Costus stenophyllus*** Standl. & L.O.Williams — Fig. 14f; Map 44

*Costus stenophyllus* Standl. & L.O.Williams (1952) 38. — Type: *P.H. Allen 6037* (holo US, 2 sheets [US00092982, US00092983]; iso EAP [EAP122343], F [F0047169F, F0047170F], NY), Costa Rica, Puntarenas, Esquinas Forest, 60 m, 27 March 1951.

Herbs, 1.5–4 m tall. *Leaves*: sheaths 5–15 mm diam, whitish base with reddish-brown margin; ligule obliquely truncate, 1–2 mm long; petiole 5–10 mm long; sheaths, ligule and petiole glabrous; lamina linear, 15–35 by 1–2 cm, lower side glaucous when young, upper and lower side glabrous, base acute, apex caudate. *Inflorescence* ovoid, 9–14 by 3.5–4.5 cm, terminating a separate leafless shoot 15–50 cm long, sheaths obliquely truncate, 2–5 cm long, glabrous; bracts, bracteole, calyx, ovary, and capsule glabrous*. Flowers* abaxially orientated to erect; bracts red, coriaceous, broadly ovate-triangular, 2–5 by 2–4 cm, callus 5–15 mm long; bracteole boat-shaped, 25–27 mm long; calyx red, 15–20 mm long, lobes triangular, c. 5 mm long; corolla yellow, c. 80 mm long, glabrous, lobes narrowly elliptic, c. 60 mm long; labellum yellow, lateral lobes rolled inwards and forming a tube 5–7 mm diam, oblong-ovate when spread out, 40–50 by c. 20 mm, 5-lobulate, central lobule reflexed, linear, c. 6 mm long, the other ones depressed ovate, c. 1.5 mm long; stamen yellow, 50–65 by 5–7 mm, far exceeding the labellum, apex slightly crenulate, anther 7–8 mm long. *Capsule* ellipsoid, 12–15 mm long.

Distribution — Costa Rica.

Habitat & Ecology — In lowland rain forests, at elevations of 0–400 m. Flowering in the rainy season.

IUCN Conservation Status — Least Concern.

Notes — *Costus stenophyllus*, endemic to Costa Rica, cannot be confused with any other Central American species of *Costus* because of its very narrow, linear leaves, an extremely short ligule, and reddish brown margined sheaths. It can also be compared against several other species that flower exclusively on basal leafless shoots and have stamens that far exceed the labellum, thus fully exposing the thecae (*C. alticolus*, *C. asteranthus*, and *C. geothryrsus*).

**67. *Costus vargasii*** Maas & H.Maas — Fig. 14g; Map 44

*Costus vargasii* Maas & H.Maas (1990) 311, f. 4 & 5. — Type: *Maas et al. 6156* (holo U, 3 sheets [U0007221, U0007222, U0007223]; iso F [F0040741F], K [K000586771, K000586772], MO [MO-202808], NY [NY00039320], USM), Peru, Cuzco, Prov. Paucartambo, road from Pillcopata to Salvación, 2 km before Atalaya, forest margin along road, 700–800 m, 24 Oct. 1984.

Herbs, to 1–3 m tall. *Leaves*: sheaths 5–25 mm diam, often glaucous; ligule (un)equally 2-lobed, 5–35 mm long; petiole 5–15 mm long, often reddish; sheaths, ligule and petiole glabrous; lamina narrowly obovate to narrowly elliptic, 17–42 by 4–15 cm, upper side often dull glaucous green, lower side sometimes purplish, upper and lower side glabrous, base acute to obtuse, apex acuminate (acumen 2–15 mm long). *Inflorescence* ovoid to cylindric, 4–10 by 3–4 cm, terminating a leafy shoot or terminating a separate leafless shoot 40–80 cm long, sheaths somewhat inflated in ripe stage, obliquely truncate, 4– 7 cm long, glabrous; bracts, bracteole, calyx, ovary, and capsule glabrous*. Flowers* abaxially orientated; bracts red, coriaceous, obovate to broadly obovate, 2.5–3.5 by 2–3 cm, apex obtuse, callus 4–8 mm long; bracteole boat-shaped, 12–22 mm long; calyx pink to red, 6–10 mm long, lobes very shallowly triangular, c. 1 mm long; corolla yellow, 53–55 mm long, glabrous, lobes narrowly elliptic, 48–50 mm long; labellum yellow, lateral lobes rolled inwards and forming a curved tube c. 10 mm diam, obovate when spread out, c. 35 by 18 mm, irregularly lobulate, lobules 2–4 mm long; stamen yellow with reddish margin, 30–35 by 10 mm, not exceeding the labellum, apex obtuse and slightly pointed, anther 6–8 mm long. *Capsul*e obovoid to ellipsoid, 10–15 mm long.

Distribution — Peru (Amazonas, Cuzco, Madre de Dios, Puno), Bolivia (Beni, Cochabamba, La Paz).

Habitat & Ecology — In primary or secondary, moist forests, at elevations of (250–)800– 1800 m. Flowering in the rainy season.

IUCN Conservation Status — Not evaluated. There are 38 collection records examined of which 34 specimens were collected within the past 50 years. This species is also reported in 3 recent iNaturalist observations (https://www.inaturalist.org/observations?place_id=any&taxon_id=504156). The species is distributed in northern Bolivia and southern Peru. Our observations indicate that this is a somewhat uncommon species but some of these areas are protected areas with no known specific threats. Population and threat data is not available but based on the number of collection records and our general field observations, this species may be Near Threatened.

Vernacular names — Peru: Chulco.

Notes — *Costus vargasii*, often occurring in quite high elevations of Bolivia and Peru, is characterized by red bracts, yellow flowers with a tubular, curved labellum, and the complete absence of any indument. Inflorescences can be either terminating a leafy or a leafless shoot and in the latter case the sheaths of the flowering shoot are often inflated.

**68. *Costus varzearum*** Maas — Fig. 14h; Map 45

*Costus varzearum* Maas (1976) 471. — Type: *Maas et al.* as *Prance 12860* (holo INPA [INPA31223]; iso COL [COL000000313], K [K000586756, K000586757], NY [NY00320356], U [U0008361], US [US00092985], W), Brazil, Acre, Rio Juruá, Igarapé Treize de Maio, 12 May 1971.

*Costus claviger* auct. non Benoist: Maas (1972) 77, pro parte minore.

Herbs, 0.3–1.5 m tall. *Leaves*: sheaths purplish, 10–15 mm diam; ligule 20–50 mm long, purplish, unequally 2-lobed, lobes rounded; petiole purplish, 5–10 mm long; sheaths, ligule and petiole glabrous to sparsely puberulous to villose; lamina elliptic to obovate or narrowly so, 13– 25 by 5–11 cm, lower side often red-purple, upper side glabrous, lower side glabrous to sparsely puberulous, base acute to obtuse, apex acute to acuminate (acumen 5–10 mm long). *Inflorescence* terminating a 3–7-leaved shoot, the leaves strongly congested towards the apex, ovoid, 4–10 by 3–4 cm; bracts, appendages of bracts, bracteole, calyx, ovary, and capsule rather densely to sparsely puberulous to villose*. Flowers* abaxially orientated; bracts pinkish red, coriaceous, ovate, 2–2.5 by 1–1.8 cm, callus 1–3 mm long; appendages green, foliaceous, horizontally spreading in living plants, ascending or reflexed and fragile when dry, broadly to shallowly triangular-ovate, 0.5–1.5 by 0.8–1.6 cm, apex rounded; bracteole boat-shaped, 15–20 mm long; calyx red, 7–10 mm long, lobes very shallowly triangular, 1–2 mm long; corolla yellow, orange, or pink, 45–55 mm long, glabrous, lobes narrowly elliptic, 35–40 mm long; labellum cream, upper part horizontally spreading, obovate, 40–45 by 25–35 mm, lateral lobes reddish striped, middle lobe recurved with a yellow honey mark, margin irregularly lobulate, lobules 1–10 mm long; stamen cream, 35–40 by 7–8 mm, not exceeding the labellum, apex red, irregularly dentate, anther 6–8 mm long. *Capsule* ellipsoid, c. 10 mm long.

Distribution — Peru (Loreto, Madre de Dios, Ucayali), Brazil (Acre).

Habitat & Ecology — In non-inundated or periodically inundated (várzea) forests, at elevations of 0–200 m. Flowering in the rainy season.

IUCN Conservation Status — Not evaluated. There are 16 collection records examined of which 14 specimens were collected within the past 50 years. This species is also reported in 4 recent iNaturalist observations (https://www.inaturalist.org/observations?place_id=any&taxon_id=343918). The species is distributed in southern Peru and western Brazil. Our observations indicate that this is a somewhat uncommon species with a unique habitat in seasonably flooded forests. Population and threat data is not available but based on the number of collection records and our general field observations, this species may be Vulnerable or at least Near Threatened.

Vernacular names — Brazil: Buncache (Kaxinawá).

Notes — *Costus varzearum*, a much cultivated species, is characterized by green appendaged bracts and purplish sheaths, petioles, and lower leaf sides, and a very long ligule up to 50 mm long.

1. 69. *Costus villosissimus* Jacq. — Fig. 14i; Map 46

*Costus villosissimus* Jacq. (1809) 55, t. 80. — [*Costus septimus* Aubl. ex Roem. & Schult. (1817) 26, nom. inval, as synonym] — Lectotype (designated here): Plumier, Msc. 5: Plate 34 ‘*Paco-Caatinga villosissima flore luteo*, *Zinziber villosissimum floribus luteis*’ (lecto P), based on material collected in St. Vincent.

*Costus hirsutus* C.Presl (1827) 112. — Lectotype (designated here): *Haenke s.n.* (lecto PR), Nicaragua (“Mexico”), locality unknown, anno 1790. For the notes about the typification see under *C. pulverulentus*.

*Costus friedrichsenii* Petersen (1893) 260, t. 15. — Lectotype (designated here): *Hort. Bot. Haun*. (lecto C), origin unknown, 1 Sept. 1891, infl. in spirit: 13 Sept. 1891.

Herbs, 1–4(–6) m tall. *Leaves*: sheaths 5–20 mm diam; ligule truncate, 2–10 mm long; petiole 2– 5 mm long; sheaths, ligule, and petiole densely to rather densely brownish villose, rarely glabrous; lamina narrowly elliptic to narrowly obovate, 15–30(–62) by 4–8(–14) cm, upper and lower side densely to rather densely brownish villose, rarely glabrous, base cordate to rounded, apex acuminate (acumen 5–15 mm long). *Inflorescence* ovoid, 6–10 by 3.5–6.5 cm, enlarging to c. 20 by 8.5 cm in fruit, terminating a leafy shoot; bracts and appendages of bracts densely to rather densely brownish villose, rarely glabrous, bracteole and calyx glabrous to sparsely puberulous, ovary and capsule glabrous to densely sericeous at the apex*. Flowers* abaxially orientated to erect; bracts green or red, coriaceous, broadly ovate, 2–4(–6) by 2.5–3.5(–4) cm, callus 0–6 mm long; appendages green, foliaceous, slightly recurved in living plants, ascending or reflexed when dry, narrowly triangular to deltate, 1–8(–16) by 1–3(–8) cm, apex acute; bracteole boat-shaped, 17–30 mm long; calyx white to red, 7–16 mm long, lobes deltate, 3–6 mm long; corolla yellow, rarely white, 60–80 mm long, glabrous, lobes narrowly obovate, 50–60 mm long; labellum yellow, rarely white, upper part horizontally spreading, broadly obovate, 60–90 by 45–60 mm, middle lobe recurved, margin glabrous or fimbriate; stamen yellow, 45–50 by 10– 15 mm, not exceeding the labellum, apex irregularly lobed, anther 8–12 mm long. *Capsule* ellipsoid, 15–25 mm long.

Distribution — Nicaragua, Costa Rica, Panama, the Greater Antilles (Jamaica), the Lesser Antilles (St. Vincent), Colombia (Antioquia, Boyacá, Caldas, Caquetá, Casanare, Cauca, Chocó, Cundinamarca, Guajira, Magdalena, Meta, Nariño, Santander, Valle del Cauca, Vichada), Venezuela (Aragua, Barinas, Bolívar, Distrito Federal, Portuguesa, Táchira, Yaracuy, Zulia), Guyana, French Guiana, Ecuador (Azuay, Esmeraldas, Guayas, Imbabura, Loja, Orellana).

Habitat & Ecology — In riverbank forests, foothill forests, lowland thickets, and swamps, at elevations of 0–600(–1600) m. Flowering in the rainy season.

IUCN Conservation Status — Least Concern.

Vernacular names — Colombia: Caña agria, Cañagria, Cañaguate. French Guiana: Ago singaafu (Boni), Wilapa psa (Wayãpi). Jamaica: Wild ginger. Panama: Caña agria, Caña de mico, Cañagria.

Notes — Good features to distinguish *C. villosssimus* are the brown, villose indument of the vegetative parts, a mostly cordate leaf base, combined with green-appendaged bracts and large, completely yellow flowers.

**70. *Costus vinosus*** Maas — Fig. 15a, 16a–e; Map 47

*Costus vinosus* Maas (1976) 470. — Type: *Dressler 4406* (holo U, 3 sheets [U0007887, U00078878, U0008040]; iso F [F0047171F], MO [MO-120591], PMA [PMA32416]), Panama, Colón, lower Río Guanche, 20 June 1973.

Herbs, 1.5–2 m tall. *Leaves*: sheaths purple-red, the basal leafless ones more or less inflated with inflexed upper margin, 25–80 mm diam; ligule 70–100 mm long, wine-red, unequally and very deeply 2-lobed, lobes ovate-elliptic to narrowly ovate-elliptic; petiole 20–30 mm long; sheaths glabrous, sometimes sparsely villose, ligule rather densely to sparsely villose, petiole glabrous; lamina narrowly elliptic to narrowly obovate, 38–55 by 14–18 cm, upper side dark green or reddish green, lower side purple-red to wine-red when young, changing to dull green with a red pattern when mature, upper and lower side glabrous, base acute, apex acute. *Inflorescence* ovoid, 8–13 by 4–7 cm, terminating a leafy shoot; bracts, bracteole, calyx, ovary, and capsule rather densely to densely puberulous to sericeous*. Flowers* abaxially orientated; bracts dark red, coriaceous, broadly ovate to broadly obovate, 3.5–5 by 3.5–5 cm, apex obtuse, callus 3–6 mm long; bracteole boat-shaped, 18–30 mm long; calyx pinkish white to red, 8–10 mm long, lobes very shallowly ovate-triangular, 2–3 mm long; corolla cream to pink, 75–80 mm long, glabrous, lobes narrowly elliptic, 60–70 mm long; labellum wine-red, upper part horizontally spreading, broadly obovate, c. 70 by 60 mm, lateral lobes wine-red, middle lobe with yellow-orange honey mark, margin irregularly lobulate; stamen dark red, c. 50 by 16 mm, not exceeding the labellum, apex irregularly dentate, anther 8–11 mm long. *Capsule* obovoid, 17–20 mm long.

Distribution — Panama.

Habitat & Ecology — in rocky vegetation in forest shade, at elevations of 0–250 m. Flowering only in the rainy season.

IUCN Conservation Status — Critically Endangered. More recently, Skinner has found additional subpopulations of this species and determined that it is not extinct in the wild.

Notes — *Costus vinosus*, endemic to Panama, is one of the most spectacular and very easily recognizable species by the presence of its inflated, reddish leaf sheaths, the wine-red lower side of the lamina and an extremely long ligule up to 10 cm long (!). According to recent observations of Skinner there is an enormous and quite puzzling variation, however, in its flowers. Skinner recognizes 3 forms in *C. vinosus* based on his observations of plants in the field and plants cultivated in his greenhouse.

**Form 1.** (Fig. 16a) Based on *Dressler 4406* (type) and *Maas, Dressler & Kennedy 1600*, both from Río Guanche. It has the typical leaf sheaths and wine-red lower side of the lamina; the corolla is cream (to pink) and the labellum is wine red, open and with 2 yellow-orange streaks as nectar guide.

**Form 2.** (Fig. 16b) Based on *Dressler s.n.*, also collected along Río Guanche (10 Aug 1971). It is widely cultivated at Marie Selby Botanical Garden in Sarasota, Florida (1986-0224), USBG 1989-0382, and Smithsonian Greenhouse - John Kress collection number 94-3727. This form shares the typical leaf sheaths and wine-red lower side of the lamina of Form 1, but the flowers are completely different: they are yellow, and the labellum is tubular (not open) and lacking any streaks or nectar guide.

**Form 3.** (Fig. 16c) Based on *Maas et al. 9568* collected sterile at the Santa Rita Ridge in 2004. Skinner visited the same locality in 2015 and collected seeds (Skinner & Black 2016) that were cultivated in Florida and are vouchered as living collections *R 3352* and *R 3353*. This form shares the typical leaf sheaths and wine-red lower side of the lamina of Form 1and 2. However, this form may be a hybrid as seeds germinated from the same plant produced offspring with a variety of forms: 1. plants with typical *C. vinosus* flowers (Form 1); 2. small plants with a yellow-orange corolla and a small labellum with red stripes (Fig. 16c); 3. taller plants without cup-shaped sheaths but with flowers similar to Form 2. These may be the result of cross fertilization in the wild with *C. allenii*.

**71. *Costus weberbaueri*** Loes. — Fig. 15b; Map 47

*Costus weberbaueri* Loes. (1929) 712. — Lectotype (designated by Maas 1972): *Weberbauer 1852* (holo B, destroyed; lecto G [G00168445]; isolecto USM), Peru, Junín, Prov. Tarma, Chanchamayo Valley, near La Merced, 700–1000 m, Dec. 1902.

Herbs, 2–4.5 m tall. *Leaves*: sheaths 20–25 mm diam; ligule truncate to obliquely truncate, 6–17 mm long; petiole 10–20 mm long; sheaths, ligule, and petiole glabrous; lamina narrowly elliptic to narrowly obovate, 32–43 by 9–14 cm, upper side glabrous, lower side glabrous, rarely densely puberulous, base acute, apex acuminate (acumen c. 5 mm long). *Inflorescence* globose to cylindric, 7–8 by 4.5–5 cm, enlarging to 30 by 8 cm in fruit, terminating a leafy shoot, the upper leaves forming a bowl-shaped involucrum at the base of the inflorescence; bracts, bracteole, calyx, ovary, and capsule glabrous*. Flowers* adaxially orientated; bracts green, coriaceous, broadly ovate, 30–45 by 30–50 mm, apex obtuse, callus 4–5 mm long; bracteole boat-shaped, (15–)20–28 mm long; calyx red, (5–)8–12(–15) mm long, lobes shallowly triangular, 1–3 mm long; corolla pinkish red to salmon-red, 50–60 mm long, glabrous, lobes narrowly obovate-elliptic, 35–50 mm long; labellum pinkish red to salmon-red, upper part of middle lobe yellowish, lateral lobes rolled inwards and forming a tube c. 10 mm diam, oblong-obovate when spread out, 35–40 by 15–20 mm, middle lobe with yellow apex, apex irregularly dentate; stamen whitish to whitish red, 25–40 by 8–10 mm, not exceeding the labellum, apex emarginate, anther 8–10 mm long. *Capsule* obovoid, 12–15 mm long.

Distribution — Peru (Ayacucho, Huánuco, Junín, Madre de Dios, Pasco, Puno, San Martín, Ucayali), Bolivia (Beni, Cochabamba, La Paz).

Habitat & Ecology — In primary or secondary forests, at elevations of 300–1000 m. Flowering in the rainy season.

IUCN Conservation Status — Not evaluated. There are 22 collection records of this reinstated species examined of which 7 specimens were collected within the past 50 years. This newly interpreted species is reported in 23 curated iNaturalist observations. The species is found along the eastern foothills of the Andes from northern Peru to northern Bolivia. Population and threat data is not available but based on the number of collection records and our general field observations which found this species to be very common within its range, this species would probably score a status of Least Concern.

Vernacular names — Peru: Sacha huiro colorado.

Notes — *Costus weberbaueri* was incorrectly synonymized by Maas (1972) with *C. laevis* sensu lato. It closely resembles another species, however, namely *C. spiralis*, with which it shares many features, a.o. the adaxially orientated, tubular, pinkish flowers, but differing by its green instead of reddish bracts and by the presence of a distinct, bowl-shaped involucrum formed by the leaves at the base of the inflorescence. *Costus weberbaueri* is restricted to the eastern foothills of the Peruvian and Bolivian Andes, whereas *C. spiralis* is occuring almost all over tropical South America, but is absent in the eastern part of the Andes.

**72. *Costus whiskeycola*** Maas & H.Maas — Fig.15c, 16f; PMap 48

*Costus whiskeycola* Maas & H.Maas in Maas et al. (2023) 118, f. 20. — Type: Colombia, Putumayo, Mocoa-Puerto Asis Road km 42, Quebrada El Whiskey, Finca Santa Marta, Hilltop forest, c. 400 m (1260 ft), 5 Nov. 1974, *Plowman & Davis 4396* (holo COL; iso PSO).

Herbs, 0.5–0.7 m tall. *Leaves*: sheaths 10–18 mm diam; ligule 10–20 mm long, obliquely truncate; petiole 5–15 mm long; sheaths, ligule and petiole glabrous, purple-red to green; lamina ovate-elliptic to narrowly ovate-elliptic, 22–29 by 10–13 cm, dark, olive green above, red-purple to dark purple below, with 5–6 dark green bands corresponding with slightly raised veins above, upper and lower side glabrous, base acute, apex acute to shortly acuminate (acumen 3–5 mm long). *Inflorescence* ovoid, c. 7 by 4.5 cm, terminating a leafy shoot; bracts and appendages of bracts, bracteole, calyx, and ovary glabrous*. Flowers* abaxially orientated; bracts red, coriaceous, broadly ovate, 3–4 by 2–3.5 cm; appendages green, foliaceous, ascending, triangular-ovate, 2.5– 5 by 2–3.5 cm, apex acute; bracteole boat-shaped, 24–30 mm long; calyx red, 12–20 mm long, lobes very shallowly triangular, 2–5 mm long; corolla white, 60–75 mm long, glabrous, lobes narrowly elliptic, 45–60 mm long; labellum white, upper part horizontally spreading, broadly obovate, 60–70 by 50 mm, lateral lobes striped with red, middle lobe recurved with a yellow honey mark, irregularly lobulate, margin crenulate; stamen white, tinged with red, 35–40 by 13– 14 mm, not exceeding the labellum, apex 3-dentate, anther 8–10 mm long. *Capsule* not seen.

Distribution — Colombia (Putumayo). Ecuador (Napo), Peru (Loreto)

Habitat & Ecology — In tropical wet forests, at an elevation of 250–400 m. Flowering period is uncertain.

IUCN Conservation Status — Not evaluated. There are only 2 collection records examined for this recently described species, which has been found in Colombia at the type location but also in Ecuador near Tena, and a population near Iquitos in Peru. There are 7 curated iNaturalist observations of this species (https://www.inaturalist.org/observations?place_id=any&taxon_id=1460794). Population and threat data is not available but our field observations indicate that this is a very uncommon species and would probably score as Endangered in a full assessment.

Notes — *Costus whiskeycola* has been widely cultivated around the world, originally grown from seeds collected by Tim Plowman when preparing *Plowman & Davis 4396* at the type locality. *Costus whiskeycola* looks quite similar to *C. erythrophyllus*, sharing many features of inflorescence and flowers, but it is markedly different in their leaves which lack the distinct plication, have a dark, olive green upper side and are thicker and have a waxy feeling in living plants. This species can also be distinguished from *C. erythrophyllus* by the length and shape of the ligules which are truncate to obliquely truncate instead of being deeply lobed.

**73. *Costus wilsonii*** Maas — Fig. 15d; Map 48

*Costus wilsonii* Maas (1976) 472. — Type: *Maas & McAlpin 1383* (holo U [U0007225]; iso COL [COL000000315], CR, F [F0047172F], K [K000586732], MO [MO-120589], NY [00320330]), Costa Rica, Puntarenas, Fila Las Cruces, near San Vito de Java, 1400 m, 22 Aug. 1974.

*Costus scaber* Ruiz & Pav. x *Costus lasius* Loes.: Maas (1972) 96.

Herbs, 0.5–2(–)4 m tall. *Leaves:* sheaths often prominently veined when dry, 5–10 mm diam; ligule obliquely truncate to slightly 2-lobed, 2–10 mm long; petiole 1–7 mm long; sheaths, ligule and petiole glabrous, sometimes sparsely to densely puberulous to villose; lamina narrowly elliptic, 8–30 by 1.5–7 cm, upper side glabrous, rarely sparsely to densely puberulous, lower side glabrous, rarely sparsely to densely puberulous to villose, base acute to rounded, apex long-acuminate (acumen with rolled inwards margin, 5–30 mm long, densely covered with long, silky hairs). *Inflorescence* cylindric to ovoid, 3–10 by 1.5–4.5 cm, enlarging to c. 15 by 6 cm in fruit, terminating a leafy shoot; bracts, bracteole, calyx, ovary, and capsule glabrous, rarely sparsely puberulous*. Flowers* abaxially orientated; bracts yellow, sometimes red, orange, or green, coriaceous, broadly ovate, 2–4 by 2–4 cm, apex obtuse, callus 2–6 mm long; bracteole boat-shaped, 12–17 mm long; calyx whitish to red, 5–7 mm long, lobes shallowly to very shallowly triangular, 0.5–1.5 mm long; corolla yellow to orange, 40–45 mm long, glabrous, lobes obovate, 30–35 mm long; labellum yellow, lateral lobes reddish striped, rolled inwards and forming a curved tube 8–10 mm diam, broadly oblong-obovate when spread out, 25–35 by 25–30 mm, irregularly lobulate, lobules red, 1–7 mm long; stamen yellow, 25–33 by 10–12 mm, not exceeding the labellum, apex yellow or reddish, obtuse, anther 4–9 mm long. *Capsule* ellipsoid to subglobose, 10–12 mm long.

Distribution — Costa Rica, Panama.

Habitat & Ecology — In cloud forests, almost always at elevations of (800–)1000–2200 m, very rarely at lower elevations. Flowering in the rainy season.

IUCN Conservation Status — Least Concern.

Notes — *Costus wilsonii*, occurring at rather high elevations in Costa Rica and Panama, can be told apart by its relatively small and long-acuminate leaves (the acumen with an rolled inwards margin) and bracts that are often yellow. The (yellow) labellum of this species is an intermediate form between spreading and tubular, it is tubular but the lateral lobes are often reddish striped, like in species with a horizontally spreading labellum. Another feature of this species is found in the venation of leaf sheaths and bracts, which is often prominent when dry.

**74. *Costus woodsonii*** Maas — Fig. 15e; Map 49

*Costus woodsonii* Maas (1972) 10, f. 47. — Type: *Von Wedel 2080* (holo MO [MO-120592]; iso GH [GH00030651], US [US00092986]), Panama, Bocas del Toro, Old Bank Island near Chiriquí Lagoon, 0–120 m, 13 Febr. 1941.

*Costus spiralis* auct. non (Jacq.) Roscoe: Woodson in Woodson & Schery (1942) 331.

Herbs, 0.5–3 m tall. *Leaves*: sheaths 7–12 mm diam; ligule truncate, 2–6 mm long; petiole 2–10 mm long; sheaths, ligule and petiole glabrous; lamina (narrowly) ovate to (narrowly) obovate 8– 26 by 4–11 cm, upper and lower side glabrous, base cordate, apex acuminate (acumen 10–20 mm long). *Inflorescence* globose, ovoid or cylindric, 3–11 by 2–3.5 cm, terminating a leafy shoot; bracts, bracteole, calyx, ovary, and capsule glabrous*. Flowers* abaxially orientated; bracts red, coriaceous, broadly ovate, 2–3 by 2–3 cm, apex obtuse, callus 4–5 mm long; bracteole boat-shaped, 12–20 mm long; calyx white to red, 4–9 mm long, lobes shallowly triangular, 1–2 mm long; corolla red to orange-red, 30–40 mm long, glabrous, lobes narrowly elliptic, 20–25 mm long; labellum yellow, lateral lobes rolled inwards and forming a curved tube 8–10 mm diam, ovate-oblong when spread out, 25–30 by 18–20 mm, irregularly lobulate, margin slightly crenulate; stamen yellow, 20–30 by 8–12 mm, not exceeding the labellum, apex obtuse, anther 7–9 mm long. *Capsul*e ellipsoid, 8–15 mm long.

Distribution — Nicaragua, Costa Rica, Panama, Colombia (Chocó, Nariño, Valle del Cauca).

Habitat & Ecology — In forested lowland thickets behind sandy beaches, at elevations of 0–50(–250) m. Flowering all year round.

IUCN Conservation Status — Least Concern.

Vernacular names — Costa Rica: Red cane.

Notes —*Costus woodsonii* is unique in the genus as it is occurring in thickets on or just behind sandy beaches; it can be recognized by usually small, ovate to obovate, completely glabrous leaf blades with a cordate base, combined with red bracts and tubular flowers.

**75. *Costus zamoranus*** Steyerm. — Fig. 15f; Map 50

*Costus zamoranus* Steyerm. (1964) 339. — Type: *Steyermark 54664* (holo F, 4 sheets [F0047184F, F0047185, F0047186F, F0047187F]; iso NY [NY00320357], US [US00092987]), Ecuador, Zamora-Chinchipe, Tambo Valladolid, along Río Valladolid, 1700 m (“2000 m”), 14 Oct. 1943.

Herbs, 1.5–3.5 m tall. *Leaves*: sheaths 8–20 mm diam; ligule truncate, 1–10 mm long; petiole 5– 20 mm long; sheaths, ligule and petiole glabrous; lamina narrowly elliptic to elliptic, 15–20 by 4–9 cm, upper and lower side glabrous, except for a few hairs near margin and apex, base acute, apex long-acute, sometimes long-acuminate (acumen c. 20 mm long). *Inflorescence* ovoid, 4–13 by 3–7.5 cm, terminating a separate leafless shoot c. 50 cm long, rarely terminating a leafy shoot, sheaths obliquely truncate, 2–5 cm long, glabrous; bracts, bracteole, calyx, ovary, and capsule glabrous*. Flowers* erect to adaxially orientated; bracts green, or red with green apex, coriaceous, broadly ovate, 3–5.5 by 2–4 cm, apex obtuse, callus inconspicuous, to c. 4 mm long; bracteole boat-shaped, 28–35 mm long; calyx red, 12–17 mm long, lobes shallowly triangular, 2–4 mm long; corolla yellow to yellowish orange, 70–85 mm long, glabrous, lobes narrowly elliptic, 60– 70 mm long; labellum dark reddish brown, upper part horizontally spreading, broadly obovate, 70–80 by 50–60 mm, lateral lobes striped with yellow, middle lobe reflexed, with yellow honey mark, margin irregularly crenate; stamen reddish, 45–55 by 9–10 mm, not exceeding the labellum, apex entire to irregularly dentate, anther 8–11 mm long. *Capsule* ellipsoid, 15–20 mm long.

Distribution — Ecuador (Loja, Morona-Santiago, Napo, Pastaza, Zamora-Chinchipe).

Habitat & Ecology — In premontane or montane cloud forest or sometimes in lowland rain forest, on clayey or sandy soils, at elevations of (300–)1000–1700 m. Flowering in the rainy season.

IUCN Conservation Status — Endangered. Since this 2015 assessment, Skinner has identified this species as very common in Napo (Ecuador) and a new assessment will likely result in a status of less conservation concern.

Notes — *Costus zamoranus* is characterized by a small ligule 1–10 mm long, a calyx 12– 17 mm long, and a (mostly) basal inflorescence with bracts green or red with green apical part, and a dark reddish brown labellum. This species is mostly found at high altitudes.

Several collections from Napo and Pastaza more or less resemble this species and may belong here, differing, however, in a shorter calyx (5–8 versus 12–17 mm long) and shorter bracteole (19–22 versus 28–35 mm long). In these collections the plants have some indument on sheaths and leaves whereas that is almost completely absent in ‘’real’ *C. zamoranus*. It involves:

ECUADOR. Napo, *Cerón M. 809* (MO, U), Reserva Biológica Jatun Sacha, Río Napo, 8 km below Misahuallí, 450 m; *Harling et al. 7016* (GB, U), Hacienda Cotapino (Concepción), 500 m,. Pastaza, *Alvarez et al. 2073* (MO), Cantón Archidona, Comunidad Pacto Sumaco, 1550 m,; *Asplund 19123* (NY, S), Mera, in forest on shore of Río Pastaza, 900 m,; *Asplund 19440* (NY, S), Vera Cruz, 900 m,; *Øllgaard 99596* (AAU), banks of Río Pastaza, S of Madre Tierra, 1000 m.

For field observations see Skinner & Jiménez León (2015). In this publication they mentioned some plants observed in the Cordillera del Condór, Río Nangaritza region, Zamora-Chinchipe, Ecuador, which were aberrant in some features like red-margined bracts, a shorter bracteole (c. 25 mm long) and a shorter calyx (c. 8 mm long). Judging from the photographs (unfortunately there is no herbarium material of it) we think that this may fall within the variation of *C. zamoranus*. This species was also found to be very common in the Narupa area of Napo Province in Ecuador.

**76. *Costus zingiberoides*** J.F.Macbr. — Fig. 15g; Map 50

*Costus zingiberoides* J.F.Macbr. (1931) 49. — Type: *L. Williams 3985* (holo F [F0041548F]), Peru, San Martín, Recreo, Yurimaguas, Lower Río Huallaga, 23 Oct. 1929.

Herbs, c. 1 m tall. *Leaves*: sheaths 4–8 mm diam; ligule 2-lobed, 1–2 mm long; petiole 1–4 mm long; sheaths, ligule and petiole densely to rather densely puberulous to villose; lamina linear, 12–22 by 1.5–2.5 cm, upper side glabrous, lower side densely puberulous, base obtuse to acute, apex long-acute. *Inflorescence* cylindric to ovoid, 5–10 by 2–3.5 cm, terminating a leafy shoot; bracts, bracteole, and calyx sparsely puberulous to glabrous, ovary rather densely sericeous, capsule glabrous*. Flowers* abaxially orientated; bracts orange-red, coriaceous, broadly ovate to depressed ovate, 1.5–2 by 2–3 cm, apex obtuse, callus 3–5 mm long; bracteole boat-shaped, 7– 13 mm long; calyx white, 7–9 mm long, lobes very shallowly triangular, c. 1 mm long; corolla bright yellow, c. 30 mm long, glabrous, lobes narrowly elliptic, c. 20 mm long; labellum yellow, lateral lobes rolled inwards and forming a tube c. 5 mm diam, narrowly oblong-elliptic when spread out, c. 20 by 6 mm, 5-lobulate, lobules yellow, c. 2 mm long; stamen yellow, 20–25 by c. 5 mm, not exceeding the labellum, apex red, obtuse, anther c. 5 mm long. *Capsule* globose, c. 8 mm diam.

Distribution — Colombia (Caquetá), Ecuador (Napo), Peru (Loreto, Pasco, San Martín).

Habitat & Ecology — In dense, moist forests, at elevations of 135–300 m. Flowering in the rainy season.

IUCN Conservation Status — Not evaluated. There are 10 collection records examined of which 6 specimens were collected within the past 50 years. This species has not been reported in any iNaturalist observations. The species is distributed in Amazonian parts of Colombia and Peru. Population and threat data is not available but based on the small number of recent collection records, this species may be Endangered or even Critically Endangered.

Notes — *Costus zingiberoides* is quite particular by having linear leaves, the only two other species with such leaves are the Costarican species *C. stenophyllus* and the Bolivian *C. alfredoi*. In *C. zingiberoides* the inflorescence is terminating a leafy shoot whereas in *C. stenophyllus* and *C. alfredoi* it is terminating a separate leafless shoot. Moreover, the calyx in *C. zingiberoides* is 7–9 mm long and in the two other species 15–16 mm.

There is a photograph of Mr. Sergio Botero taken in Colombia, Caquetá, Belén de los Andaquies, near Florencia, which may well be this species. There are also 2 sterile collections from Ecuador which probably belong here: *Kjaer-Pedersen 601* & *849* (AAU), Napo, Yasuní National Park, Estación Científica Yasuní, at Tiputini and surroundings,.

## INSUFFICIENTLY KNOWN SPECIES AND SPECIMENS OF *COSTUS*

### Costus sp. nov. “Chambi” — Fig. 15h

Herb, 2–3 m tall. *Leaves*: sheaths c. 20 mm diam; ligule slightly lobed, c. 13 mm long; petiole c. 15 mm long; sheaths, ligule and petiole glabrous; lamina narrowly elliptic, c. 52 by 10 cm, upper and lower side glabrous, base cordate, apex acuminate. *Inflorescence* cylindric, terminating a leafy shoot; bracts, appendages of bracts, bracteole, calyx, ovary, and capsule glabrous*. Flowers* abaxially orientated; bracts green, red-margined, coriaceous, broadly ovate, c. 3 by 3 cm, apex acute; appendages green, red-margined, foliaceous or possibly coriaceous, ascending, triangular, 2–3 by c. 2 cm, apex acute; bracteole boat-shaped, 30–35 mm long; calyx green, red-margined, 22–25 mm long, lobes deltate, 6–8 mm long; corolla pink, basally white, c. 80 mm long, glabrous, lobes narrowly elliptic, 50–60 mm long; labellum white to yellow, upper part horizontally spreading, ovate, c. 60 by 45 mm, lateral lobes striped with pale red, middle lobe reflexed, with yellow honey mark, margin irregularly crenate; stamen white, c. 40 by 15 mm, not exceeding the labellum, apex entire, dark yellow, anther 13–15 mm long. *Capsule* not seen.

Distribution — Peru (Cuzco).

Habitat & Ecology — In primary forest, at an elevation of c. 800 m. Flowering and fruiting: April.

Specimen examined (only photographs seen). *Chambi et al. 1098*, Peru, Cuzco, Prov. Quispicanchis, 18 km SW of Quince Mil, 800 m, 28 Apr. 2008,

Note — *Costus sp. nov. Chambi* can be distinguished by the bracts which have foliaceous appendages that are green with red margins. The bracts and the bractole are also green with red margins. Furthermore, it seems to lack any indument. While these observations have been made from photographs, we were unable to observe an herbarium specimen of this species for a full description.

### Costus sp. nov. “Gentry” — Fig. 15i

Herb, c. 0.3 m tall. *Leaves*: sheaths 10–15 mm diam; ligule unequally 2-lobed, 30–50 mm long; petiole 10–15 mm long; sheaths, ligule and petiole densely puberulous; lamina obovate, c. 36 by 16 cm, c. 10-plicate, upper side glabrous, lower side rather densely puberulous, base acute, apex acuminate (acumen 5–10 mm long). *Inflorescence* ovoid, c. 4 by 4 cm, terminating a leafy shoot; bracts densely puberulous, bracteole, calyx, ovary, and capsule not seen*. Flowers* abaxially orientated; bracts reddish, coriaceous, broadly ovate-triangular, c. 3 by 2.5 cm, callus 9–10 mm long; bracteole boat-shaped; calyx not seen; corolla yellow; labellum yellow, striped with red; stamen not seen. *Capsule* not seen.

Distribution — Colombia (Chocó).

Habitat & Ecology — Along roadsides, at an elevation of c. 220 m. Flowering and fruiting: January.

Specimen examined. — *Gentry & Rentería A. 24096* (COL, MO), Colombia, Chocó, Jequedó, 41 km W of Las Animas on Pan American Highway (under construction), c. 10 km E of Río Pato, 220 m, 12 Jan. 1979,.

Note — This species, with a very long ligule reaching 50 mm, cannot be described as the material is too incomplete.

***Costus fortalezae*** K.Schum. ex Loes. (1929) 708. — Type: *Ule 14b* (holo B, destroyed; photographs F, GH, MO, NY, etc.), Brazil, Acre, Lower Rio Juruá, near Fortaleza, Oct. 1901.

Note — A photo of the Berlin specimen of *Costus juruanus* has notations comparing it with this species and noting minor differences in the ligules, indicating that there are similarities between the two. From the Berlin photo of this specimen, it does indeed look similar to *C. juruanus*. Coauthor D. Skinner searched for this species along the Rio Juruá near a community known as Fortaleza not far from Cruzeriro do Sul in Acre, Brazil, but did not find anything matching it. Note that the name “Fortaleza” is applied to several cities and towns in Brazil, making it difficult to identify the locality with any certainty. Thus, we retain *C. fortalezae* as an insufficiently known species, but very likely it should be a synonym of *C. juruanus*.

### *Costus pulcherrimus* Kuntze

*Costus pulcherrimus* Kuntze (1898) 301. — *Costus cylindricus* Jacq. var. *pulcherrima* (Kuntze) K.Schum. (1904) 406. — Lectotype (designated here): *Kuntze s.n*. (lecto NY, 2 sheets [NY00320347, NY00320348]), Bolivia, Cochabamba, Río Juntas, 1600 m, 13–20 Apr. 1892.

Note — This very poorly preserved specimen shows only fragments of leaves and bracts, and in 1972 was placed as a synonym of *Costus laevis*. Coauthor D. Skinner visited the type locality in 2017 and found a plant that appears to match this specimen, with inflorescences produced terminally on leafy stems and bearing red unappendaged bracts and yellow, tubular flowers. It appears similar to *Costus vargasii,* but this cannot be confirmed based on available data and is thus considered an insufficiently known species.

### *Costus ulei* Loes

*Costus ulei* Loes. (1929) 709. — Type: *Ule 9192* (holo B, destroyed), Brazil, Acre, Seringal Monte Mó, Nov. 1911.

Note — D. Skinner searched for this species near the type locality along the Rio Acre in western Brazil, but did not find a plant that could be verified as being the same as this specimen. From the Berlin photo, it appears to be a small plant with very short ligules and narrow leaves, similar to *C. alfredoi*.

## EXCLUDED SPECIES

*Costus secundus* Roem. & Schult. (1817) 13. = ***Renealmia*** spec.

*Costus cernuus* Sw. ex Roem. & Schult. (1817) 25. = ***Renealmia cernua*** (Sw. ex Roem. & Schult.) J.F.Macbr.

*Costus podocephalus* Donn.Sm. (1897) 250. = ***Renealmia cernua*** (Sw. ex Roem. & Schult.) J.F.Macbr.

**3. *Dimerocostus*** Kuntze

*Dimerocostus* Kuntze (1891) 687. — Type: *Dimerocostus strobilaceus* Kuntze.

*Mulfordia* Rusby (1928) 165. — Type: *Mulfordia boliviana* Rusby (= *Dimerocostus argenteus* (Ruiz & Pav.) Maas).

Tall to gigantic herbs, often with leafy shoots at the base or the apex of the inflorescence. *Leaves*: ligule very short; lamina narrowly ovate to narrowly obovate, base acute to rounded, rarely cordate, apex often long-acuminate to caudate. *Inflorescence* a spike, often strongly elongate, terminating a leafy shoot. *Flowers* abaxially orientated; bracts coriaceous, green to yellow, often sheathing, ovate-triangular to broadly so, sometimes provided with foliaceous, apical appendages; bracteole tubular, adaxial side concave, 2-keeled or 2-winged; calyx very large, usually strongly exceeding the bracts, lobes often unequal, often with a distinct callus; corolla white or yellow; labellum large, white or yellow, rarely striped with orange or red, suborbicular (“not hooded”) or (broadly) ovate (“hooded”); stamen with the anther attached at the middle; stigma funnel-shaped, with an entire to 2-lobed dorsal appendage; ovary 2-locular, ovules organized in 2 rows per locule. *Capsule* 2-locular, ellipsoid to cylindric, coriaceous, tardily dehiscent. *Seeds* with a small, cushion-like aril.

Distribution — Five species occurring from Central America to the northern and western part of South America, at elevations of sea level to almost 2000 m.

Habitat & Ecology — Most species occur in lowland rain forests.

## KEY TO THE SPECIES OF *DIMEROCOSTUS*

1. Bracts provided with leafy appendages 2

1. Bracts without leafy appendages 4

2. Inflorescence nodding*. Flowers* white; labellum striped with red in the throat; bracts covered with a white, waxy layer. — Central and Western Colombia, 1100–2100 m 3. *D. cryptocalyx*

2. Inflorescence erect*. Flowers* yellow to cream; labellum without red stripes in the throat; bracts without a waxy layer 3

3. Lower side of leaves glabrous to hairy, but never silvery; appendages of bracts persistent. — Amazonian Colombia and Peru, 100–400 m 1. *D. appendiculatus*

3. Lower side of leaves silvery sericeous; appendages of bracts often not persistent (soon decaying). — Bolivia, Peru, 200–1900 m 2. *D. argenteus*.

4. Bracts partly sheathing, top angle 30–60 °; labellum yellow to cream, hooded. — Western part of tropical South America, in the East to French Guiana, 0–1000(–2100) m 4. *D. rurrenabaqueanus*

4b. Bracts completely sheathing, top angle 120–160 °; labellum white, not hooded. — Central America and the western part of tropical South America to Venezuela in the East, 0– 5009–1900) m5. *D. strobilaceus*

**1. *Dimerocostus appendiculatus*** (Maas) Maas & H.Maas, *comb. et stat. nov.* — Fig. 17a; Map 51

*Dimerocostus strobilaceus* Kuntze subsp. *appendiculatus* Maas, Fl. Neotrop. Monogr. 8 (1972) 25, f. 12. — Type: *Asplund 14727* (holo S, 4 sheets, [S-R-1506, S10-27114, S10-27115, S10-27116]), Peru, Loreto, Hacienda Indiana near mouth of Río Napo, 24 Nov. 1940.

Herbs, 1.5–3 m tall. *Leaves* congested towards the top of the shoot; sheaths 10–30 mm diam; ligule truncate, c. 1 mm long, at the base sometimes with a ring of erect, stiff hairs; petiole 4–10 mm long; sheaths, ligule and petiole densely to sparsely puberulous; lamina narrowly elliptic to narrowly obovate, 20–35 by 3–8 cm, upper side glabrous, lower side densely to rather densely puberulous to villose, not silvery, base acute, apex long-acuminate to caudate (acumen 15–40 mm long). *Inflorescence* ovoid to cylindric, 9–20 by 5–7 cm, to 12 cm wide in fruit, erect, often with leafy shoots arising from the base or apex; bracts, appendages of bracts, bracteole, calyx, ovary, and capsule densely to sparsely puberulous; bracts green, becoming brown in fruit, coriaceous, ovate-triangular to broadly ovate-triangular, 2–7 by 2–5 cm, without waxy layer, callus 6–12 mm long; appendages green, foliaceous, reflexed, triangular, 10-35 by 10–20 mm, persistent, apex acuminate; bracteole 15–30 mm long; calyx pale green, becoming brown in fruit, 30–35 mm long, lobes ovate-triangular to deltate, 3–6 mm long, callus 5–8 mm long; corolla cream, 65–80 mm long, glabrous, rarely sparsely sericeous towards the apex, lobes narrowly obovate, 45–60 mm long, all equal; labellum cream, throat yellow, upper part horizontally spreading, suborbicular, 70–90 by 80–130 mm, margin slightly crenate; stamen white or yellow, 50–60 by 12–15 mm, not exceeding the labellum, apex recurved, obtuse, crenate, anther 11–16 mm long. *Capsule* ellipsoid to cylindric, 25–50 mm long.

Distribution — Colombia (Amazonas), Peru (Junín, Loreto, Madre de Dios).

Habitat & Ecology — In dense forests and in thickets along rivers, at elevations of 100– 400 m. Flowering all year round.

IUCN Conservation Status – Not evaluated. There are 39 collection records examined of which 31 specimens were collected within the past 50 years. This species is also reported in 7 iNaturalist observations as a subspecies of *D. strobilaceus* (https://www.inaturalist.org/observations?place_id=any&taxon_id=507761). The species is recorded near the Amazon river in Peru and Colombia and also in Madre de Dios, Peru. Our observations indicate that this is a locally common species with no known specific threats.

Population and threat data is not available, but based on the number of collection records and our general field observations, this species would probably score a status of Least Concern.

Vernacular names — Peru: Caña agria.

Notes — *Dimerocostus appendiculatus* was formerly a subspecies of *D. strobilaceus*. The constant presence of appendages on the bracts, however, convinced us to consider it as a species of its own.

**2. *Dimerocostus argenteus*** (Ruiz & Pav.) Maas — Fig. 17b; Map 52

*Dimerocostus argenteus* (Ruiz & Pav.) Maas in Punt (1968) 37. — *Costus argenteus* Ruiz & Pav. (1798) 3, t. 4. — Lectotype (designated here): *Ruiz López & Pavón s.n.* (lecto MA [MA810650, MA810651, MA810652]; isolecto BM [BM000923840], F [F0041489F, FI011942], G [G00168453]), Peru, Huánuco, wet, shaded forests near Chinchao and Cuchero, anno 1778.

*Dimerocostus gutierrezii* Kuntze (1898) 301 (”*guttierezii*”). — *Dimerocostus strobilaceus* Kuntze subsp. *gutierrezii* (Kuntze) Maas (1972) 23. — Lectotype (effectively designated by Maas 1972): *Kuntze s.n.* (lecto NY [NY00320360]), Bolivia, Santa Cruz, Puerto Gutiérrez, San Ignacio, Río Yapacaní, 400 m, June 1892.

*Costus mooreanus* Rusby (1907) 454. — Lectotype (designated here): *Bang 2058* (lecto NY [NY00320339]; iso US [US00092969]), Bolivia, Prov. Cochabamba, Cochabamba, anno 1891.

*Mulfordia boliviana* Rusby (1928) 166, f. 1–6. — *Dimerocostus bolivianus* (Rusby) Loes. (1929) 716. — Type: *Cárdenas Hermosa 1165 A* (holo NY [NY00320361, NY00320362]), Bolivia, Beni, Rurrenabaque, 20 Nov. 1921.

*Dimerocostus bicolor* J.F.Macbr. (1930) 114. — Type: *Macbride 5001* (holo F [F0041536F]; iso G [G00168412]), Peru, Huánuco, Hacienda Villcabamba, Río Chinchao, 6000 ft, 17–26 July 1923.

Herbs, 1–4 m tall. *Leaves*: sheaths 10–25 mm diam; ligule truncate, with a salient, ciliate rim at the base, 1–3 mm long; petiole 2–6 mm long; sheaths, ligule and petiole glabrous to rather densely puberulous; lamina narrowly elliptic to narrowly obovate, 15–40 by 5–7 cm, upper side glabrous, lower side silvery sericeous, base acute to rounded, apex long-acute to acuminate (acumen 5-15 mm long). *Inflorescence* ovoid to cylindric, 10–40 by 5–6 cm, occasionally with leafy shoots at the base, erect; bracts, appendages of bracts, bracteole, calyx, ovary, and capsule glabrous to sparsely sericeous; bracts yellowish, coriaceous, triangular-ovate to narrowly triangular-ovate, 35–50 by 15–25 mm, without waxy layer, callus 1–6 mm long; appendages green, foliaceous, reflexed when dry, broadly ovate-triangular to ovate-triangular, 1–4.5 by 1–2.5 cm, often not persistent, decaying soon, base usually cordate, apex short-acuminate; bracteole 20–27 mm long; calyx yellowish green, 30–40 mm long, lobes 4–9 mm long; corolla cream, 65– 80 mm long, sericeous towards the apex, lobes narrowly obovate, 45–60 mm long, one lobe erect and much larger than the 2 other ones; labellum deep to pale yellow, hooded, upper part horizontally spreading, broadly ovate, 50–75 by 50–75 mm, lateral parts recurved, margin crenulate and fimbriate; stamen cream, 35–60 by 10–13 mm, slightly exceeding the labellum, apex recurved, obtuse, crenate, anther 14–15 mm long. *Capsule* ellipsoid, 20–25 mm long.

Distribution — Peru (Ayacucho, Cuzco, Huánuco, Junín, Madre de Dios, Pasco), Bolivia (Beni, Cochabamba, La Paz, Santa Cruz).

Habitat & Ecology — In wet forests, marshes, shady ravines, at elevations of 200–1900 m. Flowering all year round.

IUCN Conservation Status — Not evaluated. There are 56 collection records examined of which 49 specimens were collected within the past 50 years. This species is also reported in 24 recent iNaturalist observations (https://www.inaturalist.org/observations?place_id=any&taxon_id=516489). The species is widely in Peru and Bolivia. Our observations indicate that this is a common species within its range and some of these areas are protected areas with no known specific threats. Population and threat data is not available but based on the number of collection records and our general field observations, this species would probably score a status of Least Concern.

Vernacular names — Bolivia: Caña agria. Peru: Purum niña.

Notes — *Dimerocostus argenteus* can be distinguished at first glance from the other species of *Dimerocostus* by the silvery sericeous lower surface of the leaves.

**3. *Dimerocostus cryptocalyx*** N.R.Salinas & Betancur B. — Fig. 17c; Map 53

*Dimerocostus cryptocalyx* N.R.Salinas & Betancur B. (2004) 466, f. 1 & 2. — Type: *Betancur B. et al. 5976* (holo COL [COL000040001, COL000040003]; iso HUA [HUA0000020, HUA0000021], NY [NY01731846], U [U0123149, U0123150], US [US00576548]), Colombia, Antioquia, Mun. Frontino, Corregimiento Nutibara, 7–8 km between Alto de las Cuevas and La Blanquita, 1285–1400 m, 27 Jan. 1995.

Herbs, 1–5 m tall. *Leaves*: sheaths 15–20 mm diam; ligule obliquely truncate, 2–5 mm long; petiole 5–10 mm long; sheaths, ligule and petiole glabrous; lamina narrowly elliptic, 28–46 by 5–10 cm, upper and lower side glabrous, base acute to obtuse, apex long-acute. *Inflorescence* cylindric, erect when young, soon nodding, 25–45 by 4.5–7 cm; bracts, appendages of bracts, bracteole, calyx, ovary, and capsule glabrous; bracts green, chartaceous, narrowly ovate-triangular, 5–7 by 1–2 cm, covered with with a waxy, white layer, callus absent; appendages green, foliaceous, erect (the lower ones) to recurved (the upper ones), narrowly elliptic to deltate, 2–5 by 0.5–1 cm, apex acute; bracteole 18–20 mm long; calyx yellowish green, 28–35 mm long, lobes 4–10 mm long, calllus inconspicuous; corolla white, c. 60 mm long, glabrous, lobes narrowly oblong-elliptic, 32–34 mm long, all equal; labellum white, striped with red in the throat, upper part horizontally spreading, suborbicular, c. 75 by 60 mm, margin fimbriate; stamen white, c. 35 by 9–10 mm, not exceeding the labellum, apex recurved, obtuse, crenate, anther c. 11 mm long. *Capsule* ellipsoid, 25–30 mm long.

Distribution — Colombia (Antioquia, Boyacá, Caldas, Chocó, Risaralda, Santander).

Habitat & Ecology — In forests, at elevations of 1100–2100 m. Flowering all year round.

IUCN Conservation Status — Not evaluated. There are 16 collection records examined of which all were collected within the past 50 years. This species is also reported in 4 recent iNaturalist observations (https://www.inaturalist.org/observations?place_id=any&taxon_id=910959). The species is endemic to mountainous regions of Colombia where it is recorded in all three cordilleras. Our observations indicate that this is a somewhat rare species within its range and some of these areas are protected areas with no known specific threats. Population and threat data is not available but based on the number of collection records and our general field observations, this species would probably score a status of Near Threatened or Vulnerable.

Notes — *Dimerocostus cryptocalyx* is aberrant from all other species of the genus by a nodding inflorescence, a white and reddish striped labelllum and a white, waxy cover of the bracts.

Field observations — Flowers aromatic with a slight sweet odor of maple (*N.R. Salinas et al. 667*).

**4. *Dimerocostus rurrenabaqueanus*** (Rusby) Maas & H.Maas, *comb. nov*. — Fig.17d; Map 54

*Costus rurrenabaqueanus* Rusby, Mem. New York Bot. Gard. 7 (1927) 219. — Lectotype (designated here): *Cárdenas Hermosa 1882* (lecto NY [NY00320352]; isolecto BKL [BKL00000404]), Bolivia, Beni, Rurrenabaque, 1000 ft, 28 Nov. 1921.

*Dimerocostus tessmannii* Loes. (1929) 715. — Type: *Tessmann 3751* (holo B, destroyed; type photographs F, GH, MO, NY, U, US), Peru, Loreto, Parinari, Lower Río Marañon, 100 m, 8 Aug. 1924.

*Dimerocostus williamsii* J.F.Macbr. (1931) 50. — Type: *Ll. Williams 4291* (holo F; iso G [G00168415]), Peru, Loreto, Yurimaguas, Fortaleza, 155 – 210 m, Oct. – Nov. 1929.

Herbs, 2–6 m tall. *Leaves*: congested towards the top of the shoot; sheaths 10–25 mm diam; ligule truncate, 1–2 mm long; petiole 5–10 mm long; sheaths, ligule and petiole glabrous, sometimes sparsely to densely puberulous; lamina narrowly ovate to narrowly obovate, 20–40 by 4–8 cm, upper side glabrous, lower side glabrous, sometimes sparsely to rather densely puberulous or sericeous, base acute, rarely cordate, apex long-acuminate to caudate (acumen 10– 50 mm long). *Inflorescence* ovoid to cylindric, 13–25 by 5–7 cm, often with leafy shoots arising from the base or apex; bracts, bracteole, calyx, ovary, and capsule glabrous, sometimes sparsely to densely puberulous; bracts green, becoming brown in fruit, coriaceous, ovate-triangular to broady ovate-triangular, partly sheathing (top angle 30–60 °), apex acute to pungent, 2–4 by 2–3 cm, callus 3–8 mm long, without leafy appendages; bracteole 20–30 mm long; calyx green, becoming brown in fruit, 25–40 mm long, lobes ovate-triangular to deltate, 5–8 mm long, callus 3–7 mm long; corolla cream, c. 75 mm long, glabrous, lobes narrowly ovate, c. 50 mm long, one lobe erect and much larger than the 2 other ones; labellum yellow to cream, striped with orange in the throat, hooded, upper part horizontally spreading, ovate, c. 80 by 50 mm, lateral parts recurved, margin irregularly denticulate and fimbriate; stamen yellow to cream, 35–45 by 13 mm, not exceeding the labellum, apex recurved, obtuse, crenate, anther 12–15 mm long. *Capsule* ellipsoid to cylindric, 35–40 mm long.

Distribution — Colombia (Amazonas, Chocó), Venezuela (Delta Amacuro), Guyana, Suriname, Ecuador (Morona-Santiago, Orellana, Pastaza, Zamora-Chinchipe), Peru (Amazonas, Cuzco, Huánuco, Loreto, Madre de Dios, Pasco, San Martín, Ucayali), Bolivia (Beni, Cochabamba, Pando, Santa Cruz), Brazil (Acre).

Habitat & Ecology — In wet non-inundated or periodically inundated (tahuampa, várzea) forests, at edge of forests, in open places, or in swamps, at elevations of 0–1000(–2100) m. Flowering all year round.

IUCN Conservation Status — Not evaluated. There are 90 collection records examined of which 70 specimens were collected within the past 50 years. The species is recorded widely in Peru and also in Bolivia, Brazil, Colombia, Ecuador and Suriname. Our observations indicate that this may be a common species but is often confused with *D. strobilaceus* so observation data may be inaccurate. Population and threat data are not available but based on the number of collection records and our general field observations, this species would probably score a status of Least Concern.

Vernacular names — Ecuador: Gonekim (Huaorani), Noneguimonkagui (Huaorani), Ucuti muyo. Peru: Cañagria, Sachahuiro amarillo, Shinaukush, Shiwanuk, Siwanúk.

Notes — This species has been called by Maas (1972) *Dimerocostus strobilaceus* subsp. *gutierrezii*. Recently Dave Skinner studied the type collection of *D. gutierrezii* (*Kuntze s.n.* (NY) from Bolivia) consisting of an old inflorescence, but lacking any leaves. Skinner came to the surprising conclusion that it should be included in the synonymy of *D. argenteus*. Moreover, Skinner visited in 2017 the type locality of *D. gutierrezii* (Bolivia, Río Yapacani) and found there only one species of *Dimerocostus*, namely one with a silvery lower leaf side, exactly corresponding with *D. argenteus*. Because of this discovery a new name had to be given for the former subsp. *gutierrezii*. The oldest name available, namely *D. rurrenabaqueanus,* is herewith selected.

**5. *Dimerocostus strobilaceus*** Kuntze — Fig. 17e, g; Map 55

*Dimerocostus strobilaceus* Kuntze (1891) 687. —Lectotype (designated here): *Kuntze 1873* (lecto NY [NY00320358, NY00320359]; isolecto K [K000586878]), Panama, Colón, Monkhill, near Colón, 4 Nov. 1874.

*Dimerocostus elongatus* J.Huber (1906) 545. — Syntypes: *J. Huber 1384* (MG, F [F0041488F]), Peru, Loreto, Quebrada de Canchahuaya, 27 Oct. 1898; *J. Huber 1461* (not seen), Peru, Loreto, Pampa del Sacramento.*Dimerocostus uniflorus* auct. non (Poepp. ex Petersen) K.Schum.: K.Schum. (1904) 427, f. 50A, p.p., excl. type.

Herbs, 1.5–6 m tall. *Leaves* congested towards the top of the shoot; sheaths 10–30 mm diam; ligule truncate, 1–2 mm long, at the base often with a ring of erect, stiff hairs; petiole 2–5 mm long; sheaths, ligule and petiole sparsely to densely puberulous; lamina narrowly ovate to narrowly obovate, 12–55 by 3–12 cm, upper side glabrous, lower side sparsely to rather densely puberulous or glabrous, base acute, apex long-acuminate to caudate (acumen 10–50 mm long). *Inflorescence* ovoid to cylindric, 10–45 by 3–8 cm, to 10 cm wide in fruit, often with leafy shoots arising from the base or apex; bracts, bracteole, calyx, ovary, and capsule sparsely to densely puberulous; bracts green, becoming brown in fruit, coriaceous, completely sheathing (top angle 120–160 °), apex obtuse, 2–6 by 2–4 cm, callus 2–6 mm long, without leafy appendages; bracteole 20–30 mm long; calyx green, becoming brown in fruit, 25– 30 mm long, lobes ovate-triangular, 5–7 mm long, callus 3–10 mm long; corolla white, 65–80 mm long, glabrous, rarely sparsely sericeous towards the apex, lobes narrowly obovate, 45–65 mm long, all equal; labellum white, sometimes tinged with orange or yellow in the throat, upper part horizontally spreading, suborbicular, not hooded, 80–120 by 80–120 mm, margin slightly crenate; stamen white, c. 45 by 13 mm, not exceeding the labellum, apex recurved, obtuse, dentate, anther 13–15 mm long. *Capsule* ellipsoid to cylindric, 25–50 mm long.

Distribution — Honduras, Nicaragua, Panama, Costa Rica, Colombia (Amazonas, Antioquia, Boyacá, Caldas, Caquetá, Casanare, Cauca, Cesar, Chocó, Córdoba, Cundinamarca, Guaviare, Magdalena, Meta, Nariño, Norte de Santander, Putumayo, Quindío, Santander, Valle del Cauca), Venezuela (Amazonas, Apure, Barinas, Mérida, Portuguesa, Táchira, Zulia), Ecuador (Carchi, Chimborazo, El Oro, Esmeraldas, Imbabura, Los Rios, Manabí, Morona-Santiago, Napo, Orellana, Pastaza, Pichincha, Santiago-Zamora, Sucumbios), Peru (Huánuco, Loreto, Pasco).

Habitat & Ecology — In wet forests, often in open places or at forest edges, in lowland thickets and along streams, at elevations of 0–500(–1900) m. Flowering all year round.

IUCN Conservation Status — Not evaluated. There are 428 collection records examined of which 295 specimens were collected within the past 50 years. This species is also reported in 165 recent iNaturalist observations (https://www.inaturalist.org/observations?place_id=any&taxon_id=346186). The species is widely distributed from Nicaragua to Peru. Our observations indicate that this is a very common species with no known specific threats. Population and threat data are not available but based on the number of collection records and our general field observations, this species would almost certainly score a status of Least Concern.

Vernacular names — Colombia: Cagalera, Cagaloduro, Cañagria, Caña fistula, Cañaguate, Caña pangula, Conaigre, Kondrú (Embera), Mamba, Pángula, Ta-gú-chu, Vara de mico. Ecuador: Nenenkemo, Ucuti muyo, Untuntu (Huaorani). Nicaragua: Caña agria. Peru: Cañagria, Sachahuiro.

Notes — The name *Dimerocostus uniflorus* has been misapplied in the past to plants belonging to *D. strobilaceus* (see Maas 1972). There are some plants found in Peru and western Brazil, now included in *D. rurrenabaqueanus*, that have yellow corolla and yellow labellum, flower spreading, (not hooded), bracts fully sheathing, top angle > 60 °, which may belong to *D. strobilaceus,* which is normally found with white flowers in areas north of Peru. The status of these yellow flowering plants is uncertain and needs further study.

**4. *Monocostus*** K.Schum.

*Monocostus* K.Schum. (1904) 427. — Type: *Monocostus ulei* K.Schum. (= *Monocostus uniflorus* (Poepp. ex Petersen) Maas).

Monotypic, for description see under species.

1. ***Monocostus uniflorus*** (Poepp. ex Petersen) Maas — Fig. 17f, h; Map 56

*Monocostus uniflorus* (Poepp. ex Petersen) Maas in Punt (1968) 37. — *Costus uniflorus* Poepp. ex Petersen (1890) 58. — *Dimerocostus uniflorus* (Poepp. ex Petersen) K.Schum. (1904) 427, f. 50A, p.p., type only. — Neotype (designated by Maas 1972: 18): *Klug 4156* (neo GH [GH00030670; isoneo BM [BM000923842], E [E00319720], F [F0041490F], K [K000586880], MO [MO-202805], NY [NY00320355], S [S-R-3649], U [U 1212438], US [US00092984]), Peru, San Martín, Chazuta, Río Huallaga, 260 m, May 1935. A neotype was designated by Maas (1972) as the type, *Poeppig 2116 B* from Peru, San Martín, Yurimaguas, Dec. 1830, was destroyed in the Vienna Herbarium (W) in 1943.

*Monocostus ulei* K.Schum. (1904) 428, f. 51. — Lectotype (designated by Maas 1972: 18): *Ule 6333* (lecto MG [MG006196]; isolecto HBG), Peru, San Martín, near mouth of Río Mayo, Juan-Guerra near Tarapoto, Oct. 1902.

Low herbs, 0.2–0.6 m tall, with long roots. *Leaves*: sheaths 3–6 mm diam; ligule 1–2 mm long; petiole c. 1 mm long; sheaths, ligule and petiole rather densely puberulous to glabrous; lamina elliptic to obovate, or narrowly so, 2.5–8 by 1–4 cm, glabrous except for the ciliate, reddish margin, base rounded to cordate, apex shortly and abruptly acuminate (acumen 1–4 mm long). *Flowers* abaxially orientated, solitary in the axils of the upper leaves, provided with a 4–5 mm long pedicel; pedicels, bracteole, calyx, ovary, and capsule glabrous; bracteole tubular, 15–22 mm long, 2-lobed, lobes deltate, 2–4 mm long, with a 1–2 mm long callus just below the apex; calyx green, 20–30 mm long, lobes triangular, 3–5 mm long; corolla cream, 60–70 mm long, glabrous, lobes narrowly elliptic, 25–30 mm long; labellum yellow, throat striped with orange, upper part horizontally spreading, broadly obovate to almost circular, 40–50 by 50–60 mm, margin crenulate; stamen cream to yellow, with the anther attached at the middle, 25–40 by 6–8 mm, slightly exceeding the labellum, apex striped with orange, irregularly dentate, anther 4–6 mm long; stigma funnel-shaped, with an entire dorsal appendage; ovary 2-locular, flattened, ovules organized in 1 row per locule. *Capsule* 2-locular, narrowly ellipsoid, chartaceous, flattened, longitudinally dehiscent, 40–60 by 5–6 mm. *Seeds* with a lacerate aril.

Distribution — Peru (Loreto, San Martín).

Habitat & Ecology — In low forests and one collection from dry forest with cactus undergrowth, at elevations of 200–350 m. Flowering May to December.

IUCN Conservation Status — Not evaluated. There are only 8 collection records examined for this species, only 4 of which were collected within the past 50 years. It was only known in a small area of department San Martín, Peru to the southeast of Tarapoto. The earliest recorded wild collection in the Global Biodiversity Information Facility (https://www.gbif.org/occurrence/search?taxon_key=2757046) was in 1830, with the most recent collection events occurring in 1935. The region of the type specimen was visited by Specht in 2001 and Skinner in 2008 but no wild plants or populations were found. We believe this species would be assessed as Critically Endangered if formally assessed for IUCN status; it may even be extinct in the wild.

Vernacular names — Peru: Huachico, Sacha huirillo.

## Acknowledgements

We are very grateful to the curators of the herbaria mentioned under Material and Methods, for making their material available for our study. Many thanks to the horticulturalists and botanists who spent time and care cultivating our living *Costaceae* material in the greenhouses of Meise (Belgium) and Arnhem (Burgers’ Bush); we are particularly thankful to Ernst Kamphuis at Burgers’ Zoo, who eagerly accessioned and carefully cared for the many *Costaceae* specimens now held in cultivation within their indoor tropical rain forest (known as Burgers’ Bush). For many of the details included in this publication, we are greatly indebted to our naturalist and botanist colleagues who provided us with photographs of *Costaceae* for better understanding of species boundaries: Alfredo Fuentes and Stephan Beck (Bolivia), Narcísio Bigio (Brazil), Julio Betancur (Colombia), Stephan Gröger (Germany), Jana Leong-Škornicková (Singapore), Jonathan Amith (Mexico), Rosanna Maguiña (Peru), and Robin Foster (USA). During our many field trips in tropical America, PM and HM are grateful for support in Brazil (especially Douglas Daly and Sir Gillean Prance, who both invited us to undertake field work in Amazonian Brazil); Colombia (especially the late Tim Plowman); many botanists from the herbaria at Medellín including Dario Sánchez Sánchez, Saül Hoyos-Gómez, Jorge Mario Vélez, and León Morales S., who made it possible to undertake several field trips in the department of Antioquia; and the late Philip Silverstone-Sopkin of the Herbarium Cali (CUVC)); Costa Rica (especially Barry Hammel and Nelson Zamora); the Dominican Republic (especially the late Carles Roersch); and Panama (especially the late Bob Dressler and Alicia Ibañez). PM and HM are much indebted the Hugo de Vries Fonds and the Treub Maatschappij for financing their field work in Colombia in 2013 and 2018. CDS thanks Cornell University College of Agriculture and Life Sciences for supporting field, herbarium, and laboratory research and the L.H. Bailey Hortorium community for inspiring taxonomic and revisionary work within a university setting. Thanks to Gisel De La Cerda, Chodon Sass, Clarice Guan, Maria Pinilla Vargas, Julianna Harden and Jacob B. Landis who all played a role in building and organizing our neotropical *Costaceae* collections and databases for continued exploration of phylogenetic and evolutionary relationships and especially J. Harden for making the final versions for all distribution maps.

## INDEX OF EXSICCATAE

Note: Index is sorted by the surname and initials of the main collector.

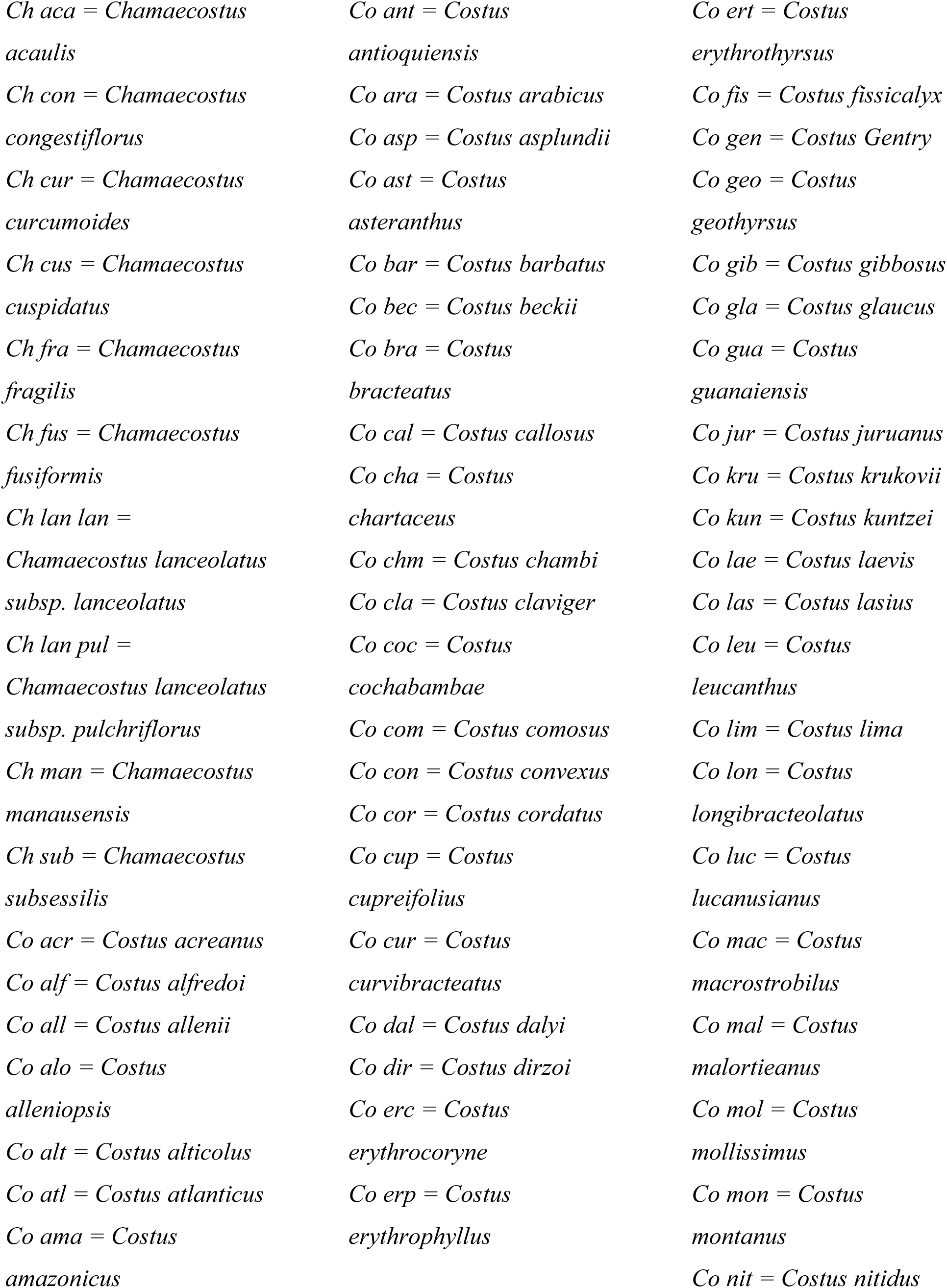
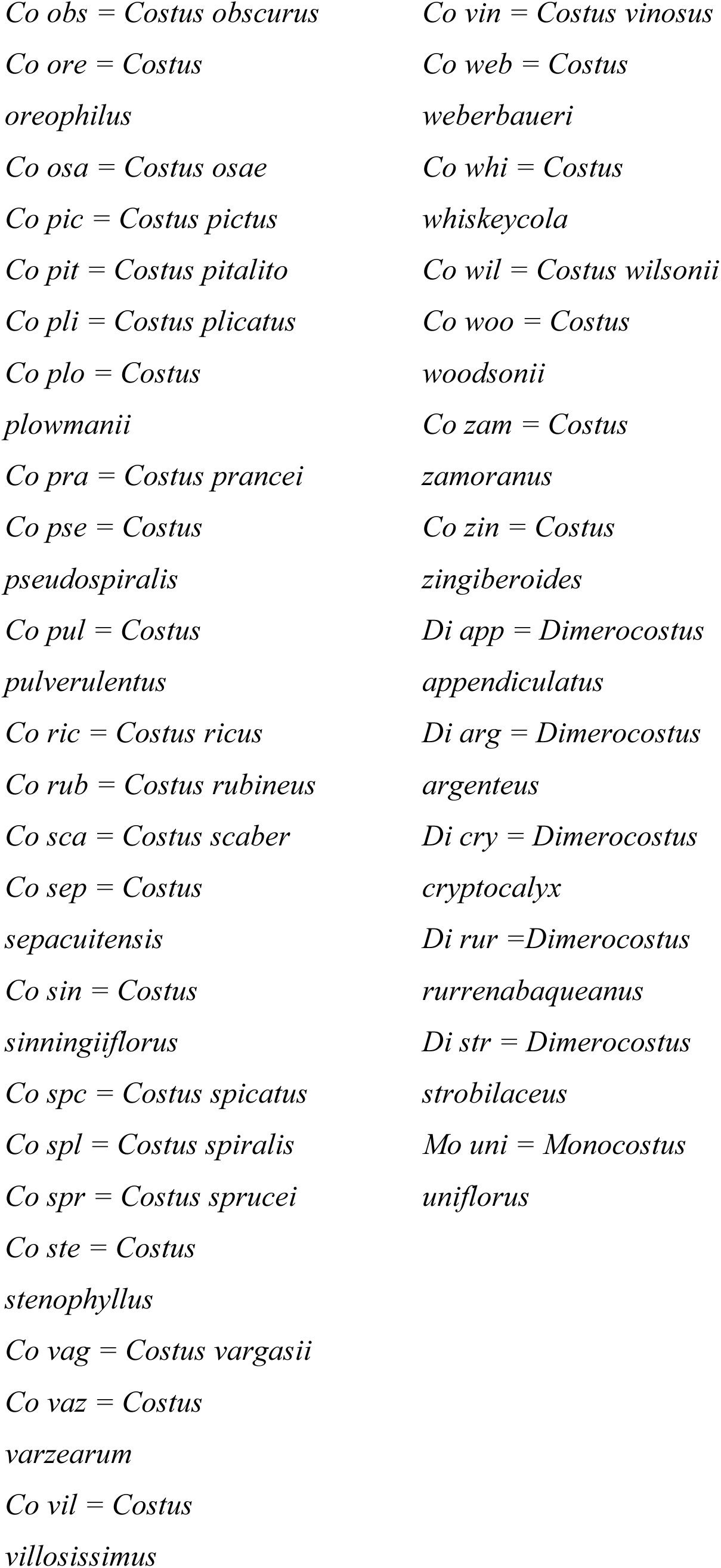

Abbott 7543: Co mal; 7560: Co spc; 8682: Co bar; 16606: Co ara; 17612: Co bar; 25045: Co vaz – Abraham 344: Co spl – Acero Duarte 22: Co sca – Acevedo-Rodríguez 3356: Co sca; 4853: Co ert; 8171: Co ara; 8581: Co jur; 8622: Co sin; 8699 : Di arg; 9182: Co spl; 9218: Co sca; 9840: Co vag; 11703: Co sca; 14993: Co ara – Ackerman 4495: Co spc; 4496: Co sca – Acosta 37: Co spl – Acosta Solís 5198: Co pul; 6142: Co lae; 6143: Co lim; 10894: Co pul; 13692: Co pul; 13973: Co lae – Acosta Vargas 134: Co sca; 1163: Co pul; 1356: Co ste; 1524: Co pul; 2165: Co woo; 6257: Co pul – Acuna 20088: Co pul – Adalardo-Oliveira 2610: Co ara – Agostini, C 1046: Co com – Agostini, G 1759: Co sca – Agra 1278: Co las – Aguilar 21: Co las; 49: Co kun – Aguilar Fernández 188: Co sca; 248: Co ste; 249: Co las; 1783: Co las; 1937: Co lim; 2008: Co kun; 2218: Co vil; 2321: Co kun; 2376: Co sca; 2516: Co sca; 3299: Co las; 5537: Co pul; 5540: Co lim; 6802: Co ric; 8020: Co mal; 12053: Co pli; 12066: Co lim; 12128: Co gla; 12163: Co vil – Aguilar Gutiérrez 10122: Co pic; 12348: Co pul – Aguilar Hidalgo 289: Co pul; 290: Co pic – Aguilar M 51: Co pul; 10560: Co pul – Aguilar R 583: Co ara; 814: Co ara –Aguinda 343: Co las; 366: Co ara; 394: Co kru; 467: Co lon – Aguirre Galviz 980: Co las – Aitken 41: Co sca – Aker 638: Co pul – Alain 23786: Co spc; 32105: Co mac – Alarcón Gallegos 141: Di str; 19374: Di str – Albán Castillo 9121: Co sca; 9141: Co ert; 9343: Co asp; 9443: Co asp; 9742: Co sca; 11010: Co asp; 11123: Co ama – Albert de Escobar 735: Co asp; 1398: Co pul; 1686: Co plo; 1750: Co pul; 2157: Co kun; 3310: Co las; 3435: Co com; 8180: Co mac; 8267: Co lim; 8270: Di str; 8300: Di str –Albuquerque 485: Co sca; 627: Co sca; 1339: Co ara; 1389: Ch aca – Albuquerque L 51: Co ert – Alegría 28: Co woo – Alencar 51: Co ara – Alexiades 1054: Di app; 1079: Co ara; 1107: Co gua; 1205: Di rur – Alfaro Vindas 248: Co las; 838: Co wil; 1692: Co wil; 4018: Co wil; 5798: Co sca – Allard 13481: Co spc; 22060: Co las; 22225: Co sin – Allemão e Cysneiro 1503: Ch sub – Allen, B 15223: Co pul; 15412: Co pul – Allen, C 247: Co mac; 298: Co sca – Allen, PH 762: Co las; 866: Co pul; 938: Co pul; 1789: Co las; 1825: Co pul; 1873: Co cur; 1952: Co sca; 2167: Co las; 2168: Co las; 2185: Co all; 2199: Co kun; 2217: Co kun; 2427: Co cur; 2619: Co vil; 3549: Co las; 3587: Co all; 3595: Co all; 4366: Co com; 4574: Co lim; 4685: Co wil; 5267: Co ste; 5269: Co pul; 5551: Co kun; 5572: Co lim; 5578: Co mac; 5598: Co pli; 5625: Co com; 6037: Co ste; 6261: Co kun; 6603: Di str; 6780: Co pic; 17274: Di str – Allorge 303: Ch con – Almeida, J 353: Co ara – Almeida, JC 452: Co mac; 3234: Co mac – Alston 8906: Co las – Altamirano Azurduy 1638: Co bec; 2372: Co ara –Altson, AHG 8832: Co pul – Altson, RA 8: Co ara – Alvarado, A 3: Co sca – Alvarado, F 186: Co sca – Alvarenga 627: Co spl – Alvarez Dávila 701: Co pul – Alvarez 701: Co pul; 758: Co lim; 768: Co kun; 799: Di str; 800: Di str; 2073: Cozam – Alverson 58: Co ant; 209: Di str – Alves 15: Co spl; 2363: Co acr; 2365: Ch aca; 2476: Co sca; 2555: Ch lan lan – Alvim 297: Co spl – Alzate, A 562: Co sca; 618: Co erp – Alzate, F 317: Co kun; 688: Co las; 3329: Co cha – Amaral, IL 291: Co ara; 323: Co lon; 510: Co kru; 620: Co sca; 1094: Co ert; 1221: Co ara; 1462: Co spl – Amaral, MCE 95-24: Co spl – Amaya 182: Co sca; 356: Co las – Amaya M 53: Co sca; 162: Di str – Amith 2386: Co pic – Amorim 949: Co ara; 2365: Ch sub; 6726: Co sca – Ancuash 375: Co sca; 565: Co ama; 1104: Co sca – Anderson, A 82: Co ara – Anderson, WR 9753: Co spl; 10570: Co sca; 11847: Co ara; 11915: Co ara –Andrade, AG 809: Co ara – Andrade, MG 102A: Co cha; 192: Co cha – Andrade, R 33011: Di str; 33050: Co lon; 33159: Co lon – André, EF 291: Co sca; 2394: Co pul – André, M 949: Co ara – Andrews 360: Co sca – Andrle 41: Co pul – Angarita 34: Di cry – Annable 3175: Co woo; 3571: Co kun – Antonio 869: Co cur; 1149: Co kun; 1788: Co mac; 2008: Co com; 2334: Di str; 2336: Co lim; 3847: Co sca; 3917: Co sca; 4058: Co sin; 4226: Co lim; 4309: Co pul; 4337: Co las; 4341: Co kun; 4369: Co lim; 4370: Co vil; 4403: Co kun; 4435: Di str; 4496: Co sca; 4626: Di str; 4654: Co sca; 4913: Co wil; 4925: Co las; 4996: Co cur; 5048: Co wil; 5067: Co cur; 5136: Co sca – Appun 762: Co ara – Arango 47: Co las – Arapa Apaza 31: Di str – Araque-Molina 18Va-81: Co ara; 18Va-82: Co sca; 18Va-83: Co ara – Araquistain 636: Co pic; 1172: Co pic; 2340: Co pul; 2343: Co sca; 2415: Co sca; 2501: Co sca; 2788: Co pul; 3026: Co pul; 3048: Co pul; 3057: Co vil; 3220: Co pul; 3228: Co pul; 3233: Co mac; 3253: Co lim; 3383: Co woo –Araujo, D 2285: Co ara; 3524: Co ara – Araujo, GM FEEP115: Co spl – Araujo Murakami 53: Co sca; 768: Co sca; 1009: Co sca; 1182: Di arg; 1213: Co sca; 1682: Di arg – Araya Mena 172: Co sca; 432: Co mal; 561: Co mal; 578: Co kun – Arbeláez S 2460: Co lae – Archer 575: Co ant; 1474: Co ant; 1476: Di str; 1918: Co las; 2075: Co kun; 2094: Co sca; 2274: Co sca; 7982: Co ara; 8269: Co ara – Arévalo 7: Co lon – Argent 6453: Co ara – Argüello 315: Co pul – Argumosa, de 137: Co ara; 138: Co ara – Arias 16: Di str; 64: Co las – Arias Ch 269: Co lae Arias Garzón: 1: Co ara; 147: Co las; 244: Co sca; 254: Co las – Arias Guerrero, S 13: Co las; 15: Co las; 29: Co kun; 34: Co las; 35: Co las; 86: Co las; 99: Co lim – Arias-G, JC 520: Co sca; 827: Ch fra; 1120: Co kru; 1153: Co sca; 1539: Co las; 1617: Co ara; 1860: Co lon – Aristeguieta 1145: Co pul; 1163: Co mac; 1589: Di str; 2744: Co sca; 3188: Co spl; 3778: Co sca; 3868: Di str; 3899: Co sca; 3901: Co ara; 3922: Co ara; 4556: Co spl; 4595: Co ara; 4603: Co sca; 4606: Co com; 4742: Co sca; 5172: Co com; 5176: Co spl; 6192: Co spl; 6345: Co mac; 6347: Co sca – Ariza 647: Co ant – Aronson 813: Co cha; 898: Co asp – Arroyo Padilla 1334: Co ara; 1795: Ch aca – Arvigo 223: Co pul; 437: Co pul; 623: Co mac – Asplund 5330: Co pul; 5422: Co ara; 5489: Co mac; 7833: Co ore; 9238: Co sca; 10273: Di str; 13365: Co lae; 14122: Co erc; 14359: Co sca; 14578: Co ara; 14727: Di app; 15162: Co lim; 15412: Co gib; 15570: Co mac; 15895: Co gib; 16421: Co gib; 16494: Co gib; 16763: Co gib; 18160: Co gib; 18179: Co lim; 18184: Di str; 19115: Co asp; 19123: Cozam; 19189: Co ama; 19431: Co ama; 19440: Cozam; 19661: Co sca; 19765: Co lon; 19784: Di str; 20068: Co ore; 20390: Co geo; 20526: Co ama – Asprilla 14: Co plo – Assumpção 72: Co pra – Assunção 107: Ch man – Asunción Leveau 6: Co sca – Atha 775: Co mac; 1004: Co pul; 6689: Co mac; 6699: Co pic – Atwood Jr 6: Co pic; 93: Co pul; 150: Co pic; 2949: Co woo; 4228: Co pul – Aubréville 61: Co spl; 158: Ch con – Augello 78: Di rur – Augusto, Bro 482: Co spc; 751: Co spc – Aulestia, C 387: Co pul; 1049: Di str; 1126: Co kun – Aulestia, M 434: Di str; 3252: Co sca – Austin 4189: Co sca – Avella M 849: Di str; 1416: Di str – Avendaño 71: Co kru – Ávila 4229: Co pul – Aviles 11: Co vil; 41: Co kun; 398: Co pul; 890: Co vil – Axelrod 4469: Co mac – Ayala 1: Co sca – Ayala Flores 93: Co sca; 493: Co ara; 499: Co sca; 633: Co sin; 1872: Co sin; 1873: Co sca; 1901: Co ara; 1962: Co erc; 2037: Co erc; 2128A: Co sca; 2145: Co sca; 2195: Co kun; 2303: Di rur; 2340: Co sca; 2394: Co sin; 2584: Co sca; 2609: Co sca; 2749: Co las; 2941: Co lon; 2984: Co las; 3018: Co sca; 3214: Co las; 3284: Co sca; 3304: Co ara; 3430: Co sca; 3497: Co sca; 3855: Co sca; 3912: Co sca; 3952: Co mac; 4490: Co sca; 5589: Co sca; 5594: Co sin; 5696: Co sca; 5882: Co sin; 5884: Co sca – Aymard C 1231: Co ara; 1411: Co mac; 2658: Co spl; 2890: Co spl; 3689: Co spl; 3961: Co sca; 5374: Co ara – Azevedo 1171: Co spl – Azofeifa Zúñiga 14: Co wil.

Bacle 46: Co spl – Badillo 1980: Co spl; 2004a: Co vil; 2023: Co com; 4332: Co spl; 4394: Co sca; 4671: Co las – BAFOG (Service Forestier) 4163: Co sca; 4434: Co sca – Bailey, EZ 83: Co pul – Bailey, LH 117: Co all; 324: Co vil; 334: Co mac; 546: Co ara; 978: Co ara; 979: Co spl; 13029: Co pul – Baker, CF 28926: Co pul – Baker, MA 5626: Co asp; 6053: Co sca – Baker, RA 4: Co woo; 62: Co bra; 221: Co cur – Balcazar, J 1026: Di rur; 1117: Co ara – Balcázar, MP 49: Co ant; 181: Co pul; 278: Co pul; 297: Di str; 444: Co pul – Balderrama 347: Co sca; 397: Co ara – Balée 1996: Ch aca; 2032: Co spl – Balick 991: Co lon; 1077: Co sca; 2005: Co pul; 2702: Co pul; 3105: Co pul; 3336: Co pul; 3510: Co pul –Balslev 10592: Co sca; 10609: Co asp; 2413: Co sca; 2810: Di str; 4554: Co cha; 62239: Co sca; 62240: Co kun; 62351: Di str; 84411: Co las; 84412: Co sca; 84514: Co sca; 84629: Co las; 84755: Co ara; 97001: Co ama; 97030: Di str; 97147: Co ama; 97301: Co ara; 97318: Co las; 97390: Co las – Bamps 5196: Co las; 5237: Co spl – Bang 912: Co vag; 1248p.p.: Co ara & Co sca; 2058: Di arg – Bangham 384: Co vil; 385: Co mac; 386: Di str – Barbosa Castillo 5267: Co sca; 5394: Co mac; 6366: Di str; 6367: Co lim; 6978: Co las; 8172: Co spl; 8205: Co cha – Barbour 2691: Co gua; 4334: Co sca; 4350: Co asp; 4781: Co acr; 5060: Di app; 5339: Co sca; 5448: Co acr – Barclay, AS 370: Co all – Barclay, HG 4733: Co kru; 4909: Co sca; 6856: Co sca – Barfod 41002: Co kun; 41012: Co pul; 41457: Co cor; 41585: Co cor; 41680: Co mac; 48093: Co lim; 48097: Co kun; 48146: Co sca; 48860: Co kun; 48907: Co kun; 48917: Co cor – Barkley 38C14: Co spl; 38C180: Co ant; 18C323: Co lim; 18C324: Di str – Barlow 26-22: Co pul; 30-153: Co pul; 30-175: Co pul – Barnard 174: Co ara; 195: Co sca; 199: Co mac – Barona 3948: Co mac; 3968: Co sca – Barrera 102: Co sca – Barreto 1020: Co ama – Barrier 282A: Co sca: 349: Co asp; 2566: Ch cur; 2614: Co ara – Barringer 2007: Co woo; 2438: Co pul; 2439: Co lim; 2447: Co mon; 3826: Co mon; 3938: Co wil – Barrios 847: Co pic – Barroso 124665: Ch sub – Bartlett, AW 8690: Co ara – Bartlett, HH 11512: Co mac; 16401: Co vil; 16435: Co mac; 16459: Co pul; 16689: Co las; 16700: Co pul; 16939: Co las – Bass 58: Co pul – Bates 1577: Co pic; 180: Co lae; 211: Co lim – Battjes 780: Co las; 781: Co sca; 783: Co las – Bayly 196: Co pic – Beach 1543: Co bar – Beck, HT 1775: Co kun – Beck, SG 440: Co vag; 558: Co sca; 559: Co gua; 1449: Co com; 1511: Co com; 1695: Co sca; 2241: Co vag; 4060: Co vag; 5172: Co ara; 6063: Co vag; 6866: Co sin; 6867: Co vag; 6868: Co sca; 7320: Co coc; 7324: Co bec; 8023: Ch aca; 8230: Co ara; 9983: Co ara 13210: Co ara; 16450: Co ara; 19191: Ch aca; 19620: Co sca; 21699: Di arg; 21706: Co sin; 25499: Co vag; 29090: Co sca; 29929: Co com; 30147: Co gua; 33048: Co bec; 36329: Co gua – Bélanger 761: Co luc – Belém 2274: Ch sub – Béliz 168: Co wil – Bell 141: Co acr; 307: Co sca – Bello Carranza 961: Co cur; 1193: Co mal – Bello 557: Co ara – Beltrán Cuartas 78: Co ant – Beltrán M 3: Co sca – Beltrán 517: Co sca; 643: Co cha; 3252: Co mac; 5655: Co ara; 5669: Di app – Belway 8735: Co sca – Benavides-Guzmán 18: Co kun – Bénitez de Rojas 321: Co spl; 647: Co vil; 831: Co ara – Bennett 3458: Co sca; 4016: Di str; 4249: Co sca; 4253: Co lon; 4409: Co sca – Benoist 534: Co cla; 673: Co spl; 875: Ch con – Berendsohn 183: Di str; 184: Co pic; 185: Co pic; JBL-694: Co com – Berg 366: Di str; 473: Co las; 779: Co ert; 1059: Co sca; 1108: Co lon; P 18519: Ch aca; P 19813: Co sca – Berlin 191: Co sca; 365: Co asp; 400: Co sca; 465: Co sca; 813: Co ert; 835: Di rur; 879: Co sca; 923: Co sca; 1523: Co ama; 1671: Co sca; 1943: Co ert; 1997: Co ama; 3582: Co ert; 3589: Co sca – Bermudez 168: Co vil – Bernacci 210: Co spl; 986: Co spl; 1709: Co spl – Bernal, R 970: Co plo; 1182: Di str; 2222: Co pul; 2415: Di str; 3414: Di str; 4361: Co las – Bernal, W 25: Co vil – Bernal M 442: Co cha; 470: Co kru – Bernardi 365: Co ara; 827: Co sca; 1074: Di str; 20600: Co ara; 6894: Co vil; 7951: Co ara; 7989: Co sca – Bernoulli 855: Co pic; 857: Co com – Berríos 175: Co sca – Berry 3863: Co vil; 5131: Co spl – Berthoud-Coulon 57: Co spl; 58: Co ara – Berton, 60: Co spl – Besse 64: Co asp; 1788: Co sca – Betancur B 503: Co spl; 710: Co spl; 771: Co las; 1190: Co sca; 1360: Di str; 1391: Co spl; 1465: Co sca; 1623: Di str; 1885: Co cha; 2355: Co com; 3079: Co pul; 4155: Co spl; 5970: Co ant; 5976: Di cry; 5984: Di str; 6006: Co kun; 6007: Co las; 6049: Co kun; 6130: Co las; 6377: Co spl; 6481: Co sca; 7307: Di str; 7500: Co ant; 7712: Co com; 7729: Co vil; 7769: Co com; 7895: Co kun; 7905: Co las; 7936: Co lim; 7957: Co sca; 8234: Di str; 8267: Co ant; 8438: Co sca; 8439: Co vil; 8451: Co lim; 8753: Co com; 8753a: Co erp; 8794: Di str; 8836: Di str; 8842: Co spl; 8964: Di str; 8997: Di str; 9023: Co com; 9054: Co sca; 9175: Di str; 9189: Co com; 9431: Co kun; 9717: Co spl; 10021: Co spl; 10022: Co mac; 10026: Co kru; 10351A: Co erp; 10376: Co com; 10377: Co erp; 11505: Co com; 11533: Co spl; 11650: Co spl; 11755: Co erp; 11771: Co com; 12180: Co sca; 12419: Co plo; 12424: Di cry; 12474: Co pul; 12506: Co kun; 12507: Co mac; 14162: Co erp; 14192: Co vil; 15284: Co ant; 16699: Co lae; 16830: Di cry; 18763: Co sca; 18776: Co spl; 19127: Co spl 9023: Co com; 9054: Co sca; 11621: Co erp; 11714: Co woo; 12506: Co kun; 14570: Co plo; 15587: Co mac; 16507: Co ant; 16651: Co ant; 17971: Co lae; 17987: Co kun; 18051: Co plo; 18117: Co pul; 18641: Co spl – Betancur J 1253: Co woo – Bhikhi 1239: Ch lan lan – Bierens de Haan 122: Co spl – Bigio 1182: Ch lan lan; 1560: Co spl; 1581: Ch sub – Bilby 33: Co pra – Billiet 1869: Co cla; 1883: Co sca; 1937: Co sca; 5727: Co ert; 5745: Co sca; 6197: Co com; 6867: Di str – Black, C R-3152: Co pul; R-3153: Co pul; R-3221: Di cry; R-3321: Co plo – Black, GA 46-15: Co lon; 46-77: Ch fra; 55-1802: Ch sub; 50-8625: Co ara; 50-9118: Co ara; 51-12850: Co ara; 51-13430: Co ara; 51-13939: Co sca; 52-14582A: Co ara; 52-15150: Co ara; 52-15433: Co ara; 54-16300: Co ara; 54-16775: Co ara; 54-17565: Co ara; 56-18837: Co las; 57-19058a: Co ara – Blackmore 4089: Co pul; 4197: Co sca – Blake 7723: Co pul – Blanc 96500: Di str – Blanchet 3: Ch cus; 1494: Co ara; 1537: Co spl; 2987: Co sca; 2988p.p.: Co sca & Co spl; 2991: Co spl – Blanco 237: Co nit – Blasco 1791: Co cha – Blasido 103: Co las; 110: Co jur; 111: Co zin – Blohm 14: Co sca – Blum 1384: Di str; 1507: Co pul; 2294: Co las; 2456: Co vil; 2501: Co all; 2507: Co woo – Boeke 1009: Co ant; 1180: Di rur – Boerboom LBB 8724: Co ara – Bohlin 1197: Co pul; 1226: Co gib; 1236: Co vil – Boldingh 3801: Co sca – Bolotin 11: Co cha – Bond 154: Co sca – Bono 4902: Co vil; 4929: Co vil – Bonpland 13: Co sca – Boom, BK 2292: Ch cus; 2295: Co mal – Boom, BM 2545: Co pul; 4168: Co ara; 7333: Co spl; 8433: Co sca – Boon 1198: Co ara – Boone 984: Ch cus – Borchsenius 61: Co kun – Bordenave 1243: Co ert; 7606: Ch con; 7608: Co sca; 7908: Co sca; 8160: Co sca; 8325: Co cla; 8387: Co cla; 8402: Ch con; 8514: Di rur – Borgo 580: Co ara – Borsboom LBB 12284: Co sca – Botero-B 1278: Co sca; 1301: Co ara – Botina-P 306: Co kru; 368: Co sca; 389: Co kun; 686: Co sca; 687: Co kun – Bourdy 2883: Co spl; 2924: Co sca; 2934: Co ert – Bourgeau 2098p.p.: Co pic & Co sca –Boyle 2556: Co alt; 3719: Di str – Braga 3214: Co las – Brahe 18C-705: Co pul – Brand Meza 26: Co sca; 258: Co pul; 332: Co pul; 352: Co all; 1055: Co lim – Brandbyge 30009: Co asp; 30119: Co lon; 30154: Co asp; 30196: Di str; 30255: Co asp; 30451: Co cha; 31696: Co ara; 31808: Co las; 31891: Co sca; 31924: Co las; 32024: Co sca; 32063: Di str; 32245: Co sca; 32488: Co sca; 32760: Di str; 33730: Co ara; 36141: Co las – Brant 1277: Di str; 1664: Co lim – Brasil 2: Co sca; 202: Co sca; 411: Ch lan lan –Breckon 2024: Co pul – Breedlove 9988: Co pul; 25587: Co pic; 25962: Co pic; 26463: Co mol; 26523: Co pul; 35003: Co pul; 47330: Co pul; 51194: Co pic – Brenes 149: Co sca; 761: Co pul; 3560: Co sca; 4165: Co sca; 4312: Co pul; 4420: Co kun; 5413: Co mon; 12676: Co sca; 12677: Co pul; 19222: Co pul; 20320: Co pul; 21279: Co sca; 21940: Co mon; 21941: Co mon; 22311: Co sca; 22606: Co mon; 22607: Co mon; 23066: Co mac – Breteler 4511: Co mac – Briceño 27: Co sca – Bristan 576: Co las; 1003: Co gla; 1020: Co vil; 1160: Co sca; 1344: Co pul; 1358: Co pul – Britton 679: Co sca; 2825: Co sca; 5433: Co pul; 6784: Co pul; 7425: Co pul – Broadway 478: Co sca; 698: Co sca; 924: Co spl; 1886: Co sca; 4377: Co sca; 7427: Co sca; TRIN 8225: Co luc; TRIN8584: Co sca; 8690: Co sca; 9123: Co mac; 9271: Co sca – Brown, CA 17346: Co kun – Brown, M 138: Di str – Brummitt 19327: Di arg – Buchtien 435: Co sca; 438: Co sin; 2249: Co sca; 3687: Co gua; 3688: Co ara; 5364: Co gua – Buenaño 208: Co lim; 334: Co pul – Bunting 523: Co lim; 578: Di str; 587: Co pul; 613: Co sca; 614: Di str; 681: Co pul; 834: Co pul; 892: Co pul; 2288: Co com; 2375: Co spl; 2713: Co sca; 3359: Co vil; 3360: Co spl; 3361: Co spl; 3495: Co spl; 3521: Co spl; 3556: Co mac; 3557: Co spl; 3624: Co ara; 3822: Co spl; 6269: Co spl; 6703: Co spl; 7284: Co spl; 7347: Co mac; 7537: Co ara; 7746: Co mac; 9405: Co pul; 9429: Co spl; 9451: Co pul; 9542: Co sca 10833: Co spl; 11002: Co mac; 11124: Co spl; 11134: Co sca; 11229: Co lim; 11526: Co spl; 11626: Co mac; 11633: Co vil; 12301: Co pul; 12318: Co pul; 12388: Co pul; 12390: Co lim; 12763: Co pul; 12792: Co lim; 13343: Co spl – Burchell 442: Co spl; 825A: Ch sub; 3068: Co ara; 3278: Co ara; 6272: Ch sub; 6365: Ch sub; 8690: Co spl; 9884: Co ara; 9937: Co spl – Burger 3880: Co pul; 4231: Co mal; 4233: Co mal; 4943: Co pul; 5125: Co pul; 5377: Co cur; 5377A: Co bar; 5426: Co kun; 5488: Co kun; 5566: Co pul; 5893: Co sca; 5893A: Co pul; 6279: Co pul; 6903: Co lim; 7004: Co mal; 7649: Co wil; 7790: Co sca; 8022: Co mal; 8095: Co sca; 8443: Co kun; 8607: Co bar; 8873: Co pul; 8975: Co kun; 9595: Co mon; 9656: Co wil; 9720: Co wil; 9853: Co mal; 10050: Co pul; 10134: Co ste; 10163: Co pul; 10359: Co woo; 10395: Co kun; 10458: Co lim; 10873: Co woo; 11052: Co mon; 11605: Co sin; 11694: Co mal; 11767: Co pul; 11973: Co mon – Buscaloni 1918: Co ara – Busey 348: Co mac; 868: Co las; 890: Co sca – BW (Boschwezen Suriname) 728: Co ara; 889: Co ara; 2218: Co ara; 2527: Co sca; 2749: Co sca; 3348: Co sca; 5595: Co sca; 6347: Ch con; 6532: Co cla.

Caballero 39: Co vil – Cabrera R 3271: Co sca; 4362: Co sca – Cabrera S 25: Co pul; 3338: Co kru; 3504: Co cha; 17605: Co ant – Cadamuro 468: Co spl – Calatayud Hermosa 3820: Co gua – Calderón, CE 2986: Co pra – Calderón, S 814: Co pic; 1308: Co pic; 1340: Co pic; 1762: Co pic; 1763: Co pic; 1764: Co pic – Calderón, Y 131: Co pul – Calderón de Rzedowski 2129: Co sca – Callejas Posada 10165: Co las; 13: Co las; 189: Co kun; 194: Di str; 756: Co ant; 2707: Co pul; 3437: Co las; 3585: Co kun; 4036: Co ant; 4124: Co las; 4130: Co pul; 4276: Co las; 4317: Co las; 4420: Co mac; 4463: Co ant; 4513: Co pul; 4689: Co vil; 4871: Co kun; 5088: Co osa; 5164: Co pul; 5494: Co plo; 5704: Co osa; 6545: Co pul; 6639: Co pul; 6647: Co pul; 6674: Co ant; 6991: Co ara; 8033: Co lim; 8209: Di str; 8777: Co las; 8792: Co mac; 8853: Di str; 9092: Co las; 9250: Di str – Calzada 233: Co sca; 340: Co dir; 2455: Co pul; 3262: Co pul; 5959: Co pul – Cámara-Leret 1774: Co sca – Camargo, AF SP 54193: Co ara – Camargo, C 30: Co spl; 31: Co ara; 81: Co ara – Camargo-G 1152: Co sca; 1175: Co kru – Camp E 1504: Co ama; E 2395: Co ant; E 3607: Co mac; E 3702p.p.: Co geo & Co pul – E 3722: Co vil – Campaña 23: Co kun; 26: Co pul – Campbell, DG Co sca 8951: Co pra; P 21204: Co sca – P 22488: Co ara –Campos, MTVA 827: Ch lan lan; 839: Co sca – Campos, S 203: Co pul – Campos de la Cruz 285: Di rur; 2987: Co lae; 3030: Co lae; 3189: Co lae; 3378: Co lae; 4047: Co lae; 4754: Co lae; 4904: Co lae – Cantonspark Baarn 40: Co ara – Carabot C 159: Co spl; 160: Co ara; 161: Co mac – Caranqui 92: Co sca; 409: Co asp; 800: Co sca – Carauta 2508: Co ara – Carballo 95: Co las; 96: Co bra – Carbonó 95: Co las; 96: Co bra; 418: Co sca; 419: Co sca; 524: Co pul; 575: Co vil; 580: Co mac – Cárdenas Hermosa 1165A: Di arg; 1851: Co sin; 1882: Di rur; 7368: Co com; 7369: Co sin – Cárdenas López 575: Co pul; 666: Co kun; 813: Co pul; 1014: Co sca; 1069: Co sca; 1238: Co sca; 1507: Co kun; 1531: Co mac; 1544: Co sca; 1615: Co sca; 1841: Co all; 1945: Co kun; 2190: Co sca; 2261: Co kun; 2725: Co las; 2734: Co mac; 2802: Di str; 3100: Co mac; 4273: Co cha; 6585: Co ara; 6602: Co spl; 7264: Co las; 7318: Co spl; 7322: Co sca; 8311: Co las; 9626: Co sca; 9991: Co las; 9994: Co sca; 10007: Co kru; 11706: Co sca; 11994: Co las; 12013: Co sca; 12055: Co sca; 12060: Di str; 12174: Co sca; 13181: Co sca; 13434: Co aspl; 13436: Di tr; 13524: Co com; 13723: Co spl; 13911: Co spl; 14803: Co spl; 14953: Co spl; 15491: Co sca; 16384: Co spl; 16746: Co spl; 18699: Co sca; 18767: Co sca; 18781: Co kru; 18785: Co mac; 20634: Co spl; 20774: Co spl; 21024: Co spl; 21110: Co spl; 21154: Co sca; 21157: Co mac; 21454: Co spl; 21595: Co ara; 21813: Co com; 21827: Co spl; 22458: Co sca; 22460: Co lon; 22806: Co spl; 23550: Co sca; 24594: Co spl; 24727: Co asp; 24780: Co cha; 24802: Co sca; 24822: Co spl; 24919: Co cha; 24928: Co cha; 40204: Di str; 40229: Di str; 40546: Co cha; 40558: Co cha; 40601: Co cha; 40685: Co cha; 40926: Co cha; 41458: Co cha; 41495: Co asp; 41574: Co sca; 41619: Di str; 41858: Co asp; 41875: Co cha; 41875A: Co sca; 41881: Co sca; 42090: Co asp; 42101: Co cha; 42104: Di str; 42112: Co asp; 42116: Di str; 42801: Co spl; 42803: Co sca; 43305: Co las; 43344: Co cha; 43475: Co asp; 43564: Co acr; 44386: Co cha; 44520: Co sca; 45377: Co sca; 45564: Co cha; 45684: Co cha; 45774: Co sin; 45778: Co ama; 46183: Co ama; 46237: Co cha; 46360: Co cha; 46466: Co spl; 46650: Co spl; 46691: Di str; 47126: Co spl; 47551: Co spl; 47562: Co ara; 47614: Co sca; 47781: Co kru; 48030: Co cha; 48297: Co sca; 48343: Co kru; 48417: Co sca; 48447: Co sca; 48491: Co cha; 48566: Co spl – Cárdenas-Henao 8: Co pul – Cardiel 112: Co vil; 174: Co sca; 263: Co ara – Cardona, F 983: Co ara; 1703: Co sca; 1748: Di str; 1829: Co vil; 1830: Co pul – Cardona, ME 83: Co ant – Cardona-O 48: Co ara – Carleton 50: Co pul – Carlson 2116: Co pul; 3289: Co pul – Carmona 112: Co ant –Carpenter 337: Co pul Carrasco-Llanes 206: Co sca – Carrasquilla 1325: Co kun; 1756: Co vil – Carreira 315: Co pra – Carrillo, L 192: Co kru; 288: Co erp; 299: Co lon; 471b: Co lon; 484: Co ara; 735: Co sca; 746: Co asp – Carrillo, Y 1: Co sca; 1a: Co ama – Carrington 2474: Co sca – Carrión 326: Ch aca; 506: Ch aca – Carroll R 2247: Co kun; R 2959: Co alo – Carvajal Ugalde 158: Co mon; 191: Co wil; 197: Co mon – Carvajal 1A: Co pul; 34A: Co kun – Carvalho 7177: Co spl – Castaño 5: Co pul; 83: Co las; 307: Co pul – Castaño Arboleda 77: Di cry; 189: Co cha; 198: Co cha; 1539: Co spl; 1614: Co mac; 1615: Co las; 1739: Co mac; 2306: Co spl; 2419: Co sca; 2456: Co spl; 2659: Co spl; 2838: Co ara; 2849: Co sca; 3123: Co cha; 3202: Co spl; 3293: Co cha; 3309: Co las; 3324: Co mac; 3380: Co sca; 3624: Co pul; 4388: Co ara; 7645: Co sca; 7650: Co ama; 7800: Co cha – Castellanos, A 119: Co pul – Castellanos, C 226: Co spl – Castillo, A 1343: Co spl; 1355: Co mac; 2231: Co spl; 2396: Co spl; 3385: Co ara; 3634: Co ara – Castillo, GA 642: Co rub – Castillo, JA 6973: Co ara – Castillo, ME 33: Co vil – Castillo C 335: Co pul – Castro, F 5111: Co ara; 5398: Co mac; 5779: Co las; 5882: Co com; 10913: Co cha – Castro, M 19: Co ara; 60: Ch aca 94: Co ara – Castro, S: 552: Co fis – Castro, SY 900: Co cha – Castro, W 66: Ch lan lan; 81: Co pra – Castro C 2328: Co pul – Castro Hernández 48: Co ric – Castro T 203: Co pul – Castroviejo CHIN 308: Co sca; 7178: Co lim; 7182: Di str; 13161: Co lim; 16085: Co mac – Cavalcante 1: Co ara; 540: Co pra; 821: Co ara; 1952: Co sca; 2384: Co ara – Cavalcanti 2137: Co spl – Cayola Pérez 451A: Co web; 1633: Co vag – Cazalet 5088: Co pul – Cazali 3362: Co pul – Cediel, J 40: Co las – Cediel, JL 1: Co ant – Cedillo Trigos 2391: Co pul – Centeno 7: Co pul – Cerón M 670: Co ama; 742: Di str; 809: Cozam; 857: Co sca; 1406: Di str; 1605: Co asp; 2088: Co sca; 2135: Co kun; 2332: Co sca; 2487: Co sca; 2667: Co lon; 3101: Co lon; 3187: Co ama; 3195: Co sca; 3686: Co lon; 4217: Co sca; 5131: Co sca; 6209: Co pul; 9650: Di str; 9779: Co ara 14298: Co ama; 16198: Co sca; 17577: Co sca; 18255: Co vil; 18323: Co pul; 18331: Co vil; 20136: Co mac; 20481: Co vil; 20528: Co vil; 20628: Co vil; 20775: Co sca; 30858: Co kun – Chacón Gamboa 29: Co mac; 41: Co gla; 46: Di str; 137: Co bra; 208: Co kun; 296: Co bra; 399: Co wil; 567: Co mal; 584: Co bra; 846: Co bra; 944: Co mal; 991: Co pul; 2079: Co pul – Chaix 35: Co spl – Chama 4277: Co jur; 4762: Co jur – Chambi 1098: Co chm – Chanderbali 295: Co sca – Chanek 67: Co pul – Chaparro 9: Co erp; 25: Co las; 71: Co las – Chapman R 3014: Co lim – Chasiluisa 5: Co kru – Chatrou 46: Co sca; 154: Co sca – Chavarría Díaz 531: Co wil – Chavarría García 50: Co woo; 289: Co mac – Chaves Chaves 352: Co wil – Chávez A, E 94: Co ert – Chávez A, R 1737: Co com – Cházaro Basáñez 369: Co sca; 3537: Co pul – Cherubini 19: Co spl – Chickering 3: Co pul; 18: Co pul; 25: Co mac – Chinchilla 79: Co pul; 168: Co pul – Chocce 408: Co ara; 437: Co ara – Chorley 90: Co pul – Christenhusz 1963: Co lae; 2084: Mo uni; 2300: Co spl; 5475: Co pul – Christensen 72504: Ch fra – Churchill 4156: Co sca; 4310: Co pul; 4486: Co cur; 4593: Co cur; 4710: Co cur; 5135: Co pul; 5138: Co kun; 5405: Co cur; 5740: Co vil; 5837: Co cur; 6253: Co pul – Cid Ferreira 1175: Co sca; 1314: Co ara; 2591: Co sca; 2806: Ch aca; 2874: Ch lan lan; 3273: Co sca; 4089: Co ert; 5182: Co kru; 5183: Co sca; 5437: Co pra; 5736: Co sca; 5832: Co ara; 5834: Co ara; 8643: Ch lan lan – Clark, JL 2707: Co pul; 3045: Co lim; 3865: Co pul; 3938: Co kun; 3967: Co lae; 4142: Co mac; 4172: Co pul; 4266: Co geo; 4364: Co pul; 4655: Co gib; 7221: Co lon; 8992: Co zam; 13045: Co ant; 13241: Di cry –Clarke 163: Co sca; 658: Co sca; 1105: Co sca; 1311: Co sca; 1754: Co ara; 1759: Co ara; 1872: Co sca; 2609: Co sca; 3391: Co sca; 3453: Co sca; 3882: Co sca; 4422: Co ara; 4959: Co spl; 6009: Co spl; 6629: Co ara; 6719: Co ara; 8032: Co spl; 8074: Co sca; 10076: Co jur; 11172: Co sca; 12169: Co spl – Claussen 180: Ch sub – Clavijo-R 220: Co sca; 415: Co fis; 553: Ch fra; 715: Co kru; 984: Co las; 1052: Co kru; 1339: Co spl; 1504: Co spl – Clewell 3228: Co kun; 3233: Di str; 4058: Co sca; 4115: Co lim; 4346: Co sca; 4562: Co pul – Cocucci 3188: Co ara – Coêlho, DF 437: Co ara; 697: Co sca – Coêlho, L 2: Ch lan lan; 275: Co spl; 1813: Co acr – Coello H 176: Co sca; 249: Co sca – Cogollo Pacheco 155: Co las; 344: Co ant; 533: Di str; 541: Co kun; 581: Di str; 1500: Di str; 1618: Di str; 1624: Co kun; 2710: Co plo; 2768: Co pul; 2801: Co cor; 4178: Co kun; 4923: Co lim; 5027: Co pul; 9050: Co cor – Colella 696: Co vil; 1399: Co sca – Coley 8246: Co sca – Collaguazo 1407: Co pul Collazos 197: Co spl; 400: Di str – Collins R 3083: Co vag – R 3282: Co vaz – Colorado 612: Co com; 752: Co sca – Colque 470: Di rur – Conrad 2884: Co pic – Conreras-H 2828: Di str – Constantino R 3232: Co plo – Contreras, E 937: Co pul; 2259: Co pul; 2708: Co mac; 3525: Co pul – Conzatti 6: Co sca – Cook, MT 16: Co pul – Cook, OF 14: Co pul; 308: Co mac; 363: Co pul; 596: Co sep – Coradin 7396: Ch sub; 7397: Ch sub – Cordeiro 1078: Ch aca; 1209: Co spl; 1234: Co ara – Cordero 58: Co pli; 96: Co kun; 156: Co ric – Cordero-P 50: Co ant; 91: Co ant; 139: Co spl; 412: Co sca; 578: Di str; 764: Co ara – Córdoba, MP 372: Co sca; 1553: Co sca – Córdoba, WA 406: Co lim – Cornejo Valverde 2670: Co bec; 2674: Co sca; 3346: Co jur; 3357: Co las; 3365: Co jur – Cornejo 425: Co gib; 1562: Co asp; 1808: Co mac; 1809: Co vil; 2988: Co pul; 3603: Co lim; 3971: Co zam; 5231: Co pul; 5507: Co lim; 5792: Co gib – Coronado González 235: Co pul; 256: Co mac; 827: Co sca; 897: Co sca; 943: Co lim; 952: Co pul; 1098: Co pul; 1365: Co lim; 1398: Co sca; 1621: Co sca; 1720: Co pul; 1749: Co lim; 1828: Co pul; 1934: Co kun; 1936: Co pul; 2040: Co pul; 2087: Co pul; 2116: Co kun; 2199: Co pul; 2279: Co pul; 2644: Co pul; 2662: Co pul; 3021: Co pul; 3243: Co pul; 3959: Co pul; 4404: Co pul; 4470: Co lim; 4488: Co pul; 4558: Co sca; 4756: Co kun; 4778: Co lim; 4913: Co pul; 4934: Co kun; 5107: Co sca; 5503: Co sca; 6707: Co sca; 7193: Co sca – Correa, E 1841: Co las – Correa, J 495: Di str – Correa, M 13: Co spl; 2285: Di str; 2603: Co ant; 2604: Co pul; 2703: Co las; 2776: Co cha; 2835: Co sca; 2904: Di str; 3007: Co kru – Correa A 1646: Co las; 1760: Co kun; 2883: Co sin; 3005: Co woo; 9682: Co pul; 9713: Co pul – Correll 12255: Co vil – Cortés 1860: Co las; 2148: Di str; 2700: Co ara – Cosentino 144: Co pul – Costa, LV 26534: Ch cus – Costa, MAS 73: Ch man; 537: Co ara; 568: Ch man; 589: Co ara – Cotton 11: Co sca; 47: Co sca – Cowan, CP 2990: Co pul; 3339: Co pul – Cowan, RS 38752: Co ert – Cowell 53: Di str; 77: Co kun; 351: Co sca – Cox 1358: Co com – Cremers 4445: Co sca; 4516: Co spl; 4566: Co cla; 5684: Co cla; 6181: Ch lan pul; 6185: Co spl; 6351: Co sca; 6366: Co ert; 6367: Co las; 6390: Co ert; 6430: Co cla; 6604: Co sca; 7042: Co spl; 7063: Ch con; 7332: Co spl; 7591: Ch con; 7659: Co ara; 8595: Ch con; 9217: Co pul; 9222: Co pul; 10497: Co ara; 10779: Co ert; 11272: Co sca; 11430: Co ert; 11503: Co ert; 11528: Co ert; 11734: Co ert; 11805: Ch con; 11931: Ch con; 11960: Co ert; 12183: Co spl; 12184: Co ert; 12200: Co spl; 12437: Ch con; 13167: Co ert; 13186: Ch con; 14043: Ch lan pul; 14497: Co cla; 14498: Co ert; 14592: Co ert; 14603: Ch lan pul; 15145: Ch con; 15147: Co ert; 15162: Co cla; 15339: Co sca – Croat 329: Co mon; 330: Co pul; 332: Co pul; 4129: Co mac; 4145: Co sca; 4295: Co kun; 4297: Co kun; 4374: Co pul; 4746: Di str; 5100: Di str; 5257: Di str; 5262: Co sca; 6388: Co mac; 6526: Co pul; 6543: Co pul; 6547: Co pul; 6598: Di str; 6644: Co pul; 6668: Co pul; 7209: Di str; 7701: Di str; 8321: Co pul; 8635: Co gla; 8711: Co pul; 9830: Co woo; 9857: Co woo; 9900: Co kun; 10026: Di str; 100333: Co pul; 100636: Co sca; 10072: Co woo; 101917: Co ara; 101953: Co sca; 101995: Co sca; 102037: Co cla; 102266: Co sca; 102296: Co ara; 102300: Co sca; 102367: Co cla; 102559: Co ert; 102665: Co spl; 102964: Co ert; 10300: Co gla; 103470: Co spl; 10727: Co vil; 10881: Co sca; 10907: Co pul; 10993: Co kun; 11115A: Co gla; 11251: Co all; 11266: Co kun; 11330: Co pul; 11354: Co woo; 11397: Co all; 11424: Co mac; 11436: Co pul; 11439: Co kun; 11455: Co kun; 11490: Co vil; 11522: Co pul; 11549: Co las; 11560: Co kun; 11586: Co kun; 11590: Co sca; 11712: Di str; 11714: Di str; 11722: Co vil; 11794: Co pul; 12233: Co kun; 12233A: Co pul; 12446: Co mac; 13337: Co cur; 14167: Co woo; 14284: Co pul; 14310: Co cur; 14868: Co sca; 14935: Co sca; 15135: Co vil; 15191: Co kun; 15261: Co all; 15318: Co all; 15529: Co sca; 15534: Di str; 15538: Co mac; 15544: Co sca; 15586: Co kun; 16198: Co all; 16471: Co nit; 16718: Co sca; 16786: Co kun; 16817: Co all; 16881: Co sca; 16898: Co woo; 16922: Co sca; 17291: Co kun; 17314: Co las; 17485: Co ara; 17569: Co sca; 17636: Co sca; 17738: Co sca; 17979: Di rur; 17994A: Di rur; 18033A: Co sca; 18143: Co sca; 18154: Co sca; 18168: Co erc; 18216: Co erc; 18260: Co sca; 18747: Co erc; 18775A: Co sca; 18816: Co sca; 19069: Co ara; 19097: Co sca; 19194: Co sca; 19334: Co sca; 19409: Di app; 19481: Di app; 19502: Co sca; 19559: Co erc; 19683: Co las; 19762: Co las; 19868: Co las; 19923: Co sca; 20242A: Co sca; 20336: Co ara; 20349: Di app; 20577: Co sca; 20667: Co las; 20850: Co sca; 20984: Co sca; 21742: Co mac; 22216: Co lim; 22341: Co kun; 22489: Co pli; 22493C: Co las; 22722: Co sca; 23319: Co pul; 23392: Co pul; 23544: Co pul; 23787: Co pic; 23876: Co pul; 24279: Co pul; 24474: Co pul; 25099: Co sca; 25115: Co all; 25312: Co cur; 25377: Co kun; 25494: Co kun; 25497: Co pul; 25524: Co pli; 25606: Co cur; 25622: Co all; 25924: Co all; 26156: Co vin; 26168: Co kun; 26210: Co all; 26579: Co wil; 26634: Co wil; 26679: Co wil; 26797: Co wil; 27055: Co sca; 27382: Co all; 27455: Co alo; 27474: Co kun; 27535: Co pli; 27552: Co all; 33348: Co cur; 33741: Co sca; 34634: Co sca; 34727: Co sca; 35081: Co pul; 35289: Co ric; 35306: Co lim; 35358: Co las; 35455: Co all; 35683: Co bra; 35746: Co mal; 35896: Co las; 35911: Co all; 36501: Co sca; 36534: Co pul; 36569: Co lim; 36583: Co cur; 36674: Co pul; 36707: Co wil; 36719: Co wil; 36872: Co woo; 37213: Co com; 37371: Co cur; 37393: Co kun; 37522: Co pul; 37554: Co all; 37583: Co vil; 37671: Co pul; 37734: Co kun; 38072: Co vil; 38104: Co nit; 38223: Co lim; 38292: Co mac; 39253: Co pic; 39701: Co pul; 39742: Co sep; 39765A: Co pul; 40042: Co pul; 40154: Co pic; 40172: Co pul; 40265: Co pul; 40332: Co pul; 41599: Co pul; 41630: Co pul; 42058: Co pic; 42085: Co pic; 42123: Co pic; 42590: Co pul; 42649: Co pul; 43166: Co woo; 43273A: Co lim; 43557: Co pul; 43822: Co pic; 44269: Co mal; 45265: Co pic; 45886: Co pic; 47106: Co mon; 48708A: Co com; 48708B: Co com; 49021: Co pul; 52086: Di cry; 52287: Co las; 54235: Co spl; 54700: Co pul; 55699: Co pul; 56061: Co all; 56062: Co cor; 56143: Co las; 56831: Co spl; 57390: Co cor; 57722: Co lae; 58585: Co sca; 58641: Co asp; 58697: Co asp; 58987: Co sca; 59103: Co asp; 59575: Co sca; 59711: Co ric; 59827: Co ric; 59880: Co las; 59981: Co cur; 60314: Co cur; 60652: Co spl; 61461: Co pul; 61656: Co erc; 62399: Co sca; 62577: Di rur; 62667: Co sca; 62674: Co dal; 62812A: Co pul; 63201: Co pul; 66254: Co wil; 66371: Co cur; 66540: Co sin; 66544: Co com; 66570: Co pul; 66868: Co pul; 66887: Co pli; 66945: Co pul; 67141: Co las; 67215: Co cur; 67340: Co sca; 68078: Co mon; 68213: Co pul; 68507: Co spc; 68544: Co ara; 68673: Co sca; 68683: Co sca; 68787: Co cur; 68796: Co alo; 69270: Co sca; 70417A: Co leu; 70430: Co lim; 70452: Co cor; 73096: Co mac; 73479: Co asp; 74689: Co pul; 75052: Co alo; 75789: Co kun; 76530: Co pul; 76686: Co cur; 76739: Co las; 76851: Co kun; 76915: Co com; 77061: Co pul; 77073: Co mac; 77135: Co pul; 78961: Co ric; 79103: Co kun; 79224: Co osa; 79688: Co plo; 80862: Co las; 80937: Co las; 83948A: Co gib; 85008: Co dal; 85017A: Co sca; 85354: Co kru; 85455: Co ara; 85498: Co sca; 85550: Co las; 85988: Co sca; 85990: Co sin; 86087: Co sca; 86155: Co vaz; 86509: Co sca; 86720: Di rur; 89356: Co acr; 89540: Co sca; 8963: Co vil; 89678: Co sca; 91313: Co asp; 91947: Co sca; 92034: Co ant; 92485: Co zam; 92754: Co pul; 94410: Co pul; 95715: Co pul; 96181: Co vil; 96907: Co ama; 97210: Co asp; 98231: Co cha; 99581: Co sca; 99704: Co leu; 99851: Co pul; 99915: Co pul; 99969: Co pul – Croizat Chaley 291: Co mac; 292: Co sca; 399: Co mac; 948F: Co ara – Crow 6037: Co woo; 9788: Co sca – Crueger 234: Co ara – Cruz 66: Co sca – Cuadros Villalobos 1561: Co spl; 3139: Co com – Cuamacás 51: Co sca; 72: Co ama; 73: Co lon – Cuascota 260: Co zam – Cuatrecasas 3537: Co spl; 3587: Co ara; 3772: Co ara; 3831: Co spl; 4204: Co vil; 4318: Co ara; 4707: Di str; 6749: Co spl; 6766: Co spl; 8396: Co ant; 8889: Co las; 8956: Co sin; 9176: Co kru; 9201: Co cha; 10604: Co erp; 10684: Co jur; 10706: Co sca; 10777: Di str; 10951: Co cha; 11118: Di str; 12865: Co jur; 12979: Di str; 13001A: Di str; 13028: Co pul; 13142: Co spl; 13162: Co sca; 13168: Co mac; 13714: Co ant; 13736: Co sca; 14234: Co lim; 14240: Co sca; 14256: Co kun; 15010: Di str; 15039: Co pul; 15947: Co woo; 16207: Co woo; 16324: Co lim; 16681: Di str; 16829: Co lim; 17604: Di str; 17637: Co kun; 19822: Co kun; 21166: Co leu; 21175: Co kun; 21466: Co sca; 23977: Co plo; 24037: Di str; 24049: Co mac; 26131: Co pul; 27132: Co sca; 27215: Co asp –Curran 419: Co mac.

Dahlgren 1: Co sca; 139: Co spl; 141: Co spl; 389: Co ara; 740: Co sca – Daly 1556: Ch lan pul; 1589: Co sca; 1694: Co spl; 1794: Ch lan pul; 1920: Co las; 6586: Co sca; 6846: Co gua; 7195: Ch aca; 7486: Co sin; 7746: Co kru; 7855: Ch aca; 7860: Ch aca; 7967: Co sca; 7982: Ch lan lan; 8040: Co sca; 8421: Co acr; 8571: Ch lan lan; 8606: Co dal; 8761: Co sin; 9040: Co sca; 9112: Ch lan lan; 9195: Co ara; 9226: Co ara; 9399: Co ara; 9990: Co acr; 10209: Ch lan lan; 10328: Co sca; 10520: Co las; 10535: Co ara; 11074: Ch lan lan; 11086: Di rur; 11087: Co sca; 11103: Ch aca; 11113: Di rur; 11518: Co ara – Damião 2770: Co lon – Daniel, Bro 1814: Co las; 5472: Co kun; 5756: Co sca – Danin 14: Di str – D’Arcy 4017: Co woo; 6101: Di str; 6185: Co mac; 6201: Di str; 6205: Co mac; 9305: Co vil; 9397: Co mac; 9682: Co cur; 10620: Co all; 12317: Co las; 13049: Co wil; 13625: Co sca; 14614: Co pul; 14934: Co sin; 14951: Co kun; 15146: Di str; 16028: Co com; 18027: Co pul – Dávalos 11: Co lim – David 1862: Co com; 2311: Co las; 2796: Co ant; 3033: Di str; 3046: Co vil; 3287: Co las; 3382: Co lim; 3457: Co las; 3494: Co ant; 3721: Co cor; 4195: Co las; 4313: Di str; 4589: Co ant; 4590: Co vil; 5071: Co pul – Davidse 2341: Co pul; 4500: Co mac; 5720: Di str; 5721: Co cha; 9431: Co pul 12291A: Co spl; 14158: Co spl; 14204: Co spl; 15561: Co spl; 15829: Co spl; 15962: Co spl; 16301: Co sca; 16819: Co ara; 18669: Co spl; 18680: Co mac; 18681: Di str; 19129: Co sca; 21431: Co spl; 21572: Co pul; 21924: Co spl; 21960: Co mac; 24068: Co wil; 24435: Co wil; 25604: Co wil; 26578: Co spl; 27493: Di str; 27913: Co ara; 28314: Co wil; 28403: Co wil; 30877: Co pul; 31157: Co mal; 34426: Co sca; 34431: Co pul; 34528: Co sca; 35915: Co pul; 36362: Co pul; 36766: Co pul – Davidson, C 3497: Di rur; 8287: Co mal; 8330: Co bra; 8388: Co sca; 8411: Co sca; 8412: Co mal; 8566: Co bra; 8696: Co kun; 8770: Co cal; 8928: Co sca; 9167: Co com; 9264: Co sin; 9395: Co rub; 9836: Co ara; 10032: Co sca – Davidson, ME 726: Co wil – Dávila 94: Co kun – Davis, EW 105: Co spl; 418: Co kun; 470: Co ama; 825: Co sca; 839: Co las; 866: Co erc; 875: Co lon; 940: Co sca; 1028: Di str; 1029: Co erc; 1065: Co spl; 1144: Co sin – Davis, TAW 283: Ch con – Dawe 291: Ch lan lan – De Bruijn 1104: Co pul; 1440: Di str – De Granville 36: Co ara; 70: Co sca; C. 152: Co ert; C. 169: Co sca; 347: Co sca; 437: Co cla; 460: Ch con; 766: Ch lan pul; 786: Co ert; 865: Co ara; 1109: Co cla; T. 1123: Co cla; 1626: Co ert; 1711: Co sca; 1847: Co sca; 1897: Co sca; 2002: Co cla; 2143: Co sca; 2209: Co spl; 2234: Co sca; 2297: Ch con; 2351: Co ert; 3283: Ch con; 3547: Co spl; 4249: Co cla; 4352: Co spl; B. 4430: Ch con; 4533: Co ara; B. 4584: Co sca; B. 4987: Co sca; 5053: Ch con; 5161: Co sca; B. 5247: Ch cur; 5425: Ch lan pul; 5494: Co sca; 5508: Co cla; 6335: Co cla; 6482: Co sca; 6617: Co spl; 6680: Ch con; 6694: Co sca; 6983: Co spl; 7036: Co sca; 7079: Ch con; 7332: Co sca; 7477: Co cla; 7707: Co ert; 7827: Co ert; 8438: Co sca; 8456: Co ert; 8466: Co cla; 9289: Co spl; 9503: Co spl; 9734: Co spl; 10043: Co ara; 10296: Ch con; 10332: Ch con; 10472: Co ert; 10591: Co ert; 10666: Ch lan pul; 10823: Co cla; 11262: Ch con; 11478: Ch con; 11479: Co ert; 11679: Co sca; 12013: Co ara; 12014: Co sca; 12433: Co spl; 12743: Co cla; 12744: Ch con; 12853: Co spl; 12939: Co spl; 13158: Co cla; 13212: Co ert; 13252: Ch con; 13349: Co cla; 13541: Co sca; 14660: Co ara; 14927: Ch con; 15074: Co spl; 15075: Co spl; 15194: Ch lan pul; 15332: Co spl; 15697: Ch cur; 15952: Co spl; 15963: Ch con; 16130: Ch lan pul; 16396: Co spl; 1646: Co cla; 16822: Co ert; 16935: Co cla; 17242: Co ert; 17320: Co ert – De Jong 32: Co sca; 33: Co sin – De la Barra 876: Co ara – De la Cruz 1464: Co ara; 3218: Co sca; 3633: Co spl; 3788: Co kru; 4116: Co ara – De la Quintana 427: Di arg – De la Sagra 248: Co pul; 583: Co pul – De Nevers 4798: Co sca; 4908: Co bra; 5058: Co pul; 5081: Co sca; 5083: Co kun; 5393: Co pli; 5736: Co bra; 5761: Di str; 6342: Co las; 6640: Co sca; 7042: Co pul; 7658: Co vin; 7982: Co sca; 7990: Co kun; 8463: Co las – De Saint-Hilaire 13: Ch sub – Deam 18: Co mac; 111: Co pul; 6034: Co pul – Del Carpio 2449: Di rur – Delascio Chitty 2227: Co sca; 2946: Co pul; 3244: Co mac; 7220: Co sca; 7280: Co sca – Delgado, L 306: Co spl; 499: Co spl; 557: Co spl; 756: Ch lan lan – Delgado, R 11: Co sca – Delinks 377: Co pul – Delnatte 282: Co sca; 572: Co ert; 729: Ch con; 1702: Co cla; 1899: Co cla – Delprete 5249: Co vil; 5250: Co com; 5275: Co las; 7695: Co sca; 7800: Co las; 7858: Co vaz; 8196: Co acr; 8212: Co sca; 8261: Co sca; 8266: Di rur; 8268: Co sin; 8624: Co sin; 9105: Ch sub – Delucchi 546: Co ara – Den Held 12: Co sca – Denslow 2469: Co pul; 2498: Co pul; 2657: Co pul – Devia Alvarez 1058: Co lae; 2480: Co cor; 2675: Co woo –Díaz, C 17: Co kun – Díaz, J 65: Co sca; 188: Co cha; 381: Co cha; 473: Co cha – Díaz, WA 356: Co mac; 913: Co spl; 954: Co spl; 2462: Co sca – Diaz-R 313: Co sca; 414: Co pul; 480: Co las; 481: Co las – Díaz Piedrahita 1526: Co ant; 3326: Co spl; 4174: Co sca – Díaz Santibáñez 6: Co ara; 96: Co asp; 482: Co las; 809: Co mac; 868A: Di rur; 1533: Co las; 2301: Co lae; 4107: Co ert; 6856: Co ert; 7350: Co ama; 7811A: Co asp; 10567: Co sca – Dieckman 305: Co mac – Diederichs 46: Co mac; 192: Co spl – Di Giovanni 37: Co spl – Dixon 56: Co lim – Döbbeler 1607: Co pul; 1608: Co lim; 1810: Co woo; 2146: Co mon; 2147: Co wil; 3030: Co cur; 3195: Co woo; 3196: Co woo; 3204: Co pul; 5154: Co kun; 5332: Co pul; 5690: Co ric; 5728: Co mal – Dodge 3461: Di str; 5878: Co mal; 9410: Co pul; 9490: Co kun; 16640: Co mac; 16915: Di str – Dodson 913: Co ore; 1163: Co pul; 1441: Co ore; 2855: Co sca; 2915: Co erc; 4301: Co pul; 5004: Co osa; 5005: Co osa; 5186: Co pul; 5189: Co lim; 5347: Co gib; 5494: Co geo; 5730: Di str; 7349: Co kun; 7449: Co geo; 9205: Co asp; 9220: Co geo; 9799: Co geo 12212: Co pul; 12676: Di str; 13014: Co osa; 13039: Co gib; 13402: Co lae; 14524: Co kun; 14524A: Co pul; 14606: Co pul –Domínguez Peña 5: Co acr; 25: Co sca – Domínquez V 173: Co pul –Donnell Smith 1827: Co pul; 2036: Co pic; 2800: Co pic; 2801: Co com; 2802: Co com; 4973: Co pul – Doreste 71: Co spl – Dorr 5780: Co spl; 5781: Co mac; 6353: Co acr; 7098: Co spl – Dowe 91: Di rur; 93: Co woo – Dressler 208: Co pic; 1553: Co pul; 2901: Co las; 2915: Co all; 2924: Co all; 2925: Co mac; 2936: Co gla; 3661: Co all; 4080: Co cur; 4167: Co ric; 4186: Co las; 4190: Co cal; 4406: Co vin; 4506: Co bra – Drouet 2075p.p.: Co ara & Co sca – Dryander 656: Co kun; 824: Di str; 1215: Co mac; 2547: Co vil; 2588: Co sca – Dryer 442: Co bar; 773: Co mon; 1175: Co cur; 1409: Co bar; 1498: Co pul – Duarte 1277: Co spl; 1521: Co spl; 3597: Di str; 5668: Ch cus; 5678: Ch sub; 5689: Ch sub; RB 65339: Co ara; 6874: Co lon; 7144: Co acr; 7146: Co acr; 7547: Ch sub – Dubs 744: Co ara; 1991: Ch aca; 2315: Ch lan lan; 2650: Ch lan lan –Ducke MG 1402: Co spl; MG 6872: Co las; MG 10511: Co spr; MG 10735: Co ara; MG 11372: Co spl; RB 14127: Ch fra; MG 15260: Co sca; MG 15649: Ch lan pul; MG 16778: Ch fus; MG 16789: Ch con; RB 18974: Ch lan pul; RB 25622: Ch man; RB 27352: Ch lan lan – Dudley 10161: Co sin; 10366: Co lim; 11872: Co lim; 12519: Co sca – Dueñas, A 325: Co lim; 739: Co pul; 822: Co pul; 1807: Co pul; 1821: Co spl – Dueñas, H 274: Co sca; 306: Co vil; 314: Co kun – Dugand 2406: Co mac – Duke 3872: Co pul; 4147: Co pul; 4586: Co vil; 4628: Co pul; 4686: Co mac; 4750: Co mac; 4758: Co pul; 4955: Co mac; 4956: Co lim; 5084: Co pul; 5100: Co pul; 5250: Co pul; 5282: Co vil; 5302: Co las; 5433: Co pul; 5614: Co pul; 8971: Co vil; 9679: Co sca; 9778: Co lim; 9899: Co lim; 10002: Di str; 10015: Co sca; 10025: Co sca; 10092: Co pul; 10155: Co pul; 10296: Di str; 10573: Co lim; 10747: Co las; 11076: Di str; 11163: Co sca; 11190: Co sca; 11311: Co lim; 11315: Co kun; 11318: Co mac; 11450: Co las; 11647: Co vil; 11648: Co lim; 11750: Co sca; 11969: Co sca; 12105: Co las; 12137: Co kun; 12331: Co vil; 13211: Co las; 13607: Co wil; 13678: Co las; 13805: Co cur; 13970: Co all; 14018: Co pul; 14044: Co pul; 14163: Co pul; 14292: Co pul; 14367: Co las; 14812: Co sca; 15826: Co vil – Dumont 7427: Co mac – Duncan 3: Co pul – Dunn 23357: Co spc; 23530: Co kun; 23701: Co bra – Duno N 148: Co spl; N 215: Co ara – Dunstan R 3233: Co lim – Duque 2696: Co las; 5224: Co sca – Duque-Jaramillo 2080: Co sca; 2115: Co acr; 2197: Co sca; 2248: Co sca; 2249: Co acr; 2250: Co kru; 2251: Co ant; 2394: Co sca; 3912A: Co ant – Duran 28: Co vil – Durkee 75-114: Co woo; 71-140: Co sca; 71-142: Di str – Dusén 11713: Co spl; 13602: Co ara; 14596: Co spl – Duss 2109: Co spc; 2109b: Co luc; 3329: Co ara; 3701: Co spc; 3704: Co spc; 3933: Co ara; 4074: Co spc – Dwyer 1702: Co vil; 2339a: Co lim; 4232: Co mac; 4477: Co cur; 7109: Co las; 7500: Co pul; 7811: Co woo; 8258: Co kun; 8692: Co kun; 8853: Co cur; 8950: Co kun; 8966: Co cur; 9776: Co lim 10232: Co pul; 10290: Di str; 10543: Co las; 10557: Co las; 10857: Co pul; 10949: Co pul; 12098: Co pul; 12525: Co pul; 15185: Co pul.

Ebinger 221: Di str; 234: Co mac; 239: Co all; 320: Co vil; 339: Co las; 557: Co sca; 560: Co pul; 64: Co pul – Echeverría, JA 49: Co woo; 1319: Co mon – Echeverry E 2023: Co ara; 2031: Co spl – Edwards 22p.p.: Co pul; 627p.p.: Co pic & Co pul – Egler 317: Co ara; 45952: Co spl – Ehrich 1c: Ch lan lan; 2c: Co sca; 3c: Co ara; 4c: Co sin; 5c: Co pra; 6c: Co mac; 7c: Co ara; 8c: Ch aca; 9c: Co acr – Ehringhaus 74: Co sca – Eilers 1022: Co pul – Eiten 257: Ch fus; 4728: Co ara; 6071: Co spl; 7861: Co spl; 8036: Co ara; 9689: Co ara – Ek 1323: Co sca – Ekman 3891: Co pul; H 5822: Co spc; 12975: Co pul; H 15292: Co spc; 16249: Co pul – Elcoro 90: Co spl – Elias 1614: Di str; 1704: Co kun; 1711: Co woo – Ellenberg 2420: Co sca; 3506: Co ama – Elmore 22: Co kun –Englesing 255: Co pul – Erlanson 483: Co mac; 5470: Co vil – Ernst 1448: Co spc; 1689: Co spc – Espina Z 17: Co ara; 45: Co ara; 406: Co sca; 502: Co pul; 603: Co pul; 640: Di str; 1218: Co leu; 1498: Co sca; 1921: Co mac; 1998: Co lim; 2184: Co las; 2186: Di str; 2473: Co kun; 2589: Co kun; 2593: Co kun; 2726: Co woo; 3728: Co kun; 3810: Co las – Espinal T 2820: Co kun – Espinosa, A 614MB: Co mac – Espinosa, R 673: Co sca – Espinoza, R 1184: Co pul – Espinoza, S 291: Co sca; 566: Co sca; 637: Co sca – Espinoza Obando 1503: Co pul – Estrada Chavarría 73: Co mac; 5385: Co woo – Estudiantes Botánica Taxonómica 1997: 73: Di str – Estudiantes Botánica Taxonómica 1998: 111: Co pul – Estudiantes Botánica Taxonómica 2000: 182: Co cha – Estudiantes ESPOCH 144: Di str; 418: Di str – Estudiantes Herbario MEDEL: 273: Co woo – Evans 2737: Co sca; 3143: Co cla – Evrard 8550: Co sca; 9732: Co acr – Ewan 16747: Di str; 16748: Co cup.

Fairchild 16: Di str; 2925: Co sca; 2958: Co sca – Fajardo 83: Co spl – Falcão 5167: Ch lan pul – Falconí 12: Co kun – Fanshawe 541: Co cla; 1089: Co ert; 1171: Co ara; 2302: Co cla – Farfán 727: Co sca; 1175: Di arg – FDBG (Forest Department British Guyana) 2274: Ch con; 3277: Co cla; 3907: Co ara; 3825: Co ert; 5038: Co cla – Fegan 3: Co pul; 24: Co kun – Feinsinger 409: Co wil – Fendler 444: Di str; 446: Co woo; 447: Co kun; 803: Co sca; 804: Co mac; 810: Co sca; 811: Co sca; 863: Co sca; 1497: Co spl; 2158: Co com – Feres 24/95: Co spi – Ferguson 19: Co pul – Fernández, Álvaro 137: Co bra; 1043: Co cur; 1179: Co pul – Fernández, Ángel 483: Co ara; 2704: Co spl; 3145: Co sca; 3992: Co sca; 5376: Co spl; 17466: Co spl – Fernández, Antonio 1756a: Co sca; 2154: Co sca; 2395: Co ara; 2412: Co spl; 2533: Co sca; 2918: Co spl; 3554: Co sca – Fernández, D 586: Co sca – Fernandez, Y 420: Co sca – Fernández-Alonso 5901: Di str; 7308: Co kun; 9226: Co sca; 11320: Co sca; 11353: Co sca; 11354: Co kun; 11386: Co sca; 11396: Co asp; 11440: Co pit; 12416: Co lim; 12917: Co sca; 15993: Co com; 16203: Co spl; 16297: Co vil; 16534: Co vil; 16586: Co sca; 16589: Co sca; 22960: Co com; 23102: Di str; 23433: Co com; 23998: Di str; 24003: Co vil – Fernández Casas 5800: Co ara; 6168: Co ara; 7957: Co vag; 8141: Co acr; 8455: Co sca; 10780: Co pul – Fernández Pérez 201: Co woo; 248: Co kun; 371: Co vil; 401: Co las; 1908: Co sca; 2259: Co sca; 5044: Di str; 5049: Co sca – Ferrari 240: Co spl; 576: Co spl; 1629: Co sca; 1631: Co sca; 1916: Co sca – Ferreira, AM 858: Co spl –Ferreira, L 124: Co sca; 139: Ch aca – Ferreyra, R 1151: Di rur; 1275: Co jur; 1617: Co sca; 4221: Co lae; 4259: Di rur; 4502: Co lae; 4589: Co sca; 5161: Co jur; 6843: Co web; 7922: Co sca; 13042: Co lae; 15763: Co pul; 15794: Co mon; 15854: Co kun; 15892: Co pul; 16008: Co kun; 16043: Co sin; 16103: Co jur; 16340: Co sca; 16917: Co sca; 17254: Co sca; 18137: Co lae; 18142: Co sca; 18153: Co sin; 18945: Co las; 19285: Co las – Ferris 5556: Co pic – Feuillet 528: Co ert; 667: Co cla; 801: Co ara; 827: Co spl; 1218: Co sca; 1314: Co spl; 2239: Co cla; 2252: Ch con; 2406: Co sca; 3781: Co ert; 4013: Co ara; 10302: Co sca – Fiaschi 3287: Co pra – Fiebrig 4929: Co ara – Figueiredo, C 299: Ch lan lan; 566: Co ara; 841: Co vaz; 1109: Co acr – Figueiredo, FOG 62: Co sca; 76: Co ara; 77: Co ara; 81: Co sca; 93: Co spl; 95: Ch con; 98: Co las; 125: Co sca – Filgueiras 1652: Co spl; 1950: Ch sub; 2376: Co spl – Finlay TRIN 2884: Co sca; TRIN 2885: Co sca; TRIN 2886: Co ara – Fleischmann 64: Co ara; 67: Di arg; 372: Co ara; 373: Co ara; 378: Co ara – Fleming 22: Co sca – Fletes Almengor 78: Co wil; 211: Co mac; 212: Co pul; 417: Co ric; 619: Co las; 714: Co ric – Fleury 152: Ch lan pul; 153: Ch con; 409: Ch con; 648: Ch con; 723: Co sca; 776: Co ara; 808: Ch lan pul; 809: Ch con; 815: Co vil; 1149: Co ara; 1261: Co ara; 1262: Co ara; 1271: Co sca; 1424: Co sca; 1839: Co sca; 2001: Co spl; 2199: Co sca – Florschütz, JP 113: Co mac; 251: Co sca; 1006: Co sca; 1054: Co ara; 1243: Co sca; 1547: Co spl; 1740: Ch con – Florschütz, PA 2309: Co sca; 2355: Co ara; 2473: Co ara; 2474: Co sca; 2533: Di rur; 2690: Co spl; 2788: Co spl; 2789: Co sca; 2790: Co spl; 2822: Co cla; 2847: Co cla; 2865: Co cla – Flynn 4078: Co sca; 4153: Di str; 4181: Co asp – Focke 123: Co sca – Foldats 325A: Co sca; 2429: Co ara; 2444: Co pul; 2740: Co sca; 3598: Co ara; 3608: Co spl; 3861: Co spl; 9179: Co spl; 9499: Co spl – Folli 1267: Co sca; 1645: Co ara; 1985: Co sca; 2696: Co ara; 4559: Co ara; 5743: Ch sub; 6751: Ch cus – Folsom 1430: Co all; 1726: Di str; 1789: Co cur; 1934: Di str; 2406: Co all; 2467: Co pul; 2553: Co kun; 2591: Co bra; 2797: Co las; 2950: Co cur; 3053: Co com; 3428: Co pul; 3506: Co sca; 3591: Co sca; 3616: Co all; 3628: Co las; 3687: Co kun; 3714: Co woo; 3744: Co pul; 3755: Co vil; 3874: Co pul; 3892: Co all; 4061: Co wil; 4354: Co cur; 4494: Co cur; 4837: Co wil; 4980: Co pul; 5470: Co com; 6374: Co pul; 6722: Co las; 6778: Co pul; 7833: Co sca; 8711: Co mal; 9853: Co bra; 9865: Co pul; 9888: Co bra; 9965: Co sca – Fonnegra G 958: Co osa: 1025: Co mac; 1731: Di str; 1741: Co las; 1760: Co pul; 1796: Co mac; 1797: Di str; 1994: Co las; 2176: Co osa; 2636: Co las; 2787: Co sca; 3070: Co las; 4464: Co las; 4502: Co las; 4759: Co sca; 4988: Co vil; 5032: Co pul; 5100: Co pul; 5136: Di str; 5455: Co plo; 5733: Co las; 5774: Di str; 6119: Co mac; 6244: Co las; 6251: Co mac; 6376: Co las; 6402: Co mac; 6451: Co mac; 6811: Di str; 6829: Co pul; 6916: Di str; 7085: Co ant; 7278: Co pul; 7299: Co lim; 7467: Di str; 7504: Co pul; 7697: Di str; 8079: Di str; 8157: Co ant; 8658: Co pul; 8839: Co sca; 8841: Co pul; 8842: Co sca; 8884: Co mac – Fonseca, G 18: Co sca – Fonseca, ML 1245: Ch sub – Fonsêca, SG 156: Co ara – Forero, E 693: Co lim; 842p.p.: Co las & Co sca; 969: Di str; 1017: Di str; 1018: Di str; 1062: Di cry; 1336: Co las; 1398: Co kun; 1400: Co las; 1438: Co sca; 1522: Co lim; 1586: Co sca; 1642: Co pul; 1845: Co sca; 1848: Co kun; 1892: Co mac; 1925: Di str; 1993: Di str; 2130: Co mac; 2410: Co pul; 2427: Co mac; 2595: Di str; 2641: Co lim; 3339: Co pul; 3340: Co kun; 3822: Co kun; 3852: Co sca; 3888: Di str; 4036: Co sca; 4085: Co kun; 4105: Co sca; 4609: Co sca; 4949: Co lim; 4950: Co lim; 5179: Co mac; 5183: Co lim; 5389: Co mac; 5458: Co sca; 5472: Co kun; 5536: Co las; 5559: Co kun; 5765: Co cal; 5772: Co las; 5814: Di str; 6161: Co ant; 6436: Co sca; 6455: Co sca; 6473: Co mac; 6503: Co kun; 6584: Co pul; 6668: Co cal; 6977: Di rur; 7086: Ch aca; 7087: Co sca; 7314: Co cor; 7364: Co cor; 7413: Co ant; 8996: Co lim; 9031: Co sca; 9232: Co lim; 9445: Co kun; 9448: Co kun; 9487: Co kun – Forero, FA 100: Co vil – Forero P 397: Co sca; 432: Co pul; 462: Co mac; 506: Co pul; 526: Co pul; 557: Di str; 572: Co sca; 629: Co las; 646: Co las; 709: Co kun; 735: Co mac; 815: Co kun –Forzza 8936: Co ara – Fosberg 29128: Co las; 29253: Co ara – Foster, MB 1311: Co com – Foster, RB 585: Co vil; 894: Co sca; 926: Di str; 948: Co gla; 1124: Co mac; 2257: Co lim; 2349: Co kun; 2446: Co sca; 2680: Co sca; 2744: Co vag; 2745: Co web; 3025: Di rur; 3195: Co sca; 3517: Di rur; 4010: Co bec; 4238: Co sin; 4249: Co las; 4379: Co lon; 5206: Di rur; 5592: Co ara; 6080: Co sca; 6579: Co acr; 6817: Co sca; 7164: Co acr; 7174: Di rur; 7393: Co jur; 7830: Co lae; 7831: Co lae; 7836: Co jur; 8003: Co jur; 8030: Co las; 8559: Co zin; 8589: Co jur; 8700: Co com; 8858: Co jur; 8950A: Co rub; 9316: Co las; 9423: Co jur; 9615: Co sin; 9644: Co ara; 10071: Co kru; 10209: Co sca; 10935: Co vag; 10937: Co obs; 11512: Co gua; 12566: Co acr; 12760: Co sin; 13658: Ch aca; 13680: Co spl – Fournet 167: Co spl; 210: Co ert; 516: Di arg – Frame 171: Co pra – Franco Mellado 2298: Di are – Franco Rosselli 1106: Co sca; 1821: Co sca; 1926: Co kun; 4879: Co plo; 5655: Di cry – Frankie 49a: Co mal; 49c: Co mal; 62a: Co pul –Freire Mayorga 3248: Co erp; 3615: Co sca; 3714: Co lon; 3744: Co lon – Freire, B 447: Co ama; 510: Co ama; 538: Co ama; 656: Co sca; 851: Di str; 965: Co sca – Freire, CV R 50963: Co spl – Freitas 66: Ch lan lan – Frey 733: Co sca – Friedrich 12: Co ant – Fróes 21624: Co las; 21630: Co acr; 25738: Co ara; 25740: Co spl; 26938: Co spl; 27096 p.p.: Co sca & Co spl; 27173: Co ara; 28612: Co spl; 29382: Co ara; 30702: Co spl; 32870: Co sca; 32965: Co ara; 33941: Co ara; 34249: Co ert; 34323: Co las; 34328: Co ert – Fuchs 21722: Di str; 21897: Co las; 21907: Co kun; 21910: Di str – Fuentes Alvarado 302: Co mon – Fuentes Claros; 3731: Di arg; 3741: Co sca; 3914: Co sca; 3921: Co sin; 5359: Co vag; 5413: Co sca; 5417: Di arg; 6127: Co gua; 6216: Co alf; 11187: Co gua; 15724: Co gua; 15797: Co gua – Funk 6095: Co sca; 6142: Co sca; 8101: Di rur; 10490: Co cal; 10524: Co pul; 10624: Co mal; 10665: Co pul; 10966: Co mal.

Gadelha Neto 3803: Co spl – Gaillard 209: Co ara – Gaitán 4: Co sca; 107: Co sca; 169: Co las; 183: Co erp – Galeano, G 78: Co kun; 282: Co kun; 907: Co spl; 1458: Co sca; 1459: Co cha; 1656: Co lon; 1696: Co sca; 1936: Co ara; 2006: Co sca; 3697: Co kun; 4592: Co las; 4833: Co kun; 5327: Co las; 5684: Cocom; 5827: Co kun; 6330: Co erp; 6424: Co com; 6463: Co com; 6466: Di str – Galeano, MP 2036: Co ant – Galen Smith 1535: Co mac – Galeotti 4991: Co pic – Galindo T 102: Co spl; 1117: Co sca; 1440: Co ant – Gallardo Ramírez 186: Co pul; 187: Co lim – Gallegos 5172: Co las – Gamboa, C 26: Co vil – Gamboa, M 222: Co ant – Gamboa-Moreno 23: Co woo – Gamboa Romero 120: Co sca; 153: Co com; 222: Co wil; 535: Co wil; 602: Co wil; 1497: Co sca – Garber 101: Di str; 83: Co mac; 90: Co mac – García 337: Co lim – García, A 198: Co spl; 228: Co com – García, L 22: Di str; 36: Co las – García, M 2074: Co pul; 2174: Co pic – García, MC 392: Co erp; 469: Co erp; 506: Co spl – García, R 634: Co sca; 1663: Co spc – García-Barriga 4636: Co kun; 5102A: Co vil; 5184: Co spl; 8203: Co ant; 8301: Co las; 10953: Co ant; 11186: Co las; 11187: Co las; 13853: Co sca; 14379: Co sca; 14467: Co fis; 14656: Co sca; 17131: Co spl; 18240: Co ant; 18319: Co mac; 18532: Co ara; 21120: Co com – García Cossio 468: Co vil – García G, JD 265: Co ara; 270: Co mac – García García 113: Co pul – Gardner 3464: Ch sub – Garnier 84: Co sca – Garreta 32: Co sca – Garvizu 313: Ch aca – Garwood 55: Co lim; 913: Co lim; 919: Co pul; 1150: Co kun; 1315A: Co pul; 1624A: Di str; 1647A: Co pul; 1675A: Co sca; 1900A: Co vil; 2214A: Co mac; 2263A: Di str; 2656A: Co kun; 2670A: Co pul; 2682A: Co vil; 2690A: Co all; 2696a: Co kun; 2697a: Co kun; 2698A: Co kun; 2938: Co kun – Garzón, G 3208: Co ara – Garzón, N 2147: Co spl; 2201: Co ara – Gasche 2: Co las; 1136: Co pul – Gastony 670: Co sca – Gaudichaud-Beaupré 325: Co spl – Gaumer 23315: Co pic – Geay 24: Co pul – Gehrt SP 46301: Co spl – Geijskes 1024: Co cla – Geiselman 89: Ch cur – Gentle 983: Co pul; 2116: Co mac; 2394: Co pul; 4810: Co mac – Gentry 1378: Co pul; 2565: Co pul; 2729: Di str; 2960: Co kun; 3805: Co pul; 4568: Di str; 4603: Co las; 4985: Co las; 5091: Co sca; 5511: Di str; 5522: Co all; 5663: Co cur; 5675: Co all; 6281: Co com; 6302: Co woo; 6454: Co pul; 6970: Co cur; 7001: Co las; 7197: Co sca; 7322: Co cor; 7731: Co pul; 7850: Co pul; 7910: Co pul; 7919: Co pul; 8002: Co pul; 8115: Co pul; 8756: Co vil; 9095: Co cha; 10122: Co pul; 10165: Co gib; 10165A: C lim; 10206: Co lim; 12105: Co pul; 13571: Co sca; 13610: Co las; 13641: Co las; 14357: Co mac; 15268: Co las; 15307: Co pul; 15666: Co sca; 15776: Co las; 16057: Co sca; 16224: Co jur; 16277: Co acr; 16322: Co sca; 16749: Co pul; 16822: Co las; 17491: Co sca; 17500: Co kun; 18239: Co kru; 18241: Co acr; 18299: Co sca; 18953: Co sca; 19017: Co sin; 20558: Co kru; 20994: Co las; 21843: Co ara; 21868: Co sca; 22096: Di str; 23122: Co asp; 23614: Di rur; 23884: Co lim; 24005: Co kun; 24039: Co las; 24096: Co Gentry; 24137: Co las; 24414: Co all; 24454: Co las; 26579: Co pul; 27110: Di rur; 27182: Co sca; 27183: Co sca; 27270: Co vag; 27286: Co las; 27304: Co obs; 27395: Co erc; 27719: Co lon; 28076: Co sca; 28664: Co cur; 28953: Co lon; 29061: Co mac; 29096: Co ara; 29211: Co sca; 30061: Co las; 30061A: Co pul; 30065: Co las; 30895: Co ama; 30960: Co ama; 31070: Co spl; 31071: Co sin; 31084: Co jur; 31086: Co jur; 31106: Di rur; 31114: Co jur; 31245: Co mac; 31247: Co jur; 31332: Co erp; 31353: Di app; 31366: Co sca; 31517: Co sca; 31792: Co acr; 32243: Co sca; 32608: Co pul; 32621: Co dir; 35700: Co leu; 36422: Co erc;36623: Di app; 36894: Co kun; 36939: Co las; 37060: Co las; 37190: Co sca; 37759: Co spl; 37984: Co sca; 37995: Di app; 38006: Co sin; 38101: Co sin; 38718: Co sin; 38720: Co sca; 39552: Co las; 39664: Di app; 40052: Di arg; 40580: Co kun; 40686: Co all; 40955: Co all; 40962: Co sca; 41421: Di rur; 42295: Co sca; 42351: Co las; 42717: Co sca; 43148: Co sca; 43591: Di rur; 43828: Co sca; 44317: Co sca; 44322: Co web; 44326: Co gua; 45809: Co sca; 45814: Co acr; 47547: Co vil; 48075: Co lae; 48184: Co vil; 48691: Co sca; 49343: Co ara; 49345: Co ara; 49791: Co spl; 52203: Co pra; 55292: Co kun; 55958: Di app; 60845: Di app; 5663: Co cur; 5675: Co all; 65759: Di app; 68928: Co sca; 69359: Co sin; 69381: Co acr; 69502: Co ara; 72077: Co sin; 72079: Di app; 72168: Co sca; 72233: Co sca; 72332: Co gib; 72952: Co pul; 72962: Co lim; 73287: Co sin; 73744: Co sca; 74248: Co sca; 75918: Co pul; 76067: Di cry; 77496: Co sca; 78160: Co acr – Gerardino 2959: Co spl – Gerlach 32-1: Co gua – Germano IAC 40332: Co spl; IAC 40333: Co spl – Gíl 5: Di str – Gillespie 1153: Co ara; 1877: Co spl; 2097: Co sca; 2191: Co ara – Gilli 119: Co gib –Gillis 9216: Co woo; 9493: Di str – Gilly 371: Co pul; 97: Co mol – Gilmartin 171: Co pul; 337: Co lim; 618: Co sca – Ginés 2052: Co pul; 4383: Co pul; 4436: Co sca; 4595: Co pul; 4918: Co sca; 5007: Co sca; 5162: Co ara – Ginzberger 826: Co ara – Giraldo, C 405: Co pul – Giraldo, R 8: Co osa; 143: Co pul – Giraldo-Cañas 141: Di str; 288: Co las; 291: Co kun; 308: Co las; 544: Co las; 591: Co las; 706: Di str; 2010: Co sca; 2014: Co kun; 3456: Co spl – Giraldo-Rodíguez 8657: Co kun; 8761: Co sca – Girardi 76: Co mac – Girault 555: Co ara; 556: Co ert – Glaziou; 4232: Co spl; 4242: Co spl; 9318: Ch cus; 9319: Co ara; 15666: Ch sub; 22180: Ch sub; 22181: Ch sub – Gleason 306p.p.: Co cla & Co spl; 493: Co ara; 522: Co sca – Glenboski C5: Co ama – Godoi 407: Co spl – Goldenberg 1348: Co spl; 1357: Co sin; 1401: Ch lan lan – Goldman 1963: Co las – Goldstein 49: Co pul – Gómez Chagala 352: Co pic; 656: Co pul – Gómez, A 330: Co woo; 411: Co vil; 441: Co lim; 471: Di str – Gómez, RF 6119: Co mac – Gómez Dominguez 2410: Co mac – Gómez-Laurito 10555: Co mal; 11488: Co mon; 12005: Co mon; 12006: Co mon – Goméz Mejia R 3229: Co all – Gómez Pignataro 18818: Co cur; 19988: Co com; 20492: Co woo; 20511: Co kun; 21201: Co cal; 22911: Co pli; 23025: Co bar – Gonto 1670: Co spl – Gonzales, A 285: Co pul – Gonzales, G 81: Co acr – González Arce 4185: Co sca – Gónzalez G 919: Di str; 2748: Co plo – González L 3788: Co pul; 4289: Co pul – Gonzalez Ortega 6859: Co pic – González, ÁC 1230: Co spl; 1368: Co mac – González, E 13: Di str – Gonzalez, G 50: Co las – Gonzalez, M 158: Co pic – González, MF 68: Co spl; 266: Co com; 267: Di str; 679: Co sca – González, R 13: Co ric – Gonzalez, S 2053: Co ara; 2133: Ch cur – González-Villareal 4118: Co pic; 4166: Co pic – Goodland 154: Ch sub – Gopaul 168: Co ert – Gordon 99C: Co pul; s.n: Co mac & Co sca – Gorostiza Salazar 1890: Co pic – Gottsberger 14-12375: Co kru; 12-24183: Ch lan pul –Goulding 1188: Co ara – Gragson 24: Co ara – Graham, EH 161: Co sca – Graham, JG 185: Co sca; 421: Co ara; 432: Di rur; 556: Co sca; 722: Di rur; 725: Co sin; 733: Co sca; 1135: Co sin; 2647: Co jur; 4272: Co jur; 4343: Co jur; 4632: Co las – Graham, SA 199: Co pul; 327: Co pul –Grández Rios 214: Co sca; 958: Co sca; 1524: Di rur; 2695: Co ara; 3217: Co erc; 3453: Co sca – Grant, JR 92-1743: Co pul; 92-1789: Co ric; 92-1905: Co com; 92-1965: Co woo; 92-2010: Co pic; 92-2265: Co mac; 96-2360: Co sca; 97-2832: Co pul – Grant, ML 10407: Co com – Grayum 1039: Co mal; 3657: Co woo; 3805: Co sin; 4008: Co ste; 5489: Co pul; 5668: Co wil; 5875: Co kun; 5909: Co com; 5918: Co pul; 5931: Co com; 8282: Co mon; 9839: Co bra; 10032: Co woo; 12580: Co wil; 12835: Co wil; 13019: Co sca – Greenman 5322p.p.: Co bar & Co woo – Grenand 4: Co sca; 64: Ch con; 86: Co ara; 594: Co vil; 732: Co ert; 740: Co sca; 1075: Co spl; 1244: Co cla; 1246: Ch cur; 1747: Co ert; 1797: Co ert; 2000: Ch con; 3196: Co spl; 3221: Co ara – Grewal 280: Co ara – Grijalva P 8: Di str; 62: Co sca; 1044: Co pul; 1071: Co pul; 1131: Co sca; 1142: Co mac; 1147: Co pul; 1160: Co pic; 1176: Co vil; 1515: Co gla; 1519: Di str; 1554: Co pul; 1656: Co pul; 1686: Co pul; 1697: Di str; 1930: Co pic; 1976: Co pic; 2916: Co pic –Groenendijk 215: Co ara – Gröger 324: Co spl – Groppo 465: Co spl – Grove 25: Co mal – Guadamuz 3620: Co pul – Guánchez Meza 1862: Co ara; 2679: Co spl; 3655: Co spl – Guareco 259: Co ara – Gudiño Jara 147: Co sca; 223: Co sca; 568: Co sca – Guedes 155: Co ara – Guerra L 19: Co sca – Guerrero, C 12: Co asp; 17: Co ama; 28: Co lon – Guerrero, W 137: Co kru – Guevara, JE 253: Co sca – Guevara-Ibarra 39: Co ant – Guillén 274: Co ara – Guillén Villarroel 1466: Co ara; 2946: Co ara; 3035: Ch aca; 3349: Co ara; 3623: Ch aca –Gutiérrez, C 468: Co sca – Gutiérrez, E 603: Co spl – Gutiérrez, L 95: Co wil – Gutiérrez, M 162: Di str; 183: Co las – Gutiérrez Villegas 17C-127: Co pul; 17C-159: Co mac; 965: Co spl; 1183: Co las; 1239: Co las; 2518: Di str; 2519: Di str; 2529: Co lon; 2734: Co erp; 35511: Co las; 35568: Co las – Guzman 799: Co lim – Guzmán-Teare 489: Co pic; 879: Co pul; 2165: Co pul; 2190: Co sca; 2192: Co pul – Gwynne Vaughan 58: Co sca.

Haber 1383: Co bar; 1697: Co sca; 1757: Co pul; 1919: Co cur; 2027: Co wil; 2725: Co wil; 3161: Co wil; 3269: Co pul; 4480: Co mon; 5435: Co mal; 5455: Co cur; 5906: Co pul; 6969: Co mon; 7110: Co cur; 8223: Co mal; 9458: Co nit; 10581: Co mon; 10798: Co wil –Hahn, L 330: Co sca; 505: Co spc; 1032: Co spc; 1836: Co spc – Hahn, R 5: Co erc; 11: Co erc; 108: Co lon – Hahn, WJ 169: Co pul; 251: Co pul; 4768: Co jur – Hallé 788: Ch con – Hamblett 611: Co woo – Hamilton 514: Co pul; 545: Co all; 550: Co pul; 805: Co wil; 822: Co wil; 826: Co wil; 1039: Co bra; 1256: Co kun; 1264: Co pul; 1344: Co pul; 1354: Co pul; 1369: Co pul; 1423: Co pul; 2778: Co alo; 3763: Co wil; 3950: Co com; 3955: Co cur; 3958: Co com; 4143: Co cur – Hammel 1659: Co sca; 1697: Di str; 1866: Co sca; 3705: Co all; 3784: Co cur; 4300: Co las; 7893: Co mal; 8708: Co sca; 8775: Co mal; 8845: Co pul; 8965: Co pul; 8966: Co sca; 8967: Co bra; 8981: Co kun; 9003: Co kun; 9057: Co sca; 9058: Co sca; 9158: Co bra; 9161: Co pul; 9243: Co sca; 9574: Co sca; 9655: Co nit; 9924: Co mal 10303: Co sca; 11438: Co mal; 11625: Co mal; 11646: Co bra; 12259: Co sca; 12444: Co pul; 12789: Co pul; 12948: Co pul; 14354: Co nit; 15871: Co gib; 16116: Co pul; 16361: Co las; 16513: Co cal; 16732: Co las; 16867: Co kun; 17117: Co vil; 17532: Co bra; 17668: Co cal; 17668A: Co bra; 17856: Co osa; 17938: Co las; 18215: Co gla; 18230: Co wil; 18247: Co pli; 18285: Co mac; 18318: Co osa; 18322: Co sin; 18406: Co ric; 18720: Co ric; 18994: Co sca; 19639: Co pul; 19665: Co woo; 20156: Co ric; 20265: Co sca; 20271: Co pul; 20381: Co pul; 20397: Co sca; 20461: Co mal; 20482: Co osa; 20492: Co pli; 20494: Co cal; 20501: Co pul; 20507: Co mac; 20513: Co lim; 20515: Co bra; 20519: Co sca; 20620: Co com; 20626: Co sca; 20694: Co cal; 20803: Co gla; 21078: Co wil; 21107: Co wil; 21145: Di str; 21212: Co ara; 21213: Co sca; 21214: Co ara; 21371: Co ert; 21642: Co cla; 21770: Ch con; 21771: Co ert; 21774: Co sca; 21778: Co sca; 23706: Co wil; 24847: Co pic; 24917: Co cur; 25016: Co sca; 26342: Co alo; 27209: Co woo – Hammit 181: Co woo – Hansen 7388: Co pul; 7568: Co pic; 7614: Co pul; 7959: Di str – Harley 10223: Co ara; 10765: Co ara; 16573: Co spl; 17876: Co spl; 18341: Co ara; 22226: Co ara – Harling 215: Co pul; 3575: Co erp; 3646: Di str; 3816: Co asp; 4296: Co kun; 4324: Co vil; 4477: Co pul; 4761: Co mac; 4784: Co vil; 4827: Co vil; 6966: Co asp; 6970: Co sca; 7016: Cozam; 7263: Co lon; 7264: Co ara; 7614: Co sca; 7697: Co asp; 9315: Co pul; 10075: Co asp; 11709: Di str; 11710: Co lon; 11717: Co ama; 11752: Co sca; 11837: Co asp; 12751: Co zam; 12797: Co ama; 12806: Co sca; 13748: Co ama; 13930: Co sca; 13960: Di rur; 13999: Co acr; 14032: Co sca; 14041: Co acr; 14352: Co pul; 14353: Co lim; 14354: Co gib; 16245: Co ama; 16514: Co erc; 16529: Co kun; 16761: Co mac; 17472: Co cha; 17699: Co asp; 18358: Co gib; 18748: Co pul; 18929: Co pul; 19264: Co vil; 19411: Co pul; 21251: Co zam; 23096: Co pul – Harlow 7: Co vil – Harmon 1928: Co sca; 2478: Co pul; 4839: Co pul; 5060: Co pul; 5796: Co pul – Harris, EM 1074: Co spl – Harris, SA TP 446: Co ara – Harrison 825: Co sca; 1711: Co ara – Hartman 12001: Co pul; 12115: Di str – Hashimoto 666: Co ara – Hassler 5974: Co ara – Hatschbach 6664: Co sp; 13528: Co spl; 15903: Co ara; 16015: Co spl; 20643: Co ara; 26068: Co spl; 26104: Co ara; 26257: Co ara; 36012: Co ara; 39011: Ch sub; 39732: Co spl; 55820: Ch sub – Haught 1779: Co pul; 1833: Co sca; 1913: Co ant; 4130: Co com; 4274: Co pul; 4292: Co ant; 4292A: Co sca; 5524: Co las – Hawkes 2079: Co pul –Haxaire 454bis: Co cla – Hayden 84: Co sca; 1020: Co cal – Hayes 113: Di str; 843: Co woo – Hazlett 1667: Co pic – Hecht 86: Ch lan pul – Heinrichs 332: Co lon; 468: Co cha – Heinsdijk 71: Co ara – Heller 10001: Co mac – Helme 743: Co sca; 804: Co ara – Henao 262: Co las – Henao-Cárdenas 380: Co ara – Henicka 246: Ch fus; 265: Ch fus; 273: Ch lan lan – Henkel 291: Co sca; 458: Co sca; 465: Co ara; 501a: Co sca; 1319: Co jur; 1906: Ch con; 2191: Co jur; 3058: Co spl; 4684: Co sca; 4767: Co ara; 4824: Co sca; 5014: Co lon; 5343: Co ara – Henrich 274: Co pul – Henry 470: Co kru – Hentrich FGIC 18: Ch con – Hequet 376: Co sca; 424: Co ara; 591: Co sca – Herbarium QCA 1316: Co pul – Heredia, L 20: Co pul – Heredia, M 14: Co ara – Heringer 949: Co spl; 1609: Co spl; 2855: Co spl; 4034: Co spl; 5080: Co spl; 5469: Co spl; 7343: Ch sub; 9407: Ch sub; 14058: Co spl; 17343: Co ara – Hernández, F 1417: Co com – Hernández, JJ 20: Co com; 58: Co mac; 86: Co vil; 161: Co las; 198: Co ant; 358: Co las; 365: Co las; 408: Co mac; 495: Co las; 525: Co las; 540: Co las; 562: Co las; 640: Co las – Hernández A 2306: Co lon; 2431: Co lon; 3090: Co asp – Hernández G 387: Co mol; 1435: Co mol; 1436: Co pul – Hernández Macías 490: Co pul; 684: Co pul – Hernández Schmidt 721: Co sca – Herrera, A 129: Co erc; 147: Di str; 188: Co asp; 250: Co kru – Herrera, H 539: Co vil; 663: Di str – Herrera C 2031: Co mon; 2792: Co cur; 2968: Co pul; 3036: Co bra; 3127: Co gla; 3979: Co osa; 4578: Co osa; 4701: Co ric; 7071: Co com; 7089: Co pul; 7638: Co bra; 9692: Co sca; 9765: Co spl; 9770: Co ara; 10072: Ch fus; 10131: Co sca; 10134: Co ara – Herrera P 15: Co pul – Hertel 35719: Co woo – Heyde 4650: Co com – Heyligers 319: Co spl – Hijman 205: Co spl – Hill 12863: Co ert; 12879: Co ara; 13138: Ch man; 20309: Co sca – Him 423: Co cur – Hinojosa Ossio 522: Co vag; 1161: Co sca – Hinton 11792: Co pic; 16011: Co pic – Hioram, Bro 2574: Co pul – Hitchcock, AS 16874: Co ara; 17023: Co ara; 17179: Co sca; 17402p.p.: Co ara & Co spl; 20512: Co mac; 20543: Co gib – Hitchcock, CL 6926: Co pul; 20466: Co pul – Hodge 1018: Co pul; 2586: Co spc; 3757: Co spc; 6972: Co osa; 7064: Co mac; 7066: Co all; 7078: Co pul – Hodges 14: Co sca; 55: Co ama; 201: Co ert – Hoehne, FC 105K: Ch lan lan; 728: Co ara; 1084: Co ara; 1634: Co ara; 2264: Co ara; 3666: Co ara; 5500: Ch lan lan; 5501: Ch lan lan; 6206: Ch sub; SP 39266: Co ara – Hoehne, W SP 74118: Ch aca – Hoff 5241: Co spl; 6234: Co ara; 6299: Co sca; 6305: Ch con; 6354: Co ara; 6436: Ch con; 6583: Co cla; 6584: Co ert; 6846: Co ert; 6931: Co ert; 7324: Co sca; 7767: Co sca – Hoffman, B 1208: Co ara; 1296: Co ara; 2253: Co ert; 2727: Ch con; 2841: Co mac; 3736: Co mac; 5560: Co sca; 5579: Ch con; 6671: Co sca – Hoffmann, C 727: Co sca –Holland 53: Co pul – Holliday 55: Co sca – Hollowell 420: Co ara; 479: Co sca – Holm 727: Co pul – Holm-Nielsen 368: Co asp; 515: Co sca; 1015: Co sca; 2678: Co pul; 4257: Co ama; 4421: Co acr; 4436: Co asp; 7007: Co pul; 19138: Co sca; 19145: Di str; 19411: Co pul; 19645: Co sca; 19657: Co sca; 19685: Co sca; 19716: Co ama; 19815: Co sca; 19842: Co erp; 19889: Co sca; 19937: Co sca; 20087: Co cha; 20124: Co erp; 20180: Co sin; 20204: Co sca; 20230: Co sca; 20453B: Co lon; 20490: Co sca; 20549: Co sca; 21044: Di str; 21073: Co lon; 21079: Co sca; 21087A: Co sca; 21177: Co sca; 21183: Co sca; 21186: Co ara; 21306: Co ara; 21357: Co sca; 21617: Co ara; 21851: Co sca; 21941: Co cha; 22005: Co cha; 22052: Co sca; 22117: Co erc; 24582: Co pul; 24589B: Co pul; 25373: Co pul; 25384: Co pul; 25582: Co kun; 25617: Co pul; 25767: Co sca; 25786: Co geo; 25789: Co sca; 25992: Co lim; 27916: Co gib; 37749: Co pul; 37825: Co pul; 188929: Co pul – Holst 2460: Co spl; 2571: Co ara; 4332: Co pul; 5769: Co pul; 8734: Co las – Holt 280: Co ara; 578: Co pul – Holton 206: Co sca – Hoover 1258: Co sca; 2423: Co pul; 3463: Co sca; 3607: Co pul; 3745: Co cor; 4127: Di str – Hopkins 138: Co spl; 534: Co sca; 705: Co ara – Horner 10: Co sca; 61: Co ara; 94: Co ara; 309: Co ara; 325: Co sca – Hort. Bot. Haun., September 1891: Co pic & Co vil – Hort. Kew, January 1844: Ch cus – Hort. Bot. Paris, October 1903: Co spc – Hostmann 139: Co ara – House 1243: Co pul – Howard 5747: Co pul; 9557: Co spc; 10555: Co sca; 11157: Co sca; 16653: Co luc; 18262: Co sca – Hoyos Gómez 285: Co pul; 296: Co sca; 375: Co pul; 402: Co vil; 453: Co pul; 553: Co sca; 772: Co woo; 785: Di str; 791: Co pul; 1085: Co sca; 2492: Co plo – Hoyos Marin 637: Co ant; 855: Co ant; 1013: Co las – Huamán 155: Co las; 333: Co las; 392: Co gua – Huamantupa Chuquimaco 2485: Co acr; 4835: Di app; 7557: Di rur; 7566: Co sca; 10624: Di rur – Huashikat 19: Co asp; 43: Co sca; 168: Co sca; 294: Co erp; 563: Co ert; 676: Co ert; 775: Co ert; 851: Co ert 1146: Co ert; 1293: Co ama; 1333: Co ert; 1530: Co ert; 1863: Co ert; 1882: Co ama; 2099: Co erp; 2209: Co ert; 2229: Co asp – Huber, J 1384: Di str; 1461: Di str; 1692: Co sca; 1807: Co las; 4501: Co sca; 4669: Co ara; 6891: Co vaz; 7010: Co leu; 7256: Co ara; 9332: Co vaz – Huber, O 416: Co sca; 1059: Co mac; 1816A: Co spl; 1817: Co ara; 5039: Co spl; 6003: Co spl; 6683: Co spl – Huertas 6959: Co sca; 7001: Co kun – Hugh-Jones 375: Co kun; 382: Co lae – Humbert 27231: Co spl – Hummel 52: Co mal – Hunnewell 16415: Co pul – Hunter, AA 17041: Co sca – Hunter, JR 17: Co mal; 7997: Co mal – Hurtado, DL 279: Co sca; 715: Di str – Hurtado, F 48: Co sca; 508: Co sca; 570: Co sca; 1504: Co sca; 2899: Di str – Hurtado, JG 74: Co las – Hutchinson 3693: Co ama.

Ibáñez, A 6489: Co woo – Ibañez, D 195: Ch aca – Ibarra Manríquez 562: Co sca; 641: Co dir; 652: Co sca; 875: Co sca; 996: Co dir; 2108: Co sca; 3353: Co dir; 3395: Co dir; 3400: Co dir; 3709: Co pul – Ibisch 970005: Co com; 990142: Co bec – Ichaso 30: Co spl – Idárraga, A 27: Co sca; 2911: Co pul; 4924: Co las; 5995: Co sca; 5999: Co gla – Idrobo 48: Co spl; 449: Co las; 757: Co mac; 834: Co gla; 844: Di str; 887: Co mac; 937: Co sca; 938: Co mac; 1420: Co sca; 1762: Co cor; 2013: Co sca; 2027: Co ant; 2392: Co kru; 2519: Co mac; 2666: Co sca; 4714: Co kru; 5314: Co lim; 6406: Co sca; 6426: Co ara; 6433: Co lon; 6718: Co ara; 6805: Co spl; 8961: Co las; 9437: Co spl; 10952: Co ant – Ijjasz 657: Co ara; 682: Co ara – INBio 106: Co pul – Infante Betancour 503: Co spl – Irwin 157: Co ara; 7086: Ch aca; 7623: Ch aca; 9066: Ch sub 10358: Ch aca; 11183: Co spl; 11898: Co spl; 14360: Co spl; 16003: Co ara; 16339: Co ara; 19140: Co spl; 19440: Ch sub; 21142: Co spl; 25422: Co spl; 34721: Co spl; 47888: Co sca; 48088: Co sca; 48229: Co sca; 48237: Co spl; 54741: Co cla; 55985: Co sca; 57522: Co ara – Ishiki 2400: Co mol.

Jack 7055: Co pul; 7643: Co pul; 8290: Co mac – Jacobs 2090: Co kun; 2099: Co sca; 2153: Co pul; 2166: Co kun; 2216: Co pul; 2294: Co bra; 2424: Co pul; 2724: Co pul; 2795: Co pul; 2914: Co mal – Jacquemin 1492: Co spl; 1502: Co spl; 1512: Co spl; 1573: Ch con; 1815: Ch cur – Jaimes-R 1247: Co cha – Jangoux, J 738: Co sca; 1492: Co sca; 1607: Co sca – Jangoux, JIG 10210: Co pul – Janovec 1947: Co sca – Jansen-Jacobs 182: Co sca; 328: Co sca; 398: Co spl; 941: Co sca; 1270: Co mac; 1520: Co sca; 2212: Co spl; 2347: Co ara; 2956: Co sca; 3309: Co sca; 4472: Co mac; 4570: Co ara; 4917: Co ara; 5207: Co sca; 5213: Co cla; 5275: Co sca; 5574: Co ara; 5575: Co spl; 5725: Co sca; 6160: Ch con; 6226: Co cla; 6356: Co sca; 7035: Co sca – Janssen 39: Ch lan pul – Janzen 10043: Co mal; 10750: Co pul; 11073: Co kun; 11540: Co com; 11608: Co vil; 11610: Co com; 11611: Co com; 11641: Co vil – Jara 536: Co vil; 629: Di str – Jaramillo Asanza 71: Co sca; 116: Co asp; 150: Co sca; 2218A: Co asp; 2219: Co asp; 2785: Co sca; 2789: Co sca; 2977: Co erc; 3062: Co sca; 3235: Co erp; 3252: Co erc; 3601: Co erc; 3612: Co ama; 4191: Co sca; 4605: Co sca; 5180: Co pul; 5180A: Co pul; 6749: Co kun; 6898: Co ara; 7590: Co kun; 8451: Co ara; 8484: Co las; 8506: Co sca; 8508: Co las 11000: Co kru; 11185: Co sca; 11412: Co sca; 11604: Co ara; 12785: Co sca; 13460: Co sca; 14869: Co sca; 14887: Co las; 14927: Co sca; 15017: Co sca; 15101: Co sca; 15961: Co sca; 15962: Co erc; 16209: Co ama; 18738: Co sca; 18738B: Co sca; 20773: Co sca; 20795: Co sca; 23298: Co lim; 24445: Co kun; 26307: Di str; 27216: Co sca; 27391: Co sca; 27924: Co sca; 28570: Co ama; 28640: Co sca; 30228: Co gib; 30714: Co erp; 30904: Co ama; 31199: Co las; 31333: Co las; 31335: Co erp; 31534: Co ama; 31718: Co erp – Jaramillo-Mejía 114: Di str; 377: Co sca; 395: Co ara; 1139: Co ara; 1215: Co sca; 2096: Co sca; 8128: Co sca – Jardim, A 56: Co pse; 1265: Co ara – Jardim, IG 868: Co sca; 1744: Co spl – Játiva 388: Co pul; 570: Di str; 731: Co mac; 2180: Co lim; 2216: Di str – Jefferson-George 7411: Co woo – Jenman 23: Co spl; 602: Co ara; 609: Co ara; 1995: Co sca; 2075: Ch con; 2076: Ch conl; 4316: Co vil; 4963: Co sca; 6038: Co vil; 6504: Co cla; 6505: Co cla; 7042: Ch con; 8690: Co ara – Jésus Jiménez 2328: Co spc; 3503: Co ara; 4167: Co spc – Jiménez, B 186: Co ara – Jimenez, M R 3075: Co lae – Jiménez, O CR 36602: Co mon – Jiménez, S 1122: Co sca – Jiménez-B 289: Co com; 322: Co spl – Jiménez Barrios 316: Co pul – Jiménez C 31155: Co pic – Jiménez-Escobar 877: Co lon; 1121: Co sca; 1158: Co ara; 1514: Co ara – Jiménez Madrigal 2317: Co pul; 2338: Co lim; 2342: Co vil – Jiménez Muñoz 693: Co las; 2102: Co pul; 2209: Co wil; 2834: Co woo; 2873: Co sca; 3434: Co mal – Jiménez-Saa 1315: Di str – Johnson, EP 132: Co pul – Johnson, W 532: Co mac – Johnston 29: Co sca; 36: Co sca; 91: Co ara; 213: Co ara; 240: Co mac; 991: Co pic 1344: Co pul; 1648: Co com – Jones 3227: Co pul – Jonker 862: Co cla; 886: Co spl – Jonker-Verhoef 98: Co sca ; 182: Co ara; 212: Co sca; 250: Co ara; 380: Co spl – Jørgensen 61236: Co sca; 61239: Di str – Juncosa 598: Co sca; 1148: Di str; 1216: Co lim; 1218: Co lim; 1328: Co kun; 1349: Co gla; 1358: Co kun; 1683: Co lim; 1709: Co las; 2027: Co sca; 2027A: Co pul; 2030: Co vil; 2065: Co mac; 2419: Co cor; 2542: Co sca; 2546: Co sca; 2548: Co kun.

Kainer 33: Co sca – Kalloo B 1566: Co mac – Kappler 1460: Co ara – Karney 17: Co sca – Karwinsky 853: Co pic; 855b: Co pul – Kasent RBAE 10: Co lim – Kayap 214: Di rur; 877: Co ama 1092: Di rur; 1400: Di rur; 1457: Co sca – Kayuk RBAE 1: Co sca – Kegel 480: Co ara; 807p.p.: Co sca – Kellerman 5284: Co pic; 7409: Co pul – Kelsall 612: Co sca – Kennedy 53: Co pul; 147: Co cor; 226: Co sca; 430: Co cur; 487: Co cur; 524: Co wil; 712: Co spl; 746: Co leu 1080: Co pul; 1116: Co all; 1217: Co pul; 1246: Co bra; 1358: Co acr; 1366: Co ama; 1463: Co pic; 1509: Co all; 1560: Co pul; 1694: Co mac; 1695: Co sca; 1733: Co ara; 1765: Co cur; 2815: Co lim; 2881: Co sca; 3033: Co cur; 3204: Co las; 3246: Co pul; 3313: Co all; 3415: Co cur; 3479: Co pul; 3483: Co all; 3526: Co sin; 3671: Co sca; 3692: Co sca; 3760: Co mon; 3788: Co bra; 4618: Co pul; 4675: Di str; 5293: Co vag; 5513: Co cla; 5516: Ch con; 5525: Ch cur – Kenoyer 233: Co pul; 235: Co mac; 236: Co kun; 237: Di str; 1560: Di str – Kernan 223: Co lim; 284: Co kun; 438: Co sca; 1134: Co las; 1139: Co ste – Kessler 354: Co sca; 7434: Co com – Keyser 1303: Co sca – Killeen 5039: Co sca; 6279: Co ara; 6318: Co ara; 7168: Ch aca; 7993: Co ara – Killip 575: Co ant 14670: Co pul; 14732: Co sca; 14907: Co ara; 14931: Co ant; 22632: Co sca; 22912: Co sin; 23030: Di arg; 23708: Co web; 23832: Co lae; 24556: Co sca; 25317: Co jur; 25831: Di arg; 26004: Co sca; 26326: Di app; 26476: Co jur; 26748: Co sca; 27235: Co sca; 27266p.p.: Co ara & Co sca; 28000: Co zin; 28736: Co sca; 29017: Co zin; 29591: Di app; 29850: Co sca; 30225: Co sca; 30659: Co ara; 33072: Co sca; 33328: Co lim; 33531: Co las; 33555: Co kun; 33572: Co pul; 34115: Co pul; 34225: Co erp; 34497: Co spl; 35031: Co sca; 35390: Di str; 35475: Co las; 35554: Co pul; 35568: Di str; 35630: Co sca; 37788: Co spl; 38691: Co woo; 38759: Co cor; 39039: Di str; 39102: Co sca – King 791: Co pic; 792: Co pul – Kirchoff 88-148: Di str; 6178: Co mal – Kirkbride Jr 26: Co all; 189: Co vil; 287: Co kun; 324: Co woo; 373: Co spl; 1162: Co kun; 1180: Co cal; 1212: Co kun; 1485: Co lim; 1486: Co kun; 2758: Co sca; 2804: Co spl – Kirsch 27: Co mon; 28: Co mon; 29: Co mon – Kjaer-Pedersen 185: Co sca; 250: Co sca; 251: Co sca; 282: Co sca; 521: Co sca; 536: Co sca; 561: Co sca; 570: Co sca; 601: Cozin; 676: Co sca; 704: Co sca; 711: Co sca; 816: Co sca; 824: Co sca; 849: Cozin; 876: Co sca; 1190: Co sca; 1191: Co sca; 1192: Co kru – Klein 2865: Co ara – Klitgaard 99438: Co cha – Klug 922: Co sin; 1016: Co sin; 4156: Mo uni – Knab-Vispo 218: Co sca; 465: Co ara – Knapp 1001: Co kun; 1006: Co all; 1025: Di str; 1100: Co cur; 1291: Co pul; 1292: Co pul; 1728: Co pul; 1912: Co mac; 2438: Di str; 3182: Co pul; 3204: Co las; 3871: Co pul; 4116: Co cur; 4519: Co cal; 4693: Co woo; 4705: Co sca; 5174: Co las; 5230: Co kun; 5460: Co com; 5524: Co cal; 5666: Co all; 5676: Co vil; 5759: Co cur; 5896: Co kun; 6615: Co sca; 6619: Co erc; 6969: Co asp; 7252: Co sca – Kodjoed 17: Co spl – Koemar 14: Co sca; 73: Co cla – Kohn 1026: Co ama; 1132: Co sca – Køie 4780: Co sca; 4818: Co mac – Kollmann 1267: Ch cus – Korning 47411: Co sca; 58737: Co kun – Kramer 2198: Co ara; 2315: Co ara; 2356: Co ara; 2598: Co spl; 2622: Co ara; 2664: Co cla; 2795: Co sca; 3158: Co sca – Krapovickas 23068: Co spl; 31529: Co ara; 31623: Co com; 31737: Ch aca; 31948: Co spl; 34895: Co sca; 42986: Co spl; 43072: Co ara – Kress 76-542: Co lim; 76-543: Co vil; 76-545: Co pul; 76-557: Co wil; 77-719: Co pul; 77-873: Co woo; 78-989: Di rur; 78-1014: Co ste; 80-1162: Co pul; 80-1236: Co all; 86-1927: Co cur; 86-1978: Co cal; 86-1988: Co cur; 86-1995: Co pul; 86-2006: Di str; 88-2538: Co sca; 89- 2556: Co ant; 89-2563: Di cry; 89-2564: Co kun; R 3028: Co spc; 90-3124: Co cha; 91-3273: Co las; 91-3299: Co kru; 91-3302: Co lon; 91-3368: Co spl; 94- 3793: Di rur; 94- 3901: Co kun; 94-4005: Co ste; 94- 4073: Co wil; 94-4138: Co wil; 94-4312: Co wil; 94-4507: Co kun; 94-4869: Di str; 94-4912: Co wil; 94-4917: Co wil; 96-5730: Co lon; 97- 6126: Co kru; 99-6356: Co cha; 6702: Co asp; 6962: Co kru – Krug SP 42074: Co ara – Krukoff 1387: Ch lan lan; 1561: Co kru; 4567: Co ara; 5886: Co acr; 7165: Co kru; 10487: Co com; 10489: Di arg; 24075: Co woo – Kuhlmann 635: Co mac; 680: Ch aca; 891: Co las; 1531: Co ara; 1650: Ch lan lan; 1679: Co spr; 1707: Co ara; 1916: Ch fus; 3125: Co sca – RB 63016: Ch cus – Kujikat 364: Di rur – Kuntze 850: Co sca; 1873: Di str; 2140: Co kun – Kupper 947: Co bar – Kuyper 7: Co ara – Kvist 520: Co sin; 522: Co sca; 747: Co ara; 883: Co sca ; 1425: Co sca; 40116: Co pul; 40173: Co pul; 40312: Co kun; 40372B: Co pul; 40373: Co pul; 40583: Di str; 40700: Co pul; 49103: Co pul; 49113: Co lim; 60285: Co kun.

La Frankie 85-38: Co cal; 85-48: Co lim – La Rotta 461: Co sca; 645: Co las – Labala 608: Co ara – Labiak 6250: Co pra – Lægaard 52243: Co pul – Lamotte 286: Co ara; 287: Co sca; 288: Co sca; 372: Di rur; 469: Co ara; 653: Di rur – Langenheim 3050: Co ant – Lanjouw 106: Co ara; 196: Co ara; 388: Co ara; 413: Co cla ; 1318: Co sca; 1327: Co sca; 1369: Co ara; 2046: Co ara; 2880: Co cla; 3202: Co ara; 3429: Co sca – Lankester 240: Co mon – Larpin 751: Co ert – Laskowski 1451: Co kun – Lasser 1646: Co spl; 3668: Co sca; 3956: Co ara; 4070: Co mac – Lau 2613: Co sca; 2614: Di rur; 2615: Co pul; 2741: Ch cus – Lawesson 43354: Co ara; 43437: Co sca; 43469: Co ara; 43512: Co ara; 43551: Co ara; 44330: Co sca; 44456: Co sca – Lawrance 477: Di str – Lazor 2438: Co woo; 2517: Co pul; 5255: Co vil; 5782: Di str – LBB (Lands Bosbeheer Suriname) 8585: Ch con; 10561: Co spl; 10562: Co spl; 10613: Co spl; 10628: Co spl; 10740: Co ert; 10825: Co ert; 10905: Co cla; 10959: Ch con; – Le Clézio 227: Co pul – Leal 336: Co ara – Lechler 2475: Co sin – Ledezma 13: Co ara; 893: Ch aca; 953: Co ara – LeDoux 2608: Di str – Leitão Jr 32934: Co spl; 32935: Co spl; 34495: Co ara; 34535: Co ara – Lellinger 518: Co pul; 1127: Co pul – Lemée 7129: Co spl – Lemos 6981: Co sca – Lems 3: Co pul; 5061: Co lim – Leng 305: Co sca – Lent 238: Co mal; 519: Co lim; 668: Co pul; 864: Co mon; 995: Co cur; 1257: Co sin; 2740: Co wil; 3168: Co mon; 4083: Co cur – Lentz 1932: Co pul – Léon, Bro 9430: Co pul – León, HA 65: Co vil; 99: Co sca; 241: Co vil; 384: Co mac; 397: Di str; 441: Co pul; 701: Di str – León, J 319: Co mon – Lépiz Villalobos 23: Co mon; 59: Co kun; 250: Co mon; 281: Co cur; 326: Co cur; 406: Co com; 441: Co vil; 442: Co pul; 443: Co kun – Lescarboura 56: Co ara – Lescure 197: Ch cur; 377: Co sca; 521: Ch con; 538: Co spl; 565: Ch cur; 2175: Di str – Lévy 74: Co pic – Lewis, M 38024: Co sca – Lewis, WH 1626: Co vil; 1706: Co pul; 1934: Co las; 2032: Co pul; 2231: Co pul; 2293: Co las; 2304: Co kun; 2811: Co woo; 2935: Co pul; 3210: Di str; 3218: Co woo; 4122: Co sca; 9998: Co lon; 10609: Co cha; 10941: Co sca; 11561: Co sca; 11675: Co sca; 11988: Co sca; 12323: Co sca; 12423: Co ert; 12549: Di rur; 12604: Co sca; 12845: Co erc; 12881: Co sca; 13004: Co sca; 13820: Co lon; 13836: Co cha; 13883: Co cha; 14023: Co cha; 14467: Di rur; 18422: Co sca; 38024: Co sca – Liebmann 14722: Co pic; 14723: Co pul; 14724: Co sca; 14725: Co sca; 14726: Co pul – Liesner 204: Co sca; 1031: Co sca; 1104: Di str; 1121: Co sca; 1387: Co woo; 1890: Co kun; 1970: Co ric; 2101: Co las; 2854: Co kun; 2982: Co lim; 2983: Co sca; 2987: Co pul; 3181: Co las; 5672: Co ara; 5827: Co sca; 5875: Co mac; 7436: Co spl; 7679: Co pul; 7865: Co pul; 8347: Co spl; 9254: Co spl; 9397: Di str; 10423: Di str; 10608: Di str; 10819: Co jur; 11098: Co sca; 11447: Co sca; 11918: Co sca; 12414: Di str; 12641: Di str; 13092: Co spl; 13250: Co ara; 14031: Co ara; 14128: Co pul; 14456: Co mon; 14540: Co cur; 14933: Co wil; 15103: Co sca; 15124: Co pul; 15425: Co nit; 15465: Co mon; 15960: Di str; 16493: Co ara; 16992: Co ara; 17440: Co sca; 17849: Co sca; 19604: Co spl; 20402: Co spl; 20521: Co spl; 24499: Co spl; 24507: Co spl; 25936: Co spl; 26224: Co sca; 26258: Co pul – Lima, J 140: Ch lan lan; 248: Ch lan lan – Lima, JAL 134: Co spl – Lima, L 204: Co acr; 227: Co kru – Lima, LFG 272: Co spl – Lima, SA 53-1525: Co ara – Lindeman 22: Co sca; 46: Co cla; 50: Co cla; 95: Co ara; 97: Co ert 147: Co cla; 388: Co cla; 460: Co spl; 557: Co cla; 595: Co cla; 663: Co ara; 693: Co cla; 3545a: Ch con; 3867: Co ara; 4442: Co ara; 4486: Co ara; 5156: Co sca; 5633: Co spl; 5756: Co ara; 6826: Co spl; –– Lindman A 2485: Ch aca; A 2489: Ch aca; A 2767: Co ara – Linneo Foronda 1435: Co ara – Liogier 11469p.p.: Co sca & Co spc; 11618: Co sca; 14464: Co sca; 14470: Co spc – Lisboa 661: Co sca ; 1264: Ch lan pul – Little 16612: Co ara; 16789: Co ara – Lizarralde 353: Co spl – Llano- Almario 35: Co pul – Lleras P 16602: Co sca; P 16877: Co las; P 16880: Co las; P 17073: Co las; P 17078: Co sca; P 17356: Co sca – Loaiza 109: Di str; 116: Co las; 385: Co las – Lobo, M 74: Co pul – Lobo, MGA 51: Co ert; 179: Co spr; 316: Ch lan pul – Löfgren 2510: Co ara – Lohmann 525: Co ara – Løjtnant 13252: Di rur; 13295: Co sca; 13440: Di str; 14538: Co sca; 15622: Co pul; 15674: Co pul; 15781: Co pul – Lombardi 10476: Ch lan lan – Londoño, AC 1561: Co sca; 1586: Co sca – Londoño, PA 15: Co kun – Long, LE 55: Co kun; 56: Co lim; 57: Co pul; 69: Co mal – Long, RW 3241: Co pul – López 4365: Co sca; 4419: Co sca – López A 2016: Co plo; 2202: Co mac; 2394: Co kun; 2396: Co cor; 6415: Co las; 7482: Co las – López B 118: Co ant – López C 1152: Di str; 2805: Co ama; 2822: Co asp; 2918: Di str; 2994: Di str; 3596: Co com; 3730: Co spl; 3959: Co mac; 4089: Co erp; 4281: Co lon; 4304: Co las; 4556: Co las; 5939: Co sca; 6053: Co sca; 6274: Co lon; 6279: Co las; 6384: Co sca; 6712: Co spl; 7053: Co las; 7078: Co las; 7214: Co spl; 7677: Co spl; 8453: Co ara; 8487: Co fis; 8510: Co erc; 8524: Co las; 8526: Co erp; 8527: Co sin; 10215: Co spl; 10777: Co las; 10808: Co lon; 10961: Co sca; 10986: Co kru – López Cruz 436: Co com – López Luna 93: Co pul – López M 8660: Co las – López, F 55: Co osa – López, M 22: Co pul – López, R 62: Co sca – López-Palacios 2739: Di str; 3173: Co mac; 3224: Co spl; 3256: Co mac; 3346: Co ara; 3416: Co spl; 4589: Co sca; 4606: Co ara; 4614: Co ara; 4615: Co mac – Loredo 1852: Co sca; 1864: Co lim – Lot 699: Co dir – Loureiro INPA 48031: Co las – Lowell 435: Co sca; 490: Co asp – Lowrie 445: Ch aca; 485: Ch lan lan; 526: Ch aca; 604: Ch lan lan – Lozano-Contreras 354: Co sca; 357: Co sca; 357A: Co las; 610: Co sca; 641: Co ara; 655: Co ant; 1743: Co ara; 1783: Di str; 3795: Co pul; 3811: Co sca; 3842: Co sca; 3914: Co mac; 5187: Co kun; 5268: Co las; 5329: Co sca; 5694: Co kun; 5862: Co vil; 6157: Co pul; 7140: Di str; 7397: Co ant – Lugo Sanchez 1456: Di str; 1593: Co lon; 1975: Co sca; 2000: Co lon; 2321: Di str; 2756: Co asp; 2833: Co sca; 2839: Di str; 2843: Di str; 2845: Co lon; 2869: Co sca; 3368: Co sca; 3839: Di str; 4202: Co las; 4325: Co las – Luna 75: Co atl – Lundell 470: Co pul; 2667: Co pul; 4288: Co pul; 6275: Co pul; 15869: Co pul; 16132: Co pul – Luteyn 606: Co mon; 759: Co pul; 842: Co wil; 934: Co vil; 1020: Co pul; 1061: Co vil; 1101: Co mac; 1190: Co pul; 1278: Co kun; 1279: Co vil; 1294: Di str; 1325: Co las; 1351: Di str; 1368: Co pul; 1430: Co woo; 1456: Co vil; 1606: Co all; 1655: Co all; 1824: Co vil; 4759: Co mac; 4762: Di str; 8523: Di str; 8535: Co sca.

Maas 596: Co ant; 670: Co las; 681: Co sca; 708: Co all; 716: Di str; 756: Co kun; 764: Co lim; 769: Co pul; 834: Co cur; 864: Co nit; 873: Co bra; 876: Co mal; 900: Co mon; 1039: Co pic; 1052: Co sca; 1074: Co wil; 1086: Co sin; 1095: Co mal; 1130: Co bra; 1195: Co nit; 1196: Co sin; 1218: Co wil; 1241: Co bar; 1264: Co cur; 1291: Co bra; 1295: Co mal; 1306: Co sca; 1306A: Co pul; 1331: Co cur; 1343: Co mon; 1345: Co nit; 1356: Co kun; 1358: Co vil; 1371: Co cur; 1372: Co bar; 1383: Co wil; 1430: Co gla; 1431: Co pli; 1432: Co las; 1436: Co pli; 1444: Co ste; 1451: Co pli; 1461: Co kun; 1462: Co gla; 1463: Co pli; 1471: Co wil; 1535: Co gla; 1544: Co sca; 1548: Co cal; 1569: Co all; 1571: Co woo; 1600: Co vin; 1602: Co all; 1617: Co com; 1619: Co pli; 1629: Co pul; 1651: Co cur; 1652: Co alo; 1655: Co com; 1710: Co all; 1726: Co pli; 1727: Co gla; 1729: Co all; 1732: Co las; 1741: Co vil; 1744: Co woo; 1773: Co bra; 1783: Co ant; 1900: Co plo; 1918: Co vil; 1919: Co lim; 1920: Co leu; 1965: Co cor; 1965A: Co cor; 1969: Co spl; 2015: Co kun; 2216: Co cla; 2224: Co ert; 2300: Ch con; 2465: Ch con; 2760: Co alo; 2901: Co lim; 2915: Co gib; 2916: Co geo; 2917: Co pul; 3166: Ch con; 3172: Co spl; 3173: Co spl; 3355: Co spl; 3360: Co cla; 3397: Co sca; 3398: Co ara; 3400: Co mac; 3443: Co spl; 3445: Co vil; 3830: Co sca; 3898: Co mac; 4553: Di rur; 4557: Co sin; 4609: Co jur; 4709: Co lim; 4739: Co kun; 4744: Di str; 4759: Co leu; 4768: Co pul; 4795: Co gib; 4906: Co wil; 5006p.p: Co alo; 5026: Co cur; 5332: Co mac; 6000: Co zin; 6017: Mo uni; 6052: Co vag; 6086: Co ast; 6095: Co gua; 6156: Co vag; 6158: Co web; 6224: Co mac; 6274: Co erc; 6313: Di app; 6315: Co ara; 6360: Co lon; 6447: Co spc; 6450: Co spc; 6507: Co pul; 6510: Co lim; 6516: Co kun; 6527: Co leu; 6529: Co geo; 6530: Di str; 6758: Co mac; 6970: Co spl; 6972: Co sca; 69-83: Co sca; 7439: Co sca; 7440: Co spl; 7823: Co vil; 7862: Co sca; 7867: Co ric; 7880: Co las; 7883: Co ric; 7890: Co kun; 7914: Co bra; 7935: Co woo; 7935A: Co woo; 7939: Co pul; 7951: Co mon; 7954: Co cur; 7969: Co nit; 7998: Co sin; 8000: Co cur; 8096: Co ert; 8256: Co kru; 8271: Co las; 8372: Co spc; 858: Co pul; 8598: Co lon; 8608: Co ama; 8656: Co acr; 8660: Ch aca; 8697: Co ara; 8731: Co sca; 8751: Co pse; 8884: Co com; 8885: Co bra; 8999: Co dal; 9036: Co kru; 9153: Ch lan lan; 9155: Co las; 9279: Co dir; 9299: Co zin; 9300: Co jur; 9301: Co gla; 9305: Ch con; 9306: Co cla; 9317: Co ert; 9329: Ch cur; 9334: Co ert; 9335: Co spl; 9381: Co cur; 9409: Co bra; 9475: Co wil; 9501: Co osa; 9507: Co wil; 9512: Co cur; 9514: Co com; 9515: Co wil; 9524: Co nit; 9529: Di str; 9541: Co alo; 9544: Co las; 9563: Co all; 9568: Co vin; 9583: Co ert; 9795: Co spc; 9931: Co mol; 9933: Co gua;10488: Co ant; 10534: Co plo; 10540: Co pic; 10609: Co ert; 10622: Co cla; 10623: Co ert; 10629: Co bra; 10631: Co ant; 10632: Co bar; 10633: Co jur; 10635: Co gib; 10637: Co pul; 10638: Co sca; 10646: Co gla; 10647: Co kun; 10650: Co all; 10653: Di str; 10654: Co leu; 10656: Co vil; 10657: Co sca; 10658: Co cor; 10671: Co cor; 10672: Co gla; 10674: Co pul; 10675: Co plo; 10682: Co leu; 10685: Co kun; 10688: Co cal; 10689: Di cry; 10690: Co ama; 10691: Co asp; 10693: Co lon; 10694: Co sca; 10707: Co con; 10708: Co asp; 10732: Co ama; 10733: Co pit; 77-83: Co acr; 69-83: Co sca; 80-128: Co dir; 71-153: Co vaz; 69-162: Co com; 80-164: Co jur; 68-169: Co pul; 68-219: Ch con; 68-227: Ch cus; 68-228: Co mal; 68-233: Co sca; 68-236: Co pic; 69-236: Co spl; 68-251: Co pul; 68-252: Co pul; 68-254: Co pul; 84-281: Co zin; 67-331: Co spl; 67- 334: Co spl; 67-335: Co spl; 72-347: Co kru; 72-359: Co erp; 68-405: Ch con; 74- 507: Co woo; 74-529: Co vil; 74-553: Co leu; 74-618: Co whi; 68-773: Co cur; 66-1790: Co ara; LBB 10886: Co ara; P 12860: Co vaz; P 12952: Co sin; P 12972: Co dal; P 12978: Co sca; P 13091: Co las; P 13155: Co acr; P 13165: Co vaz; P 13213: Co dal – Macbride 5001: Di arg – MacBryde 1369: Di str – MacDougal 1020: Co kun; 1024: Co mal; 1035: Co pul; 1051: Co mal; 3729: Co ant; 4020: Co cal – Maceda 281: Co sca – Macedo 23: Ch aca 1316: Ch aca; 1993: Ch aca – Machado 33: Ch lan lan – Macía 1529: Co sca – Maciel 538: Ch lan pul – Madison 1189: Co cha; 3165: Co sca; 3230: Co sca; 3515: Co sca; 4623: Co ant; 5028: Co leu; 5038: Di str; 5075: Co cor; 5273: Co geo; 7276: Di str – Madrigal 563: Co lim; 581: Co woo; 620: Co mac; 628: Di str; 809: Co mac; 819: Di str; 866: Co kun; 873: Co woo – Madriñán R 189: Co kun; 196: Co sca; 208: Co sca – Madsen 86796: Co con – Maguire 23628: Co cla; 23926: Co sca; 28056: Co mac; 35815: Co ara; 36740: Co sca; 40064: Co spl; 40824: Ch con; 47024: Ch con; 47070: Co spl; 53513: Co mac; 53995: Co ara; 54242: Co spl; 57147: Ch sub; 61518: Mo uni; 61519: Co sin – Malavassi V 2: Co pic – Maldonado Goyzueta 1617: Co sca – Malme 2630: Ch aca – Mamani M 424: Co ara; 567: Co ara; 846: Co ara ; 1091: Ch aca; 1123: Ch aca – Marcano- Berti 61-1-76: Co ara; 192-981: Di str; 194-981: Co spl; 195-981: Co mac; 198-981: Di str; 204- 981: Co spl; 215-979: Co vil; 260-981: Co pul; 262-981: Co pul; 271-981: Co mac; 272-981: Co spl; 915: Co sca; 937: Co sca; 939: Co sca; 942: Co sca; 946: Co sca; 1093: Co mac – Marcondes-Ferreira 1067: Co spl – Marín Alpízar 34: Co ste; 9: Co kun – Marin, E 103p.p.: Co ara & Co sca; 810: Co spl; 956: Co spl; 961: Co mac – Marín, G 481: Co ara – Marín, S 187: Co pic – Marín-C, CA 182: Co spl; 347: Di str; 348: Co sca; 407: Co erp; 491: Co spl; 626: Co spl; 1957: Co cha; 1980: Di str; 2015: Co ama; 2032: Co sca; 2160: Di str; 2202: Co las; 2468: Di str; 2493: Co cha; 2568: Co sca; 2607: Co sca; 2650: Di str; 2798: Di str; 2992: Co sca – Marín-C, NL 1800: Co spl; 2573: Co cha – Markgraf 3762: Co ara – Marrugo-Rondón 46: Co com – Marshall 6459: Co pul – Martin, GJ 564: Co pul – Martin, RT 1090: Co las; 1162: Co ara; 1281: Co sin; 1834: Co web – Martinelli 6808: Co ara; 7064: Co ara – Martinet 1567p.p.: Co sca – Martínez Calderón 70: Co pic; 3015: Co dir – Martínez Crovetto 10845: Co ara – Martínez Meléndez 1335: Co pic – Martínez Salas 1984: Co pul; 3160: Co mol; 7205: Co pul; 7294: Co pul; 7428: Co pul; 7503: Co mac 10824: Co sca; 12712: Co sca; 15593: Co pul; 15676: Co pul; 15796: Co pul; 20721: Co com; 22721: Co sca; 31068: Co pic – Martínez Sequeira 229: Co pul – Marulanda 690: Co pul; 715: Co mac; 725: Co sca; 1211: Co mac; 1539: Co ara – Mashu RBAE 13: Co asp; 34: Di str – Mason 2377: Co mac – Mattos 10065: Ch con; 10176: Co spl – Matuda 353: Co com; 16379: Co pic; 17278: Co com – Maxon 6510: Co vil; 6990: Di str; 7017: Co pul; 7469: Co pic – Mc Vaugh 15972: Co pic – McCook 1026: Co vil; 1038: Di str – McDade 77- 706: Co pic – McDaniel 16188: Co las; 17865: Co sca; 17946: Co las; 18823: Co ara; 18906: Co ara; 18908: Co sca; 18915: Co las; 19048: Co erc; 19439: Co las; 19504: Co sca; 20372: Co sca; 20504: Co ara; 21074: Co sca; 22501: Co erc; 24212: Co ara; 27561: Co sca; 32906: Co sca – McDonagh 115: Co kun; 115a: Co sca; 468: Co pul – McDowell 106: Co sca; 126: Co mal; 555: Co mal; 752: Co mal; 971: Co sca; 1073: Co bra; 1891: Co sca; 2280: Co spl; 2454: Co spl; 2468: Co sca; 3372: Co sca; 3387: Co ara; 3413: Co ert; 3597: Co cla; 4179: Ch con; 4180B: Ch con; 4185: Co sca; 4238: Co ert; 4322: Co mac; 4469: Co ert; 4939: Co ert –McElroy 9: Co ama ; 281: Di str; 336: Co sca – McLean TRIN 6379: Co mac – McPherson 6820: Co pul; 7634: Co all; 9651: Co kun; 9701: Co cur; 9901: Co wil; 10813: Co sca; 10843: Co pul; 10954: Co las; 11223: Co all; 11562: Co pul; 11703: Co pul; 12252: Di str; 20760: Co kun; 20830: Co sca – Medeiros 270: Co sca; 368: Co ara; 515: Co ara; 666: Di rur; 667: Co gua – Medina, A 46: Co erp – Medina, E 276A: Co ara; 906: Co mac; 943: Co spl – Medina, N 181: Co ara – Meebold 28214: Co vil – Meier 3278: Co spl; 4965: Co sca; 8260: Co com; 8449: Co sca; 9209: Co mac; 9779: Co sca; 9896: Co mac; 11137: Co mac; 11389: Co spl; 14326: Co sca – Mejía, C 189: Co pic – Mejía, MM 6672: Co spc; 6857: Co sca; 691: Co spc; 7891: Co spc; 7955: Co sca; 9528: Co spc; 9598: Co spc; 9808: Co spc; 9936: Co spc; 10158: Co spc; 10488: Co sca; 11221: Co spc; 23341: Co sca – Mélinon 96: Ch cur; 153: Co sca – Mellado 2298: Di arg – Mello Barreto 1458: Co ara; 8612: Co spl – Mello filho 194: Co sca; 2980: Ch cus – Mello Netto R 50948: Co spl – Mello 14: Co sca; 3417: Co ara – Melnyk 100: Co spl; 108: Co mac – Menacho Viruez 912: Ch aca; 957: Co ara – Méndez 8: Co las – Méndez-Vargas 6375: Co pul – Mendonça 32: Co spl; 2206: Ch sub; 2898: Ch sub; 2983: Co spl; 3832: Co spl; 3974: Co ara – Mendoza 3843: Co com – Menéndez L 319: Co pic – Menezes 69: Co spl – Merardo Martínez 13: Co vil; 13A: Co pul; 16: Co lim – Merello 571: Co mal – Mermillier 258: Co spl – Merrill King 6059: Co sca; 6091: Co sca; 6138: Co sca; 6145a: Co sca – Mesa 46: Co kun – Mexia 1854: Co pic – Meyer, H 438: Co ara – Meyer, KM 326: Co jur – Meyer, T 5409: Co ara – Meyrat 228: Co sca; 328: Co mac – Meza 50: Co sca; 71: Co sca; 73: Co sin; 228: Di rur – Mileski 400: Ch lan pul – Miller jr, GS 1046: Co sca – Miller, JS 612: Co pic; 6299: Co pic; 765: Co vil; 800: Di str; 843: Co all; 926: Co kun; 987: Co las 1091: Co pul; 1372: Co mac; 1575: Co spl; 2320: Co sca; 5519: Co pul; 5605: Co erp; 5610: Ch cus – Milliken 108: Co sca; 1700: Co mac – Minorta-C 1066: Co spl – Miralha 296: Ch man – Miranda Bastos 85: Ch con – Miranda-Moyano 145: Co sca; 252: Co sca; 405: Co sca – Mitchell 71: Co pul – Miyagi 513: Co spl – Molina Rosito 1802: Co pul; 2135: Co pul; 6001: Co pul; 6959: Co sca; 7119: Co sca; 8081: Co pic; 8376: Co sca; 8444: Co pul; 10484: Co pul; 10898: Co pic; 10952: Co pul; 11423: Di str; 11426: Co sca; 12060: Co pul; 12129: Co pic; 14986: Co pul; 15042: Co vil; 15181: Co pul; 15790: Co pul; 15865: Co pul; 17225: Co pul; 18059: Co lim; 20823: Co pul; 30566: Co pic; 33657: Co pic; – Molina 2363: Di str – Molino 2227: Ch lan pul; 2432: Ch con – Moncaio 123: Co spl; 22: Co spl – Monro 687: Co pic; 5886: Co cur – Monsalve Benavides 1533: Co kun – Monteagudo Mendoza 4059: Co gua; 5027: Co gua; 7658: Co rub; 9943: Co erp; 10931: Co las; 11957: Di arg; 15220: Di arg; 16204: Di arg; 16686: Co spl; 16895: Di arg; 16970: Di arg – Monteiro 483: Co las; 494: Co dal; 497: Co acr; 601: Co dal; 654: Co las; INPA 53410: Ch lan lan; INPA 53453: Co ara; INPA 53463: Ch lan lan – Montero 2005: Co las; 5308: Co ara – Montes 2516: Co ara – Montes Guarín 128: Co las – Montoya Jiménez 390: Co las; 391: Co las; 849: Co las; 850: Co las; 1037: Co las; 1327: Co ama; 3114: Co sca; 3115: Co sca; 3567: Co vil – Mood R 3184: Co cur – Moonen 210: Co ert – Moore, BH 3654: Co pic – Moore, CE 38: Co pul – Moore, HE 8108: Co pul – Moore, Spencer Le Marchant 679: Ch aca; 813: Co ara – Mora Castro 315: Co pul; 1011: Co pul; 1325: Co nit; 1457: Co kun – Mora Maroto 158: Co pul – Mora O APA 198: Co sca; 1083: Co kun – Mora V 49: Co wil; 183: Co wil; 324: Co mon; 433: Co mon – Moraes R 385: Co sca; 1327: Co sca; 1693: Co sca – Moraga López 208: Co las – Moraga Medina 236: Co pul – Morales Can 4365: Co pul – Morales Quirós 21: Co wil; 129: Co pul; 144: Co lim; 284: Co kun; 300: Co pul; 411: Co sin; 1556: Co wil; 1566: Co vil; 2099: Co ric; 2746: Co mon; 2747: Co mon; 2805: Co pul; 2855: Co wil; 5659: Co sca; 6440: Co sca; 6463: Co wil; 8076: Co sca; 13084: Co pul; 13228: Co sin; 13309: Co mal; 13872: Co lim – Morales, ME 600: Co sca – Morales, P 426: Co ant – Morawetz 85: Co sca; 88: Co las – Moreno B 103: Co pul; 485: Co sca; 93: Co mac – Moreno M 20: Co vil; 54: Co lim; 54A: Co sca – Moreno, PP 842p.p.: Co lim; 1433p.p.: Co pic; 1487: Co pic; 2635: Co pic; 4104: Co pul; 4738: Co pic; 6285: Co pic; 6461: Co pic; 9517: Co pic 10205: Co pul; 10302: Co pic; 10570: Co sca; 11951: Co woo; 12184: Co pul; 12203: Co pul; 12219: Co woo; 12315: Co woo; 12355: Co woo; 12506: Co woo; 12549: Co pul; 12615: Co pul; 12801: Co kun; 12807: Co kun; 12918: Co mal; 13121: Co pul; 13214: Co woo; 13255: Co woo; 13772: Co pic; 14711: Co sca; 14803: Co pul; 14913: Co pul; 15064: Co pul; 15159: Co sca; 15168: Co pic; 15224: Co woo; 15253: Co pic; 16183: Co pul; 16533: Co pic; 16881: Co pic; 17141: Co pul; 17198: Co pul; 17302: Co pul; 17345: Co pul; 17538: Co pul; 19419: Co pul; 21120: Co pul; 23063: Co bra; 23201: Co pul; 23212B: Co mal; 23673: Co pul; 23813: Di str; 24092: Di str; 24107: Co pul; 25958: Co pul; 26251: Co mac; 26252: Co kun; 26275: Co mal; 26308: Co pul; 26758: Co pul; 28434: Co pul – Moreno, R 40: Co spl – Moretti 694: Ch lan pul; 697: Co ara – Mori 164: Co sca; 1964: Co sca; 2194: Co las; 2663: Co pul; 3119: Co cur; 3818: Co cur; 3835: Co kun; 4264: Co pul; 4276: Co sca; 4284: Co lim; 4575: Co sca; 5017: Co pul; 5493: Co las; 6027: Co pli; 6124: Co alo; 6699: Co cur; 6819: Di str; 6994: Co vil; 7203: Co wil; 7660: Co kun; 7865: Co cur; 9351: Ch sub; 9551: Co sca 10753: Ch sub; 18151: Ch con; 18420: Co cla; 18441: Co cla; 18855: Co cla; 18955: Co sca; 18995: Co ert; 21098: Co sca; 22367: Co ara; 22477: Co sca; 22478: Co ara; 22638: Co sca; 23014: Co cla; 24001: Co sca; 24219: Co sca; 25530: Co cla; 25538: Co ert – Morillo 3410: Co spl; 8824: Co sca; 9143: Co sca; 9282: Co pul – Morrone 2025: Co ara – Morton 2466: Co pic; 5115: Co sca – Mosén 2967: Co ara; 2968: Co spl – Mota 277: Co sca; 295: Co ara – Mowbray, RN 69423: Co ama; 69975: Co sca; 70326: Di str – Mowbray, T 1432: Co pul – Moya 132: Co lon; 191: Co ara; 271: Co sca; 311: Co sca; 363: Co sca; 432: Co lon; 487: Co sca; 488: Co erp; 608: Co sca; 685: Co lon – Müller 8496: Ch lan lan – Muñoz 3: Co sca – Murillo 1539: Co erp; 1691: Di str; 1715: Co vil; 1937: Co sca – Murillo Rodríguez 15: Co vil – Murphy 119: Co sca; 146: Co sca; 182p.p.: Co ara & Co erc; 497: Di str – Mutchnick 454: Co ara; 1060: Co sca – Mutis 2534: Di str; 4577: Co pul; 4581: Di str; 4655: Co lim.

Naessany 56: Co sin; 57: Co sca – Nagata 420: Co woo; 421: Co pul; 2480: Co sca; 2481: Co pic; 2588: Co jur; 2589: Co erp; 2743: Co sca; 2770: Co las; 2771: Co erp; 2810: Co lim; 2811: Co gib; 2824: Co kru; 2878: Co jur; 2996: Co pic; 3011: Co vaz; 3012: Co bra; 3013: Co mac; 3014: Co ara; 3115: Co vaz; 3364: Co kun; 3452: Co erp; 3453: Co vaz; 3454: Co erp; 3455: Co erp; 3456: Co woo; 3550: Co sca; 3781: Co vaz; 3782: Co ara; 3783: Co sca – Nakagomi 18: Co spl – Naranjo 131: Co sca; 148: Di str; 332: Di rur; 334: Di str; 335: Co asp; 404: Di rur; 736: Co sca – Nascimento, J 62: Ch lan pul – Nascimento, JR 726: Ch man – Nascimento, OC 693: Co spl; 1097: Ch lan pul – Nash 575: Co sca; 1218: Co ara – Navarro V 800: Co wil – Navarro-L 508: Co fis; 968: Co las; 1049: Co las; 1113: Di app; 1122: Co erc – Nee 7191: Di str; 7193: Co vil; 7655: Co kun; 7924: Co sca; 8928: Co woo; 11066: Di str; 11502: Co las; 11692: Co woo;17843: Co sca; 17866: Co sca; 18453: Co pul; 19983: Co pul; 22759: Co dir; 22762: Co sca; 25100: Co sca; 27684: Co pic; 27855: Co pul; 27865: Co kun; 28424: Co sca; 34198: Co com; 35973: Di arg; 36081: Co sca; 36085: Co ara; 37356: Di arg; 38030: Co com; 38076: Di arg; 39247: Co com; 40559: Di rur; 40565: Co ara; 40884: Co sca; 40938: Di arg; 41707: Co spl; 41925: Di arg; 41962: Co com; 42842: Co pra; 43251: Co sca; 43259: Co ara; 44928: Co com; 44937: Co ara; 44981: Di arg; 45216: Co sca; 45217: Co sin; 45233: Di arg; 45989: Di rur; 45999: Co sca; 46000: Di rur; 46050: Co ara; 46413: Di rur; 46495: Co sca; 46507: Co com; 46823: Co pul; 46925: Co pul; 46926: Co mac; 46931: Co pul; 47332: Co pul; 48392: Co ara; 48421: Co sca; 48526: Co gua; 50169: Co com; 51554: Co sin; 51661: Co sca; 51683: Co sca; 53969: Co sca – Neill 214: Co pul; 784: Co pul; 11581: Co pul; 1897: Co lim; 3333: Co lim; 3564: Co pul; 3570: Co lim; 3671: Di str; 3988: Co pul; 4458: Co pul; 6281: Co sca; 6288: Di str; 8791: Co ama; 9133: Co sca – Nelson, BW 510: Ch lan lan; 514: Co sca – Nelson, CH 2745: Co pul; 3052: Co sca; 4205: Co kun; 9063: Co pic; 9147: Co lim; 9216: Co pul; 9226: Co pul – Nelson, EB 4279: Co pul; 4460: Co woo; 4824: Co lim – Nelson, EW 911: Co pic – Nepokroeff 718: Co pul; 723: Co kun – Neves Armond R 50979: Co ara – Nevling 1464: Co pul; 1526: Co mol; 2580: Co pul – Newton Santos R 50981: Co ara – Nicolalde-Morejón 1319: Di str – Nielsen 80813: Ch fra – Niemeyer 8: Co sca; 29: Co com – Novelo Retano 76: Co pul – Núñez Vargas 9544: Co acr; 9654: Co sca; 9961: Di arg; 11945: Co lae; 12080: Co acr; 12976: Di rur; 14049a: Di arg; 14644: Di rur; 19059: Di arg; 20039: Co jur; 20873: Co jur; 21718: Co sin; 21757: Co jur; 23785: Co obs – Núñez, F 717: Co pul – Núñez, T 71: Di str; 306: Co gib; 712: Co sca – Nusbaumer 3928: Co spl.

Obando 284: Co ant – Occhioni 104: Co spl – Odonne 79: Co lae; 742: Co ert – Oersted 14720: Co woo; 14721: Co woo – Ohba 1024: Co sca; 1224: Di str – Ojeda 33: Co sca – Oldeman 12: Co sca; 17: Co sca; 40: Co sca; 151: Co spl; 255: Co ara; 298: Co spl; 307: Co spl; T. 495: Ch con; B. 553: Ch cur; T. 557: Ch con; T. 736A: Co sca; B. 959: Co ara; 1181: Co ert; 1412: Co cla; B. 1564: Co spl; B. 1704: Co sca; B. 1715: Co ara; 1928: Co ert; B. 2057: Co cla; 2668: Co ert; 2944: Co sca; B. 3072: Co sca; 3183: Co ert; B. 3187: Co ara; B. 3981: Ch con – Oldenburger 192: Co ara; 740: Co ara; 1300: Co ara; 1301: Co ara – Oliveira, E 66: Co spl; 160: Co sca; 215: Co las; 732: Co ara 1324: Co ara; 1392: Co ara; 1507: Co spl; 1638: Co spl; 1977: Co ara; 4317: Co spl; 4768: Co spl – Oliveira, FCA 182: Co spl; 1117: Ch sub; 1173: Ch sub; – Oliveira, GM 3096: Ch lan pul – Oliver 3688: Co sca – Øllgaard 34594: Co sca; 34619: Co asp; 34641: Co sca; 34692: Co asp; 34837: Di rur; 34867: Co cha; 34911: Di rur; 34957: Co sca; 35014: Co ara; 35265: Co erp; 35416: Co sca; 37749: Co pul; 37825: Co pul; 38942: Di str; 39107: Co kun; 57071: Co ama; 57169: Co pul; 57228: Co cor; 57302: Co kun; 57415: Co kun; 57651: Co kun; 90385: Co con; 99596: Cozam – O’Neill 8518: Co pul; 8523: Co pul; 100808: Co pul – Oppenheimer 1215: Co pul – Ordoñez 48: Co mac – Orellana 1117: Co sca – Orozco 395: Di str; 478: Co las; 503: Co las; 666: Co sca; 709: Co lim; 813: Co las; 1136: Co leu – Ortega O 1162: Co pul – Ortega U 330: Co sca – Ortega, F 615: Di str ; 1263: Co mac – Ortiz S 39: Ch aca; 66: Co spl – Ortiz V 521: Co rub – Ortiz, A 800: Co kun; 803: Co kun – Ortiz, F 26: Co mac; 32: Co pul; 79: Co pul; 83: Co pul; 165: Co pul; 275: Co pul; 357: Co pul; 544: Di str; 545: Co pul; 1283: Co sca; 1642: Co pul; 1728: Co lim; 1887: Co lim; 2149: Co pul – Osores de Diaz 219: Co asp; 220: Co sca – Osorio 2: Co ant – Ostenfeld 75: Di str; 92: Co vil – Otero 102: Co woo; 742: Co ant – Oviedo 23: Co sca.

Pabón E 141: Co spl; 279: Co spl; 527: Co ara; 836: Co fis – Pabst, GFJ 9215: Co ara – Padilla 331: Co pic – Paguaga 170: Co pul – Paixão 516: Co ara – Palacio 3: Co mac – Palacios Cuenca; 658: Co cha; 782: Co lon; 2900: Co sca; 3458: Co sca; 6931: Co kun; 7450: Co pul; 10270: Co sca – Palacios H 168: Co cor – Palacios, PA 307: Co sca; 488: Co cha; 1333: Co sca; 2671: Co cha – Palma 415: Co sca – Paniagua Zambrana 823: Co ara; 1016: Co sin; 1066: Co vag; 2149: Ch aca; 2183: Ch aca; 4803: Co sca – Parada-Gutierrez 19: Co sca; 1311: Co ara; 1320: Co sca – Parrado 89: Co sca – Pastore 676: Co spl – Patiño-Ch 114: Co cor; 114A : Co kun – Paz R 158: Co ara; 201: Co sca – Peck 892: Co pul – Peckolt 99: Ch cus – Pedersen 10702: Co ara – Pedraza 1239: Co asp – Peele; 344: Ch cus 898: Co pic; 1410: Co pic – Peña 614: Ch lan lan; 628: Di rur – Peña-Chocarro 65: Co pse – Peñaloza 4936: Co sca – Peñaranda 436: Co sca – Pengel 1: Co spl – Penland 179: Co ore – Pennell 47: Co ara; 3295: Co pul; 3304: Co las; 8486: Co ant; 8609: Co pul – Penneys 294: Co wil; 555: Co mon – Pennington, RT 430: Co sca – Pennington, TD 56SD: Co gib – Peñuela 66: Co vil – Perea 95: Co sca; 341: Di arg; 408: Co sca; 426: Di arg; 4582: Co sca – Pereira, BAS 2009: Ch sub; 2251: Ch sub – Pereira, E 552: Co ara; 4074: Co ara; 8202: Co spl; 30535: Co spl – Pereira-Silva 7401: Co spl; 8758: Co spl – Pérez Arbeláez 757: Co cha; 760: Co kun; 760A: Co mac – Pérez, AJ 504: Co pul; 1203: Co sca; 6281: Co erp; 9713: Co sca – Pérez, V 92: Di str – Pérez-Levis 17: Co spl – Pérez-Zabala 370: Co mac; 2576: Co ant; 2585: Co ant; 2586: Co ant; 2588: Co pul; – Perino 3061: Co pul; 3111: Co sca – Perry 1177: Co sca – Pesha B 53: Co acr – Pessôa, C 431: Co sca – Pessoa, E 1346: Co atl; 70: Co atl – Pessoal do CPF 6118: Co mac – Pessoal do Museu Goeldii MG 9606: Co ara – Peters 10: Co ara; 167: Co sca; 168: Co ara – Peterson 6310: Co woo; 6385: Di str; 6629: Co lim; 6643: Co woo; 6673: Di str; 6966: Co lim; 7053: Co pul; 7100: Co lim; 7182: Di str; 8542: Co woo – Pfeifer 2046: Co pul – Philcox 3289: Co ara; 3395: Co spl; 4000: Ch aca; 4061: Ch aca – Philipson 1312: Co spl; 1613: Co mac – Phillips 88: Co vaz; 366: Di arg – Pickel 942: Co sca; 2759: Co spl; 3966: Co sca – Pilger 438: Co ara – Pinheiro 1742: Ch sub; 1747: Ch cus; 1771: Co ara – Pinilla 69: Co las; 100: Co acr – Pinilla-Moreno 386: Co sca; 393: Co ara; 475: Co com; 613: Co sca – Pinto E 72: Co las; 79: Co sca; 88: Co las; 117: Co lim; 251: Co spl; 789: Co spl; 808: Co spl; 913: Co sca; 94: Di str; 940: Co spl; 968: Co sca 1123: Co spl; 1416: Co spl; 1453: Co spl; 1604: Co spl; 6149: Co mac; 6394: Co sca; – Pinzón 181: Co ara – Piper 5443: Co mac; 5444: Co vil; 5778: Di str – Pipkin 182: Di str – Pipoly 3751: Co lim; 3863: Co sca; 3991: Co pul; 4198: Co lim; 4323: Di str; 4327: Co lim; 4547: Co lim; 4847: Co pul; 5121: Co pul; 5976: Co sca; 6878: Co sca; 8067: Co sca; 8115: Co sca; 8218: Co sca; 8331: Co sca; 8665: Co ara; 11270: Co ara; 11639: Co ara; 11647: Co sca; 11727: Co ara; 13165: Co sca; 13756: Co sca; 13900: Co lon; 13902: Co erc; 14134: Co erc; 14168: Co sca; 15149: Co ara; 15206: Di app; 15323: Co sca; 15671: Di app; 16242: Co sca; 16917: Co plo; 17659: Co plo; 21093: Co plo; 21526: Co kun – Pirani 2732: Co spl; 3122: Co spl; 3909: Ch sub – Pires, JM 724: Co ara; 953: Co las; 961: Co acr; 1026: Co acr; 1281: Co ara; 1453: Co ara; 3482: Co ara; 3559: Co ara; 3737: Co ara; 3824: Co spr; 3852: Co kru; 6143: Co ara; 11765: Co sca; 50384: Co ara; 50430: Co sca; 50482: Co sca; 50814: Co sca; 52137: Co spl – Pires, MJP 782: Ch lan pul; 812: Co ara; 871: Co ara; 899: Ch con; 982: Co ara – Pirie 99: Co ert – Pitman 247: Co sca; 894: Co lim; 1028: Co pul – Pittier 1622: Co pul; 2307: Di str; 2359: Co mon; 2373: Co pul; 3776: Co vil; 3779: Co mac; 3894: Co kun; 4051: Co pul; 4084: Co pul; 4137: Co woo; 6690: Co vil; 7610: Co pul; 7796: Co pul 10884: Co ara; 12091: Co mac; 12419: Co pul; 13443: Co pul; 13844: Co spl; 14166: Co mon; 15609: Co mac; – Plotkin 205: Co sca; 284: Ch con – Plowman 2137: Di str; 2273: Di str; 2274: Co mac; 2298: Co acr; 2335: Co erc; 2336: Di app; 2337: Co kru; 2357: Di app; 2429: Co kru; 2463: Co lon; 2464: Co erc; 2474: Co ara; 2504: Co kru; 2529: Co sca; 2580: Co mac; 2581: Co sca; 3555: Co com; 3999: Co asp; 4366: Co sca; 4385: Co asp; 4396: Co whi; 4454: Co gib; 4455: Co gib; 4475: Co asp; 4709A: Co web; 4725: Di arg; 4766: Co gua; 4996: Co gua; 5011: Co acr; 5027: Co gua; 5029: Co lae; 5049: Di rur; 5054: Co obs; 5054A: Co obs; 5054B: Co lae; 5660: Co lae; 5702: Co lae; 5707: Co sin; 5755: Co sin; 5773A: Co jur; 5775: Co obs; 5791: Co obs; 5915: Di arg; 5988: Co ama; 6050: Mo uni; 6351: Co acr; 6363: Co las; 6404: Co sca; 6417: Co ara; 6470: Co kru; 6475: Co las; 6500: Co erc; 6517: Co kru; 6520: Co lon; 6560: Co ara; 6613: Co lon; 6755: Co sca; 6983: Di app; 6987: Co erc; 7502: Di rur; 7522: Co las; 7595: Co obs; 7754: Co spl; 8126: Ch aca; 8338: Ch aca; 8442: Co spl; 8448: Ch aca; 8505: Ch lan pul; 9628: Ch lan pul; 9778: Ch lan pul; 11362: Co las; 11468: Co sin; 11560: Co jur; 11585: Co lae; 11608: Co sca; 11655: Co las; 11668: Co las; 12000: Co jur; 12122: Co kru; 12605: Co sca; 13305: Co osa; 13306: Co asp – Poeppig 2: Co ara; 1815: Co sca; 1942: Co las; 2116B: Mo uni – Poiteau 1: Co sca; 79: Co sca; 159: Co sca – Pollard 78: Co pul – Pomari 24: Co spl – Ponce 32: Co pul – Poncy 200: Ch cur; 1189: Co ara; 2772: Co sca; 2831: Co spl; 2833: Co cla; 2857: Ch lan pul – Poole 1923: Co spl – Porter, DM 4648: Co cur; 4750: Co sca – Poulain 76: Co las – Poulsen 1224: Co erp; 78438: Co las; 78511: Co sca; 78585: Ch fra; 78793: Ch fra; 79303: Ch fra; 79893: Co las; 80149: Ch fra; 80496: Ch fra; 80518: Ch fra; 80809: Co las; 80973: Co las – Pounds 409: Co wil – Poveda Álvarez 1212: Co kun – Pozada 2091: Co sca; 2477: Di app; 2481: Co sca; 2494: Co sca; 3051: Co sca; 3054: Co sca; 3200: Co las – Prado G 10: Co pul; 51: Co pul; 68: Co vil; 69: Co pul; 157: Co pul – Prado M 13: Co pul – Prance 1712: Co sca; 2012: Co las; 2019: Co ert; 2288: Co sca; 2486: Ch lan lan; 2773: Co sca; 2917: Co vaz; 3151: Co pra; 3227: Ch lan lan; 3445: Co kru; 3529: Co sca; 4303: Co sca; 4325: Co mac; 4583: Co spl; 5218: Co pra; 5725: Co kru; 6163: Co ara; 6672: Co pra; 6917: Co spl; 7416: Ch aca; 7420: Ch lan lan; 7444: Ch lan lan; 7623: Ch aca; 7918: Ch lan lan; 7973: Co kru; 8103: Co sca; 8175: Co kru; 8227: Co pra; 8338: Ch aca; 8459: Co pse; 8478: Co kru; 8866: Co pra; 8867: Co ara; 8923: Ch lan lan; 8937: Co kru; 8975: Ch aca; 9268: Co sca; 9906: Co spl; 10213: Co sca; 10350: Co ara; 10730: Co ara; 10865: Co spl; 10918: Co sca; 11161: Co ara; 11253: Co ara; 11529: Ch man; 11966: Co sca; 11971: Co sca; 11982: Co ara; 12064: Co vaz; 12064a: Co erp; 12210: Co kru; 12230: Co dal; 12265: Co pra; 12265A: Co las; 12334: Co sin; 13399: Co sca; 13405: Co ara; 13460: Co sca; 13527: Co sca; 13684: Co kru; 14455: Co acr; 14675: Co sca; 14690: Co acr; 15068: Co sca; 15300: Co pra; 15418: Co ara; 15879: Co spl; 15917: Co spl; 16276: Co ara; 16463: Co kru; 16516: Co las; 16692: Co sca; 16824: Co ara; 16854: Co las; 18232: Co sca; 23874: Co las; 23923: Co sca; 24069: Co kru; 24155: Co las; 24189: Co ara; 24198: Co ara; 24412: Co sca; 24523: Co ara; 24656: Co sca; 24823: Co spl; 25141: Co sca; 25218: Ch aca; 25327: Ch lan lan; 25343: Co sca; 25477: Co ert; 25527: Co las; 25875: Co spr; 25906: Co ara; 25912: Co sca; 26376: Ch lan pul; 26393: Co sca; 27979: Co las; 28724: Co spl; 28788: Co spl; 29621: Co spl; 30218: Co sca; 58784: Co sca; 58886: Ch con; 59344: Co spl; 59644: Ch aca – Prévost 1311: Co ara; 1403: Ch con; 1414: Ch con; 1441: Co cla; 1822: Ch con; 1895: Ch con; 1987: Ch con; 2030: Co ara; 2118: Co ert; 2245: Ch cur; 2381: Co sca; 2722: Ch con; 2725: Co ert; 2728: Co spl; 3211: Co ert; 3836: Co spl; 4867: Co ert – Proctor Cooper 212a: Co woo – Proctor 16568: Co vil; 17261: Co sca; 21113: Co spc; 26163: Co sca; 27125: Co pul; 27389: Co pul; 32259: Co pul; 44752: Co mac – Pruski 3418: Co ara; 3487: Co ert – Puga 10: Co vil – Puig 1091: Co pic – Pulido 107: Co ant – Pulle 163: Co sca; 271: Co spl; 470: Co ara – Purpus 7831: Co sca; 7832: Co com; 8036: Co sca; 10751: Co sca; 16105: Co pic – Pursell 8154: Co ara; 9178: Co sca – Putcher 220: Co sca.

Quarles van Ufford 102: Co com – Quentin 831: Co spc; 832: Co spc – Quesada Quesada 120: Co cur; 487: Co pul; 627: Co pli – Questel 1469: Co spc – Quevedo Sopepí 831: Co sca; 2473: Ch aca – Quinet 1697: Co sca; 2460: Co spl – Quintana, C 969: Co sca – Quintana, D de la 427: Di arg – Quintero 1: Co vil – Quipuscoa Silvestre 235: Co vag – Quisbert 1360: Co acr – Quizhpe 1333: Co sca.

Raaijmakers 1: Co las; 54: Co ste – Raghoenandan UVS 17883: Ch lan pul – Ramamoorthy 1790: Co mol – Rambaran 60: Co ara; 61: Di rur – Ramia 571: Co sca – Ramírez Arango 284: Di str; 728: Co ant; 870: Co ant; 997: Co las; 1006: Co ant; 1763: Co las; 7155: Co spl; 7777: Co sca; 7811: Co mac; 7877: Co las; 8365: Co com; 9122: Co ara; 9371: Co ara – Ramírez Morillo 400: Co spl – Ramírez, BR 18552: Co kun – Ramírez, L R 3295: Co pic – Ramírez, N 93: Co com – Ramírez, V 176: Co pul – Ramírez-G 540: Co cha; 615: Co cha; 616: Co cha –Ramos, CE 279: Co spl – Ramos, J 682: Ch aca – Ramos C 183: Co pul –Ramos-Pérez 1395: Co plo; 1896: Co ant; 2418: Di str; 2419: Co asp; 2483: Co sca; 3052: Co ant – Ramos S 39: Co pul – Rankin de Merona 8: Co spl – Ratter 2630: Ch sub; 4567: Co pul; 5667: Co sca; 5668: Co ara; 6410: Ch sub – Raven 21666: Co kun; 21685: Co osa; 21792: Co wil; 21815: Co kun – Raynal 15508: Co mac; 18504: Co cla; 18508: Co ert; 18509: Co ert; 18624: Ch con; 19924: Co ara – Read 1011: Co pul; 1048: Co pul; 1060: Co spc; 8065: Co sca – Redden 2254: Co ert; 3497: Co sca; 4887: Co mac; 5179: Co sca; 5341: Co sca; 5460: Co ert; 5757: Co ert – Reed 41: Co pul – Regnell III 1208: Ch aca; III 1212: Co spl; III 1213: Co ara – Reijenga 519: Co ara; 651: Co sca; 670: Co ara; 935: Co ara – Reina 1181: Co ant; 1301: Co pul – Reitz 3798: Co ara; 4406: Co spl; 6730: Co ara; 7006: Co ara – Rentería Arriaga 36: Co kun; 52: Co cor; 56: Di str; 470: Co ant; 699: Co kun; 740: Di str; 816: Di str; 1490: Co ant; 1608: Di str; 1645: Di str; 1646: Co lim; 1843: Co ant; 1844: Di str; 1908: Co ant; 2112: Di str; 2123: Co ant; 2295: Di str; 2802: Co las; 3031: Co las; 3702: Di str; 3929: Co plo; 4201: Co sca; 4235: Co pul; 4343: Co sca; 4612: Co sca; 4645: Co pul; 4659: Co lim; 4771: Co lim; 4772: Co mac; 4781: Co sca; 4835: Co sca; 4976: Co las; 5048: Co lim 10648: Co mac; 10772: Co leu; 10778: Co sca; 10808: Co leu; 10816: Co sca; 10866: Di str – Restrepo 230: Co las; 231: Co spl; 761: Co spl; 771: Co vil – Revilla Cardenas 143: Co sin; 259: Co ara; 433: Di app; 770: Co ara; 1094: Co sca; 1265: Co erc; 1611: Co ara; 1804: Co sca; 1868: Co sin; 1886: Co sca; 1888: Di app; 1941: Co ara; 2192: Co ara; 2283: Co ara; 3023: Co las; 3739: Co ara – Rey-C 438: Co vil; 468: Co vil – Reyes, D 256: Co sca; 260B: Co ama; 260C: Co sca; 328: Co asp; 408: Co sca; 4154: Co ama; 901: Co sca – Reyes Escobar 29: Co pic – Reyes-García 2328: Co mac; 2540: Co pul; 6258b: Co pic – Reynel Rodríguez 5036: Co sca – Reynoso Santos 102: Co pic – Richard, H 418: Co cla; 683: Co las – Riera 1787: Ch cur – Rimachi Y 717: Co ara; 1185: Co erc; 1206: Di app; 1762: Co sca; 2764: Co ara; 3198: Co ara; 3290: Co sca; 4040: Co las; 4221: Co ara; 4325: Co spl; 4440: Co mac; 5277: Co erc; 5420: Co sin; 5679: Co erc; 5814A: Co sca; 5869: Co sca; 5941: Co erc; 6409: Co erc; 7767: Co ara; 8442: Co erc; 8460: Co lae; 9843: Co erc; 10355: Co mac; 10360: Di app; 10460: Co ara; 10995: Co acr; 11001: Co las; 11044: Di app; 11373: Co sin; 11493: Co sin; 11818: Co sca – Rimbach 40: Co pul; 291: Co lim – Rincón, R 63: Di str – Rincón Gutiérrez 592: Co alt – Rios, M 60: Co pul; 62: Co pul; 91: Co kun; 189: Co pul; 492: Co sca – Ríos, MA 2601: Co las; 5605: Co lon – Ritter 1425: Co ara; 3152: Co ara – Rivera C 412: Co las – Rivera D 64: Co kun – Rivera Díaz 314: Co lim; 537: Co spl; 1223: Co spl; 3170: Co com – Rivera E, C 12: Co vil – Rivera Elizondo 816: Co kun; 1508: Co mal – Rivera-P 315: Co kun; 316: Co asp – Rivera Reyes 1610: Co alt – Rivero, E 63: Co sca – Rivero, R 2617A: Co pul – Rivet 849: Co lim – Riviere 310: Co kun – Robbins 5623: Co lim; 5624: Co pul – Robertson 25: Co sin; 26: Co sin – Robinson 60: Co lae; 255: Co sca; 376: Co pul – Robles González 1426: Co woo; 1723: Co kun; 1843: Co pul; 1873: Co pul; 1874: Co bra; 1880: Co pul; 2210: Co pul; 2225: Co bra; 2601: Co kun; 2792: Co pul – Robleto Téllez 328: Co pul – Rocha 17: Di rur; 19: Di str; 21: Co vil – Rodrigues da Silva 192: Co spl – Rodrigues 1628: Co ara; 2131: Co mac; 2996: Co pra; 4318: Co ara; 5115: Co ara; 5587: Co ara – Rodriguez, C 79: Co lon; 82: Di app; 83: Co sca; 86: Co ara; 87: Co kru; 88: Co kru; 89: Co erc; 90: Co kru; 91: Co sca; 98: Co fis; 766: Co kru – Rodriguez, L 3716: Co spc; 4338: Co spc – Rodríguez, M 16: Co las; 28: Co sca; 46: Co las; 3027: Co cha – Rodríguez, N 722: Co cup; 1117: Co sca; 1158: Co sca – Rodríguez, WD 8133: Co sca – Rodriguez 228: Co com – Rodriguez B 4: Co las – Rodríguez C 336: Co ara – Rodríguez Delcid 105: Co pic – Rodríguez González 10311: Co wil; 11444: Co cur; 2257: Co woo; 2433: Co kun; 3360: Co pul; 3558: Co las; 4684: Co mon; 4891: Di str; 4967: Co vil; 5062: Co mon; 5358: Co kun; 6296: Co cur; 7007: Co ric; 760: Co ste – Rodríguez Olmos 572: Ch aca – Rodríguez R 224: Co ert; 228: Co asp; 267: Co sca – Rodríguez Ramírez 307: Co wil – Rodríguez-Suárez 2: Co cor – Rodríguez Vargas 156: Di arg – Roe 905: Co pul; 928: Co mol – Rohweder 1232: Co pic; 1233: Co pic – Rojas 2011: Co ara; 10148: Co ara; 10883: Co ara – Rojas Gonzales 51: Co ert; 146: Co ama; 1403: Co sca; 1988: Co lae; 2567: Di arg; 2726: Co jur; 2763: Co lae; 2819: Co lae; 2861: Di arg; 3411: Co lae; 3487: Co lae; 3574: Co rub; 4786: Co com – Rojas Leiva 245: Co pul – Rojas Mora 64: Co pul; 132: Co pul – Rojas-Zamora 136: Co ant – Roldán 3468: Co ant; 3658: Co spl; 3867: Co sca; 3899: Co mal; 3930: Co ant – Rombouts 74: Co spl; 145: Co ara; 452: Co ara; 741: Ch lan pul; 929: Co ara – Romero Castañeda 336: Co com; 2702: Di str; 3079: Co ant; 3407: Co spl; 3901: Co spl; 4019: Co sca; 4164: Co erp; 4463: Co vil; 5432: Co kun; 6237: Co sca; 6429: Co pul; 11164: Di str – Romoleroux 2726: Co kru – Ronquillo 889: Co sca – Roque 1372: Co sca – Rosa 3599: Co spl; 4036: Ch lan pul; 4537: Ch lan pul – Rosales 854: Co pic; 983: Co pic; 999: Co pic; 1036: Co pic; 1047: Co pic; 1216: Co pic; 1964: Co pic; 2254: Co pic; 2699: Co pic; 2700: Co pic – Rosales Castellanos 1073: Co sca; 1578: Co mac; 1609: Co ara – Rosário 1771: Co sca; 1773: Co ara; 1905: Co ara; 1908: Co sca; 2131: Co sca – Rosas R 1411: Co sca – Rose Innes 274: Co pul – Rose 4335: Co sca – Rosero 127: Co sca; 257: Co sca – Ross 12-58: Co pic – Rowlee 45: Co pul; 325A: Co mac; 401: Co vil; 675: Co bra – Rubiano 76: Co cor – Rubio 651: Co pul ; 1012: Co pul; 1292: Co kun; 1345: Co sca; 2007: Co gib – Rudas Lleras 1209: Co erc; 1633: Co sca; 2108: Co ara; 2184: Co sca; 2502: Co ara; 3027: Di str; 3242: Co sca; 3450: Ch fra; 3627: Di rur; 7319: Co spl – Rueda 449: Co sca; 1495: Co pul; 1527: Co woo; 1602: Co bra; 1603: Co kun; 1855: Co pul; 2072: Co pul; 2075: Co pul; 2691: Co pul; 2928: Co mal; 3130: Co woo; 3214: Co mal; 3286: Co pic; 3608: Co lim; 3813: Di str; 4191: Co woo; 4653: Co pul; 4738: Co kun; 4795: Co woo; 4991: Co pul; 5174: Co sca; 5198: Co pul; 5258: Co sca; 5353: Co pic; 5415: Co sca; 5417: Co sca; 5456: Co pic; 5563: Co lim; 5648: Co lim; 5709: Co pul; 5722: Co mal; 5775: Co woo; 5846: Co pic; 5872: Co pul; 5935: Co kun; 5961: Co pul; 6078: Co pul; 6230: Co pul; 6249: Co woo; 6480: Co pul; 6489: Co pul; 6585: Co pic; 6603: Co lim; 6617: Di str; 6653: Co pic; 6684: Co pul; 6714: Co pul; 6734: Co lim; 6841: Di str; 6942: Co pic; 7234: Co pic; 7252: Co pul; 7281: Co pic; 7379: Co pul; 7628: Co pic; 7676: Co pul; 7693: Co pul; 7695: Co pic; 7713: Co lim; 7714: Co pic; 8172: Co pul; 8199: Co pul; 8618: Co pul; 8741: Co mal; 8812: Co kun; 8932: Co woo; 9198: Co pic; 9349: Co mal; 9395: Co mal; 9707: Co mal; 9783: Co pul; 10084: Co pul; 10169: Co lim; 10205: Co pul; 10344: Co pul; 10500: Co lim; 11213: Co pul; 12168: Co pul; 12318: Co pul; 13226: Co pic; 14186: Co sca; 14204: Co kun; 14406: Co pul; 15009: Co sca; 15041: Co lim; 15115: Co kun; 15118: Co lim; 15250: Co sca; 15370: Co lim; 15493: Co pul; 16173: Co sca; 16530: Co pul; 16538: Co mac; 16650: Co pul; 16879: Co sca; 17034: Co pul; 17084: Co pul; 17237: Co pul; 18095: Co pic; 18739: Co pic; 19401: Co mac; 19402: Co lim – Ruiz, C 29: Co kun – Ruiz, D 173: Co pul – Ruiz, DP 15: Co ant – Ruíz, J 1269: Co las – Ruiz, J. 1269 p.p.: Co las; 5869: Co ara – Ruiz, R 216: Co pul; 217: Co pul – Ruiz Boyer 887: Co mon – Ruiz López B 23: Co lae; B 24: Co mac; B 25: Di arg – Ruíz-Terán 24: Co mac – Ruíz Zapata 3819: Co pul; 4692: Co vil – Rusby 1295: Co vag; 1399: Ch aca; 2225: Co gua; 2229: Ch lan lan – Russell 551: Co cur; 768: Co cur; 865: Co pul – Rutkis 277: Co ara; 389: Di str – Rutten 136: Co pul.

Saavedra 2A: Co ama – Sabatier 4956: Co cla; 5186: Co cla; 5713: Co sca – Saborío 98: Co com – Sáenz 41: Co woo – Sagástegui 5729: Co lae – Sagot 564: Co spl; 565: Co ara; 565a: Co ara; 566: Co sca – Salas TT 51: Co sca – Salaun 42: Ch fus – Salazar 137: Co ant – Salazar Pinilla 14: Co vil – Saldias Paz 1035: Co ara; 1364: Co sca; 2387: Co sca; 2766: Co spl; 5161: Co ara – Sales de Melo 58: Co spl – Salick 8161: Co kun; 7869b: Co kun; 8049A: Co pul; 8049B: Co kun – Salinas, H 6566: Co jur – Salinas, NR 11: Co cha; 12: Co cha; 13: Co ara; 14: Co spl; 18: Co spl; 33: Co ant; 203: Di str; 278: Co cor; 341: Co erp; 500: Co plo; 564: Co kun; 576: Co ant; 609: Co spl; 612: Co erc; 667: Di cry; 637: Di app; 678: Co kru; 710: Co spl – Sampaio 5847: Co sca; 7927: Co ara – Samper 1: Co ara – Samuels 33: Co sca; 186: Co ara; 355p.p.: Co ara & Co sca; 402: Co ara – Sánchez, C 1059: Co lae – Sanchez, J 18: Co sca – Sanchez, J 18: Co sca – Sánchez, M 287: Co sca – Sánchez, OD 18: Co com – Sánchez, R MCF 398: Co spl; 1051: Di cry; 1389: Co kun – Sánchez-Gómez 209: Co las – Sánchez M 499: Co sca – Sánchez S 31: Co ant; 575: Di str; 691A: Co lim; 931: Di str; 980: Co ant; 1051: Di cry; 1502: Co ant; 1951: Co las; 5658: Co ant – Sánchez-Vega 8057: Co sca – Sandino 96: Co pul; 107: Co pul; 136: Co pic; 137: Co pul; 182: Co pul; 234: Co sca; 235: Co pul; 1287: Co pul; 1620: Di str; 1678: Co lim; 2163: Co woo; 2381: Co lim; 2796: Co lim; 2820: Co sca; 3367: Co pul; 4052: Co woo – Sandoval, EA 297: Co com; 1223: Co com – Sandoval, M 266: Co com – Sandwith 238: Co sca; 447: Co ara – Sanín 4314: Co ant – Sanint 849: Co leu; 1473: Co sca – Sanoja 277: Co sca – Santa 63: Co vil; 186: Co las – Santamaría A 440: Co lim; 827: Co pul; 1295: Co kun; 1577: Co mon; 2065: Co vil; 3506: Co wil; 4397: Co wil; 5127: Co mal; 6178: Co kun; 6530: Co cur; 7804: Co com; 8864,1: Co mon – Sant’Ana 268: Co sca; 830: Co spl – Santos, AA 2367: Ch sub; 3350: Co sca – Santos, JU 1: Co ara; 43: Co ara; 205: Co ara; 707: Ch sub – Santos, NR 50981: Co ara – Santos, RR R 1474: Co spl – Santos, TS 828: Ch sub; 893: Ch cus; 1157: Ch cus; 1301: Ch sub; 1548: Co ara; 1654: Ch cus; 1661: Ch sub; 1974: Co ara; 2380: Co ara; 3004: Co spl; 3005: Co sca; 3237: Co sca; 3247: Co ara – Saravia T 2423: Co las – Sarmiento, F 859: Co com; 872: Co spl; 1095: Co spl – Sarmiento, JC 276: Co sca – Sarthou 812: Co spl; 865: Co cla – Sasaki 1848: Co spl – Sastre 10: Co spl; 45: Co ara; 49: Co sca; 71: Co sca; 77: Co las; 84: Co ara; 103: Co ara; 228: Co ara; 360: Co spc; 463: Co kun; 464: Co spl; 465: Co spl; 466: Co asp; 535: Co sca; 548: Co sca; 579: Co kru; 580: Co kru; 581: Co erc; 603: Co sca; 649: Co sca; 662: Co kru; 768: Di str; 913: Co spl; 968: Co sca; 981: Co spl; 1037: Co spl; 1110: Co spl; 1400: Co sca; 1428a: Co sca; 1428b: Co ara; 1446: Co ara; 1466: Ch con; 1475: Co ert; 1584: Co spl; 1659: Co spl; 1752: Co spl; 1794: Co sca; 2183: Co spl; 2483: Co spl; 2485: Co pul; 3233: Co spl; 3562: Co sca; 3958: Ch con; 3977: Co ara; 4040: Co sca; 4095: Co cla; 4099: Co cla; 4364: Co ara; 4391: Co sca; 4558: Co spl; 4559: Co sca; 4583: Co cla; 4607: Ch con; 4627: Co sca; 4648: Ch con; 4738: Co spl; 4927: Co sca; 5026: Co sca; 5753: Co ert; 6302: Co ert; 6484: Co cla; 9176: Co sca – Saunders 278: Co pul; 351: Co sca; 555: Co sca; 642: Co pul; 695: Co pul; 1153: Co lim – Sauvain 86: Co sca – Schäfer 9207: Co ert; 9222: Co ert – Scharf 28: Ch con – Schenck 684: Co sca –Schessl 44 1-4: Co ara – Schiefer 705: Co spl – Schimpff 525: Co lim; 557: Co pul; 983: Co vil; 1018: Co pul – Schipp 262p.p.: Co pic & Co pul; 416: Co mac; 507: Co kun – Schmalzel 10: Co pul; 144: Di str; 172: Di str; 633: Co mac; 637: Co vil; 816: Co pul; 839: Co kun; 853: Co vil; 872: Co kun; 930: Di str; 1123: Co kun; 1217: Co pul; 1614: Co wil – Schmutz 4907: Co mal – Schnee 1002: Co vil; 1039: Co spl; 1134: Co com – Schnell, CE 986: Co sca; 1009: Co mal – Schnell, R 11406: Co sca; 11786: Co ara – Schott 1: Co lim – Schubert 2208: Co ara – Schultes 2005: Co sca; 3540: Co cha; 3631: Di str; 3891: Co las; 5566: Co cha; 12565: Co ara; 13263: Co mac; 13569: Co ara; 13681: Co sca; 13910: Co spl; 14940A: Co ara; 16240: Co sca; 18962: Di str; 19092: Co asp; 24095: Co sca; – Schulz 21: Co lae; LBB 10563: Co ara; LBB 10627: Co ara – Schunke Vigo 1302: Co lae; 1304: Di rur; 1791: Co sca; 2241: Co sca; 2271: Co jur; 2894: Co com; 2954: Co gua; 3412: Co jur; 3418: Co jur; 3765: Co las; 3841: Co sca; 4311: Co las; 4382: Co sin; 4512: Di rur; 4544: Co web; 5076: Co web; 5527: Co las; 5643: Co spl; 5857: Co sca; 6124: Di rur; 6336: Co las; 6351: Co jur; 6542: Co sin; 6685: Co web; 6716: Di rur; 7124: Co sca; 7348: Co sin; 7396: Co las; 7925: Co jur; 8113: Co las; 8191: Co jur; 8291: Co las; 8378: Co kru; 8505: Co sin; 8571: Co web; 8672: Co las; 9775: Co sca; 9783: Mo uni; 9894: Co jur; 9979: Co las 10195: Co ant; 10309: Co las; 10351: Co web; 10397A: Co jur; 10577: Co web; 10798: Co sca; 10827: Co las; 11058: Di rur; 11598: Co las; 11983: Co jur; 12007: Co las; 12218: Co sin; 12259: Co jur; 12370: Co jur; 12565: Co las; 13588: Co sca; 14066: Co sca; 14118: Co jur; 14119: Co sin; 14121: Co sca; 14136: Co las; 14554: Co lae; 14752: Co vaz; 14766: Co ara; 14798: Co ara; 14820: Co acr; 14852: Co ara; 14855: Co sca; 14947: Co vaz; 15285: Di rur; 15345: Co vaz; 15482: Co sca; 15567: Co dal; 15998: Co web; 16000: Co sin; 16076: Co sca; 16185: Co dal; 16200: Co jur; 16355: Co lae; 16596: Co lae; 16608: Co lae – Schunke 384: Co lim; 435: Co sca – Schwacke 7165: Co spl – Seibert 121: Co vil; 330: Co wil; 419: Co pul; 454: Co las; 508: Co las; 582: Di str; 584: Co pul; 593: Co vil – Seidel 1132: Co vag; 2606: Co ara; 2690: Co bec; 2775: Co sin; 3027: Co com; 3029: Co sca; 3158: Ch aca; 7525: Co gua; 7547: Co sca; 7553: Co bec; 7726: Co sin; 7737: Di arg; 7763: Co vag; 8400: Co sca; 9373: Co ara – Seifriz 256: Co sca – Sendulsky 591: Co spl; 592: Co ara – Sepúlveda, F 190: Co ant; 644: Di str – Sepúlveda, P 529: Co lae; 682: Co las – Serato 208: Co spl – Sermeño 130: Co com – Serres 243: Co spl – Seymour 2977: Co woo; 3774: Co pul; 5624: Co vil; 5909: Co pul – Shafer 7694: Co pul; 13709: Co pul – Shank 4191: Co pul – Shannon 137: Co pic – Shattuck 183: Co vil; 313: Di str; 472: Co pul; 691: Co pul; 890: Co vil – Shea Semple 77: Co pul – Sheviak 2567: Co erp – Short 137: Co pul – Sigle 513: Co gua – Silland 211: Co sca – Silva, AM 4629: Ch sub – Silva, AS 242: Co las – Silva, ASL 627: Co ara – Silva, DG 2: Co ara – Silva, IA 1660: Co ara – Silva, JA 414: Co spl; 447: Co mac – Silva, LAM 368: Ch cus; 1053: Ch cus; 1804: Ch cus; 5718: Co mon – Silva, MF 246: Ch fra; 584: Co sca; 949: Ch fra; 1386: Co spl; 1424: Co ert; 1824: Co ara – Silva, MFF 411: Ch lan lan – Silva, MG 488: Ch lan lan; 3086: Ch fus – Silva, NT 1815: Ch lan pul; 1816: Ch fus; 1946: Co ara; 2333: Ch fus; 2334: Ch lan pul; 57750: Ch sub; 60852: Co kru – Silveira, ALP 559: Co pra; 596: Co pra – Silveira, M 1015: Co ert; 1070: Co jur; 1178: Di rur; 1446: Ch lan lan; 1455: Ch lan lan; 1616: Ch lan lan; 1618: Co vaz – Silverstone-Sopkin 1402: Co mac; 3085: Co ant; 3470: Co ant; 3528: Co ant; 4091: Co kun; 460: Co cha; 5162: Co sca; 5498: Co ant; 5521: Co sca; 5744: Co pul; 5810: Co sca; 5811: Co ant; 6986: Co pul; 7147: Co ant; 7834: Co ant; 8424: Co kun; 8963: Co ant; 9307: Co pul; 10902: Co lae; 12233: Co leu; 12251: Co cor; – Simmonds 3: Co mac; 17: Co sca; 27: Co mac; 28: Co sca; 32: Co sca; 60: Co mac; 65: Co sca; 74: Co mac; 79: Co sca; 133: Co sca; TRIN 14311: Co ara – Simonis 129: Co ara; 177: Co ara; 190: Co ara – Simpson, DR 453: Co pul – Simpson, GG 81: Co spl – Sinaca Colín 1515: Co dir; 1564: Co dir – Sintenis 1740: Co mac – Skinner R 1562: Co mal; R 2256: Co ert; 2847: Co whi; R 2952: Co vin; R 2970: Co ric; R 2971: Co sca; R 2972: Co mon; R 2973: Co cur; 3026: Co whi; R 3030: Co bra; R 3032: Co cur; R 3035: Co kun; R 3065: Co pul; R 3066: Co pul; R 3067: Co pul; R 3069: Co mac; R 3072: Co kun; R 3073: Co con; R 3074: Co vil; R 3076: Co sca; R 3077: Co ama; R 3078: Co asp; R 3079: Co acr; R 3080: Co acr; R 3081: Co sca R 3086: Co kun; R 3088: Co nit; R 3089: Co pul; R 3090: Co wil; R 3092: Co wil; R 3094: Co pul; R 3116: Co sin; R 3117: Co sca; R 3118: Co erc; R 3119: Di app; R 3120: Co sca; R 3122: Co ara; R 3123: Co sin; R 3124: Co ara; R 3125: Co asp; R 3126: Co lon; R 3127: Co las; R 3128: Co spl; 3131: Co rub; 3135: Co kun; 3136: Co alo; 3139: Co com; 3140: Co sin; 3141: Co sin; 3149: Co sin; 3150: Co sin; 3199: Co leu; 3200: Co ant; 3201: Co kun; 3209: Co mac; 3210: Co kun; 3211: Co pli; 3213: Co bar; 3214: Co ara; 3219: Co cla; 3225: Co vil; 3231: Co pul; 3245: Co vil; 3250: Ch fus; 3260: Co acr; 3262: Di rur; 3264: Co vag; 3280: Co bec; 3297: Co sca; 3301: Ch cus; 3306: Co erp; 3307: Co erp; 3308: Co com; 3312: Co com; 3313: Co com; 3315: Co com; 3323: Co gib; 3342: Co kun; 3343: Co geo; 3344: Co gib; 3346: Co gib; 3347: Co geo; R 3352: Co vin; R 3353: Co vin; 3355: Co kun; 3356: Co sin; 3363: Co gla; 3364: Co gla; 3366: Co kun; 3374: Co woo; 3380: Co sin; 3387: Co com; 3391: Ch aca; 3395: Co sin; 3396: Co com; 3397: Co com; 3400: Co alt; 3409: Co com; 3410: Co sin; 3421: Co rub; 3422: Co rub; 3423: Co rub; 3424: Co gua; 3426: Co sin; 3427: Ch aca; 3428: Co sin; 3429: Co gua; 3438: Co sin; 3440: Co mac; 3442: Co mac; 3443: Co sep; 3444: Co sep; 3446: Co alt; 3462: Co cup; 3463: Co whi; 3470: Co dal; 3479: Co dal; 3482: Co dal; 3483: Co dal; 3489: Co gua; 3492: Co dal; 3494: Co com; 3495: Co com; 3497: Co alo; 3498: Co alo; 3500: Co com; 3502: Co com; 3503: Co alo; 3504: Co alo; 3505: Co com; 3516: Ch aca – Skog 5627: Ch con; 7392: Co ert – Skorupa 155: Co spl – Skutch, AF 1313: Co com; 2690: Co kun; 2775: Co pul – Smith, A H 195: Co mon; 1476: Co mon; P 2560: Co mal – Smith, AC 2869: Co ara; 3452: Co sca; 3539: Co mac – Smith jr, CE 3261: Co vil; 3291: Di str; 3410: Co kun – Smith, CL 17: Co kun; 27: Co pul – Smith, DA 1037: Co mon – Smith, DN 2005: Co jur; 2946: Co rub; 3721: Co lae; 3848: Co sca; 3943: Co jur; 5360: Co rub; 5423: Di arg; 6375: Di arg; 6512: Co mac; 6635: Co sca; 6666: Di arg 12908: Co sca; 12968: Co sin; 12973: Co sca; 13010: Di rur; 13016: Co ara; 13234: Co sca; 13236: Co sin; 13649: Co ara; 13774: Co bec – Smith, E 1311: Co com – Smith, GL 10391: Co sca – Smith, HH 328: Co sca; 2329: Co pul; 2330: Co vil; 2632: Co com; 2632a: Co com; 2658: Co sca; 2767: Co lim – Smith, JF 447: Co wil – Smith, LB 5724: Co spl; 15348: Co spl 4816: Ch sub – Smith, RF 1725: Co sca; 4173: Co mac – Smith, SF 120: Co sca; 1625: Co ara; 1651: Co sca – Sobel 2370: Co asp; 4845: Co sca – Soejarto 592: Co las; 950: Co lim; 2720A: Co pul; 2977: Co sca; 3182: Di str; 3213: Co ant; 9303: Di str – Soeprato 7G: Co ara; 113: Co ara; 209: Co ara; 275: Co ara – Sohmer 9433: Co pic – Solano Peralta 835: Co wil; 1254: Co mal; 1415: Co mal; 1498: Co kun; 1504: Co sca; 3180: Co wil – Solano V 2075: Co pul – Solano, PJ K 266: Ch cur; K 267: Co cla; K 277: Co ert; K 414: Ch con – Solheim 877: Co pul; 895: Co pul; 1367: Co alt; 1740: Co pic – Solís Rojas 197: Co pul; 202: Co mal; 223: Co lim; 227: Co lim; 232: Co pul; 571: Co woo – Solomon 3251: Co sca; 3363: Co sca; 3897: Co sca; 6197: Co acr; 6296: Co sca; 6556: Co vag; 6962: Co ara; 7341: Co gua; 7811: Co acr; 7812: Co sca; 7941: Co ara; 7978: Co pse; 8166: Co sca; 8801: Co vag; 8802: Co vag; 9148: Co vag 12030: Co vag; 12913: Co vag; 13958: Co web; 14041: Co sca; 14183: Di arg; 14579: Co sca; 14809: Co gua; 14826: Co vag; 16783: Co ara; 17056: Co sca; 17382: Co vag – Solos 91: Co com – Soto Solís 52: Co nit; 73: Co wil – Souza, MAD 674: Co ara – Souza, SAM 582: Co sca; 822: Ch lan pul; 858: Co spl; 1068: Ch aca; 1177: Ch aca; 1179: Ch fus; – Souza, VC 5659: Co spl; 5883: Co spl; 9077: Co spl; 9320: Co spl; 11116: Co spl – Soza 345: Co lim – Sparre 13120p.p.: Co asp & Co sca; 13145: Co sca; 13590: Co lim; 13862: Co gib; 14450: Co lim; 14452: Co geo; 14554: Co ara; 15515: Co mac; 17109: Co pul; 17213: Co osa; 17857: Co ara; 18211: Co kun; 18212: Co sca; 19073: Co geo – Spellman 1683: Co pul; 1684: Co mac; 1911: Co pul; 1917: Co pul; 1966: Co pic; 2001: Co pic – Sperling 5449: Co vag – Sperry 538: Co mal; 553: Co pul; 574: Co mal; 579: Co mal; 740: Co mal; 740A: Co kun; 856: Co sca; 870: Co pul – Splitgerber 116: Co ara; 124: Co ara; 317: Co sca – Spruce 102: Co ara; 439: Co spr – St. George Expedition 320: Co kun; 612: Co sca – Stahel 260: Co spl – Ståhl 1807: Co sca; 3128: Co asp; 6729: Co pul; 6782: Co geo; 6897: Co geo; 7228: Co lim; 7278: Co pul – Standley 7639: Co pul; 8300: Co pic; 19365: Co pul; 19409: Di str; 19625: Co mal; 19781: Co com; 23702: Co pul; 25710: Co pul; 25970: Co vil; 26642: Di str; 26842: Co pic; 27476: Co pul; 27892: Co vil; 28127: Di str; 28672: Co kun; 28692: Co vil; 29175: Co vil; 29324: Co vil; 29549: Di str; 30394: Co vil; 31408: Co pul; 33596: Co bar; 34565: Co mon; 36877: Co sca; 37426: Co pul; 39534: Co mon; 40853: Co pul; 41017: Co pul; 41068: Co mac; 42116: Co mon; 44078: Co mon; 44246: Co gla; 45930: Co sca; 46344: Co sca; 48858: Co vil; 48871: Co kun; 51656: Co mon; 52705: Co sca; 53294: Co sca; 53672: Co pul; 54708: Co pul; 56615: Co pul; 58283: Co com; 58296: Co pic; 60714: Co pic; 63606: Co com; 64556: Co com; 64571: Co com; 66845: Co com; 66956: Co com; 67103: Co com; 67993: Co com; 68029: Co com; 68413: Co com; 68745: Co com; 70278: Co pic; 70693: Co sca; 71690: Co pul; 72516: Co pul; 72769: Co pul; 72838: Co pul; 78519: Co com; 78529: Co pic; 87184: Co com; 87508: Co pic; 87952a: Co com; 88389: Co com; 89509: Co com; 89564: Co pic; 90115: Co pic; 90200: Co pul – Stannard 487: Di str; 767: Co sca – Staples 834: Co kun – Starr 49: Co pul – Starry 30: Co pul; 128: Di str; 211: Co vil – Stauning 349: Co pic – Stehlé, H 427: Co spc – Stehlé, M 4980: Co spc – Stehmann 4712: Co sca – Stein 1072: Co sca; 1338: Co pul; 1705: Co sca – Steinbach 5543: Co ara; 7386: Ch aca – Steiner 1067: Co mal – Stergios 2748: Co sca; 3017: Co spl; 3262: Co sca; 3560: Co sca; 3802: Co sca; 5024: Co sca; 5029: Co ara; 5041: Co sca; 5730: Co ara; 5865: Co mac; 5868: Co sca; 5967: Co sca; 6058: Co ara; 6068: Co ara; 6068A: Co sca; 6205: Co ara; 6894: Co vil; 6990: Co pul; 7656: Co ara; 8661: Co vil; 8739: Di str; 9406: Co ara; 10676: Co ara; 10676A: Co spl; 10991: Co sca; 11068: Co sca; 11155: Co mac; 11791: Co mac; 11803: Co spl; 12061: Co sca; 12719: Co ara; 13850: Di str; 14050: Co sca; 14326: Co ara; 15078: Co sca; 15236: Co ara; 15271: Co sca; 16127: Co mac; 19604: Co spl – Stern, P 328: Co erc – Stern, WL 357: Co mac; 449: Di str; 675: Co mac; 911: Co pul – Stevens, FL 690: Co pul; 1135: Co vil; 1224: Co mac; 1292: Co sca – Stevens, WD 1111: Co pul; 2414: Co pic; 3277: Co pic; 4817: Co kun; 4818: Co pul; 4958: Co pul; 5048: Co mal; 6364: Co kun; 7019: Co lim; 7079: Di str; 7082: Co pul; 7162: Co lim; 7808: Di str; 7893: Co woo; 8114: Di str; 8459: Co pul; 8671: Di str; 8684: Co pic; 8685: Co pul; 8817: Co lim; 9590: Co sca 10490: Co woo; 10638: Co vil; 11956: Co lim; 11959: Di str; 12117: Co pul; 12184: Co pul; 12488: Co pul; 12564: Co lim; 12590: Di str; 12657: Co kun; 12848: Co mac; 13004: Co pul; 13043: Co lim; 13051B: Co pul; 13095: Co pul; 13328: Co mal; 13575: Co pul; 13578: Co cur; 13737: Co kun; 14557: Co sca; 15133: Co sca; 16652: Co pul; 16738: Co pul; 16825: Co lim; 17590B: Co pic; 18664: Co pul; 18749: Co kun; 18781: Co pul; 19243: Co pul; 19285: Co lim; 19783: Co woo; 20001: Co woo; 20114: Co woo; 20682: Co woo; 20772: Co woo; 20873: Co woo; 20927: Co sca; 21090: Co sca; 21118: Co sca; 21237: Co pul; 23113: Co pul; 23889: Co sca; 23890: Co pli; 23891: Co bra; 23966: Co mal; 24061: Co pul; 24075: Co woo; 24081: Co kun; 24292: Co mal; 24626: Co kun; 25547: Co pul; 27088: Co pul; 28156: Co lim; 28208: Di str; 28272: Co pul; 28509: Co sca; 30544: Co pul; 31451: Co mac; 31650: Co pul; 33517: Di str; 35246: Co sca; 36890: Co mac; – Stevenson Diaz 455: Co mac; 496: Co mac; 2002: Co sca – Stevenson 1222: Co pul; 1243: Co wil – Steward 476: Co pra; P 20366: Co sca – Steyermark 17112: Co pul; 31799: Co pic; 33231: Co pic; 33250: Co com; 33426: Co com; 33859: Co pic; 35228: Co pic; 37117: Co com; 37680: Co kun; 38106: Co pul; 38906: Co sca; 39976: Co pul; 41844: Co sca; 41888: Co pul; 44870: Co pul; 45291: Co pul; 45948: Co pul; 46779: Co pic; 46780: Co com; 47650: Co pic; 48990: Co pul; 51033: Co pic; 51099: Co pic; 54664: Co zam; 56718p.p.: Co pul & Co spl; 61083: Co sca; 87163: Co sca; 87167: Ch con; 87706: Di rur; 87765: Co ara; 89517: Co spl; 90738: Co spl; 91805: Co com; 94487: Co pul; 95681: Co sca; 96074: Co mac; 96421: Co sca; 96491: Co mac; 97806: Co sca; 99616: Co pul; 99670: Co lim; 99970: Co mac 100189: Co ara; 101423: Di str; 101869: Co spl; 102089: Co sca; 102136: Di str; 102671: Co ara; 102672: Co spl; 105347: Co mac; 105408: Co sca; 106194: Co vil; 106998: Co sca; 107200: Co spl; 107676: Co com; 107752: Co mac; 111946: Co sca; 112890: Co sca; 113848: Co ara; 113881: Co mac; 113889: Co spl; 114437: Co sca; 114739: Co ara; 114745: Co sca; 114805: Co sca; 114826: Co sca; 114860: Co ara; 115072: Co ara; 118730: Co spl; 119062: Co jur; 119067: Co spl; 119142: Co mac; 119228: Co spl; 120705: Co mac; 122473: Co spl; 122736: Co lim; 123428: Co sca; 123429: Co lim; 124251: Co sca; 125740: Co spl; 127111: Co com; 127433: Co spl; 131548: Co mac; 131549: Co spl; 131746: Co ara – Stier 149: Di str; 164: Co vil – Stoffers 302: Co sca – Stone 2756: Co pul; 3964: Co sca – Stork C 110: Co mon; 2260: Co mal; 3111: Co pul – Strudwick 3052: Co sca; 3090: Co sca; 3492: Co ara – Suárez 357: Co las; 819: Co kun; 946: Co lim; 952: Co las; 984: Co sca; 1248: Co kru; 1307: Co sca; 1312: Co las – Suárez-B 84: Co kru – Sucre 8293: Co sca – Suescún 187: Co sca; 208: Di str – Sugden 146: Co vil; 301: Co vil; 594: Co kun – Sullivan 40: Co vil; 42: Co las; 356: Co cur; 472: Co las; 540: Co all; 615: Co pul; 649: Co kun; 740: Co vil; 1165: Di arg – Sutton 10: Co pul; 207: Co pul – Sytsma 958: Co sca; 1234: Co vil; 1244: Di str; 1274: Co mac; 1289: Co kun; 1550: Co all; 1555: Co sca; 1595: Co kun; 1596: Di str; 1659: Co all; 1970: Co las; 1971: Co las; 2040: Co sca; 2497: Co sca; 2549: Co sca; 2607: Co sca; 2623: Co sca; 2626: Co bra; 2656: Co sca; 3074: Co kun; 3129: Co sca; 3487: Co lim; 3514: Di str; 3842: Co las; 3853: Co las; 3913: Co mac; 3992: Co sca; 4035: Co las; 4077: Co cur; 4166: Co kun; 4276: Co sca; 4310: Co kun; 4329: Co las; 4373: Co bra; 4488: Co bra; 4509: Co sca; 4565: Co cur; 4610: Co alo; 4665: Co com; 4671: Co cur; 4786: Co pul; 4881: Co cur.

Tadri-Zocher 122: Co spl – Tamayo 1303: Co spl – Tameirão Neto 2591: Ch cus – Tate 79: Co sca; 441: Co sca; 522: Co ara; 523: Co sca – Taylor, EL E 1020: Co ara – Taylor, J 11662: Co mal; 11666: Co kun; 11685: Co pul; 17970: Co kun – Taylor, N 31: Co sca – Teixeira 389: Co ert; 414: Co sca; 1399: Ch lan lan – Téllez 76: Co com – Téllez Valdés 5654: Co pul – Temponi 8: Co sca – Terán Aguilar 1020: Co sca; 1886: Co sin; 1983: Co sca; 2499: Co bec; 4271: Co sca – Terry 1366: Co wil; 1532: Co pul – Ter Steege 551: Co cla –Tessmann 3292: Co sca; 3324: Co mac; 3751: Di rur; 4262: Co ert; 4813: Co erp; 4903: Co ama – Teunissen LBB 13193: Co ara; LBB 15328: Di rur – Thomas, E 766: Co sca; 943: Co bec – Thomas, J 1460: Di str; 1541: Co spl – Thomas, MB 605: Co spl – Thomas, WW 3099: Co sca; 3309: Co sca; 4384: Ch aca; 4760: Ch lan lan; 9329: Ch cus; 10552: Co spl; 10858: Ch sub; 11304: Co spl; 11937: Ch sub; 13401: Ch sub; 13510: Co sca; 15691: Co spl – Thompson 783: Co cha – Thoms 15: Co vil – Tillett 45336: Co spl – Timaná 1300: Co erp; 1357: Di app; 1425: Co erp; 1429: Co sca; 1665: Co erp; 1678: Co sca; 1951: Co sca; 2146: Co sca; 2147: Co sca; 2162: Co sca; 2176: Co sca; 2218: Co sca; 2286: Co sca; 2358: Co sca – Tinjacá 9A: Co sca; 17: Di str – Tipaz 572: Di str – Tirado 36: Co sca; 870: Co pul; 1954: Co kun – Tiwari 352: Co sca –Tjon-Lim-Sang 204: Co cla; 219: Co cla – Toasa 8671: Co lon; 8828: Co sca – Tobar 57: Co woo – Tobón 82: Co sca – Toledo, R 7: Co pul – Toledo G 23: Co ara; 145: Co ara; 567: Di rur; 923: Co sca –Ton 3568: Co pic; 5944: Co pul; 6071: Co pul – Tonduz 282: Co com; 1301: Co bar; 6659: Co lim; 8755: Co pul; 9422: Di str; 11386: Co lim; 12554: Co cur; 12926: Co lim; 12926A: Co kun; 14546: Co pul – Toriola- Marbot 216: Co ara; 229: Co ara; 259: Co ara; 1277: Co ert; 1278: Co ert – Torke 577: Co ara – Toro 300: Co spl – Torres, AM 1046: Co acr – Torres, LM 142: Co ara – Torres, M 1027: Co las Torres Colín 7807: Co pul – Torres R 2996: Co sca –– Tostain 210: Co spl; 318: Co spl; 1154: Co sca; 1340: Co sca; 1456: Co spl; 1680: Co sca; 1696: Co cla; 1823: Ch con; 2021: Co cla; 2656: Co ert; 2755: Co ert; 2768: Ch con; 2970: Co spl – Toval 336: Co pul – Townsend, WR 2: Co sca – Treacy 169: Co lon; 565: Co lon – Tredwell 104: Co sca; 127: Co sin – Tresling 32: Co ara – Tressens 5582: Co ara – Triana 652: Co cha; 653: Di str; 654: Co kun; 656: Co sca; 668: Co sca; 1638.2: Di str – Troya 26: Co pul – Trujillo, B 1798: Co mac; 1906: Co spl; 2244: Co ara; 2387: Co sca; 2388: Co sca; 3411: Co com; 3698: Co mac; 3717: Co sca; 3718: Co sca; 4887: Co com; 6031: Co sp; 10610: Co ara; 10859: Co mac; 14054: Co lim; 14276: Co ara; 14306: Co ara; 14545: Co spl – Trujillo, E 799: Co sca – Trujillo, P 4272: Co las; 5050: Co vil; 5566: Co pul; 6227: Co las; 7006: Co las – Trujillo-C 392: Co mac; 903: Co cha; 1789: Co com; 1830: Co kru – Tsugaru B 2295: Co spl – Tuberquia 3042: Co pul; 3147: Co ant; 3149: Co pul – Tuinherbarium Botanische Tuinen Utrecht 68-197: Co ara; 68-250: Co mon – Tulleken 156: Co sca; 237: Co ara; 261: Co ara; 421: Co sca; 570: Co ara – Tunqui 28: Co sca; 281: Co asp; 430: Co ert; 654: Co asp; 677: Co ert; 683: Co asp; 726: Co asp; 749: Co ert; 801: Co ert; 932: Co ert – Tyson 900: Co kun; 1115: Co vil; 1656: Co pul; 1719: Di str; 3677: Di str; 4033: Co las; 4097: Co las; 4290: Co mac; 4323: Co las; 4414: Co vil; 4628: Co pul; 4831: Co pul; 5530: Co lim; 5553: Co mac; 6727: Co mac.

Ugent 66: Co mac; 76: Co pul – Ule 95br: Co ara; 5740: Co jur; 6180: Co las; 6188: Co erc; 6333: Mo uni; 6475: Co lae; 8097: Co ara; 9194: Co sca; 9195: Co acr; 9196: Co ara; 9197: Ch aca; 9198: Ch lan lan – Uliana 1342: Co pra – Ulitzka 8C: Co ara – Ulloa U 4: Co pul; 86: Co kun; 91: Co pul; 99: Co kun; 106: Co kun; 111: Co gib; 136: Co gib; 136A: Co kun; 173: Co pul – Underwood 902: Co pul – United Fruit Company 229: Co pul – Urbina 7: Co vil; 12: Co lim – Urbina H 12: Co pul; 55: Co vil – Uribe, W 119: Di str – Uribe P, C s.n: Co cha – Uribe Uribe 1495: Co ant; 1627: Co vil; 1920: Co las; 2189: Co erp; 3999: Di str – Urrego 2016: Co sca – Usteri 4c: Co ara – Utley 1053: Co pul; 1141: Co kun; 1199: Co mac; 2374: Co mon; 2775: Co wil; 2803: Co kun; 4795A: Co wil; 5467: Co woo – UVS (Suriname) 17883: Ch lan pul.

Valcarcel 653: Co lon – Valdespino Quintero 444: Co cur; 468: Co pul; 663: Co all – Valencia, J 274: Co spl – Valencia, R 323: Co las – Valenzuela Gamarra 162: Co sin; 418: Co sca; 2506: Co acr; 4228: Di app; 7882: Cocom; 8459: Co gua; 9788: Co sca; 10471: Co gua; 11972: Co jur; 12291: Co las; 16159: Co ama – Valera 7: Co sca – Valerio 11: Co pul – Valerio Rodriguez 1903: Co pic – Van Andel 93: Co sca; 627: Co sca; 629: Co ara; 861: Co ert; 1003: Ch con; 1503: Co ara; 1581: Co ert; 3406: Co luc; 4809: Co sca; 4854: Co ara; 5161: Co spl; 6162: Co sca – Van Beek 34: Co spl – Van Borssum Waalkes 10169: Co mal – Van der Eynden 902: Co sca – Van der Laan 22: Ch cus; 124: Ch cus – Vanderveen 572: Co pul – Van der Wal 17: Co las; 91: Ch fra – Van der Werff 4649: Co mac; 4716: Co sca; 4960: Co spl; 6690: Co cur; 6871: Co cur; 7096: Co wil; 7800: Co vil; 10323: Co asp; 18147: Co jur; 18237: Co jur; 22757: Di arg – Van Donselaar 2537: Co sca – Van Dulmen 74: Co sca – Van Hattum 7005: Ch cus – Van Rooden 3: Di str; 4: Co spl; 17: Co mac; 18: Co mac; 19: Co mac; 2: Co spl; 20: Co ara; 24: Co spl; 27: Co mac; 28: Co sca; 29: Co mac; 30: Co spl; 31: Co mac; 32: Co spl; 33: Co spl; 114: Co lim; 116: Co sca; 117: Co sca; 118: Co kun 574: Co lim; 575: Di str; 665: Co sca; 775: Co sca; 812: Co dir – Van Severén 86: Co pul – Vanni 3530: Co ara – Vareschi 1674: Co ant; 3142: Co pul – Vargas, VA 40: Co cha; 85: Co lon – Vargas, WG 6124: Co all; 9336: Di str; 10846: Di str; 10898: Co vil; 11005: Di str; 11010: Co pul; 11277: Co plo; 17866: Co ant; 18945: Co asp; 18950: Co sca; 21564: Co pul; 21597: Co leu; 22066: Co com – Vargas, X 20: Co ant – Vargas Caballero 550: Co sca; 2032: Co com; 5770: Co sca; 6068: Co sca; 6082: Co ara; 6085: Ch aca; 6105: Co ara; 6130: Ch aca; 6168: Co ara; 6237: Co ara – Vargas Calderón 3471: Co com; 3798:Co sin; 7702: Co acr; 12285: Co jur; 13490: Co acr; 14044: Di rur; 14662: Co vag; 15188: Co sca; 16084: Co acr; 16862: Co acr; 16940: Co sca; 17846: Co vag; 18759: Co lae; 20812: Co jur – Vargas-Figueroa 855: Co ant – Vargas López 546: Co asp; 3656: Co asp; 4640: Co cor; 4984: Co pul; 5734: Co pul; 5836: Co pul – Vargas N 321: Co pul – Vargas Ramírez, L 654: Co acr; 1248: Co ara; 1252: Co sca – Vargas Ramírez, ORA 63: Co pli; 1605: Co kun; 1758: Co lim – Vargas Salazar 1811: Co pul; 2704: Co sca; 2811: Co mal; 3194: Co pul; 3398: Co kun; 3748: Co pul; 4241: Co pul – Vasco Gutiérrez 94: Co sca – Vasconcellos Sobrinho RB 93956: Co sca – Vásquez Aspinwall 190: Co sca – Vásquez Martínez 328: Co erc; 357: Co ara; 358: Co sca; 537: Co sin; 729: Di app; 873: Di rur; 888: Co sca; 966: Co sin; 1171: Co sca; 1468: Co sin; 1615: Co jur; 1727: Co erp; 2039: Co sca; 2262: Co acr; 2263: Di rur; 2265: Co sca; 2363: Co vaz; 2891: Di app; 3531: Co erc; 3631: Co sca; 3845: Co sca; 3937: Co sca; 4338: Co ara; 4338A: Co sca; 4652: Co sin; 4806: Co sin; 4860: Co ara; 4871: Di rur; 5129: Co sin; 6150: Co las; 8350: Co erc; 8428: Co sca; 8521: Co sin; 8741: Co ara; 8937: Co sca; 9296: Co sca; 11239: Co sca; 11271: Co sin; 11508: Co ara; 12564: Di str; 12565: Co las; 12580: Co sca; 14077: Co erc; 15854: Co sca; 15930: Co sin; 15973: Co las; 18558: Co asp; 18649: Co sca; 18657: Co ert; 21475: Co sca; 21646: Co ama; 21756: Co ama; 21813: Co ama; 22198: Co sca; 22201: Co sca; 22452: Co asp; 22662: Co ama; 22667: Co ert; 26104: Co asp; 27509: Co sca; 27722: Co jur; 27782: Co rub; 27905: Co lae; 29787: Co las; 29832: Co las; 31337: Co sca; 31340: Co las; 32140: Co kru; 34917: Di rur; 37113: Co spl – Vásquez, AI 32: Co kun; 41: Co woo – Vaughan 602: Co cur; 614: Co gla – Vázquez, A 4658: Co pul – Vázquez B 48: Co pul; 195: Co pul – Vázquez Torres 503: Co sca; 607: Co pul – Velasco-Ramirez 25: Co ant; 26: Co ant – Velásquez 5758: Co com; 5759: Co ant – Velástegui 96: Co kun – Velayos 6345: Co cha – Vélez, G 312: Co spl – Vélez, J 7073: Co ant – Vélez-Puerta 5003: Di str; 5004: Co ant – Veloz E13428: Di str – Ventura V 212: Co pic; 5254: Co pic – Vera Sánchez 2463: Co com; 2465: Co com – Verboom 5089: Co pul – Vergara 34: Di str; 35: Co spl – Versteeg 496: Co sca – Vester 643: Co las; 716: Co spl; 717: Co las – Veth 14: Co sca; 64: Co ara; 223: Co ara; 287: Co cla – Veyret 4413: Co spl – Vicentini 544: Co ara – Vidal 5443: Co spl – Vidal-Hernández 24: Co woo – Viegas 5456: Co ara – Vieira, MGG 233: Co sca; 359: Co sca; 369: Co dal; 395: Co ert; 440: Ch aca; 466: Co sca; 544: Ch lan lan; 575: Ch lan lan; 855: Co las; 858: Ch lan lan; 1064: Ch lan lan – Vieira, RF 568: Co ara; 798: Co spl; 1605: Co spl; 1945: Co spl – Vieira, S 267: Co sca – Vilca 317: Co sca – Villacorta M 2940: Co com – Villanueva 1844: Co ant – Villarroel Segarra 16: Co sca; 352: Co ara – Villeda 244: Co sca – Villela 64: Co pul – Villiers 6057: Co spl – Vincelli 255: Co kun; 395: Co pul; 943: Co spl – Vogel 96: Co pul; 251: Co las – Vogl 775: Co com; 777: Co com; C 1343: Co com – Von Eggers 12375: Co pul; 1289: Co mac; 13497: Co pul; 14186: Co mac; 14857: Co geo; 15012: Co pul; 1740: Co mac; 5204: Co pul; 5607: Co sca – Von Luetzelburg 227: Co ara; 7241: Ch cus; 20149: Co spl; 21450: Co sca; 21451: Co sca; 22197p.p.: Co sca; 22198: Co sca; 22295: Co spl; 22651: Co spl; 22655: Co spl – Von Scherzer 9903: Co mon – Von Sneidern A1285: Co cha; 5207: Co pul; 5347: Co cha; 5805: Co sca – Von Tuerckheim II 76: Co pul; II 385: Co kun; 7686: Co pul; 8015: Co kun– Von Wedel 25: Di str; 28: Co lim; 138: Co woo; 219: Co kun; 475: Co mac; 491: Co lim; 670: Co mac; 740: Di str; 817: Co lim; 1507: Co kun; 1560: Di str; 1632: Co lim; 2000: Co woo; 2080: Co woo; 2093: Co kun; 2505: Di str; 2506: Co kun; 2507: Co lim; 2520: Co mac; 2586: Co pul; 2664: Di str; 2797: Co lim; 2899: Co woo; 2939: Co woo; 2959: Co lim – Vriesendorp 339: Co erc – Vrieze 4301: Co pul.

Wachenheim 145: Co cla; 164: Co sca; 178: Ch con – Wachter 29: Co vag; 78: Co sca; 119: Co gua – Wagner, M 128: Co sca – Wagner, RJ 877: Co mal; 884: Co mac; 1608: Co mac –Walker 4: Di str – Wall 6: Co sca – Wallnöfer 5827: Co pul; 13-17988: Co las; 14-41287: Co sca; 19- 181188: Co sca – Walter 802: Co spl; 946: Co spl; 3489: Co spl – Ward 5864: Co pic – Warming 502: Ch sub; 521: Co spl; 773: Co sca – Warner 108: Co las – Watson, AG 44: Co ara – Watson, S 299: Co sca – Weberbauer 1852: Co web; 1856: Co lae – Webster 9529: Co sca; 16514: Co las; 22243: Co woo; 27437: Co kun; 28313: Co kun – Weddell 2810: Ch sub; 2832: Ch sub – Welch 19852: Co mac – Wendland s.nn.: Co mal – Went 60: Co ara; 69: Co ara – Werdermann 2499: Co sca – Wessels Boer 621: Co cla; 733: Co sca; 822: Co ert; 876: Co ert; 975: Co ara; 1022: Co cla; 1147: Ch lan pul; 1576: Co cla; 1959: Co spl; 1960: Co mac; 1992: Co pul; 2014: Co spl; 2439: Co ant; 2440: Co ant; 2457: Co mac; 2462: Co lim –White, OE 647: Co sca; 1801: Co sca; 1808: Co sca – White, P 126: Co vil; 147: Di str – White, S 689: Co kun – Whitefoord 119: Co lim; 145: Co vil; 206: Di str; 303: Co lim; 510: Co gla; 806: Di str; 1098: Co pul; 1545: Co pul; 1836: Co pul; 3860: Co sca; 10334: Co pul; 1061048: Co sca – Whitson 177: Co mal; 178: Co kun; 204: Co mal; 475: Co sca; 491: Co pul – Widgren 1205: Co ara – Wied zum Neuwied 19: Co spl – Wilbert 174: Co sca; 28471: Co sca; 68328: Co sca – Wilbur 8590: Co lim; 10700: Co pul; 11006: Co wil; 15047: Co all; 15051: Co kun; 30135: Co mal; 30198: Co pul; 33452: Co mal; 34142: Co bra; 34367: Co pul; 34392: Co sca; 34400: Co bra; 34701: Co lim; 34706: Co kun; 34744: Co sca; 34784: Co mal; 34797: Co pul; 36902: Co sca; 37015: Co pul; 37210: Co pul; 37226: Co kun; 37242: Co pul; 37329: Co pul; 37364: Co sca; 37511: Co pul; 37512: Co pul; 37671: Co pul; 37817: Co pul; 37822: Co pul; 37863: Co pul; 38863: Co mal; 39695: Co kun; 39745: Co pul; 39961: Co pul; 40055: Co mal; 40137B: Co kun; 40423: Co pul; 63467: Co sca; 64462: Co pul; 66106: Co sca; 66135: Co pul; 66157: Co pul; 66301: Co mal; 66328: Co mal; 66809: Co pul; 70054: Co mal – Wiley 42: Co pul; 566: Co mac – Williams, DE 1094: Di arg – Williams, Ll 772: Co sca; 968: Co ara; 977: Co sca; 1272: Co sin; 1661: Co jur; 2497: Co ara; 3004: Co las; 3924: Co zin; 3985: Co zin; 4291: Di rur; 4570: Co sca; 6529: Co lae; 8613: Co pul; 8873: Co pul; 9176: Co pul; 9242: Co pul; 9333: Co pul; 11251: Co ara; 11799: Co sca; 12238: Co mac; 14035: Co spl – Williams, LO 18796: Co pul; 18935: Co ste; 23846: Co pul; 28906: Co mon; 29108: Co pul; 42061: Co pul – Williams, RS 431: Co lim; 689: Di str – Wilson 400: Co pul; 646: Co pul; 18082: Co pul; 18089: Co pul – Windisch 136: Co ara; 1456: Ch aca; 2112: Co ara – Wingfield 6407: Co sca – Witingthon 34: Co spl – Wolfe 319: Co pul; 12106: Di str; 12117: Co sca; 12122: Co sca; 12266: Co las – Wood, CW 296: Co spl – Wood, TM 1037: Co kru – Woodson 202: Co cur; 282: Co wil; 544: Co cur; 785: Co vil; 786: Co lim; 788: Co vil; 789: Co mac; 857: Co pul; 885: Co pli; 886: Co pul; 896: Co mac; 966: Co pul 1000: Co cur; 1169: Co wil; 1231: Co las; 1358: Co mac; 1678: Co vil; 1835: Co lim; 1926: Co lim; 1929: Co kun; 1951: Co pli – Woodworth 338: Di str; 698: Co kun – Woolston 1091: Co ara – Worthington 13552: Co woo – Wright 528: Co sca; 1514: Co pul – Wullschlaegel 517: Co sca; 571p.p.: Co ara & Co sca; 1812: Co spl – Wurdack 2165: Di rur.

Yale Dawson 14941: Co spl – Yánez, AP 263: Co pul; 289: Co pul; 855: Di str; 1543: Co kun; 1570: Co pul; 1594: Di str; 2427: Co sca – Yanez, M 37: Co spl – Young, HJ 210: Co sca; 278: Di app – Young, KR 87: Co sca; 743: Co sin; 781: Co sca; 946: Co jur – Yuncker 3566: Di str; 4785: Co sca; 6120: Co pic; 8277p.p.: Co pul & Co sca; 8512: Co sca – Yusti-M 45: Co ant. Zak 1594: Di str; 4946: Co sca; 5278: Co ama; 5291: Co ama; 5512: Co pul; 5589: Co pul; 5609: Di str; 5641: Co pul – Zambrano B 13: Co pul; 15: Co pul – Zamora Villalobos 3896: Co bra; 3905: Co kun; 3908: Co bra – Zanoni 11990: Co sca; 12001: Co spc; 15956: Co spc; 16425: Co spc; 17631: Co sca; 17792: Co sca; 21077: Co spc; 21183: Co sca; 21284: Co spc; 23111: Co spc; 23427: Co spc; 23455: Co spc; 29484: Co spc; 29490: Co spc; 30551: Co spc; 30585: Co sca; 31576: Co spc; 34134: Co sca; 34144: Co sca; 38863: Co spc; 43051: Co spc – Zapata C 419: Co mac; 1831: Co pul – Zappi 896: Ch fus; 1174: Co sca – Zardini 9293: Co ara; 16495: Co ara; 17089: Co ara; 18600: Co ara; 20562: Co ara; 23149: Co ara; 24807: Co ara; 26530: Co ara; 28510: Co ara; 30220: Co ara; 30397: Co ara; 30745: Co ara; 32181: Co ara; 32841: Co ara; 33154: Co ara; 33177: Co ara; 33431: Co ara; 33499: Co ara – Zarucchi 1051: Co kru; 1307: Co spl; 2677: Ch aca; 2765: Ch lan lan; 3266: Co sca; 5059: Co las; 6658: Co ant – Zaruma 37: Co sca – Zent, EL 1996: Co spl; 2000: Co mac; 2349: Co vil – Zent, S 686-52: Co mac – Zenteno 628: Co sca – Zenteno-R 9995: Ch aca; 10064: Ch aca – Zomer 116: Co pul – Zuluaga, A 554: Co leu – Zuluaga R 79A: Co sca; 91: Di str; 224: Co pul; 861: Co pul; 1097: Co lim; 1703: Co mac – Zumárraga 107: Co sca – Zuniga, W 624: Co sep – Zúñiga Villegas 174: Co mon; 196: Co pul – Zurita 67: Co sca; 356: Co vag.

## INDEX

Accepted names are in roman type. New names are in **bold** type. Synonyms are in *italics*. Species that are insufficiently known (insuff) or nomina dubia (nom.dub.) and nomina nuda (nom. nud.) are indicated as such. References to pages are given in square brackets.

Abarema jupunba

*Alpinia comosa* Jacq.

*Alpinia spicata* Jacq.

*Alpinia spiralis* Jacq.

*Amomum petiolatum* Lam.

Brosimum

Calophyllum

Chamaecostus C.D.Specht & D.W.Stev.

Chamaecostus acaulis (Sp.Moore) T.André & C.D.Specht

Chamaecostus congestiflorus (Rich. ex Gagnep.) C.D.Specht & D.W.Stev.

Chamaecostus curcumoides (Maas) C.D.Specht & D.W.Stev.

Chamaecostus cuspidatus (Nees & Mart.) C.D.Specht & D.W.Stev.

Chamaecostus fragilis (Maas) C.D.Specht & D.W.Stev.

Chamaecostus fusiformis (Maas) C.D.Specht & D.W.Stev.

Chamaecostus lanceolatus (Petersen) C.D.Specht & D.W.Stev. subsp. lanceolatus

Chamaecostus lanceolatus (Petersen) C.D.Specht & D.W.Stev. subsp. pulchriflorus (Ducke)

C.D.Specht & D.W.Stev.

Chamaecostus manausensis Maas & H.Maas

Chamaecostus subsessilis (Nees & Mart.) C.D.Specht & D.W.Stev.

Chamaedorea sp.

Costaceae

Costoideae

*Costus acaulis* Sp.Moore

Costus acreanus (Loes.) Maas

Costus alfredoi Maas & H.Maas

Costus allenii Maas

Costus alleniopsis Maas & D.Skinner

Costus alticolus Maas & H.Maas

Costus amazonicus (Loes.) J.F.Macbr.

*Costus amazonicus* (Loes.) J.F.Macbr. subsp. *amazonicus*

*Costus amazonicus* (Loes.) J.F.Macbr. subsp. *krukovii* Maas

*Costus anachiri* Jacq.

Costus antioquiensis Maas & H.Maas

*Costus arabicus* auct. non L.: Aubl. (1775) 2.

Costus arabicus L.

*Costus argenteus* auct. non Ruiz & Pav.: Gagnep. (1902)

*Costus argenteus* Ruiz & Pav.

*Costus arrabidae* Steud. (= *C. arabicus* Vellozo).

Costus asplundii (Maas) Maas

Costus asteranthus Maas & H.Maas

Costus atlanticus E.Pessoa & M.Alves

*Costus bakeri* K.Schum.

Costus barbatus Suess.

Costus beckii Maas & H.Maas

*Costus bracteatus* Gleason, non Rowlee

Costus bracteatus Rowlee

*Costus brasiliensis* K.Schum.

Costus callosus Maas & H.Maas

*Costus cernuus* Sw. ex Roem. & Schult.

Costus chartaceus Maas

*Costus ciliatus* Miq.

Costus claviger Benoist

Costus cochabambae Maas & H.Maas

Costus comosus (Jacq.) Roscoe

*Costus comosus* (Jacq.) Roscoe var. *bakeri* K.Schum.

*Costus congestiflorus* Rich. ex Gagnep.

*Costus congestiflorus* Rich. ex Gagnep. var. β *glabrior* K.Schum.

*Costus congestus* Rowlee

*Costus conicus* Stokes

Costus convexus Maas & D.Skinner

Costus cordatus Maas

Costus cupreifolius Maas

*Costus curcumoides* Maas

Costus curvibracteatus Maas

*Costus cuspidatus* Nees & Mart.

*Costus cylindricus* auct. non Jacq.: Roscoe (1825) t. 78.

*Costus cylindricus* Jacq.

*Costus cylindricus* Jacq. var. y *acreanus* Loes.

*Costus cylindricus* Jacq. var. β *anachiri* (Jacq.) Petersen

*Costus cylindricus* Jacq. var. β *pulcherrima* (Kuntze) K.Schum.

*Costus cylindricus* Jacq. var. γ *ciliatus* (Miq.) Petersen

Costus dirzoi García-Mend. & G.Ibarra

*Costus discolor* Roscoe

Costus douglasdalyi Maas & H. Maas

*Costus dussii* K.Schum.

Costus erythrocoryne K.Schum.

Costus erythrophyllus Loes.

Costus erythrothyrsus Loes.

Costus fissicalyx N.R.Salinas, Clavijo & Betancur B.

*Costus flammulus* K.M.Kay & P.Juárez

*Costus formosus* C.V.Morton

Costus fortalezae K.Schum. *insuff*.

*Costus fragilis* Maas

*Costus friedrichsenii* auct. non Petersen: Woodson (1942) 329.

*Costus friedrichsenii* Petersen

*Costus fusiformis* Maas

*Costus gagnepainii* K.Schum.

Costus geothyrsus K.Schum.

Costus gibbosus D.Skinner & Maas

*Costus giganteus* Kuntze, non Welw. ex Ridley

*Costus glabratus* Sw.

*Costus glabratus* Sw. var. *niveo-purpureus* (Jacq.) Petersen

Costus glaucus Maas

*Costus gracilis* Loes.

Costus guanaiensis Rusby (as “guanaiense”)

*Costus guanaiensis* Rusby var. *asplundii* Maas

*Costus guanaiensis* Rusby var. *macrostrobilus* (K.Schum.) Maas

*Costus guanaiensis* Rusby var. *tarmicus* (Loes.) Maas

*Costus guianicus* Loes.

*Costus hirsutus* C.Presl

*Costus igneus* auct. non N.E.Br.: K.Goebel (1931) 161, f. 142.

*Costus igneus* N.E.Br.

Costus juruanus K.Schum.

*Costus juruanus* K.Schum. var. *strigosus* Maas

*Costus kaempferoides* Loes.

**Costus krukovii** (Maas) Maas & H.Maas

Costus kuntzei K.Schum.

Costus L.

Costus laevis Ruiz & Pav.

*Costus lanceolatus* Petersen

*Costus lanceolatus* Petersen subsp. *pulchriflorus* (Ducke) Maas

Costus lasius Loes.

*Costus latifolius* Gagnep.

*Costus laxus* Petersen

Costus leucanthus Maas

Costus lima K.Schum.

*Costus lima* K.Schum. var. *scabrimarginatus* Maas (as “scabremarginatus”)

*Costus lima* K.Schum. var. *wedelianus* Woodson

Costus longibracteolatus Maas (as “longebracteolatus”)

*Costus longifolius* Rusby

Costus lucanusianus J.Braun & K.Schum.

Costus macrostrobilus K.Schum.

Costus malortieanus H.Wendl.

*Costus malortieanus* Wendl. var. *amazonicus* Loes.

*Costus maritimus* Standl. & L.O.Williams

*Costus maximus* K.Schum.

*Costus mexicanus* Liebm. ex Petersen

*Costus micranthus* Gagnep.

Costus mollissimus Maas & H.Maas

Costus montanus Maas

*Costus mooreanus* Rusby

Costus nitidus Maas

*Costus niveo-purpureus* Jacq.

*Costus niveus* G.Mey. (as “nivea”)

*Costus nr. 4* Aubl. (nom. nud.)

*Costus nr. 5* Aubl. (nom. nud.)

*Costus nr. 6* Aubl. (nom. nud.)

*Costus nr. 7* Aubl. (nom. nud.)

*Costus nutans* K.Schum.

Costus obscurus D.Skinner & Maas

Costus oreophilus Maas & D.Skinner

Costus osae Maas & H.Maas

*Costus paucifolius* Gagnep.

*Costus phlociflorus* Rusby

*Costus pictus* D.Don

*Costus pilgeri* K.Schum.

*Costus pilosissimus* (Gagnep.) K.Schum.

*Costus pisonis* Lindley

Costus pitalito C.D.Specht & H.Maas

Costus plicatus Maas

Costus plowmanii Maas

*Costus podocephalus* Donn.Sm.

Costus prancei Maas & H.Maas

*Costus productus* Gleason ex Maas var. *productus*

*Costus productus* Gleason ex Maas var. *strigosus* (Maas) Maas

Costus pseudospiralis Maas & H.Maas

*Costus pubescens* Sp.Moore

*Costus pubescens* Sp.Moore forma *fibrillosus* Loes. (as “fibrillosa”)

*Costus puchucupango* J.F.Macbr.

Costus pulcherrimus Kuntze

*Costus pulchriflorus* Ducke

Costus pulverulentus C.Presl

*Costus pumilus* Petersen

*Costus pumilus* Petersen var. *pilosissimus* Gagnep.

*Costus quartus* Roem. & Schult. (nom. nud.)

*Costus quasi-appendiculatus* Woodson ex Maas

*Costus quintus* Roem. & Schult. (nom. nud.)

*Costus ramosus* Woodson

Costus ricus Maas & H.Maas

*Costus ruber* C.Wright ex Griseb.

Costus rubineus D.Skinner & Maas

*Costus rurrenabaqueanus* Rusby

*Costus sanguineus* Donn.Sm.

*Costus scaber* auct. non Ruiz & Pav.: Loes. (1931) 89.

Costus scaber Ruiz & Pav.

*Costus scaber* Ruiz & Pav. x *Costus lasius* Loes.

*Costus scaberulus* Rich. ex Gagnep.

*Costus secundus* Roem. & Schult.

Costus sepacuitensis Rowlee

*Costus septimus* Aubl. ex Roem. & Schult. (nom. nud).

*Costus sextus* Roem. & Schult. (nom. nud.)

Costus sinningiiflorus Rusby (as “sinningiaeflorus”).

*Costus skutchii* C.V.Morton

*Costus sp. B* Maas

**Costus sp. nov. Chambi**

**Costus sp. nov. Gentry**

*Costus spec. A* Maas (1972) 124.

Costus spicatus (Jacq.) Sw.

*Costus spicatus* auct. non Jacq.: Standl. (1928) 118, pl. 15.

*Costus spicatus* Sw. β *anthocono purpurascente*… Horan.

Costus spiralis (Jacq.) Roscoe

*Costus spiralis* (Jacq.) Roscoe var. *villosus* Maas

*Costus spiralis* (Jacq.) Roscoe var. γ *hirsutus* Petersen

*Costus spiralis* (Jacq.) Roscoe α *jacquinii* Griseb.

*Costus spiralis* (Jacq.) Roscoe β *pisonis* Griseb.

*Costus spiralis* (Jacq.) Roscoe γ *roscoei* Griseb.

*Costus spiralis* auct. non (Jacq.) Roscoe: Woodson (1942) 331.

*Costus splendens* Donn.Smith & Türckh.

Costus sprucei Maas

*Costus steinbachii* Loes.

Costus stenophyllus Standl. & L.O.Williams

*Costus subsessilis* Nees & Mart.

*Costus tarapotensis* J.F.Macbr.

*Costus tarmicus* Loes.

*Costus tatei* Rusby

Costus ulei Loes. *insuff*.

*Costus uniflorus* Poepp. ex Petersen

*Costus validus* Loes.

Costus vargasii Maas & H.Maas

Costus varzearum Maas

*Costus verschaffeltii* (as “verschaffeltianus”)

*Costus villosissimus* auct. non Jacq.: Rowlee (1922) 287, in part.

Costus villosissimus Jacq.

Costus vinosus Maas

*Costus warmingii* Petersen

Costus weberbaueri Loes.

Costus wilsonii Maas

Costus whiskeycola Maas & H.Maas

Costus woodsonii Maas

Costus zamoranus Steyerm.

Curcuma

Cyathea sp.

Cymbopetalum

Deppea sp.

Dialium

Dictyocaryum lamarckianum

**Dimerocostus appendiculatus** (Maas) Maas & H.Maas

Dimerocostus argenteus (Ruiz & Pav.) Maas

*Dimerocostus bicolor* J.F.Macbr.

*Dimerocostus bolivianus* (Rusby) Loes.

Dimerocostus cryptocalyx N.R.Salinas & Betancur B.

*Dimerocostus elongatus* J.Huber

*Dimerocostus gutierrezii* Kuntze (as “*guttierezii*”)

Dimerocostus Kuntze

**Dimerocostus rurrenabaqueanus** (Rusby) Maas & H.Maas

Dimerocostus strobilaceus Kuntze

*Dimerocostus strobilaceus* Kuntze subsp. *appendiculatus* Maas

*Dimerocostus strobilaceus* Kuntze subsp. *gutierrezii* (Kuntze) Maas

*Dimerocostus tessmannii* Loes.

*Dimerocostus uniflorus* (Poepp. ex Petersen) K.Schum.

*Dimerocostus williamsii* J.F.Macbr.

Ficus

*Gissanthe* Salisbury

*Glissanthe* Steud.

Hedyosmum mexicanum

Heliconia sp.

Heliconiaceae

*Iacuacanga, aliis Paco Caatinga* Piso

Iriartea deltoidea

*Jacuanga* Lestiboudois

Lauraceae

Liquidambar macrophylla

Marantaceae

Matayba sp.

Melastomataceae

Monocostus K.Schum.

*Monocostus ulei* K.Schum.

Monocostus uniflorus (Poepp. ex Petersen) Maas

*Mulfordia boliviana* Rusby

*Mulfordia* Rusby

Myrtaceae

Oenocarpus bataua

*Paco Caatinga clava rubente maior* Plum.

*Paco caatinga floribus amplioribus niveis et purpureis* Plum.

*Paco-caatinga caule spirali minor* Plum.

*Paco-Caatinga villosissima flore luteo* Plum.

Posoqueria latifolia

Renealmia cernua (Sw. ex Roem. & Schult.) J.F.Macbr.

Renealmia sp.

Salvia divinorum

Siparuna andina

Sloanea tuerckheimii

Solenophora sp.

Tapirira

Zingiberaceae

Zingiberales

*Zinziber villosissimum floribus luteis* Plum.

## Figures, Plates, Maps and corresponding Legends

**Map 1.**
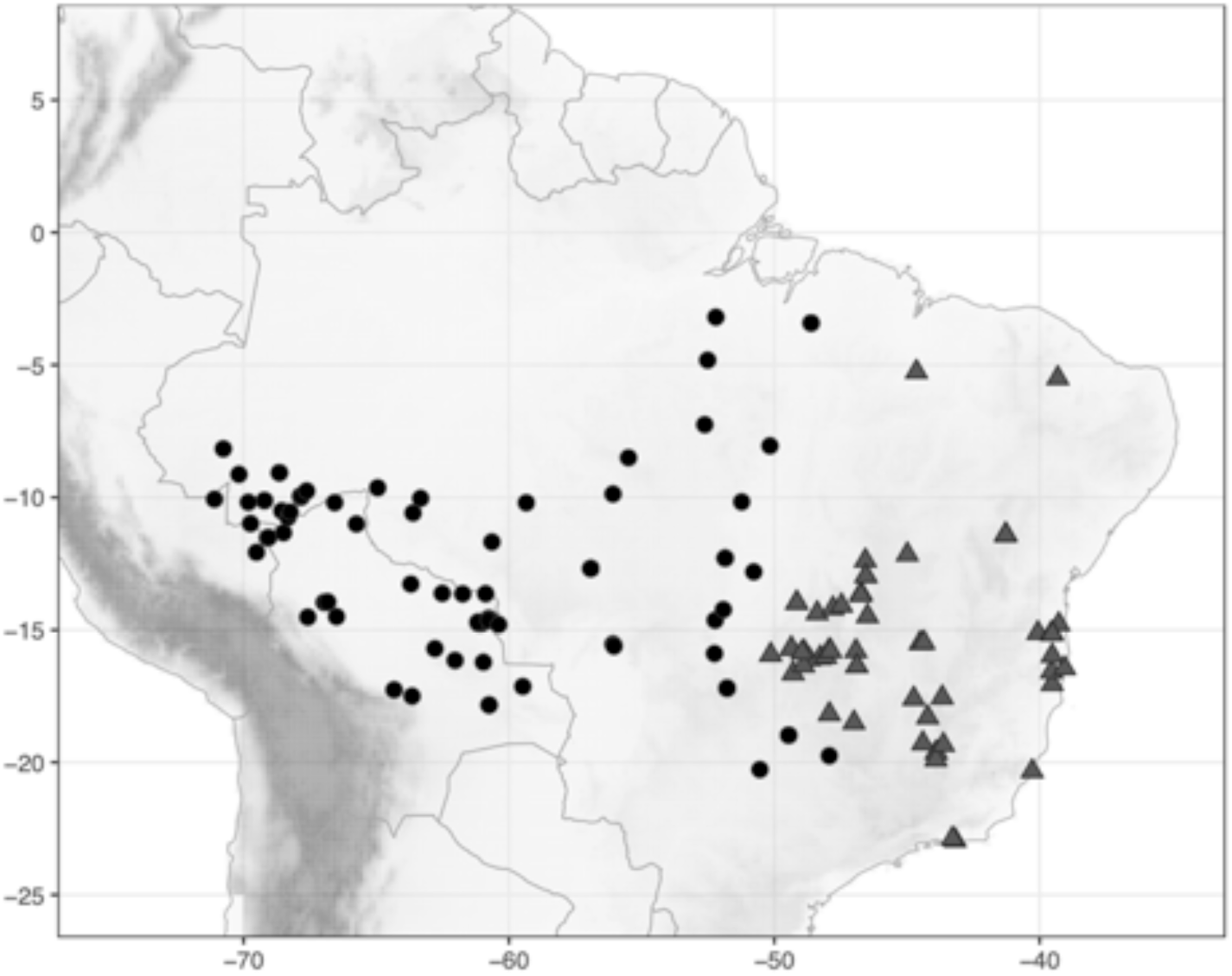
Distribution of *Chamaecostus acaulis* (S.Moore) T.André & C.D.Specht (●) and *Ch. subsessilis* (Nees & Mart.) C.D.Specht & D.W.Stev. (▲).

**Map 2.**
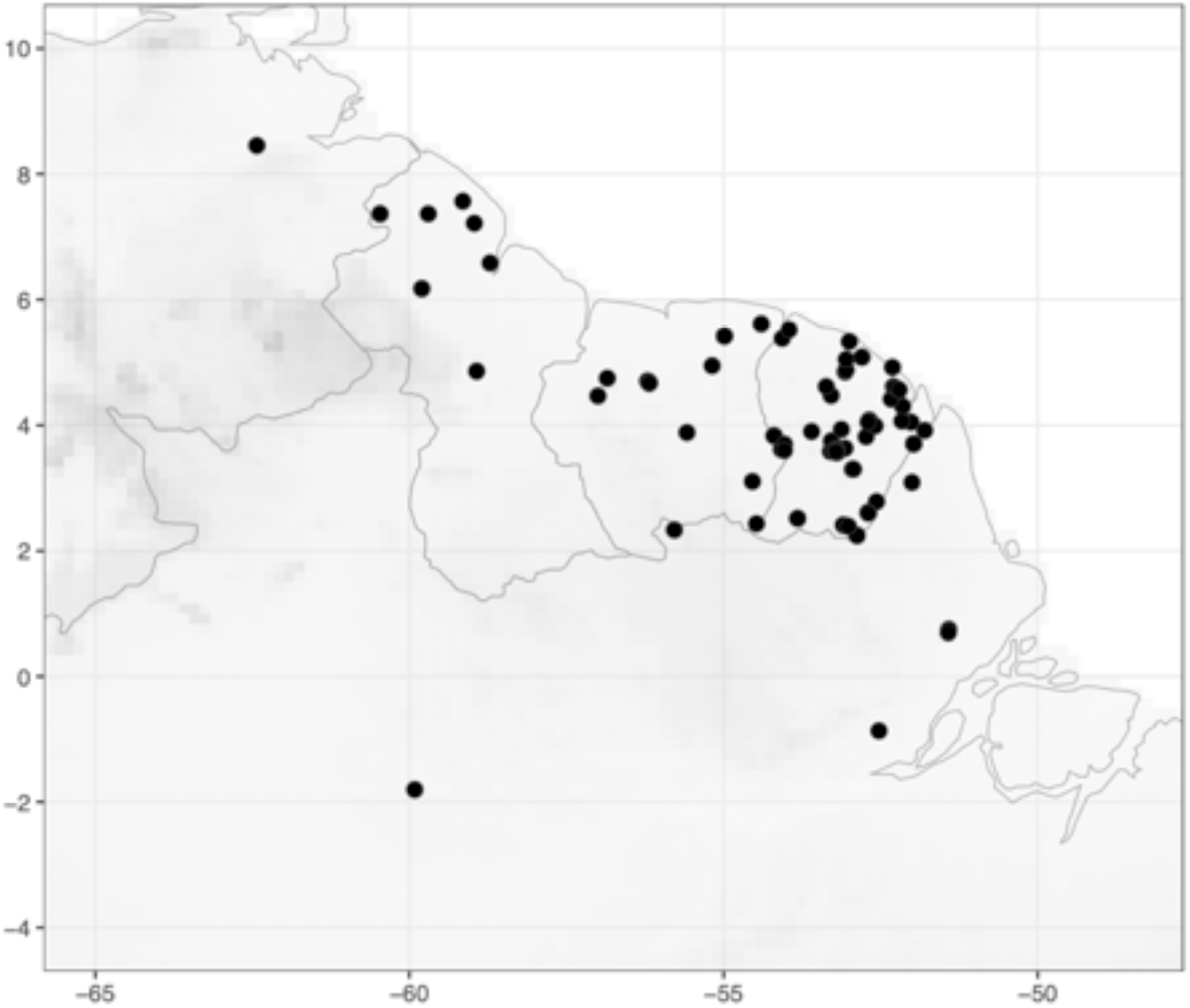
Distribution of *Chamaecostus congestiflorus* (Rich. ex Gagnep.) C.D.Specht & D.W.Stev. (●).

**Map 3.**
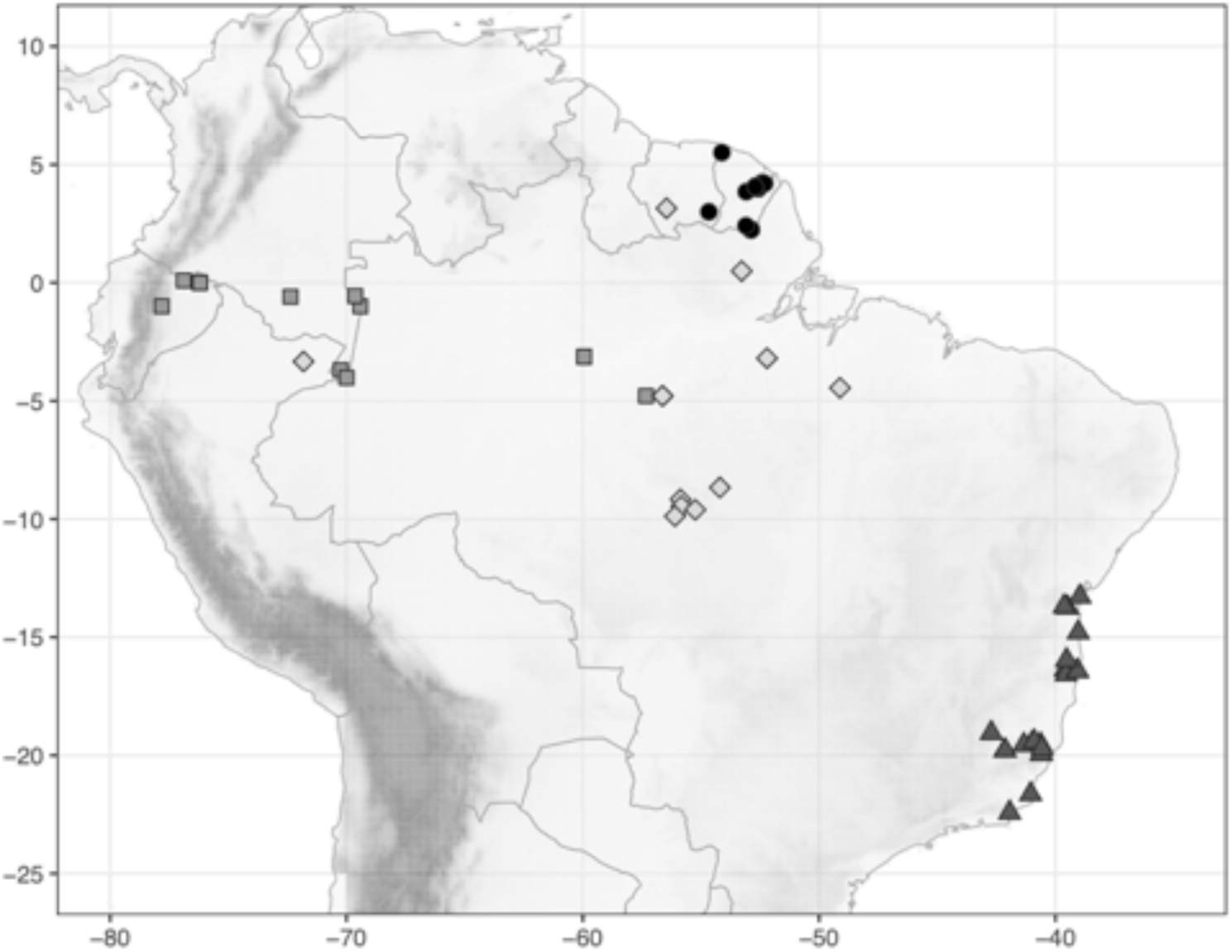
Distribution of *Chamaecostus curcumoides* (Maas) C.D.Specht & D.W.Stev. (●), *Ch. cuspidatus* (Nees & Mart.) C.D.Specht & D.W.Stev. (▲), *Ch. fragilis* (Maas) C.D.Specht & D.W.Stev. (◼), and *Ch. fusiformis* (Maas) C.D.Specht & D.W.Stev. (◆).

**Map 4.**
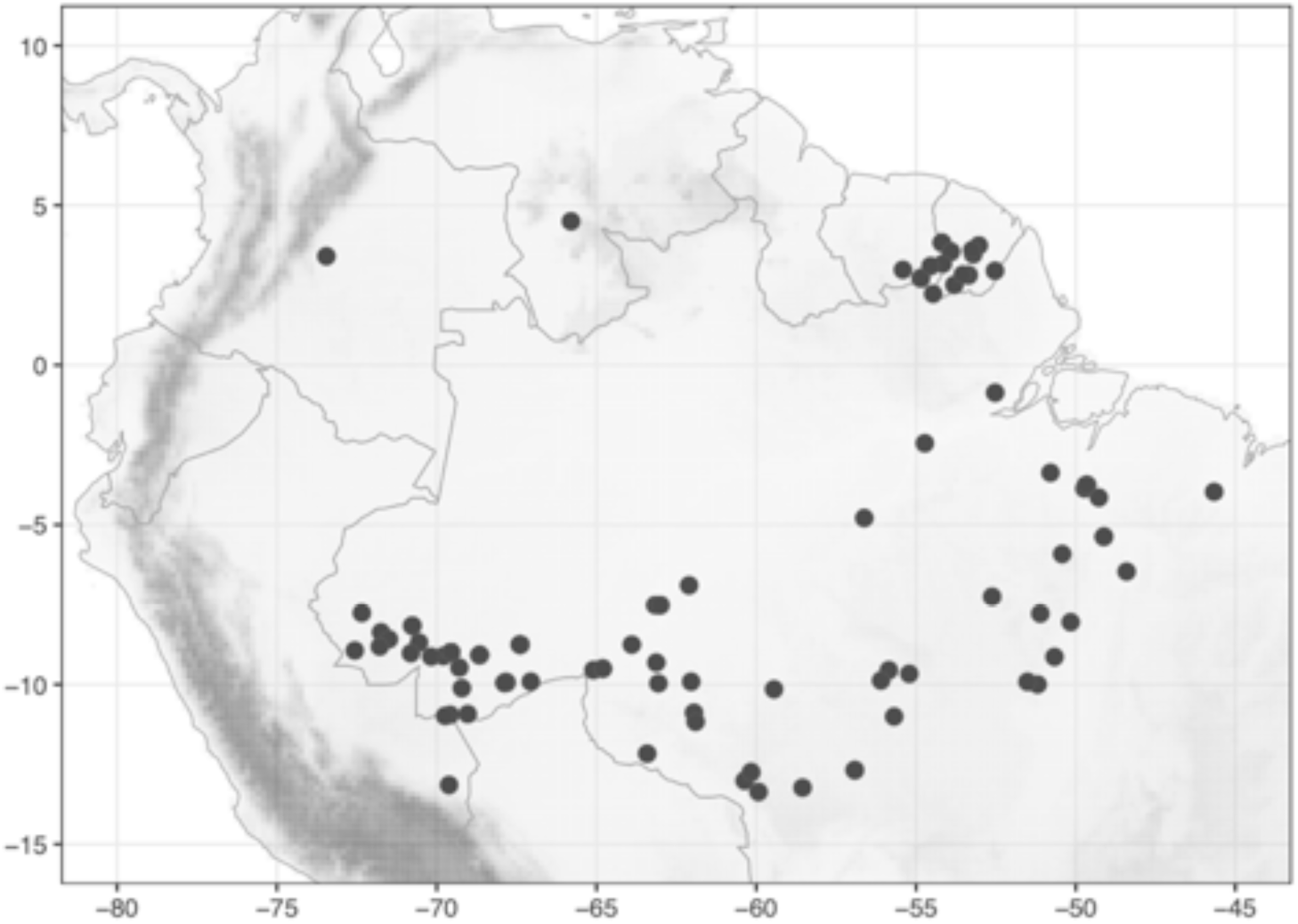
Distribution of *Chamaecostus lanceolatus* (Petersen) C.D.Specht & D.W.Stev. (●).

**Map 5.**
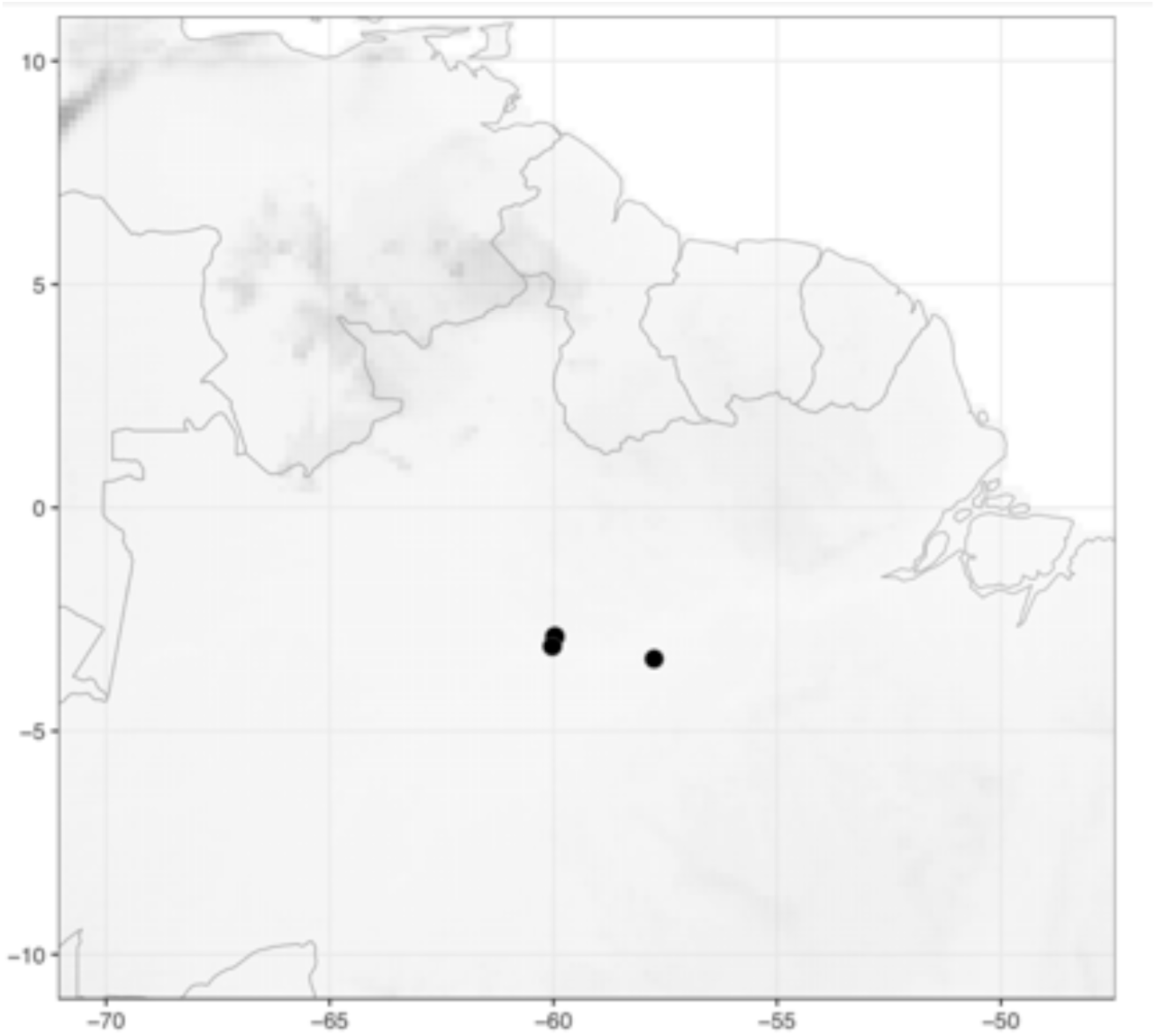
Distribution of *Chamaecostus manausensis* Maas & H.Maas (●).

**Map 6.**
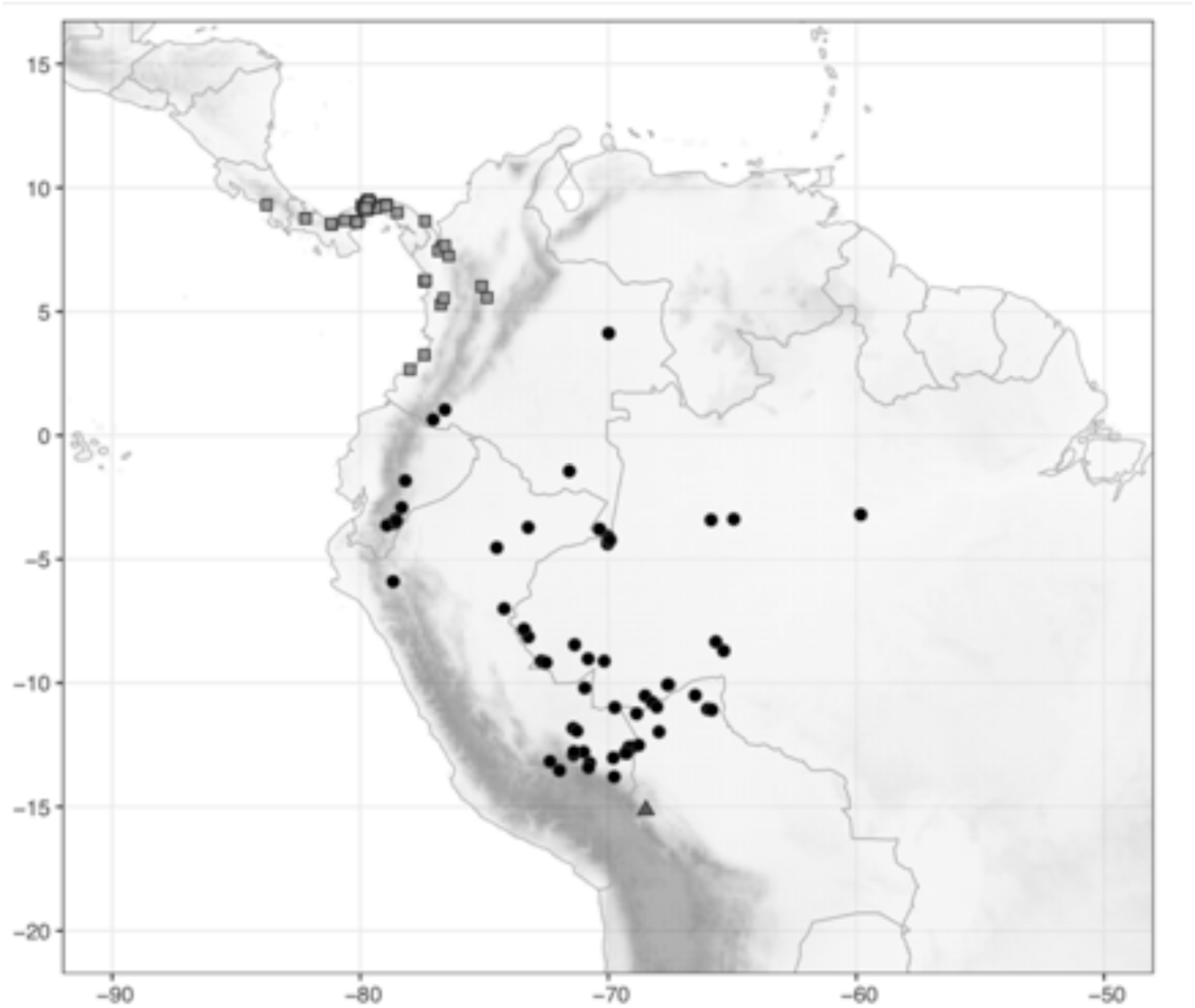
Distribution of *Costus acreanus* (Loes.) Maas (●), *C. alfredoi* Maas & H.Maas (▲), and *C. allenii* Maas (◼).

**Map 7.**
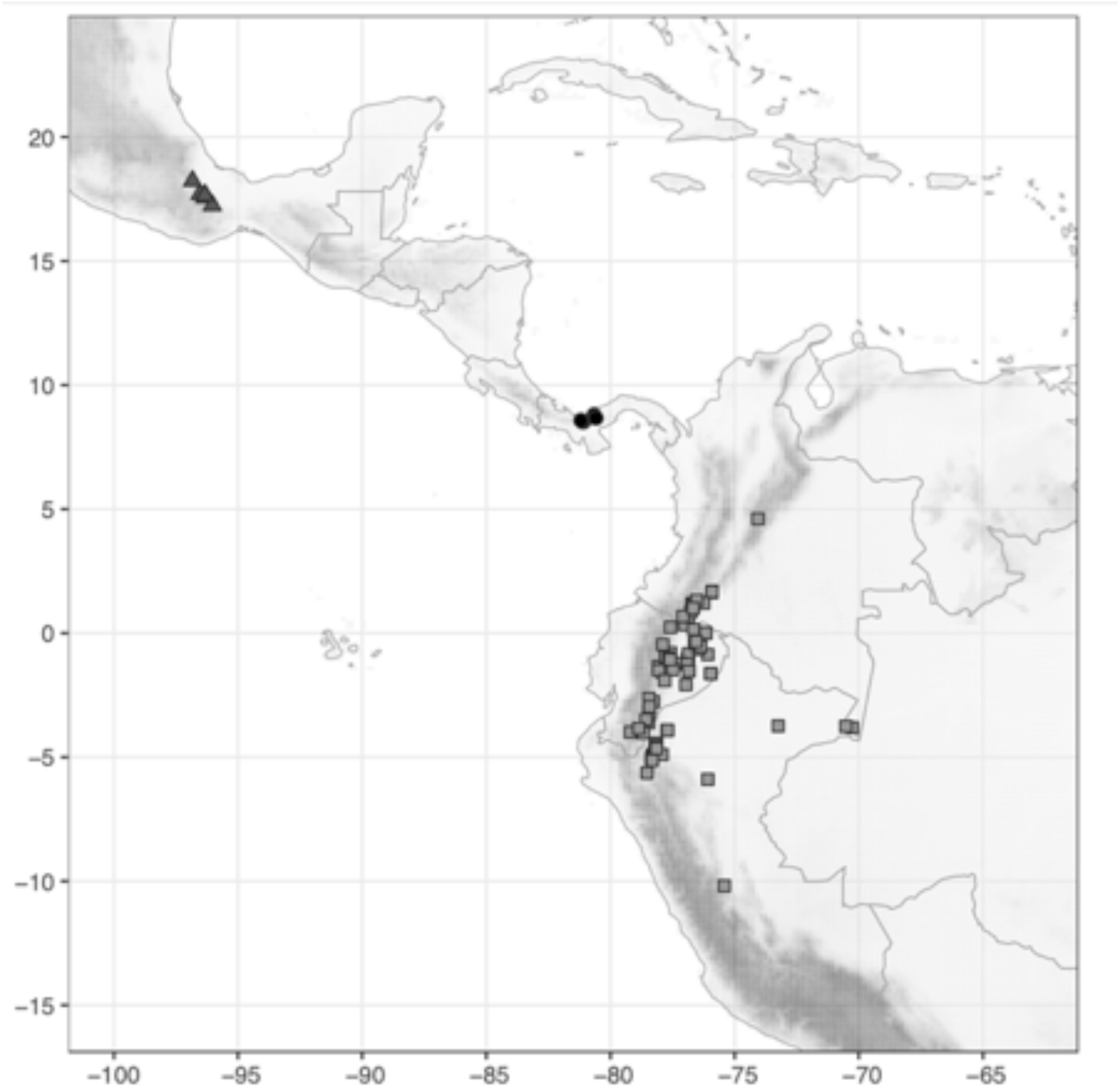
Distribution of *Costus allenopsis* Maas & D.Skinner (●), *C. alticolus* Maas & H.Maas (▲), and *C. amazonicus* (Loes.) J.F.Macbr. (◼).

**Map 8.**
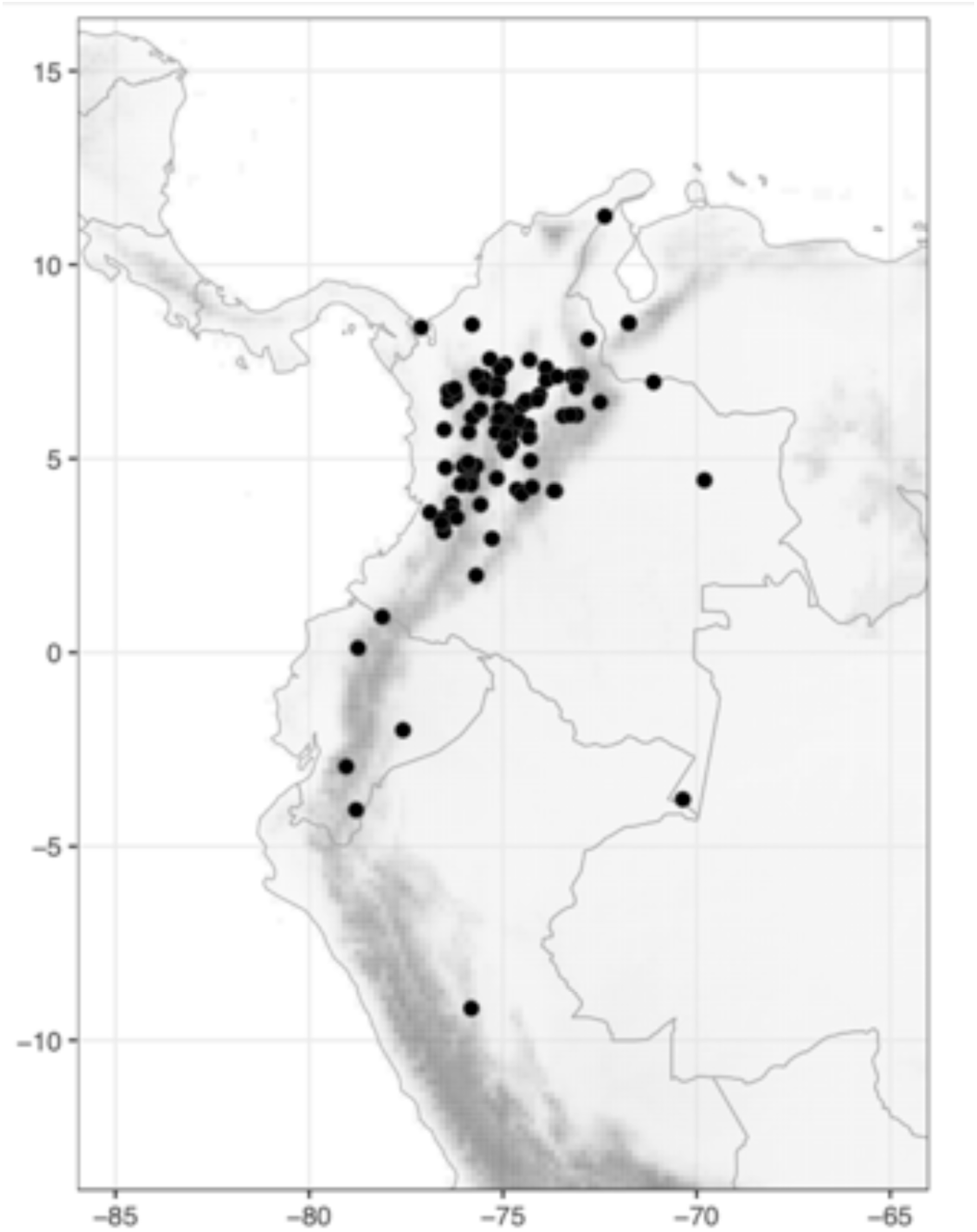
Distribution of *Costus antioquiensis* Maas & H.Maas (●).

**Map 9.**
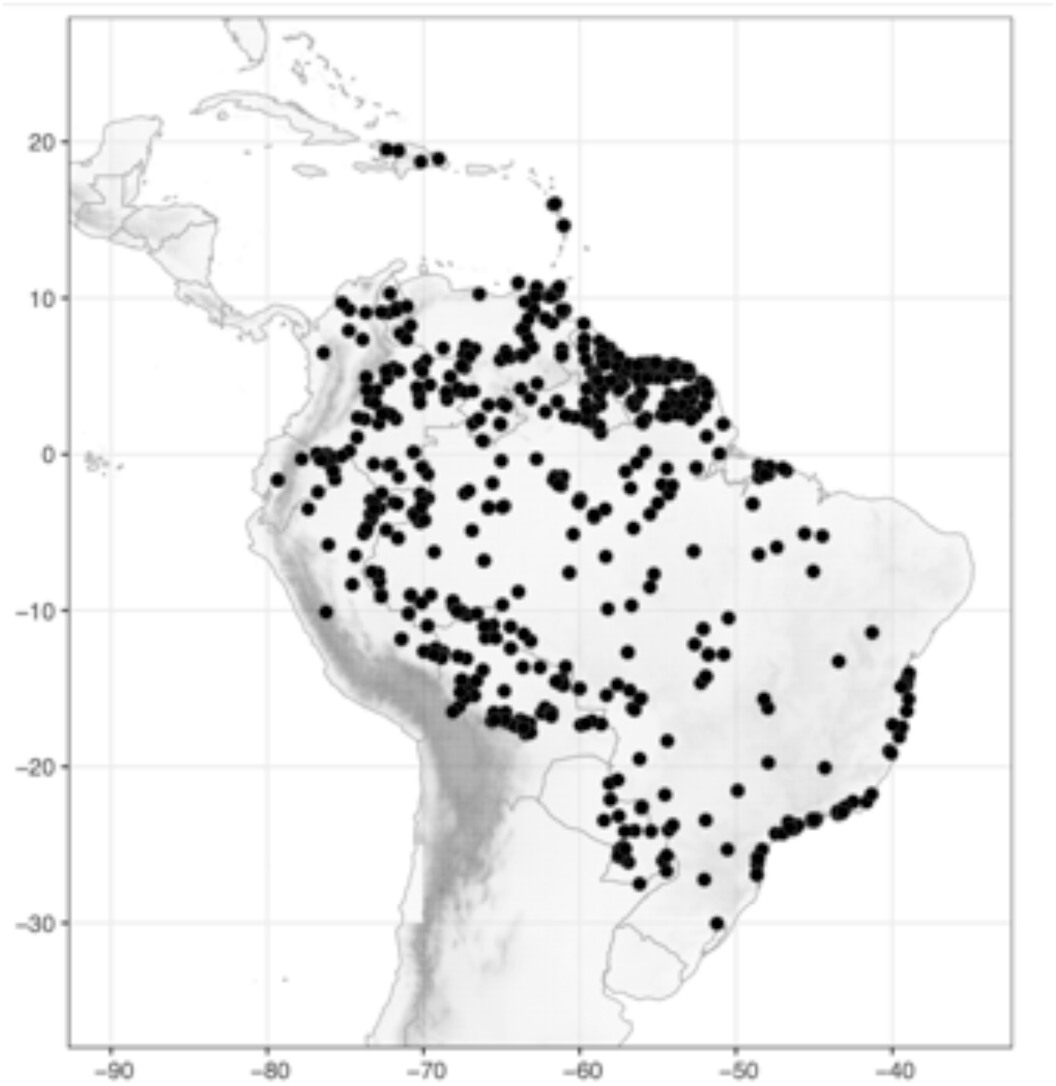
Distribution of *Costus arabicus* L. (●).

**Map 10.**
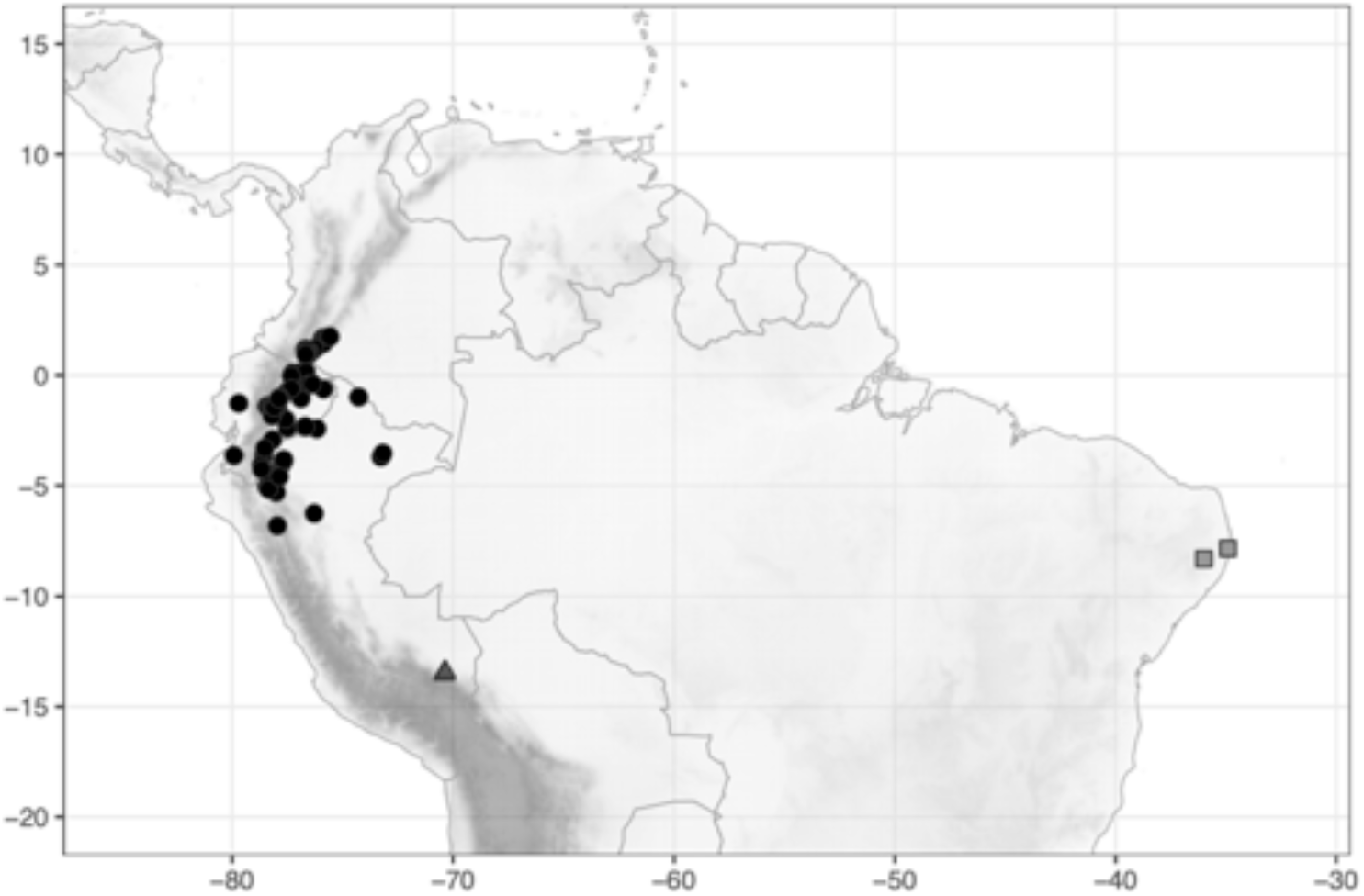
Distribution of *Costus asplundii* (Maas) Maas (●), *C. asteranthus* Maas & H.Maas (▲), and *C. atlanticus* E.Pessoa & M.Alves (◼).

**Map 11.**
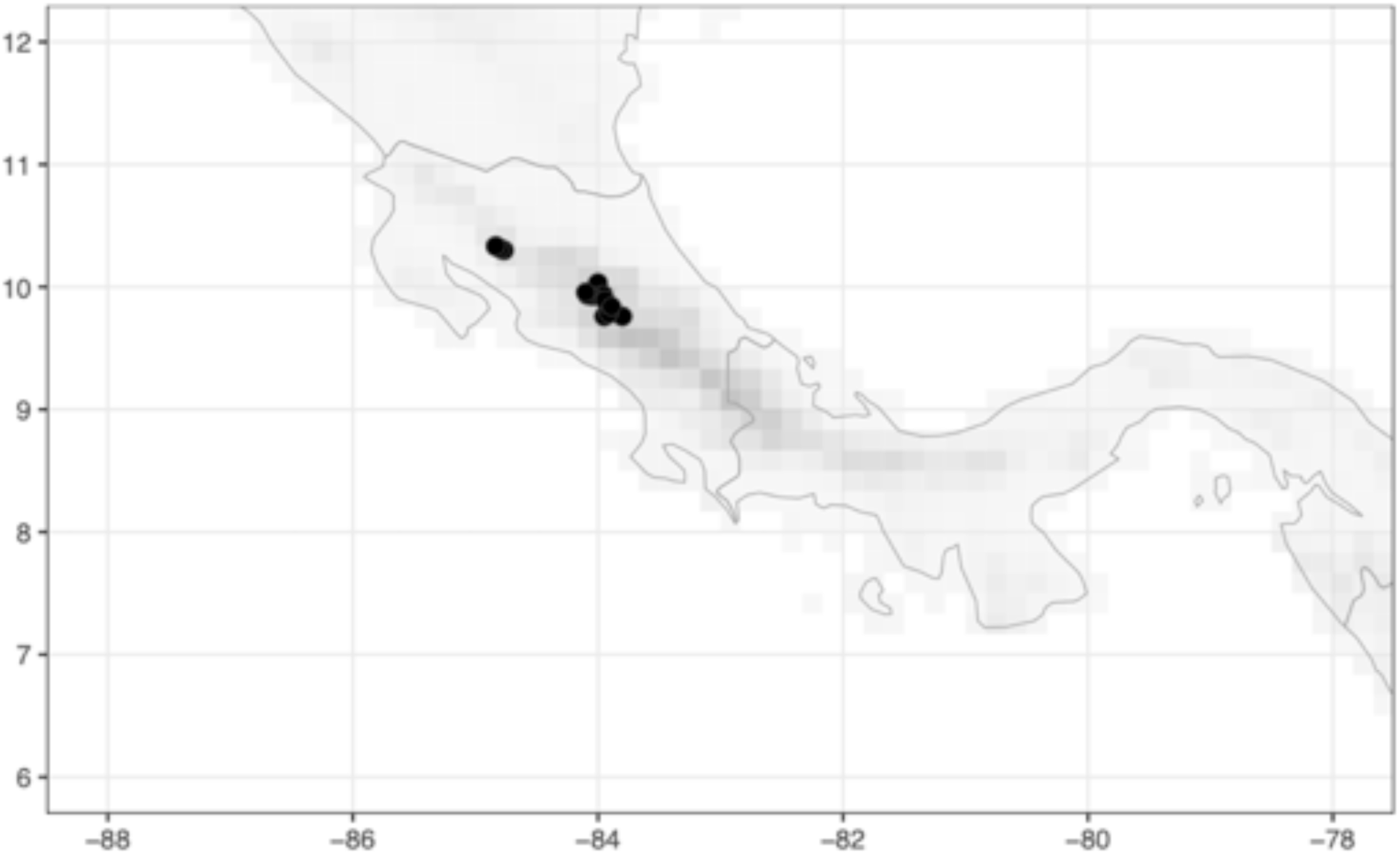
Distribution of *Costus barbatus* Suess. (●).

**Map 12.**
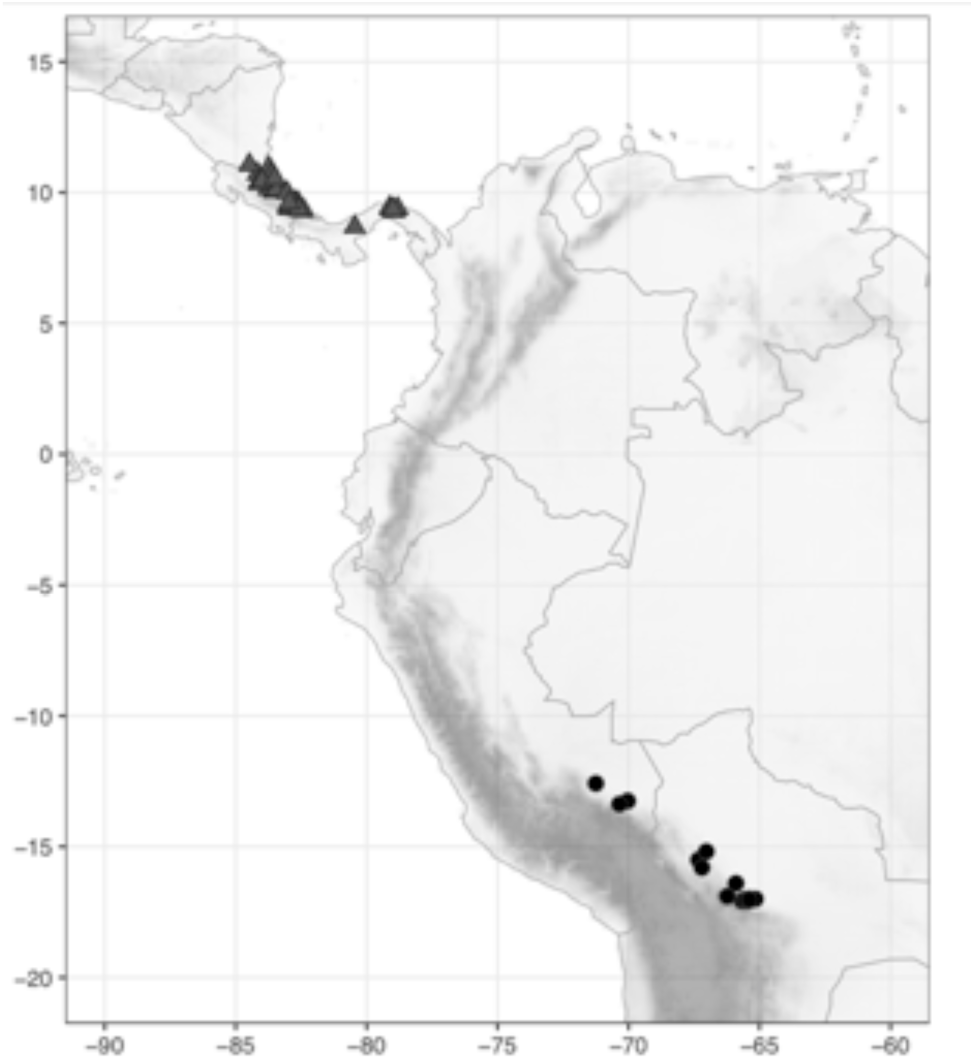
Distribution of *Costus beckii* Maas & H.Maas (●) and *C. bracteatus* Rowlee (▲).

**Map 13.**
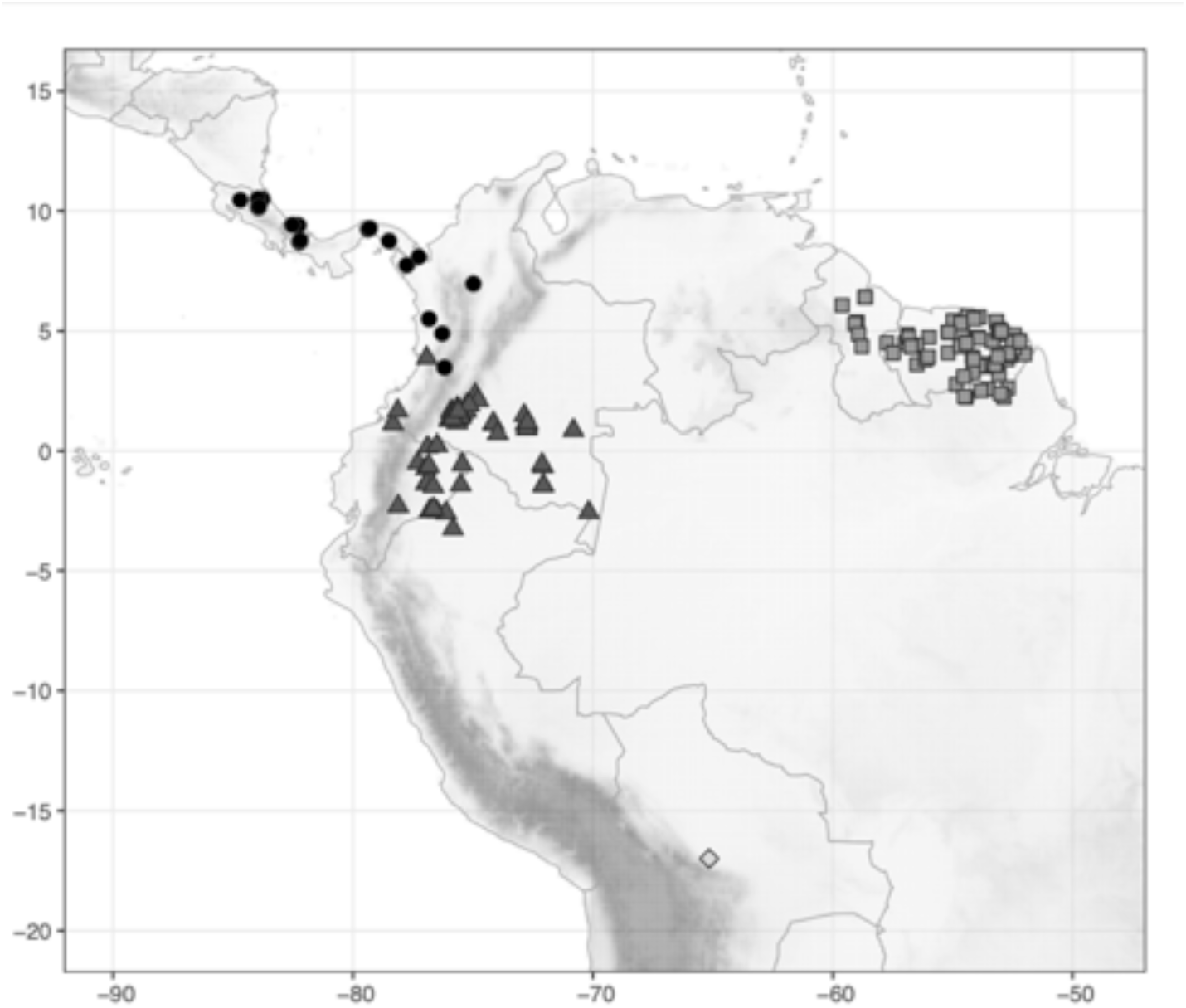
Distribution of *Costus callosus* Maas & H.Maas (●), *C. chartaceus* Maas (▲), *C. claviger* Benoist (◼), and *C. cochabambe* Maas & H.Maas (◆).

**Map 14.**
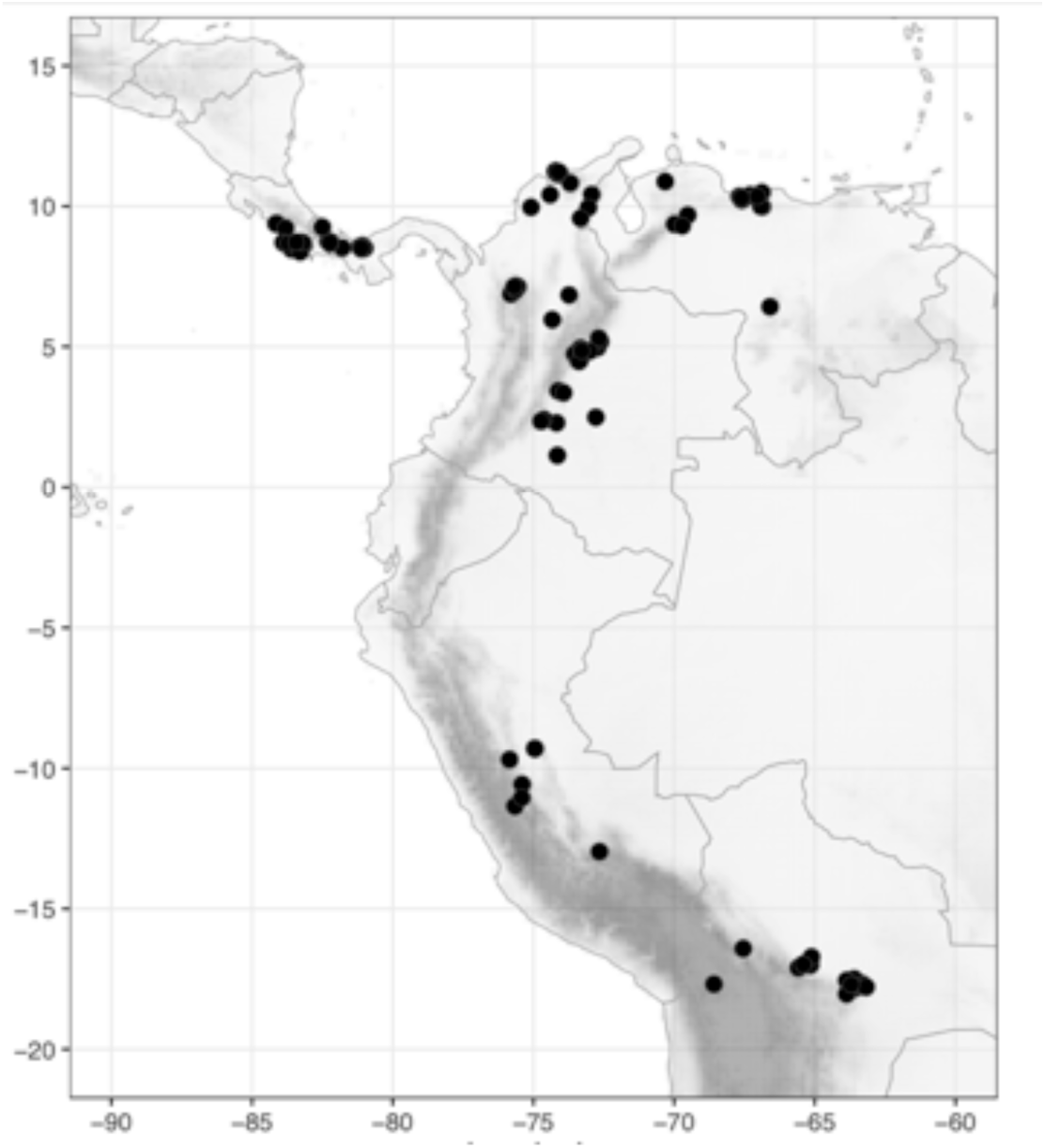
Distribution of *Costus comosus* (Jacq.) Roscoe (●).

**Map 15.**
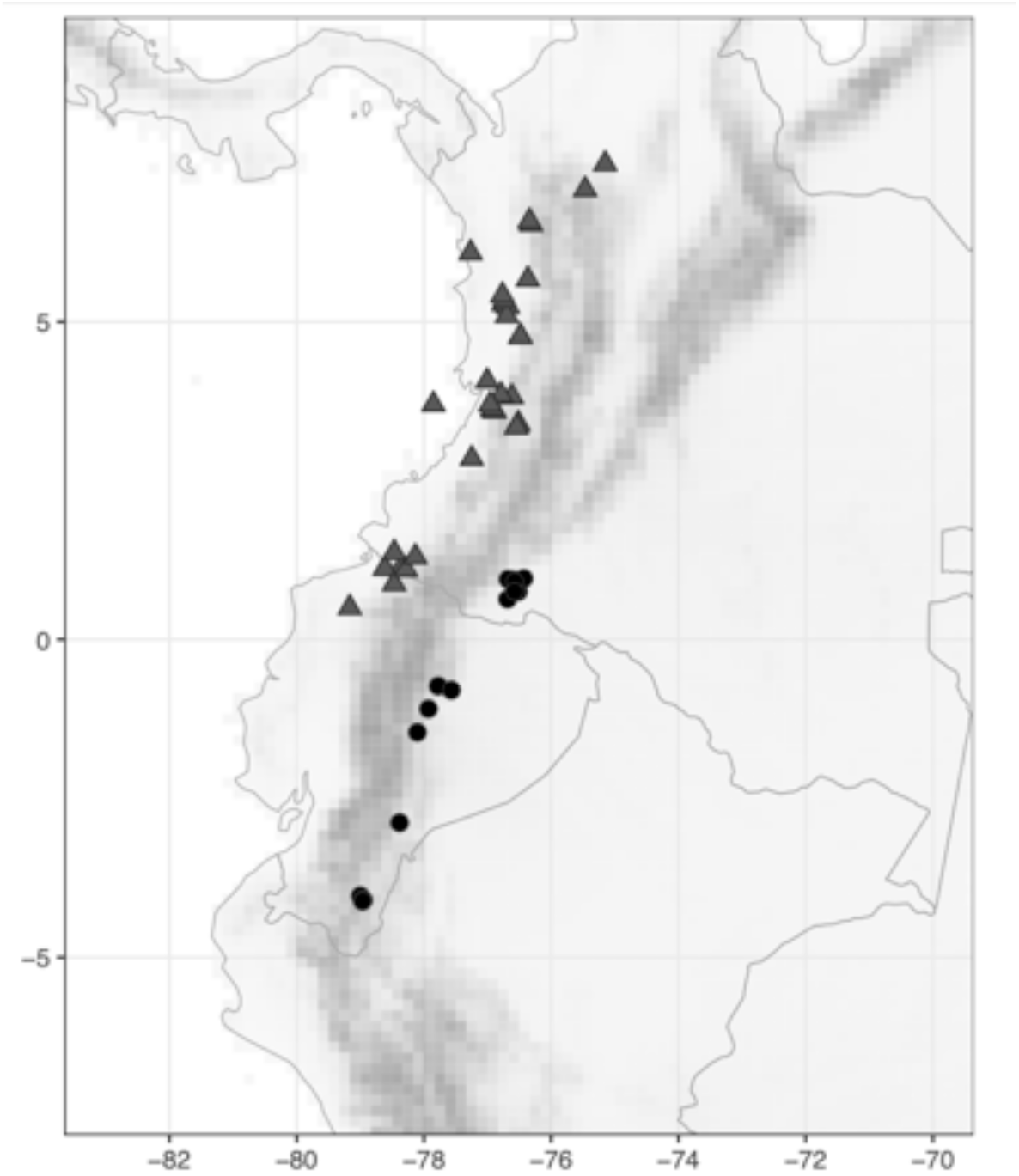
Distribution of *Costus convexus* Maas & D.Skinner (●) and *C. cordatus* Maas (▲).

**Map 16.**
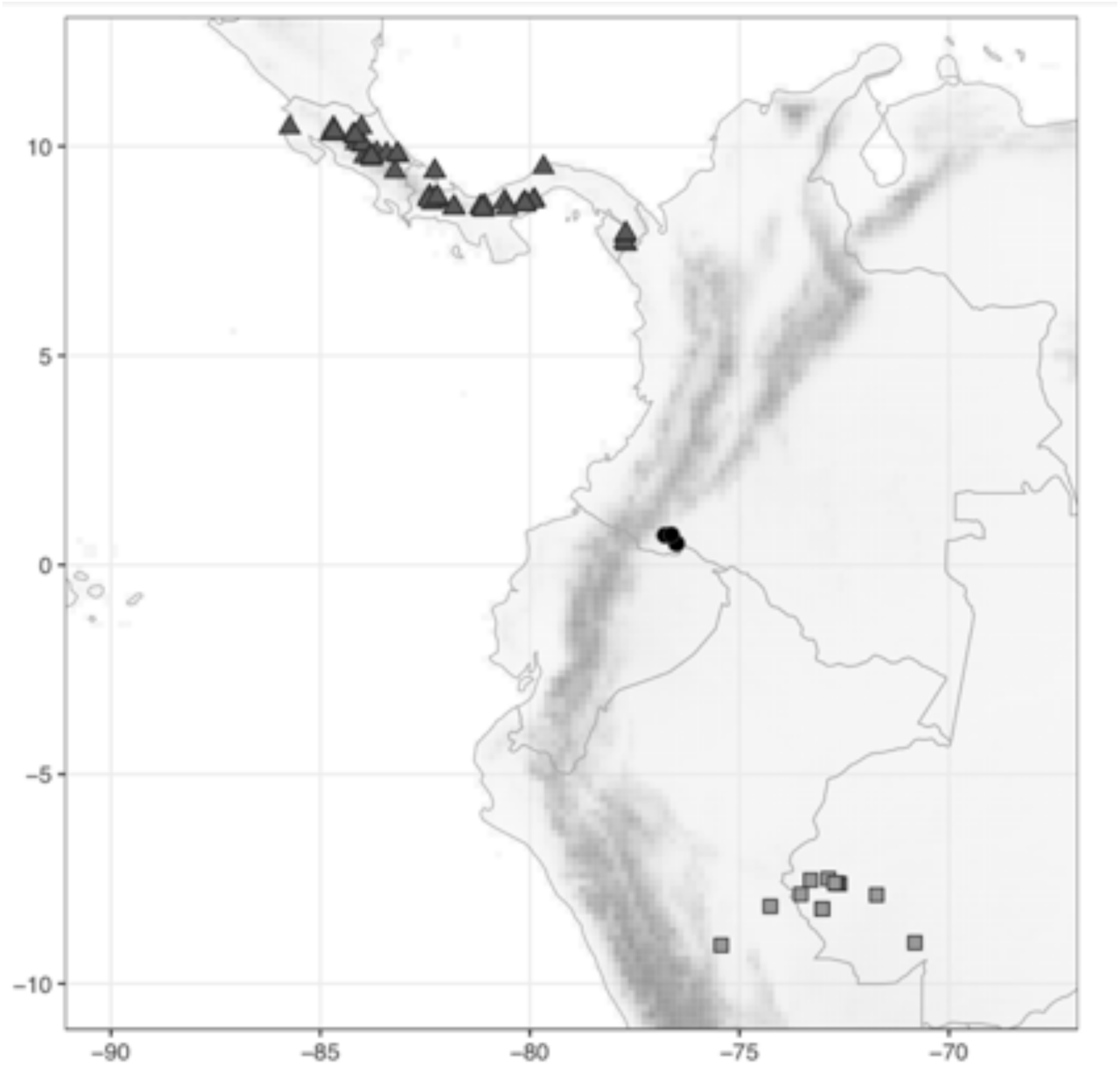
Distribution of *Costus cupreifolius* Maas (●), *C. curvibracteatus* Maas (▲), and *C. douglasdalyi* Maas & H. Maas (◼).

**Map 17.**
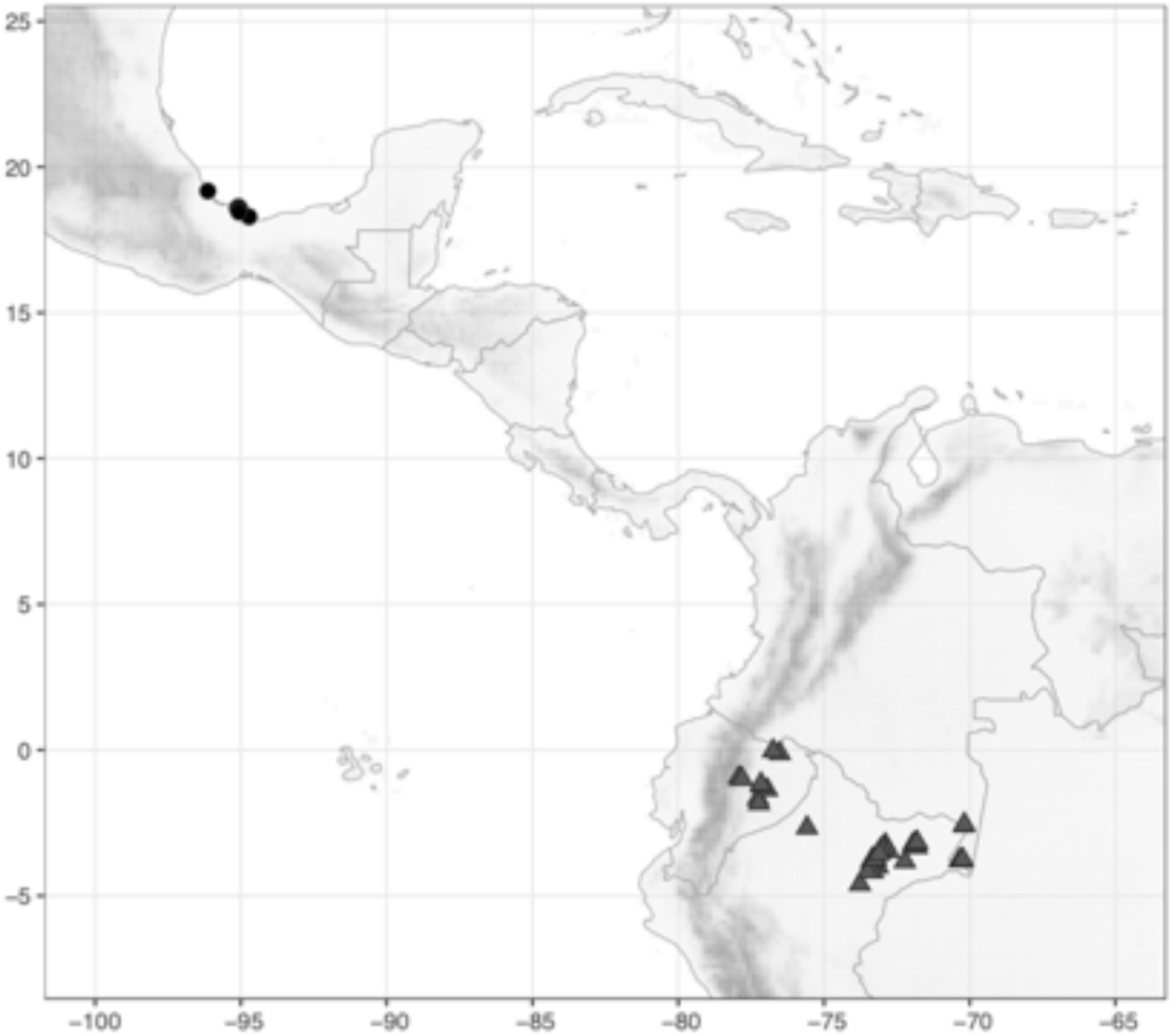
Distribution of *Costus dirzoi* García-Mend. & G.Ibarra (●) and *C. erythrocoryne* K.Schum. (▲).

**Map 18.**
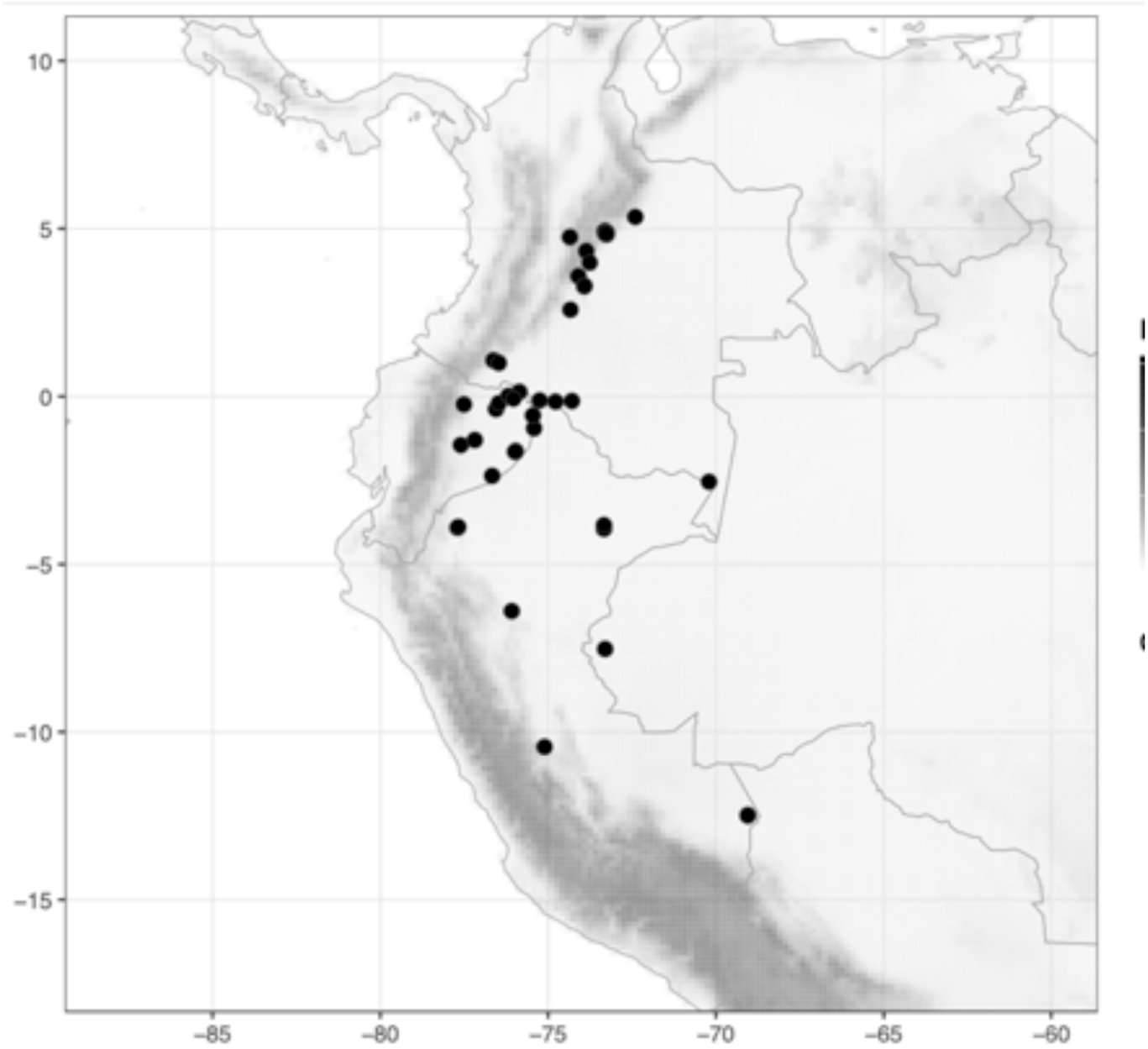
Distribution of *Costus erythrophyllus* Loes. (●).

**Map 19.**
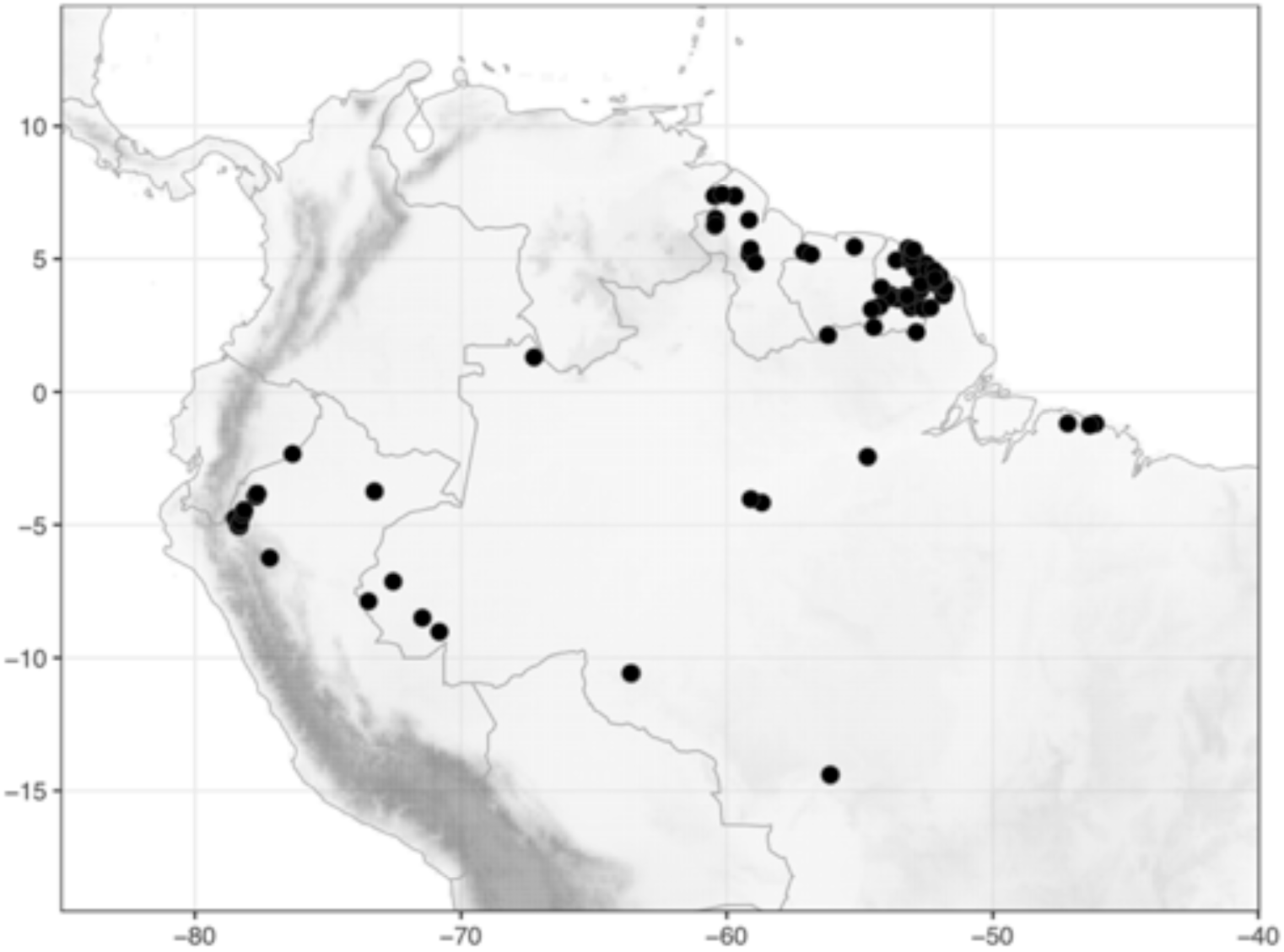
Distribution of *Costus erythrothyrsus* Loes. (●).

**Map 20.**
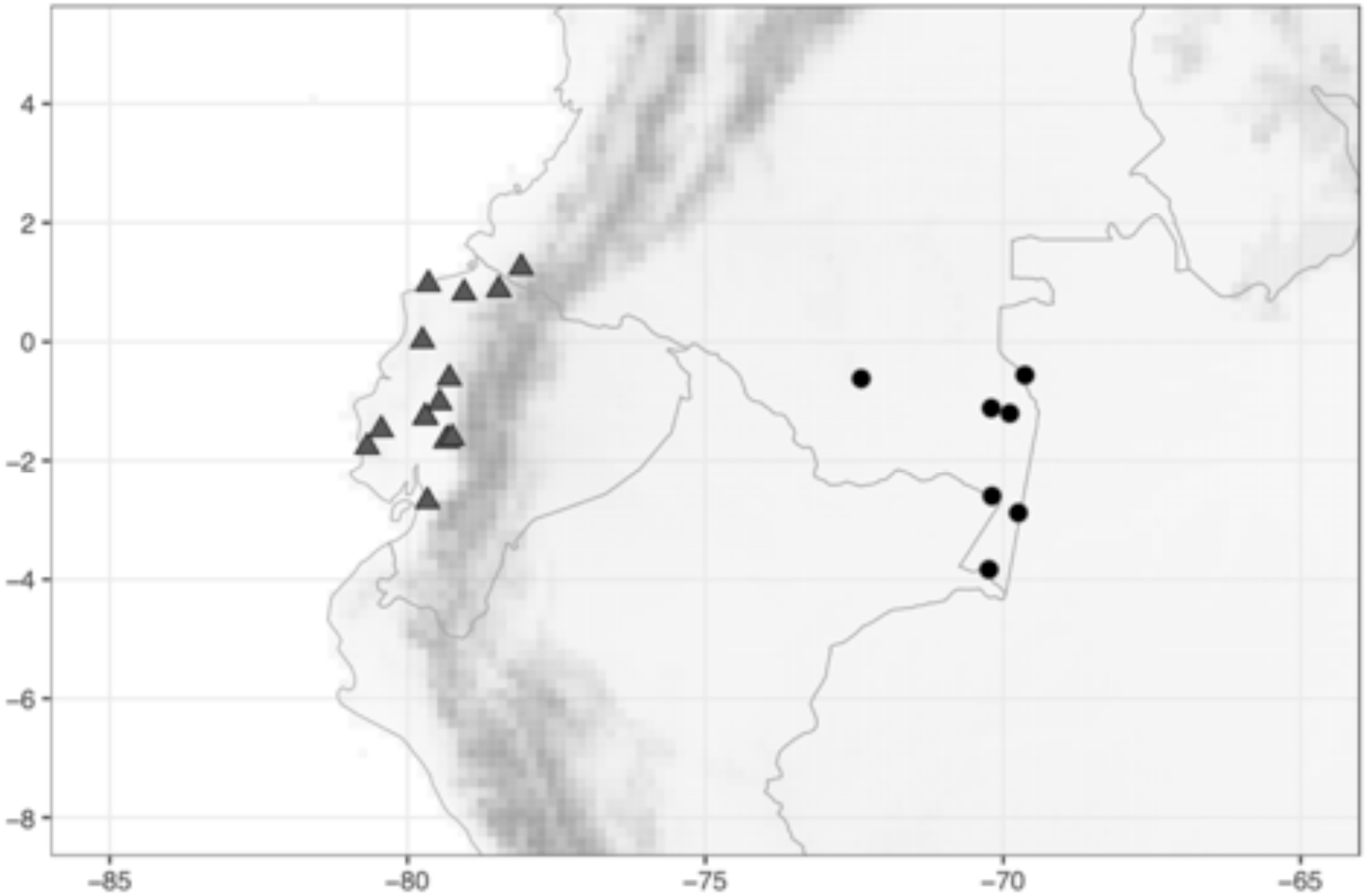
Distribution of *Costus fissicalyx* N.R.Salinas, Clavijo & Betancur B. (●) and *C. geothyrsus* K.Schum. (▲).

**Map 21.**
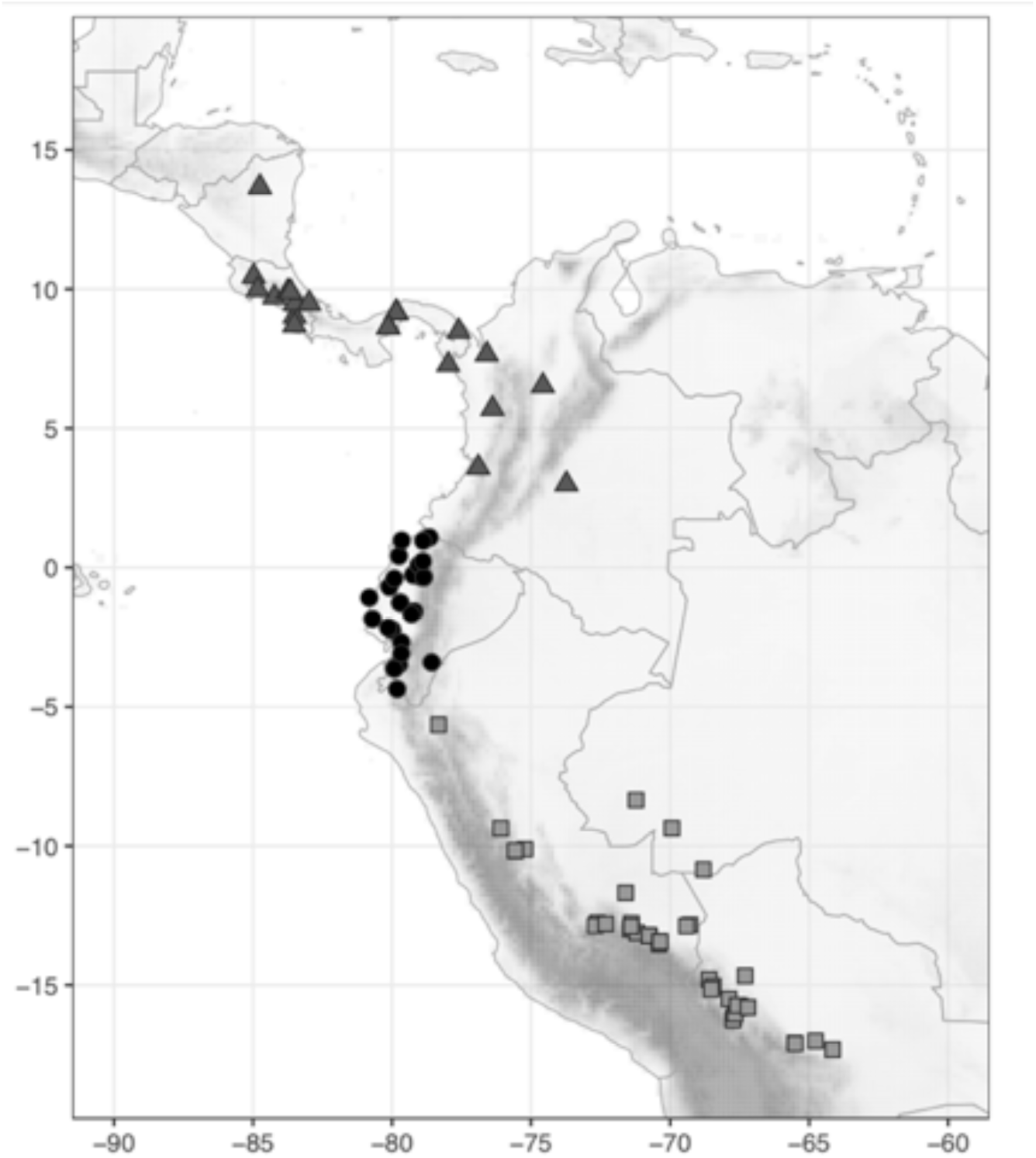
Distribution of *Costus gibbosus* D.Skinner & Maas (●), *C. glaucus* Maas (▲), and *C. guanaiensis* Rusby (◼).

**Map 22.**
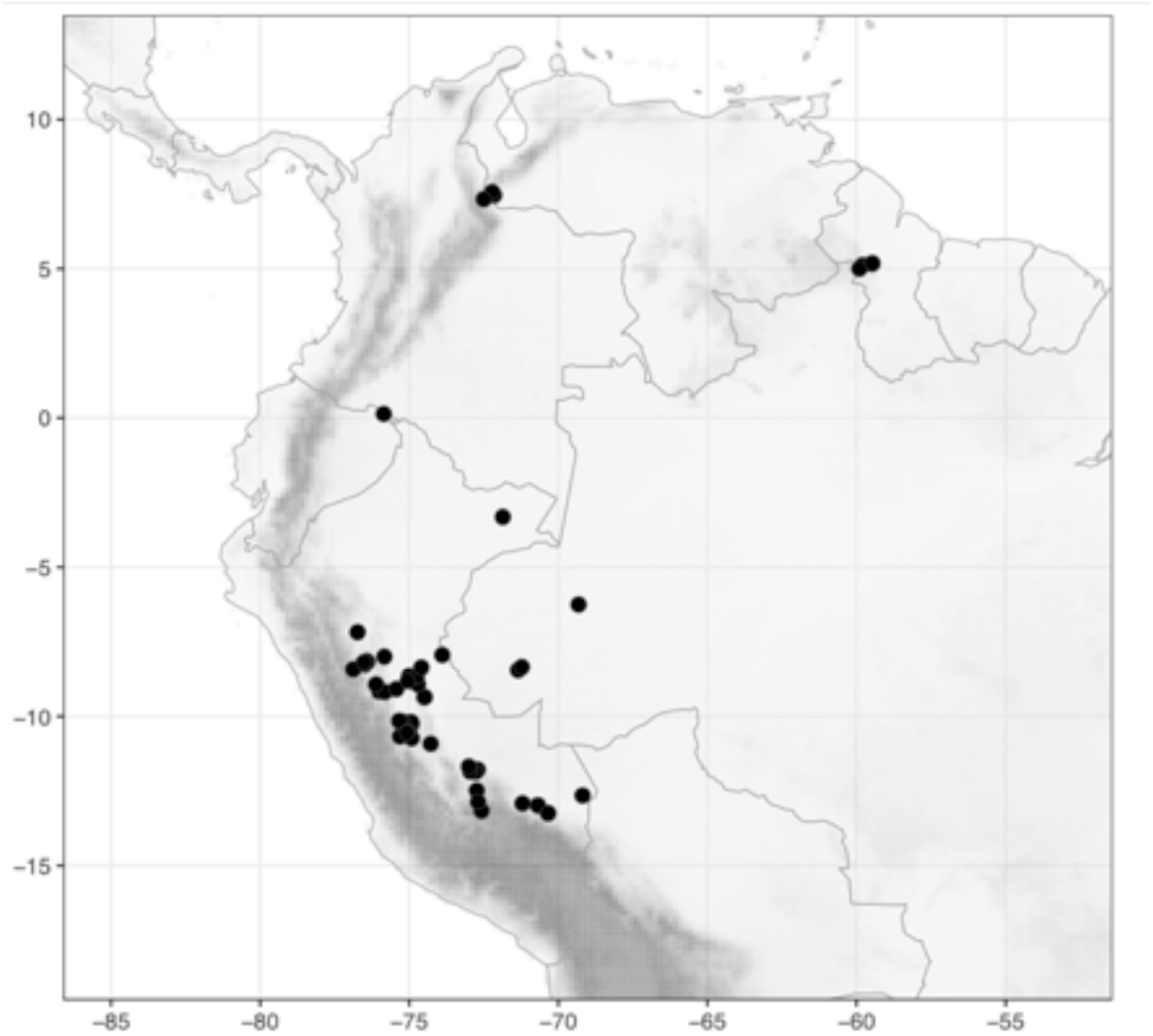
Distribution of *Costus juruanus* K.Schum. (●).

**Map 23.**
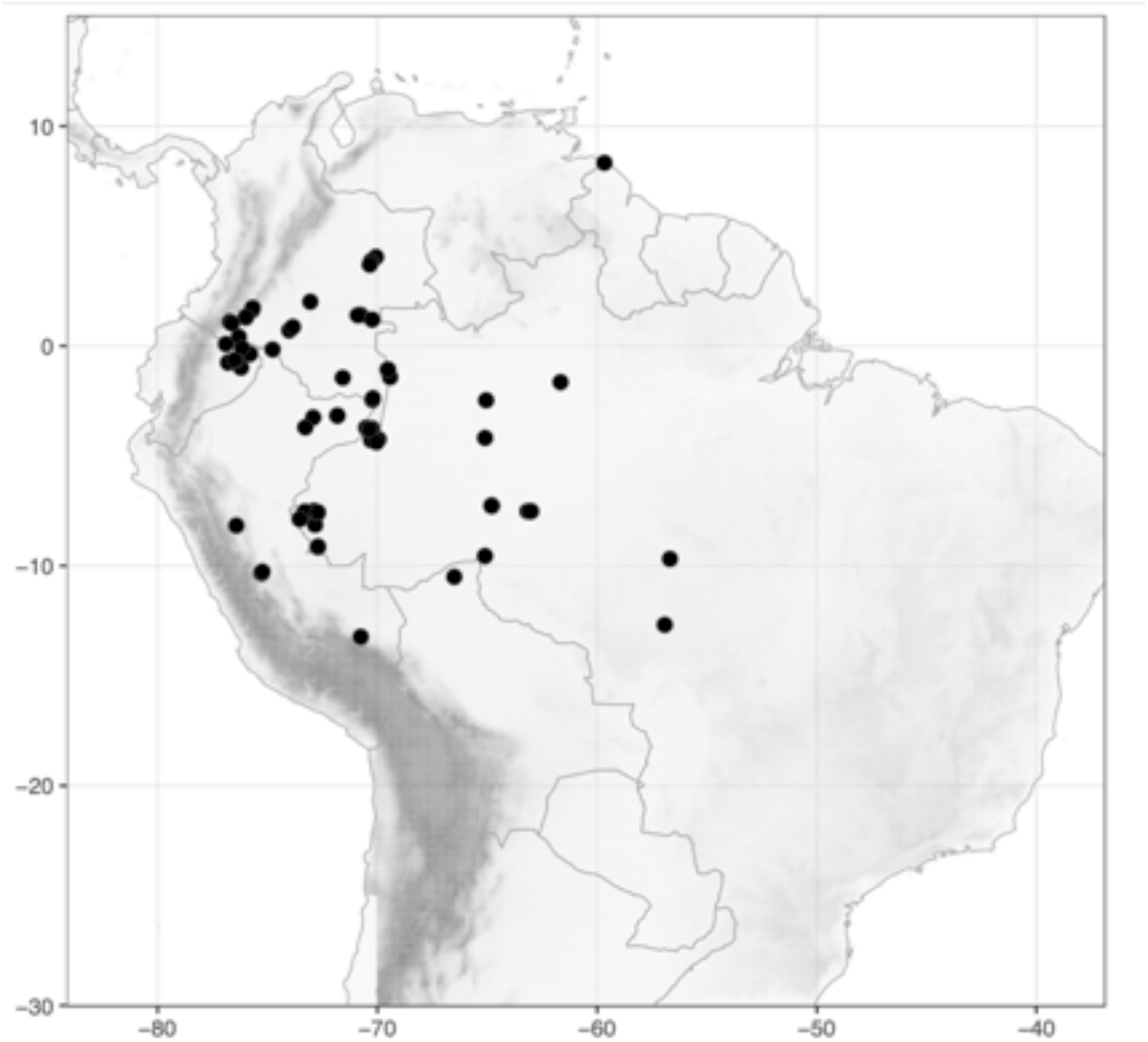
Distribution of *Costus krukovii* (Maas) Maas & H.Maas (●).

**Map 24.**
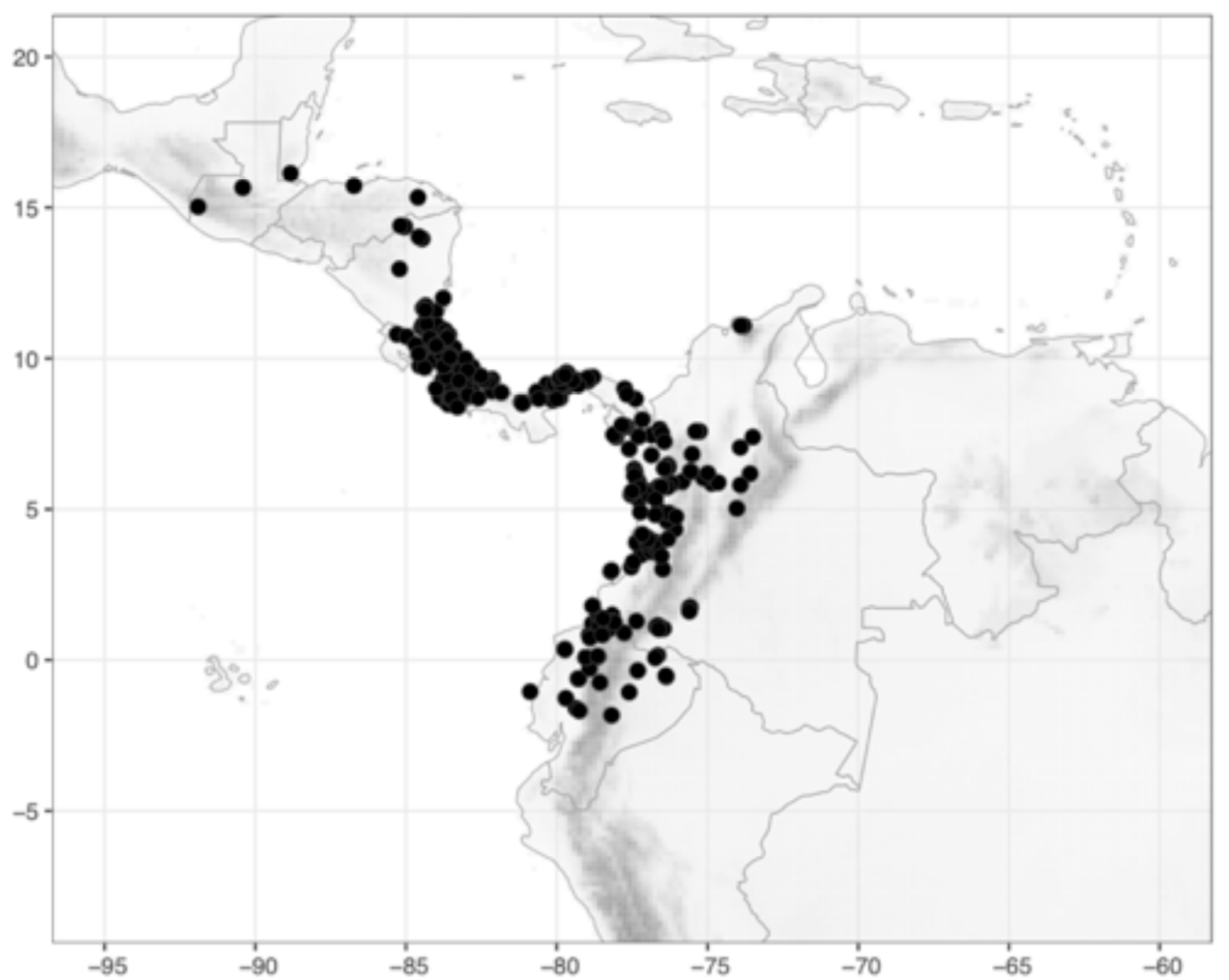
Distribution of *Costus kuntzei* K.Schum. (●).

**Map 25.**
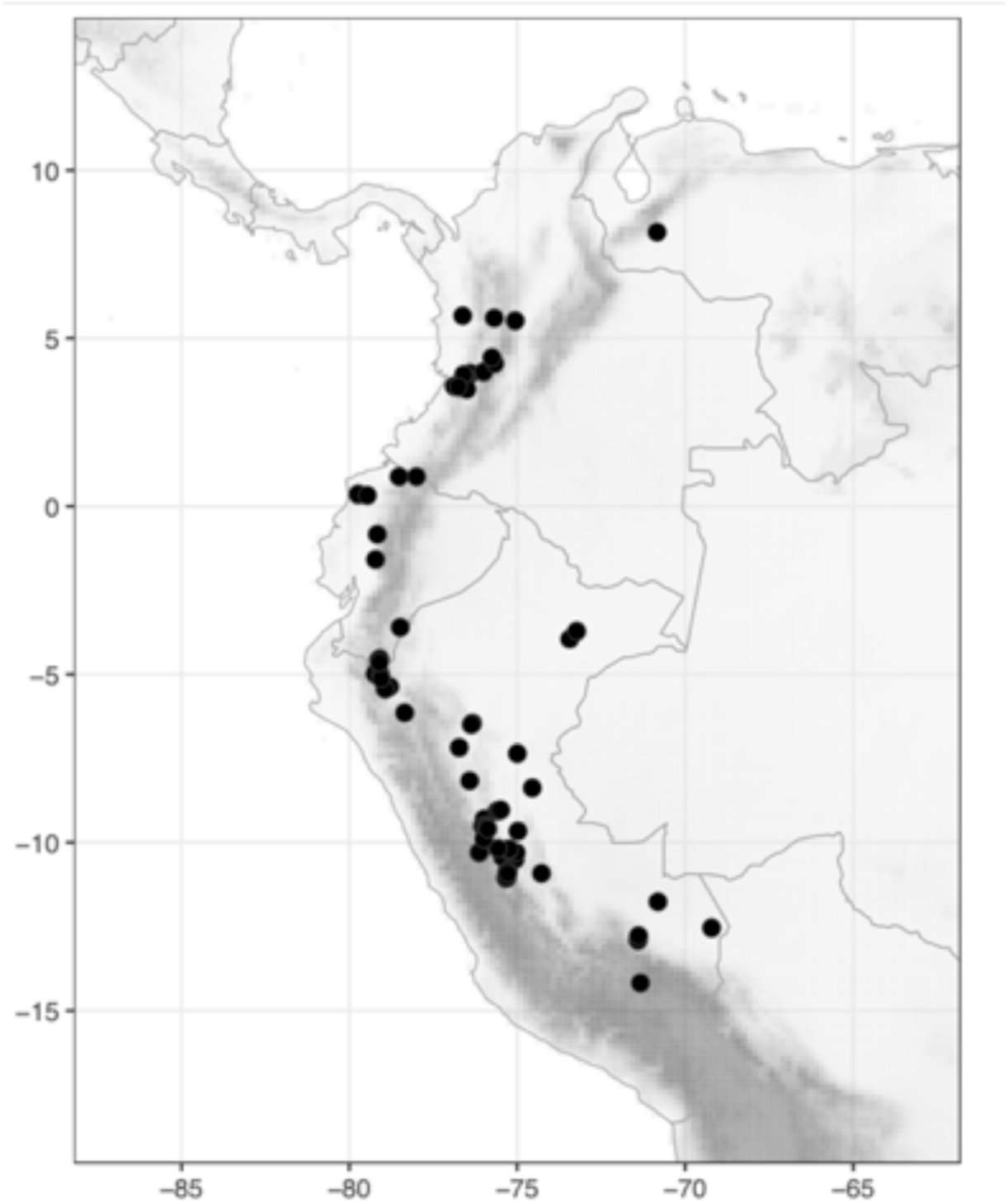
Distribution of *Costus laevis* Ruiz & Pav. (●).

**Map 26.**
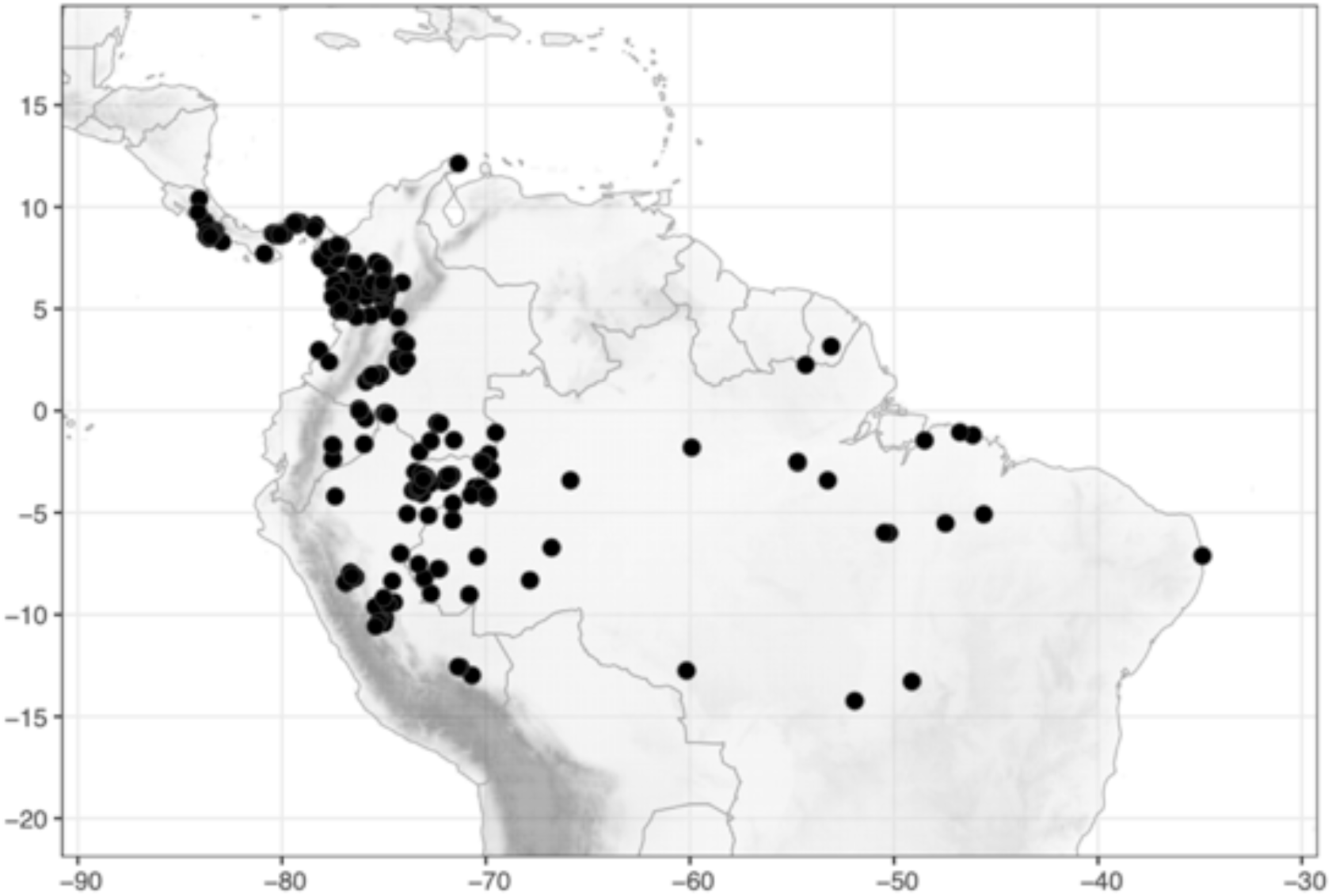
Distribution of *Costus lasius* Loes. (●).

**Map 27.**
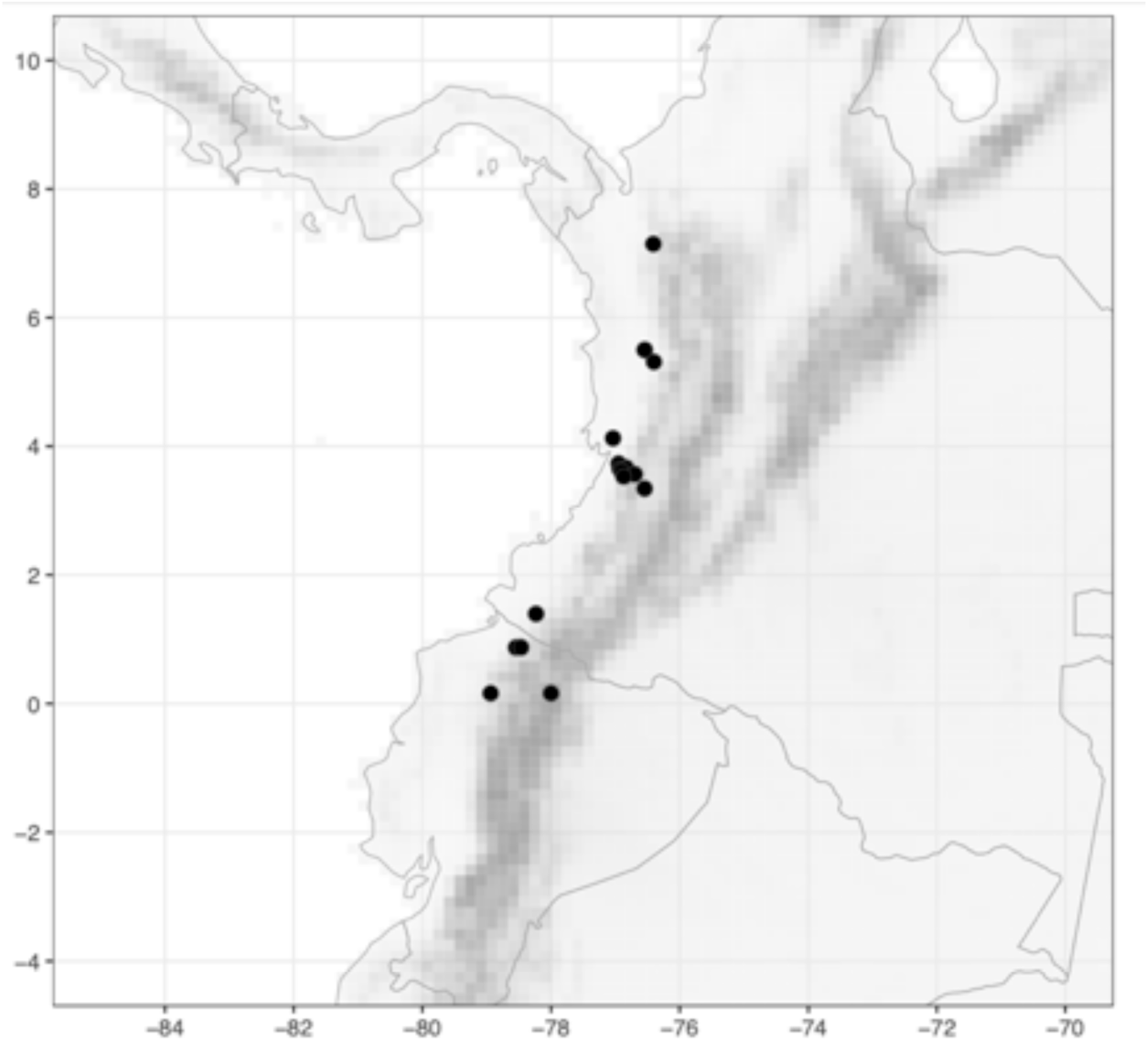
Distribution of *Costus leucanthus* Maas (●).

**Map 28.**
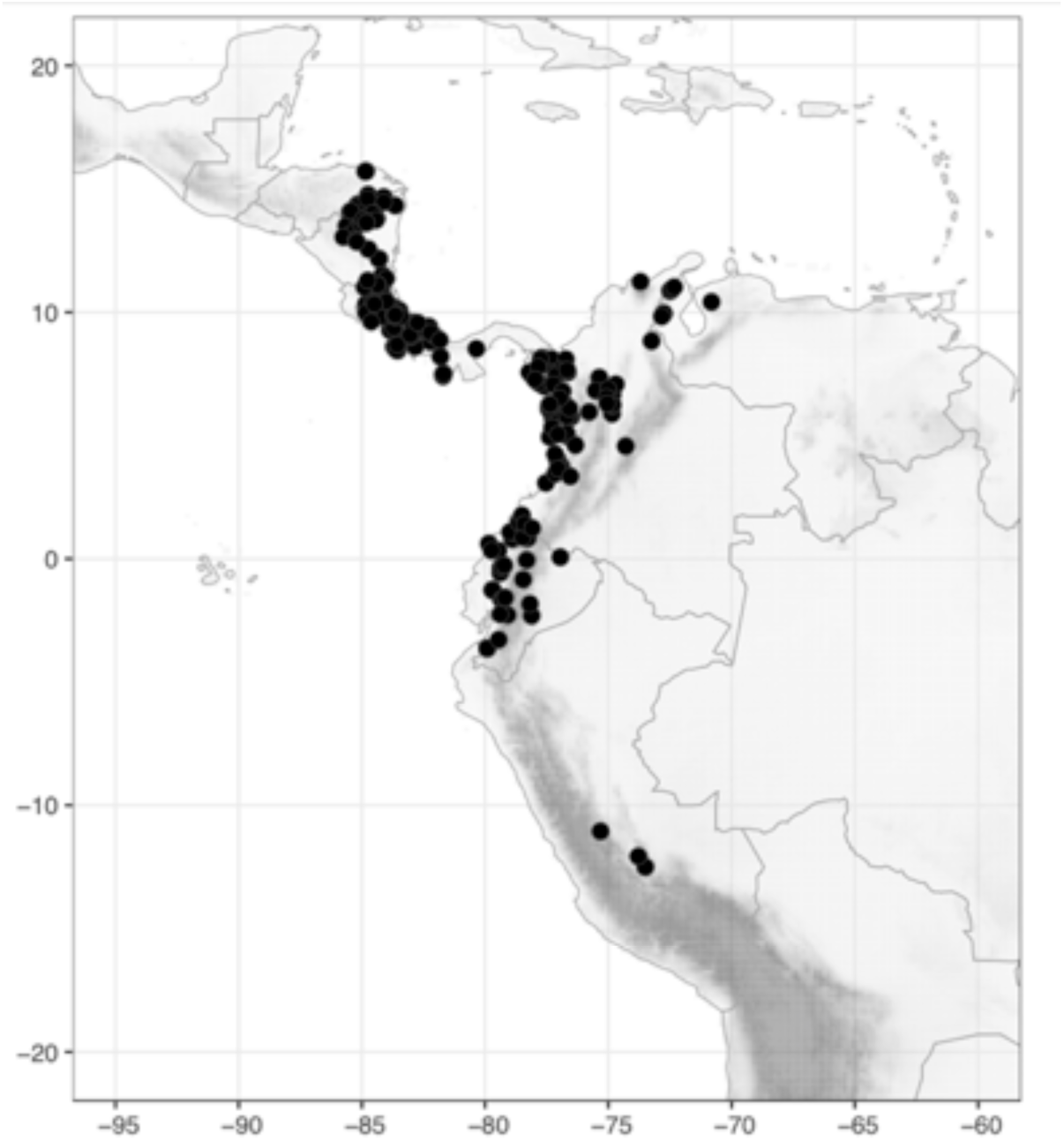
Distribution of *Costus lima* K.Schum. (●).

**Map 29.**
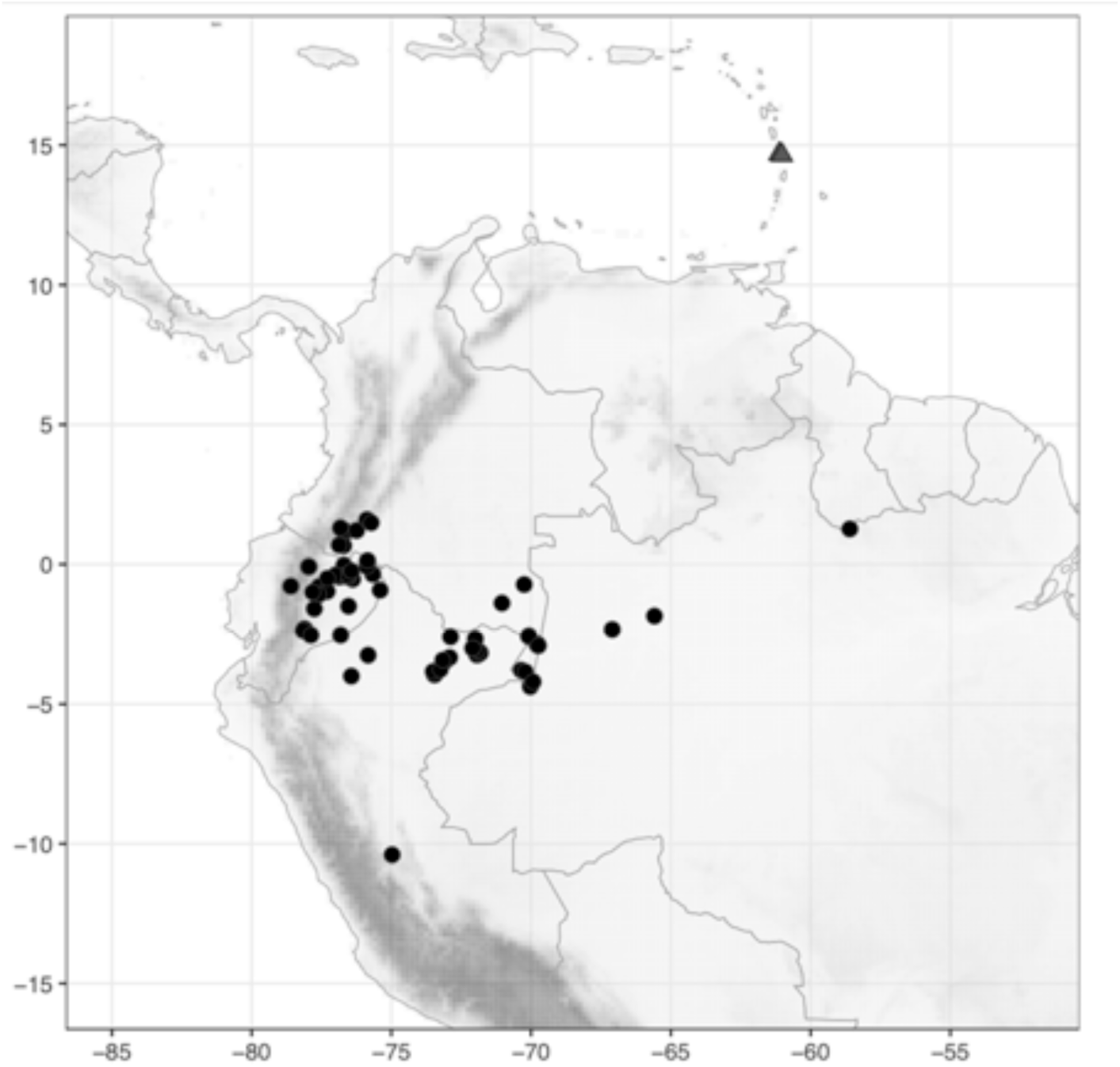
Distribution of *Costus longibracteolus* Maas (●) and *C. lucanusianus* J.Braun & K.Schum.(▲).

**Map 30.**
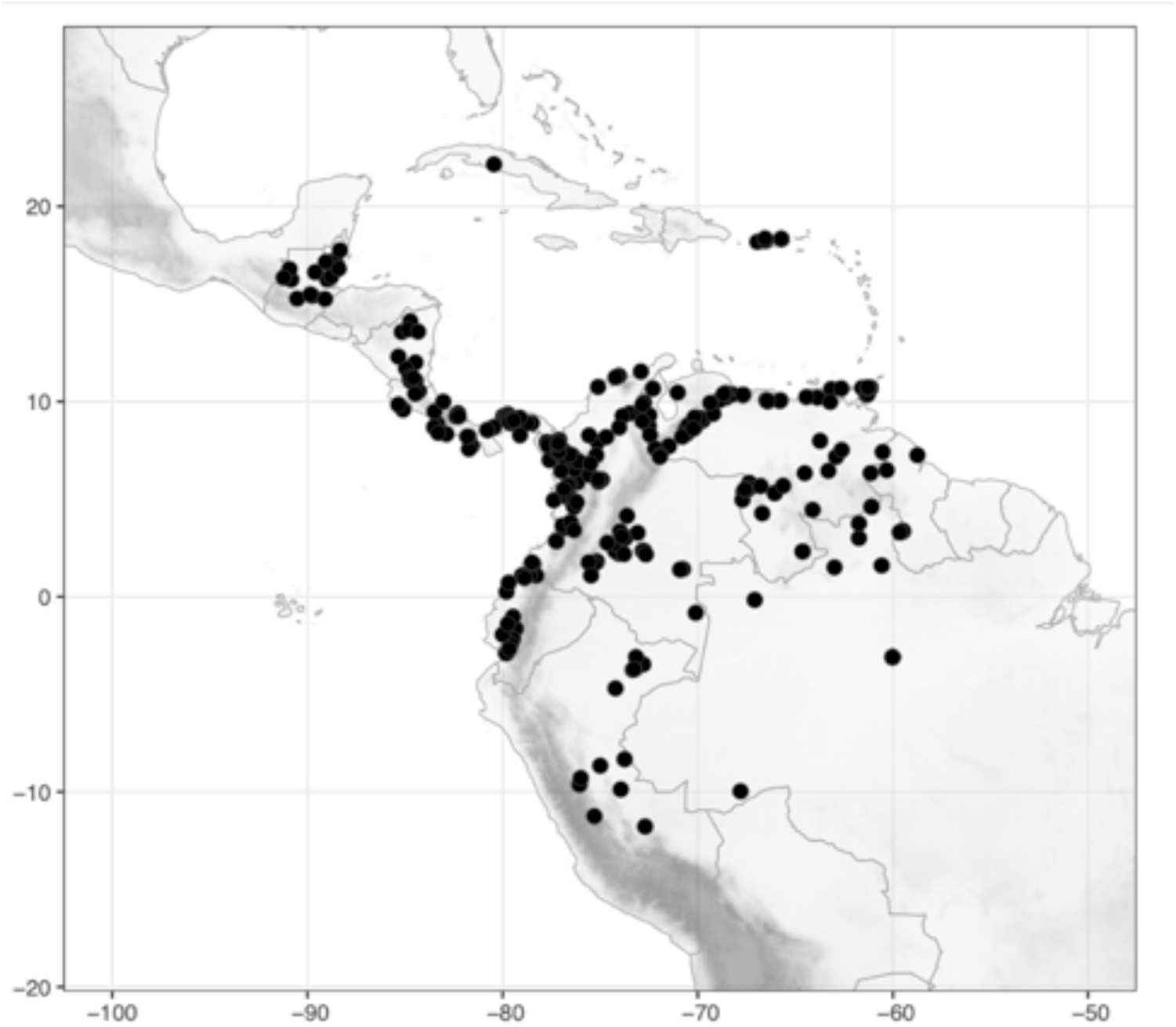
Distribution of *Costus macrostrobilus* K.Schum. (●).

**Map 31.**
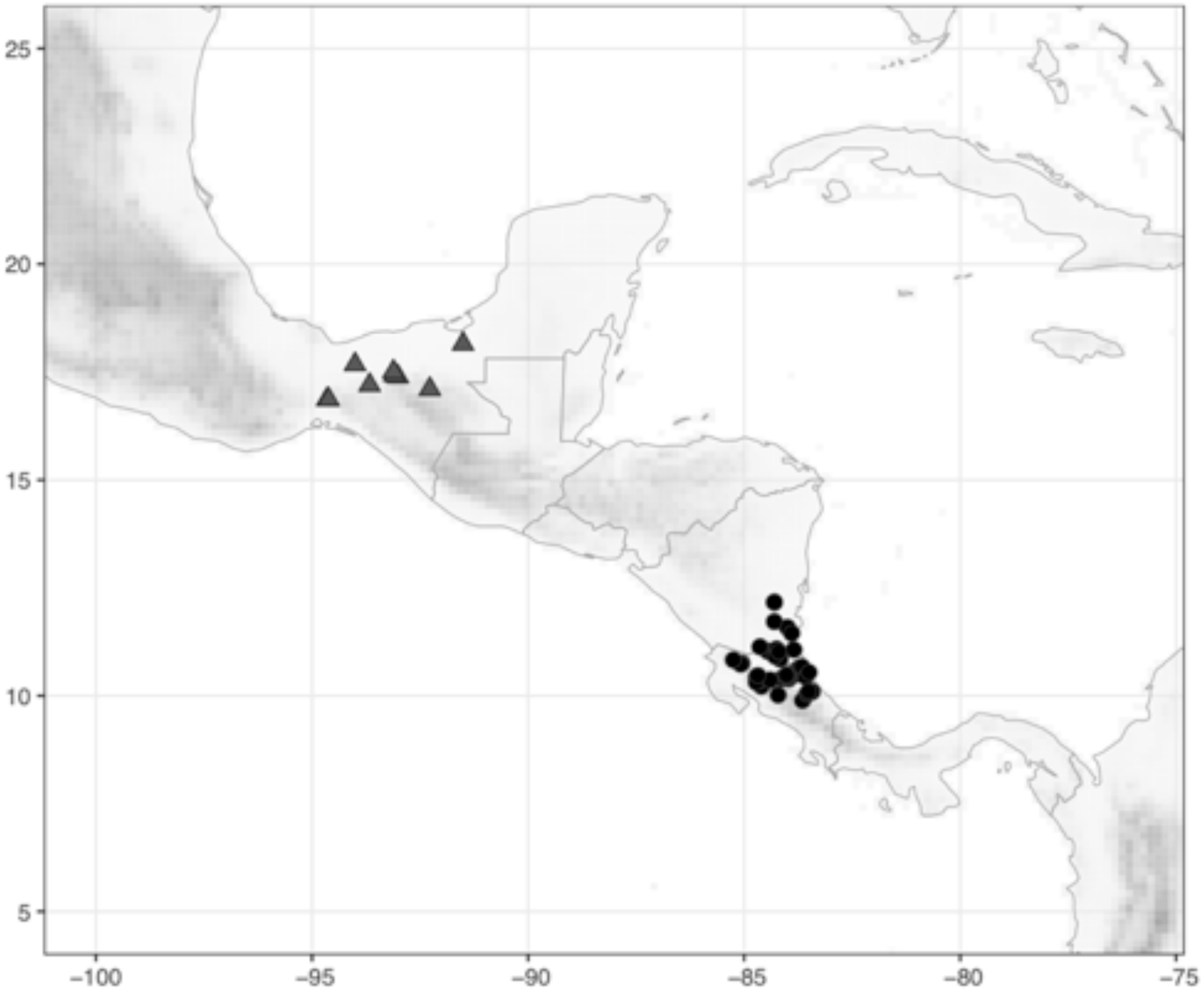
Distribution of *Costus malortieanus* H.Wendl. (●) and *C. mollisimus* Maas & H.Maas (▲).

**Map 32.**
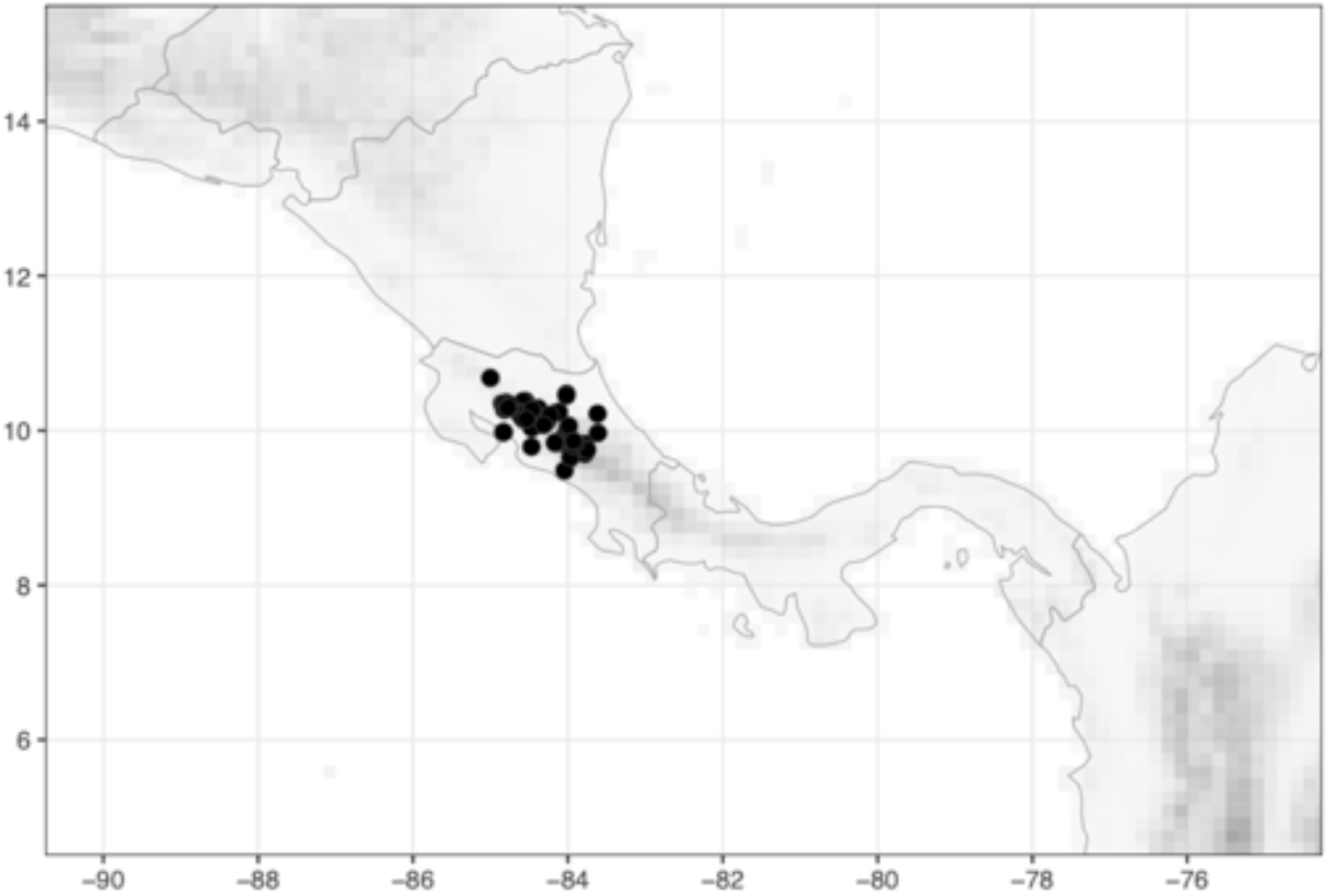
Distribution of *Costus montanus* Maas (●).

**Map 33.**
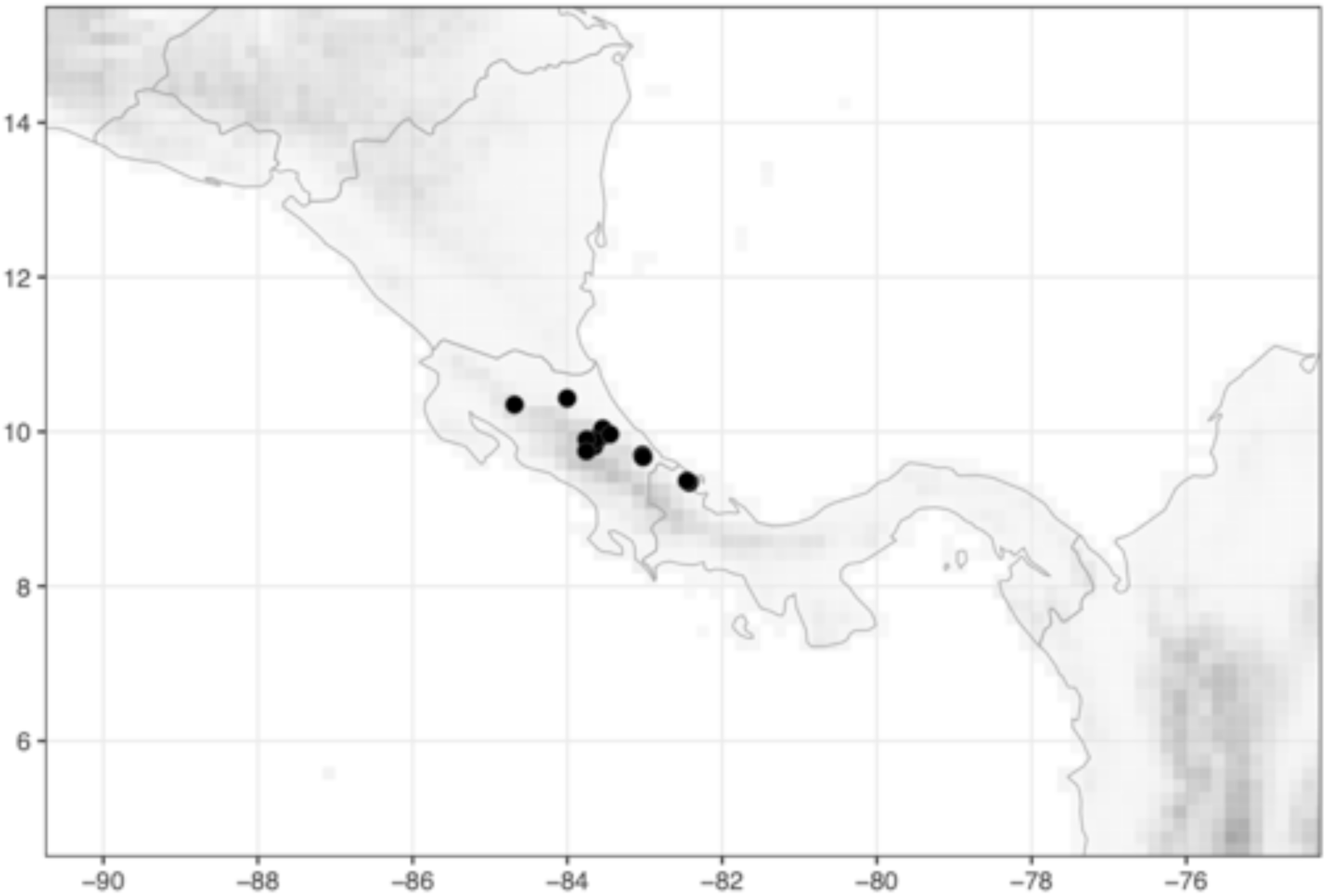
Distribution of *Costus nitidus* Maas (●).

**Map 34.**
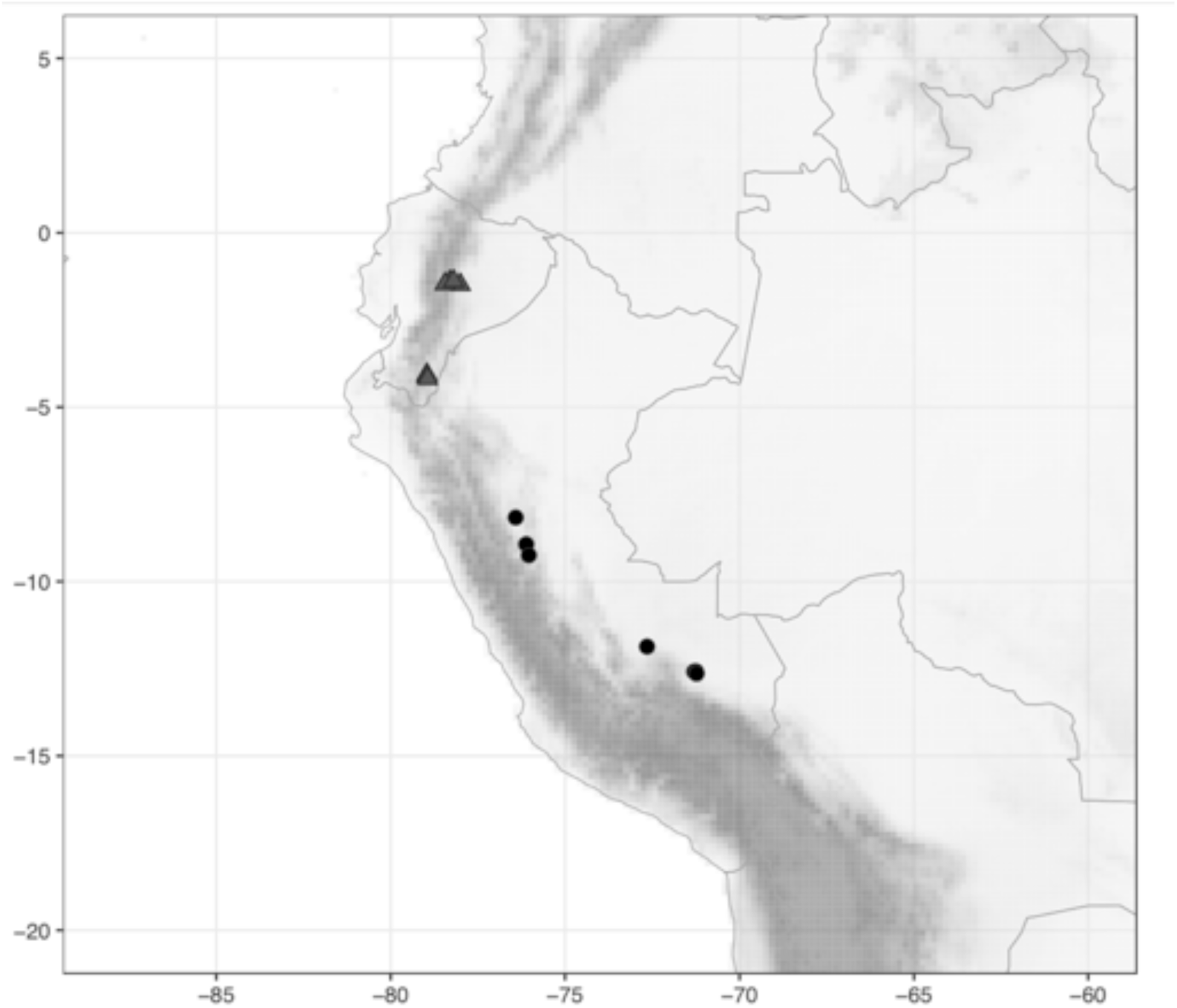
Distribution of *Costus obscurus* D.Skinner & Maas (●) and *C. oreophilus* Maas & D.Skinner (▲).

**Map 35.**
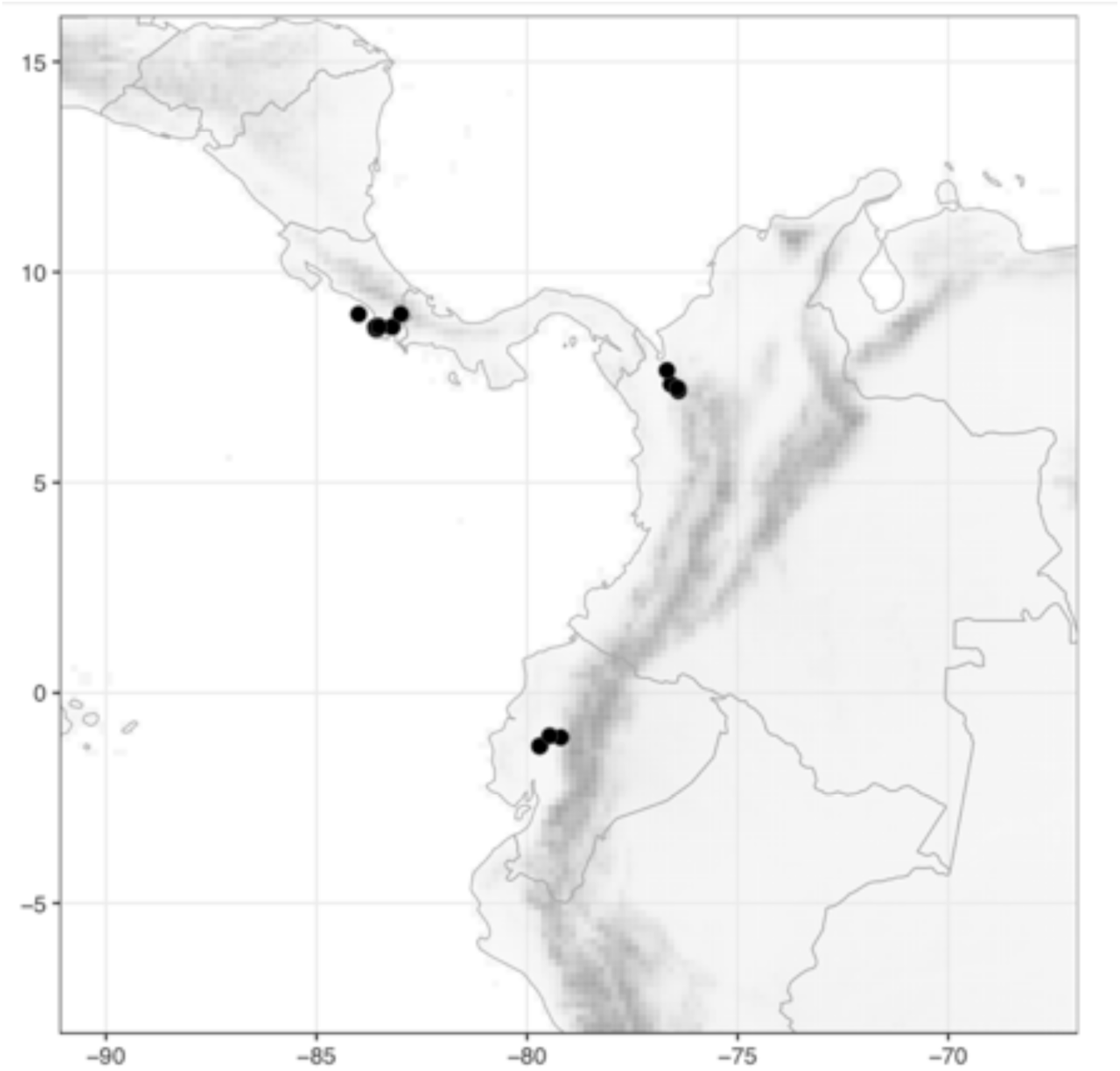
Distribution of *Costus osae* Maas & H.Maas (●).

**Map 36.**
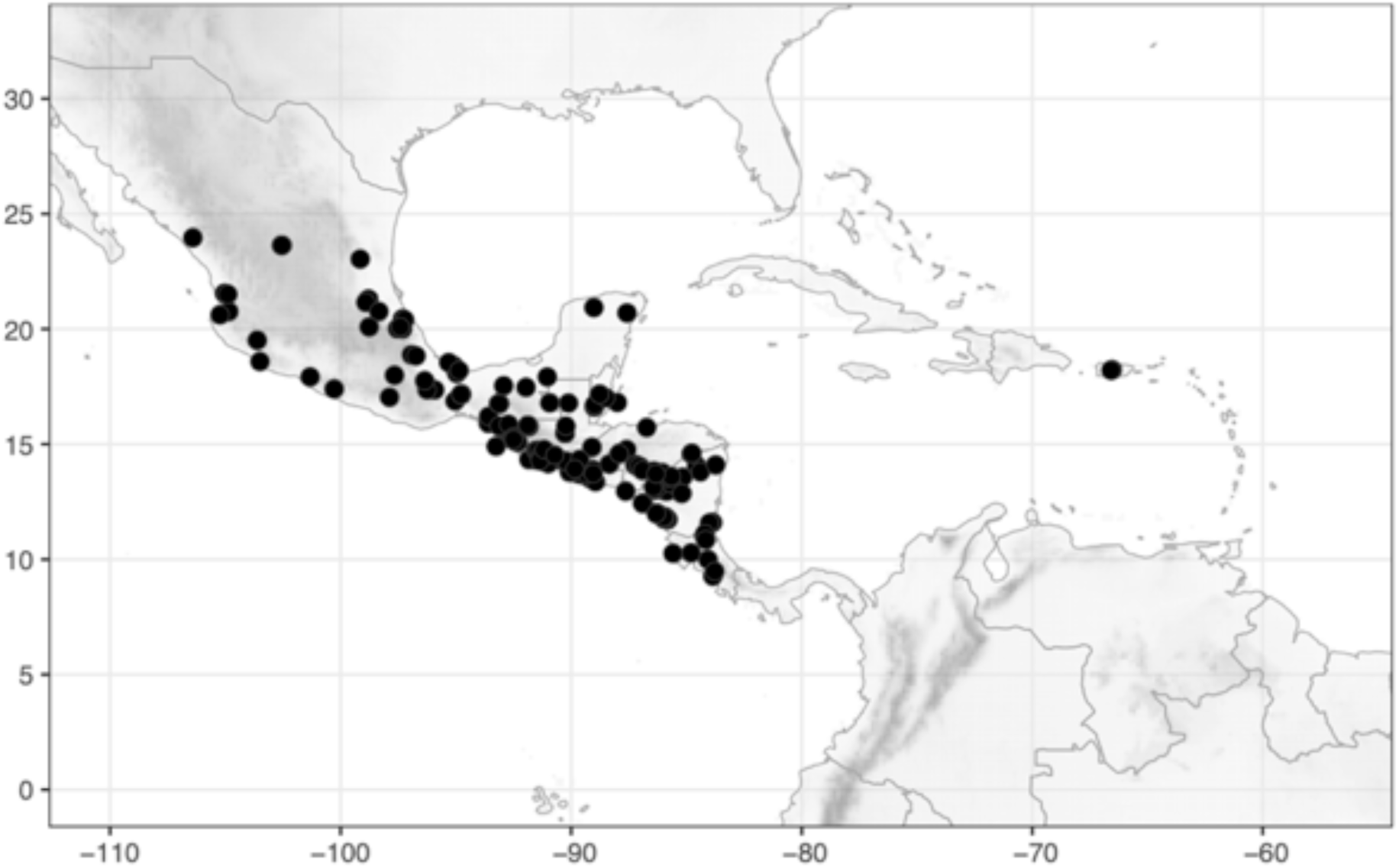
Distribution of *Costus pictus* D.Don (●).

**Map 37.**
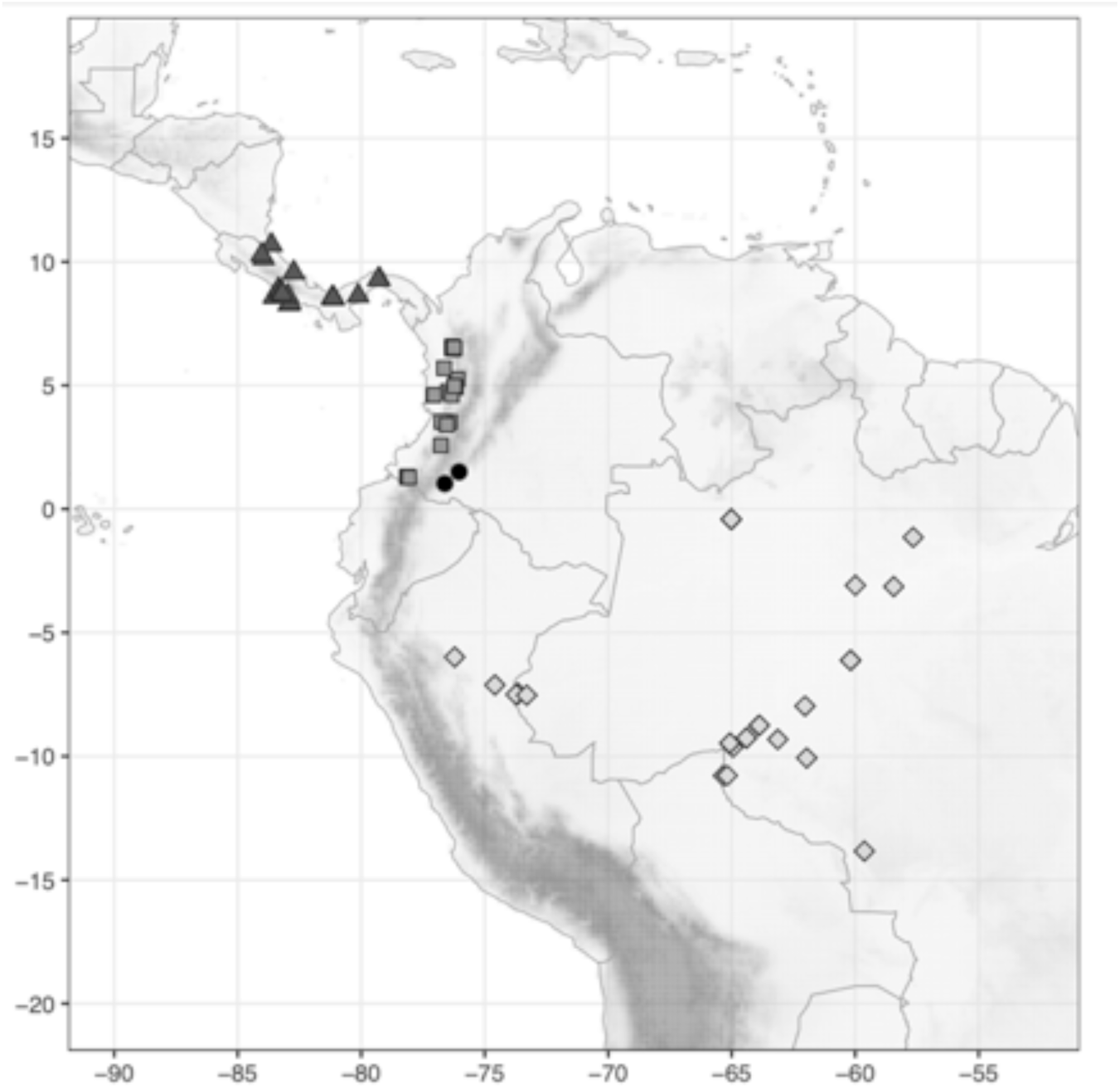
Distribution of *Costus pitalito* C.D.Specht & H.Maas (●), *C. plicatus* Maas (▲), *C. plowmanii* Maas (◼), and *C. prancei* Maas & H.Maas (◆).

**Map 38.**
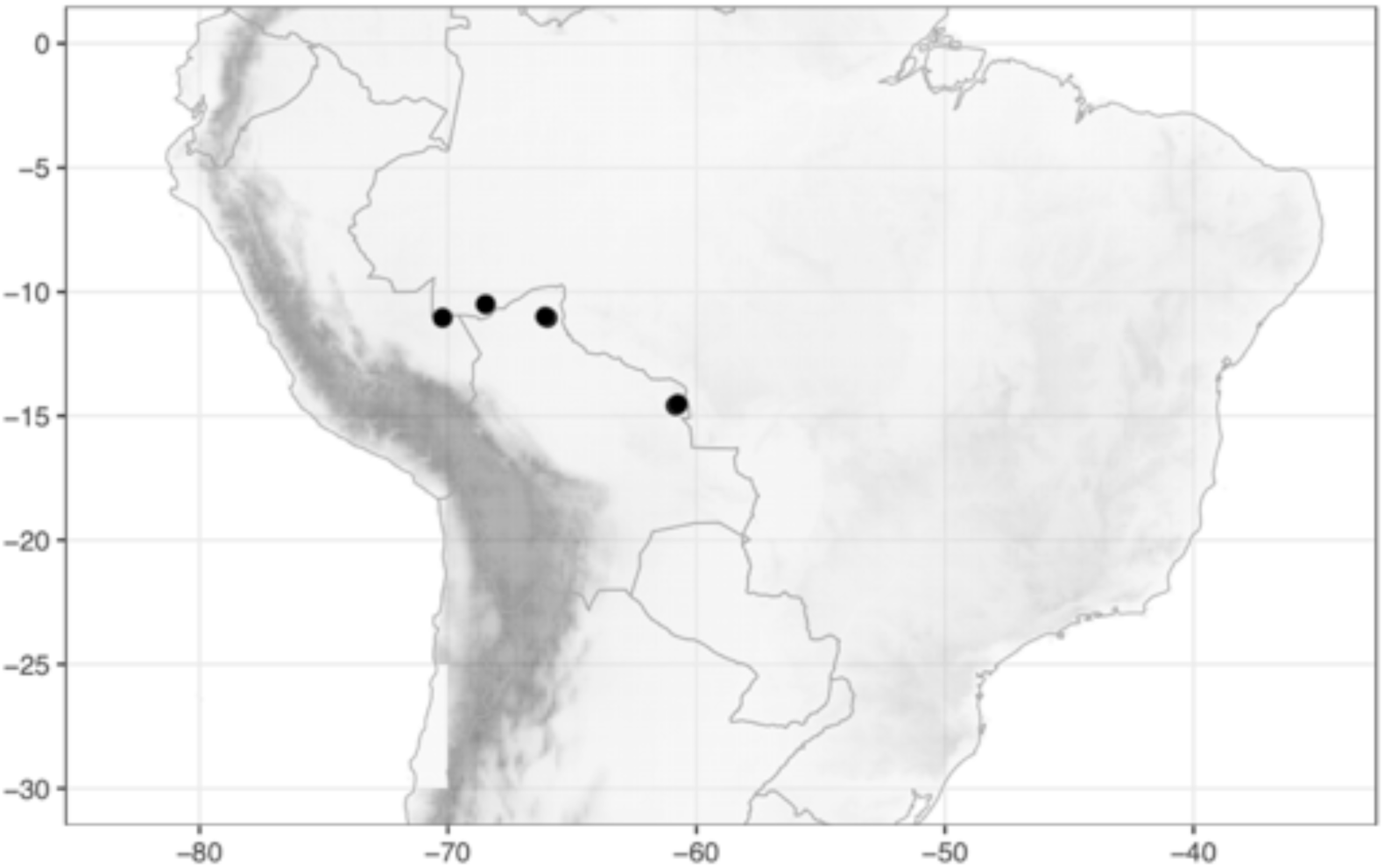
Distribution of *Costus pseudospiralis* Maas & H. Maas (●).

**Map 39.**
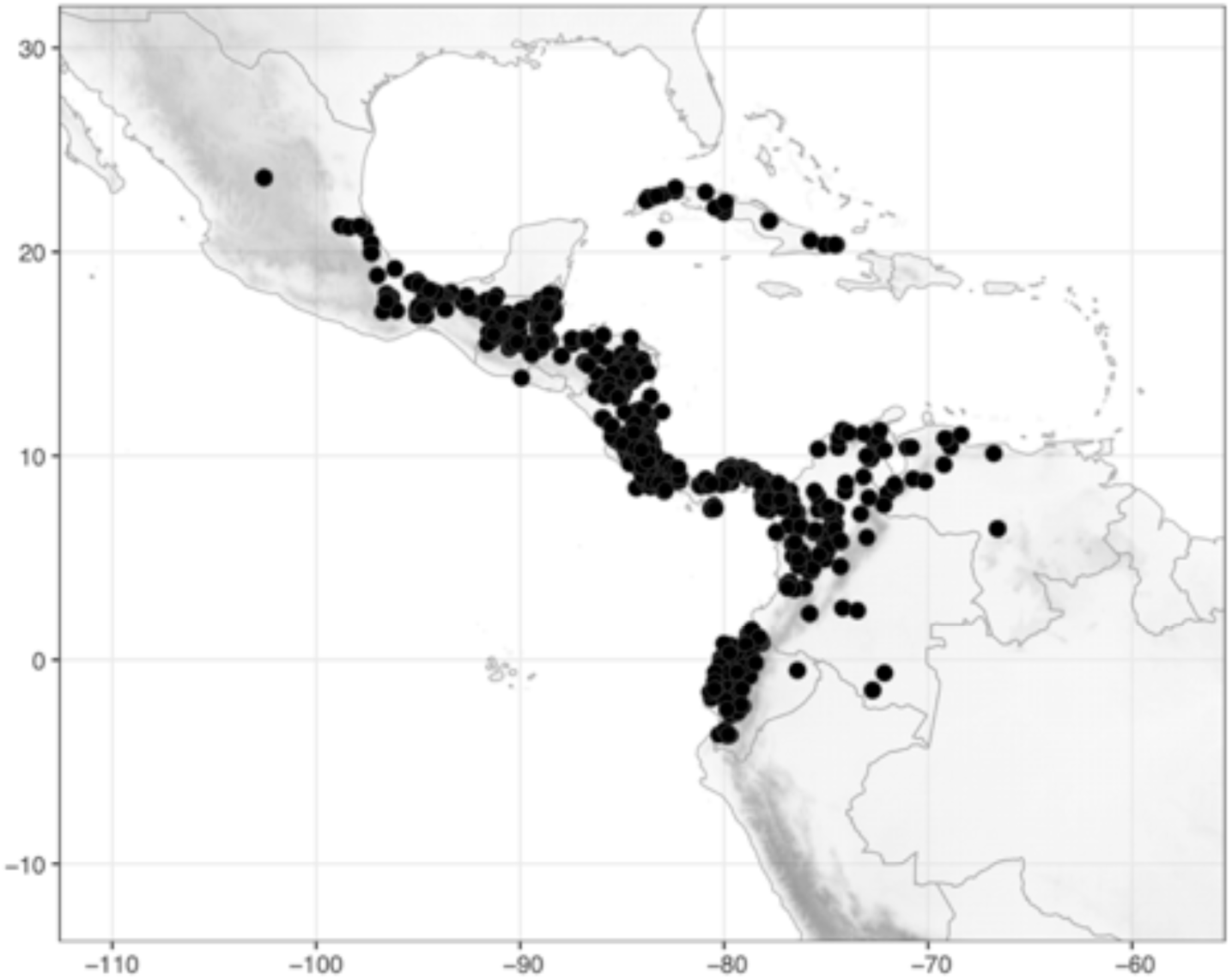
Distribution of *Costus pulverulentus* C.Presl (●).

**Map 40.**
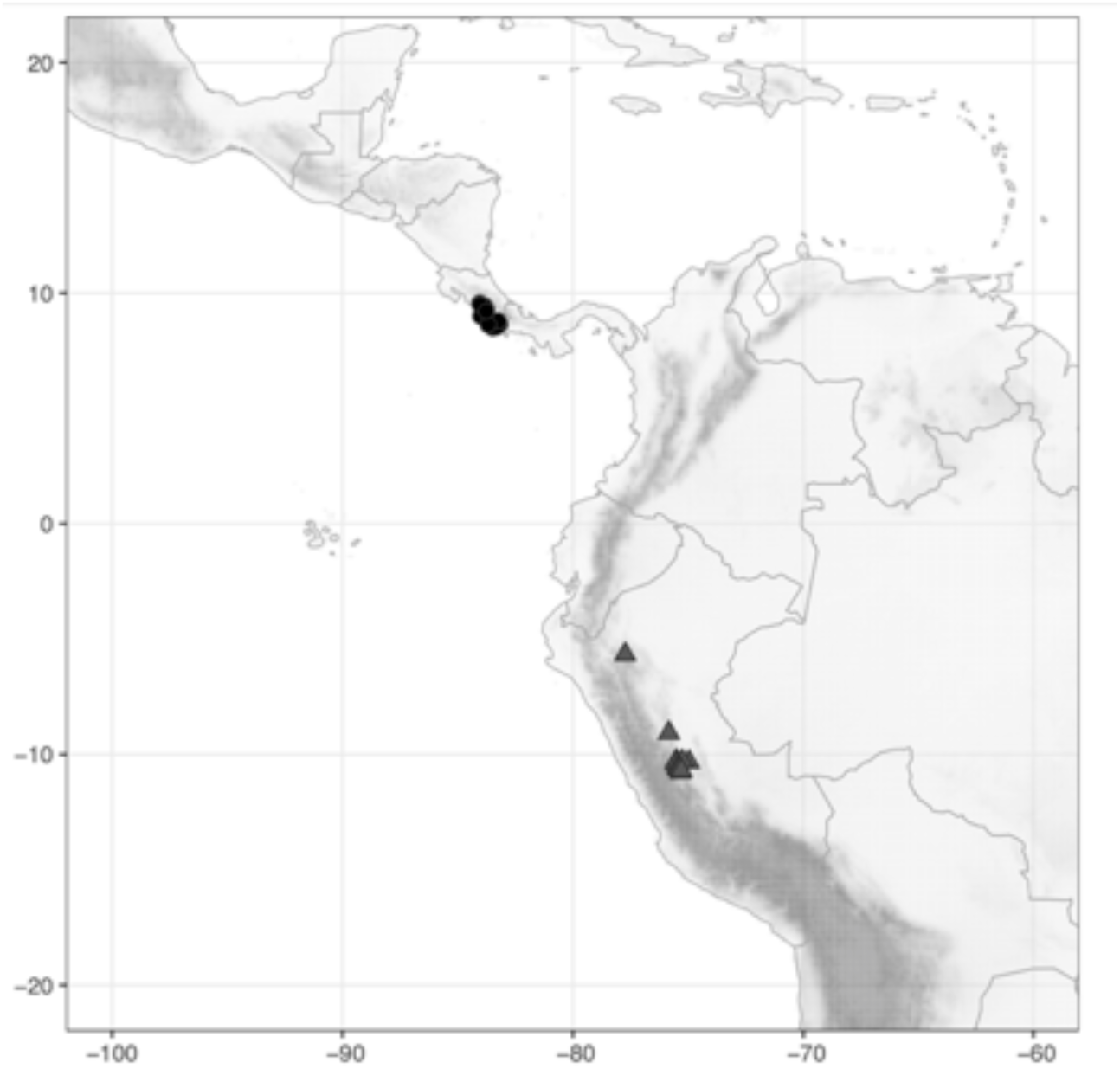
Distribution of *Costus ricus* Maas & H.Maas (●) and *C. rubineus* D.Skinner & Maas (▲).

**Map 41.**
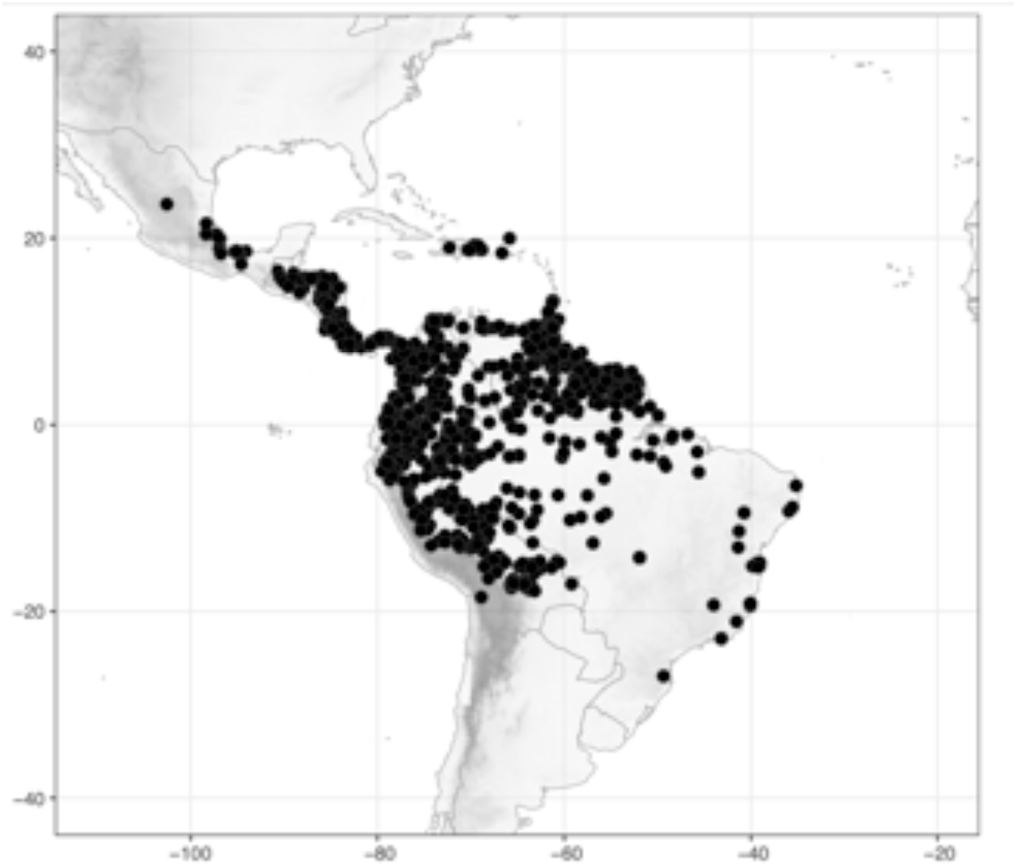
Distribution of *Costus scaber* Ruiz & Pav. (●).

**Map 42.**
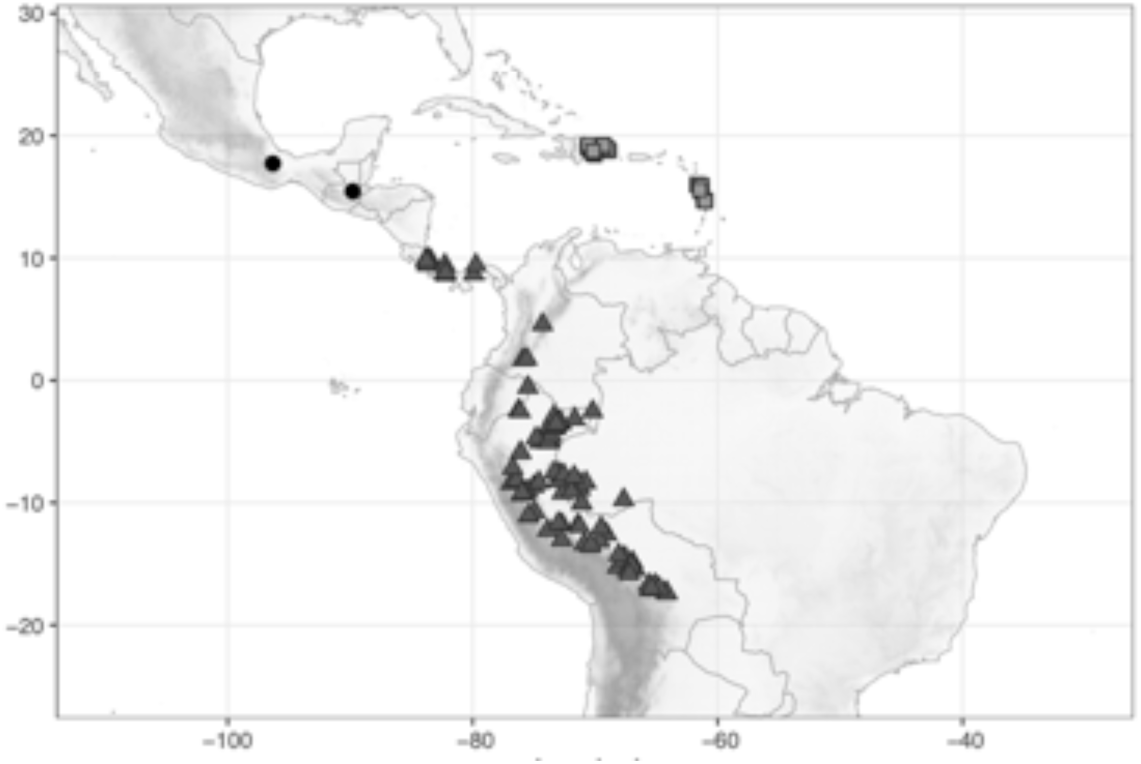
Distribution of *Costus sepacuitensis* Rowlee (●), *C. sinningiflorus* Rusby (▲), and *C. spicatus* (Jacq.) Sw. (◼).

**Map 43.**
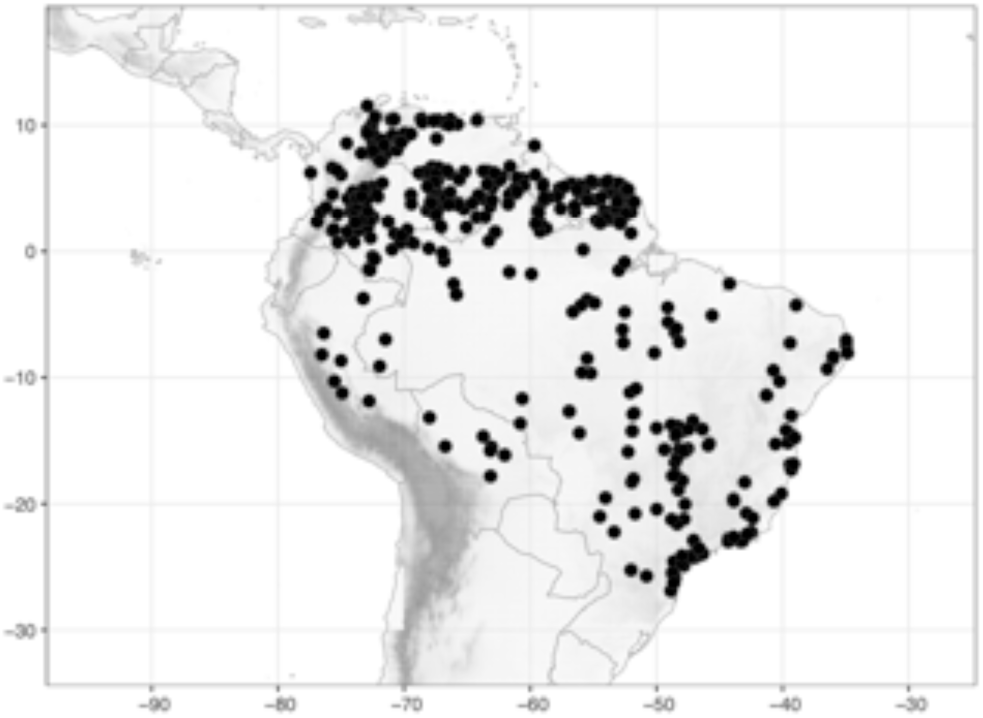
Distribution of *Costus spiralis* (Jacq.) Roscoe (●).

**Map 44.**
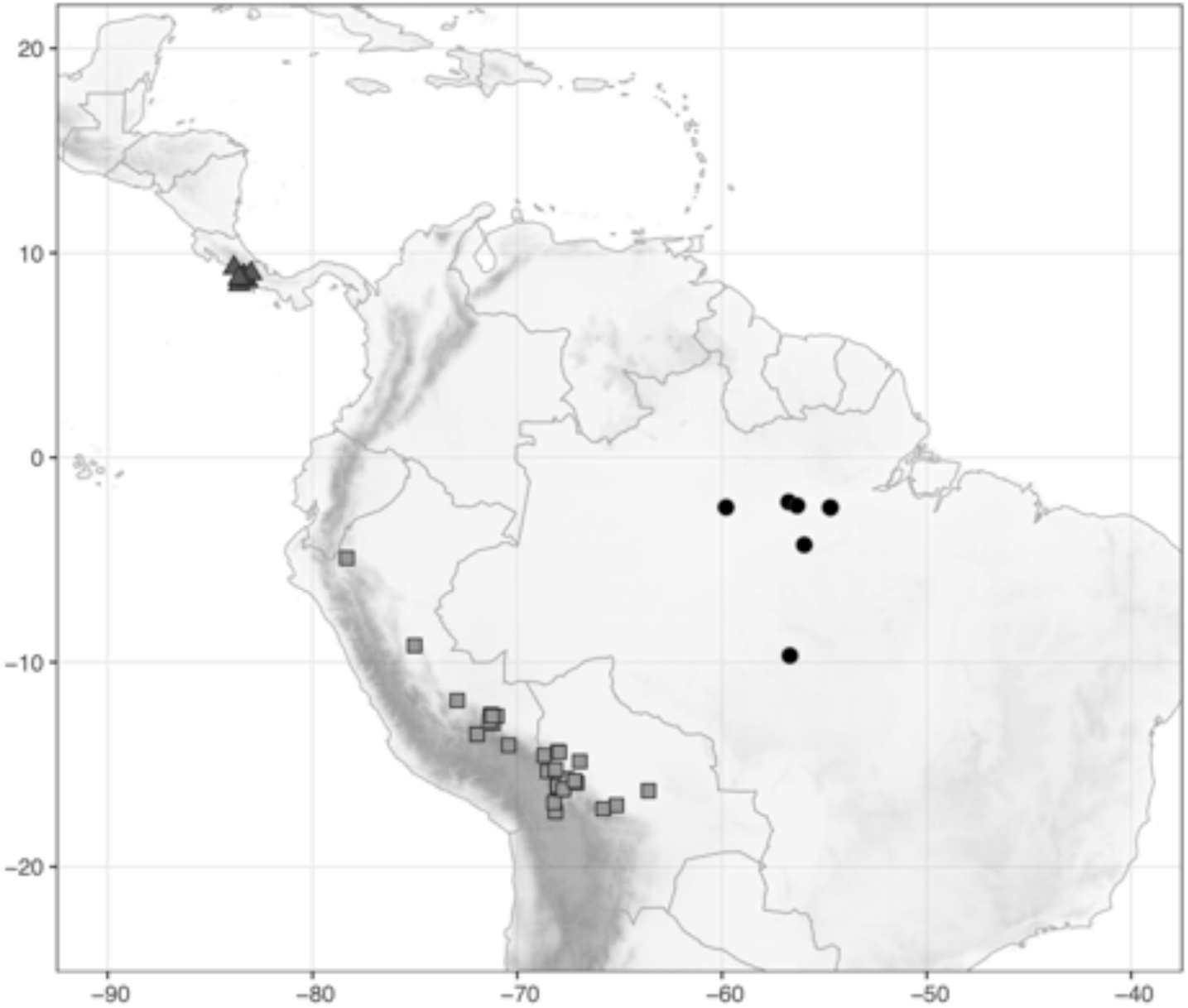
Distribution of *Costus sprucei* Maas (●), *C. stenophyllus* Standl. & L.O.Williams (▲), and *C. vargasii* Maas & H.Maas (◼).

**Map 45.**
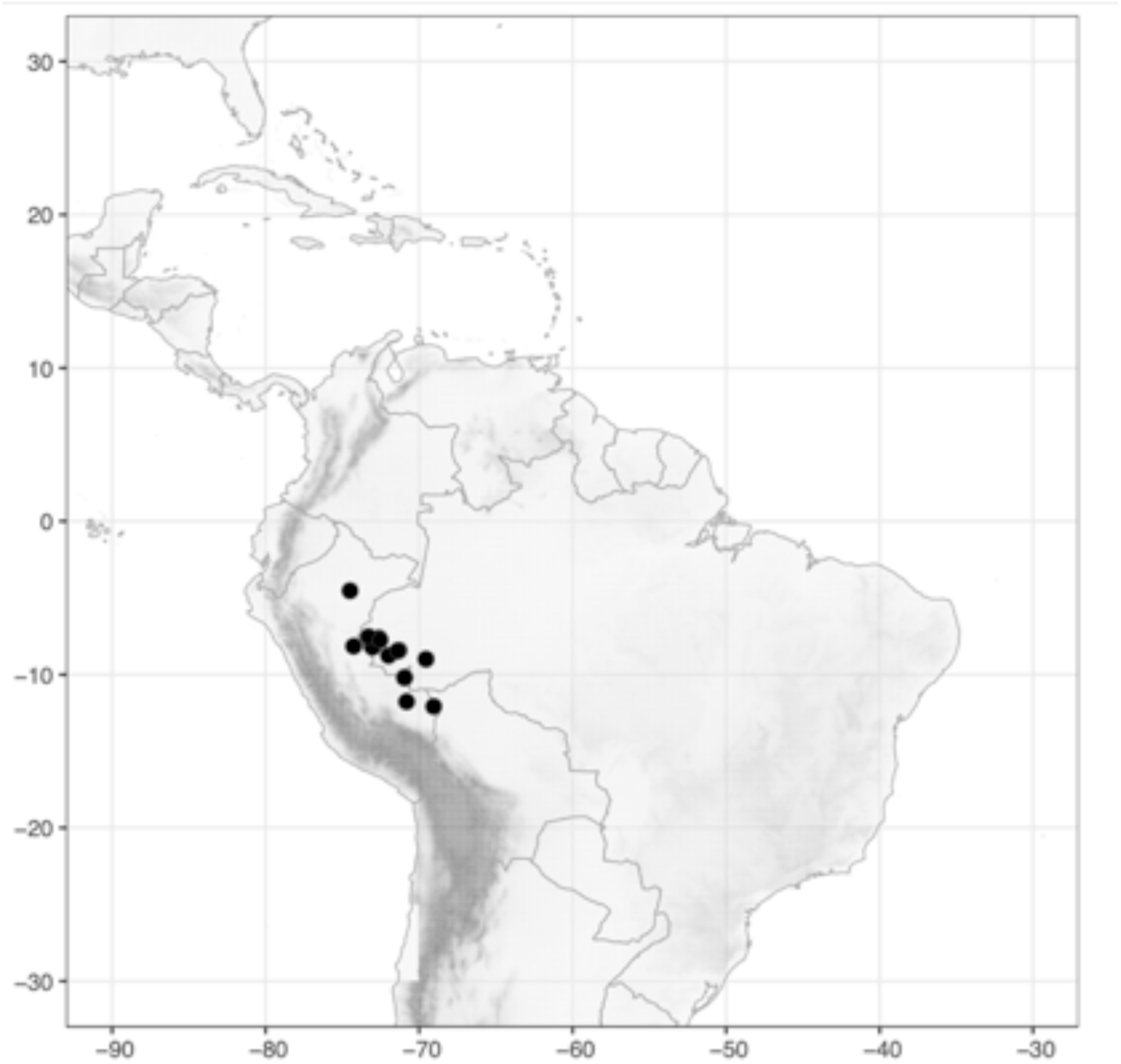
Distribution of *Costus varzearum* Maas (●).

**Map 46.**
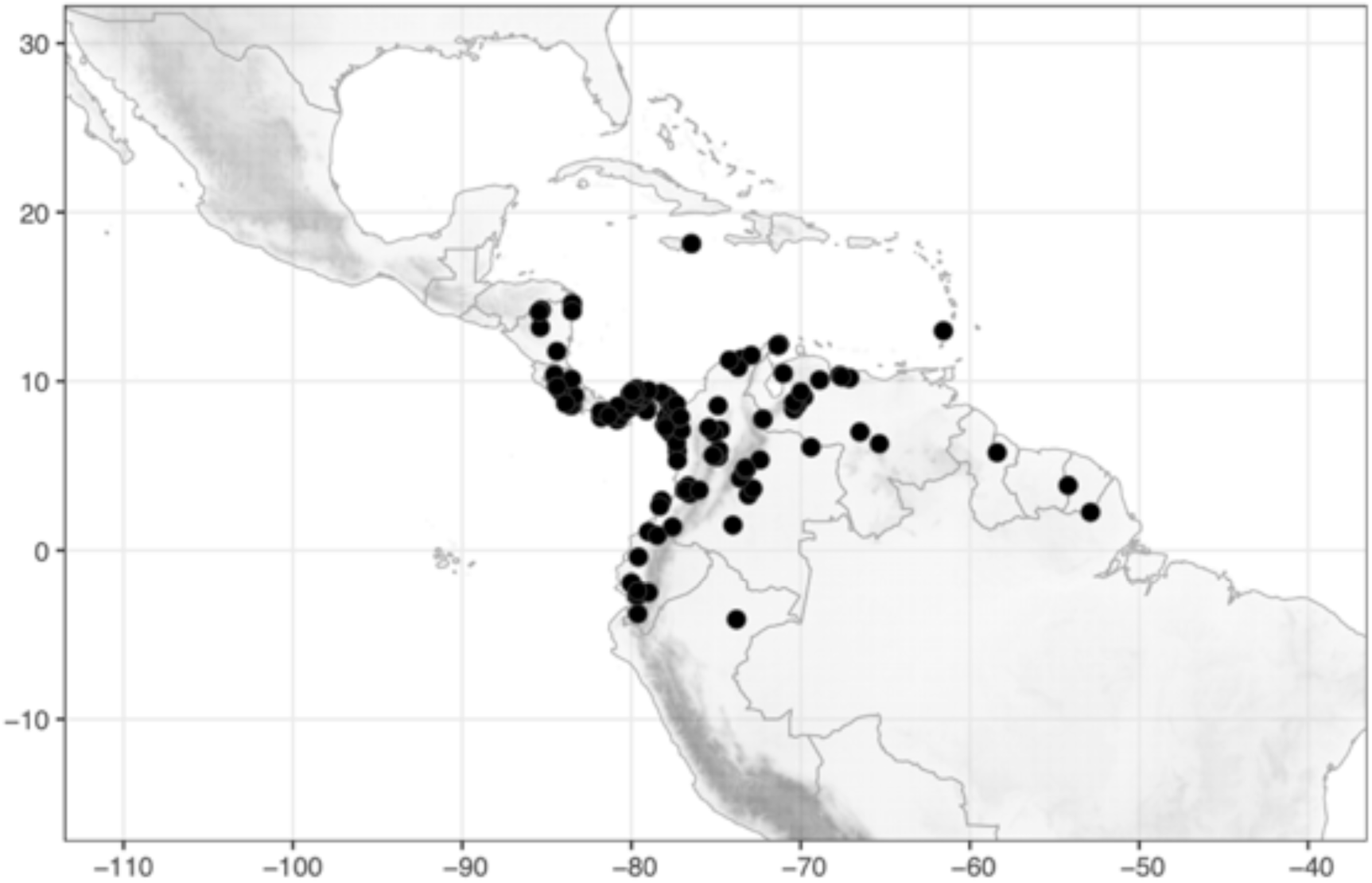
Distribution of *Costus villosissimus* Jacq. (●).

**Map 47.**
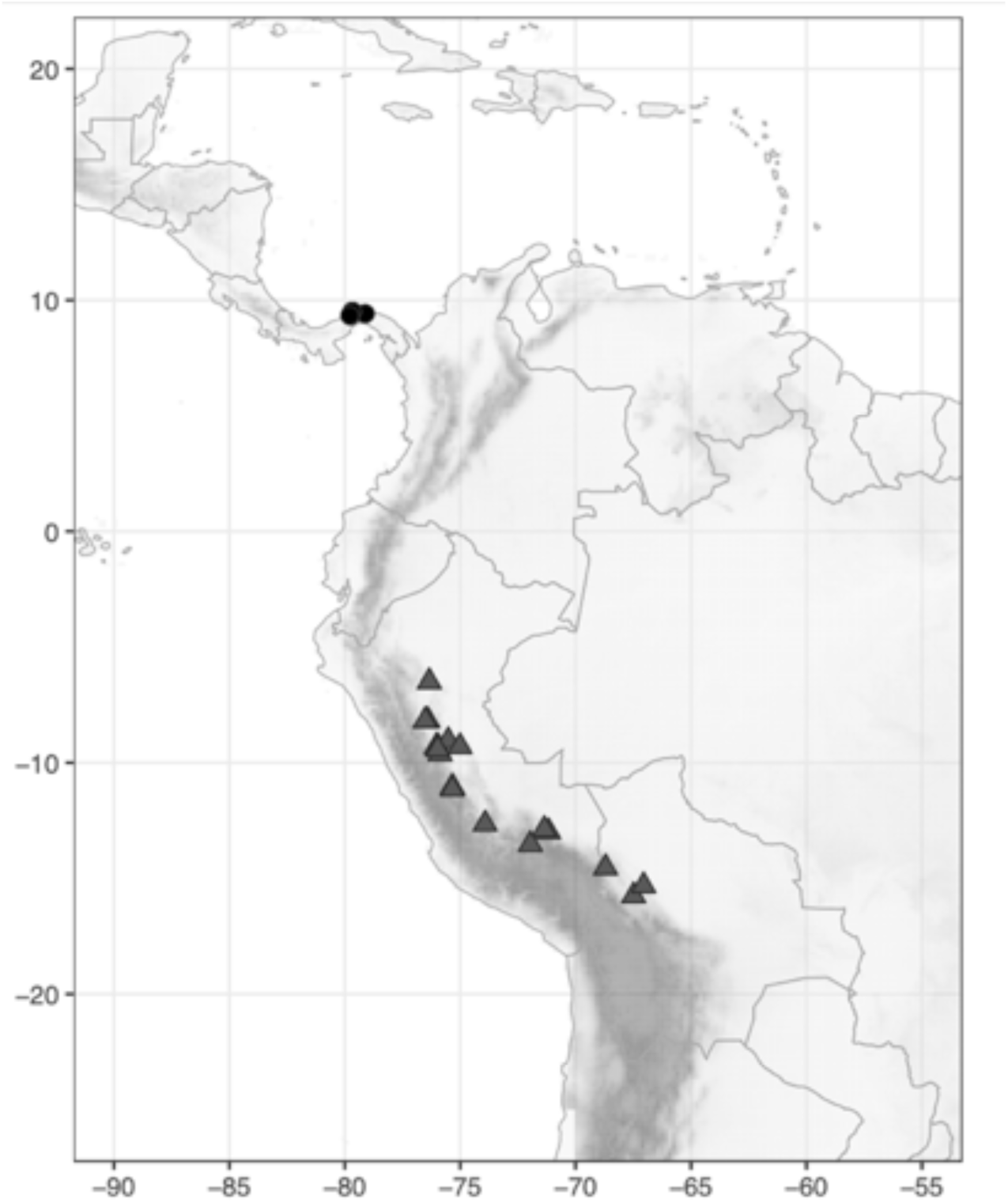
Distribution of *Costus vinosus* Maas (●) and *C. weberbaueri* Loes. (▲).

**Map 48.**
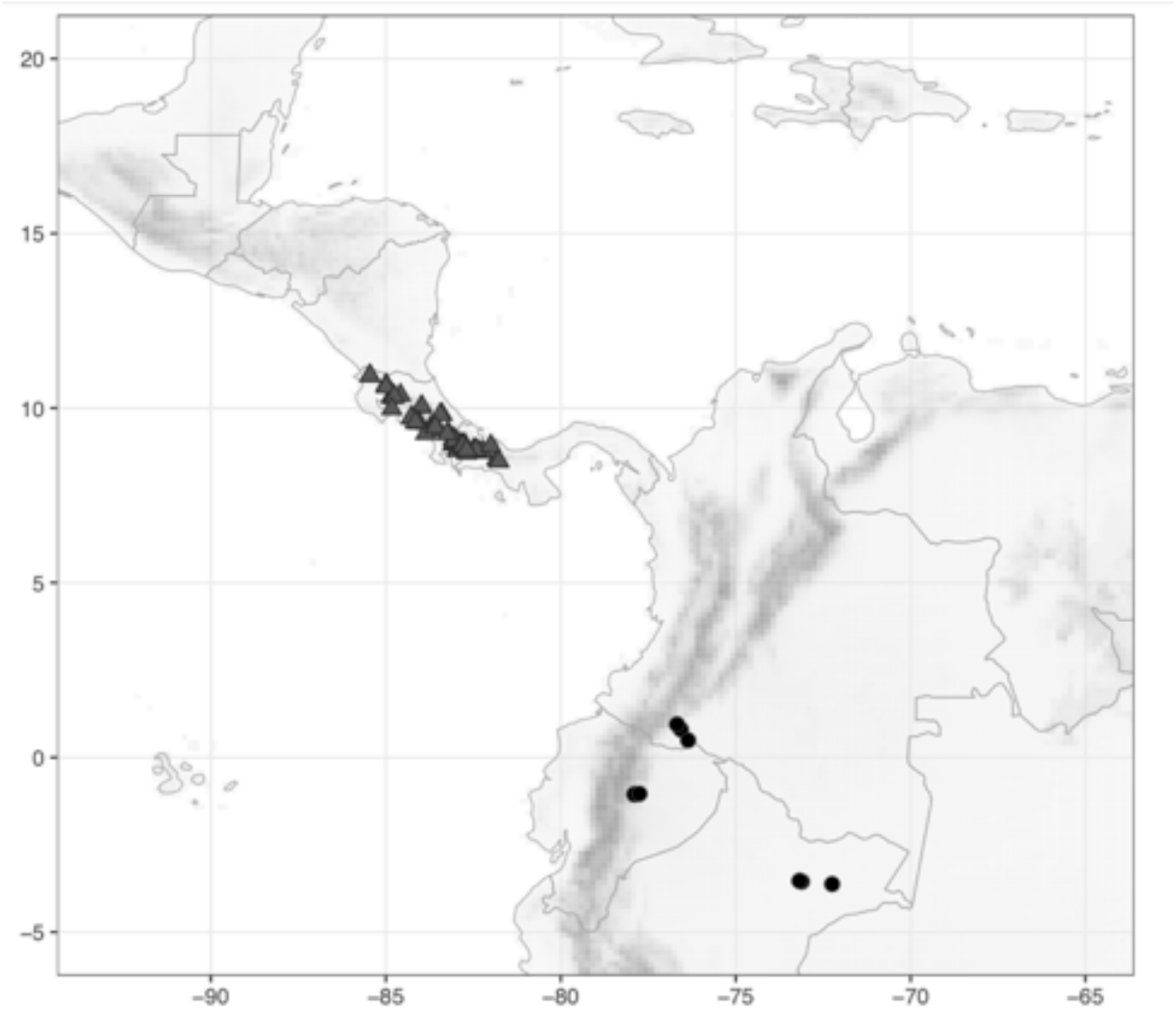
Distribution of *Costus whiskeycola* Maas & H.Maas (●) and *C. wilsonii* Maas (▲).

**Map 49.**
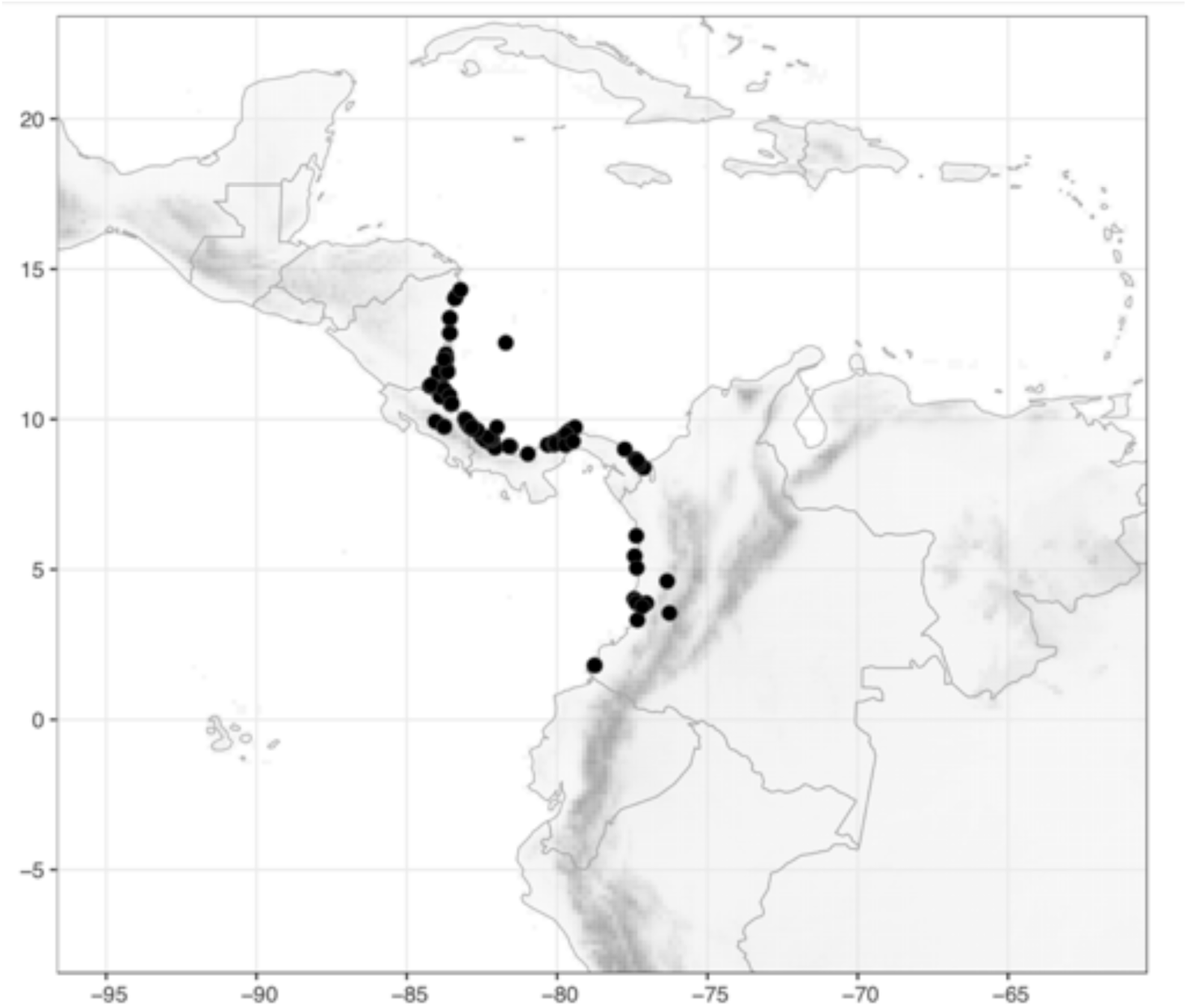
Distribution of *Costus woodsonii* Maas (●).

**Map 50.**
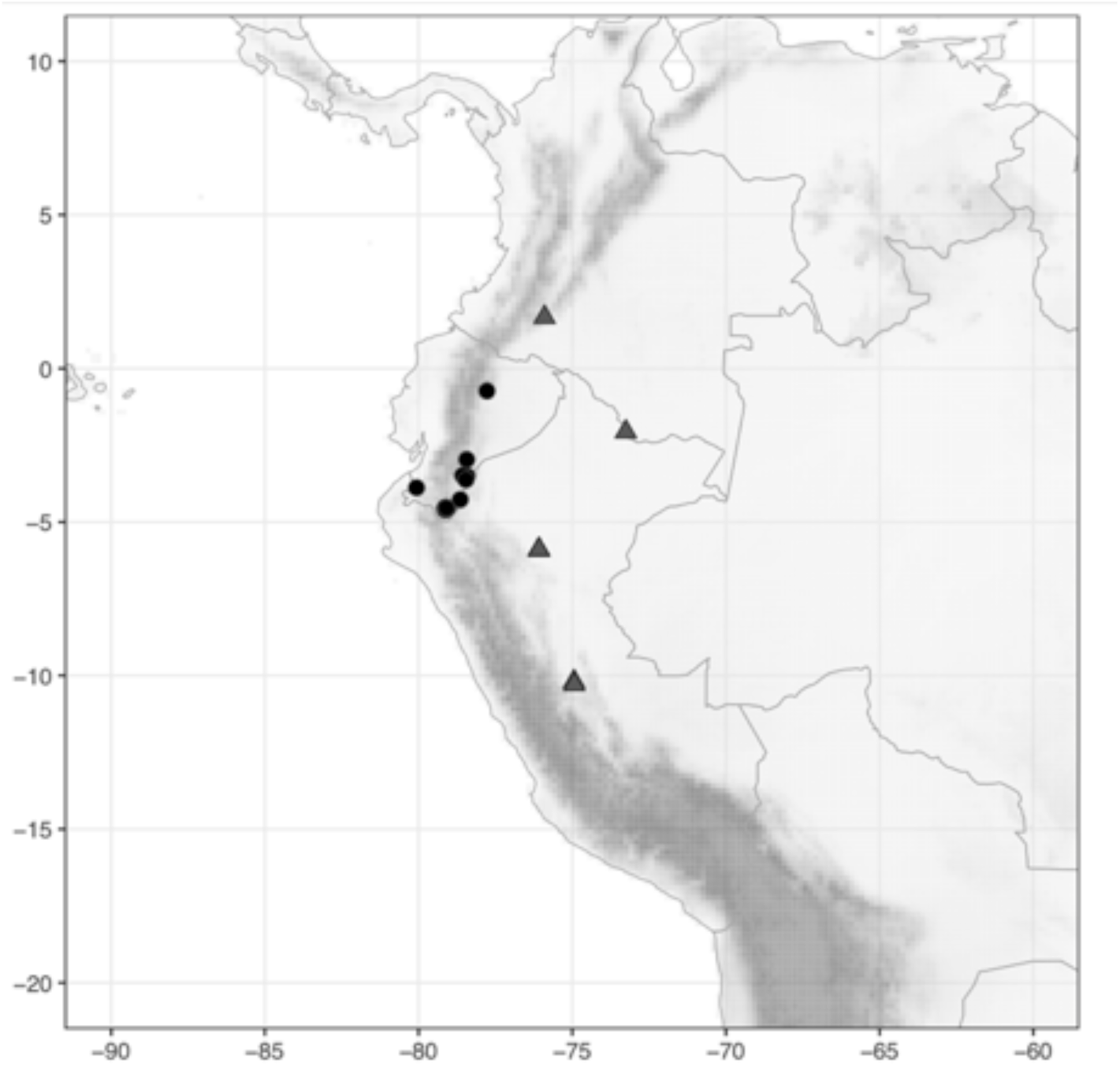
Distribution of *Costus zamoranus* Steyerm. (●) and *C. zingiberoides* J.F.Macbr. (▲).

**Map 51.**
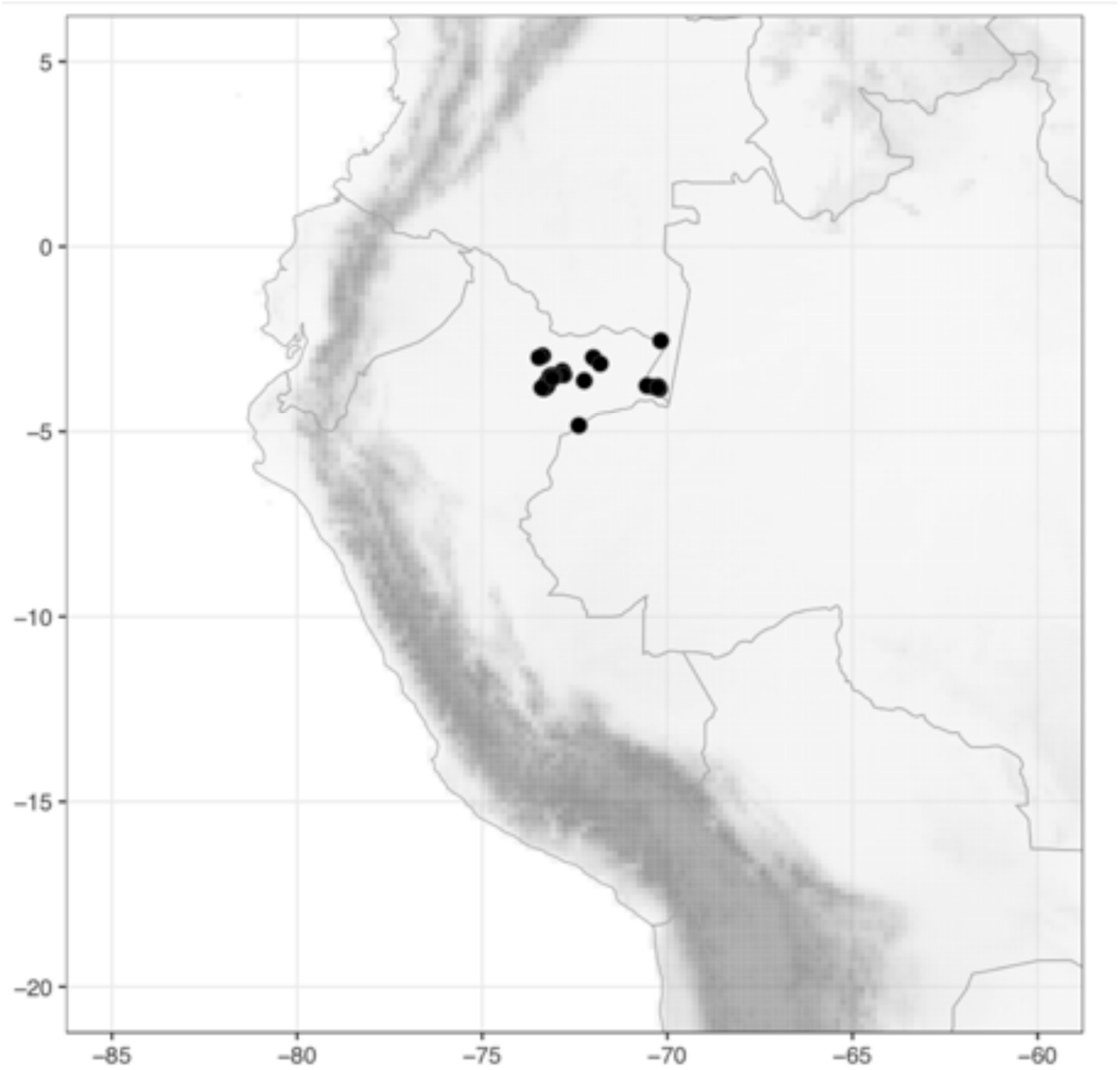
Distribution of *Dimerocostus appendiculatus* (Maas) Maas & H.Maas (●).

**Map 52.**
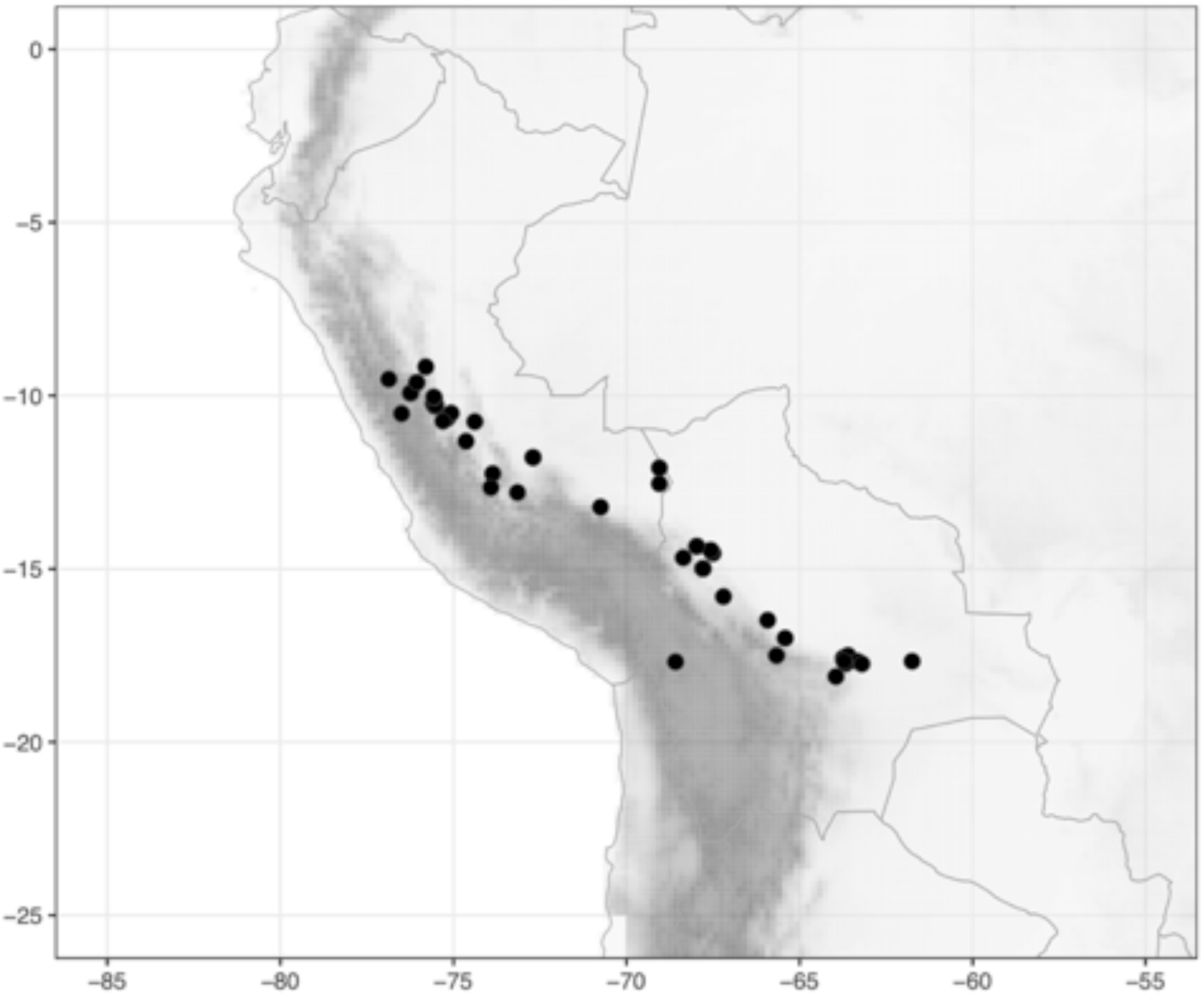
Distribution of *Dimerocostus argenteus* (Ruiz & Pav.) Maas (●).

**Map 53.**
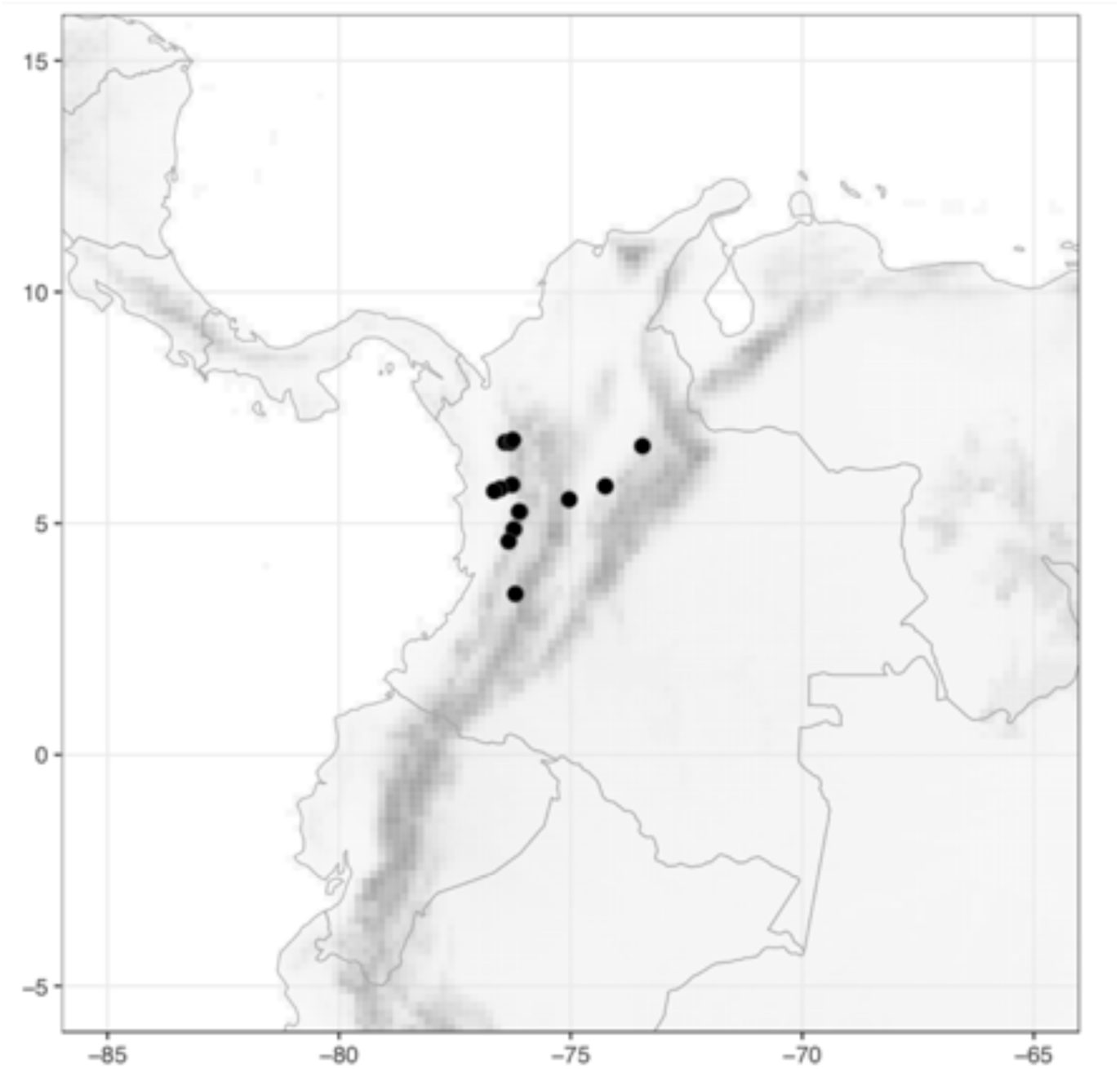
Distribution of *Dimerocostus cryptocalyx* N.R.Salinas & Betancur B. (●).

**Map 54.**
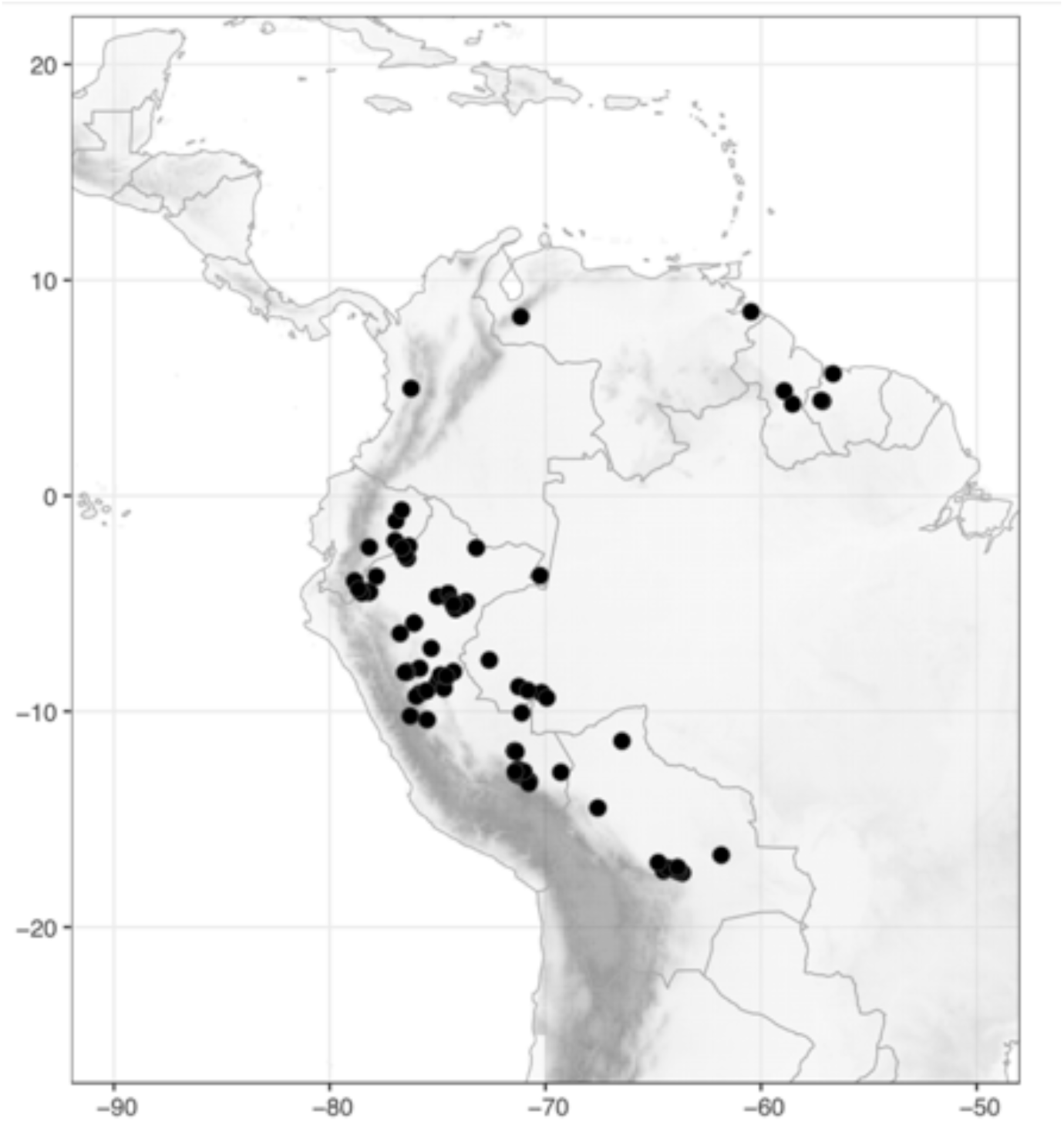
Distribution of *Dimerocostus rurrenabaqueanus* (Rusby) Maas & H.Maas (●).

**Map 55.**
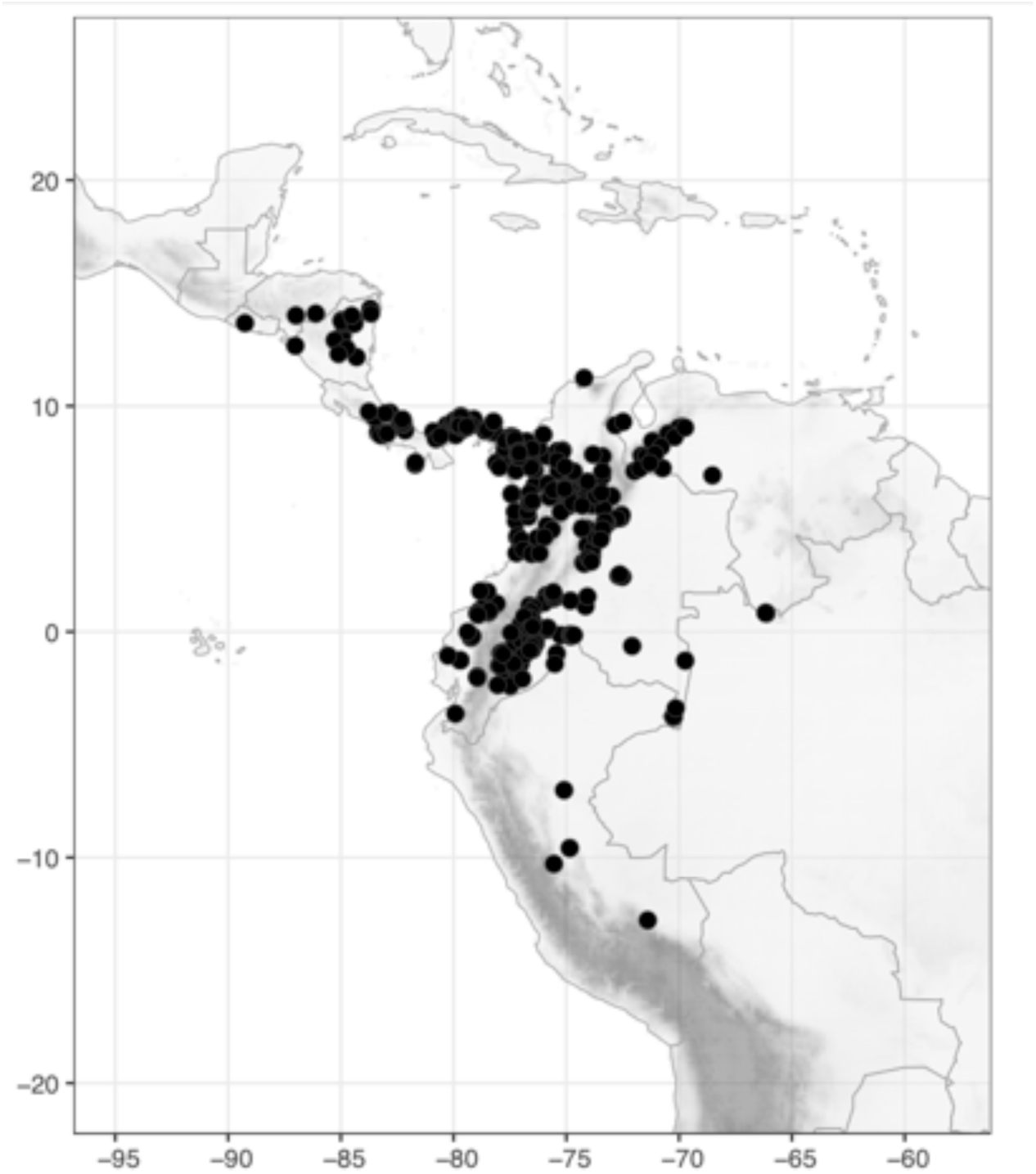
Distribution of *Dimerocostus strobilaceus* Kuntze (●).

**Map 56.**
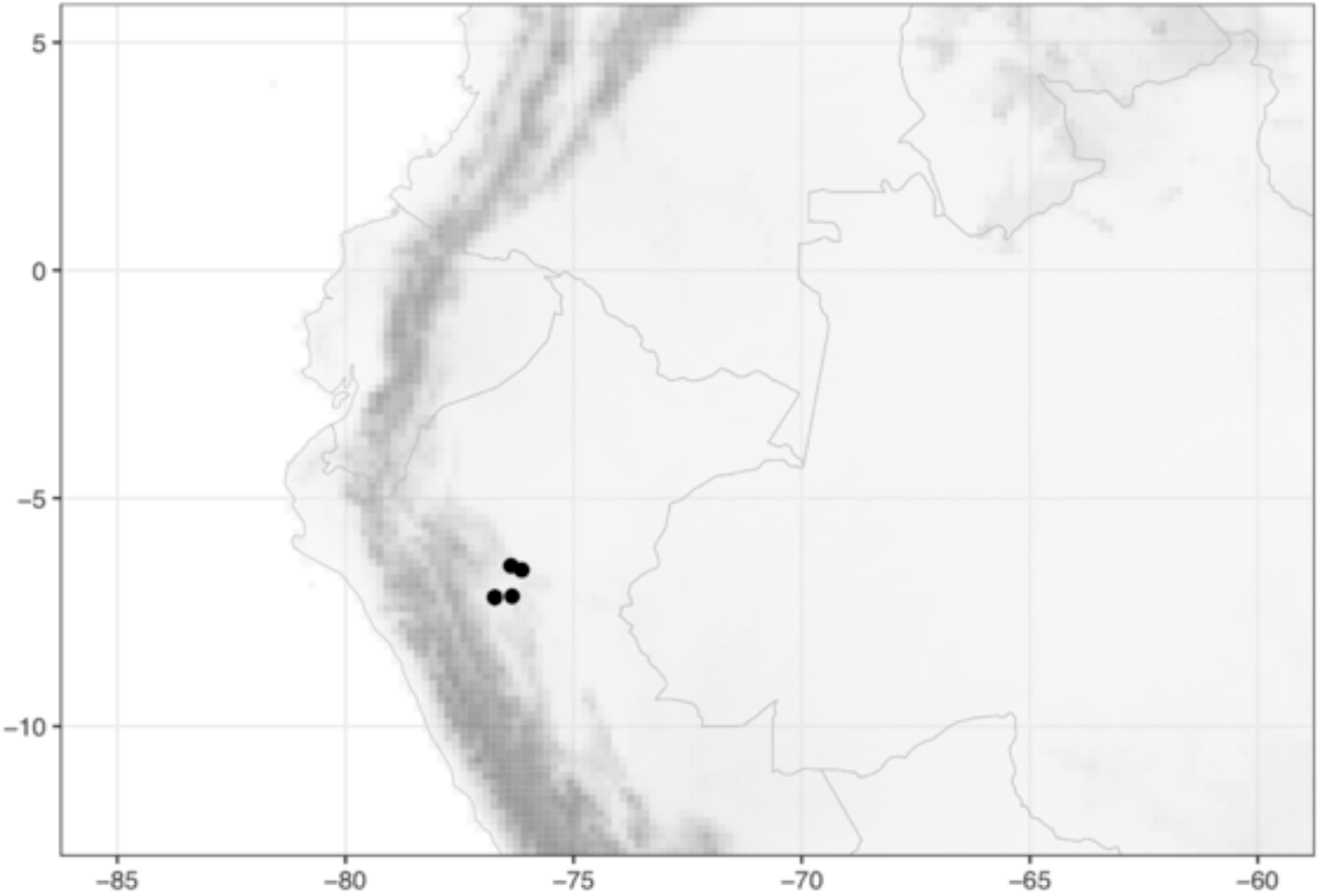
Distribution of *Monocostus uniflorus* (Poepp. ex Petersen) Maas (●).

